# Twists and turns of the salicylate catabolism in *Aspergillus terreus*

**DOI:** 10.1101/2020.07.23.217166

**Authors:** Tiago M. Martins, Celso Martins, Cristina Silva Pereira

## Abstract

In fungi, salicylate catabolism was believed to proceed only through the catechol branch of the 3-oxoadipate pathway, as shown *e.g.* in *Aspergillus nidulans*. However, the observation of a transient accumulation of gentisate upon cultivation of *Aspergillus terreus* in salicylate media questions this concept. To address this we have run a comparative analysis of the transcriptome of these two species after growth in salicylate using acetate as a control condition. The results revealed the high complexity of the salicylate metabolism in *A. terreus* with the concomitant positive regulation of several pathways for the catabolism of aromatic compounds. This included the unexpected joint action of two pathways: the nicotinate and the 3-hydroxyanthranilate, possibly crucial for the catabolism of aromatics in this fungus. New genes participating in the nicotinate metabolism are here proposed, whereas the 3-hydroxyanthranilate catabolic pathway in fungi is described for the first time. The transcriptome analysis showed also for the two species an intimate relationship between salicylate catabolism and secondary metabolism. This study emphasizes that the central pathways for the catabolism of aromatic hydrocarbons in fungi hold many mysteries yet to be discovered.

**IMPORTANCE:** Aspergilli are versatile cell factories used in industry for production of organic acids, enzymes and pharmaceutical drugs. To date, organic acids bio-based production relies on food substrates. These processes are currently being challenged to switch to renewable non-food raw materials; a reality that should inspire the use of lignin derived aromatic monomers. In this context, Aspergilli emerge at the forefront of future bio-based approaches due to their industrial relevance and recognized prolific catabolism of aromatic compounds. Notwithstanding considerable advances in the field, there are still important knowledge gaps in the central catabolism of aromatic hydrocarbons in fungi. Here, we disclosed a novel central pathway, defying previous established ideas on the central metabolism of the aromatic amino acid tryptophan in Ascomycota. We also observed that the catabolism of the aromatic salicylate greatly activated the secondary metabolism, furthering the significance of using lignin derived aromatic hydrocarbons as a distinctive biomass source.

## Introduction

In recent years, the production of organic acids through bio-based processes has resurfaced as sustainable alternative to petroleum-based production. This is the case of the biotechnological production of *e.g.* succinic acid that is already produced at an industrial scale (1). The bio-based production of organic acids is currently challenged as to shift from food (corn, beet, cane, etc.) to renewable non-food feedstocks that may include production by engineered microorganisms (2). Aspergilli are used industrially for the production of organic acids (*e.g.* citric acid, itaconic acid) and secondary metabolites (*e.g.* the cholesterol-lowering drug lovastatin), hence seen as ideal candidates for novel production processes (3). Lignin, a heterogeneous aromatic plant polymer, is the second most abundant after cellulose. It constitutes a major underutilized residue from various industrial processes (*e.g.* >100 million tons per year from Kraft pulping alone) (4). Lignin high recalcitrance hinders many biotechnological approaches (5), despite that recent progresses in biorefinery can ensure its depolymerization into numerous and structurally diverse aromatic hydrocarbons (5, 6). In nature, lignin depolymerization is mostly accomplished by potent oxidative enzymes secreted by fungi that possess multiple strategies for the catabolism of these aromatic hydrocarbons (7). In fact, fungi can use countless peripheral pathways for their catabolism, all of which converge towards a small number of intermediates that undergo ring-cleavage in the central pathways. So far, the better described central pathways are of the intermediates catechol, protocatechuate, hydroxyquinol, homogentisate and gentisate (8). In the last decade, many of the catabolic genes composing these central pathways have been assigned (8–13). Searching for these genes in the genomes of Dykaria revealed that the five central pathways are present in both Ascomycota and Basidiomycota, sharing simultaneously high similarity but also potentially some speciation (*e.g.* a distinctive Basidiomycota protocatechuate 3,4-dioxygenase) (8). In particular, the existence of specific dioxygenases for gallate, hydroquinone, homoprotocatechuate and pyrogallol, as described for bacteria, remains unclear to date, despite that these compounds are known to be channeled into the known central pathways (10, 14, 15).

Aspergilli generally display in their genomes the orthologous genes for all the better described central catabolic pathways, with the exception of the gentisate pathway, which is either present, partially present or absent (8, 16). In addition, the gentisate 1,2-dioxygenase orthologous gene is constantly organized in a cluster that comprises either all the other pathway genes (designated gentisate cluster) or all except the maleylpyruvate isomerase gene (designated gentisate-*like* cluster) (8). In several fungal genomes the two clusters are present (*e.g. Aspergillus niger*) suggestive of different and unknown functional roles. *Aspergillus terreus* has a gentisate-*like* gene cluster in its genome, whereas *A. nidulans* does not have any kind of a gentisate gene cluster or a gentisate 1,2-dioxygenase orthologous gene. In the present study we undertook a comparative transcriptome analysis of these two species upon their growth in salicylate as sole carbon and energy source using acetate as the control condition. The study was complemented by expression analyses of targeted genes and chromatographic quantifications of specific intermediates.

## Materials and Methods

### Strains and growth conditions

*Aspergillus nidulans* FGSC A4 or A1147 and *A. terreus* FGSC A1156 asexual spores were harvested and maintained as frozen suspensions at −80 °C (17). Cultures were initiated with 10^6^ spores/mL and incubated with orbital agitation (200 rpm) in the dark at 37 °C. Batch cultivations were performed in 250 mL Erlenmeyer flasks with a working volume of 50 mL. A low nitrogen minimal medium was used containing per liter 3 g NaNO_3_, 0.01 g ZnSO_4_·7H_2_O, 0.005 g CuSO_4_·5H_2_O, 0.5 g MgSO_4_·7H_2_O, 0.01 g FeSO_4_·7H_2_O, 0.5 g KCl. Filter sterilized salts were added to an autoclave sterilized 100 mM potassium phosphate (pH 6.0) solution. The carbon sources were added either directly to the phosphate solution for sterilization (60 mM sodium acetate - control) or to the mineral media after filter sterilization (20 mM sodium salicylate). Solid minimal medium was jellified with 5 g/L agarose, and cultures grown at 30°C in the dark, on top of a PVDF membrane (Hybond-P, 0.45 µm, GE Healthcare) (18).

### Metabolites identification and quantification

Culture media were analyzed by ultra-performance liquid chromatography (UPLC) for the identification and/or quantification of the aromatic compounds and their transformation products following an established protocol (19).

### Transcriptome Analysis

Total RNA extraction of liquid grown mycelia was performed essentially as previously described (17), using a Tissuelyser LT (Qiagen) for cell disruption. The quality and quantity of RNA was determined by capillary electrophoresis using HS RNA kit and 5200 Fragment Analyzer (Agilent Technologies). For single-end RNA sequencing (RNA-seq), libraries were generated using the Smart-Seq2® mRNA assay (Illumina, Inc.) according to the manufacturer’s instructions. Twelve samples were indexed and sequenced on the Illumina NextSeq550 (20M reads per sample). Generated FastQ files were analyzed with FastQC and any low-quality reads were trimmed with Trimmomatic (20). All libraries were aligned to the corresponding model fungus; either *A. nidulans* FGSC A4 genome assembly (ASM14920.v2) or *A. terreus* NIH2624 genome assembly (ASM14961.v1) with gene annotations from Ensembl Fungi v. 37 using HISAT2 v. 2.1.0 (21) and only matches with the best score were reported for each read. All RNA-seq experiments were carried out in three biological replicates. Differential expression analysis was performed using DESeq2 v. 1.24.0 (22). Principal component analysis (PCA) plot and MA-plots were also generated using the DESeq2 package. The genes that showed more than log_2_ 1-fold expression changes with *p*-adj value <0.05 are defined as significant differentially expressed genes in this analysis. Transcript abundance was defined as the number of reads per kilobase of transcript per million mapped reads (RPKM). The full genomes were scanned using InterProScan v. 5.32-71.0 (23) to build a protein domain database for further enrichment analyses. The enrichment analyses were performed with the FunRich software v. 3.1.3 (24), using the constructed protein domain databases as the terms library and the hypergeometric test with a *p*-value < 0.05.

### Reverse transcription quantitative PCR analysis (RT-*q*PCR)

Total RNA extraction was performed as reported above and cDNA synthesis as previously described (17). Oligonucleotide pairs were designed using Primer-BLAST (25) and supplied by STAB Vida (Oeiras, Portugal) (Table S1). The RT-*q*PCR analysis was performed in a CFX96 Thermal Cycler (Bio-Rad), using the SsoFast EvaGreen Supermix (Bio-Rad), 250 nM of each oligonucleotide and the cDNA template equivalent to 10 ng of total RNA, at a final volume of 10 µl per well, in three biological replicates. The PCR conditions were: enzyme activation at 95 °C for 30 s; 40 cycles of denaturation at 95 °C for 5 s and annealing/extension at 60 °C for 15 s; and melting curve obtained from 65 °C to 95 °C, consisting of 0.5 °C increments for 5 s. Data analyses were performed using the CFX Manager software v. 3.1 (Bio-Rad). The expression of each gene was taken as the relative expression in pair-wise comparisons of each condition relative to the acetate control. The expression of all target genes was normalized by the expression of the 60S ribosomal protein L33-A gene, AN2980 or ATEG_01624.

**Table 1.** Observed regulation for secondary metabolism biosynthesis genes in salicylate media. Genes were considered upregulated when the fold change ≥ 1 and downregulated ≤ −1 and for both when *p*-value ≤ 0.05. Polyketide synthase (PKS), nonribosomal peptide synthetase (NRPS), hybrid (PKS and NRPS), and aromatic prenyltransferases (PTases).

| Species | *Aspergillus nidulans* |  |  | *Aspergillus terreus* |  |  |
| --- | --- | --- | --- | --- | --- | --- |
| Genes regulation | Total | Up | Down | Total | Up | Down |
| Biosynthesis cluster genes | 446 | 260 | 39 | 398 | 118 | 51 |
| Core biosynthesis genes | 96 | 57 | 13 | 111 | 41 | 11 |
| PKS | 32 | 23 | 3 | 32 | 13 | 3 |
| NRPS | 33 | 17 | 6 | 39 | 13 | 4 |
| Hybrid | 1 | 1 | 0 | 1 | 1 | 0 |
| PTases | 7 | 6 | 0 | 11 | 4 | 1 |
| Terpene | 11 | 6 | 1 | 10 | 1 | 1 |
| β-lactone | 6 | 2 | 2 | 4 | 2 | 1 |
| Cyclic peptide | 6 | 2 | 1 | 13 | 6 | 1 |
| Siderophore | 0 | 0 | 0 | 1 | 1 | 0 |

### Gene structure prediction

Automated gene structure prediction of new genes or of poorly annotated genes was done using FGENESH with or without similar protein-based gene prediction (26). Gene structure was manually curated (Table S2) and used in the transcriptome analysis.

### Identification of gene clusters

Metabolic gene clusters were identified within annotated genomes through sequence homology searches using MultiGeneBlast v. 1.1.13 (27). A recent customized database created from online GenBank entries was done using the same software. Identification of secondary metabolite biosynthesis gene clusters was done using antiSMASH v. 5.0 (28) and the results were incorporated in available information (Table S3 and S4), namely from pre-computed results at MycoCosm obtained by using SMURF (29, 30).

### Data availability

Gene transcript RPKM values are contained in Tables S5 and S6, and raw data with metadata were deposited in the Sequence Read Archive accession number SUB7137746.

## Results and discussion

Cultivation of *Aspergillus terreus* in salicylate (20 mM) resulted in the transient accumulation of gentisate up to *ca*. 3 mM (Figure 1). This observation conflicts with salicylate catabolism only through the catechol branch of the 3-oxoadipate pathway as systematically shown in other aspergilli, including in *A. nidulans* ((9) and references therein). Some yeasts have been shown to use the gentisate pathway for the catabolism of 3-hydroxycinnamates, 3-hydroxybenzoate and gentisate (2,5-dihydroxybenzoate) (12). The genes of the gentisate pathway are clustered in the genomes of fungi (8), yet in the gentisate-*like* cluster the maleylpyruvate isomerase gene is absent as observed in *A. terreus* (ATEG_06711 to ATEG_06714). This suggests that during the catabolism of salicylate in *A. terreus*, gentisate catabolism also takes place through a yet unknown pathway. To better understand the catabolism of salicylate in *A. terreus* we analyzed its transcriptome response and, that of *A. nidulans*, grown in salicylate. The comparative analysis revealed that both species share a global common response to salicylate compared to the control conditions in acetate (Figure 2). In the presence of salicylate, 2703 encoding genes were found differentially upregulated and 1489 downregulated in *A. nidulans*; and 2980 upregulated and 2328 downregulated in *A. terreus* (see Figures S1 and S2 for PCA and MA plots, Tables S5 and S6). The first striking feature is that many of the upregulated genes could not be annotated in KEGG (*ca*. 80%) while half of the downregulated ones could (*ca*. 50%). This is likely related to the fact that well-known processes/pathways were globally downregulated in salicylate medium, such as, carbohydrate metabolism (*e.g.* tricarboxylic acid (TCA) cycle, glycolysis, pentose phosphate pathway), energy metabolism (*e.g.* oxidative phosphorylation) and genetic information processing (*e.g.* protein processing in the endoplasmic reticulum). As expected, the acetate utilizing genes (*fac*A to *fac*C, *acu*D to *acu*N and *mae*A to *mae*B) were downregulated, including the ones from the glyoxylate shunt (Table S7). Globally, the amino acid metabolism was greatly affected in both species. Specifically, the aromatic amino acids metabolism was upregulated and their biosynthesis was downregulated (Table S5 and S6). The peripheral pathways of the catabolism of aromatic compounds show resemblance to the metabolism of phenylalanine, tyrosine and tryptophan, suggestive of its upregulation. In agreement, and expectably, the xenobiotics biodegradation and metabolism was also upregulated. The analysis of the protein domains demonstrated, that those upregulated and enriched are from families of oxidoreductases, hydrolases, transporters, transcriptional regulators, and transferases (Figure 2).

**Figure 1.**
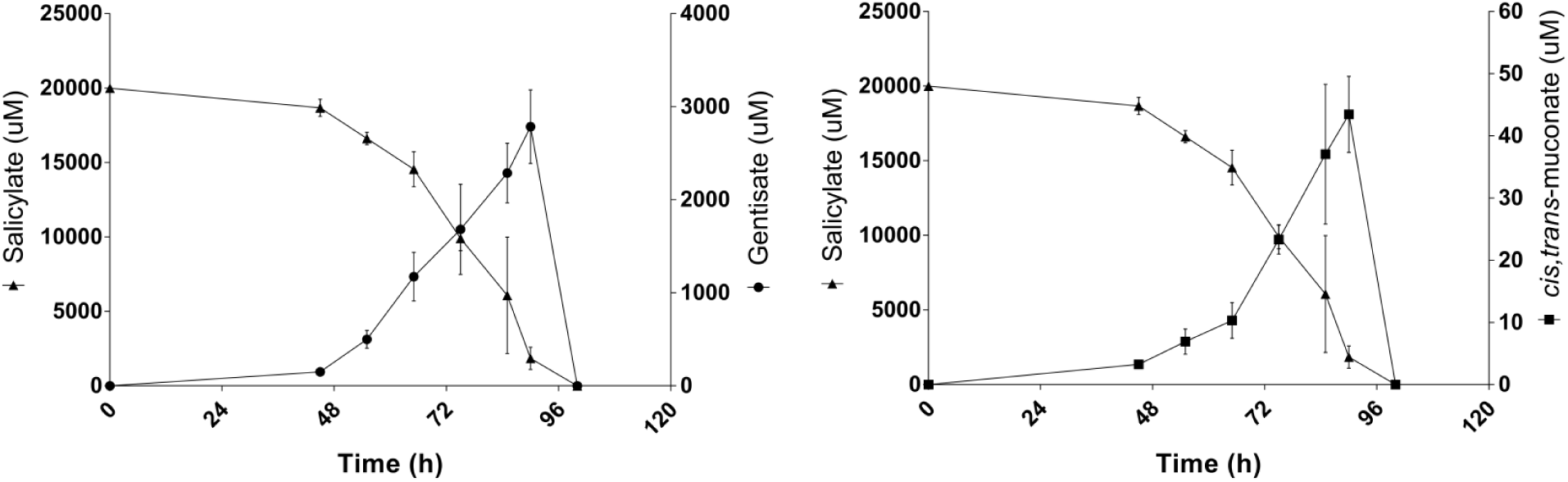
Depletion of salicylate and formation of gentisate and *cis,trans*-muconate in *Aspergillus terreus* liquid media cultures at discrete cultivation time-points (chromatographic analyses). 2,3-Dihydroxybenzoate, of which the retention time is very similar to gentisate, was detected but at very low concentration therefore not quantified. Values represent means ± standard deviation of three biological replicates.

**Figure 2.**
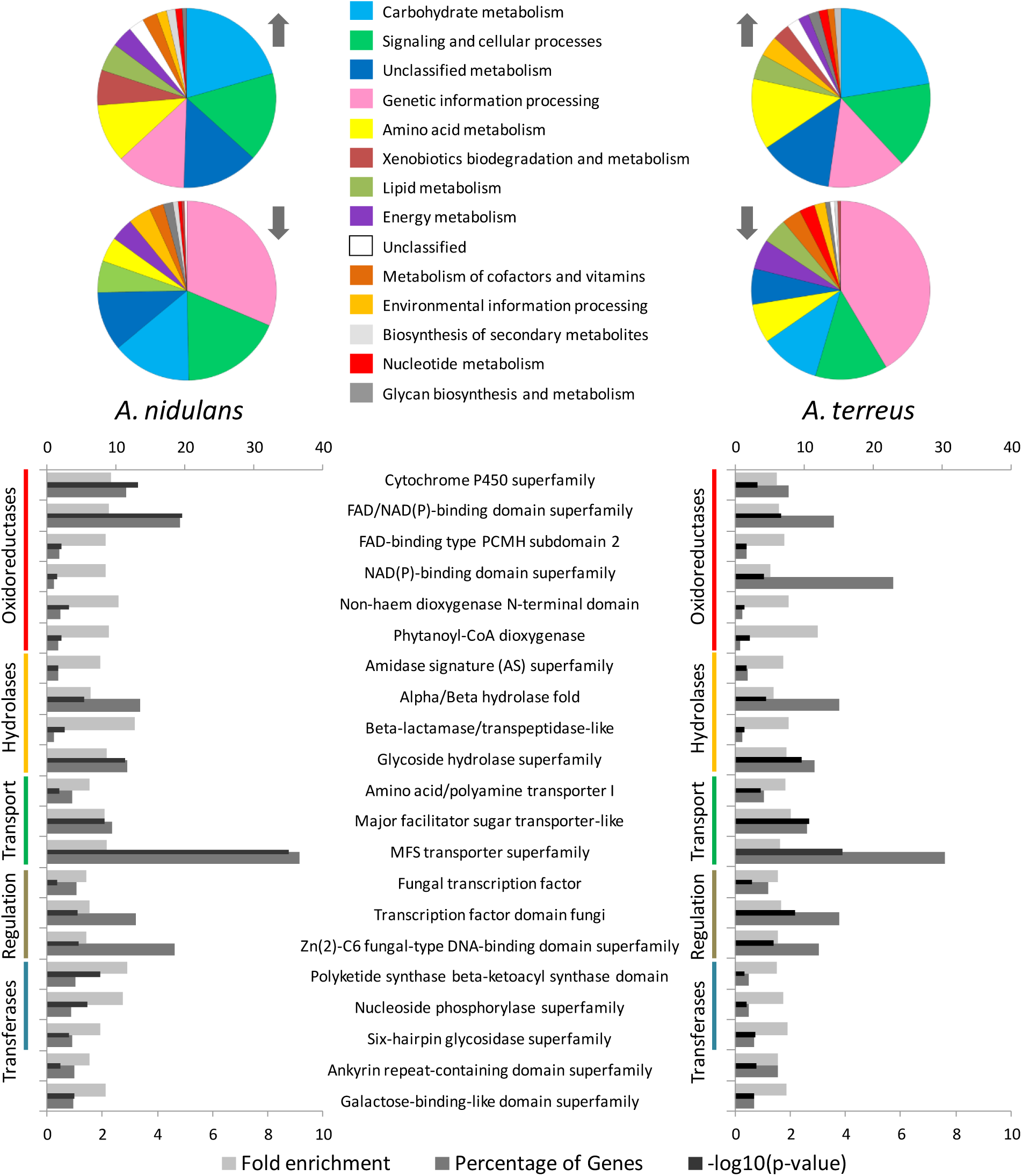
Analysis of the RNA-seq data collected for *Aspergillus nidulans* and *A. terreus* grown in salicylate (compared to acetate) comprising functional characterization and protein domains enrichment of the observed differentially expressed genes. KEGG functional categories were slightly simplified by creating super categories that merge the most similar ones. The predicted protein domains of the upregulated and significantly enriched genes in salicylate (*p*-value ≤ 0.05) for both species are shown. Fold enrichment and percentage of upregulated genes is displayed in the bottom axis and the log_10_(*p*-value) in the top axis.

On the contrary, none of the downregulated genes encode for abundant enriched protein domain families, with the exception of FAD/NAD(P)-binding domain (IPR023753) found in diverse oxidoreductases (*e.g.* thioredoxin and glutathione reductases, dihydrolipoyl dehydrogenase).

### Secondary Metabolism

Both aspergilli species grown in salicylate underwent similar transcriptome responses in secondary metabolism, with the upregulation of nearly half of the core biosynthesis genes (Table 1, Tables S3 and S4). In addition, only *ca.* 10% of the genes annotated in secondary metabolism underwent downregulation. Specifically, in *A. nidulans*, five out of ten *lae*A-like methyltransferase genes (*llm*A, *llm*C, *llm*E, *llm*I and *llm*J) with potential regulatory roles in secondary metabolism (31), were found upregulated, while the remaining five were unaltered. In *A. terreus*, out of the eight *lae*A-like genes, three were upregulated (ATEG_01440, ATEG_06013 and ATEG_09440), four were unaltered (ATEG_00678, ATEG_01936, ATEG_04864 and ATEG_00912) and only one was downregulated (ATEG_07798). Collectively, the concerted response observed in secondary metabolism reveals its close relation with the catabolism of the aromatic salicylate. To better understand this link, we then analyzed with greater detail some secondary metabolite gene clusters.

In salicylate, the monodictyphenone gene cluster of *A. nidulans* was upregulated, whereas that of sterigmatocystin was downregulated, consistent with that observed before by us using proteomics analyses (9). Besides the monodictyphenone gene cluster, the other known prenyl xanthone biosynthesis genes were also found moderately to highly expressed and upregulated. This included the aromatic DMATS-type prenyltransferases (PTases) *xpt*A (AN6784) and *xpt*B (AN12402), the GMC oxidoreductase superfamily *xpt*C (AN7998), and the short-chain dehydrogenase/reductase (SDR) gene AN7999. The last was reported before to have no obvious effect on prenyl xanthone biosynthesis (32), however the major expression and upregulation of AN7999 observed here contradicts this idea. Other DMATS-type PTases were also found moderately to highly expressed and upregulated, namely AN11080, AN11194/AN11202 and AN10289. The first gene belongs to the nidulanin A cluster and the second to viridicatin/aspoquinolone, using as precursors either kynurenine or anthranilate, and the amino acids phenylalanine and valine (33, 34). To study the function of the yet uncharacterized AN10289 gene that underwent here major expression and upregulation, salicylate can be used as an inducer. The same applies to the study of NRPS AN8433 gene, which is located in a cluster that contains also a tyrosinase and two cytochrome P450 monooxygenases genes, here also greatly expressed and upregulated by salicylate. While monodictyphenone biosynthesis relies on malonyl-CoA as precursor (32), formed from plentiful acetyl-CoA made available by the catabolism of salicylate, the biosynthesis of the other prenyl xanthones (also undergoing upregulation) use precursors that are not direct metabolic products, further questioning the nature of salicylate induction.

Salicylate also upregulated many secondary metabolite biosynthetic clusters in *A. terreus*, including the asterriquinones and the aspulvinones clusters that comprise the DMATS-type PTases ATEG_09980 and ATEG_01730, respectively. These clusters use as precursors indole-3-pyruvate and 4-hydroxyphenylpyruvate, respectively (35–37), which belong to the tryptophan and the phenylalanine pathways also undergoing upregulation in salicylate. A putative cyclic peptide (ribosomally synthesized and post-translationally modified peptide, RiPP) gene cluster comprising seven genes, underwent massive expression and upregulation in salicylate. One of its genes, the ATEG_07504 encodes a precursor protein with seven repetitions of the motif XANDVD[EA]ISTRS[QE]WFYNK[KR]. The motif has flanking cleaving sites, namely XA for serine peptidase of the S9 family (*i.e.* STE13) and KK or KR for the S10 family (*i.e.* KEX), analogous to those found in the Ascomycota α pheromone and in other cyclic or non-cyclic peptides (38). The presence of a serine peptidase gene (ATEG_07509) in this cluster is suggestive of the participation of the encoded S10 family peptidase (IPR001563) in the initial processing of the RiPP precursor protein. Only one serine peptidase S9 family (ATEG_09875) (located outside of the gene cluster) was observed to be greatly expressed and upregulated in salicylate, likewise indicative of its participation in the biosynthesis. The cluster contains also three UstY homologs with the DUF3328 domain (encoded by ATEG_07505 to 07) that catalyzes sequential cyclization events. One cyclization likely involves oxidation of a tyrosine residue (39). Another may require prior dehydration of serine and threonine residues (present in the conserved motif), possibly mediated by the activity of protein kinase ATEG_07508 (IPR011009) also encoded in the cluster. This step is required for nucleophilic attack of cysteine thiols and formation of thioether bonds, as previously reported in the biosynthesis of lanthipeptides (40). Finally, the last gene out of the seven of the RiPP cluster, the ATEG_07503, encodes for a aminomethyltransferase (IPR013977) that likely mediates a N-methylation of the cyclic peptide, possibly to increase its proteolytic stability and membrane permeability (41). ATEG_07503 orthologous gene is found in *Aspergillus fumigatus*, specifically Afu8g06470, regardless of being placed in a different putative RiPP cluster. Interestingly, the RiPP cluster of *A. terreus* is only similar to the ones present in distantly related Ascomycota *Drechmeria* and *Venturia* genera, what suggests that the produced cyclic peptide may confer competitive advantages in specific niches.

Our observations highlighted the interconnection of secondary metabolism and salicylate catabolism in aspergilli, possibly a consequence of shared regulatory and metabolic routes, including availability of specific precursors. Our initial findings deserve further consideration in the near future; especially as the use of simple hydrocarbon aromatics, such as salicylate, may be used to activate unknown biosynthesis clusters, guiding the identification of the formed metabolites.

### Salicylate Catabolism in *Aspergillus nidulans*

As previously reported, salicylate catabolism in *A. nidulans* most likely results, either directly or with 2,3-dihydroxybenzoate as intermediate, in catechol formation (9). The formed catechol, a central substrate for a ring-cleavage dioxygenase, is channeled to the respective branch of the 3-oxoadipate pathway, here found mostly upregulated (Figure S3 and Table S8). However, only the two genes of the non-oxidative decarboxylation pathway and the catechol 1,2-dioxygenase (AN7418, *dhb*D and AN4532) underwent major expression, while the remaining ones only minor expression (AN2114, AN4061, AN4531 and AN10495). We have previously observed that some of these less expressed genes are not essential for growth in salicylate (9). Accordingly, we previously suggested that the hydroxyquinol variant of the 3-oxoadipate could constitute an alternative path (9). In the present study, we observed by RT-*q*PCR expression analyses of the ΔAN4531 mutant grown on salicylate that this alternative path is unlikely (Figure S4). Herein, we noticed that the dark colored metabolite, likely 3-oxoadipate enol-lactone that accumulated during the growth of this mutant (9), disappeared after prolonged growth incubations. These observations suggest that muconate and their catabolic derivatives may be metabolized by broad substrate specificity enzymes. The upregulation of several oxidoreductase and hydrolase gene families (Figure 2) most likely contributes to redundancy in the metabolism of muconate, and may offer metabolic plasticity in similar contexts. Besides the two salicylate monooxygenase genes of the catabolic pathway (AN2114, AN7418), 45 other aromatic-ring FAD-binding monooxygenase genes (IPR002938) were found moderately to highly expressed and upregulated, including seven phenol monooxygenase genes (IPR038220). A major facilitator superfamily (MFS) transporter gene (AN10845) that underwent upregulation in salicylate is located contiguous to *dhb*D. These two genes show shared synteny within aspergilli and penicillia, suggestive that this MFS transporter may be a 2,3-dihydroxybenzoate transporter. This requires validation since numerous other genes of this superfamily were also positively regulated.

One remarkable observation is that we found also a cluster of four genes highly expressed and upregulated in salicylate. It comprises the following genes: the putative salicylate 1-monooxygenase (AN2114) previously identified by us (9), a domain of unknown function (DUF3425) gene (AN2115), a NAD-dependent epimerase/dehydratase (AN2116) and an amydohydrolase (AN2118) (Table S8). The last gene has sequence homology to *ors*B (orsellinic cluster) and *dhb*D, both encoding for enzymes that catalyze the decarboxylation of aromatic hydrocarbons. This cluster is only present in the genomes of aspergilli belonging to the *Nidulantes* section (Figure S5), being absent in most aspergilli including *A. terreus*, hence suggestive of a high specific functional role. One hypothesis is that this putative salicylate 1-monooxygenase plays other functions than the hydroxylation of salicylate in this fungus, possibly acting in other peripheral pathway or in secondary metabolism.

### Salicylate Catabolism in *Aspergillus terreus*

The known pathways in microorganisms for the catabolism of salicylate either involve the catechol branch of the 3-oxoadipate pathway or the gentisate pathway (7). We observed in *A. terreus* that salicylate was hydroxylated to 2,3-dihydroxybenzoate, similarly to what occurs in *A. nidulans,* but also to gentisate in significant amounts (Figure 1). The orthologous or predicted genes for these pathways are present in the genome of *A. terreus*. The RNA-seq data revealed that these genes were mostly unaltered in salicylate medium in the respective time point (Figure 3 and Table S9). However, massive upregulation of the non-oxidative decarboxylase gene (ATEG_06350) and of key genes coding in the catechol branch (ATEG_09602; ATEG_03095) took place at early time points along the incubation in salicylate medium as verified by their RT-*q*PCR expression levels (Figure 4). Corroborating this result, muconolactone, *cis*,*trans*-muconate and 2,3-dihydroxybenzoate (Figure 1) were detected in low amounts in the extracellular media. The upregulation of the gentisate-*like* cluster (ATEG_06711 to ATEG_06714) was less expressive and occurred at later time points compared to the catechol branch (Figure 4). Salicylate 3-monooxygenase (AN7418) orthologous gene is not present in the genomes of *A. terreus* and other aspergilli. In addition, the identity of salicylate 5-monooxygenase is even more obscure as it has not been previously reported in fungi. Alike that observed here for *A. nidulans*, 32 aromatic-ring FAD-binding monooxygenases (IPR002938) were found moderately to highly expressed and upregulated, including six phenol monooxygenases (IPR038220), hindering a precise assignment of the salicylate monooxygenases at this stage.

**Figure 3.**
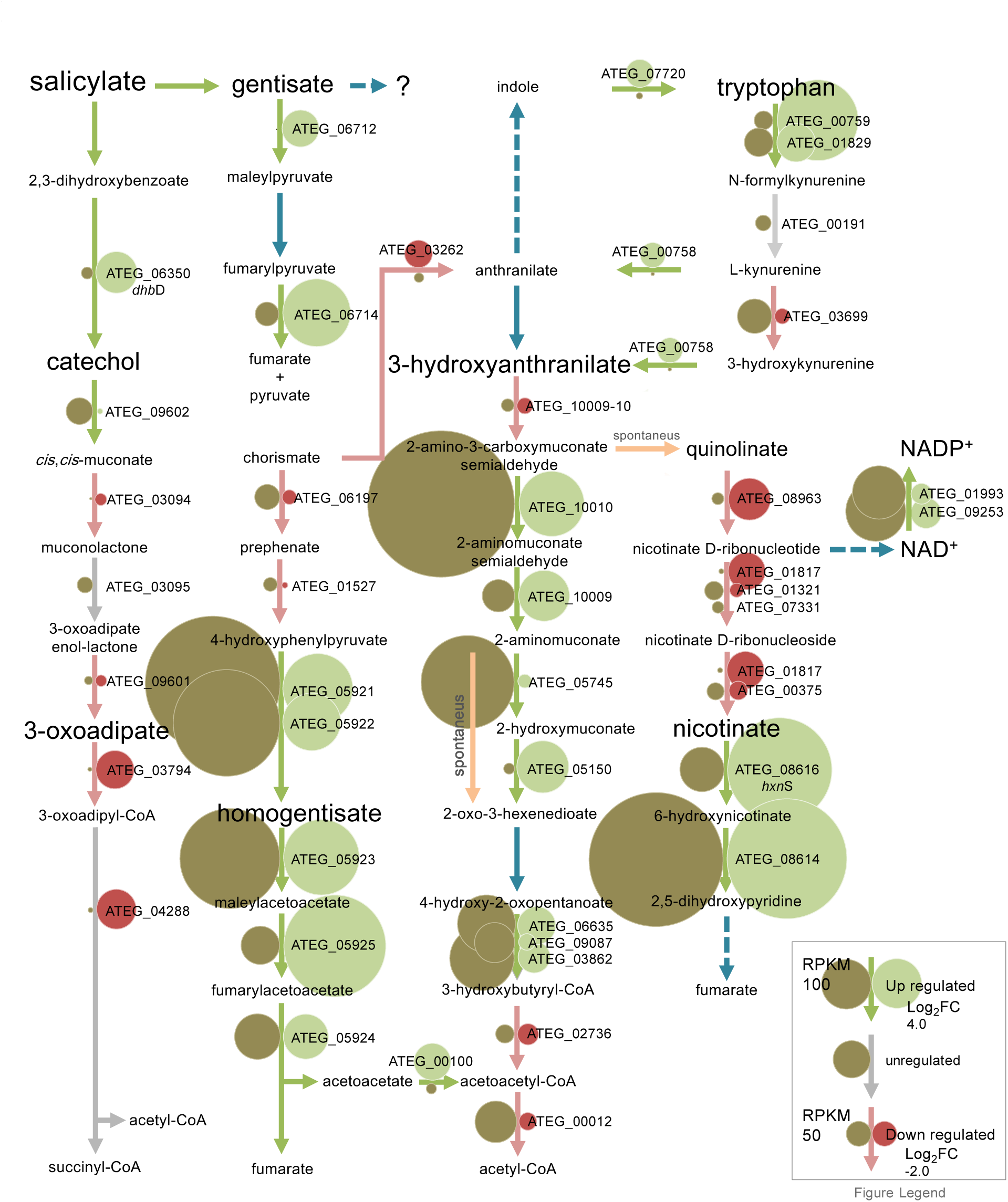
Diagram of the metabolism of salicylate in *Aspergillus terreus*. The differentially expressed genes (RNA-seq) regulation and abundance is represented by circles with variable size and color (see inserted legend).

**Figure 4.**
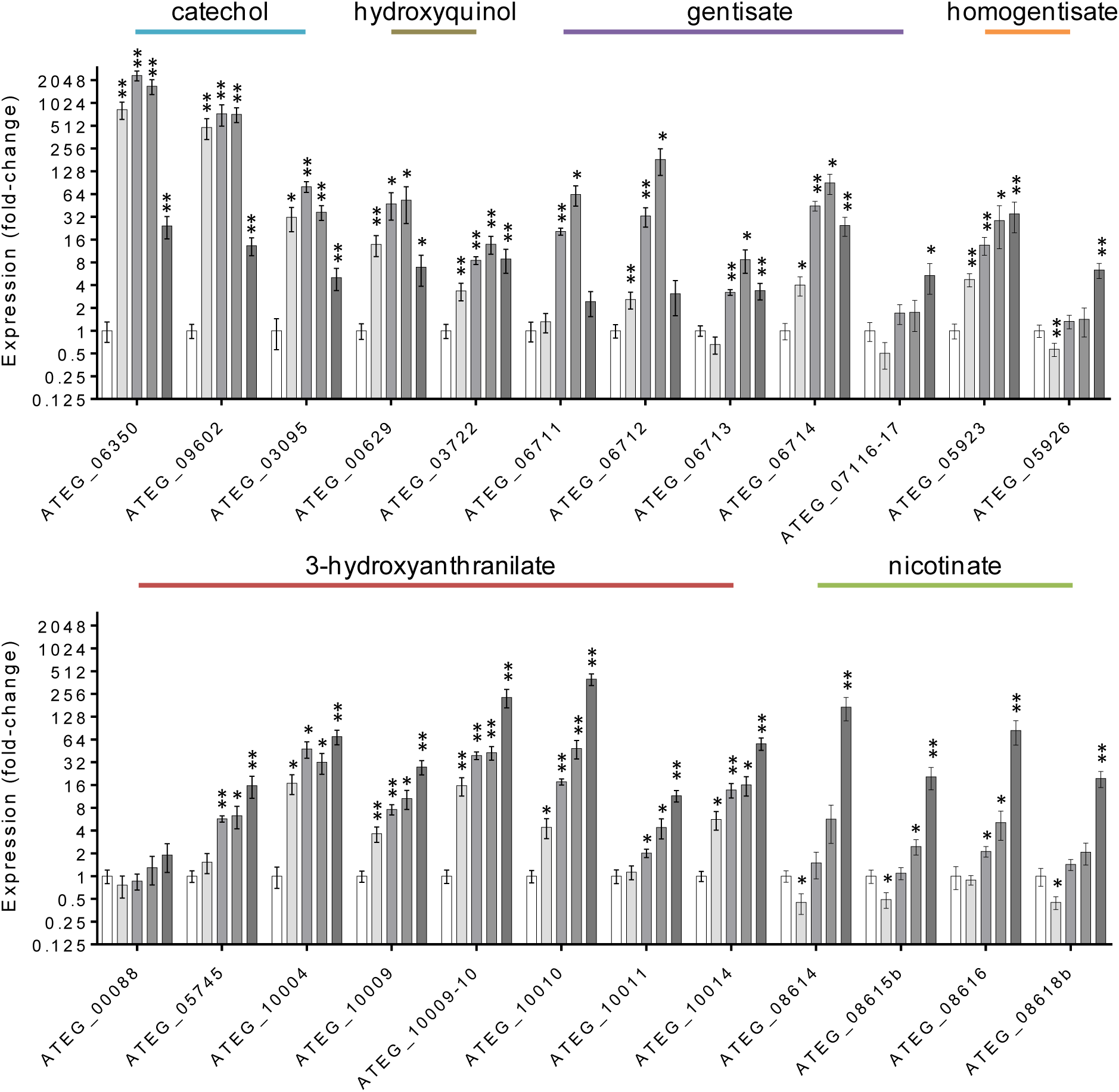
Expression levels of genes coding in the metabolic pathways of *Aspergillus terreus* in salicylate media along cultivation (grey bars: 55h, 75h, 90h and last time point used for RNA-seq) compared to the control in acetate media (white bar: time point used for RNA-seq). The central intermediates of each metabolic pathway are indicated. Values represent means, ± standard error of the mean of three biological replicates (* *p*-value < 0.05 and ** < 0.005).

The RNA-seq data showed the upregulation of several genes of the phenylalanine, tyrosine and tryptophan catabolism, as well as three gene clusters, namely the homogentisate cluster (ATEG_05922 to ATEG_05926), the nicotinate-inducible cluster (ATEG_08614 to ATEG_08620 and ATEG_05673) and a cluster unknown in fungi (Figure 3, Table S9). The first two have been previously described in *A. nidulans* (42, 43). The last cluster is here described for the first time in fungi, which we designate as the 3-hydroxyanthranilate catabolic cluster. Three of the comprised genes of this cluster putatively encode for three consecutive enzymes of the tryptophan kynurenine pathway (44), namely 3-hydroxyanthranilate dioxygenase (3HAO, ATEG_10009-10, not annotated in the public databases; Table S2), aminomuconate semialdehyde dehydrogenase (AMSDH, ATEG_10010) and 2-amino 3-carboxymuconate 6-semialdehyde decarboxylase (ACMSD, ATEG_10009). This cluster, as well as some upstream pathway genes, was observed to be upregulated in salicylate (Figure 3, Table S9). Three types of indoleamine 2,3-dioxygenases (IDO) present in aspergilli can catalyze L-tryptophan to kynurenine at the start of the pathway, although with different biochemical characteristics and unknown functional roles (45). IDOα (ATEG_03482) was found downregulated and highly expressed, while IDOβ (ATEG_00759) and IDOγ (ATEG_01829) were found upregulated and with even higher expression levels. However a fourth IDO gene (ATEG_07359) is present in the genome of *A. terreus*, which exhibited almost no expression in salicylate and acetate media. The 3-hydroxyanthranilate dioxygenase genes in fungi have been merely associated to an alternative path for NAD^+^ biosynthesis (46). Our data contradict this assumption, constituting a first evidence of the existence of the catabolic tryptophan kynurenine pathway forming 2-aminomuconate in fungi (Figures 3 and 4). The reaction product of 3-hydroxyanthranilate dioxygenase is 2-amino 3-carboxymuconate semialdehyde. This aldehyde decays non-enzymatically to quinolinate, which is the precursor of NAD^+^. 2-Aminomuconate semialdehyde can non-enzymatically form picolinate, a bidentate chelating agent, of which the metabolism remains largely unknown (44). The 3-hydroxyanthranilate catabolic gene cluster here identified is present across Ascomycota, including some species of *Aspergillus*, *Fusarium, Metarhizium,* and *Penicillium* genera (Figure 5), but absent in Saccharomycotina. In some aspergilli, two similar clusters are present (*e.g.* in *A. oryzae)* with the second cluster showing structural conservation and higher sequence homology to the cluster found in *Metarhizium* and *Penicillium*; indicative of a more recent horizontal gene transfer event. The cluster is absent in *A*.

**Figure 5.**
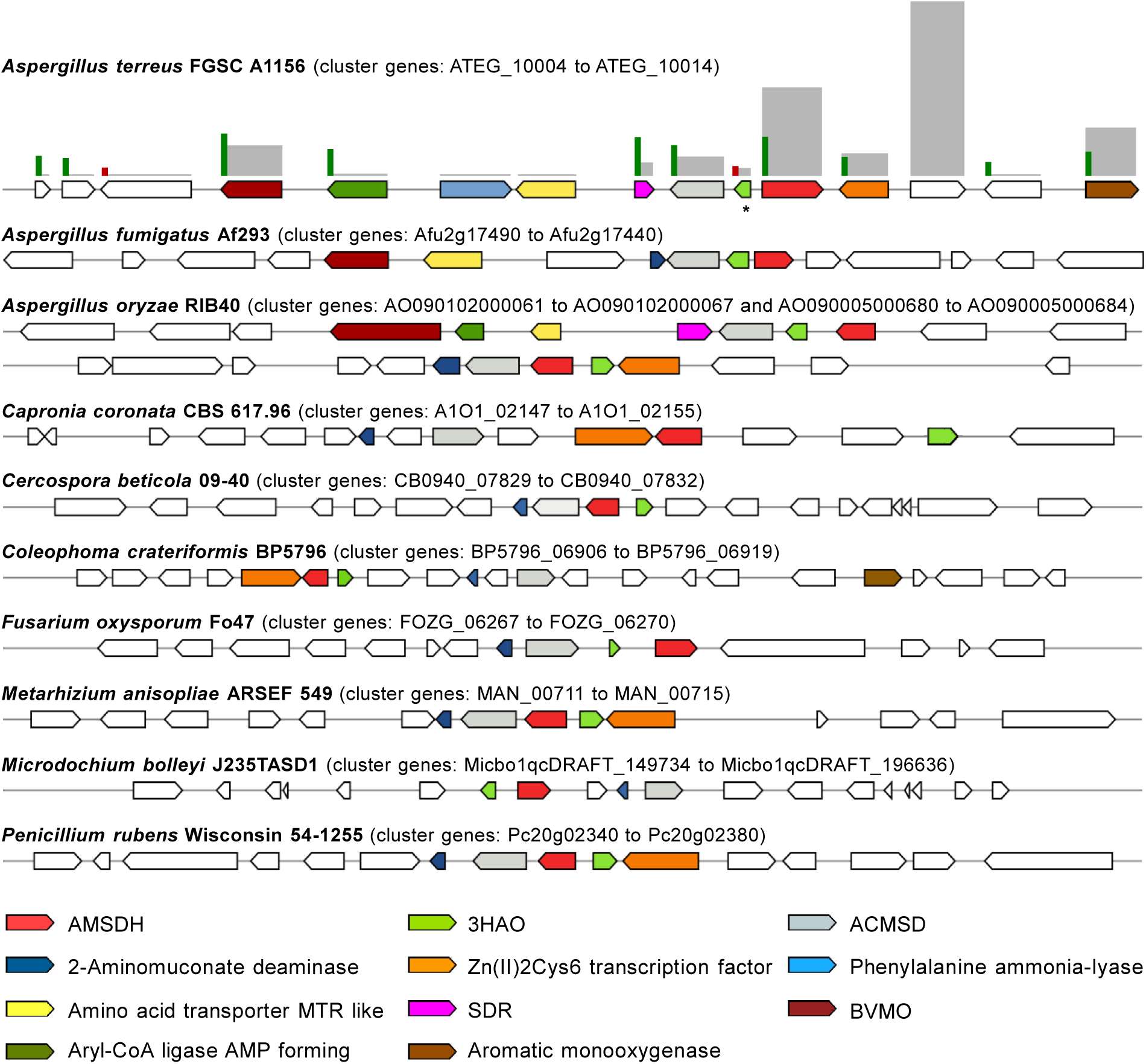
Representation of the 3-hydroxyanthranilate catabolic gene cluster in some Ascomycota. The cluster includes the following genes: 3-hydroxyanthranilate-3,4-dioxygenase (3HAO), 2-amino-3-carboxymuconate-6-semialdehyde decarboxylase (ACMSD), 2-aminomuconate semialdehyde dehydrogenase (AMSDH) and a novel putative 2-aminomuconate deaminase (AMDA, 2-hydroxymuconate-forming) genes, except in *A. terreus* that does not contain the last gene. Some aspergilli syntenic genes, namely a Baeyer-Villiger monooxygenase (BVMO), a short-chain dehydrogenase/reductase (SDR) and an amino acid transporter, are also represented. Clusters were identified within annotated genomes through sequence homology searches (MultiGeneBlast software). Genes within the cluster are represented using a color code and asterisk indicates a non-annotated gene. The differentially expressed genes (RNA-seq) regulation and abundance is represented by scale sized bars and color (upregulated in green, downregulated in red and RPKM in grey).

*niger* and *F. graminearum,* where L-tryptophan catabolism proceeds via anthranilate, 2,3-dihydroxybenzoate and the catechol branch of the 3-oxoadipate pathway (47, 48). A fourth gene of the YjgF/YER057c/UK114 family (IPR006175) is recurrently present in the cluster, but not in *A. terreus* (Figure 5). Proteins of this diverse family are expected to have various metabolic roles extending from endoribonuclease activity and 2-aminomuconate deaminase to enamine/imine deaminase of the branched-chain amino acid (BCAA) biosynthesis pathway (49). Its orthologous gene in *A. terreus* (ATEG_05745) was found moderately expressed and upregulated in salicylate (Figures 3 and 4), hence possibly it encodes for a 2-aminomuconate deaminase. It has been shown that this deamination reaction may also occur non-enzymatically forming the same products: 2-hydroxymuconate and its tautomer 2-oxo-3-hexenedioate (50). In line with this, herein we noticed that a putative 4-oxalocrotonate tautomerase gene (ATEG_05150) was found upregulated, suggestive that the encoded protein (IPR014347) may mediate 2-oxo-3-hexenedioate formation (Figure 3). This compound may undergo further reduction to 2-oxoadipate or decarboxylation and hydration to 4-hydroxy-2-oxopentanoate. The subsequent decarboxylation of 2-oxoadipate to glutaryl-CoA by the oxoglutarate dehydrogenase complex (encoded by ATEG_03911 and ATEG_09000) could not be verified by the RNA-seq data. We observed that the TCA cycle in *A. terreus* (and in *A. nidulans*) was found highly downregulated in salicylate media, and the predicted glutaryl-CoA dehydrogenase (ATEG_08866) was also found downregulated (Table S6).

Surprisingly, several genes of the leucine catabolic pathway were found extremely expressed and upregulated in salicylate, namely the E1 and E2 components of the branched-chain alpha-keto acid dehydrogenase complex (ATEG_03862, ATEG_06635 and ATEG_09087), isovaleryl-CoA dehydrogenase (ATEG_06574) and 3-methylcrotonyl-CoA carboxylase (ATEG_06573 and ATEG_06576). However the remaining genes of the leucine catabolism were downregulated or only moderately expressed (Figure S6 and Table S9). Taken together, it remains unclear if those genes that underwent upregulation have a role in the catabolism of the branched-chain 2-oxoacid 4-hydroxy-2-oxopentanoate, namely in its decarboxylation to 3-hydroxybutyryl-CoA. The predicted 3-hydroxybutyryl-CoA dehydrogenase (ATEG_02736) and the downstream acetyl-CoA acetyltransferase (ATEG_00012) were found downregulated, although highly expressed (Figure 3). Importantly, the catabolism of branched chain amino acids was previously shown to be upregulated in conditions where 2-hydroxymuconate was produced (51).

In aspergilli the 3-hydroxyanthranilate catabolic gene cluster shows synteny to genes likely coding in the peripheral pathways (Figure 5), some of which upregulated by salicylate (ATEG_10004, ATEG_10005 and ATEG_10008). Two of these genes, encoding for an Aryl-CoA ligase (AMP forming) (ATEG_10005) and a SDR (ATEG_10008), show low homology to characterized proteins. However, the FAD-binding monooxygenase ATEG_10004 gene which underwent here major expression, contains the sequence motif of a Baeyer-Villiger monooxygenase (52), a distinct class of flavoproteins that catalyze the insertion of an oxygen atom in a C–C bond of a ketone forming an ester. In addition, another FAD-binding monooxygenase gene (ATEG_10014), which was observed also greatly expressed and upregulated, shows discrete homology to the salicylate 1-monooxygenase (decarboxylating) gene. Interestingly, this monooxygenase gene is located in the vicinity of the 3-hydroxyanthranilate catabolic cluster.

The six orthologous genes of the *A. nidulans* nicotinate-inducible gene cluster (43) are part of a highly expressed and upregulated gene cluster of nine genes in *A. terreus* grown in salicylate. The three genes which were not previously annotated: two FAD-dependent monooxygenase genes (ATEG_08614 and ATEG_08615b) and a SDR (ATEG_08615a) showed major upregulation in *A. terreus* (Figure 6). ATEG_08614 shows discreet homology to the bacterial 6-hydroxynicotinate 3-monooxygenase (Q88FY2) that forms 2,5-dihydroxypyridine as part of nicotinate catabolism, but also to 3-hydroxybenzoate 6-monooxygenase (Q9F131) and orsellinic acid 1-monooxygenase (J4VWM7). ATEG_08615b is a phenol hydroxylase (IPR012941). Their major co-regulation in salicylate is suggestive of participation in the nicotinate metabolism. In addition, its cluster preservation in aspergilli and across Ascomycota, being however divided in two clusters in the genome of *A. nidulans* (Figure 6), strengthens the hypothesis of their role in the nicotinate metabolism. Expanding the cluster analysis across numerous fungal species (Figure 6) revealed the presence of two additional co-localized genes, a deacetylase gene (ATEG_05673) and an amidase gene (ATEG_05674) that are not located in the nicotinate-inducible cluster in the majority of aspergilli including in *A. terreus* and *A. nidulans*. In salicylate, only the deacetylase gene (ATEG_05673) was found highly expressed and upregulated, suggestive of a role in the nicotinate metabolism. In general, the nicotinate-inducible genes of *A. nidulans* showed to be somehow upregulated in salicylate media, though to a much lesser extent compared to *A. terreus* (Figure 6). This together with the moderate co-upregulation of the single copy 3-hydroxyanthranilate dioxygenase (3HAO gene, AN11252 orthologous to ATEG_00088) in *A. nidulans* (Table S8) pushes forward the idea of an alternative pathway for the metabolism of aromatic compounds. The existence of such metabolic route possibly explains why the metabolism of the quorum sensing molecule phenylethyl alcohol yielded the pyridine ring of NADH and NADPH in *Candida albicans* (53).

**Figure 6.**
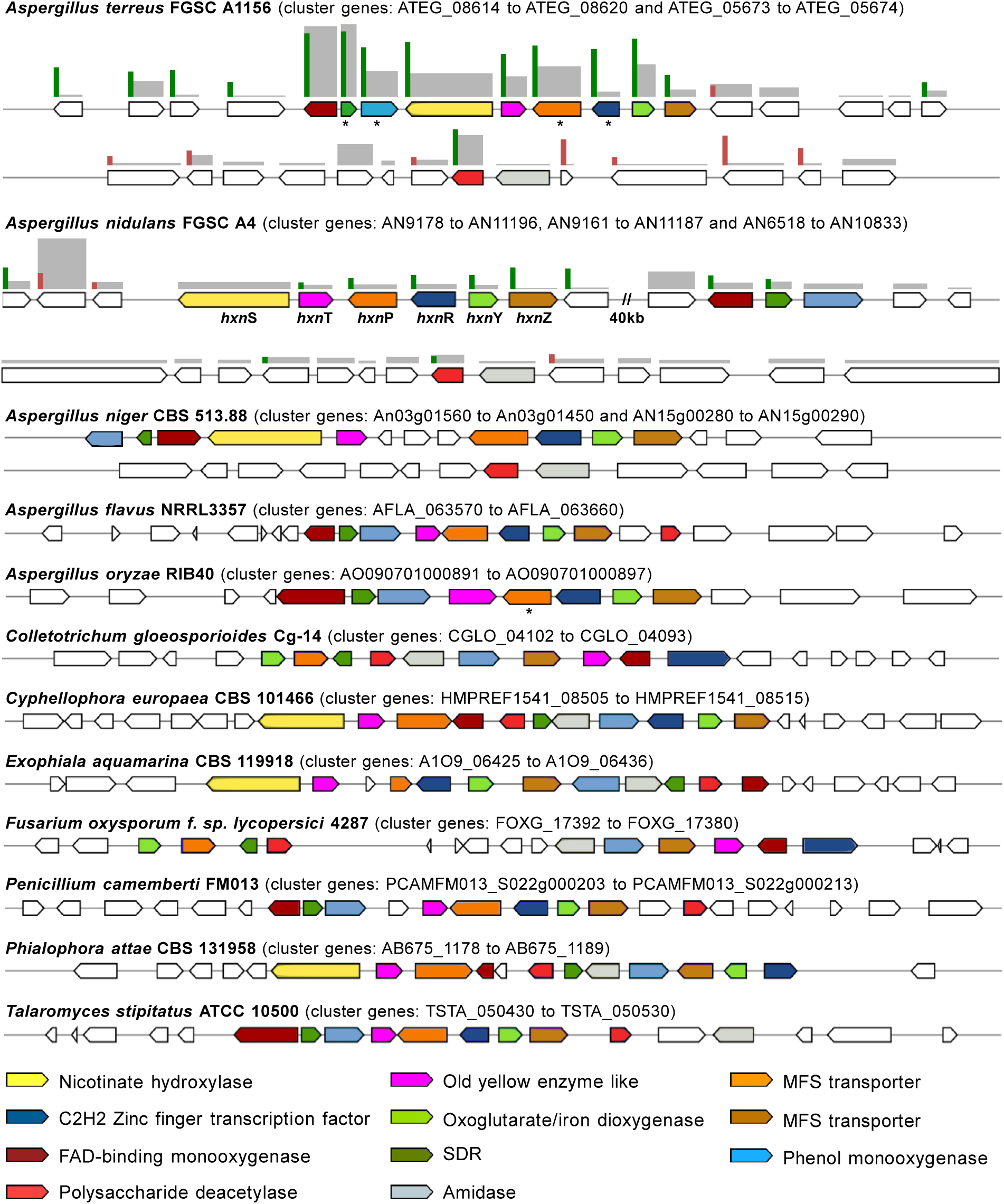
Representation of the nicotinate gene cluster in Ascomycota. The cluster includes the following genes: nicotinate hydroxylase (*hxn*S) and the remaining five nicotinate-inducible orthologous genes of *Aspergillus nidulans* (*hxn*T, *hxn*P, *hxn*R, *hxn*Y, *hxn*Z) as well as five additional genes that are recurrently clustered in Ascomycota genomes. Clusters were identified within annotated genomes through sequence homology searches (MultiGeneBlast software). Genes within the cluster are represented using a color code and asterisk indicates a poorly or non-annotated gene. Double slash indicates a gap within the same chromosome. The differentially expressed genes (RNA-seq) regulation and abundance is represented by scale sized bars and color (upregulated in green, downregulated in red and RPKM in grey).

The gentisate-*like* cluster contains a FAD-binding monooxygenase gene (ATEG_06711) that did not undergo initial upregulation (Figure 4). A similar trend was noticed for the gentisate 1,2-dioxygenase orthologous gene (ATEG_06712). These observations exclude their foreseen involvement in the hydroxylation of salicylate to gentisate and its subsequent ring-cleavage. We have identified another gentisate 1,2-dioxygenase gene (ATEG_07116-17), which is not currently annotated (Table S2) that showed no altered expression levels during cultivation in salicylate (Figure 4). Both *A. terreus* and *A. nidulans* were unable to grow at the expenses of gentisate (5 or 10 mM) regardless of capacity to co-metabolize gentisate (depletion of 5 mM in less than 3 days with acetate pre-grown cultures). In addition, gentisate may also be transformed to hydroxyquinol by salicylate 1-monooxygenase (54). However, the expression analysis of the two hydroxyquinol 1,2-dioxygenase genes (ATEG_00629 and ATEG_07465), orthologous of the two recently biochemically characterized genes of *A. niger* (11), as well as of the predicted maleylacetate reductase gene (ATEG_03722), showed no great alteration (Figure 4 and Table S9).

The homogentisate pathway for the catabolism of phenylalanine and tyrosine (chorismate as common precursor) was found mostly upregulated in salicylate (Figures 3 and 4). Several routes can be used for the metabolism of their corresponding precursors, phenylpyruvate and 4-hydroxyphenylpyruvate, likely involving a small group of enzymes capable to mediate analogous reactions (55). In line with this idea, we observed that a pyruvate decarboxylase PDC1-*like* gene (ATEG_06881) and several alcohol and aldehyde dehydrogenase genes (Table S9) that may code for enzymes mediating production of fusel alcohols and acids (*e.g.* tryptophan to tryptophol) (56), underwent up regulation in salicylate. These observations are suggestive that the Ehrlich pathway was likely upregulated. This is reinforced by concurrent upregulation of phenylacetate 2-hydroxylase and 3-hydroxyphenylacetate 6-hydroxylase orthologous genes (ATEG_09785 and ATEG_08499) that encode for monooxygenases of the fusel acids phenylacetate and 4-hydroxyphenylacetate or their products.

## Conclusions

Analysis of the transcriptome associated with the catabolism of salicylate in two species of *Aspergillus* showed its intimate relationship with the secondary metabolism, particularly obvious for metabolites requiring action of aromatic prenyltransferases. Exogenous aromatic compounds and their metabolites can provide the building blocks for the synthesis of many secondary metabolites, as well established *e.g.* for penicillin (*i.e.* phenylacetate) (55). We observed that the metabolism of salicylate yield gentisate in *A. terreus*, likely also producing gentisyl alcohol, an intermediate of secondary metabolites such as patulin (57, 58). This observation suggests that salicylate (or gentisate) usage as carbon sources may improve the production of target metabolites. Similarly, salicylate upregulated several catabolic pathways in *A. terreus,* including that of tyrosine that can also provide abundant and diverse precursors for the secondary metabolism. Tyrosine catabolic genes and secondary metabolite core biosynthetic genes showed similar regulation in penicillia (59). Understanding the relationship between salicylate and secondary metabolism in fungi may provide unexpected findings on plant-fungi interaction, specifically how fungi counteract salicylate-dependent plant defense mechanisms (60).

Collectively our data showed the high complexity of the metabolism of salicylate in *A. terreu*s with concomitant upregulation of several pathways for the catabolism of aromatic compounds at distinct timings (Figure 4). On the contrary, *A. nidulans* used only a major route (Figure S3), similar to that previously reported for many fungal species ((9) and references therein). One unexpected idiosyncrasy of *A. terreus* here observed for the first time, refers to the formation of significant amounts of gentisate in the salicylate medium, and subsequently its metabolism. The pathways used by *A. terreus* for gentisate catabolism remain uncertain. The observed upregulation of the homogentisate pathway, of the nicotinate catabolism and of the 3-hydroxyanthranilate catabolic pathway in salicylate medium could not be anticipated, deserving more attention in the near future. This idea is strengthened because in *A. nidulans* these pathways remained mostly unregulated (or some are even absent) and the formation of gentisate was also unnoticed. The joint action of the nicotinate metabolism and the 3-hydroxyanthranilate catabolic pathway in *A. terreus* should be seen as a novel route for the degradation of aromatic compounds. One of the most striking observations is that the gentisate-*like* gene cluster in this fungus is apparently not involved in the catabolism of the formed gentisate, suggestive of a yet unknown role. In addition, 2-hydroxymuconate is one possible intermediate of the 3-hydroxyanthranilate catabolic pathway, as observed elsewhere in pyrogallol catabolism (61), suggestive of a convergence of the two pathways. Naturally, this may provide a straightforward strategy for solving the assignment of the genes associated to the catabolism of 2-hydroxymuconate in fungi.

To date the catabolism of tryptophan in fungi was thought to involve anthranilate and the catechol branch of the 3-oxoadipate pathway. On the contrary, our data on *A. terreus* and evidence of the 3-hydroxyanthranilate catabolic pathway genes in some Ascomycota genomes, suggest that the tryptophan kynurenine pathway in these species shares similarity with that of higher eukaryotes. This opens the exciting possibility of using fungi as model organisms for the investigation of diseases related with tryptophan catabolism, *e.g.* study of the toxic effects of quinolinate in the progress of Huntingtońs Disease (44). The 3-hydroxyanthranilate catabolic pathway shows lower occurrence in Ascomycota genomes compared to the other central aromatic catabolic pathways. Why a shorter number of Ascomycota possess this catabolic plasticity, raises intriguing questions about its role and evolutionary traits, especially as some aspergilli have two similar 3-hydroxyanthranilate catabolic gene clusters.

Finally, the sustainable bio-based production of organic acids requires usage of renewable non-food biomass, with lignin moving towards the pole-position. The transformation of lignin related aromatic monomers into value-added chemicals may be conceived by the so called “biological funneling” (5). To date a delocalized cytosol biosynthesis of TCA cycle intermediates (3) has been used for their production from sugars. Since the aromatic hydrocarbon catabolism in fungi occurs also at the cytosol (62, 63), it is possible to foresee engineered fungal strains as tools to convert these compounds into central intermediates of the catabolism of aromatics, and subsequently into organic acids. Our study provides another piece of the puzzle of the catabolism of aromatics in fungi, revealing new catabolic paths (some unexpected) and their putative genes and propose future paths to further dissect their functional roles, evolutionary traits and technological relevance.

## Acknowledgements

This work was financially supported: by Project LISBOA-01-0145-FEDER-007660 (Microbiologia Molecular, Estrutural e Celular) funded by FEDER funds through COMPETE2020 - Programa Operacional Competitividade e Internacionalização (POCI) and by national funds through Fundação para a Ciência e a Tecnologia (FCT); and by project “PinusResina” n° PDR2020-101-031905, funded by PDR2020 through Portugal2020. TM and CM are grateful to FCT for the working contract financed by national funds under norma transitória D.L. n.° 57/2016 and the fellowship SFRH/BD/118377/2016, respectively.

## Supplemental Table headings and Figure legends

**Table S1.**
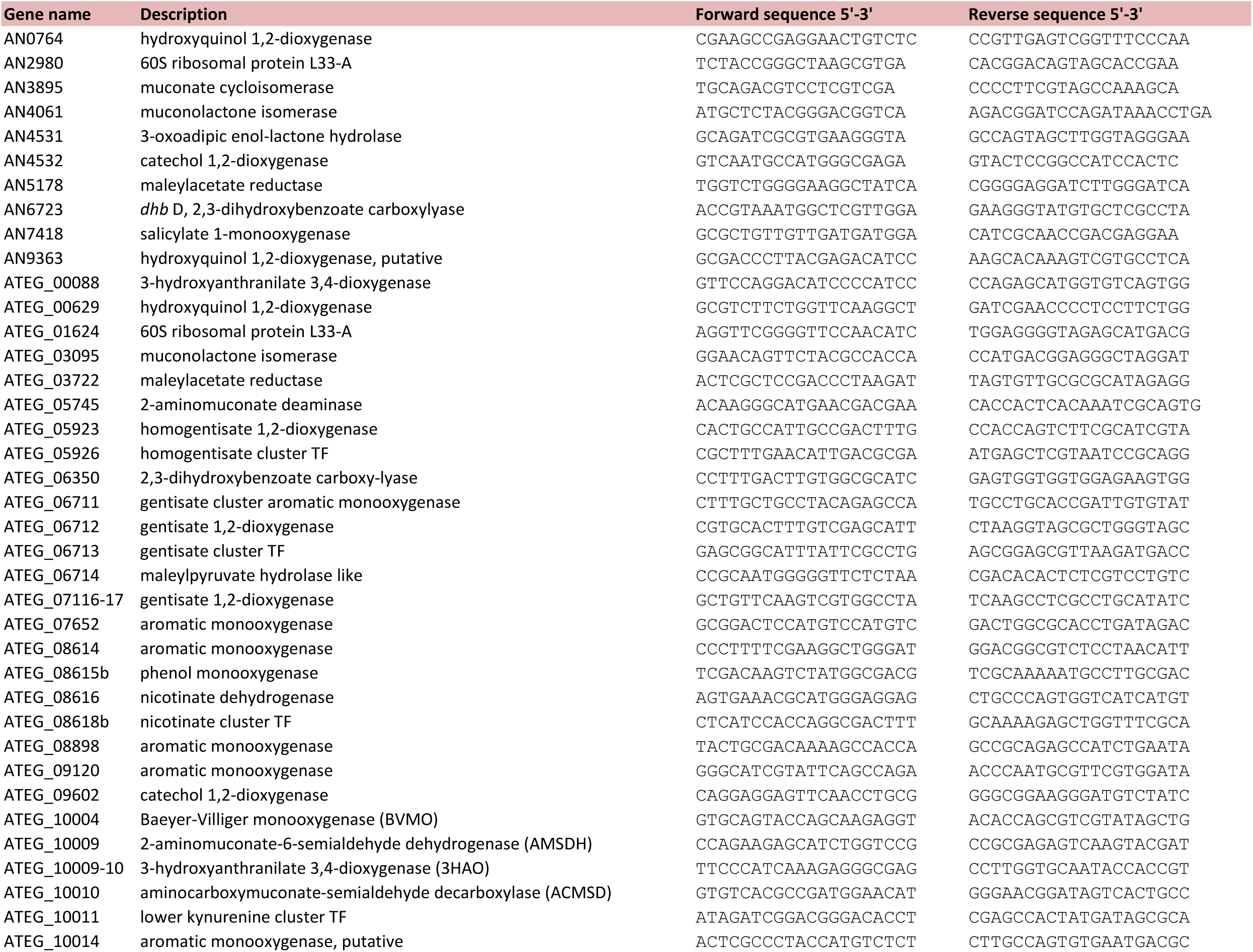
List of oligonucleotides used in RT-*q*PCR.

**Table S2.**
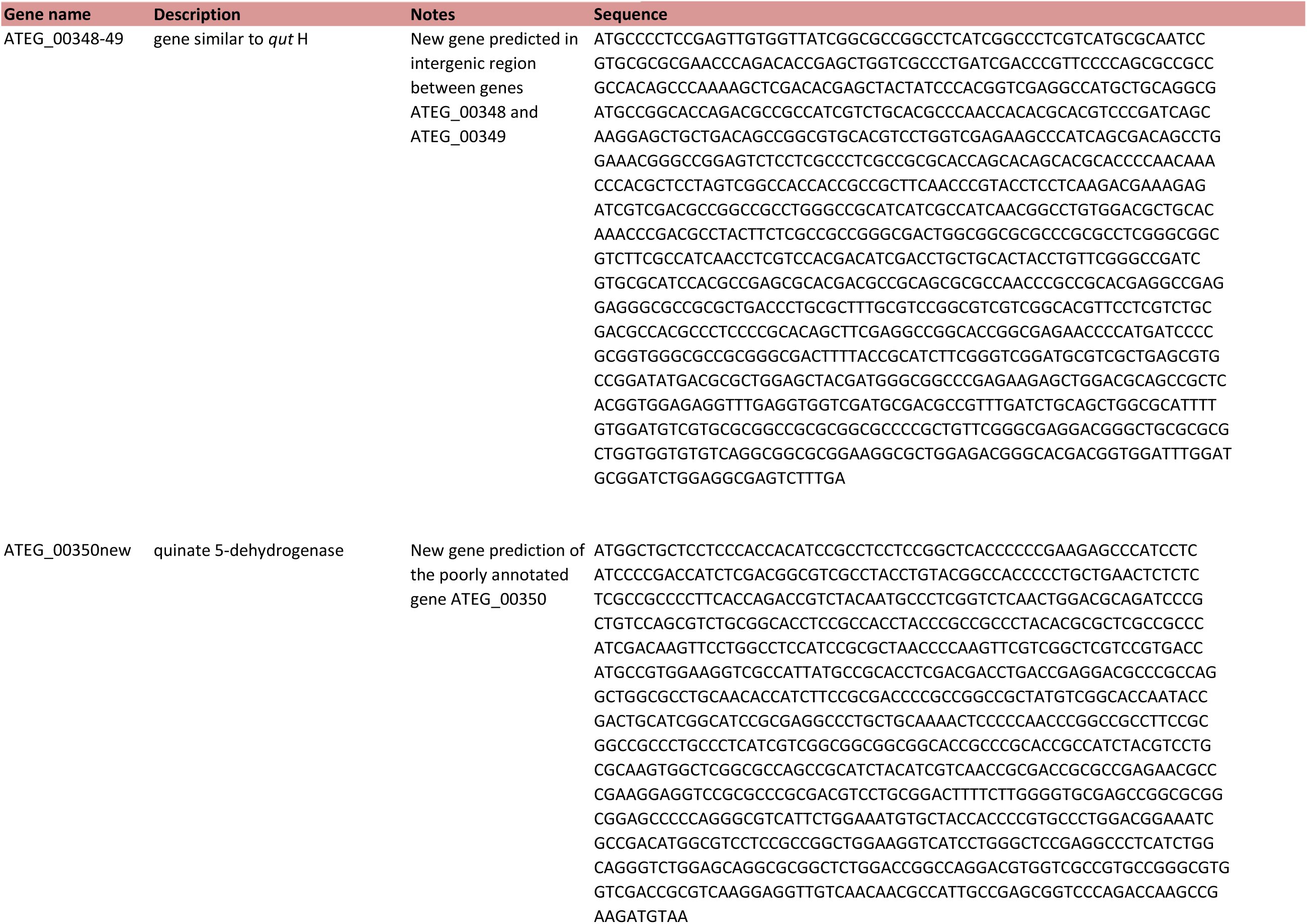
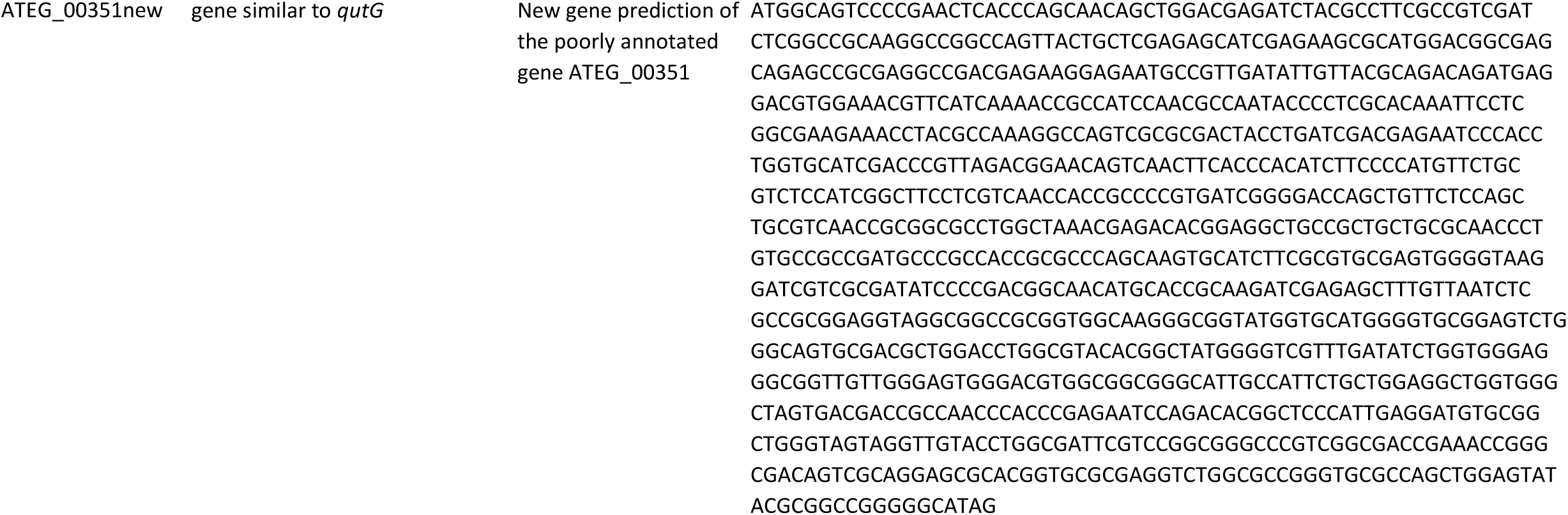
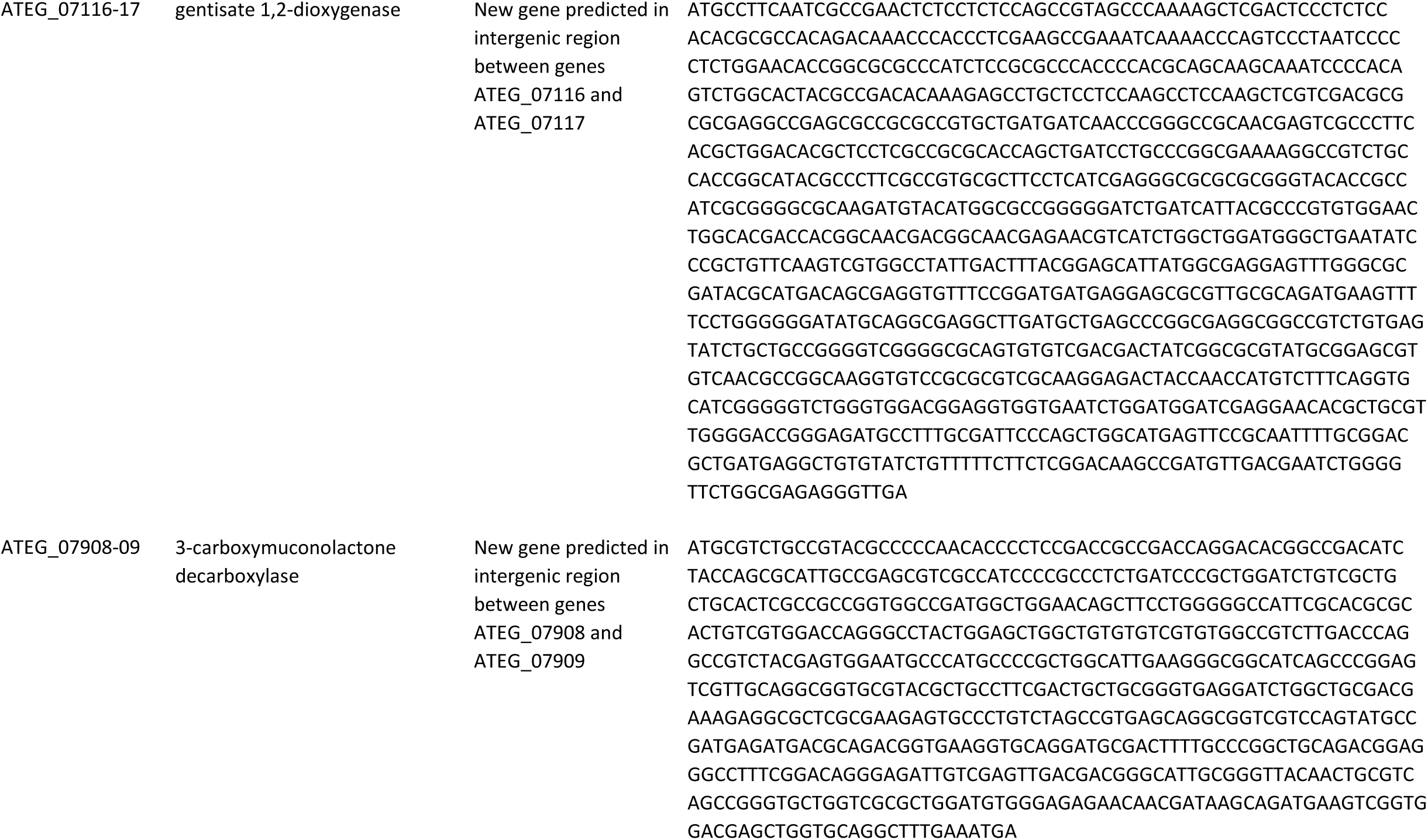
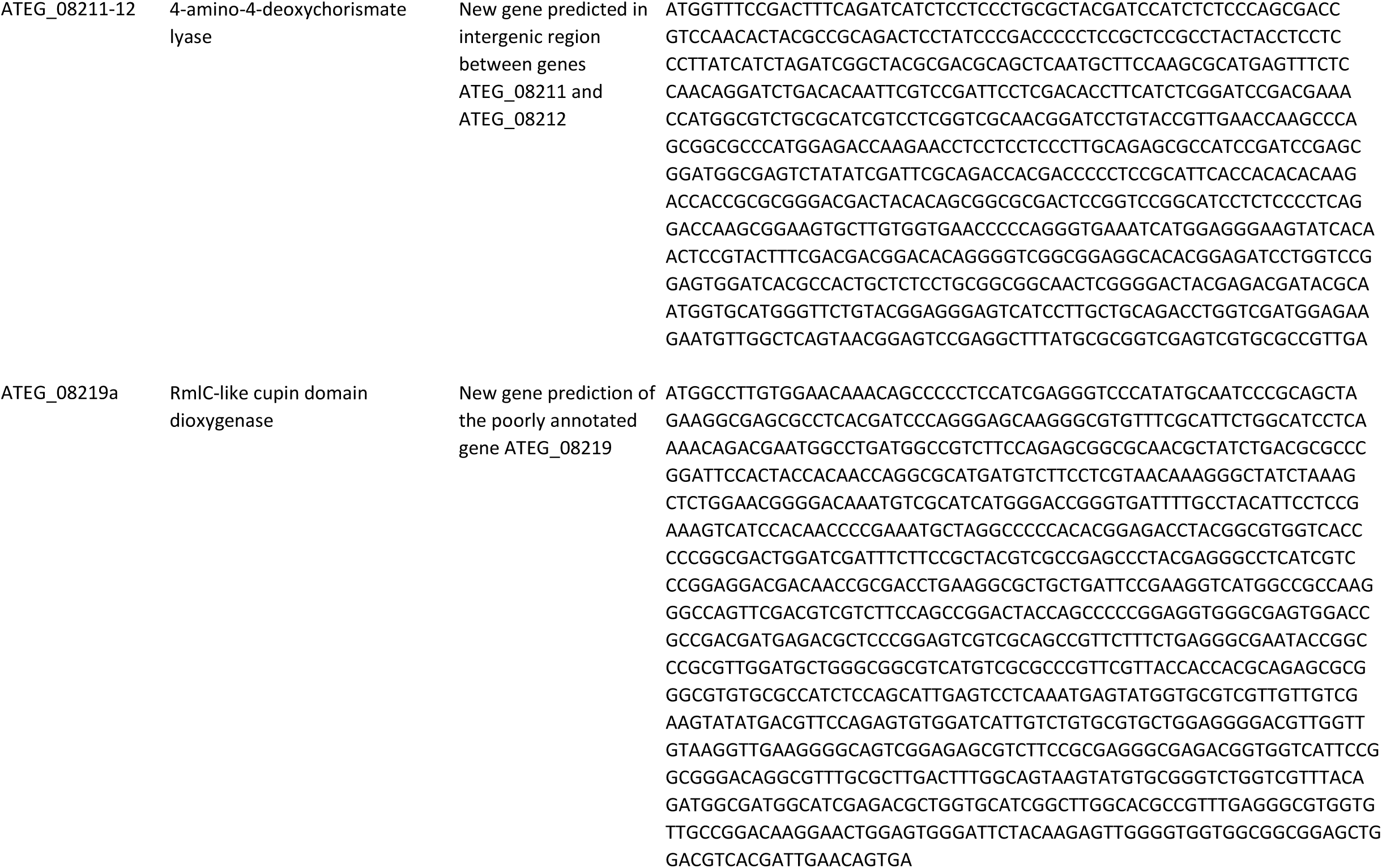
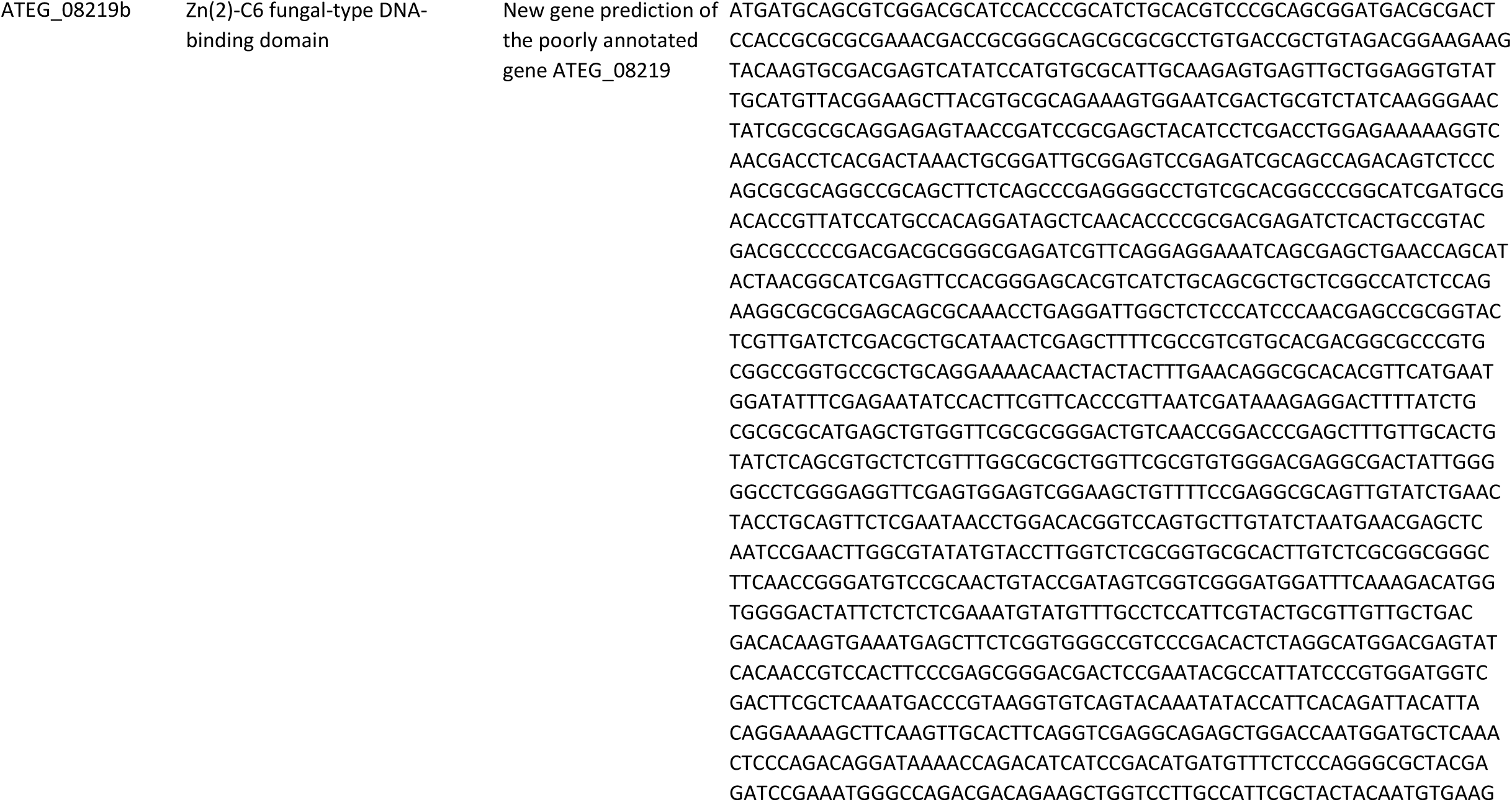
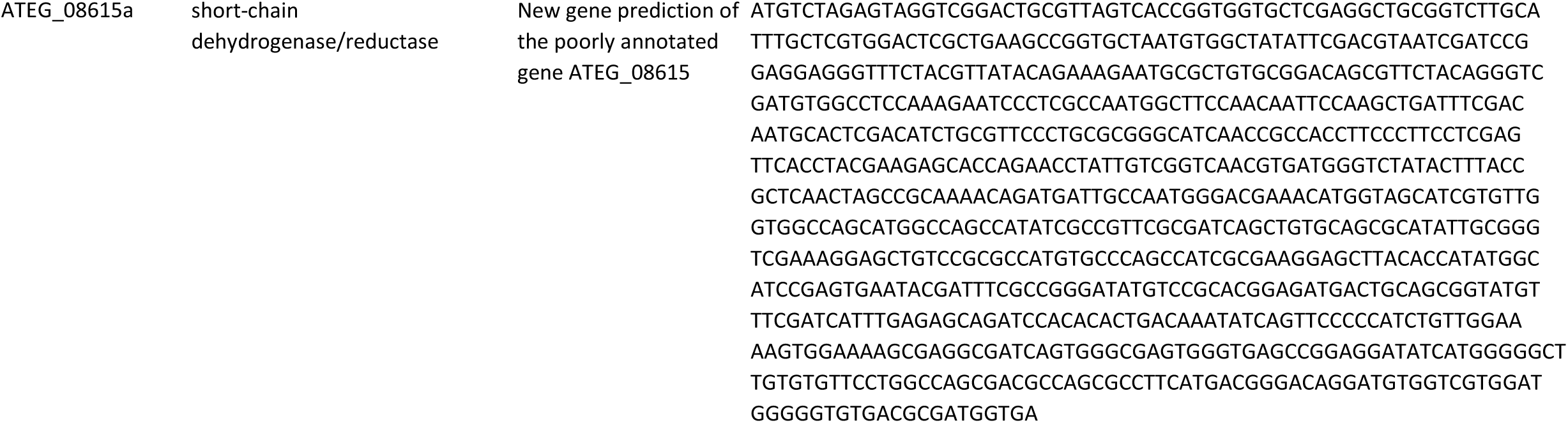
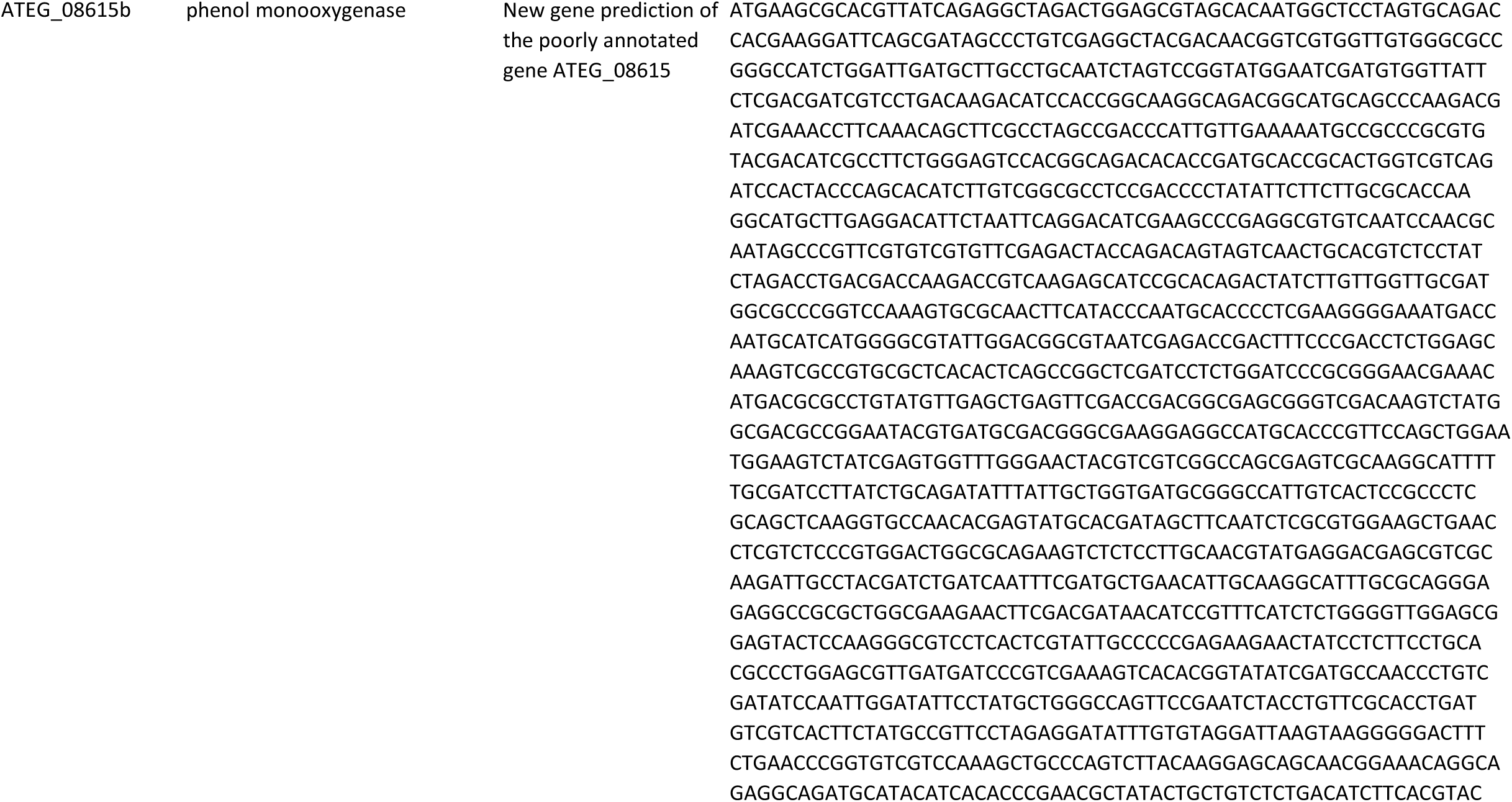
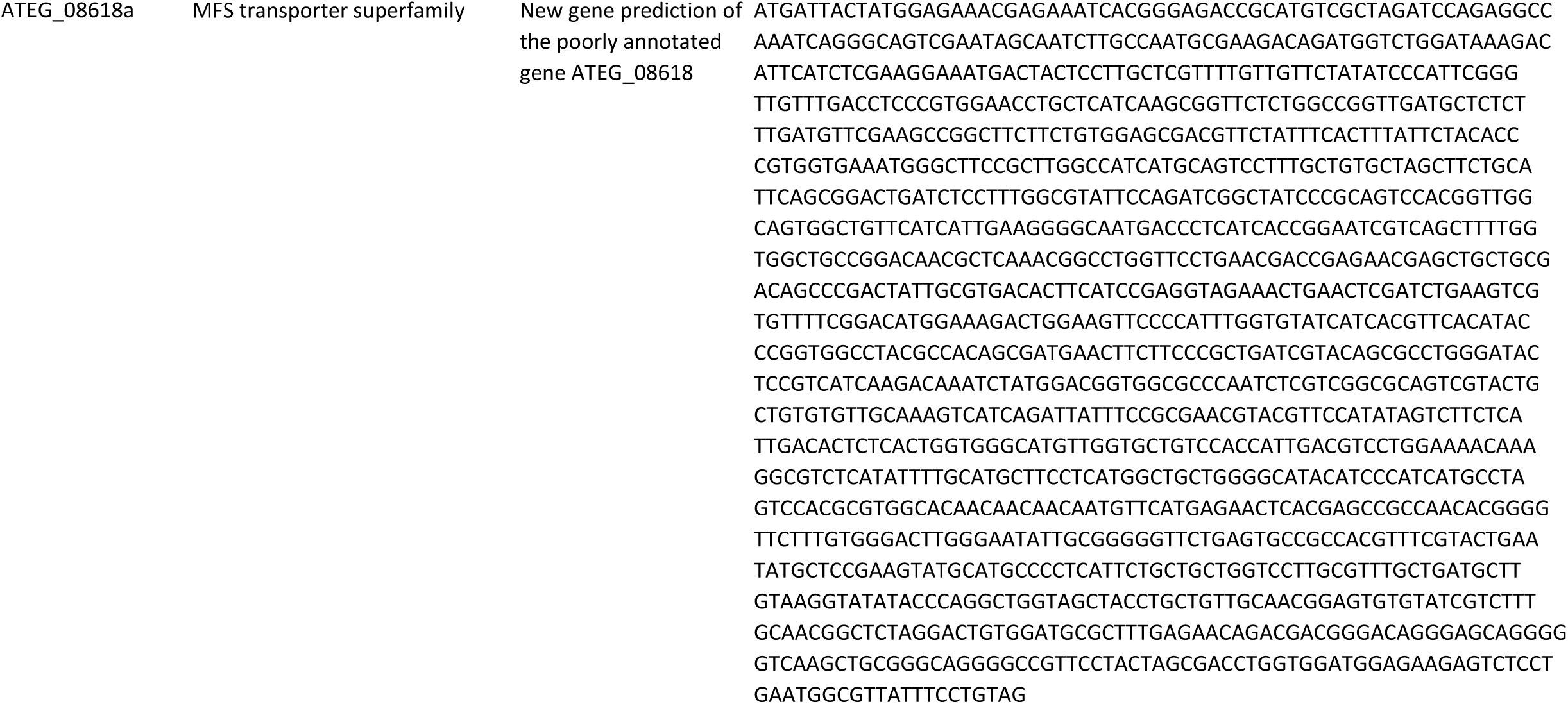
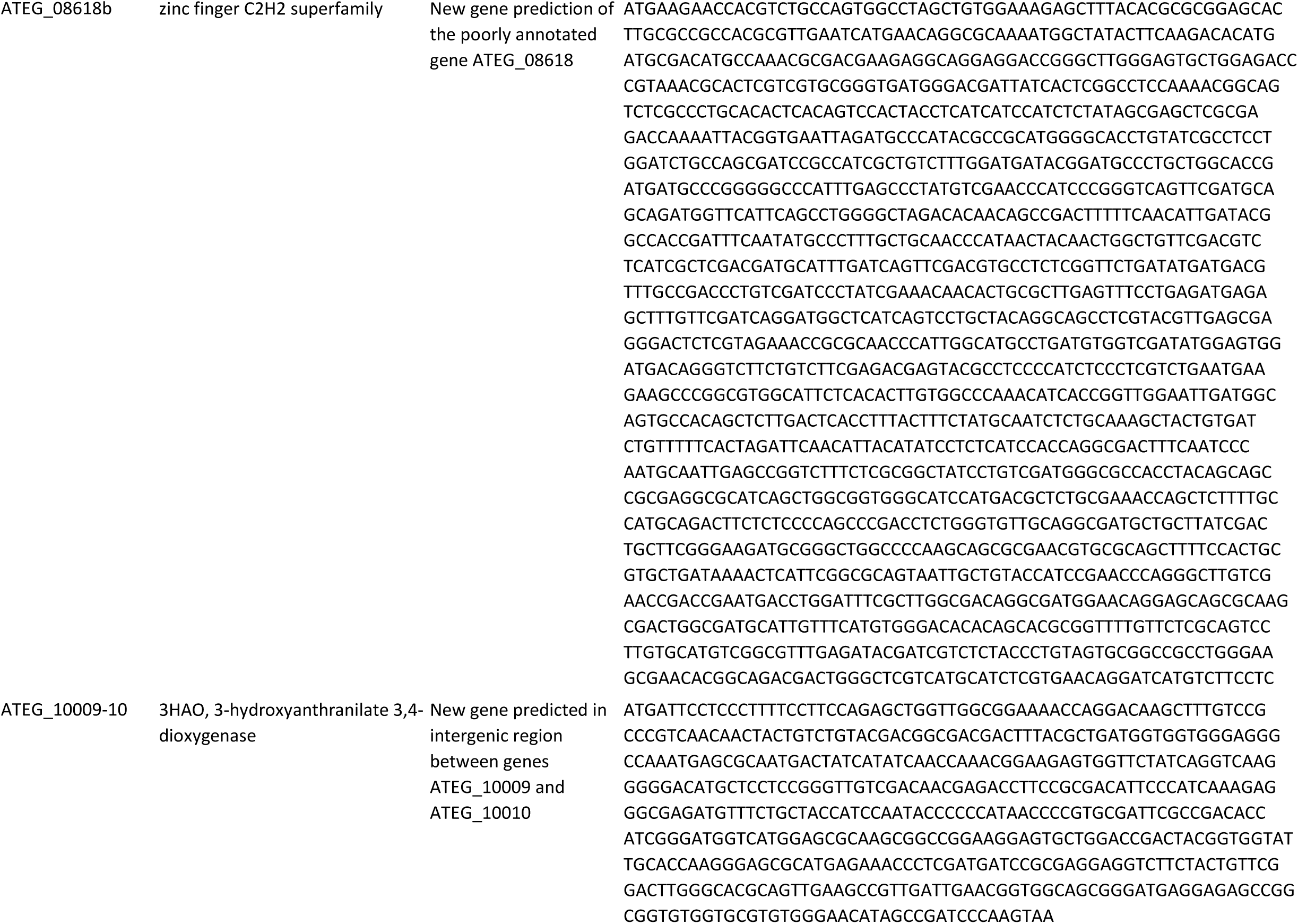
Gene sequences of predicted new genes or of poorly annotated genes.

**Table S3.**
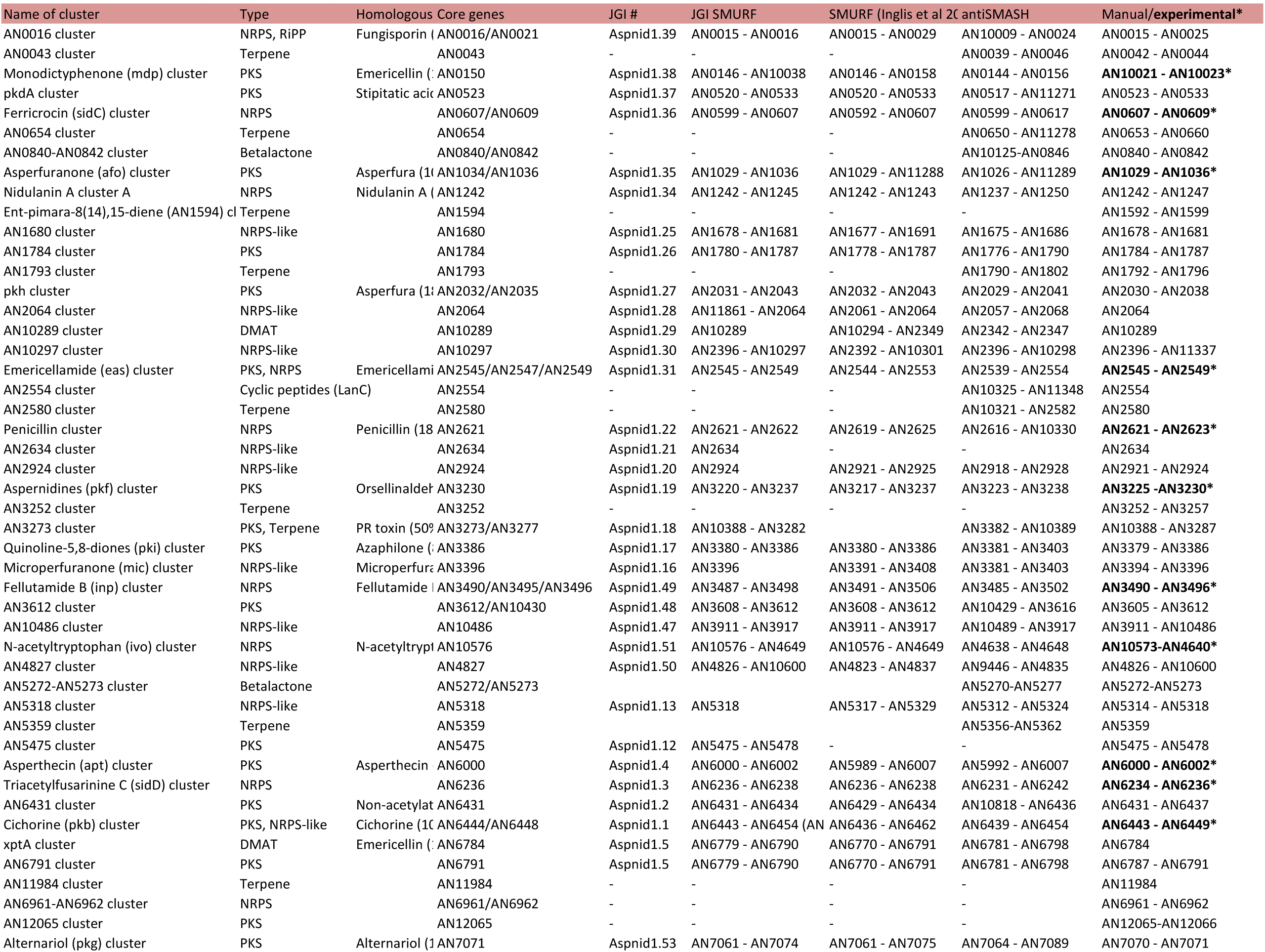
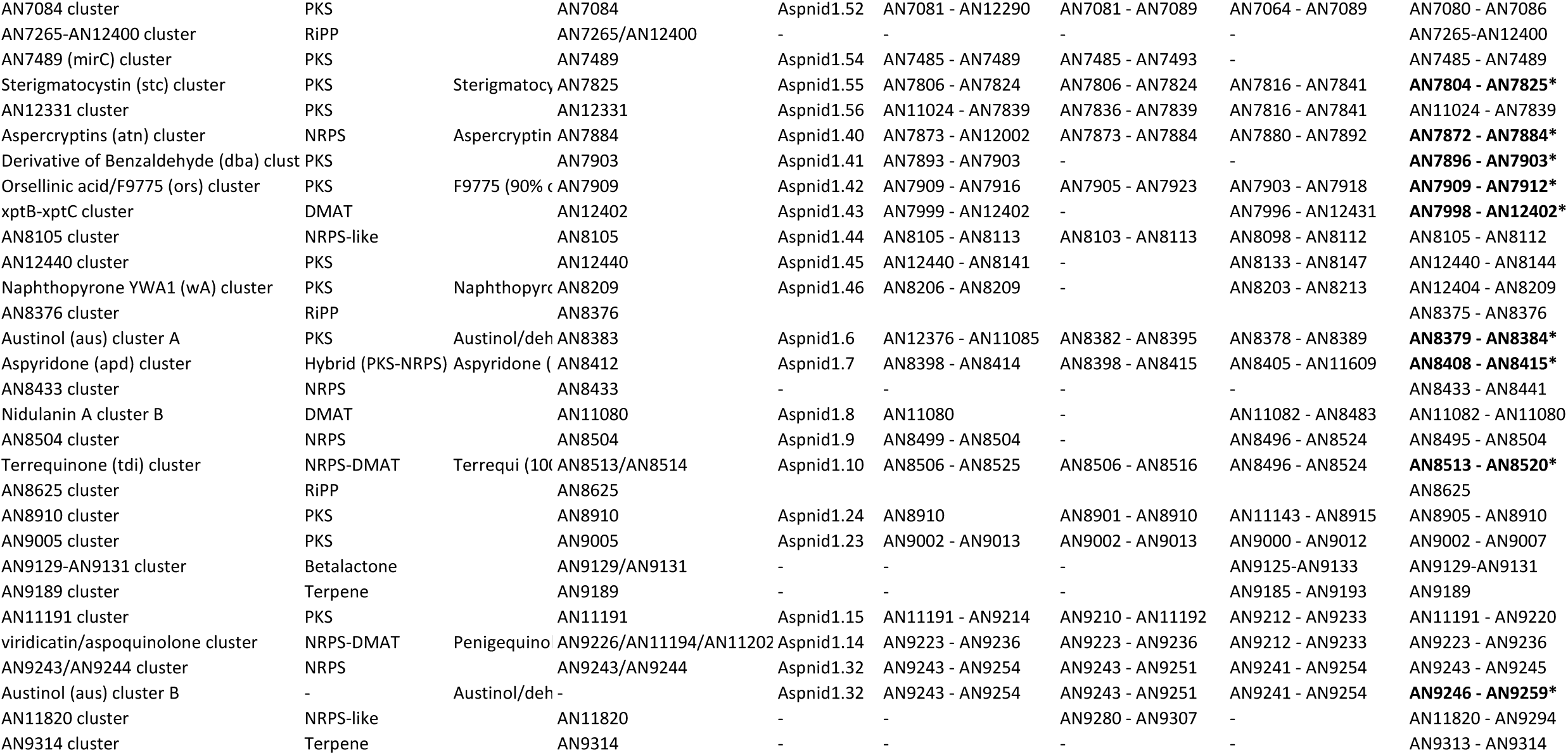
Secondary metabolite gene clusters of *Aspergillus nidulans*. Boundaries were manually curated (mostly from Inglis et al. 2013) from data obtained by using antiSMASH and from JGI MycoCosm.

**Table S4.**
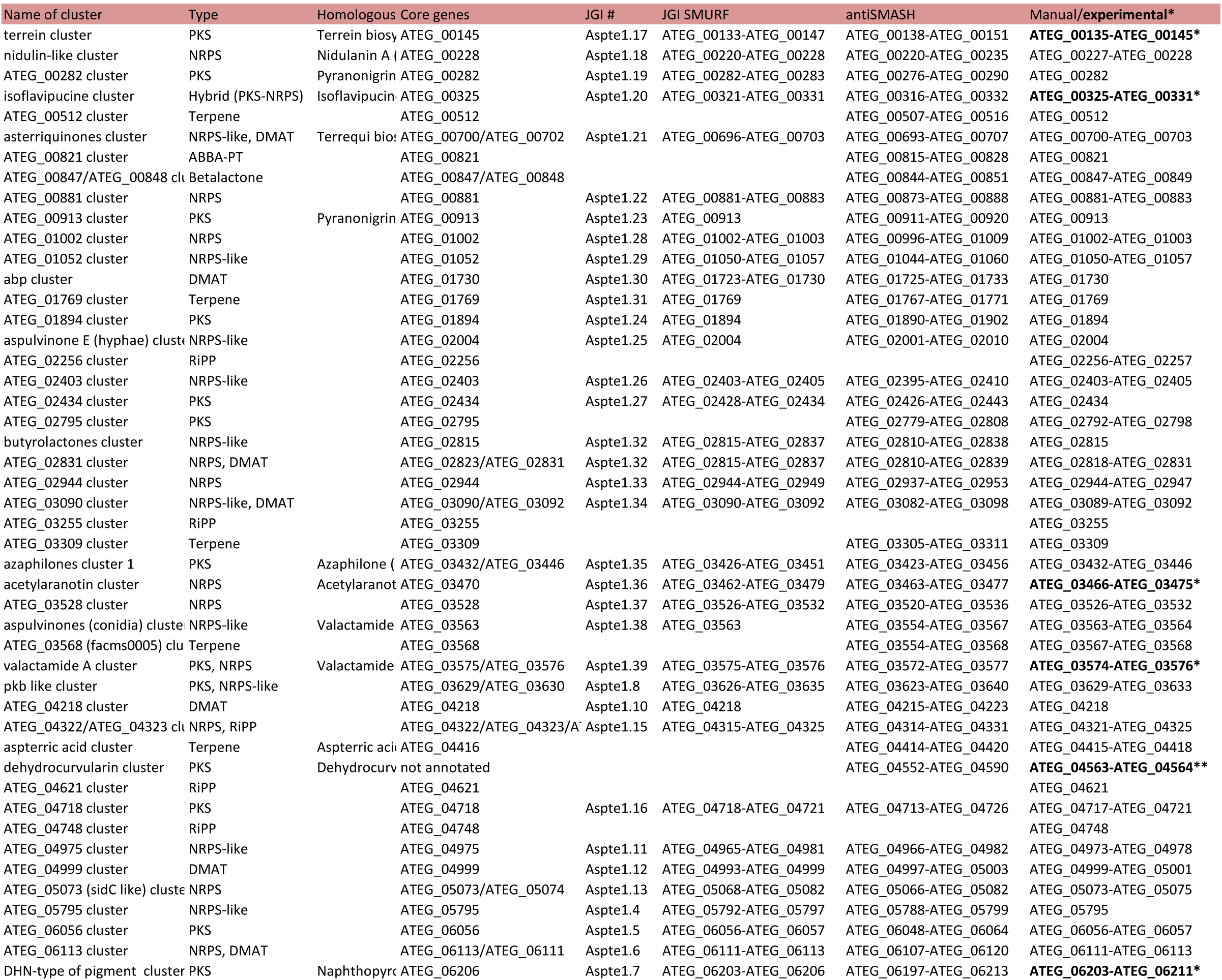
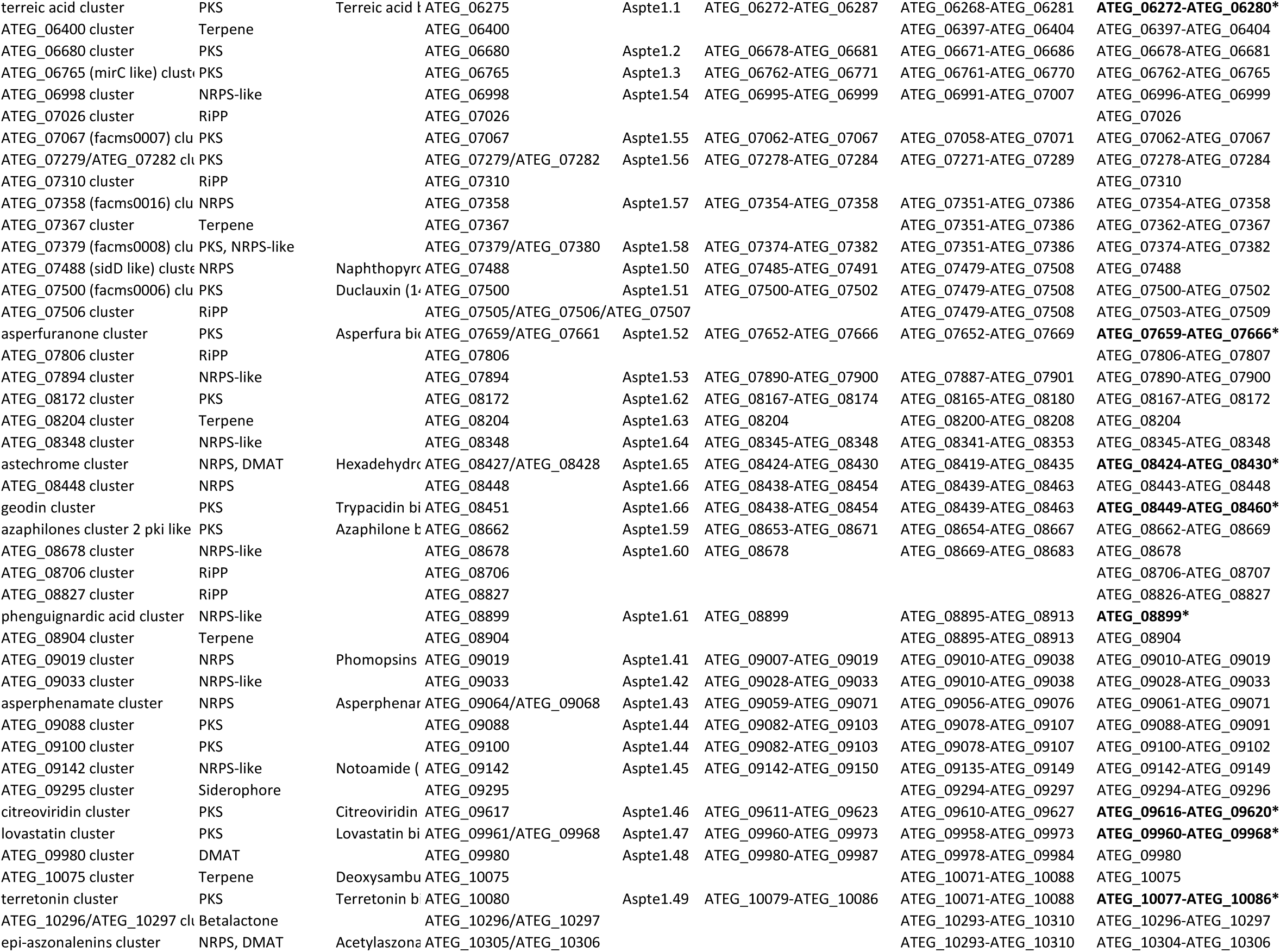
Secondary metabolite gene clusters in *Aspergillus terreus*. Boundaries were manually curated from data obtained by using antiSMASH and from JGI MycoCosm.

**Table S5.**
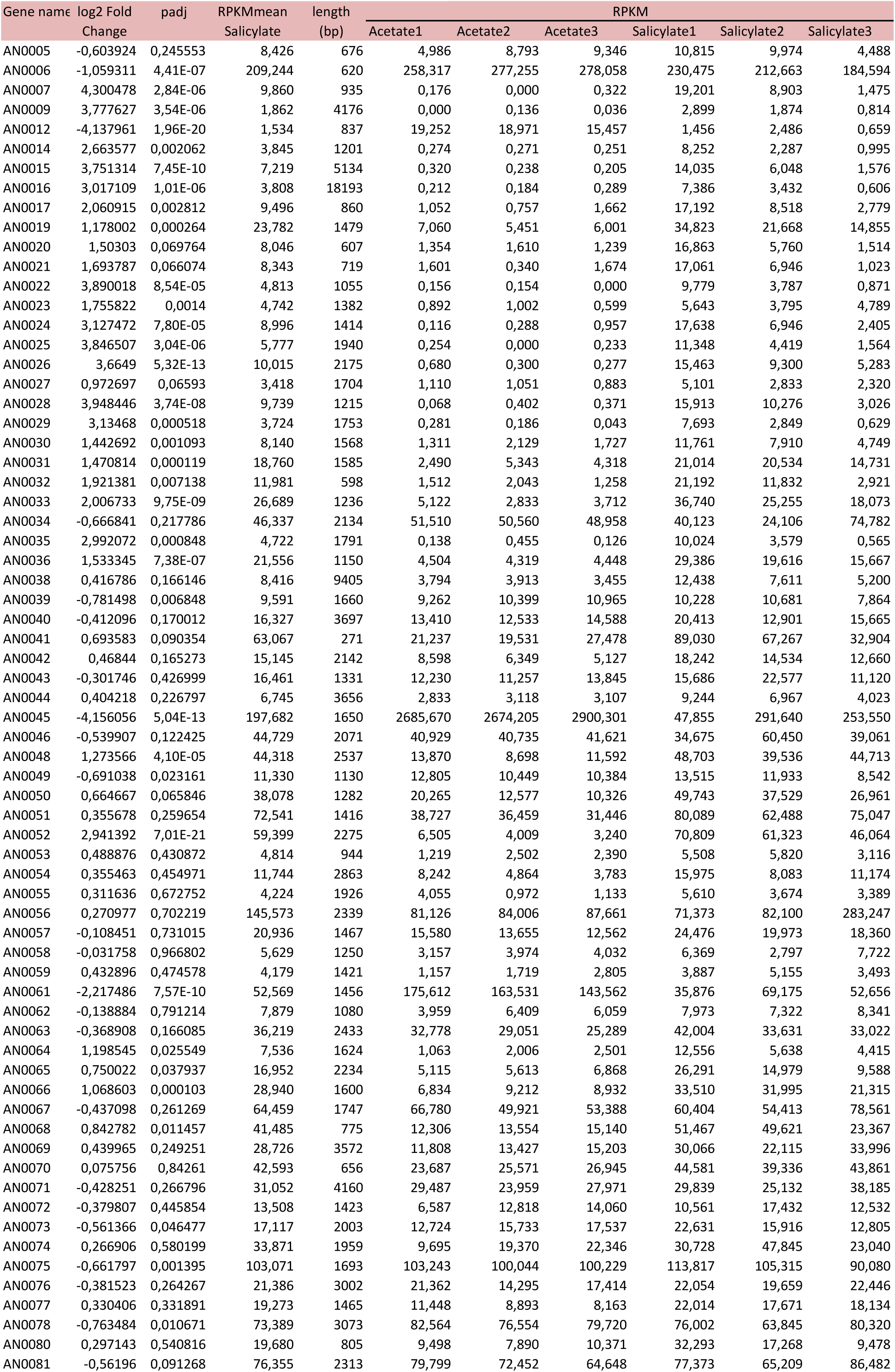
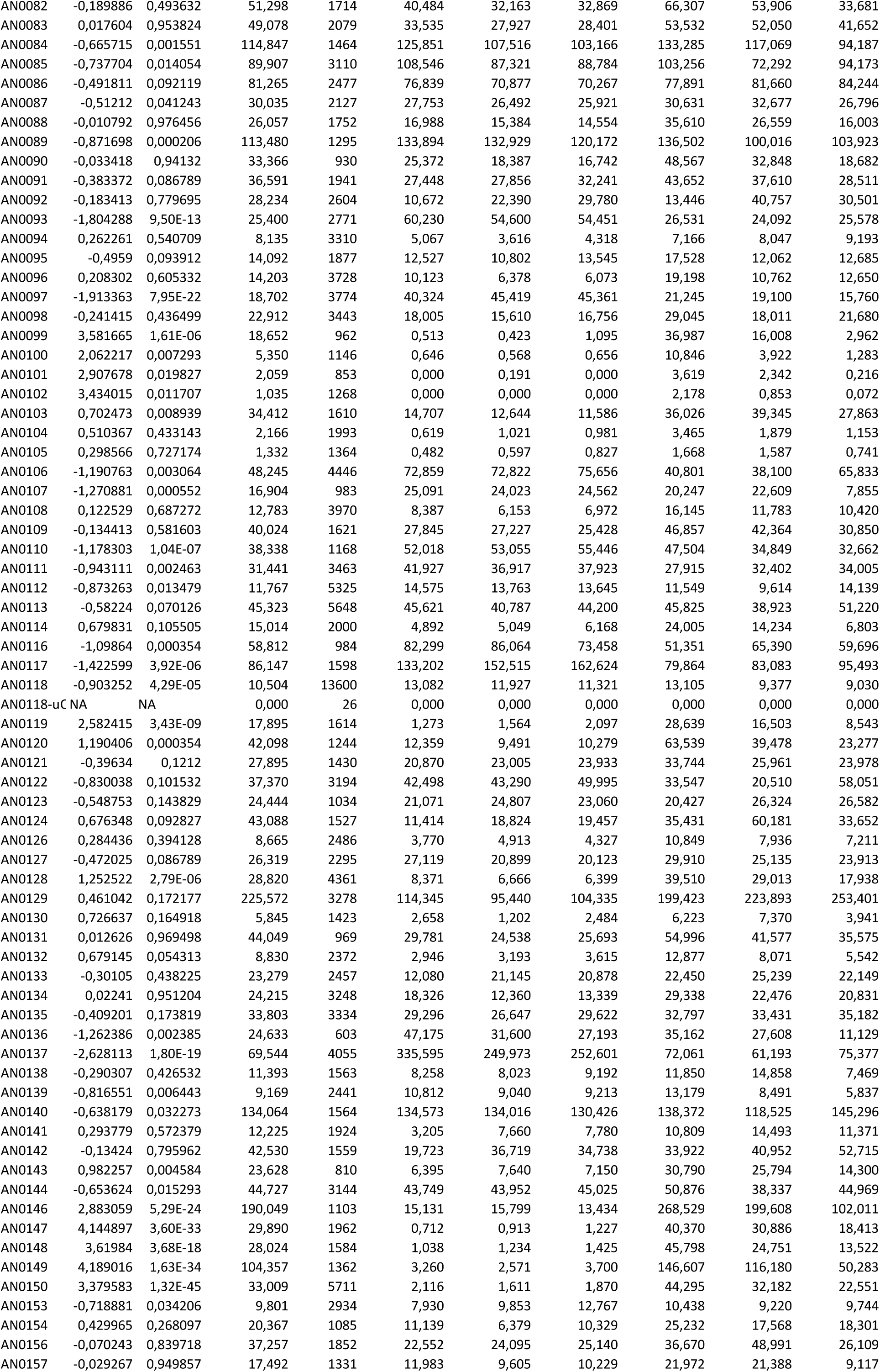
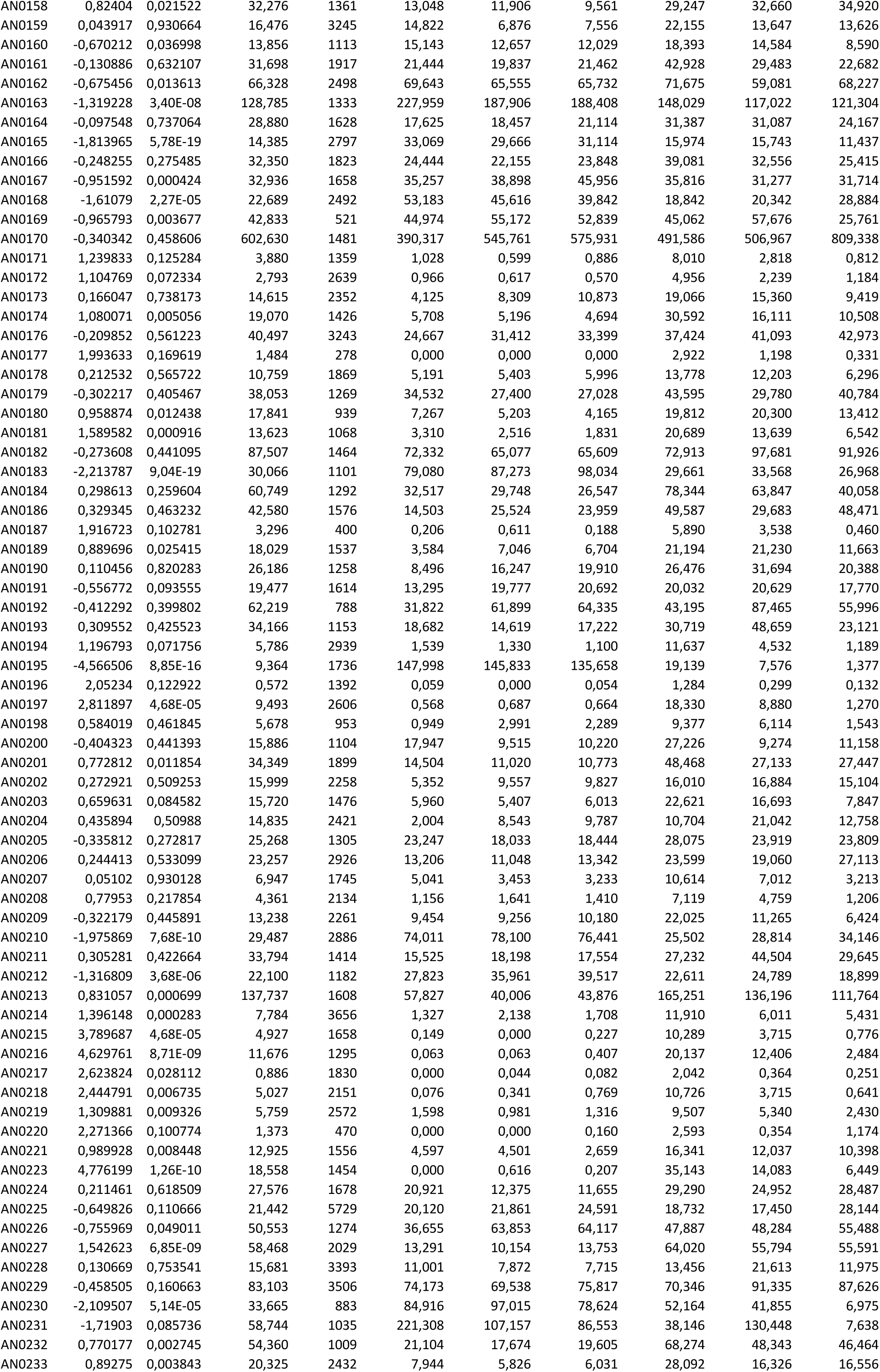
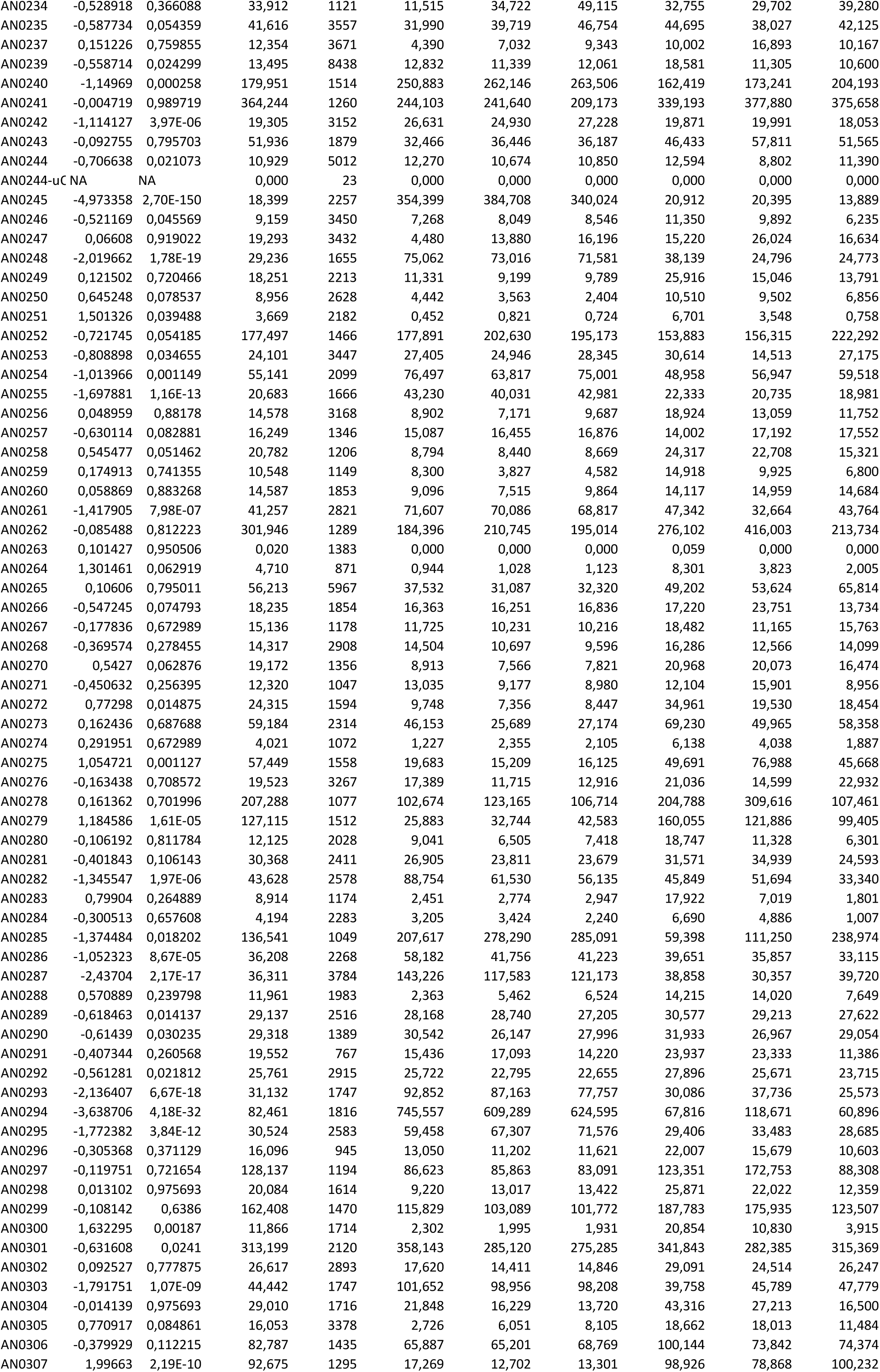
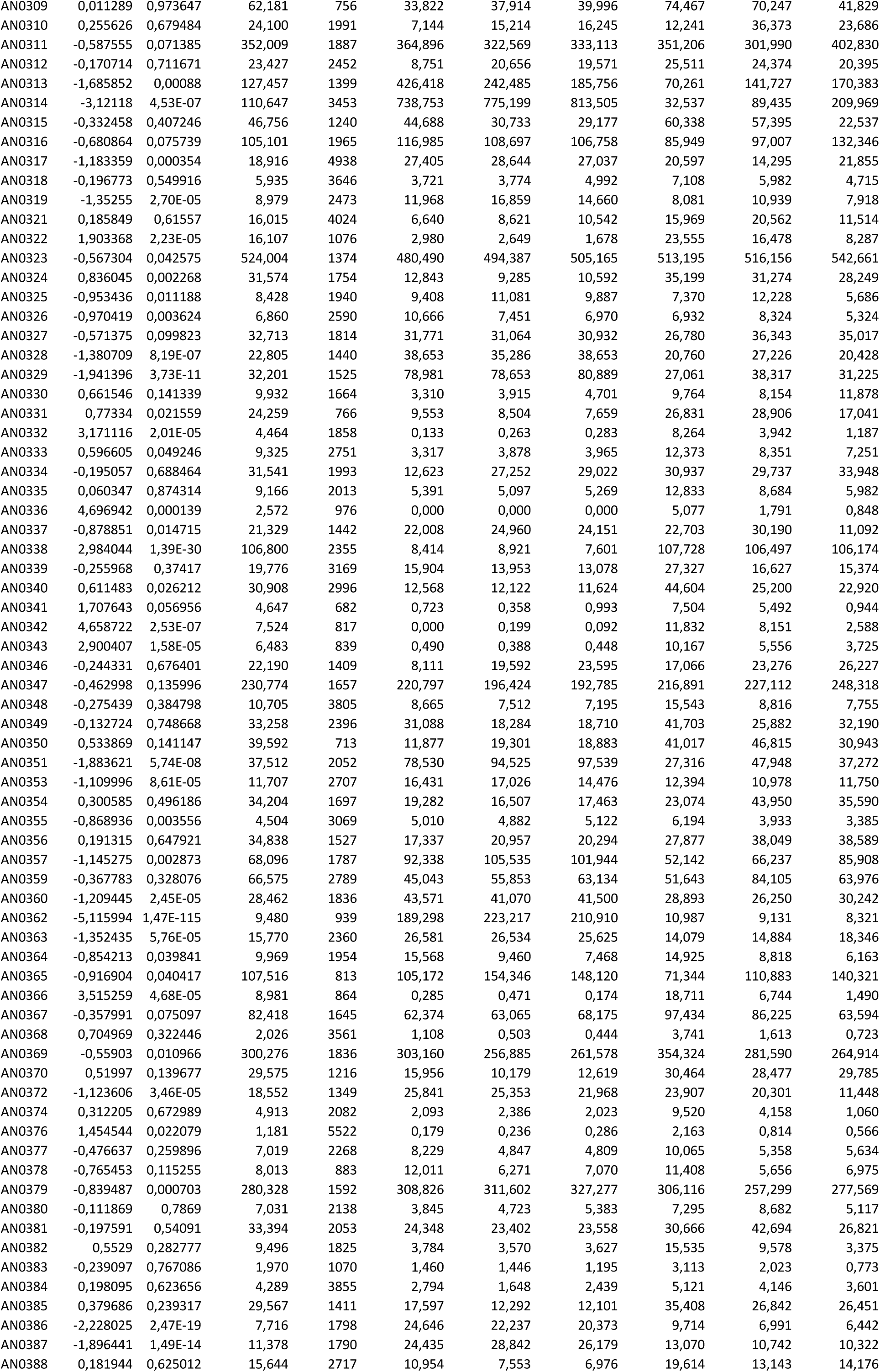
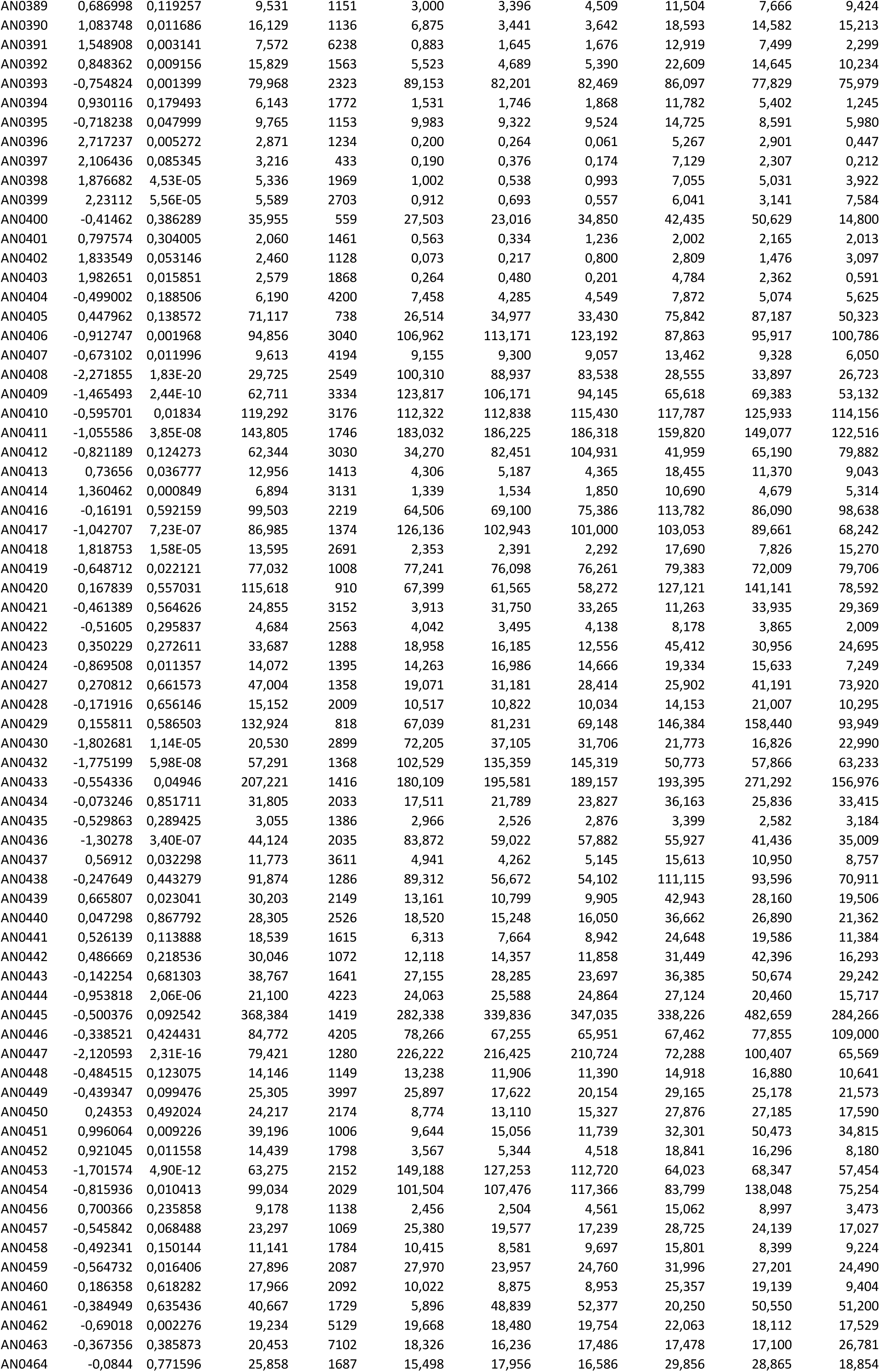
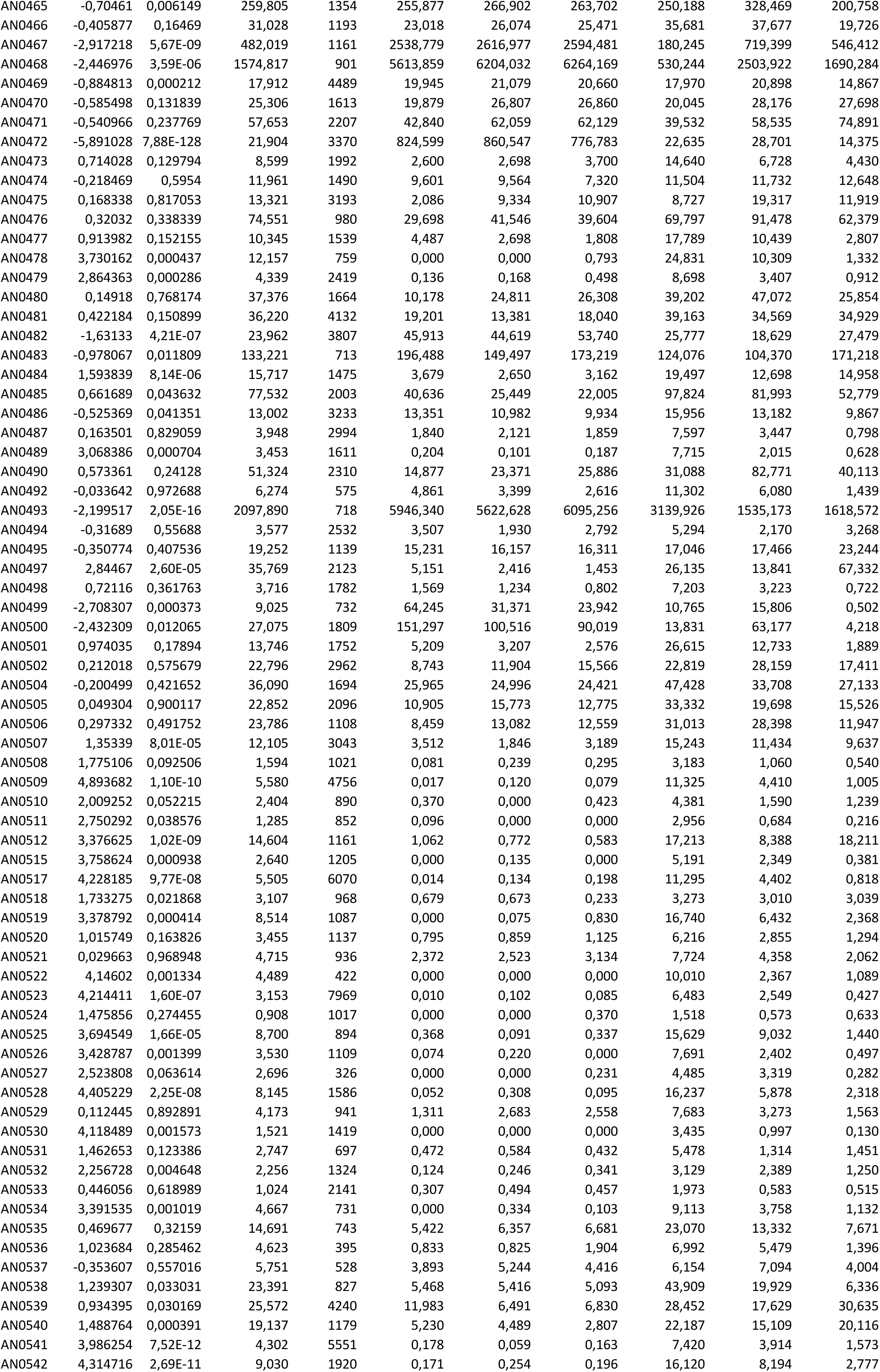
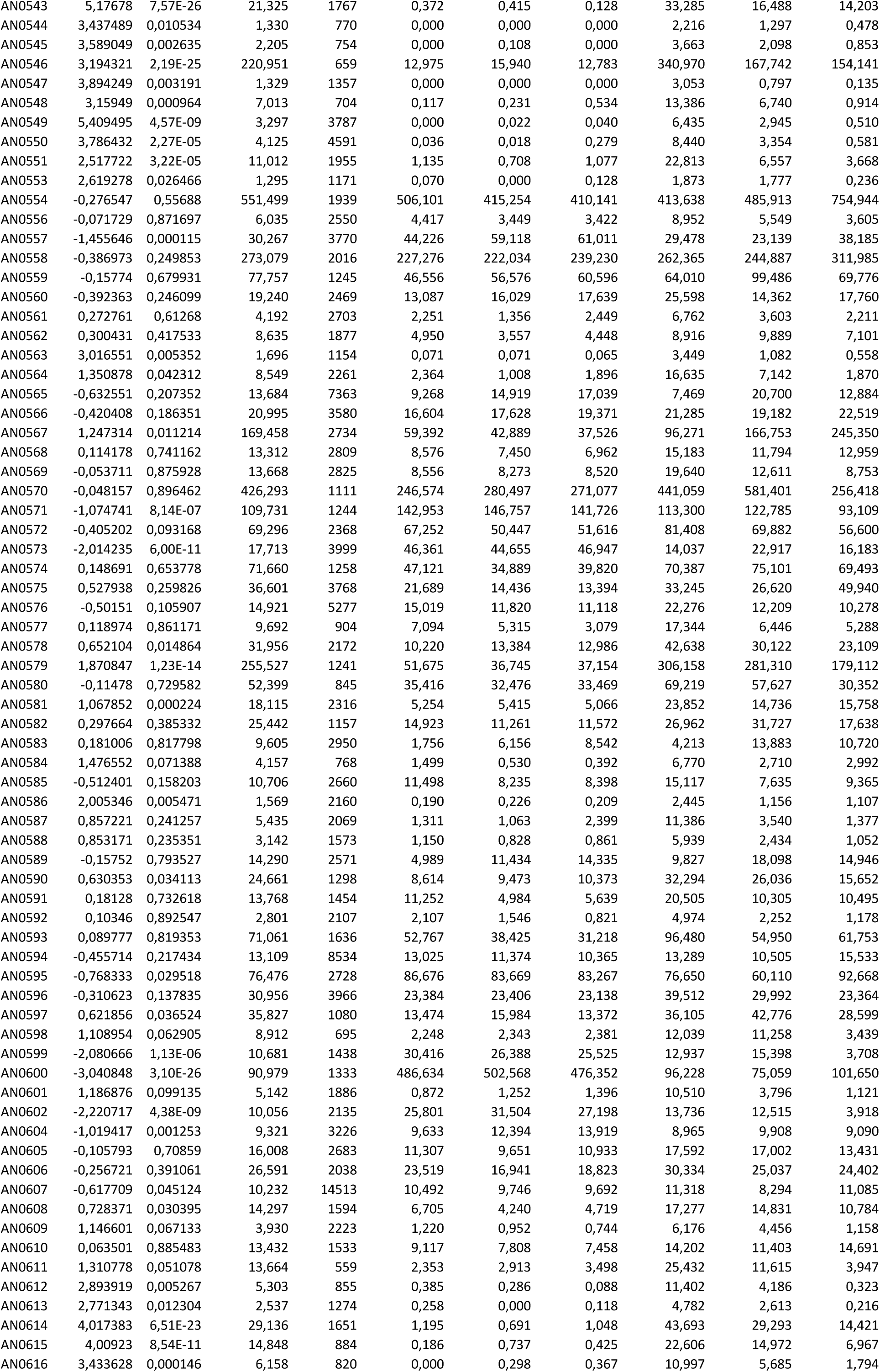
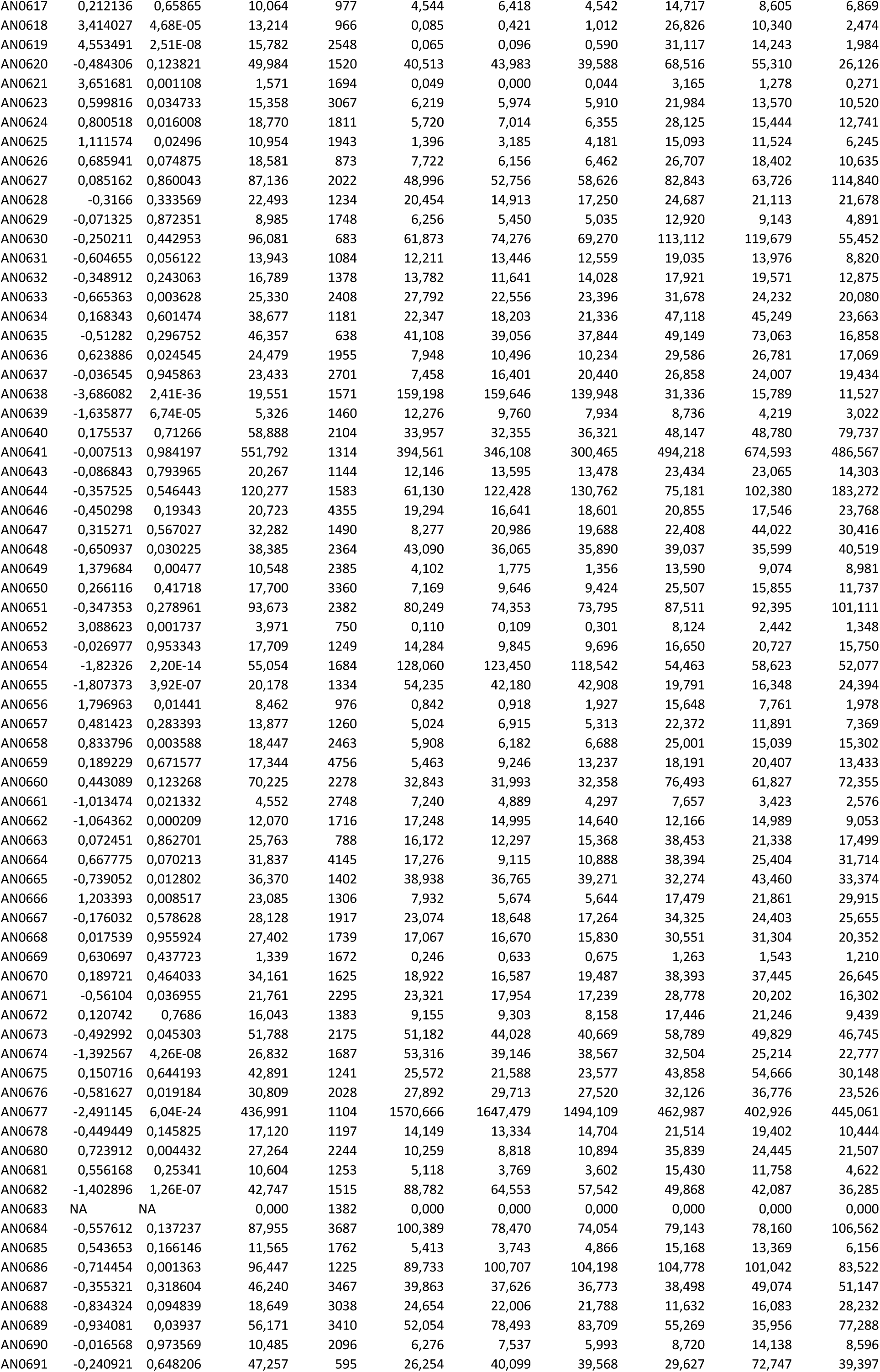
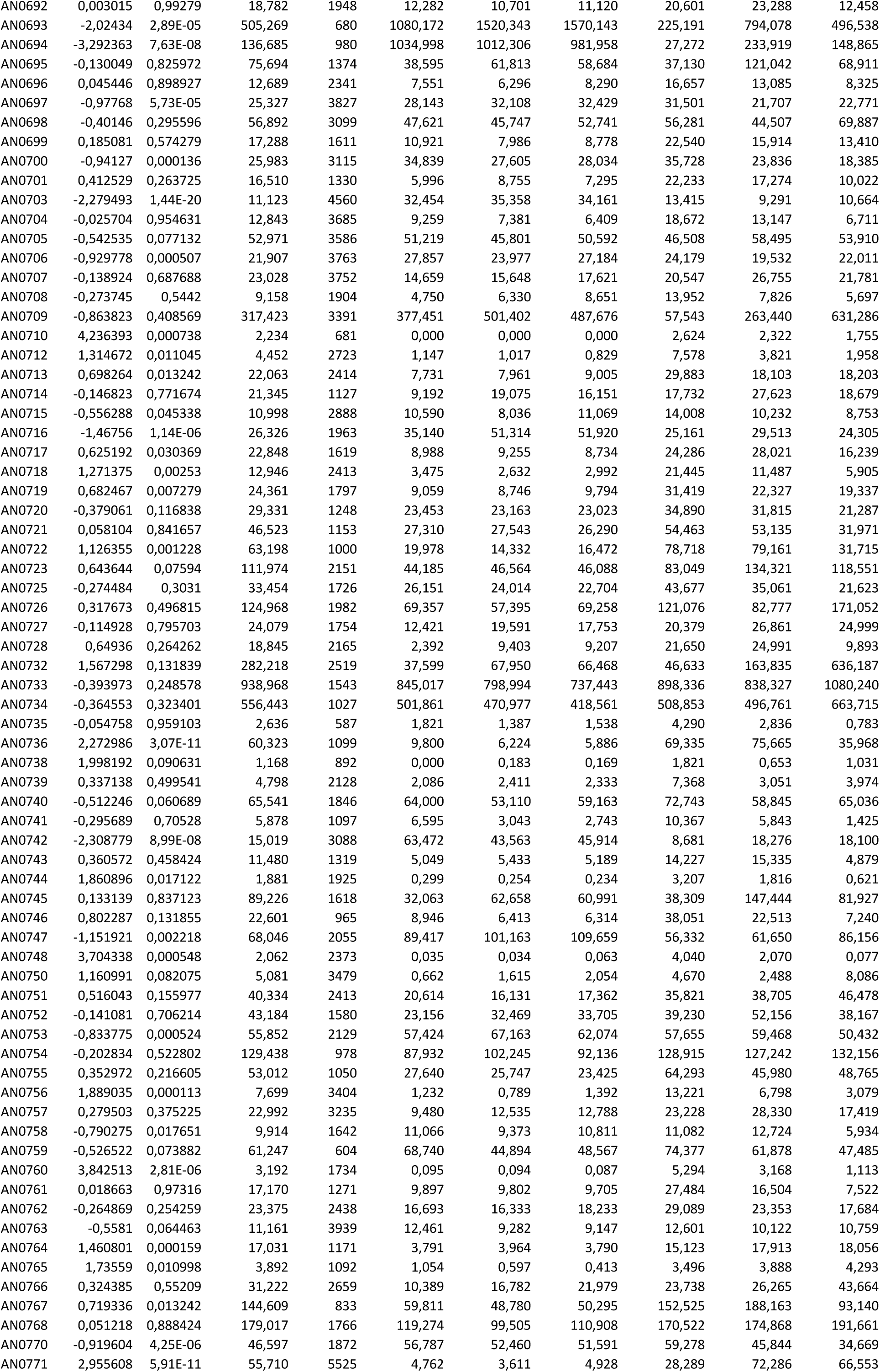
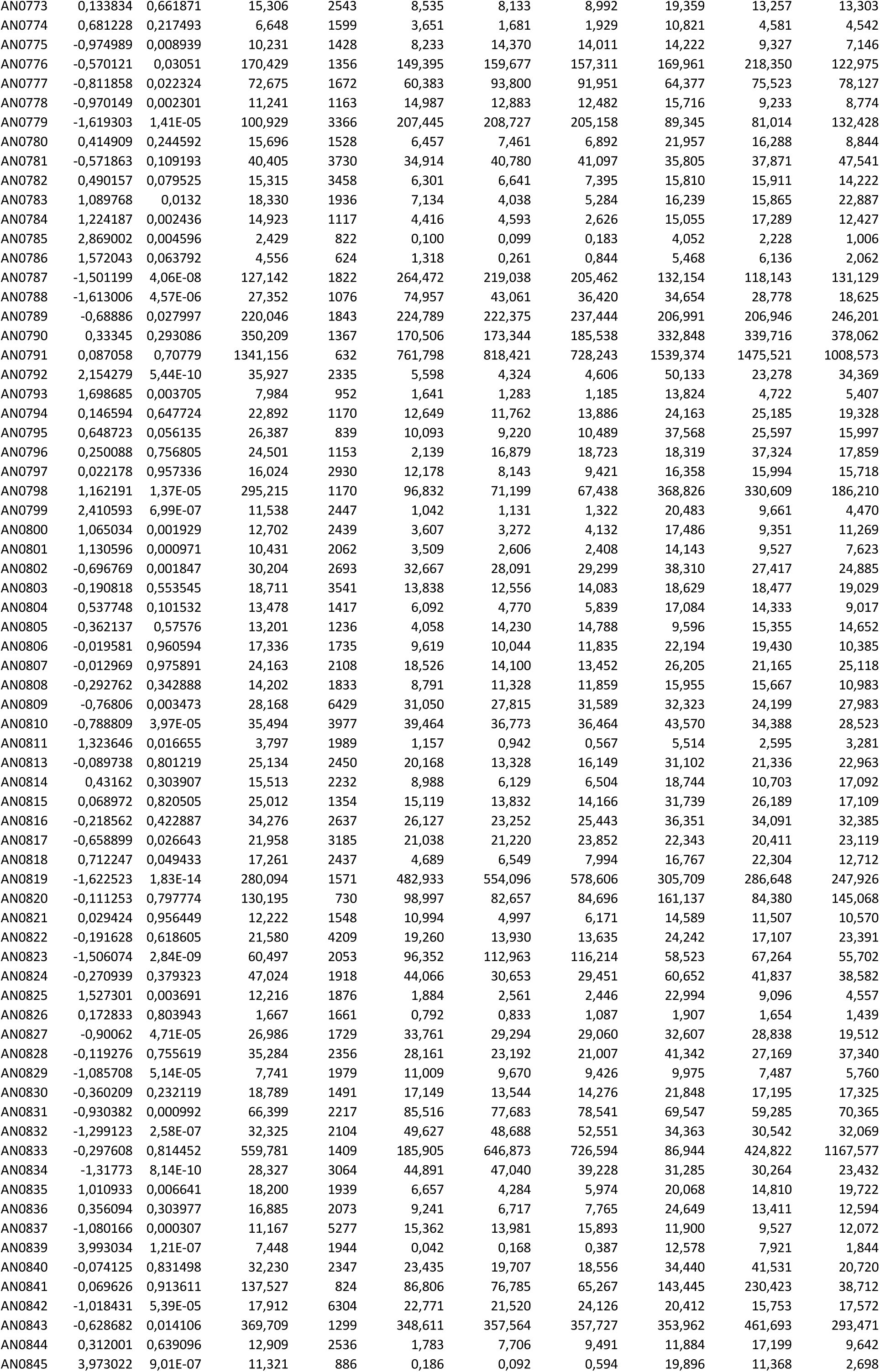
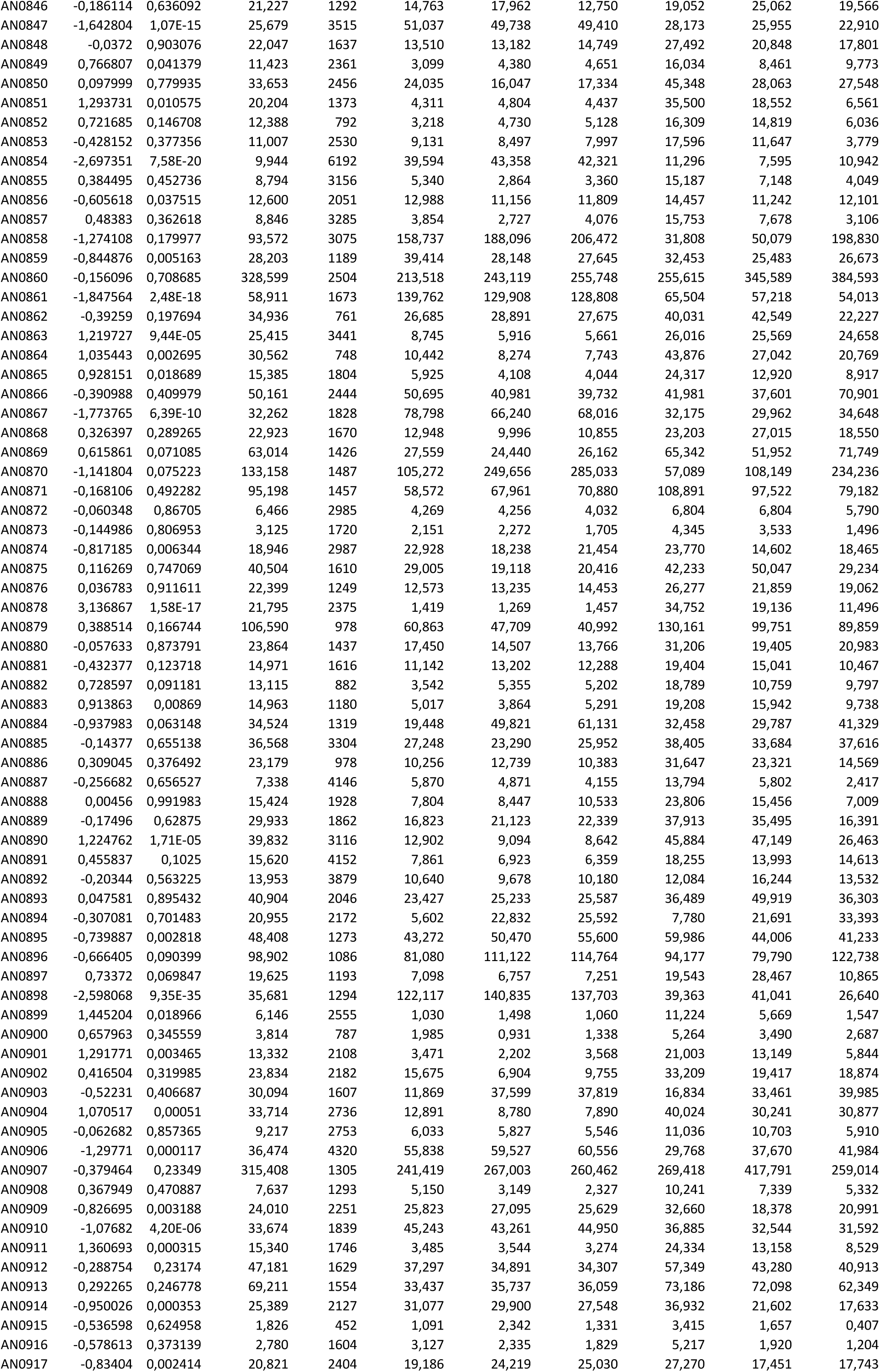
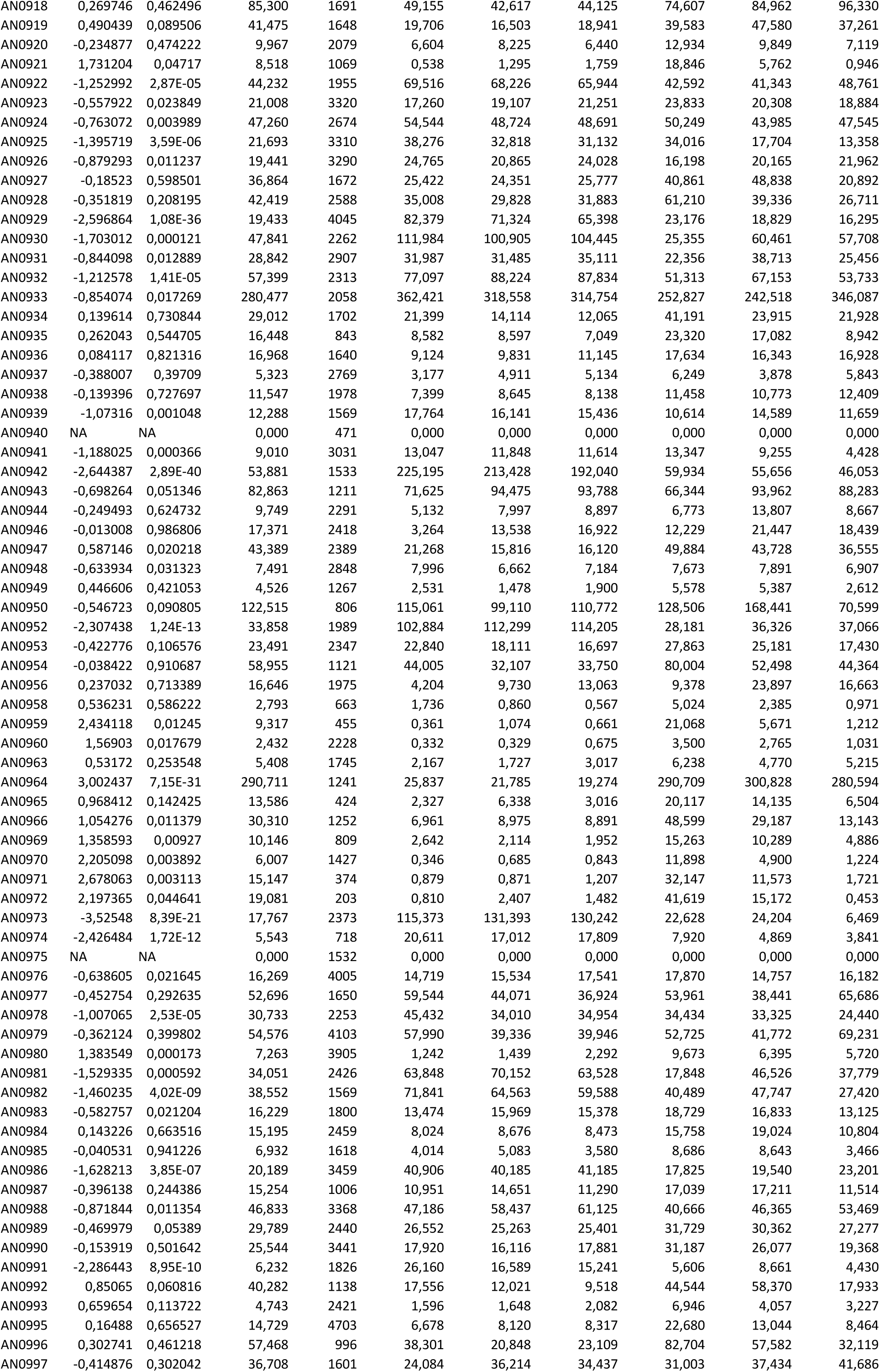
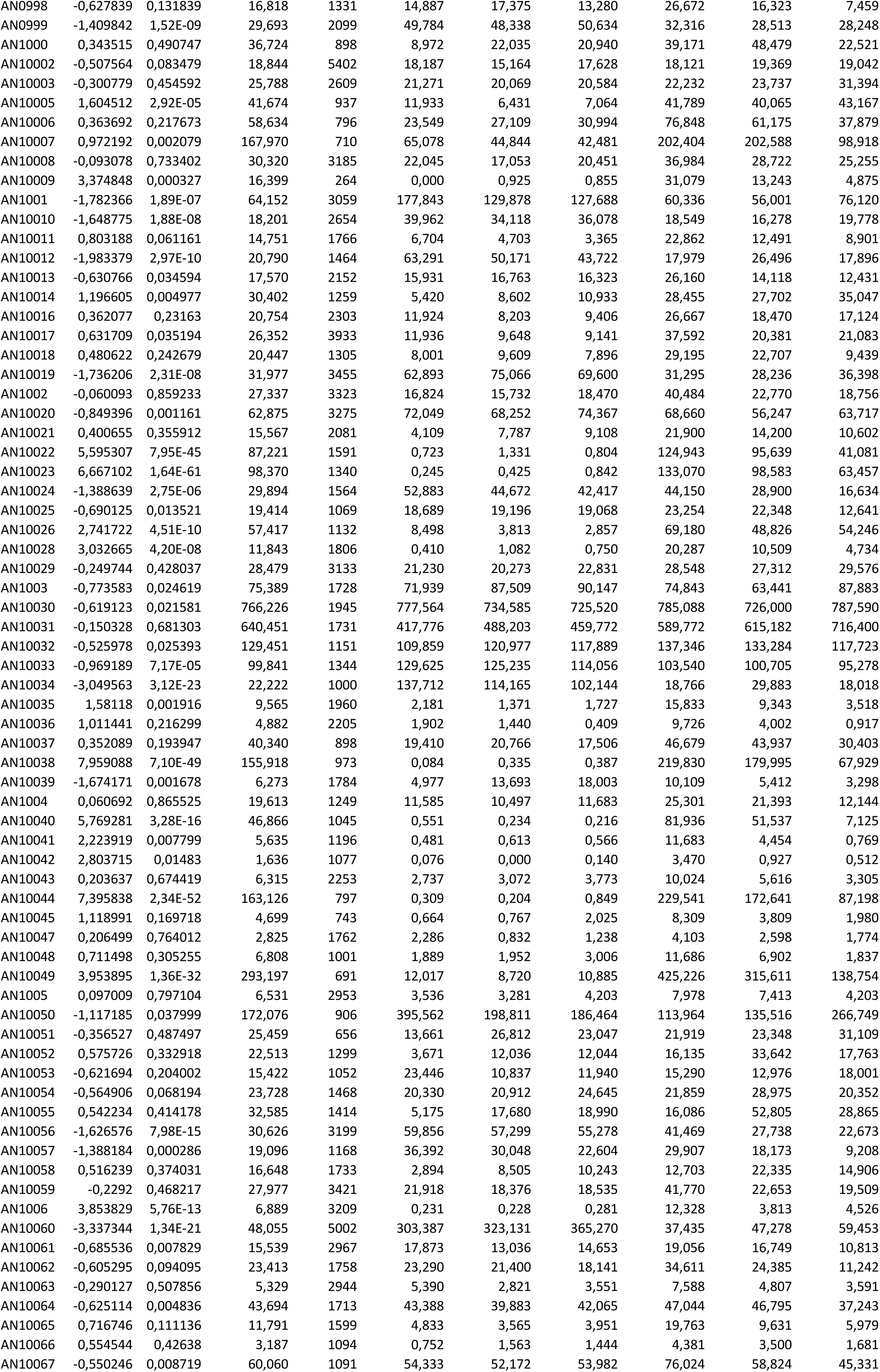
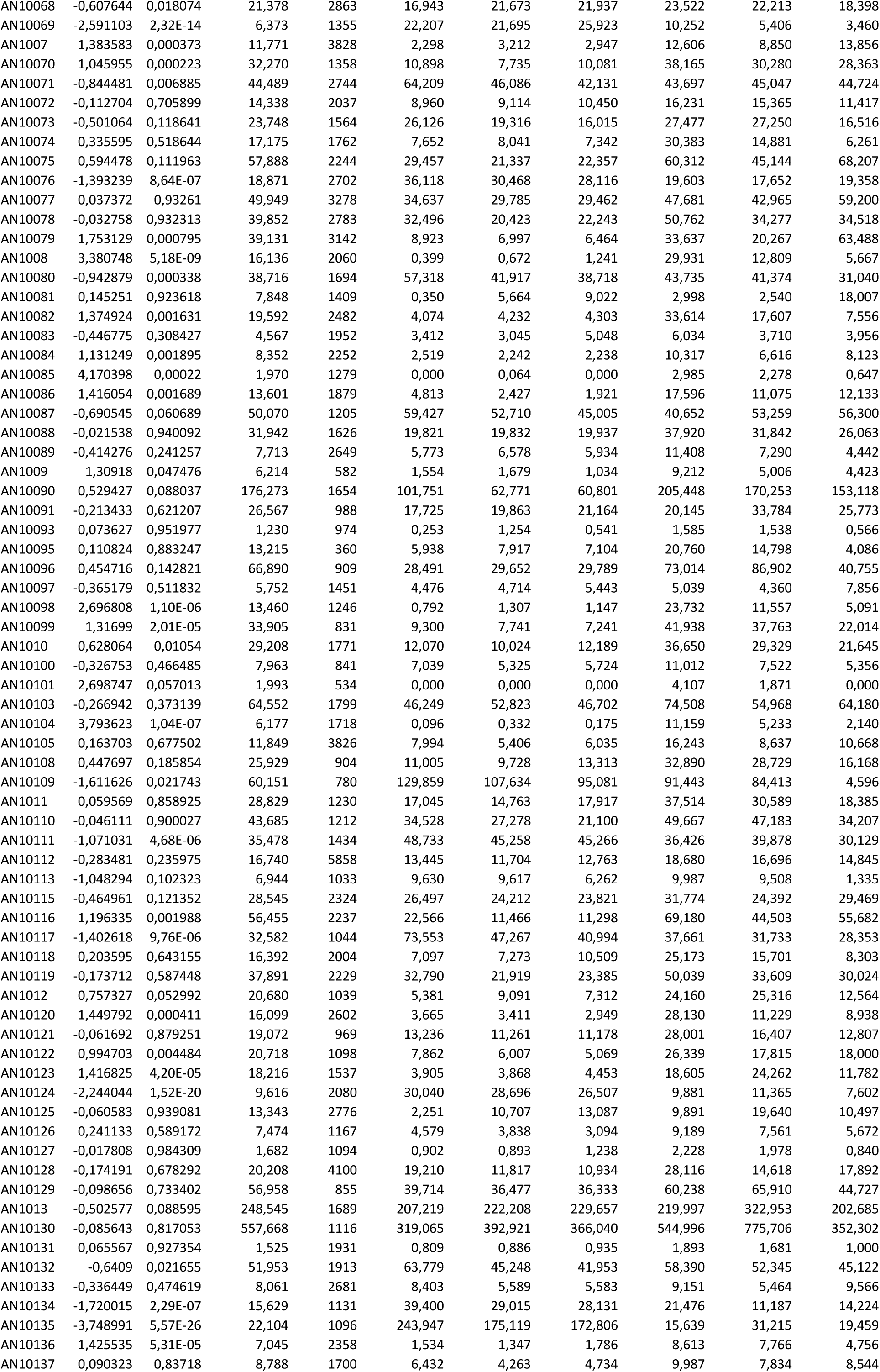
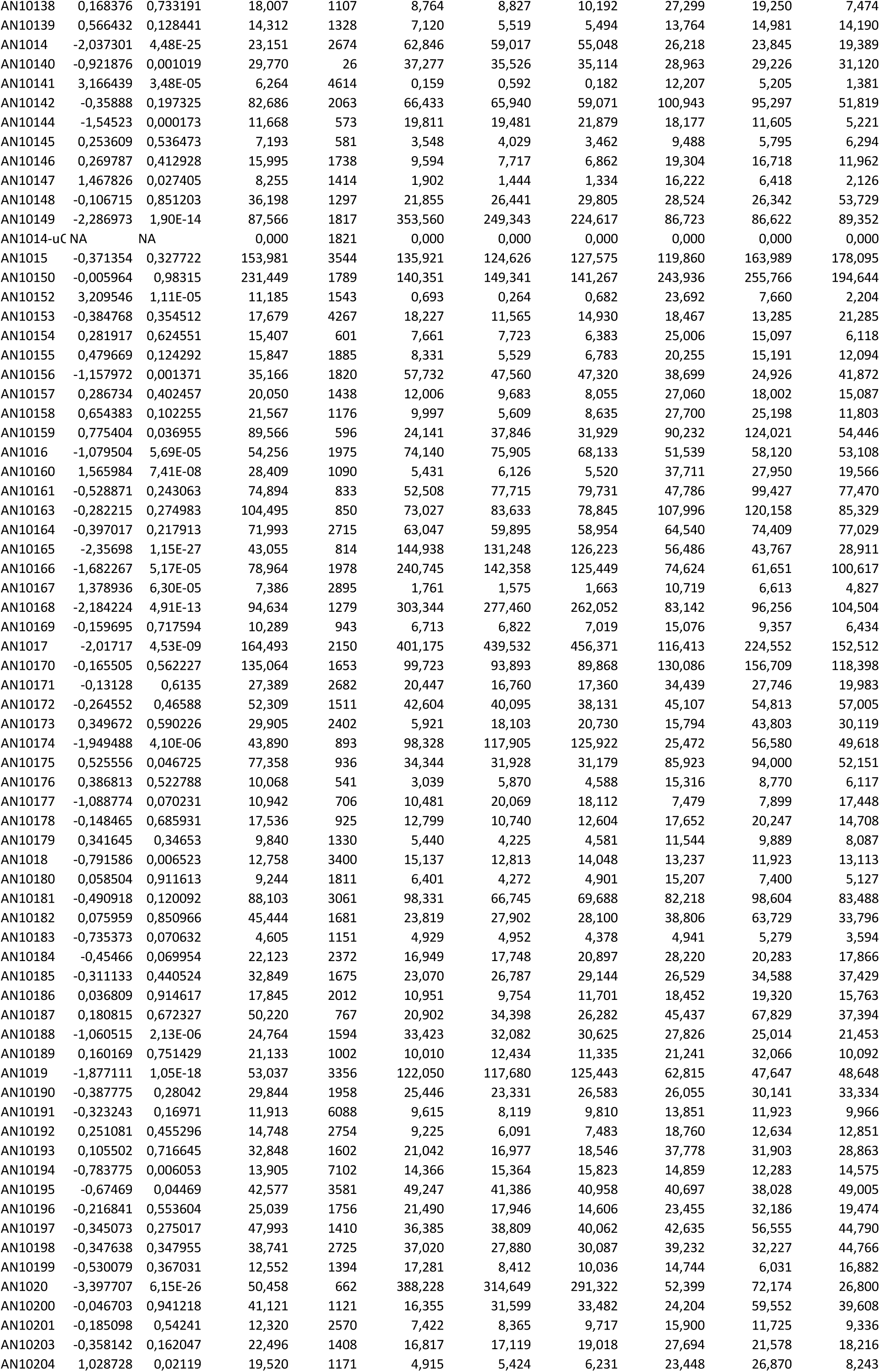
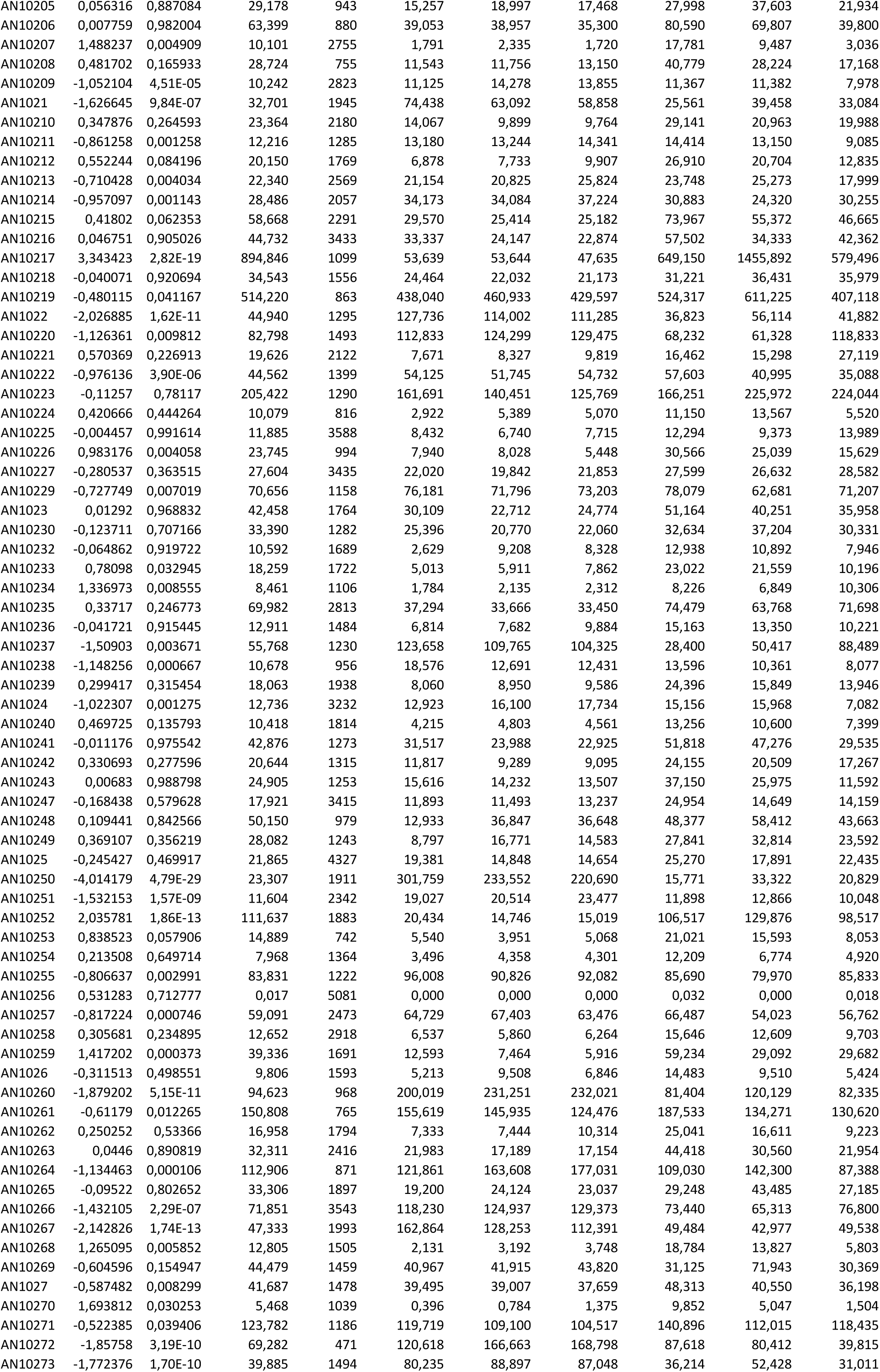
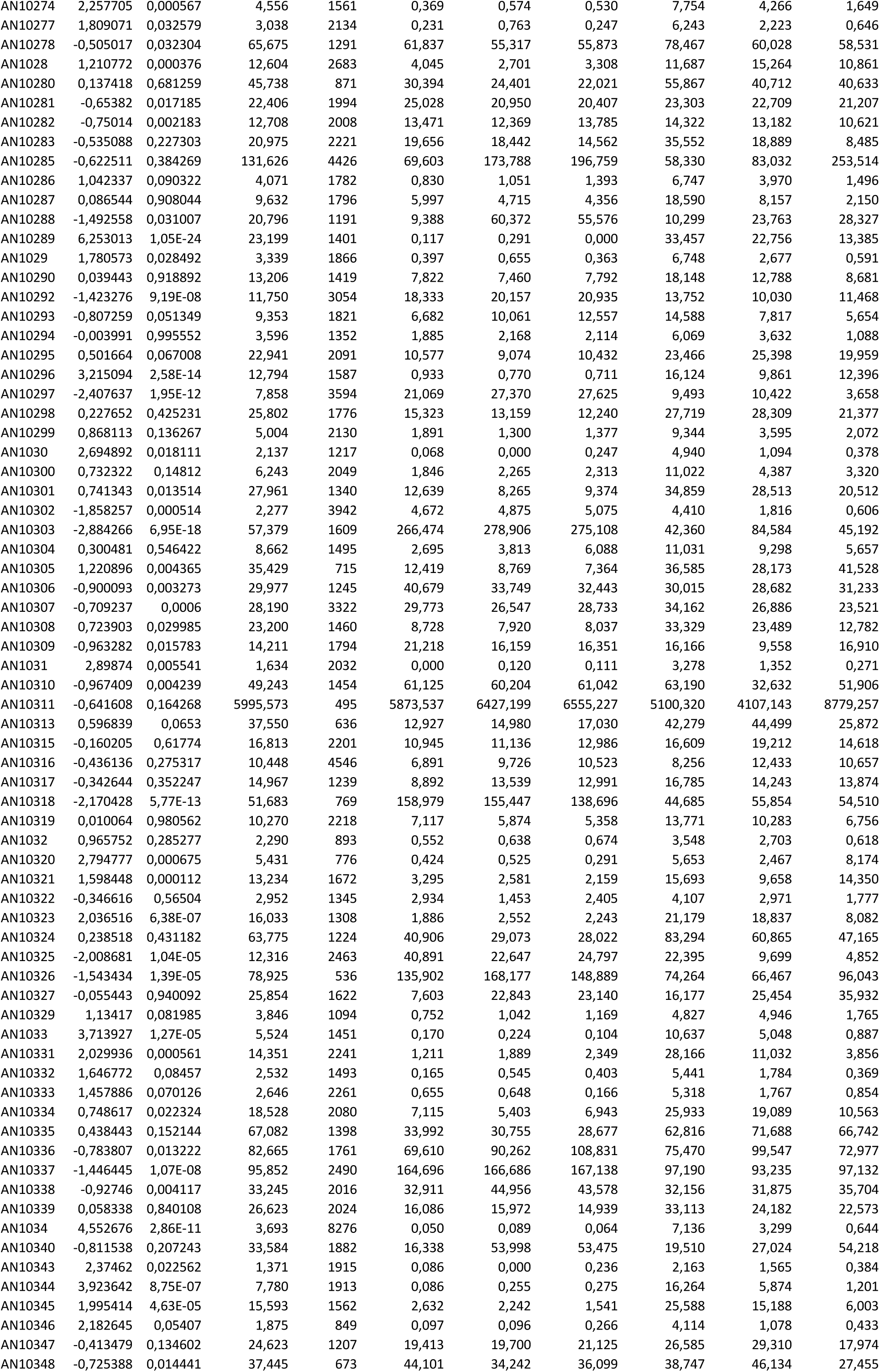
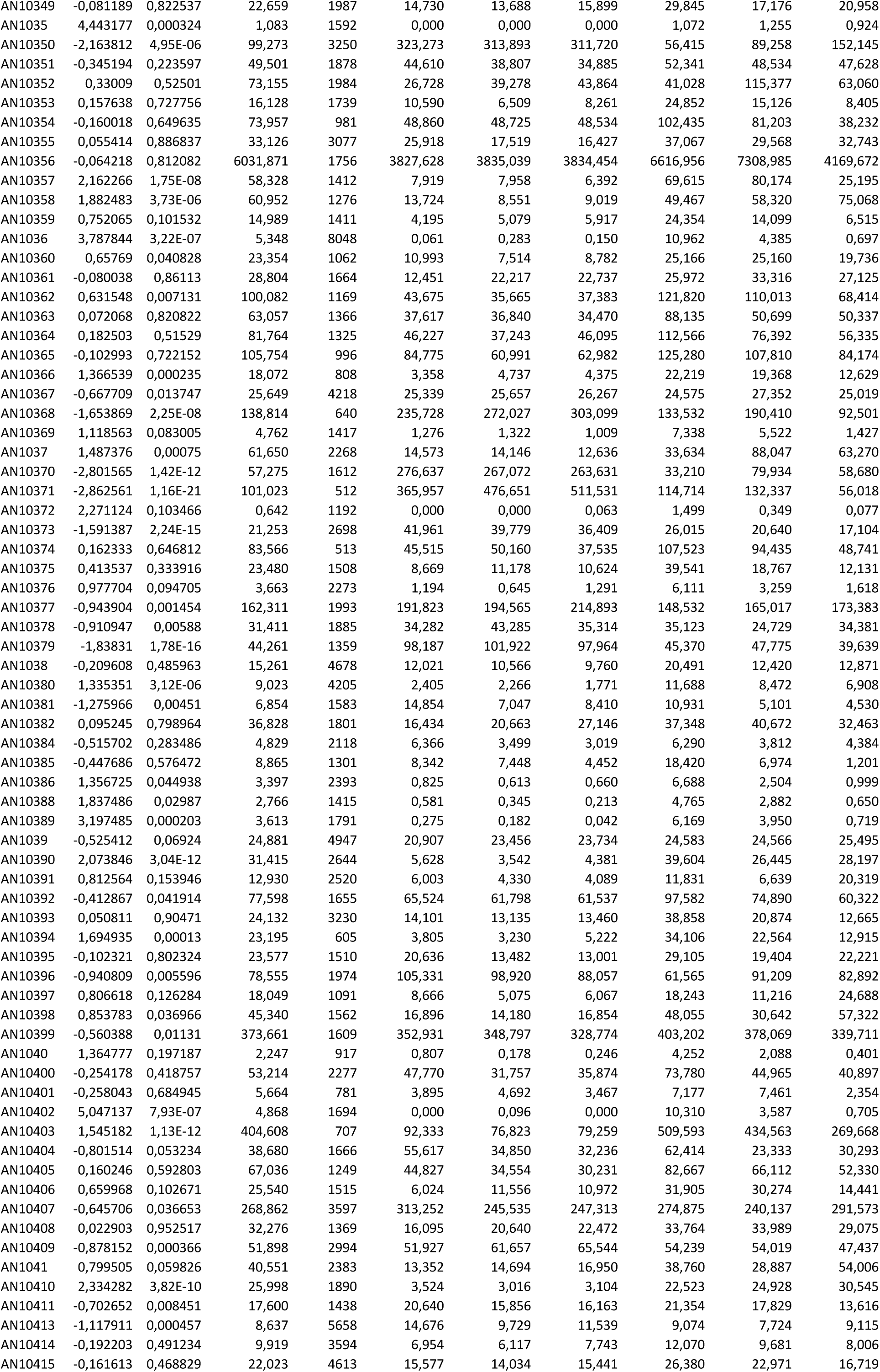
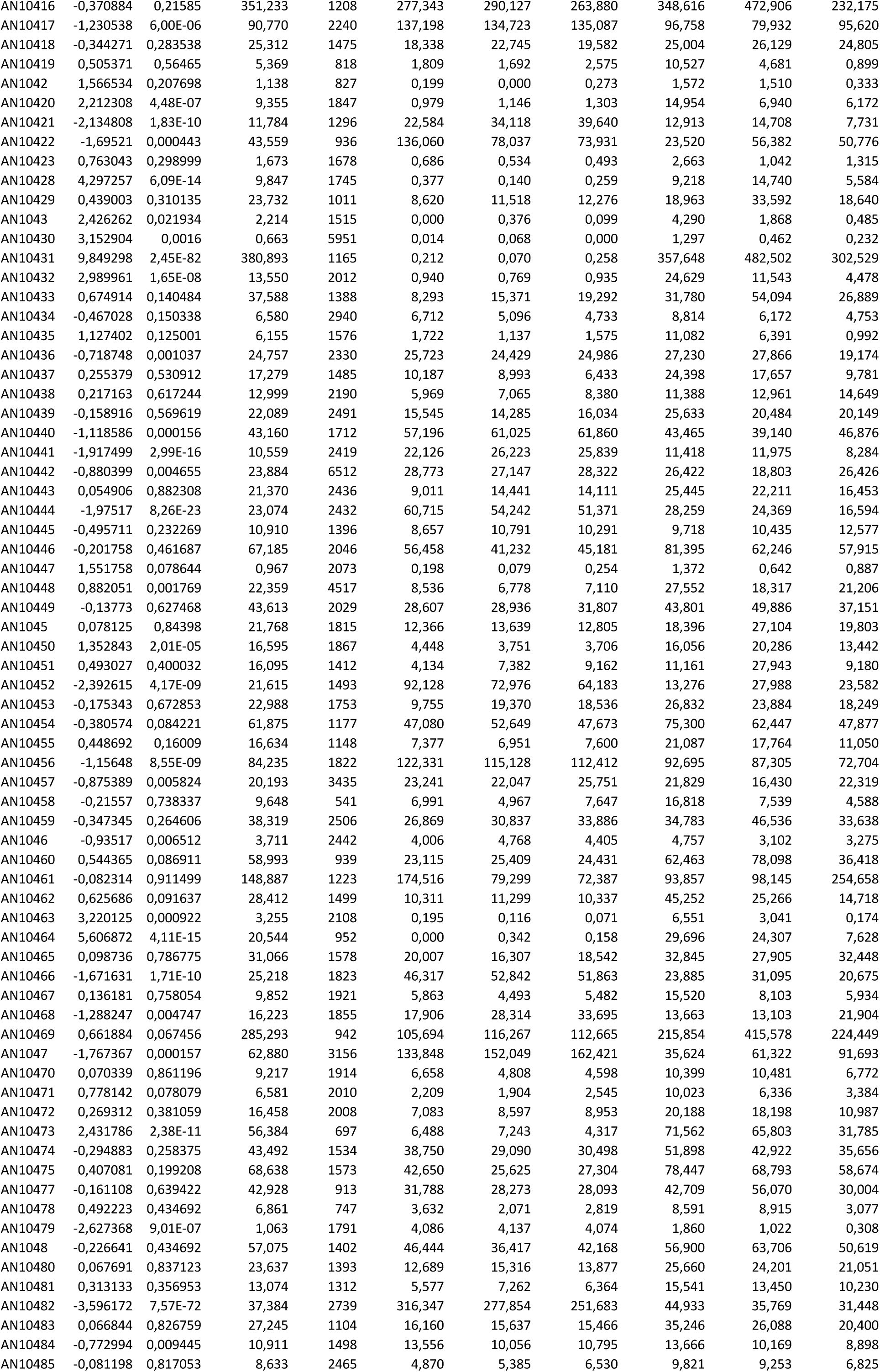
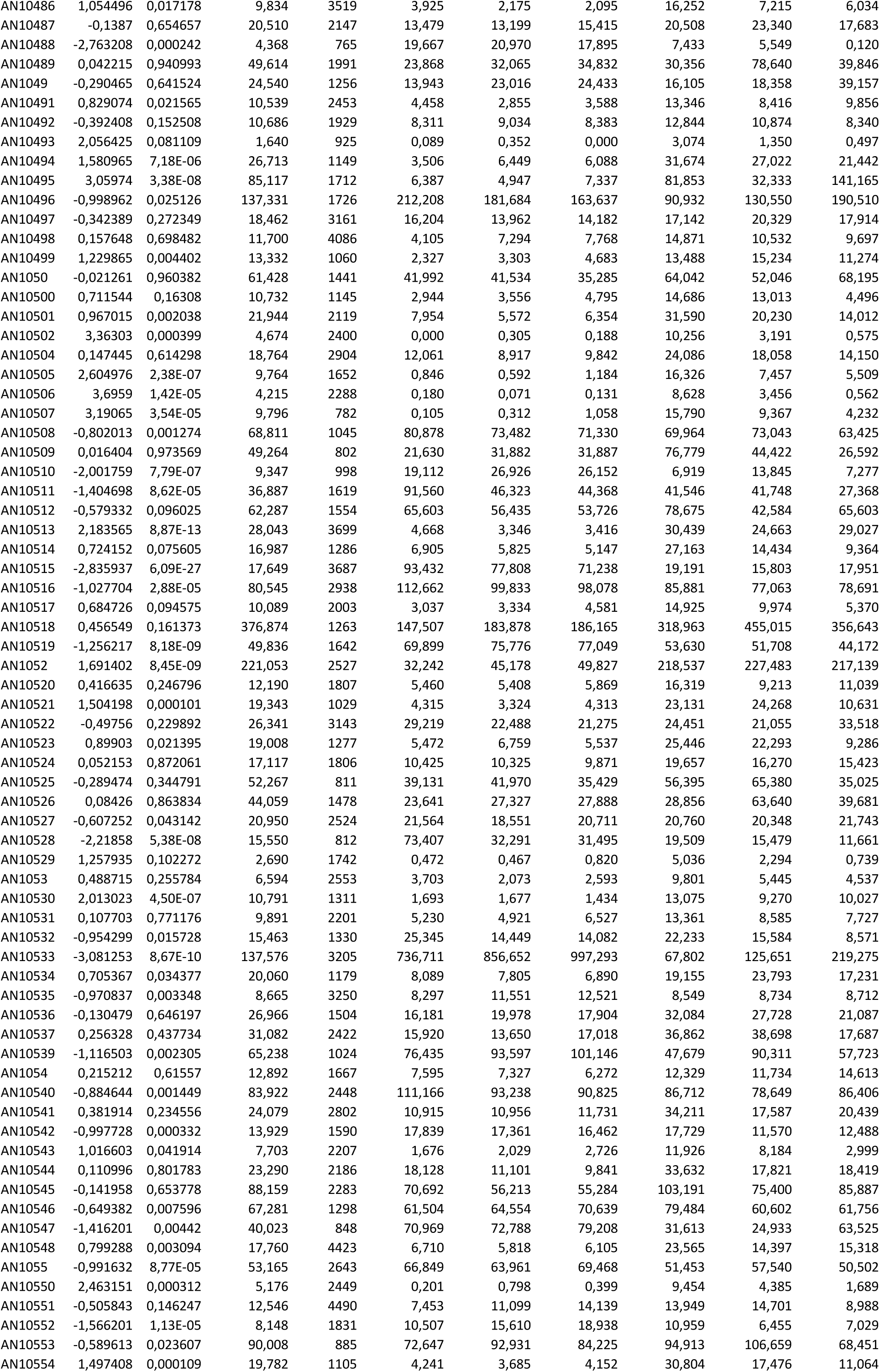
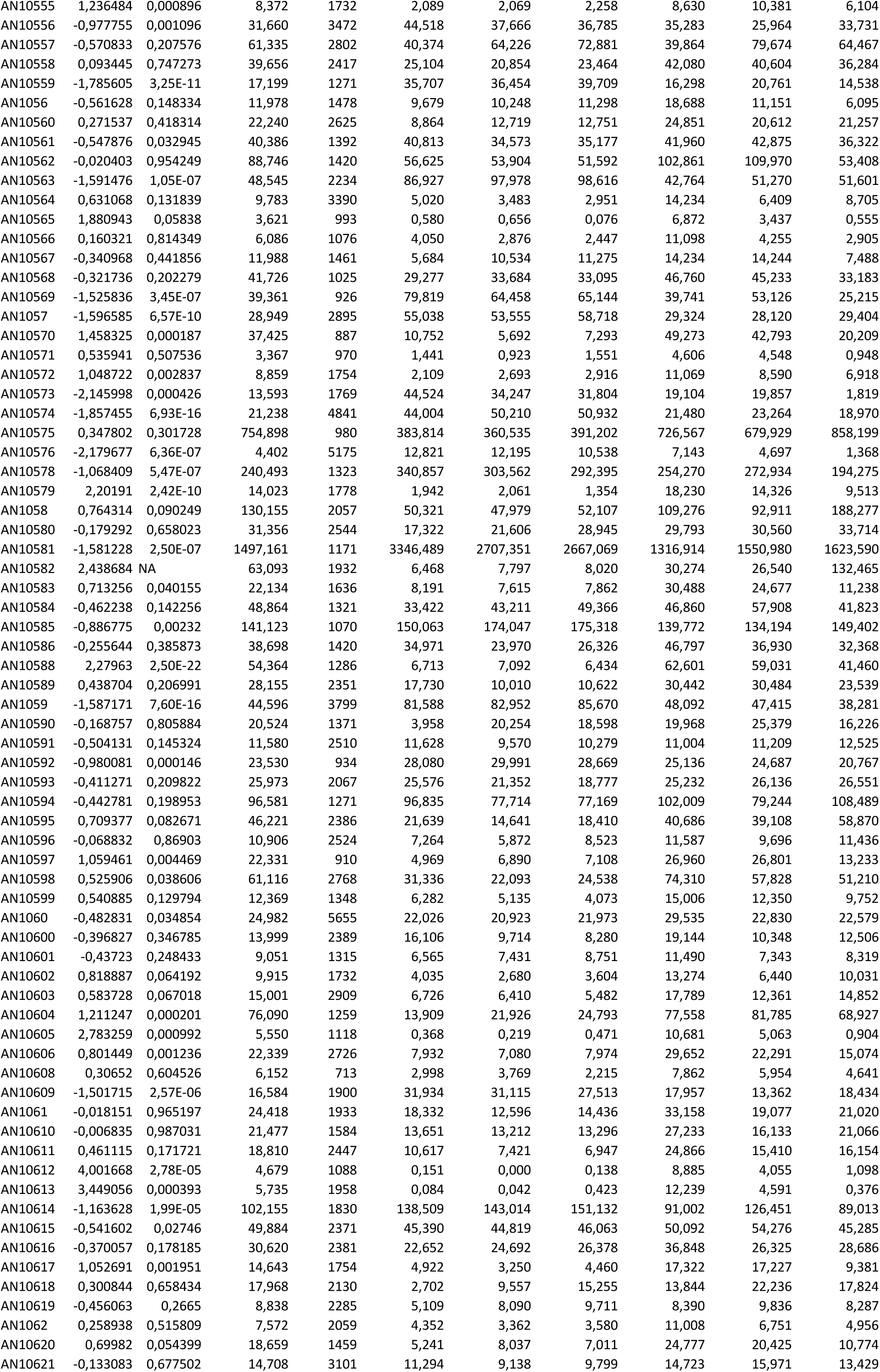
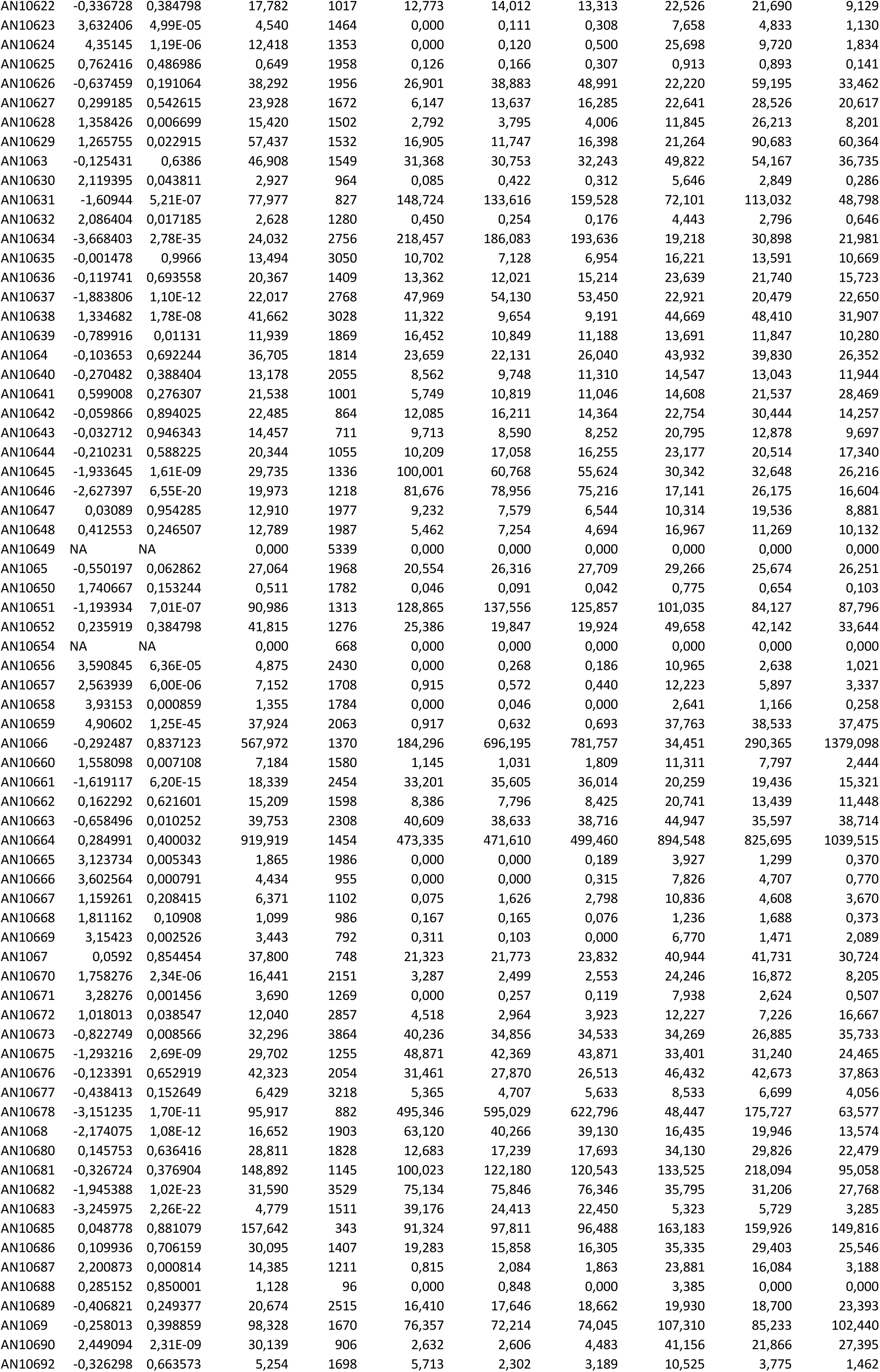
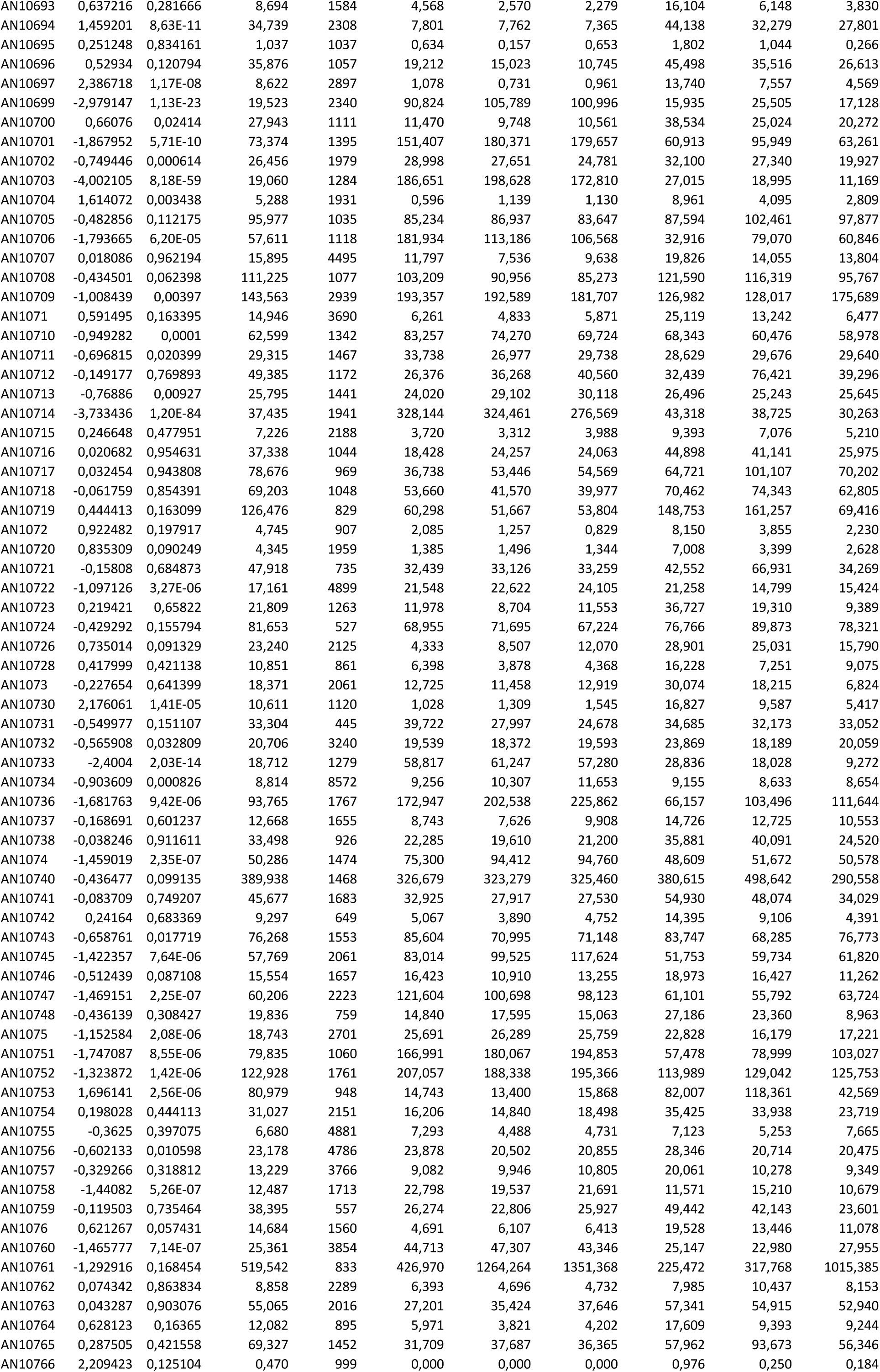
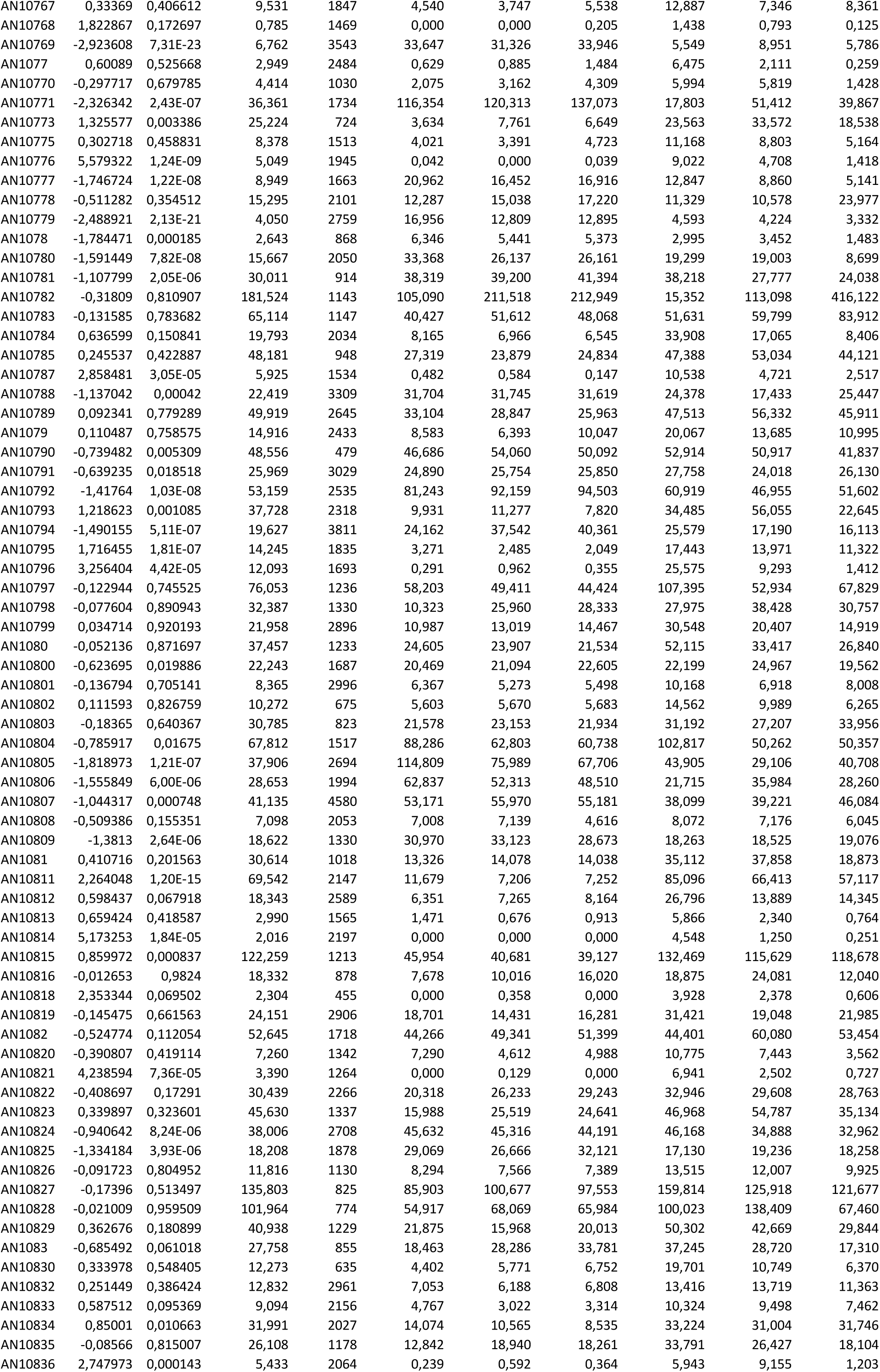
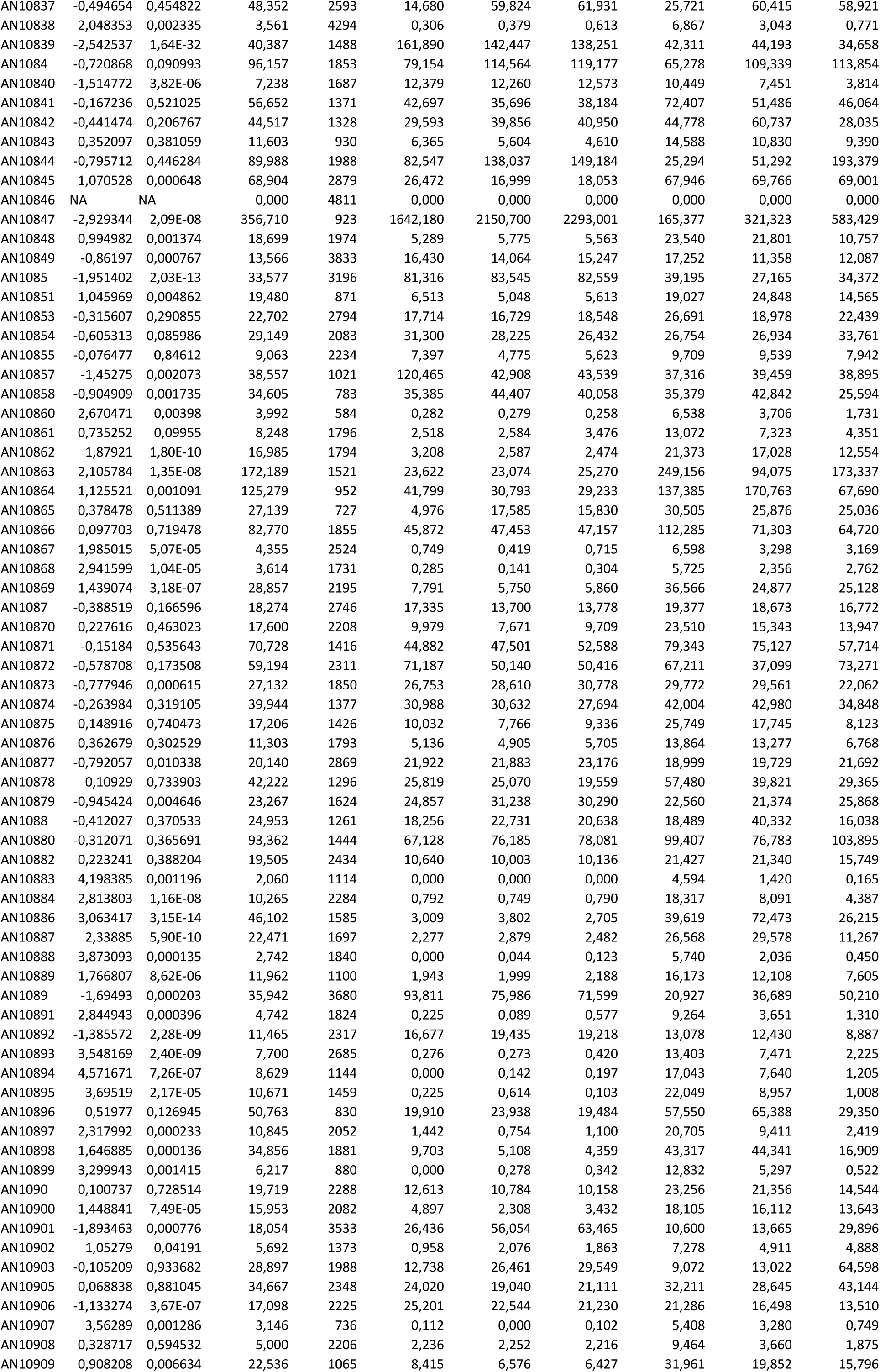
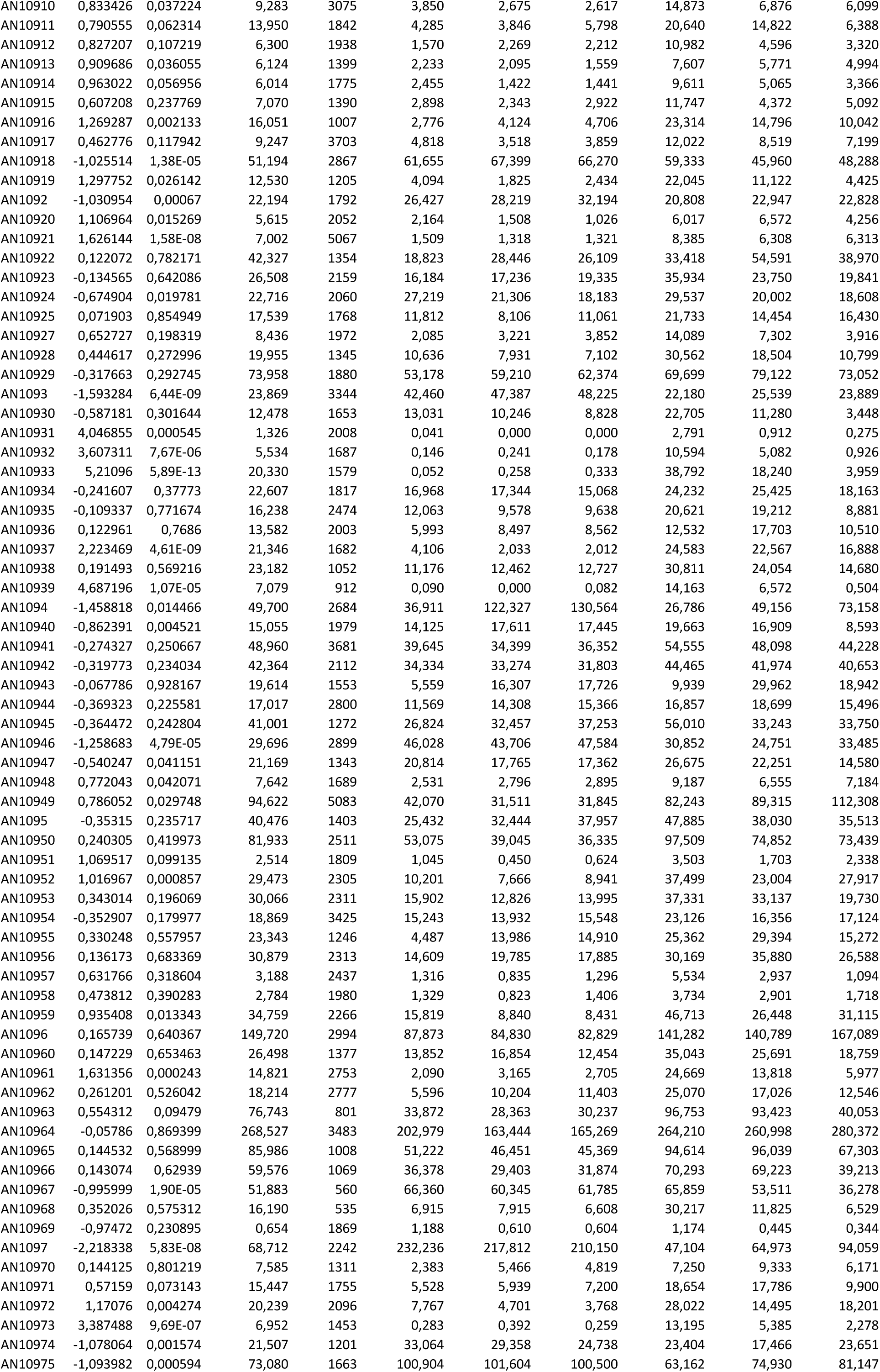
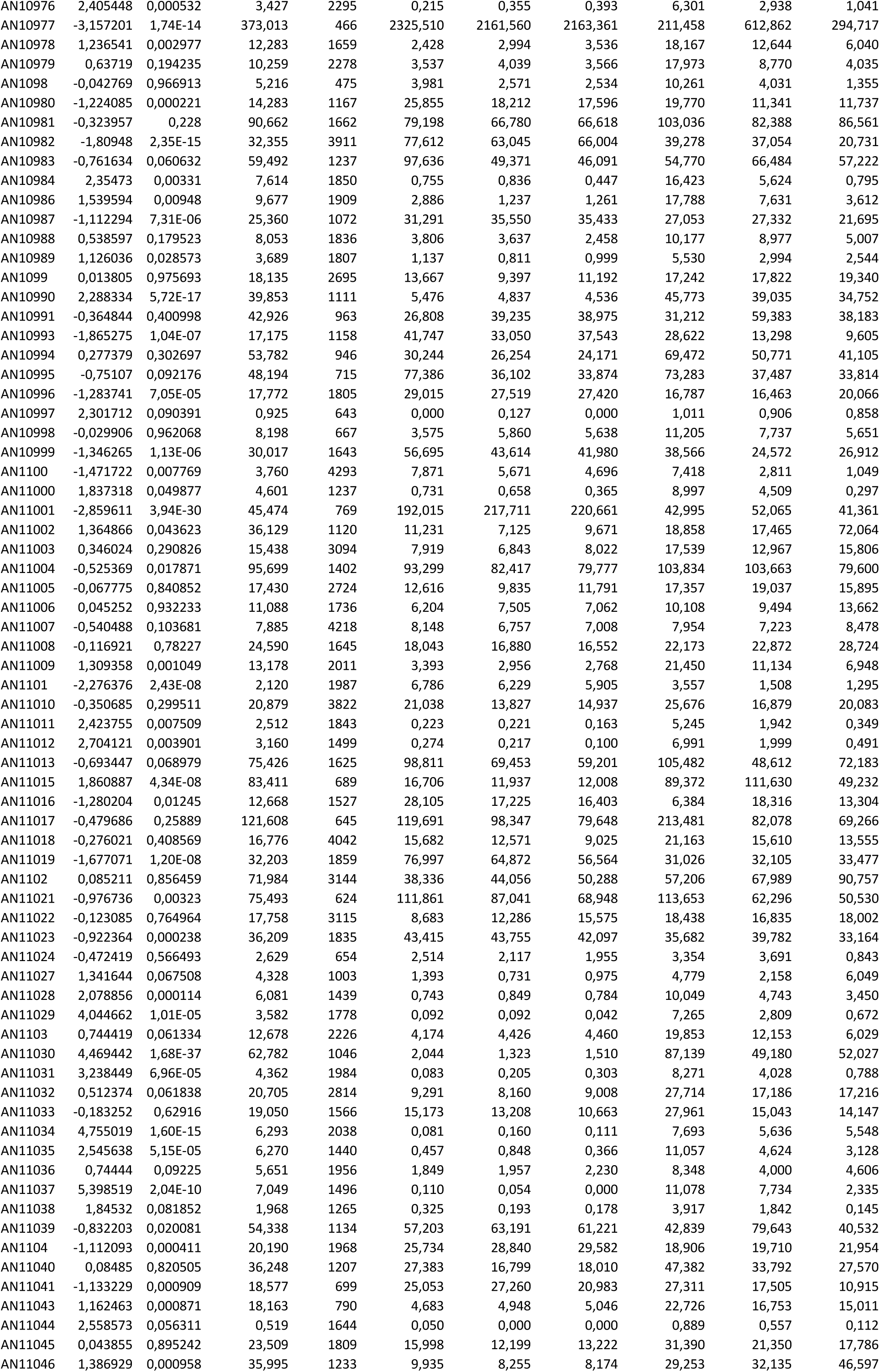
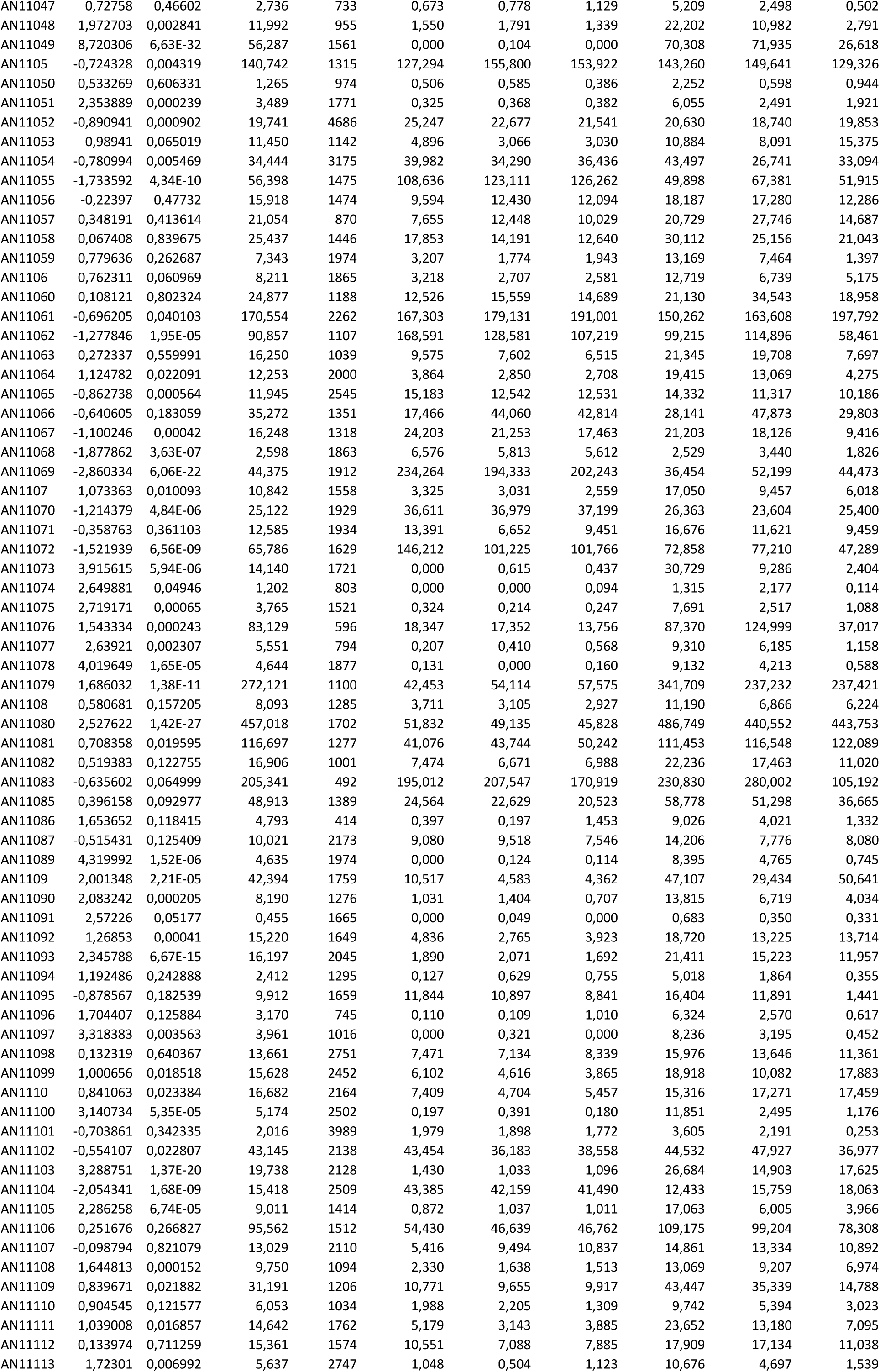
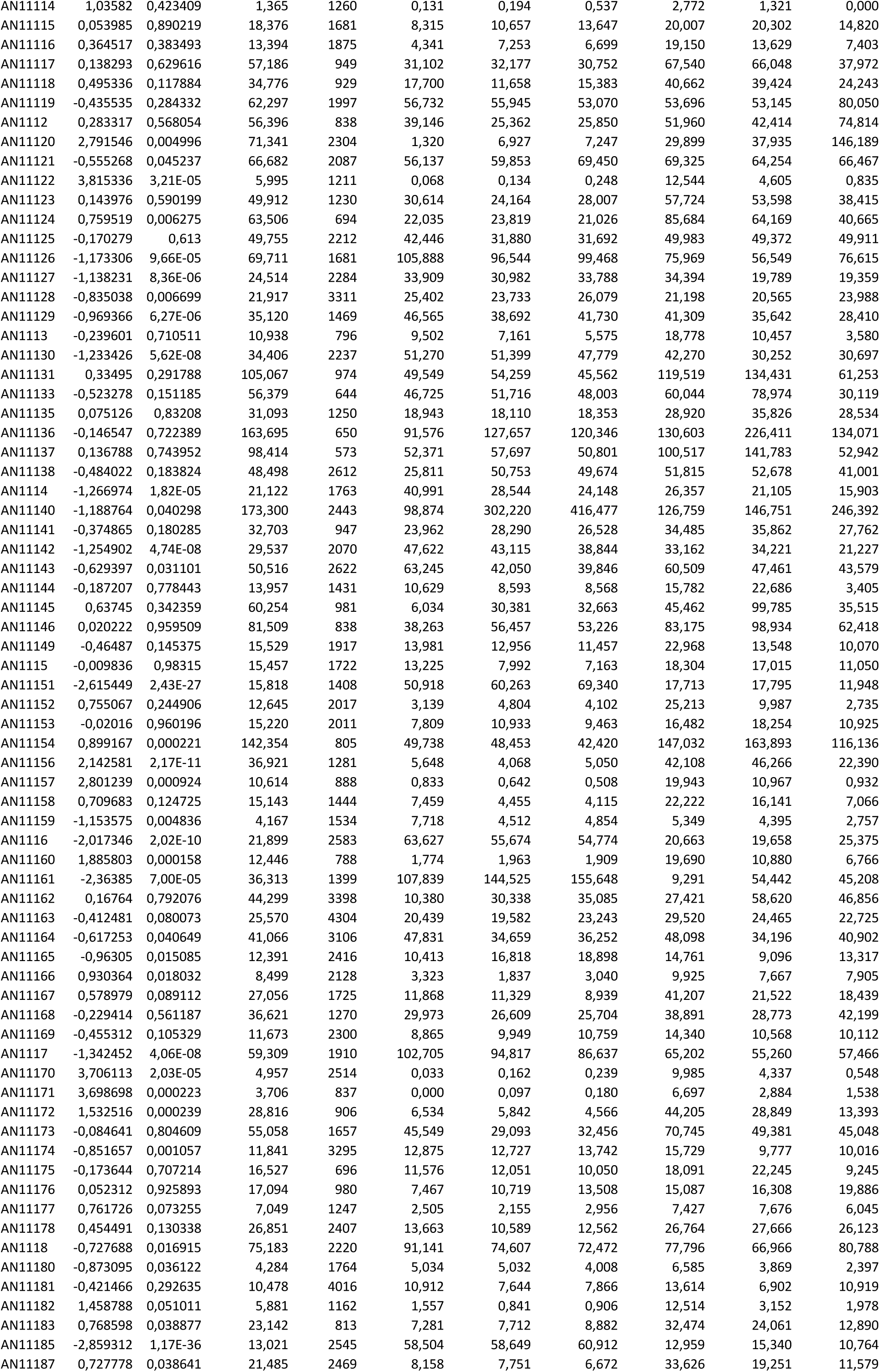
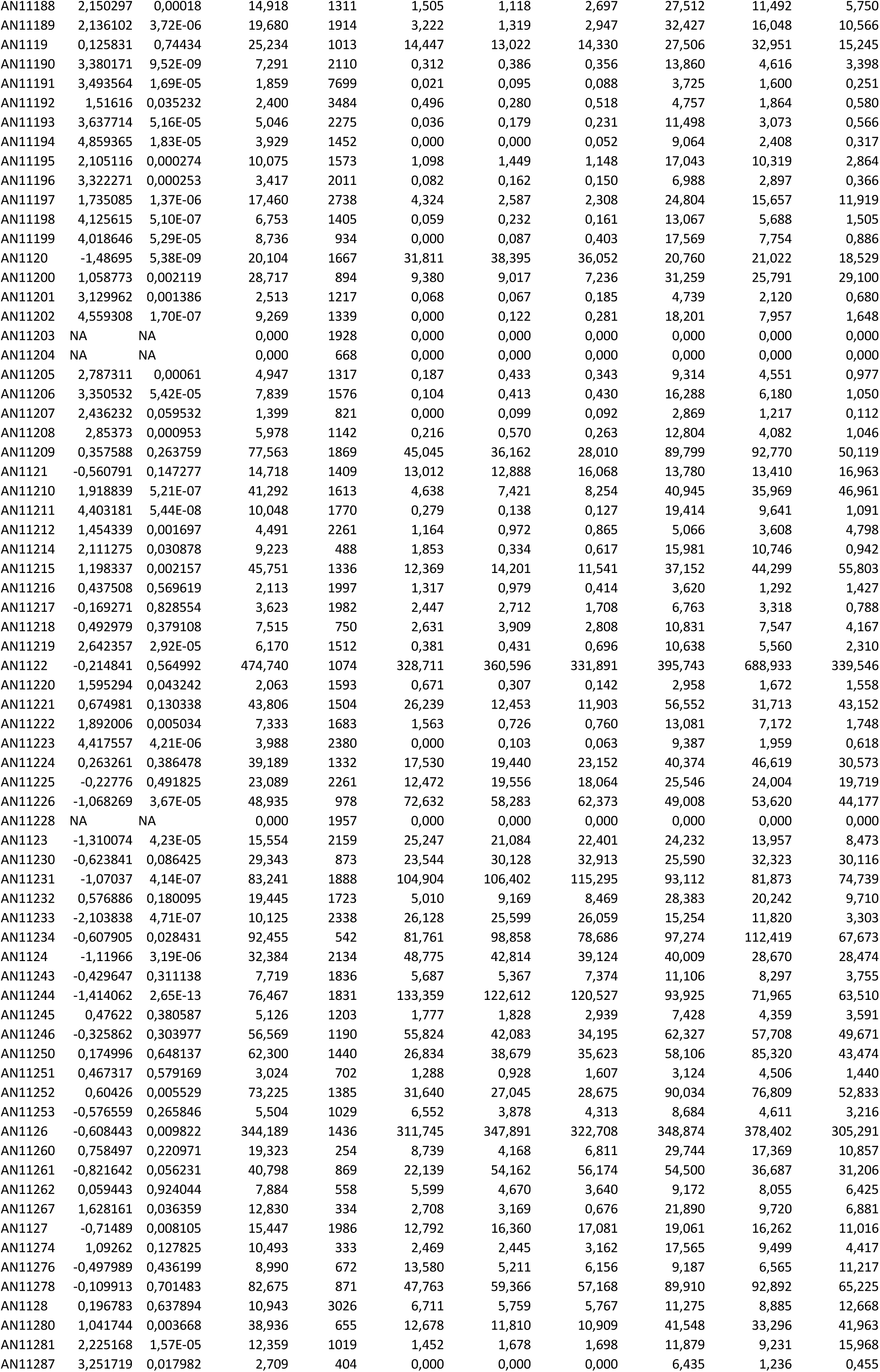
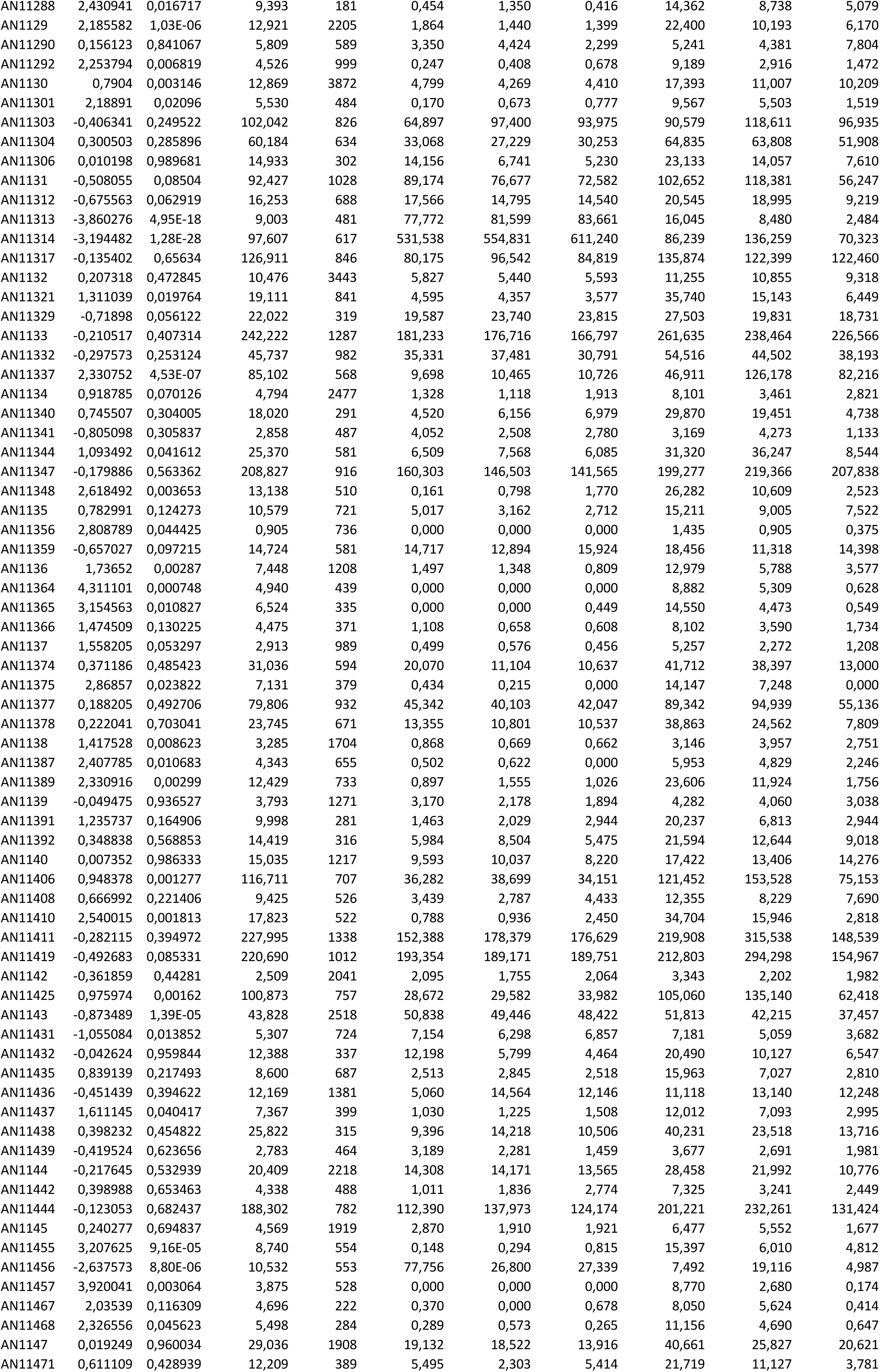
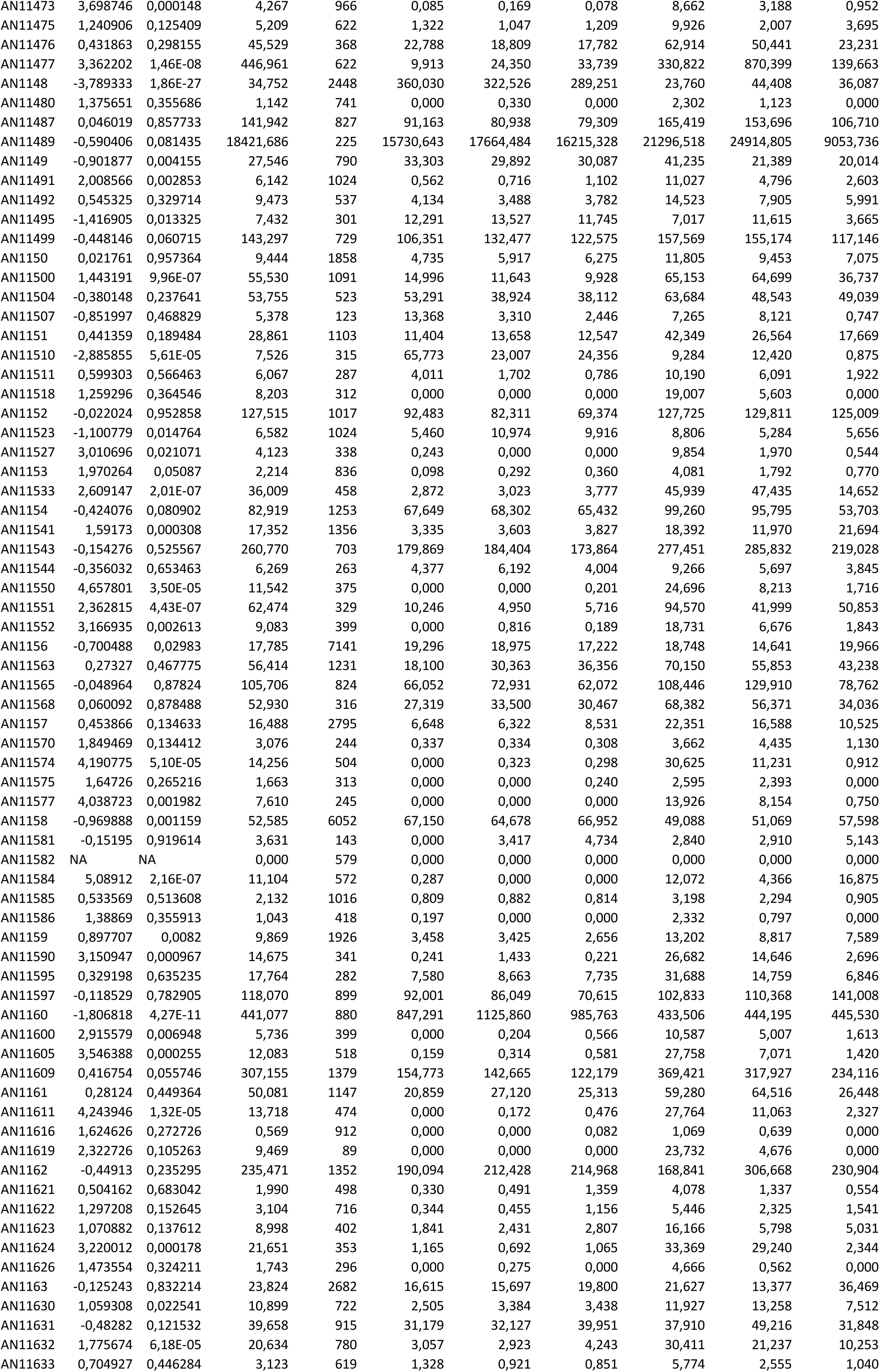
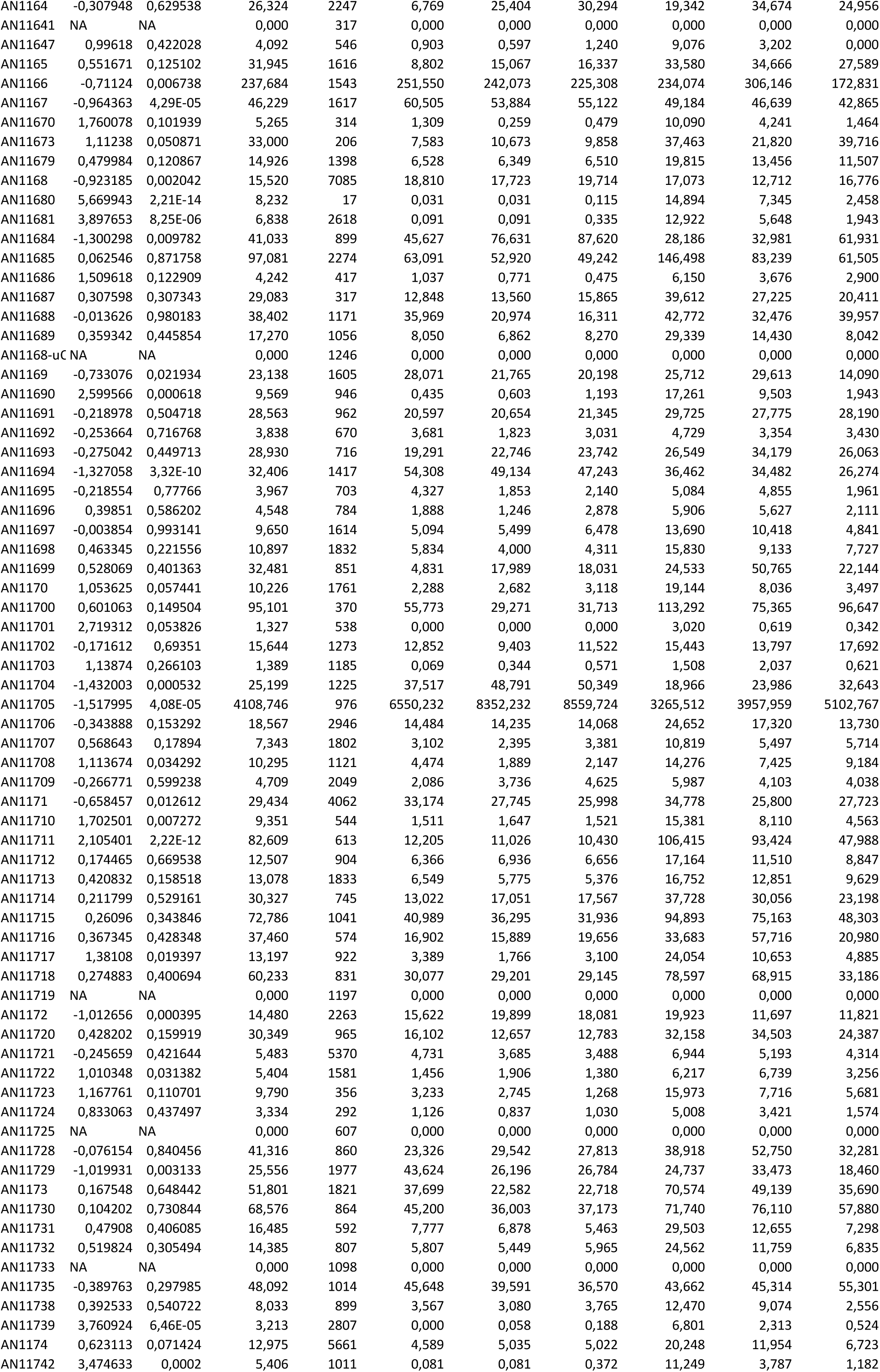
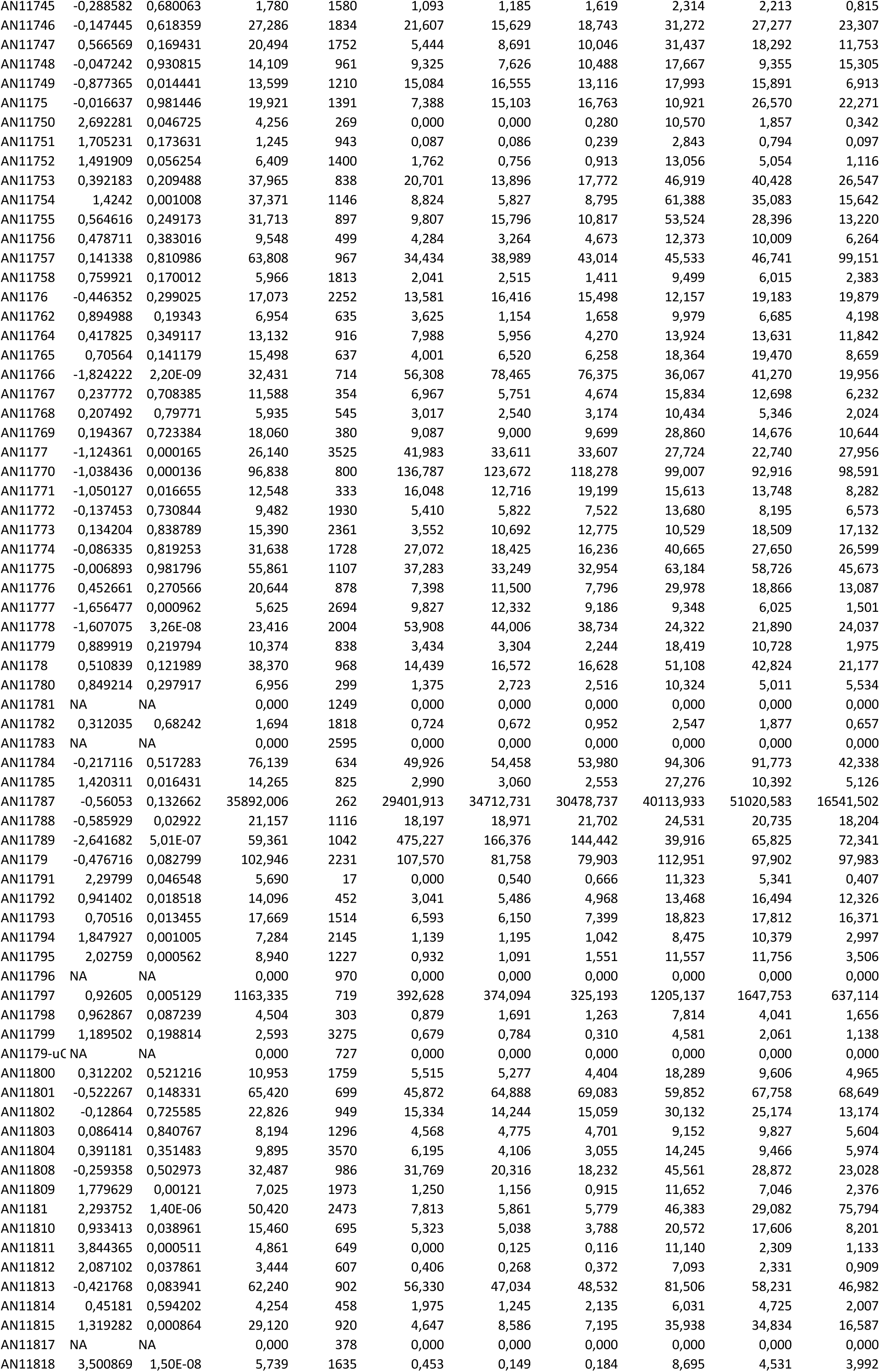
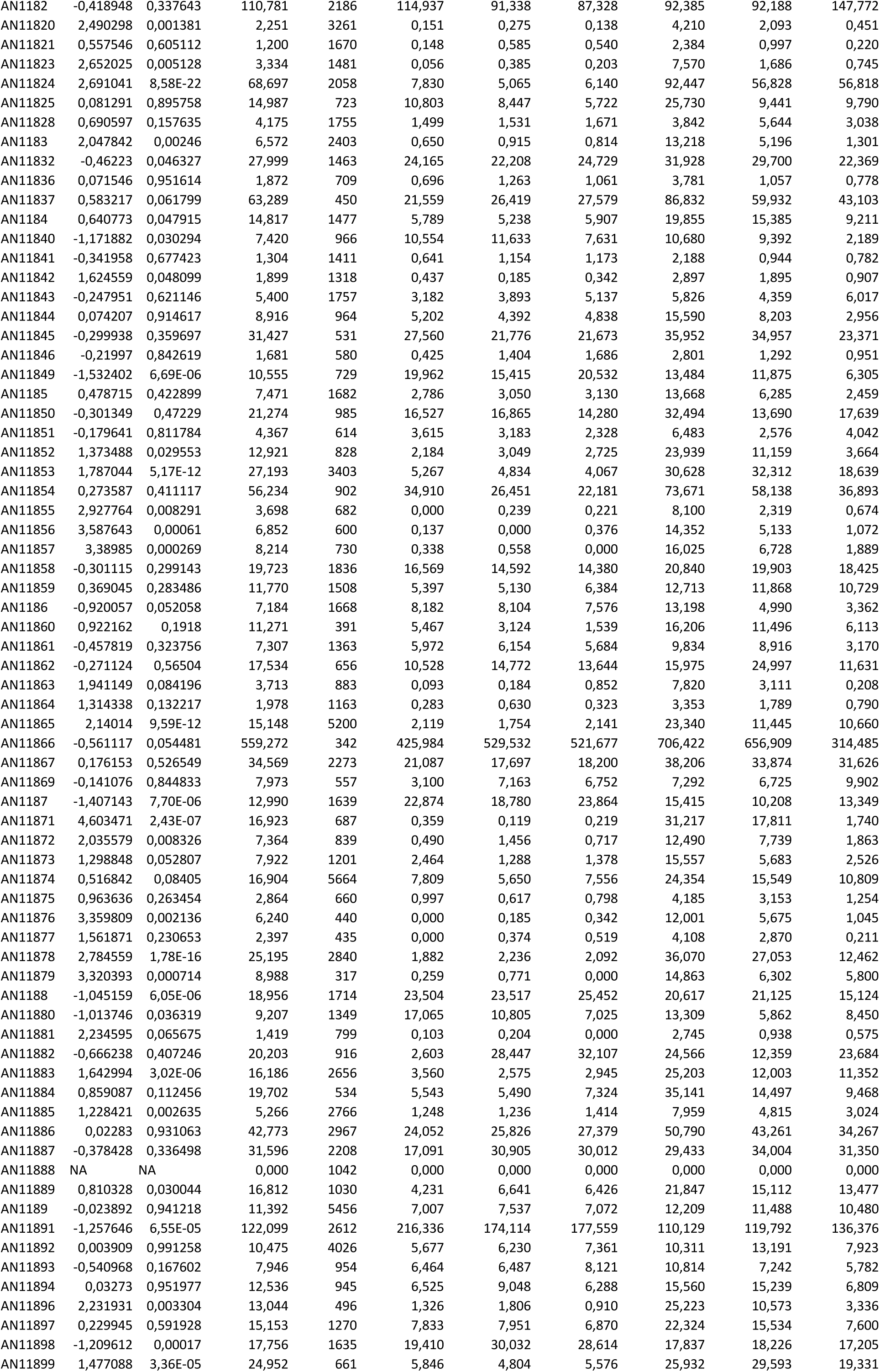
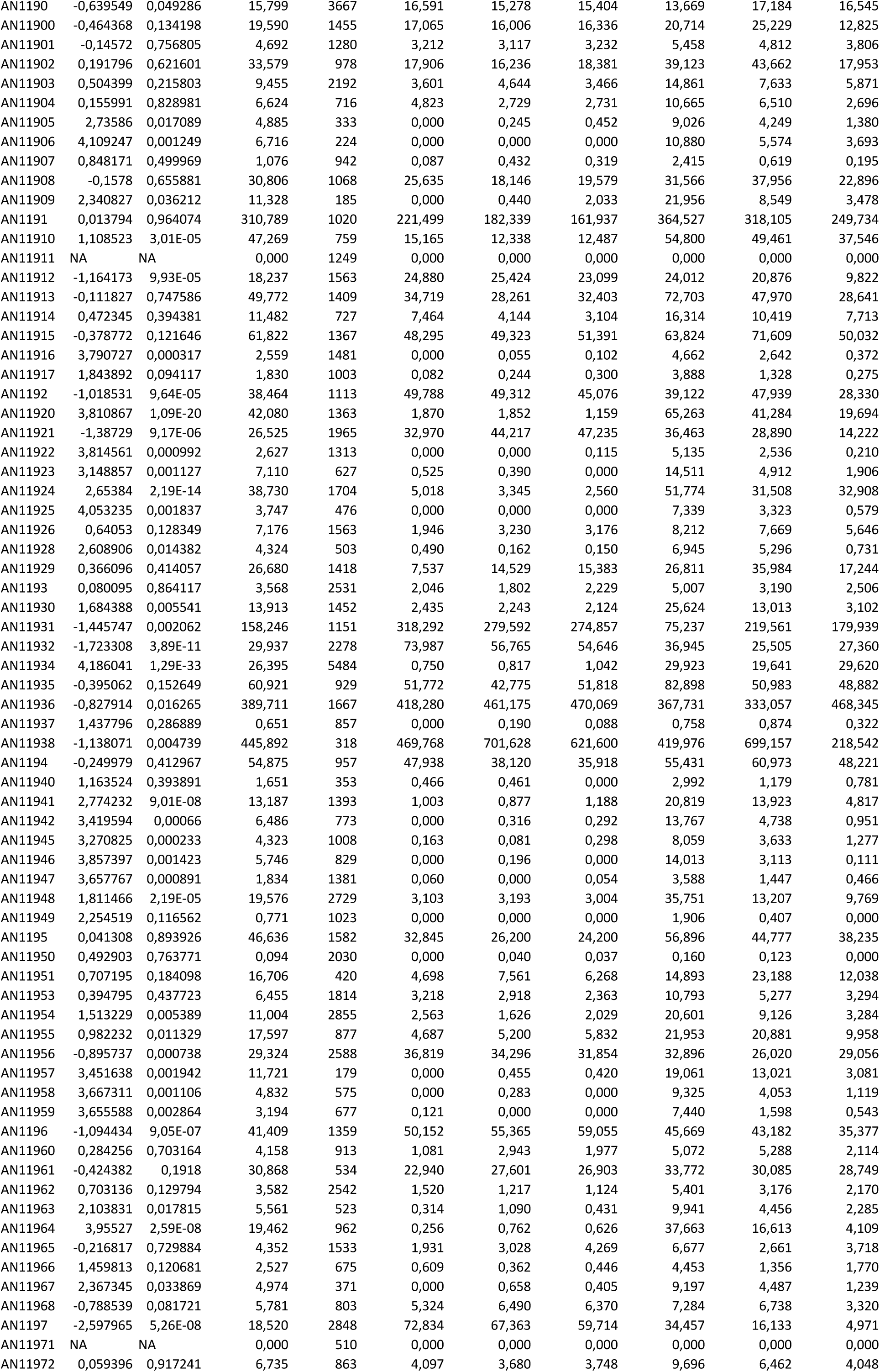
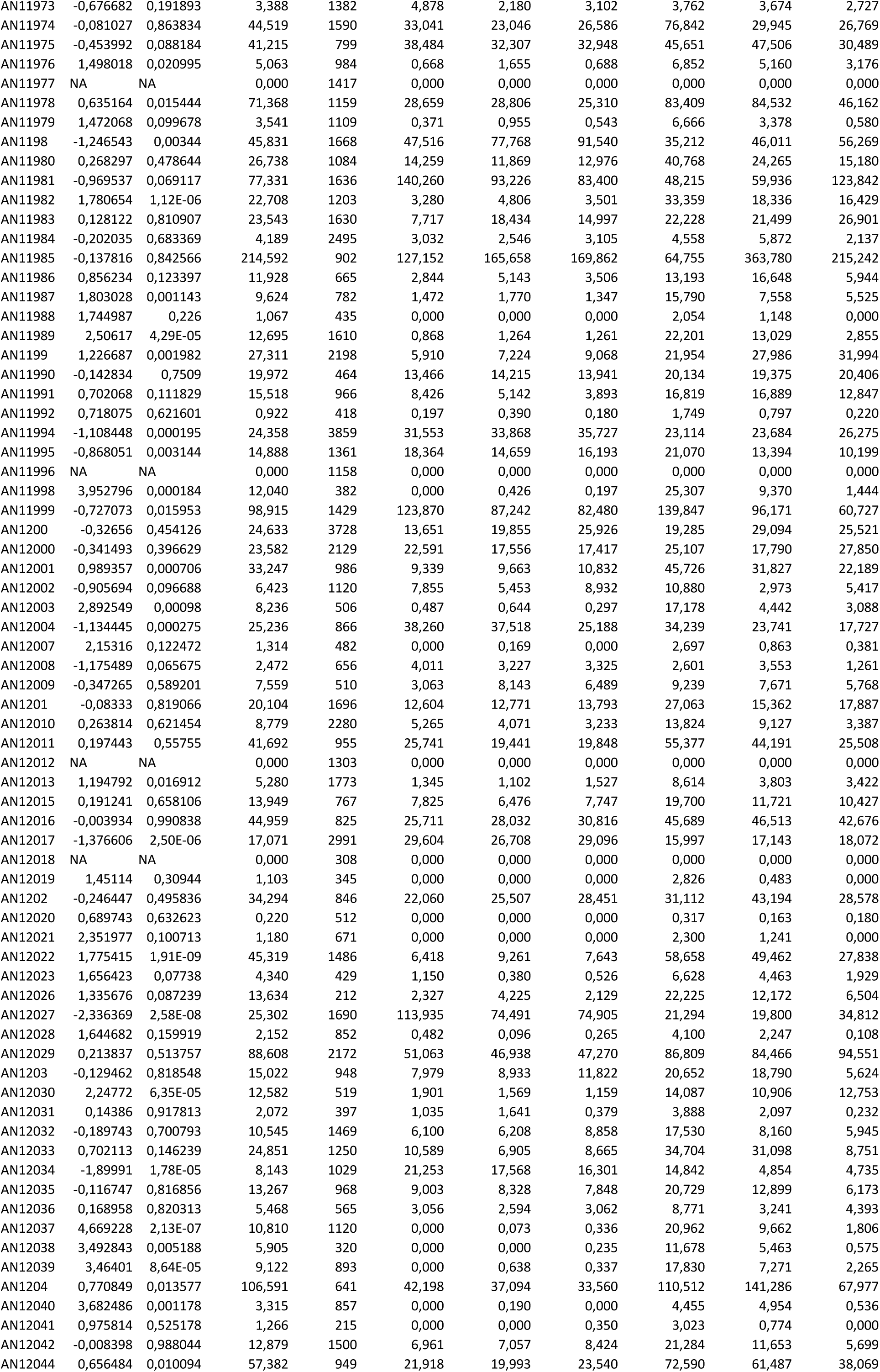
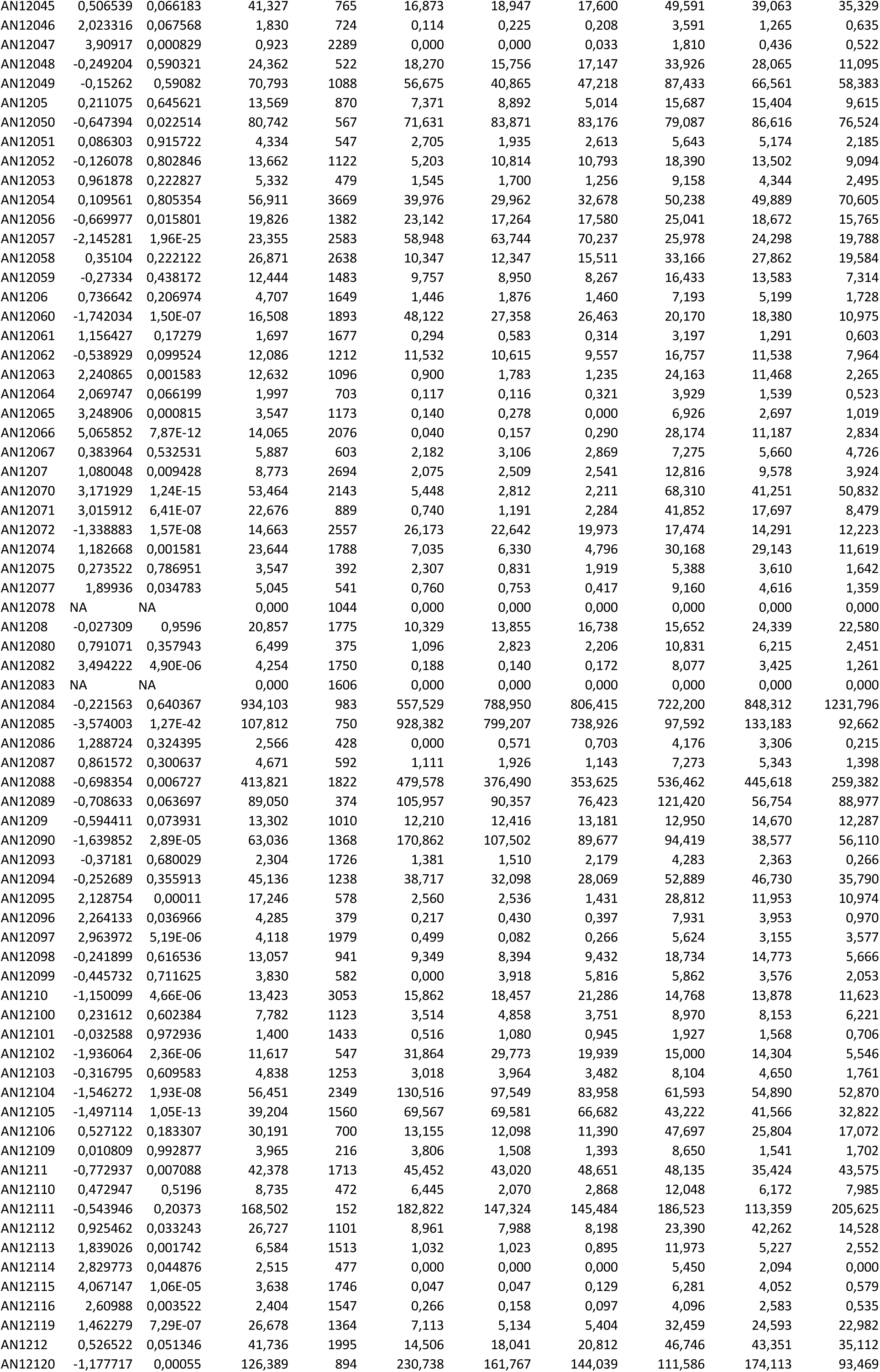
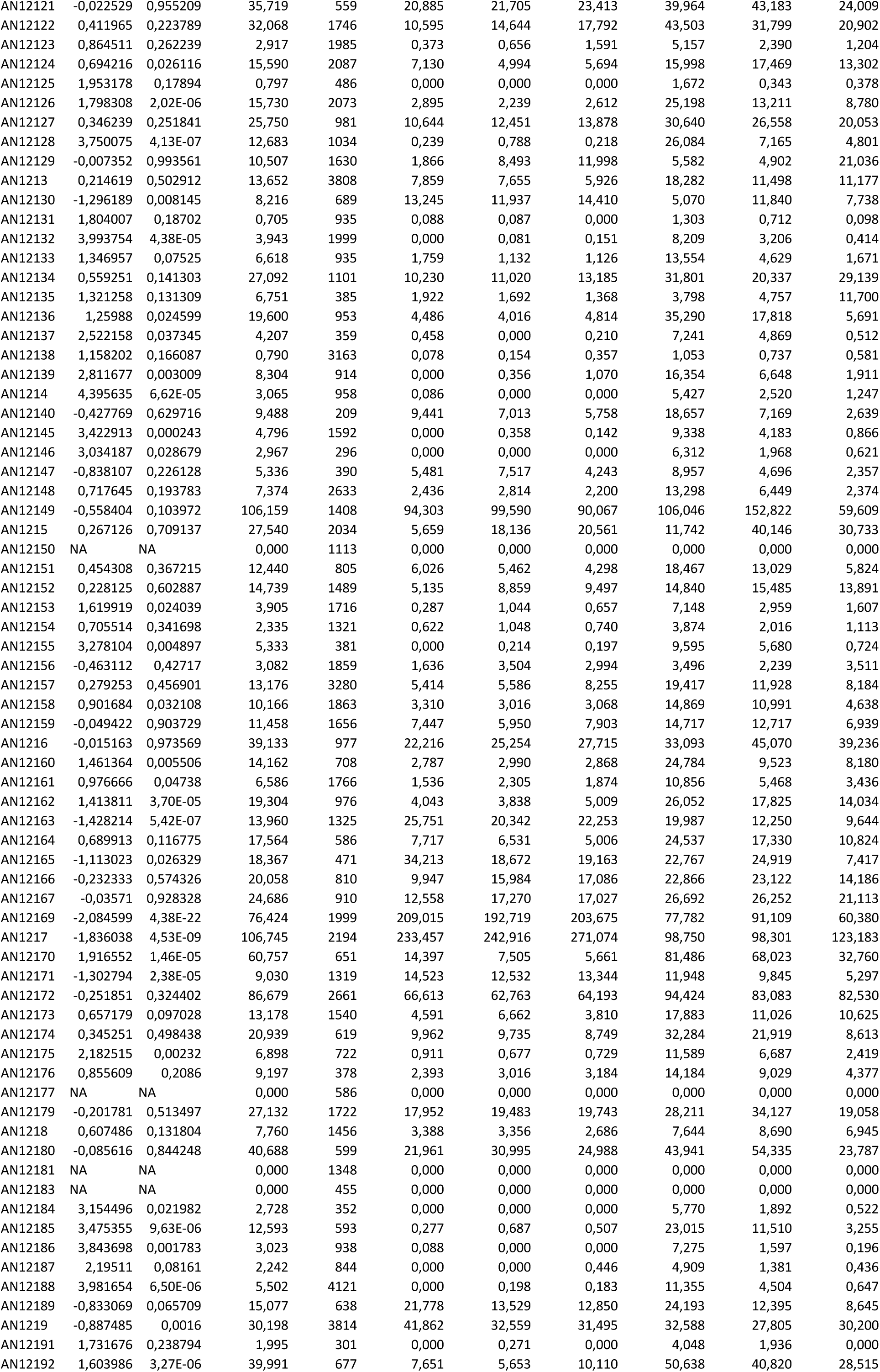
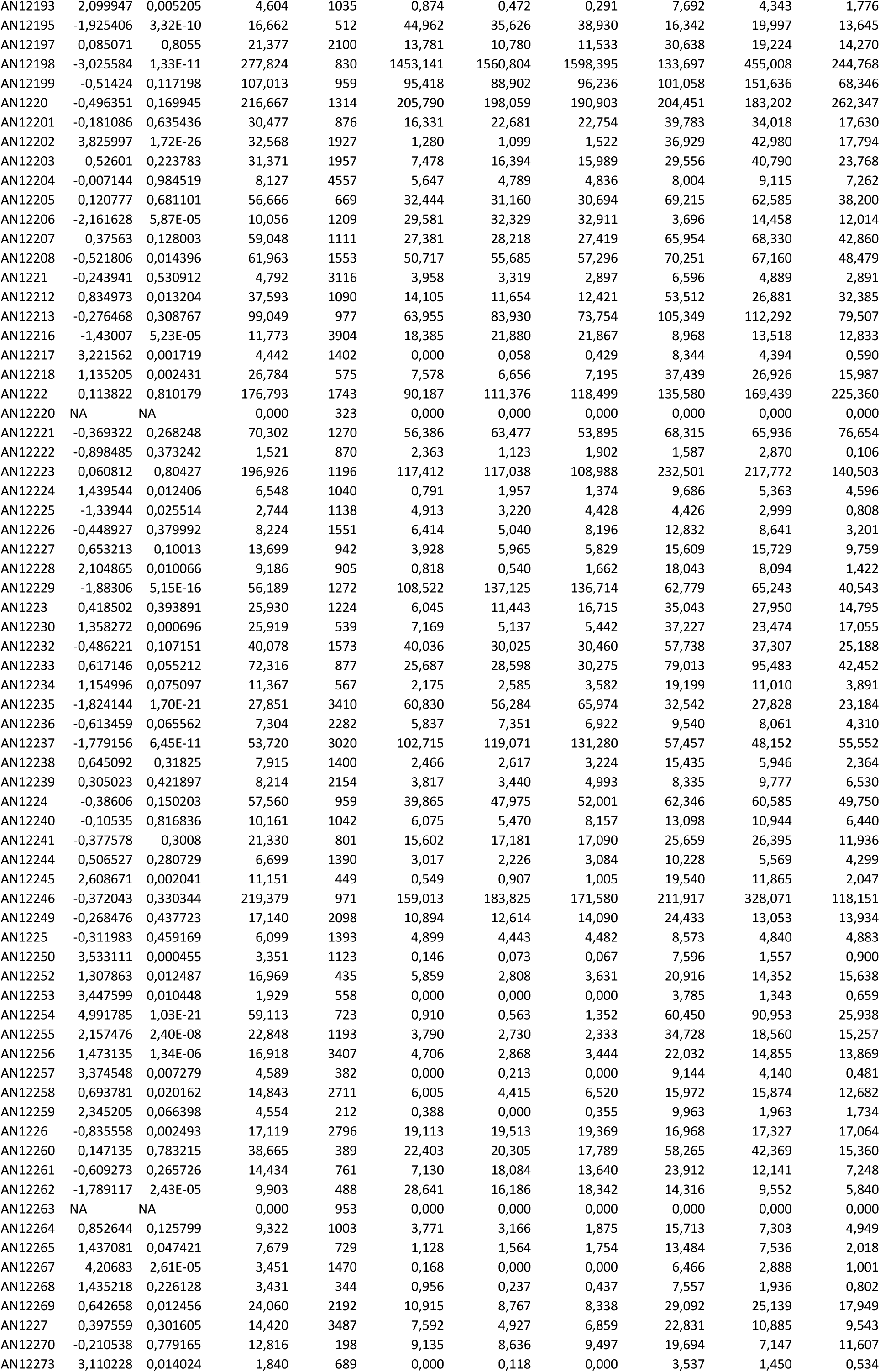
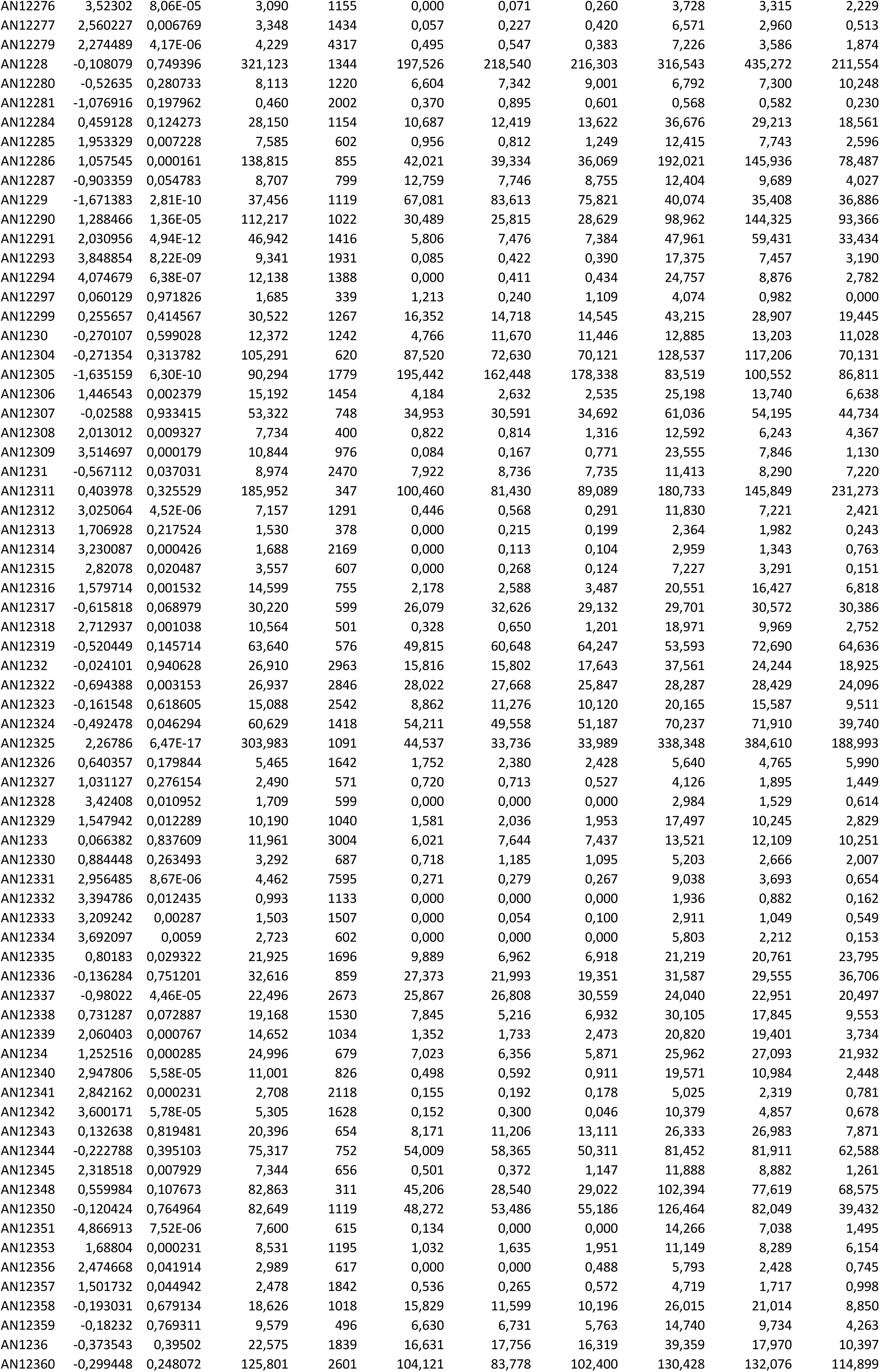
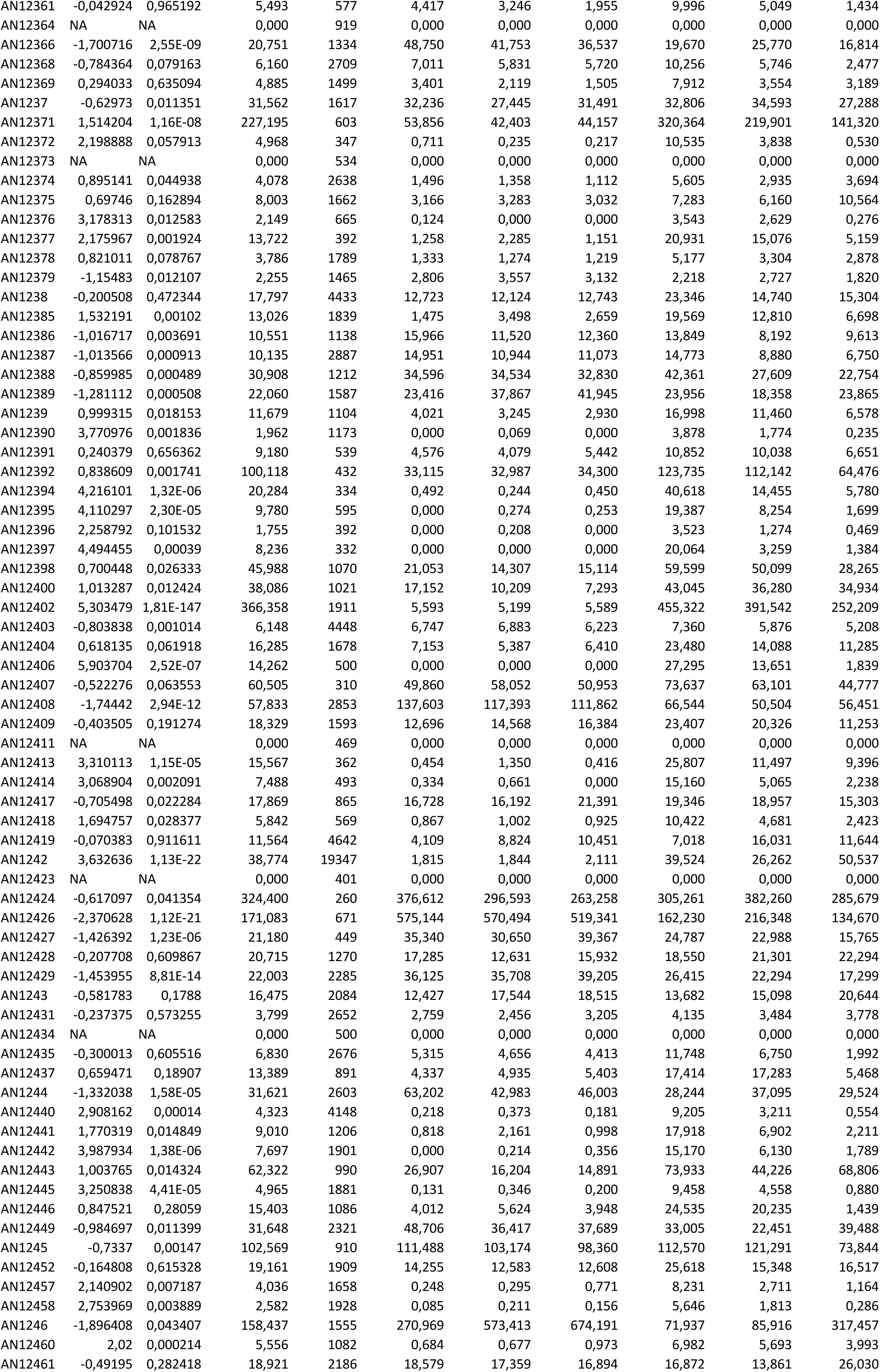
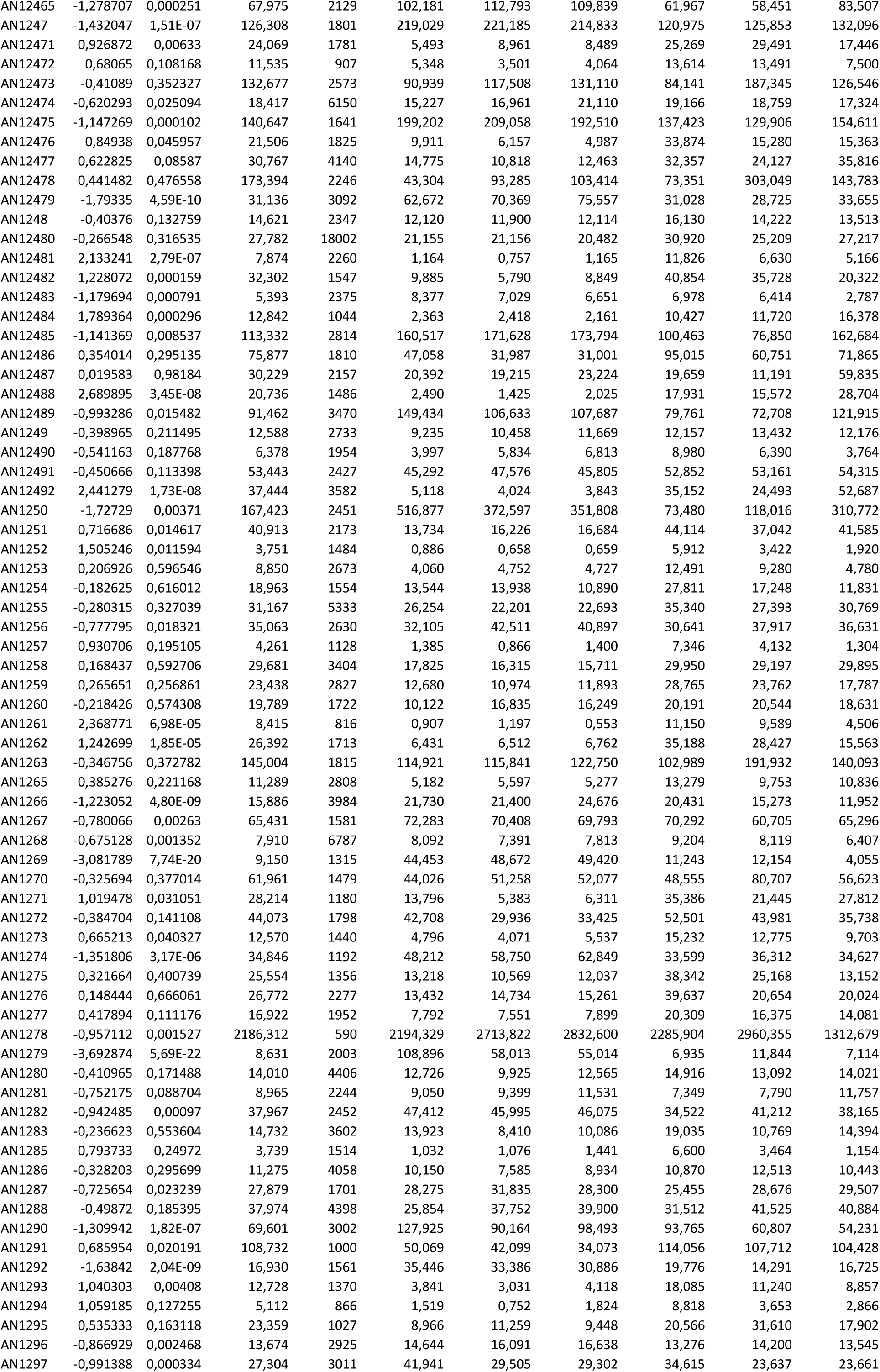
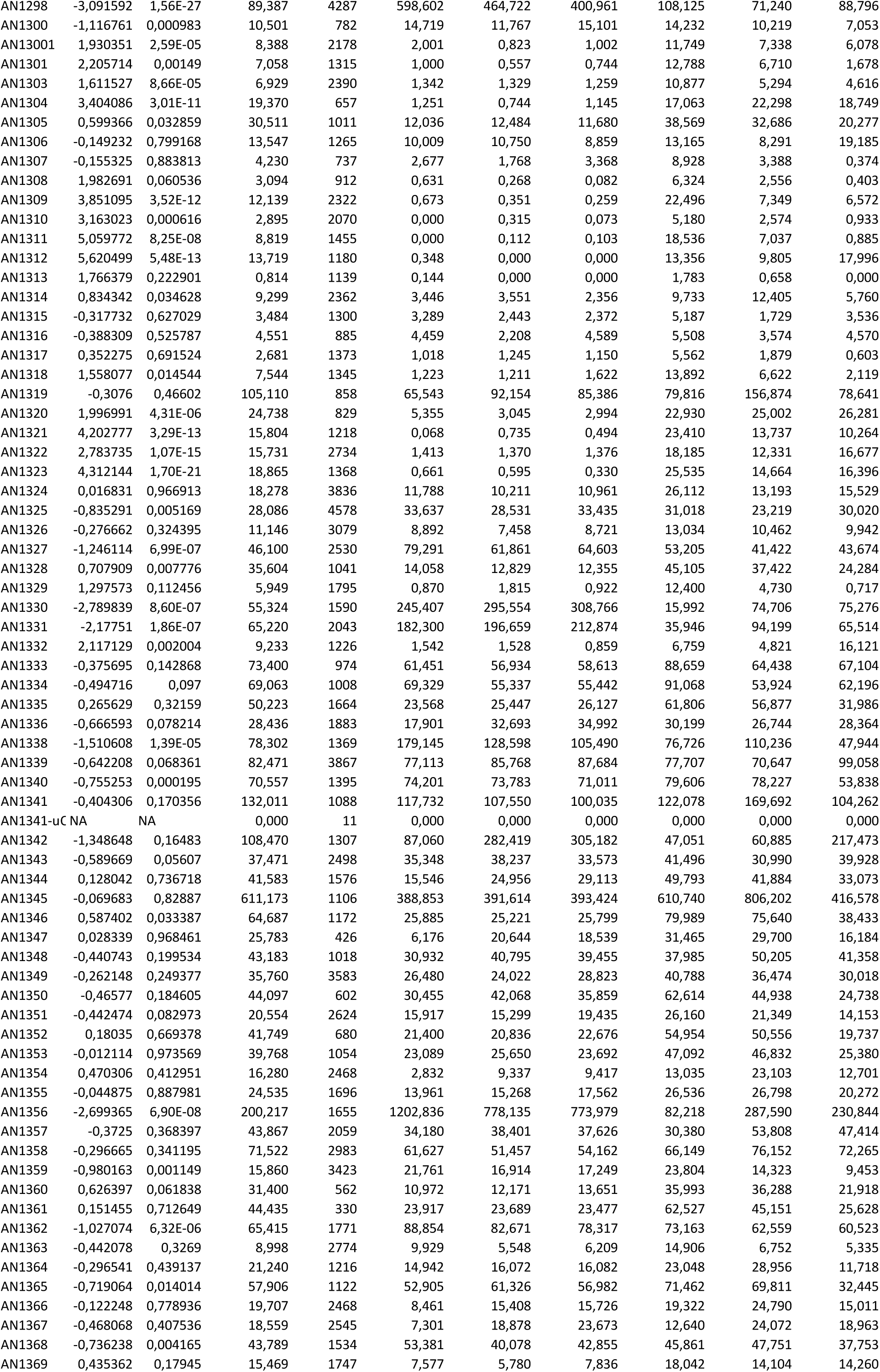
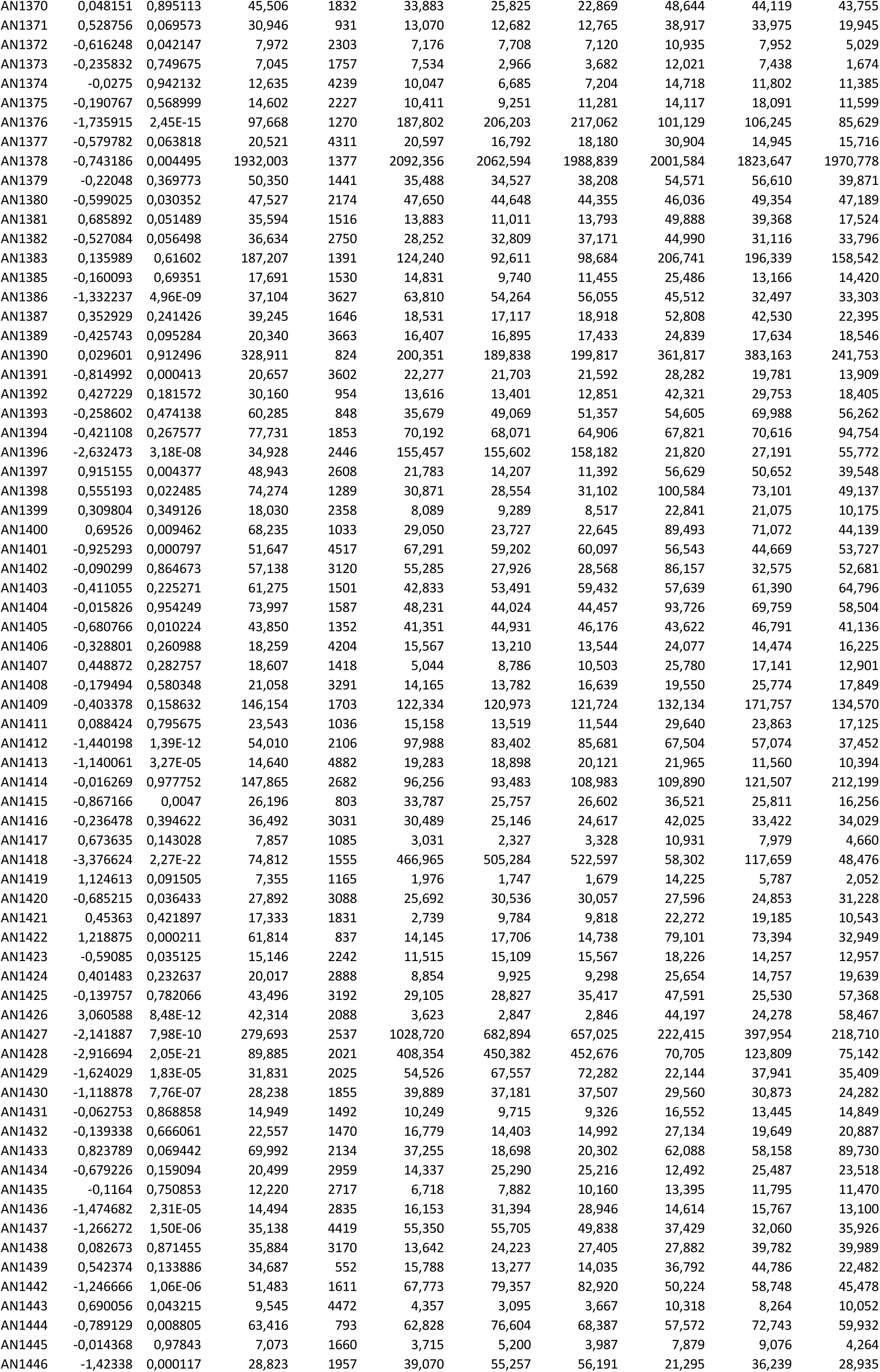
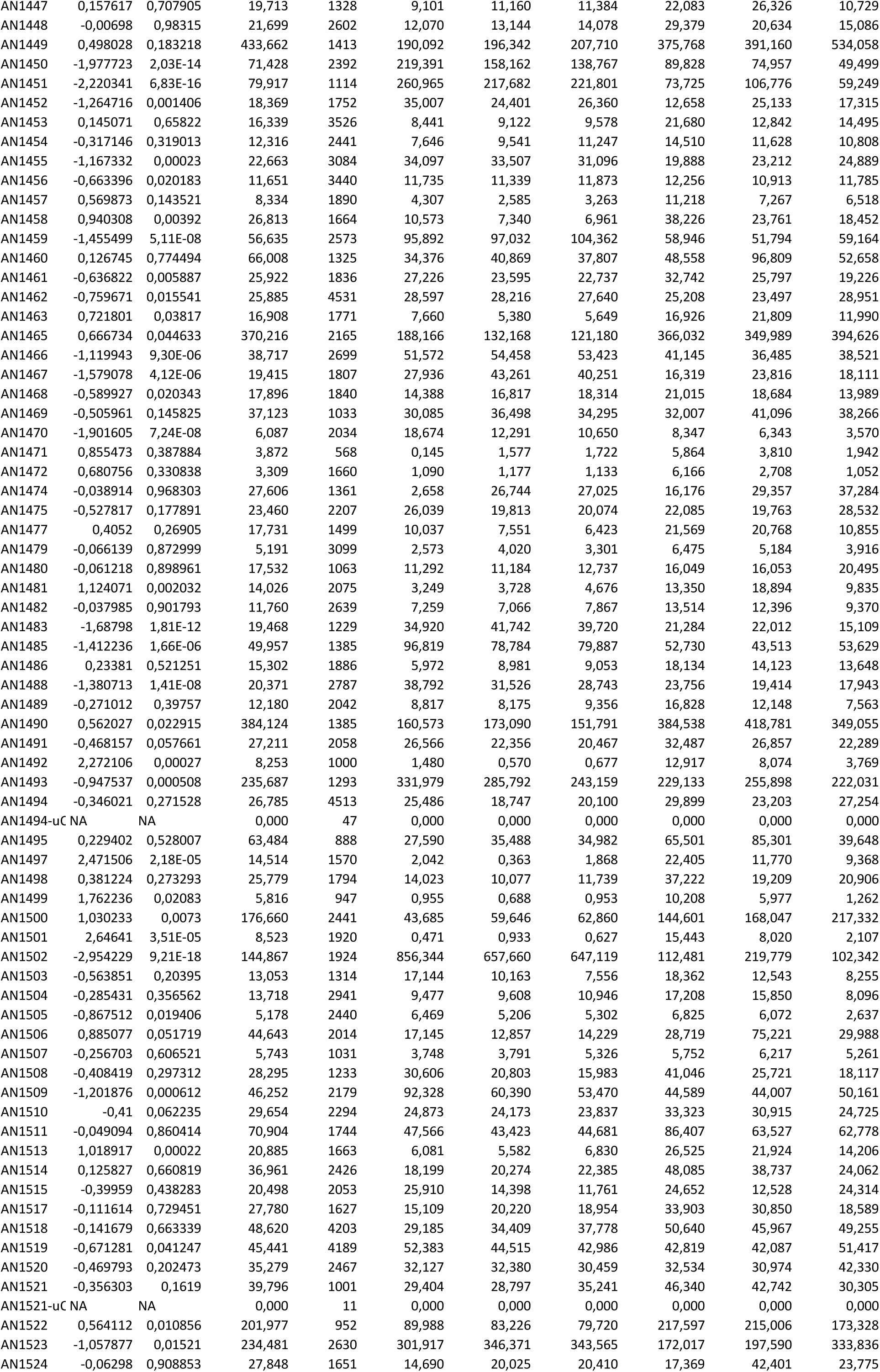
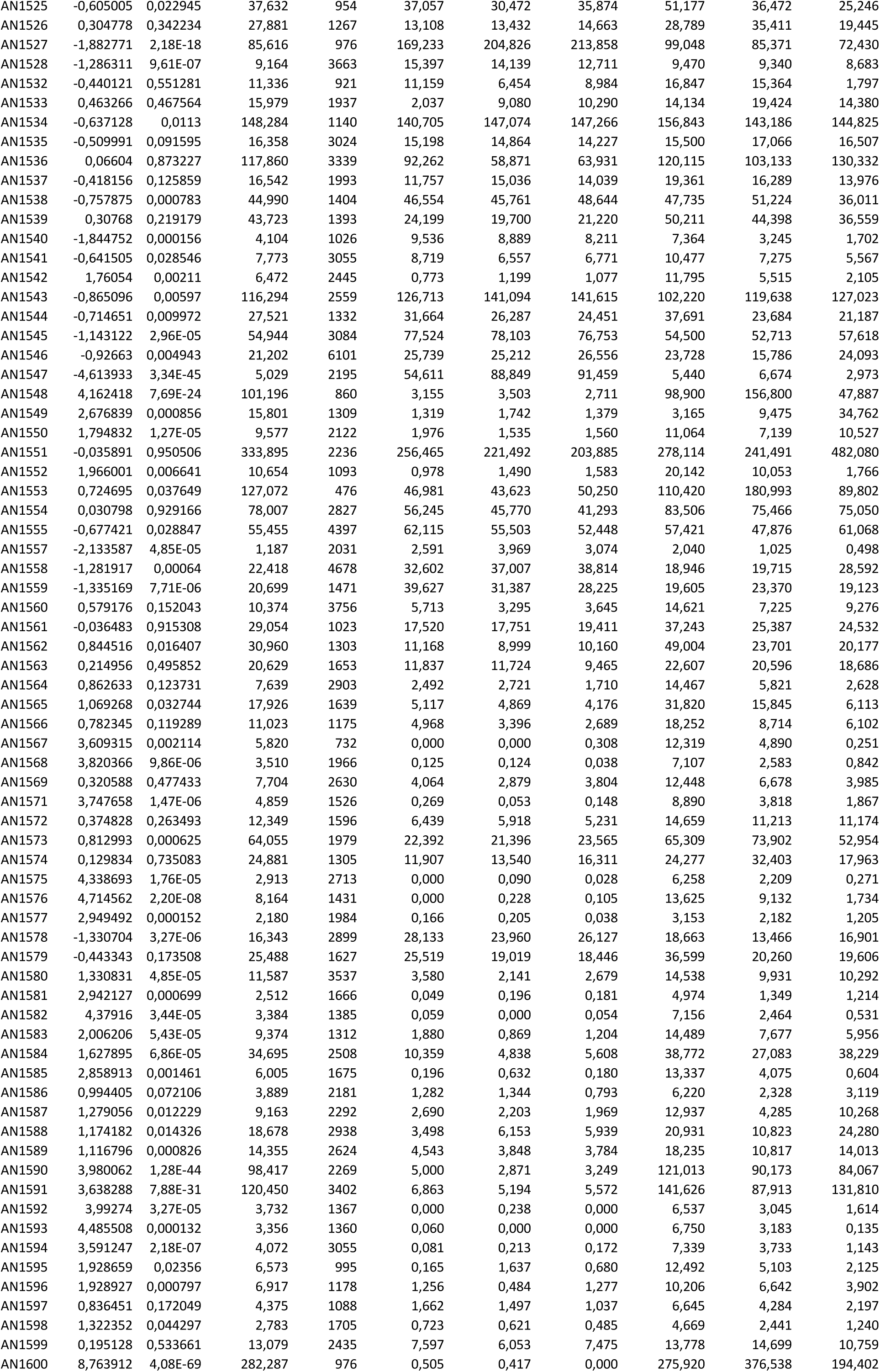
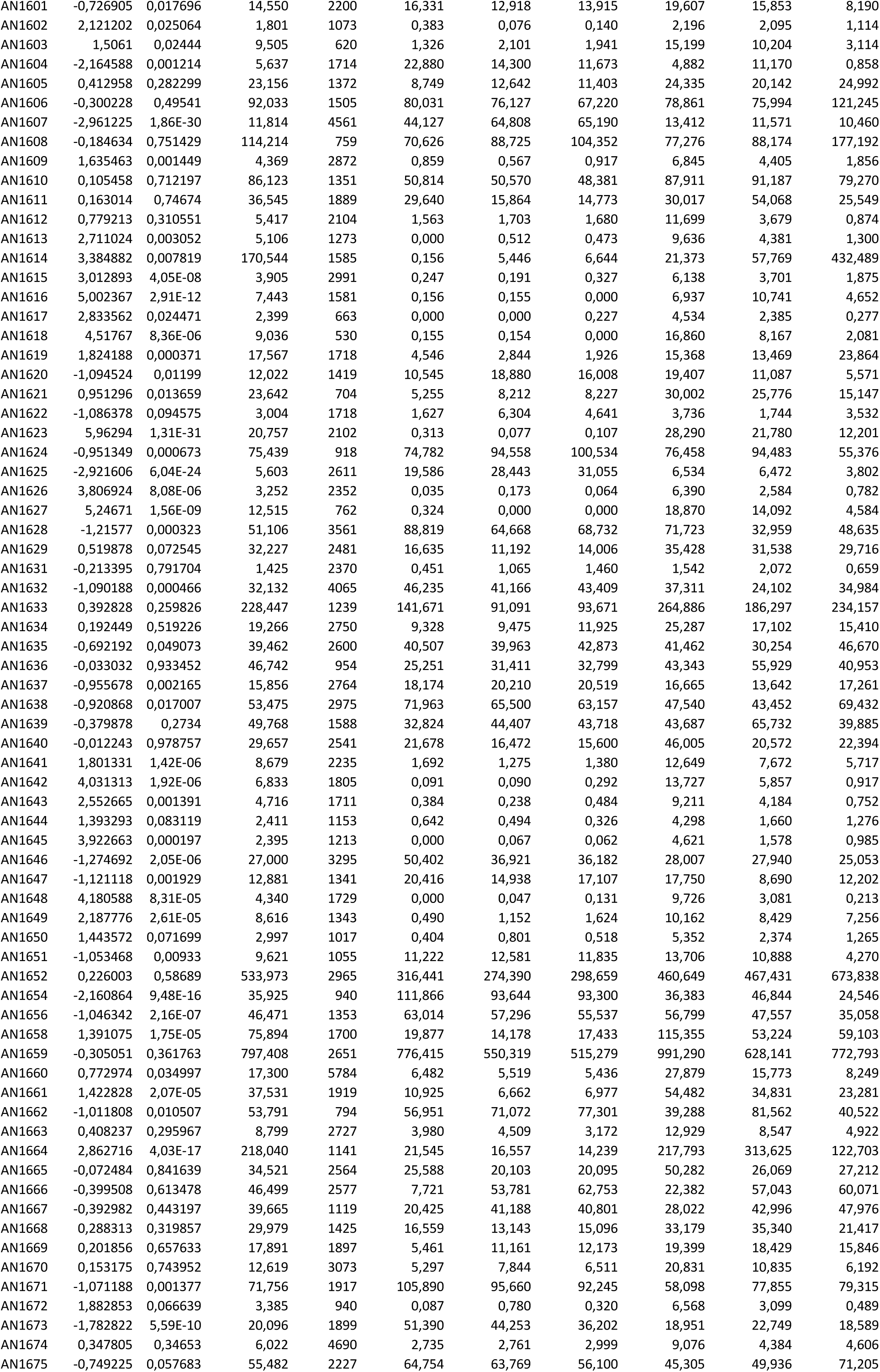
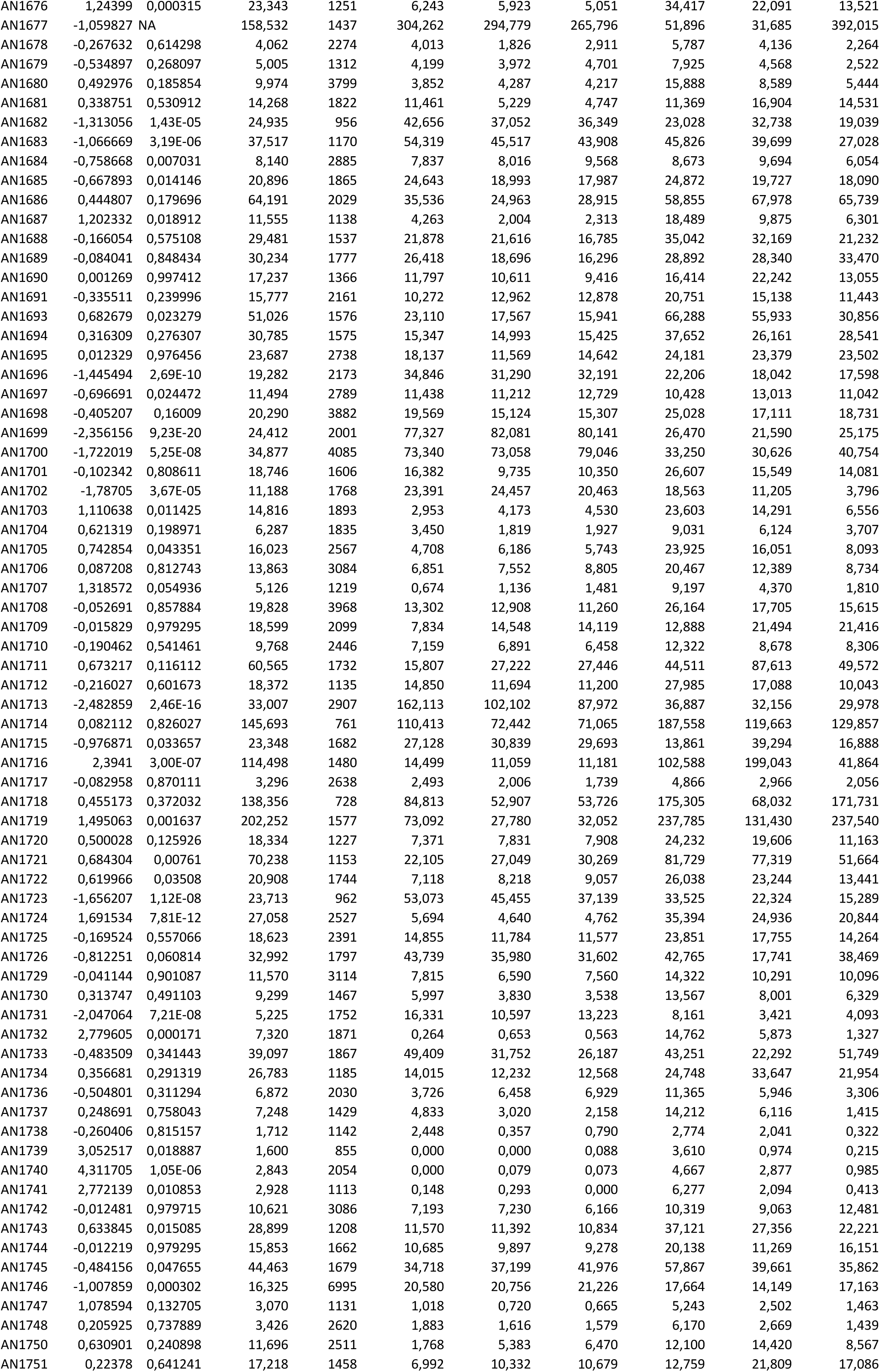
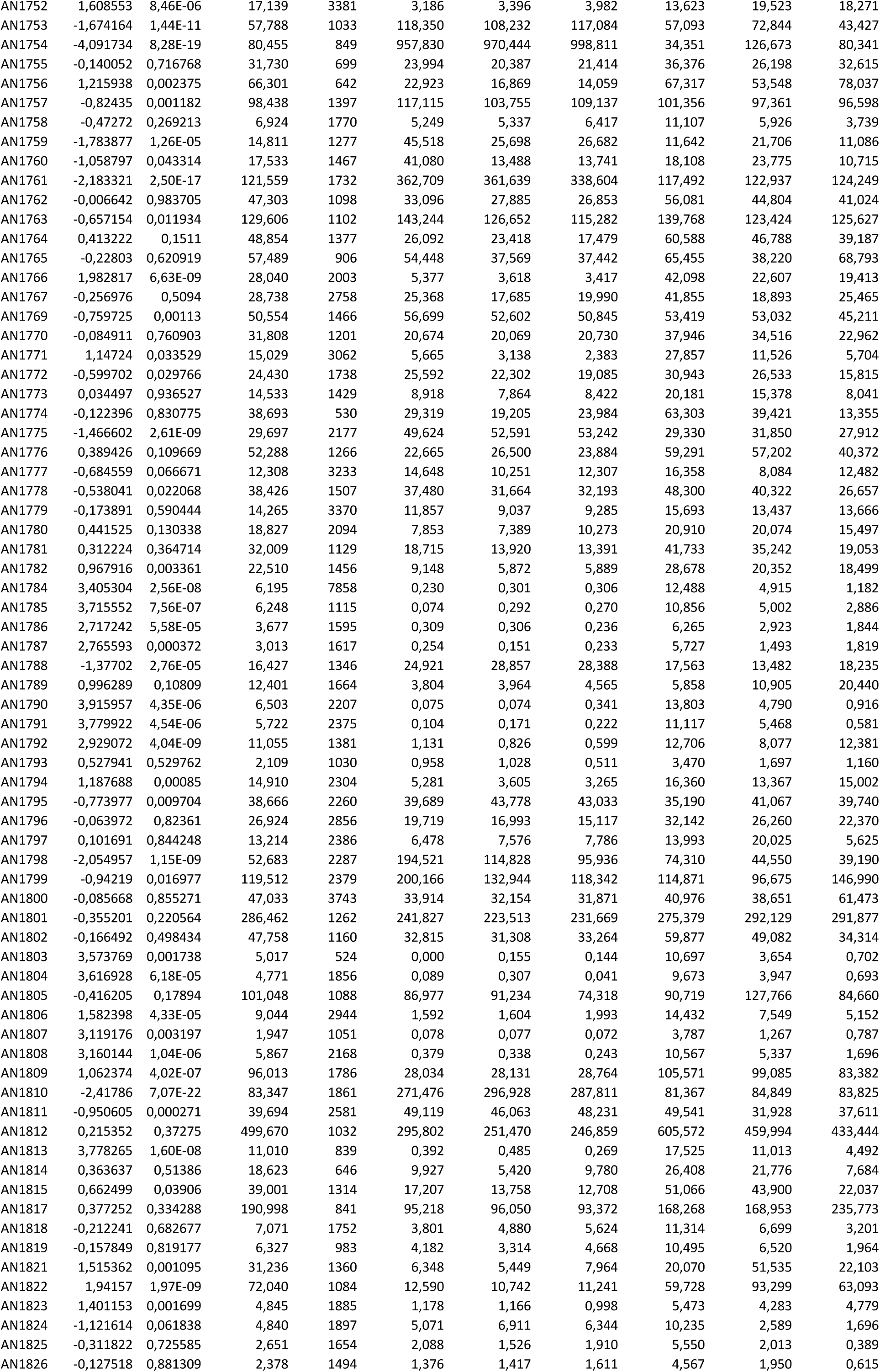
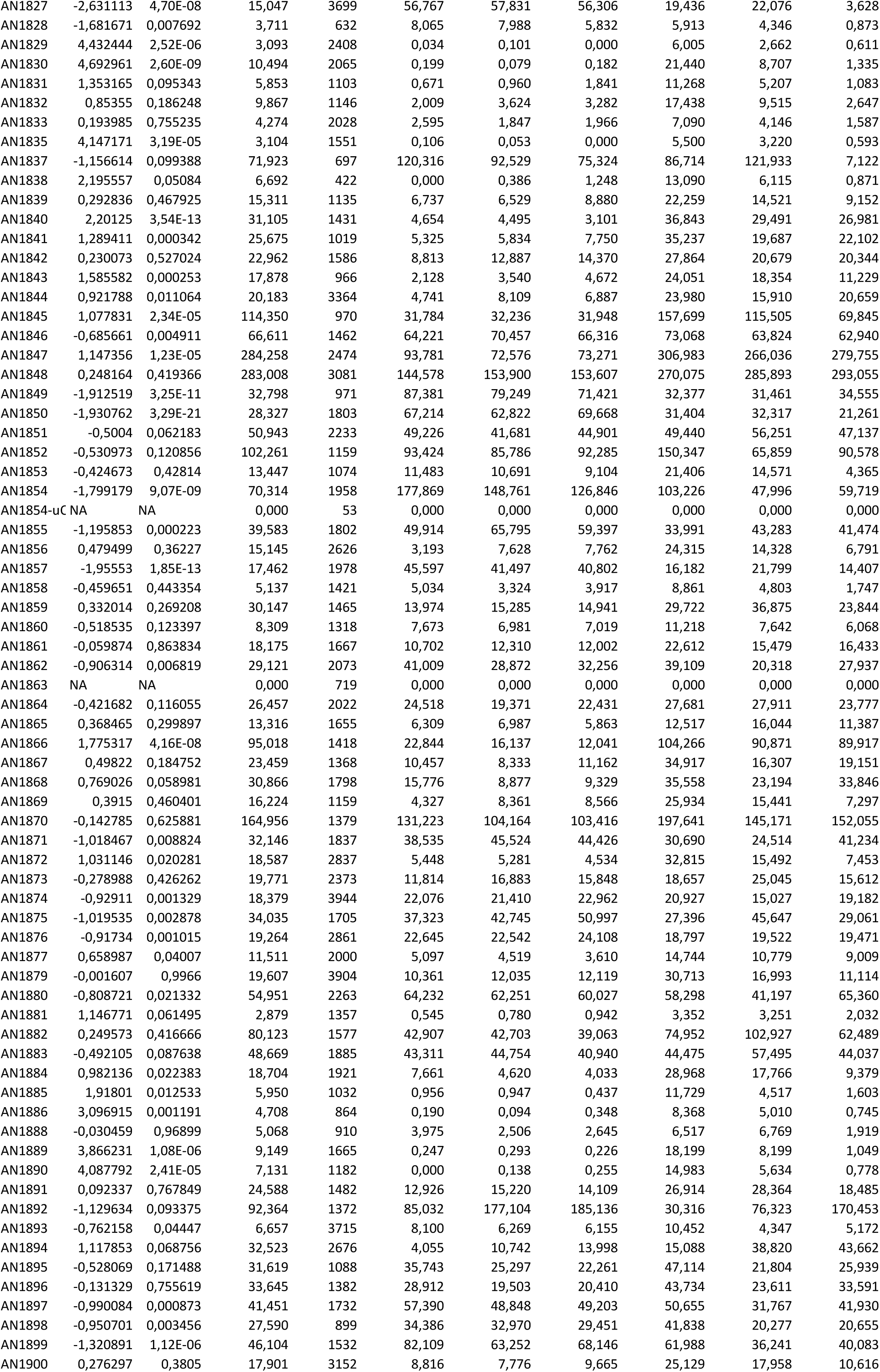
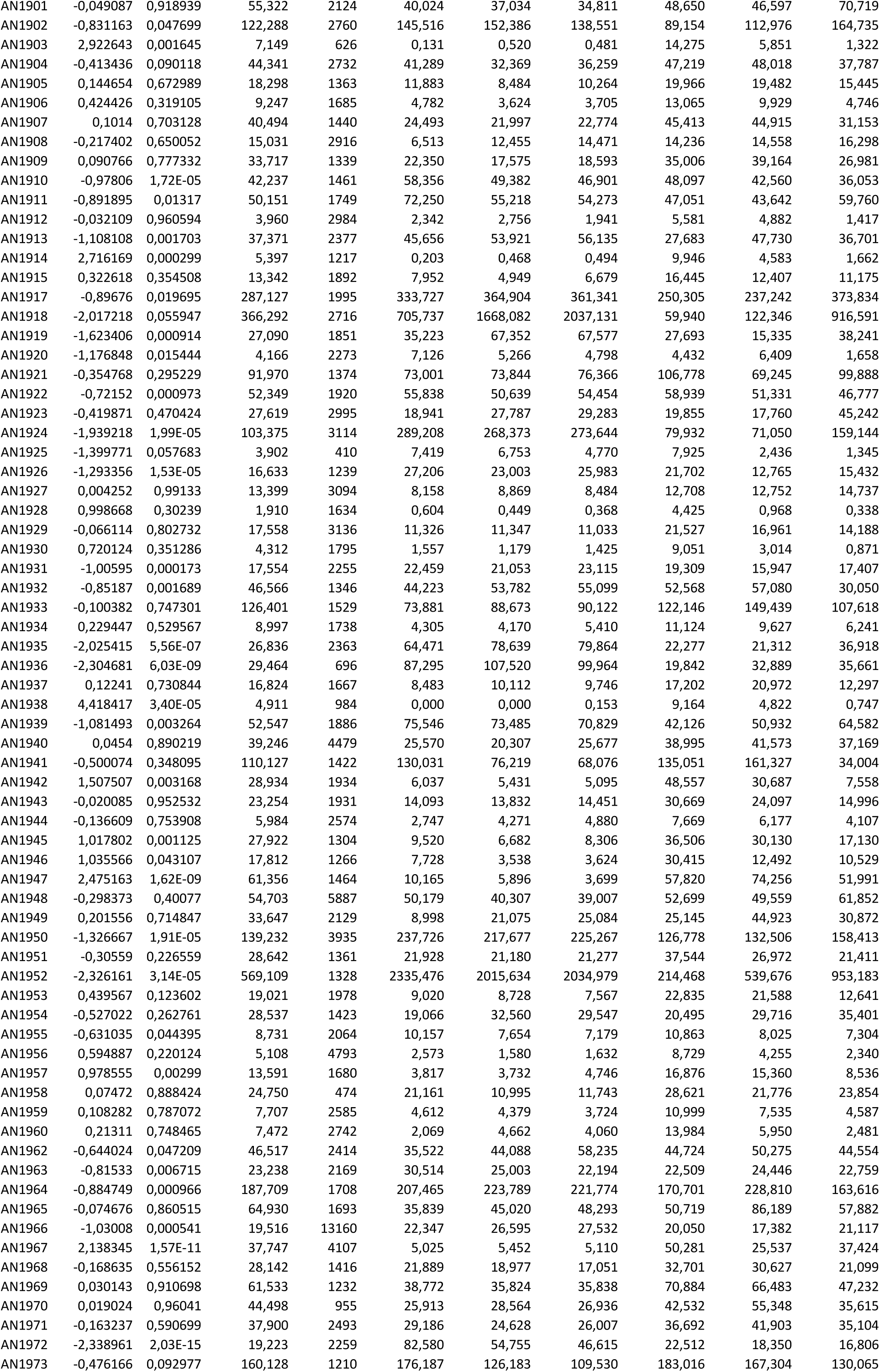
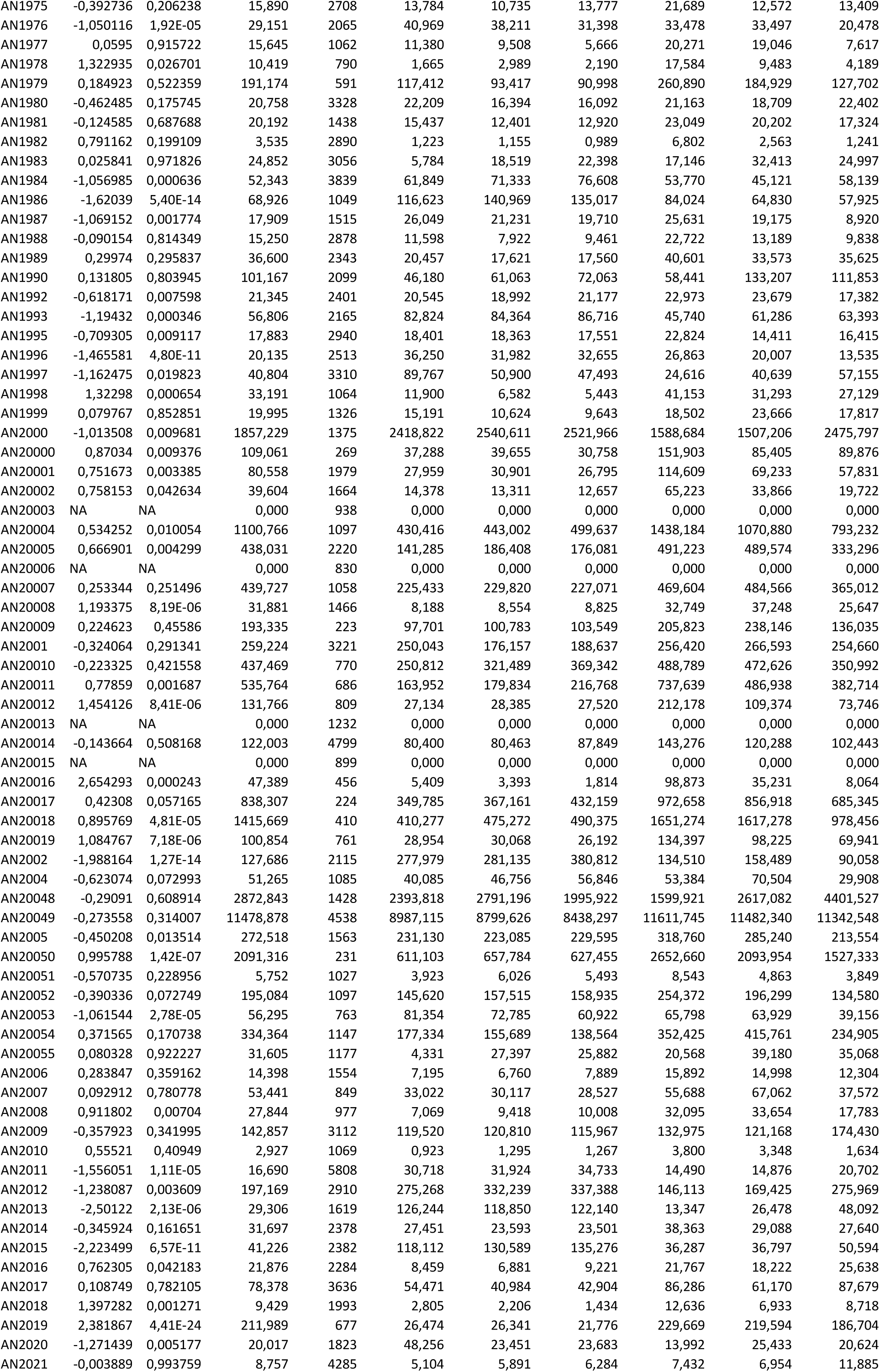
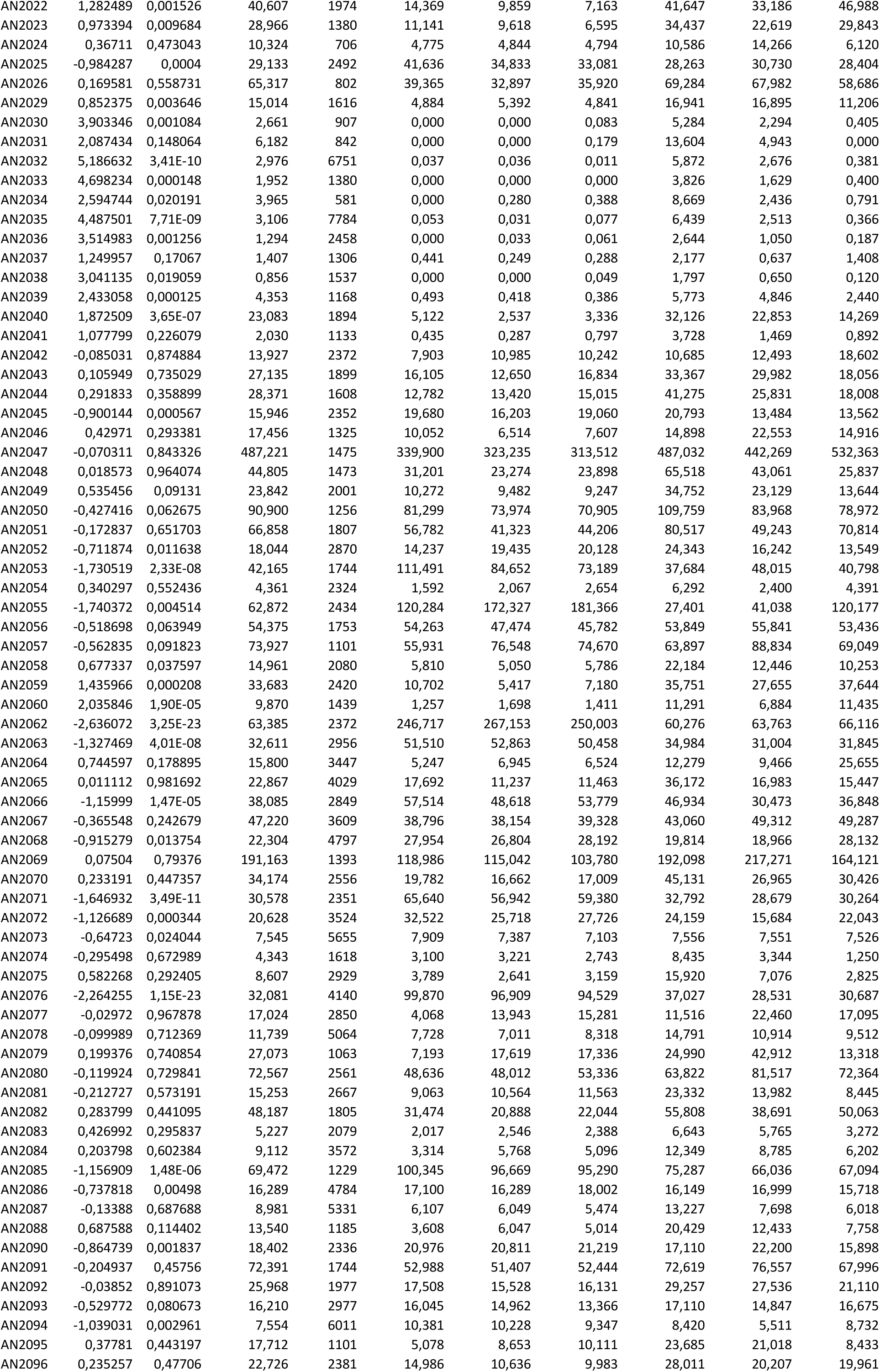
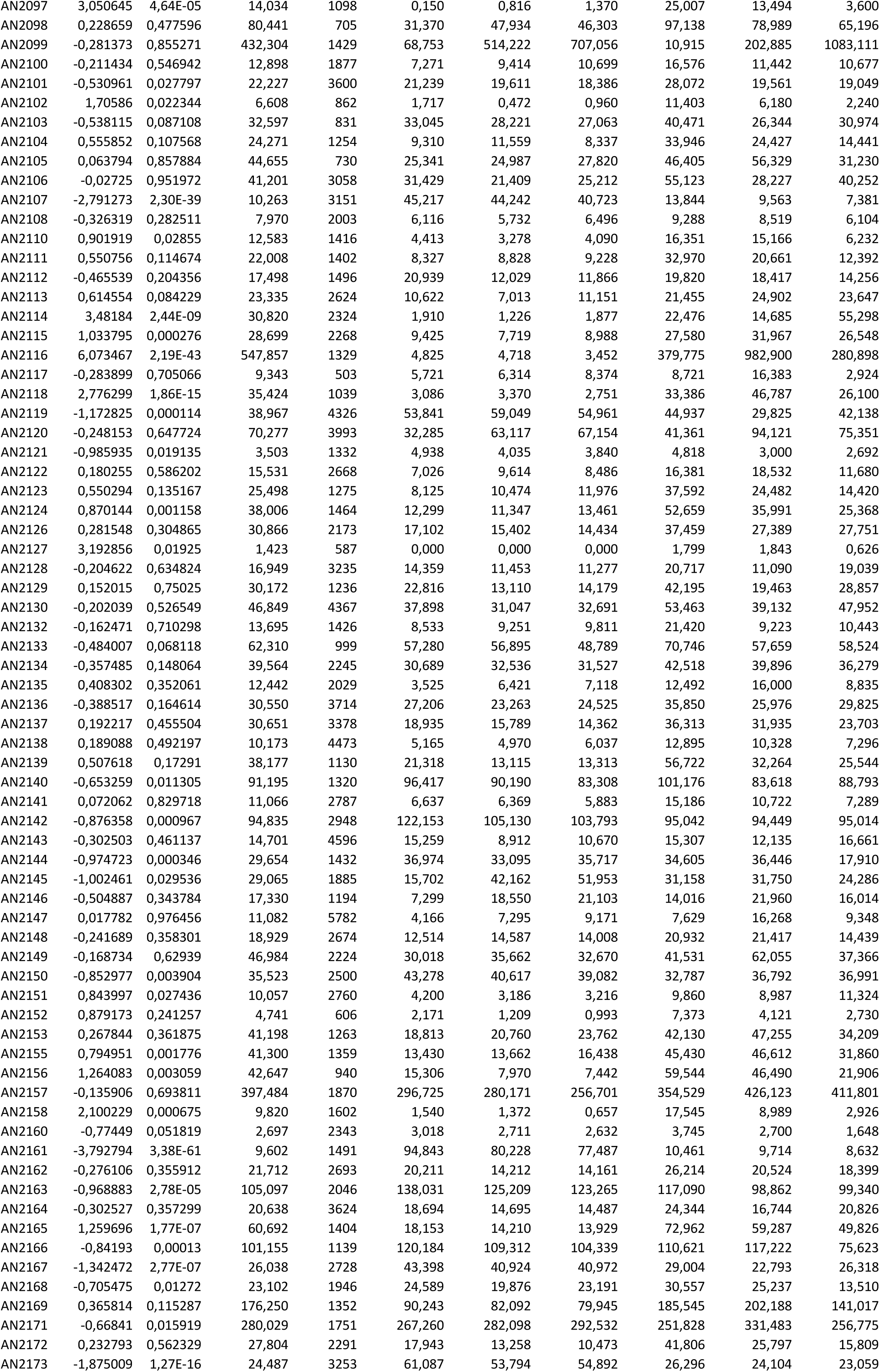
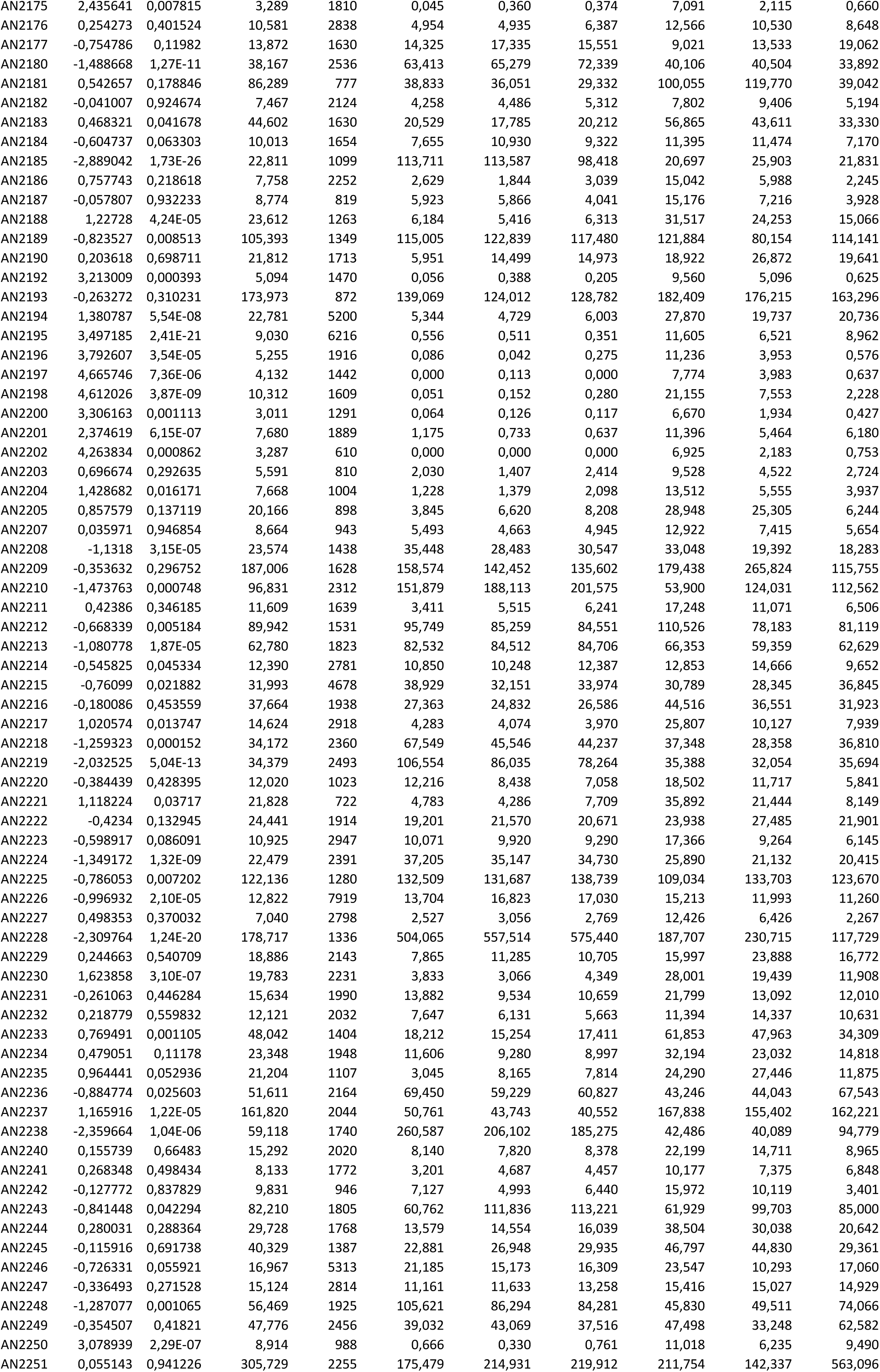
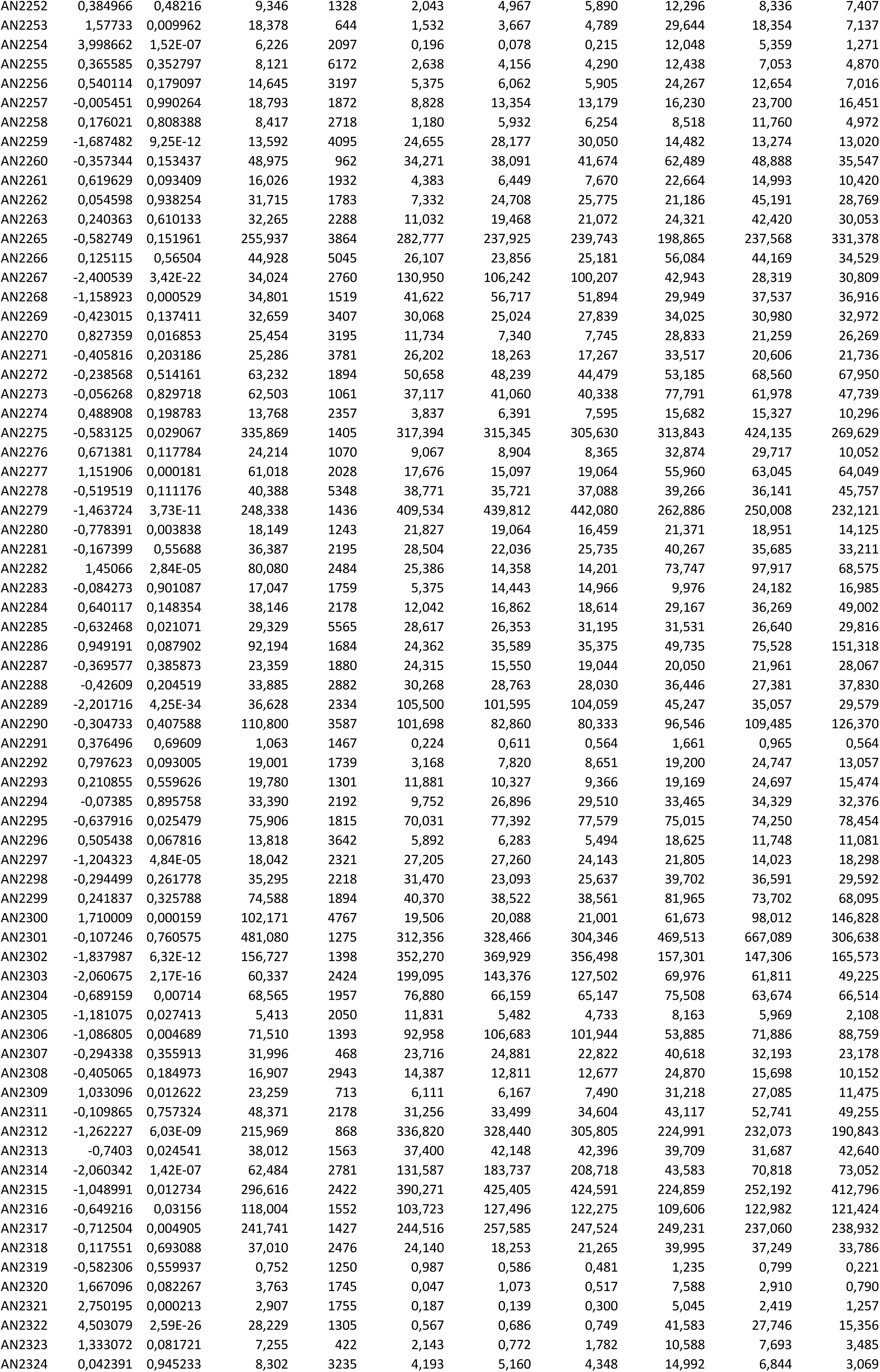
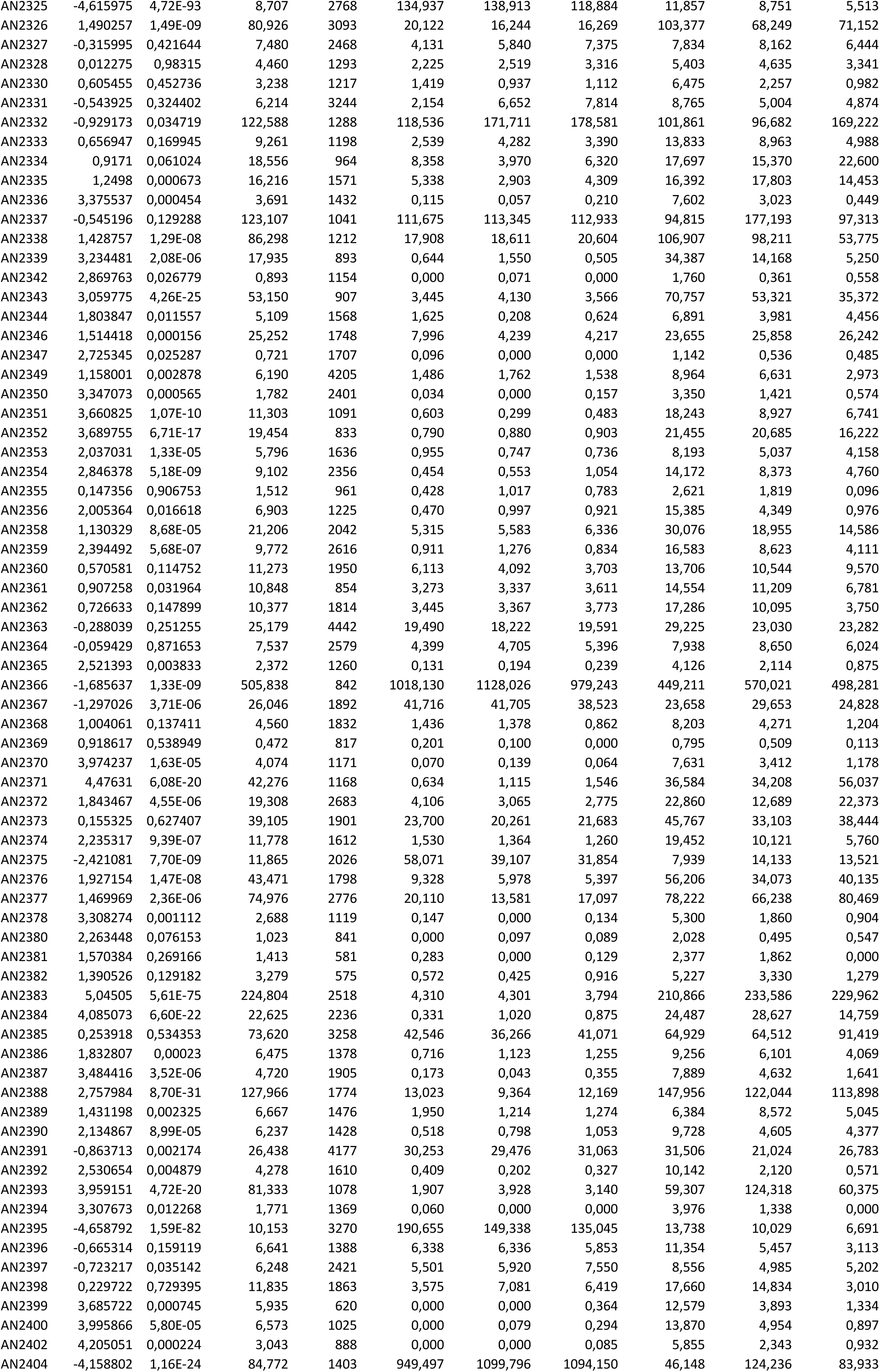
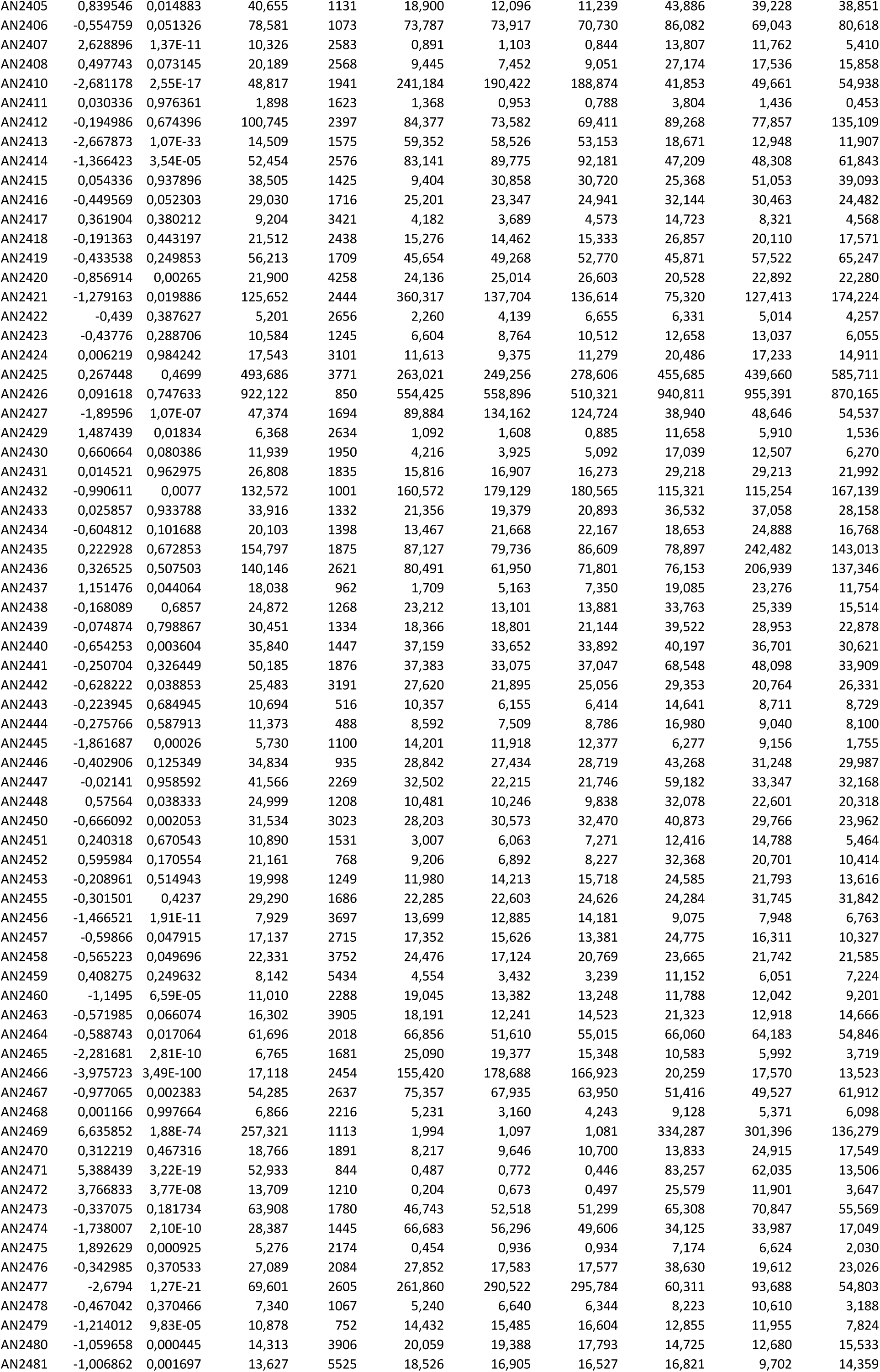
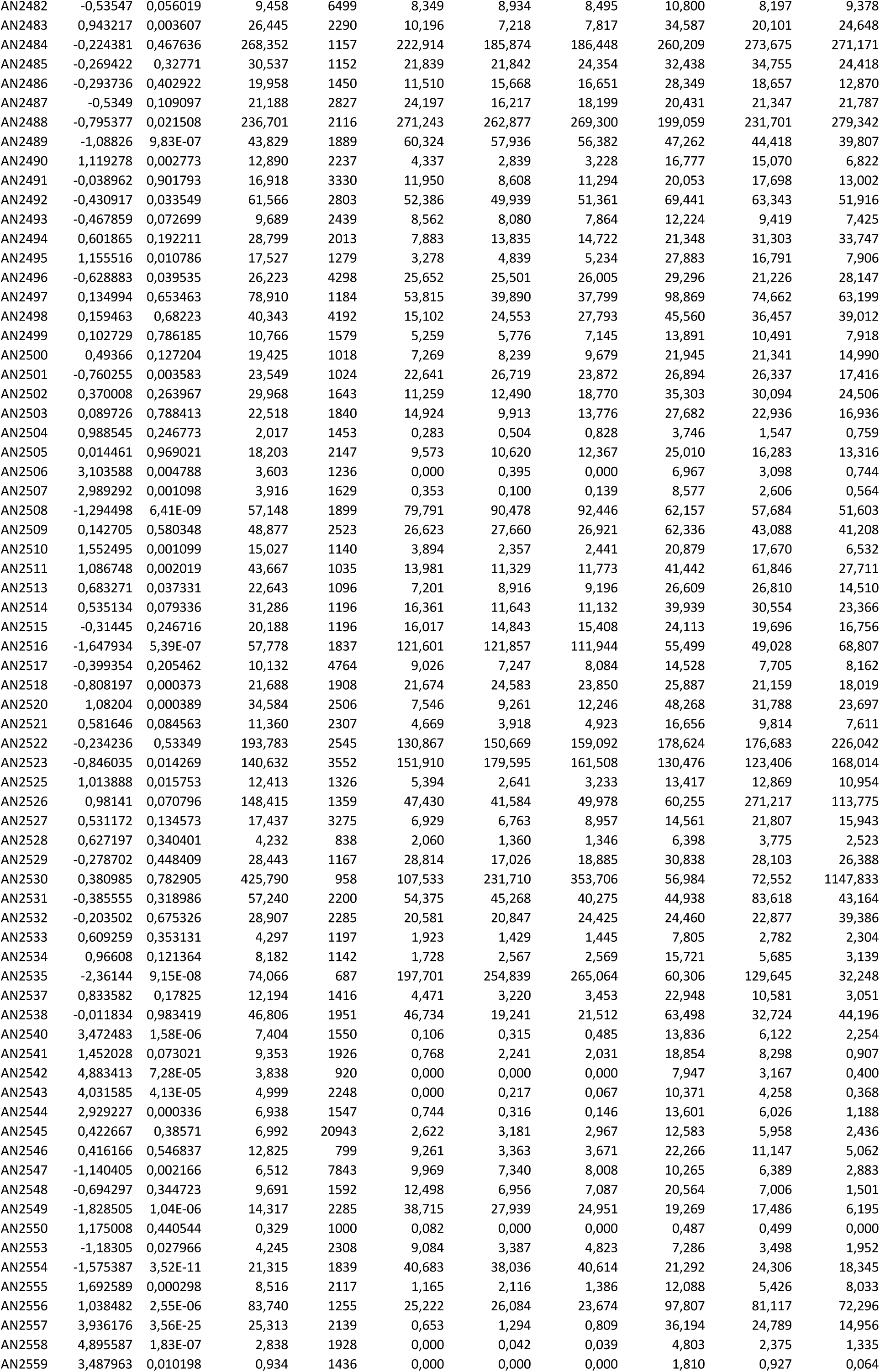
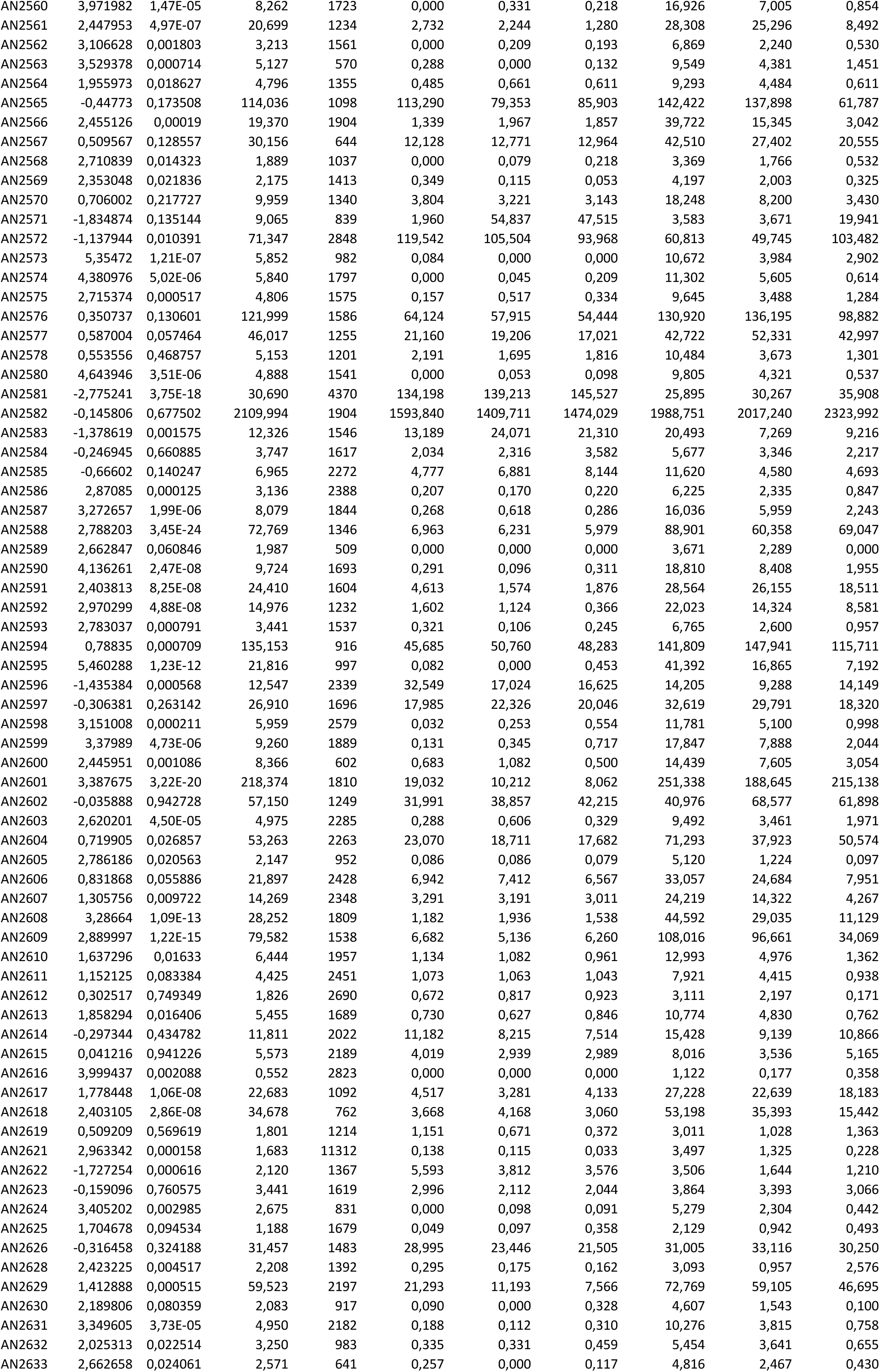
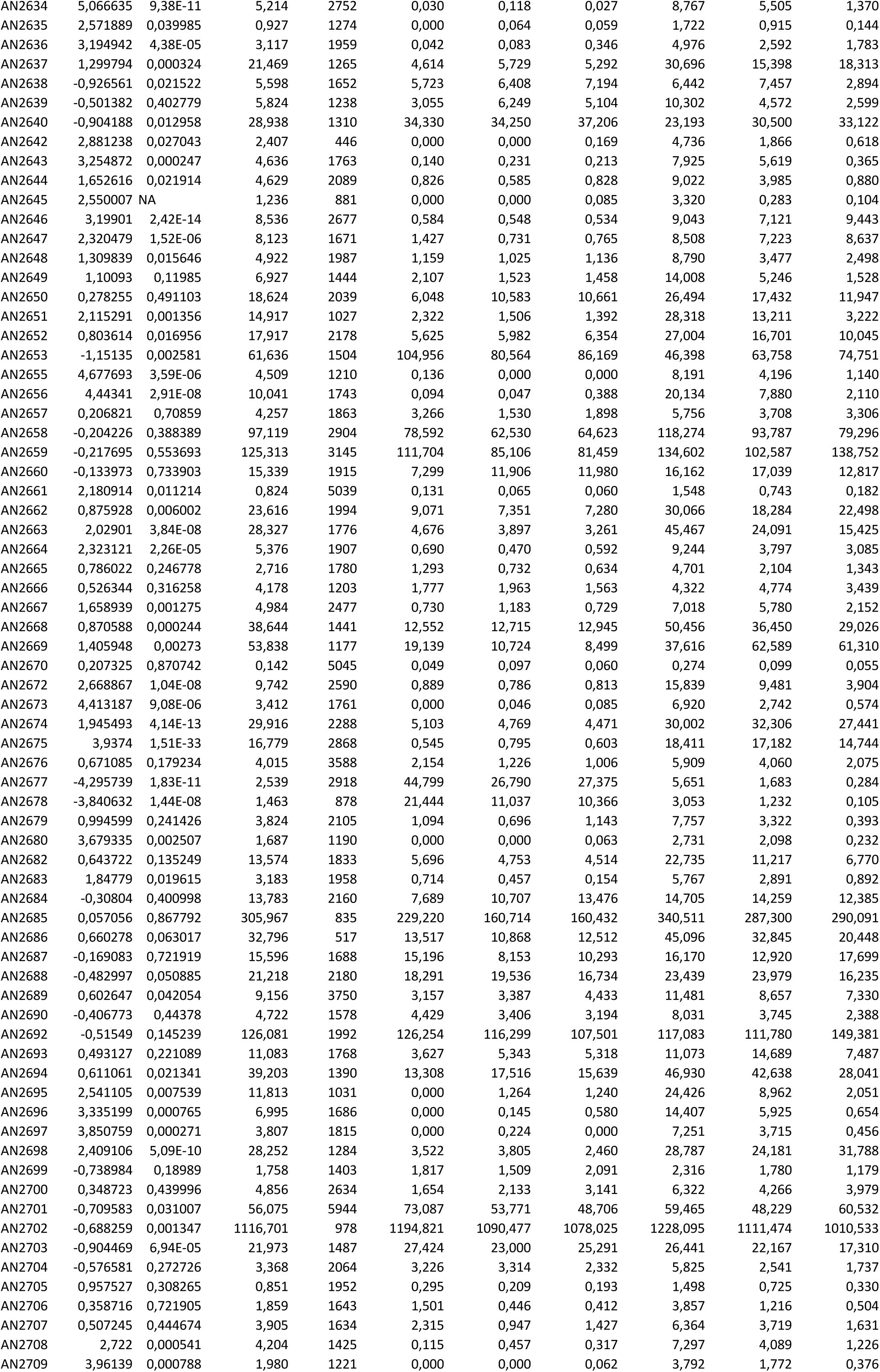
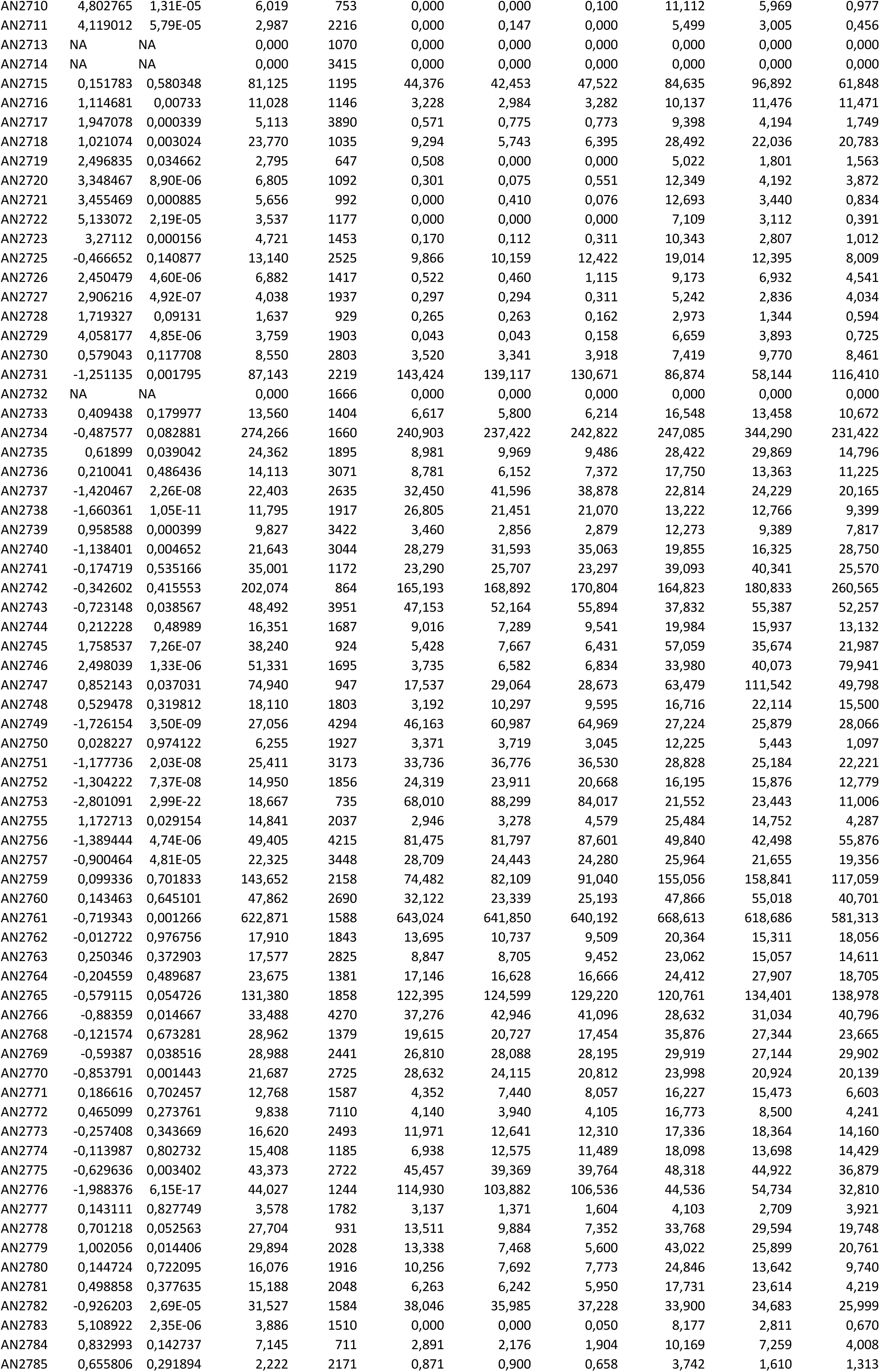
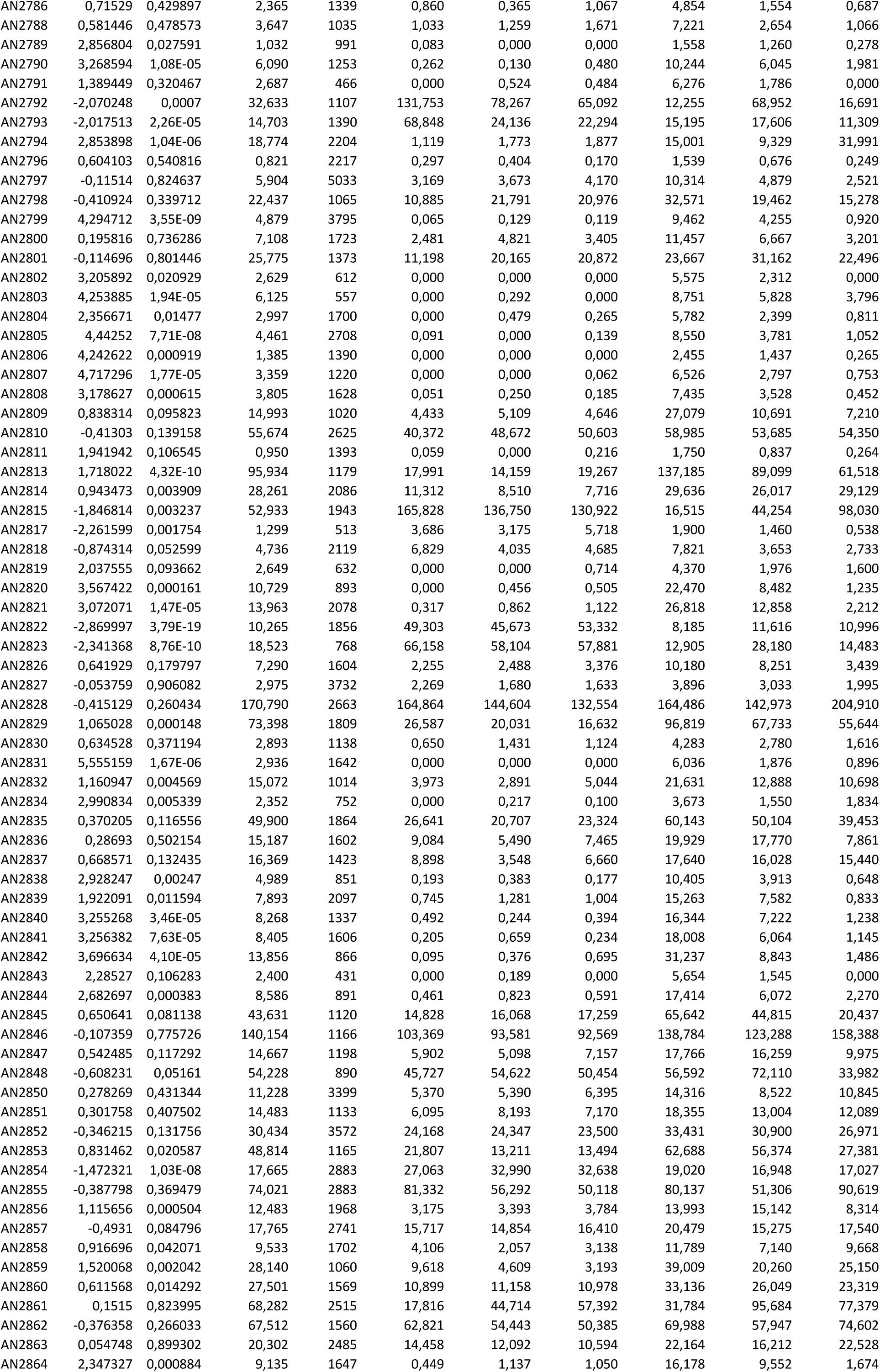
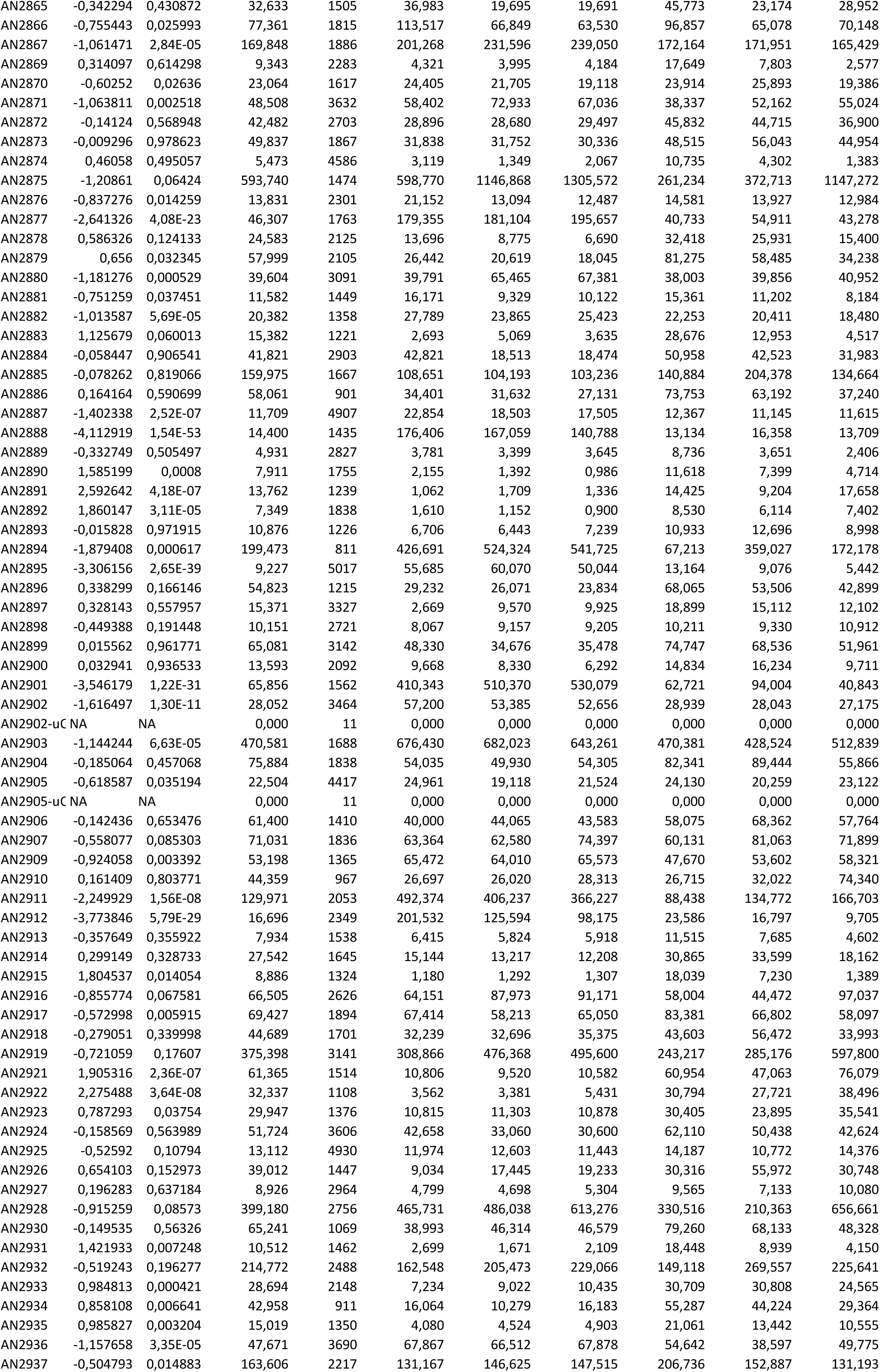
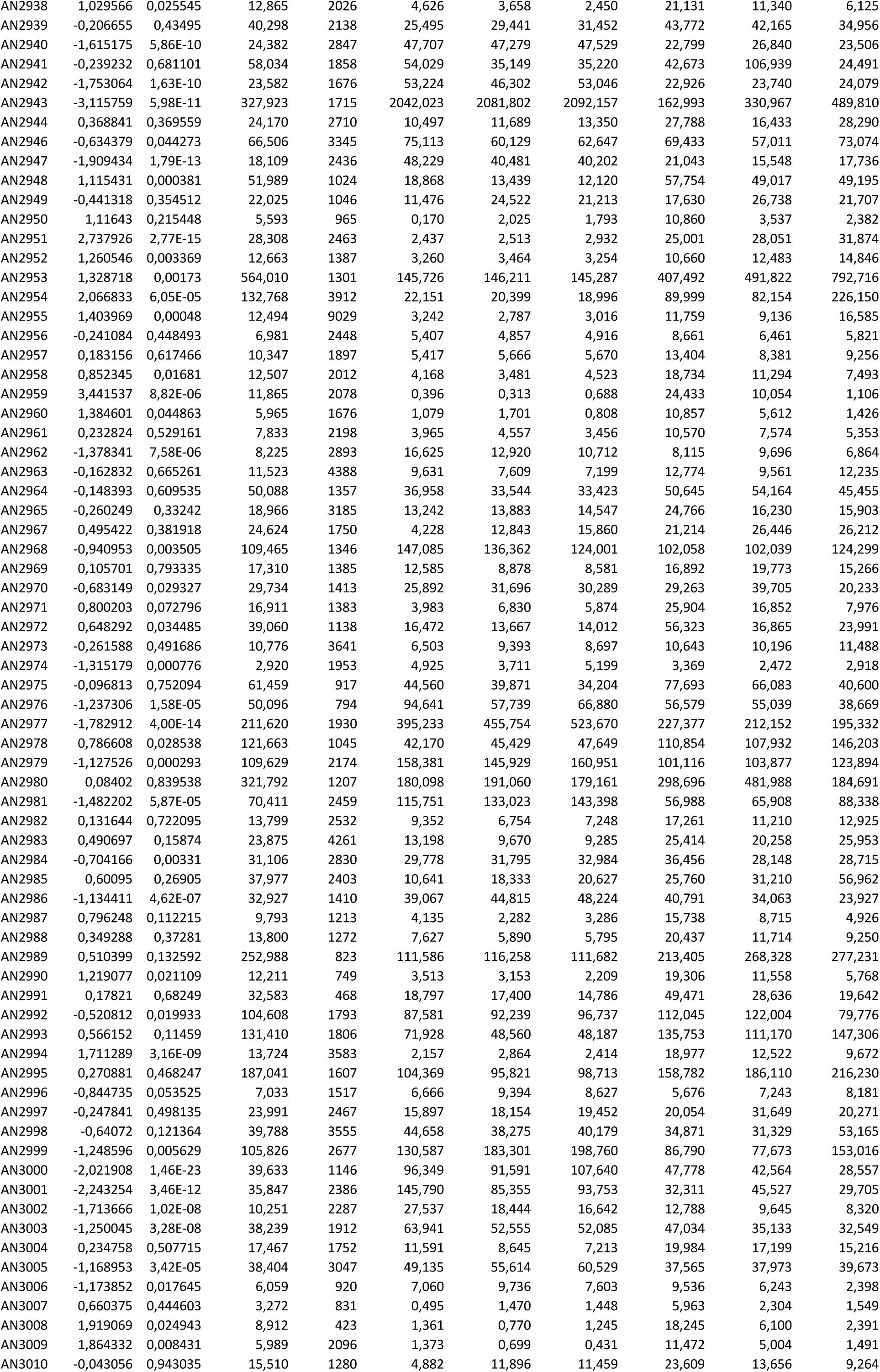
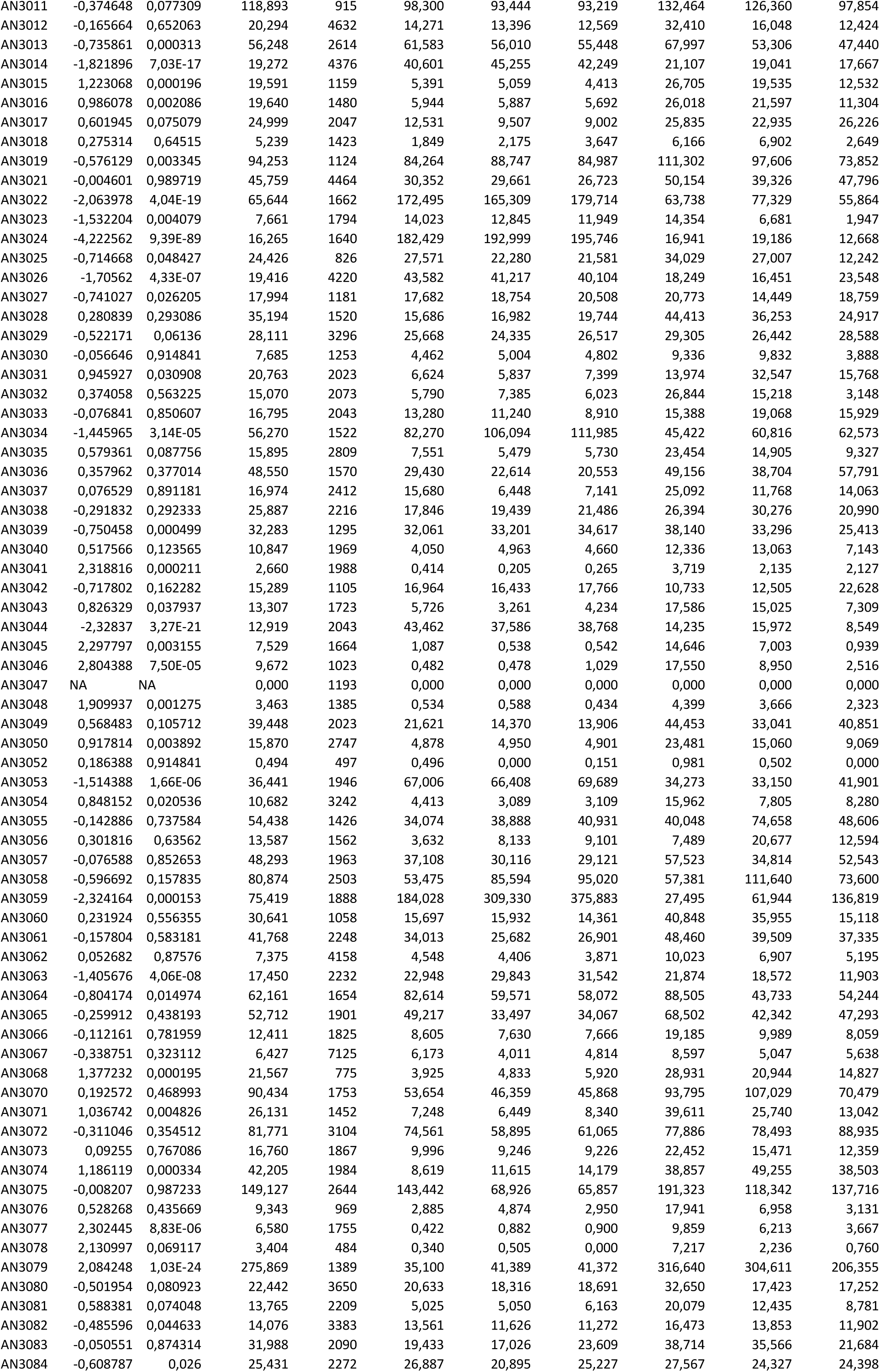
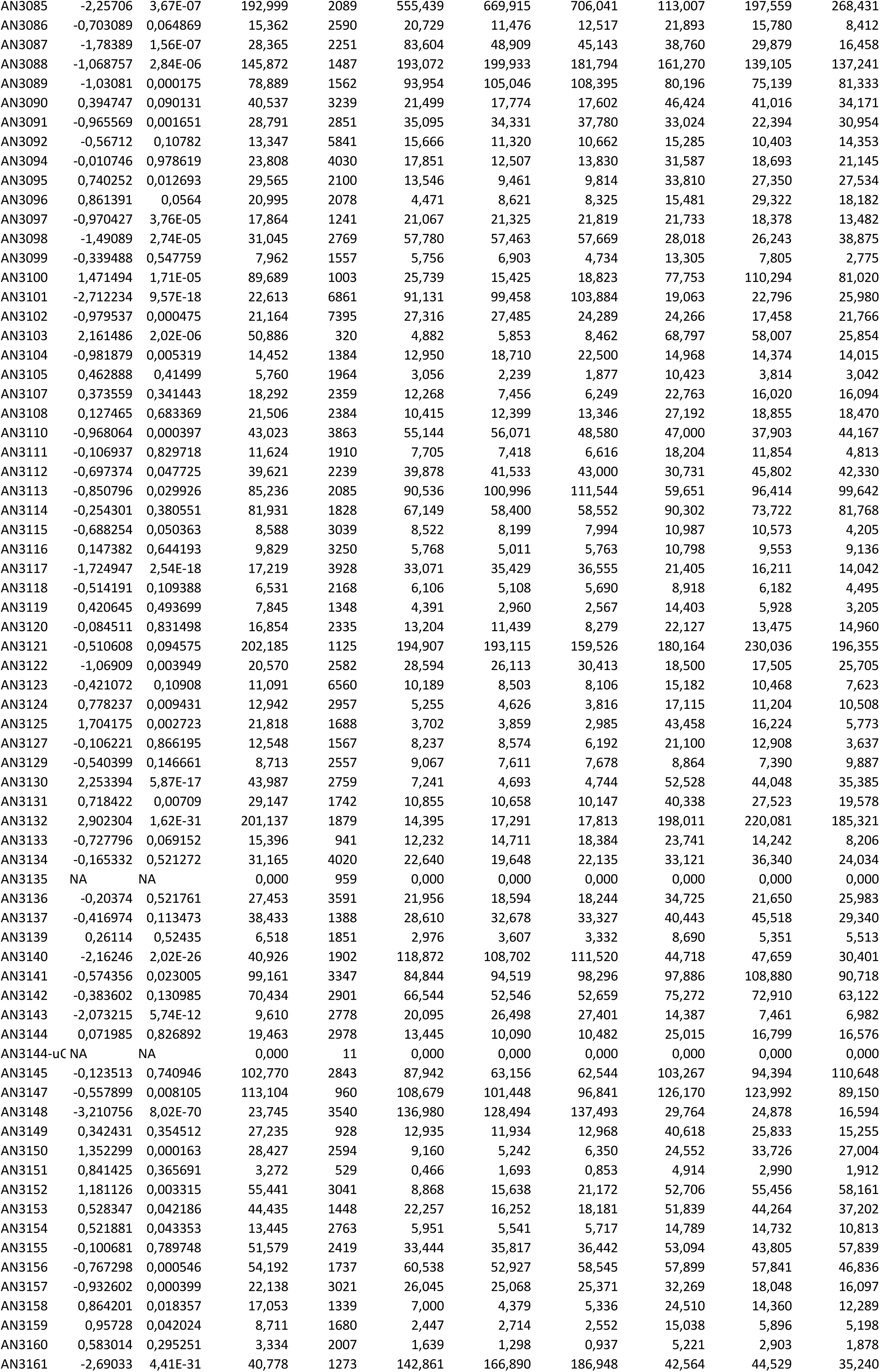
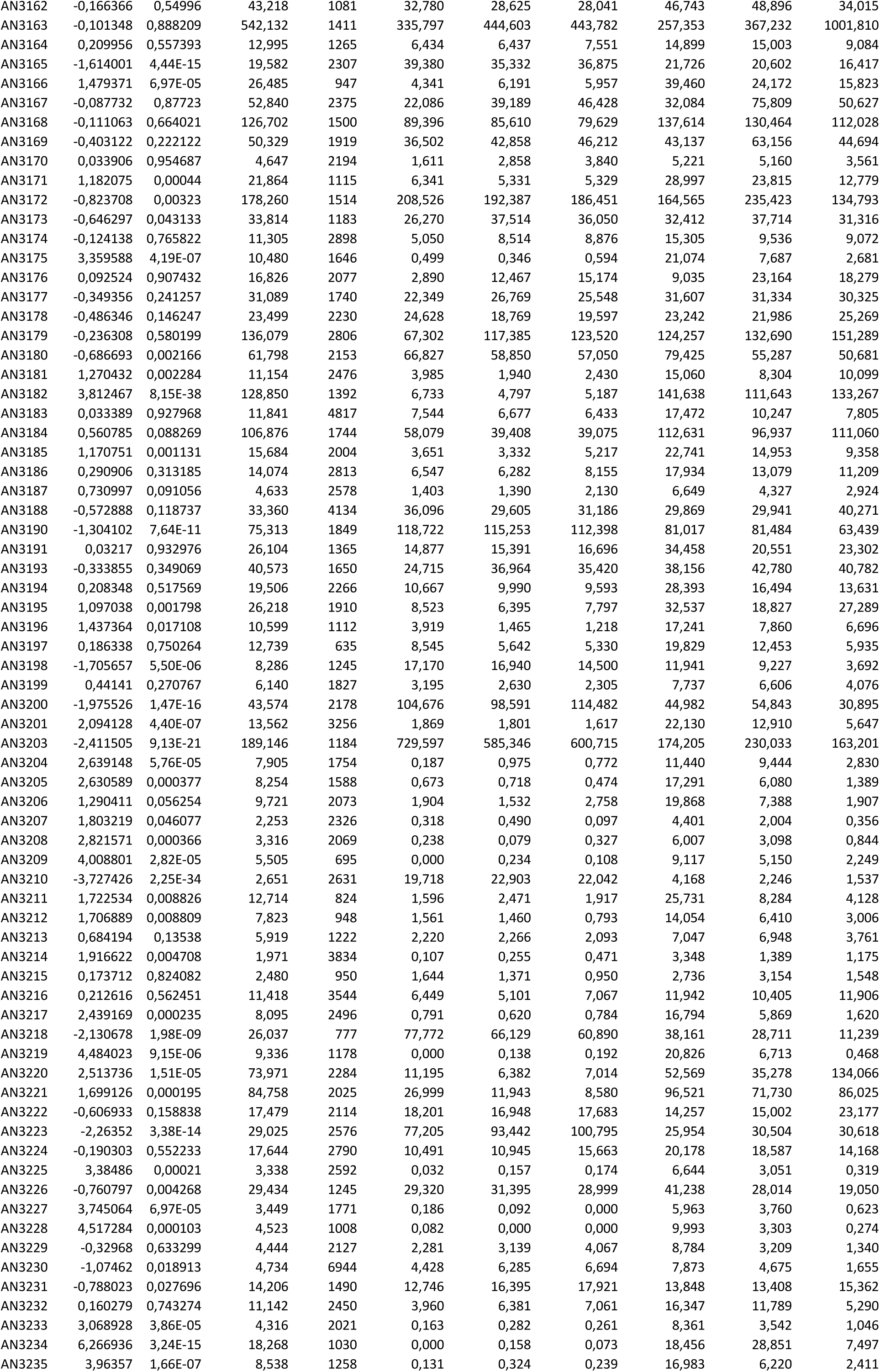
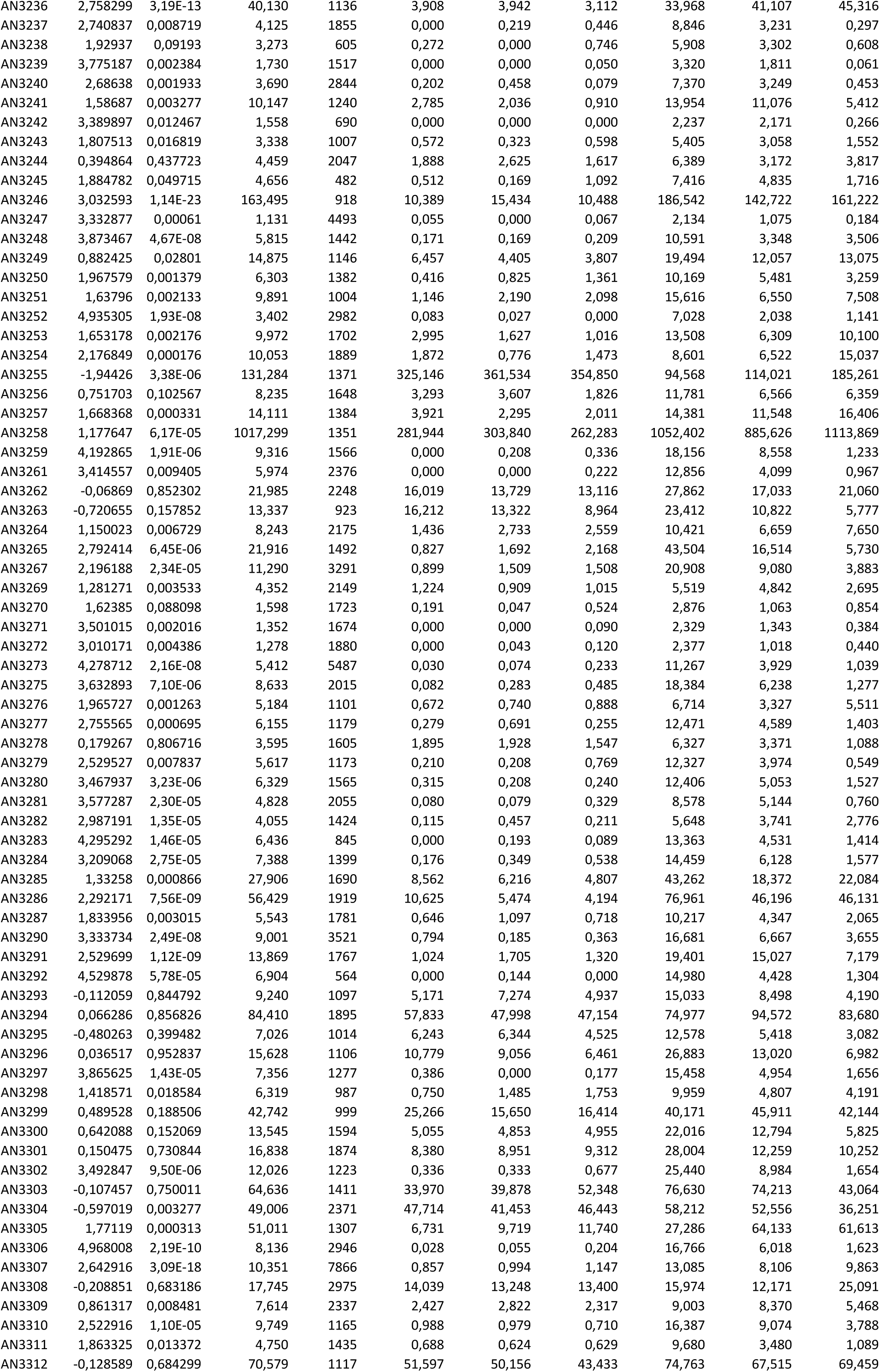
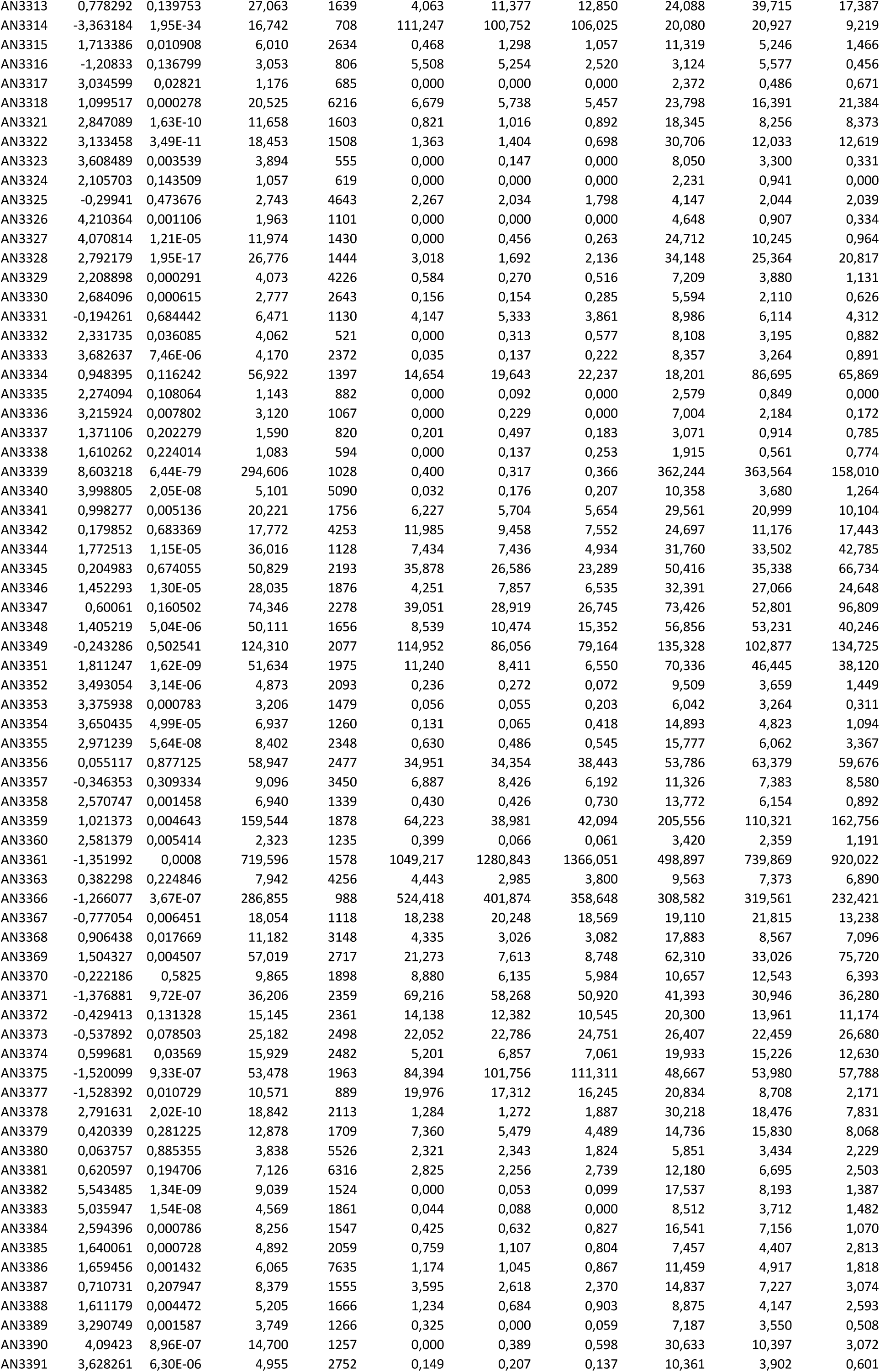
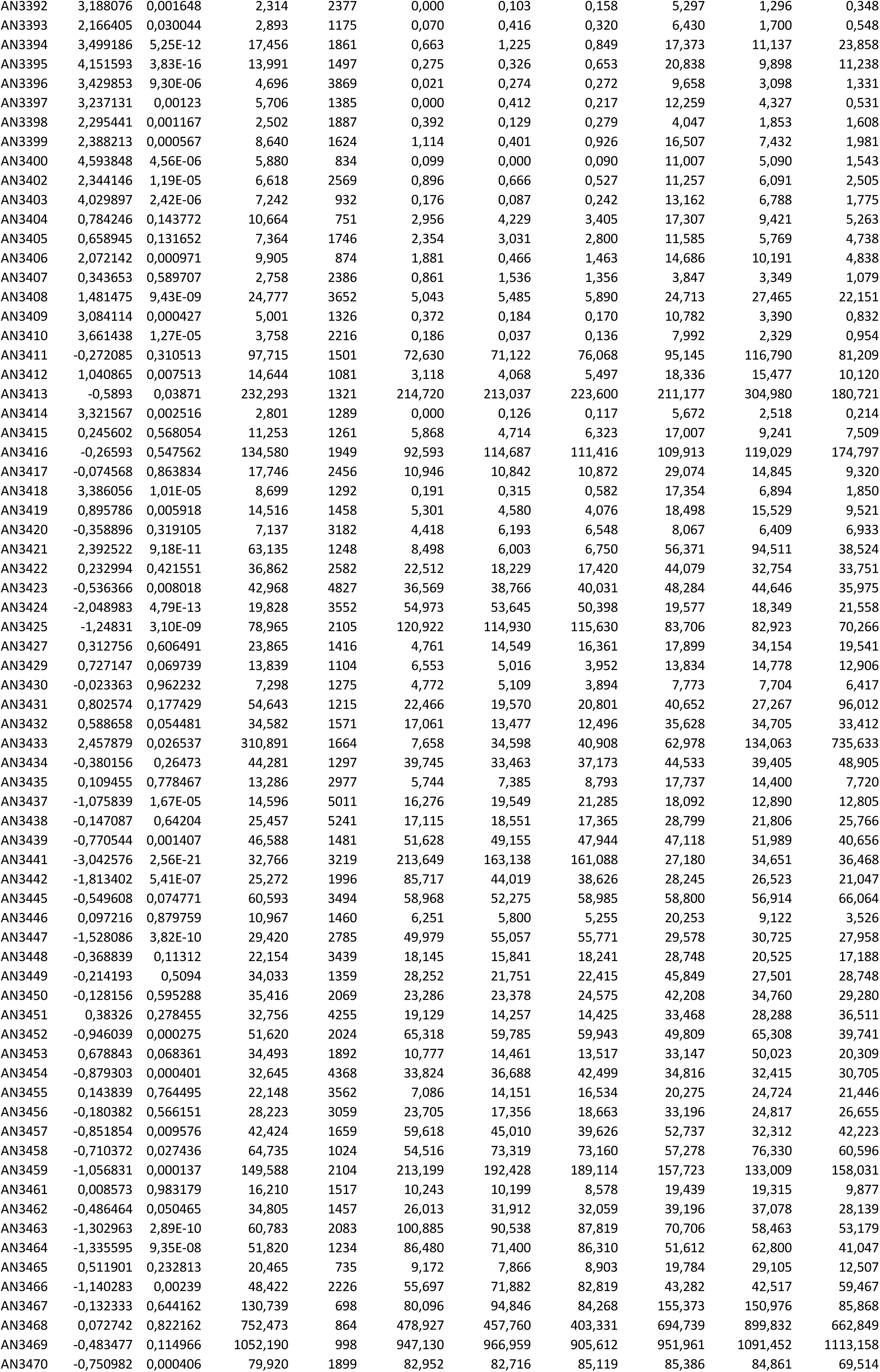
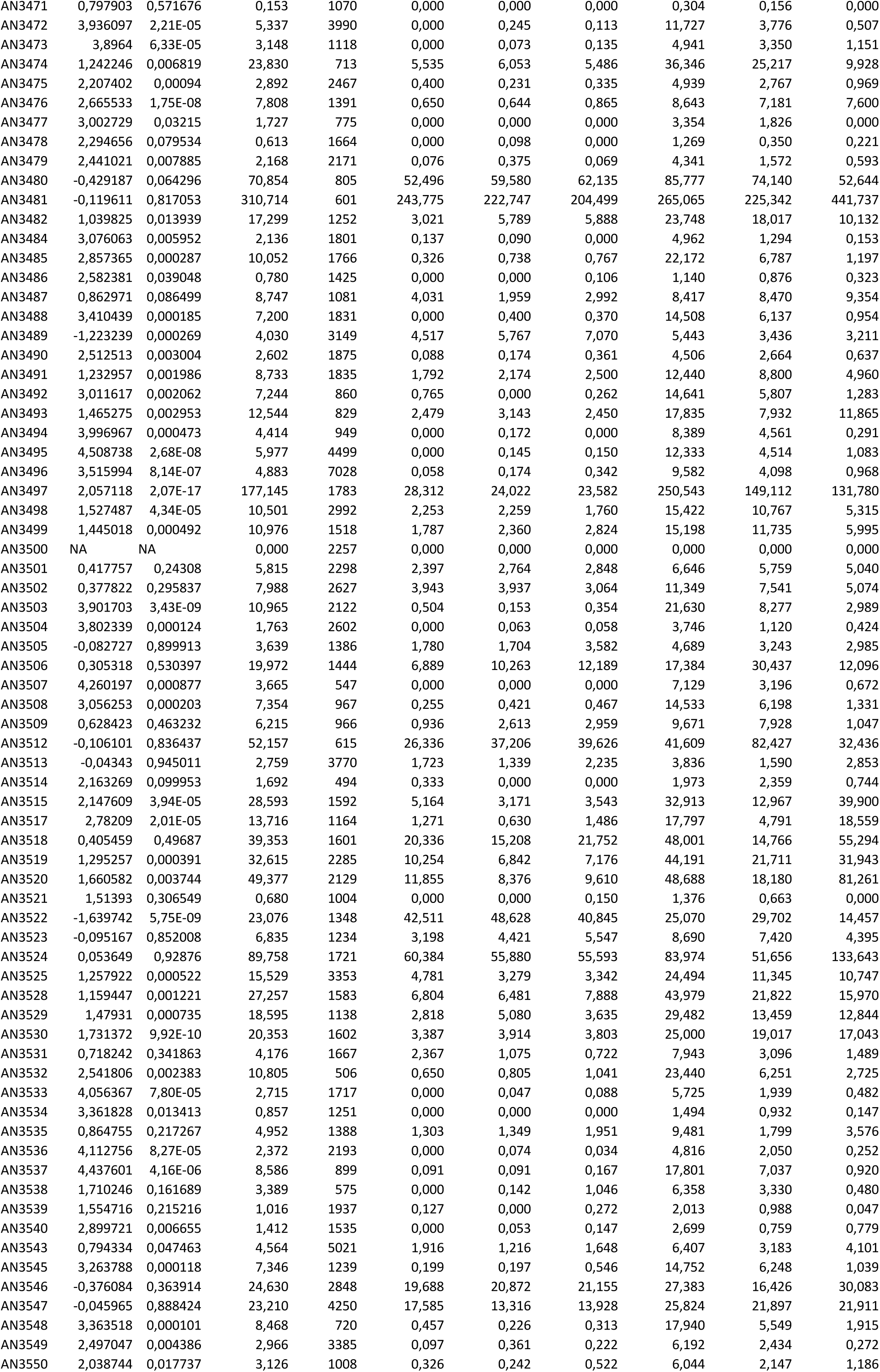
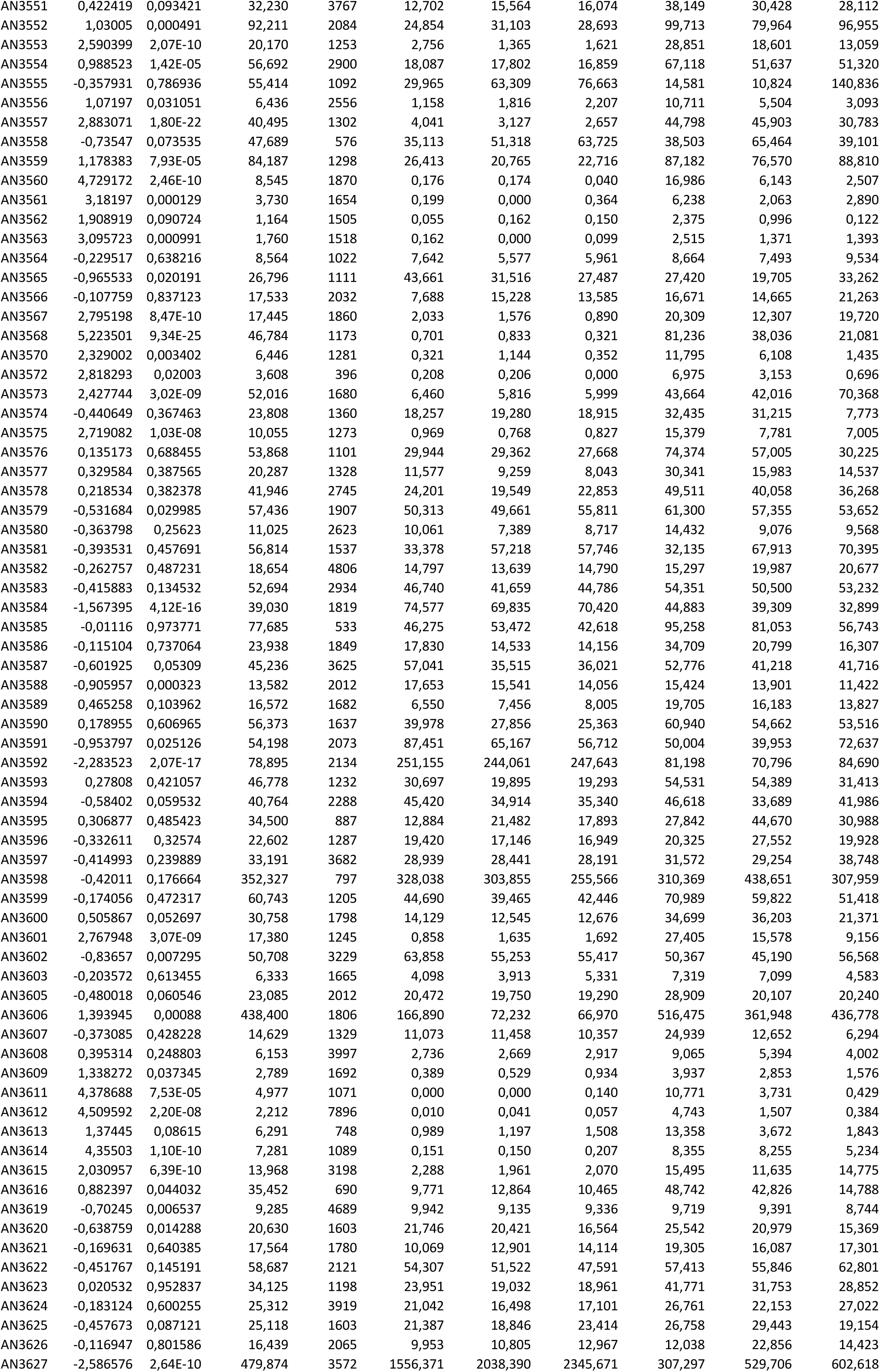
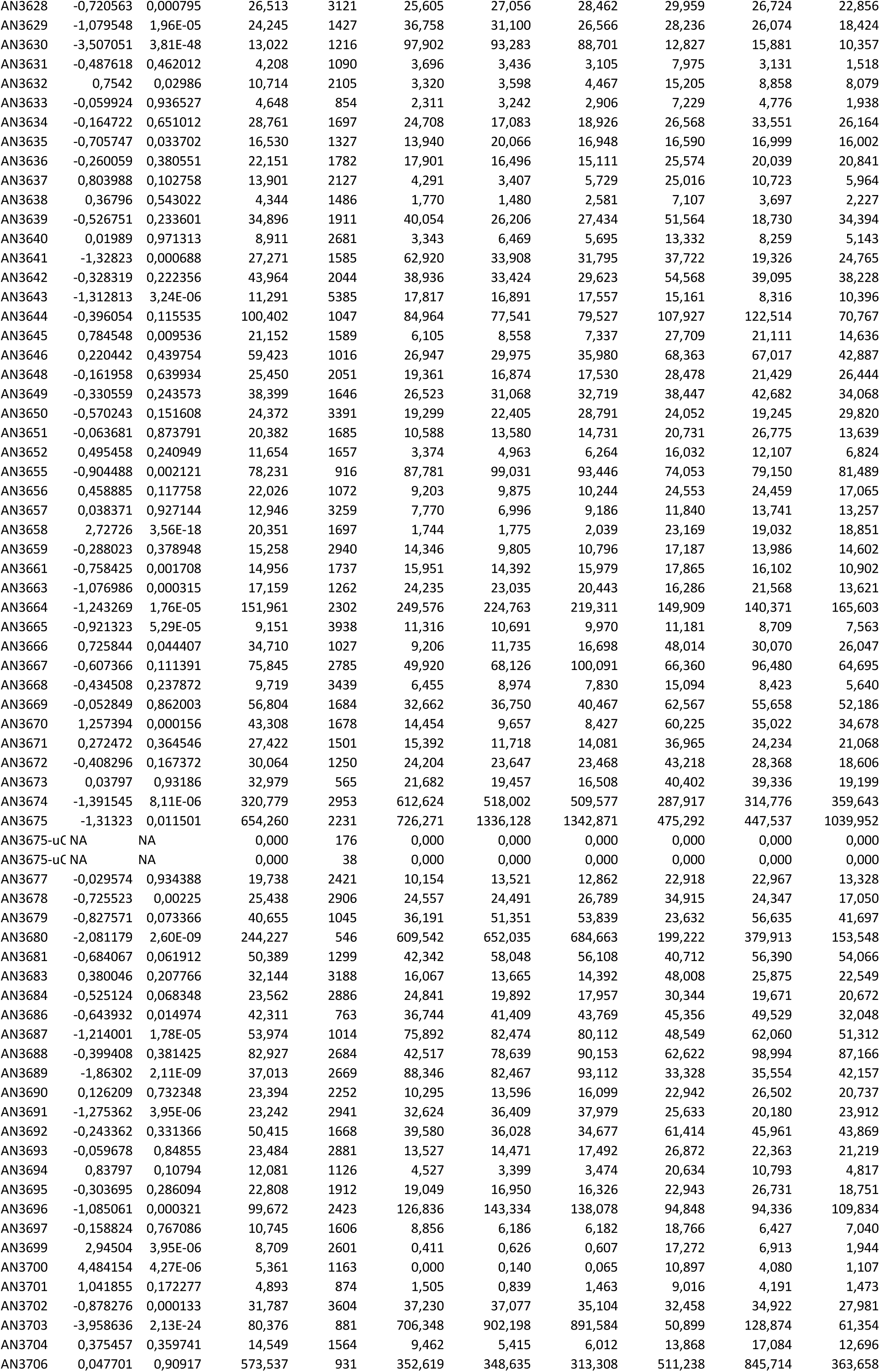
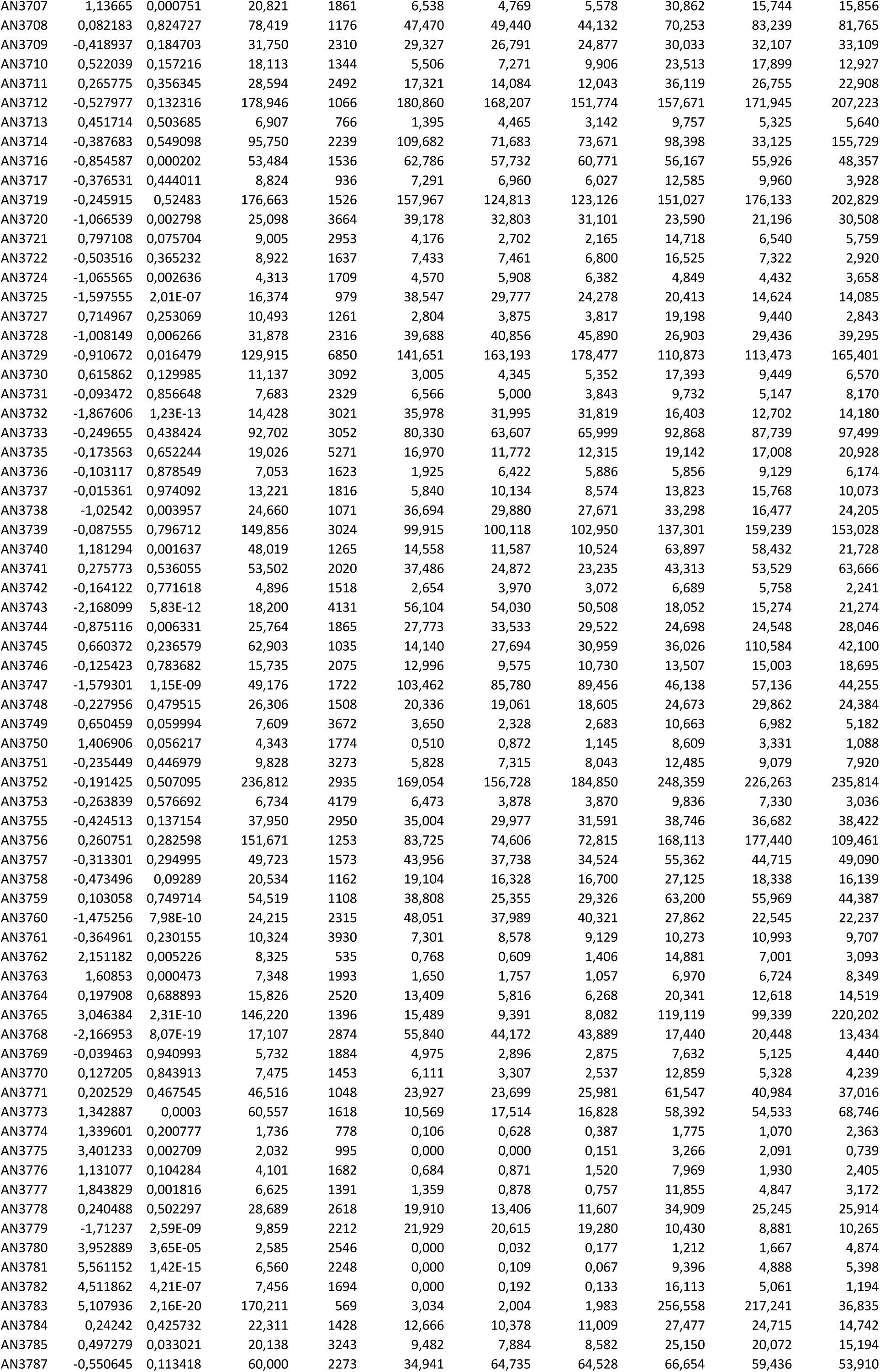
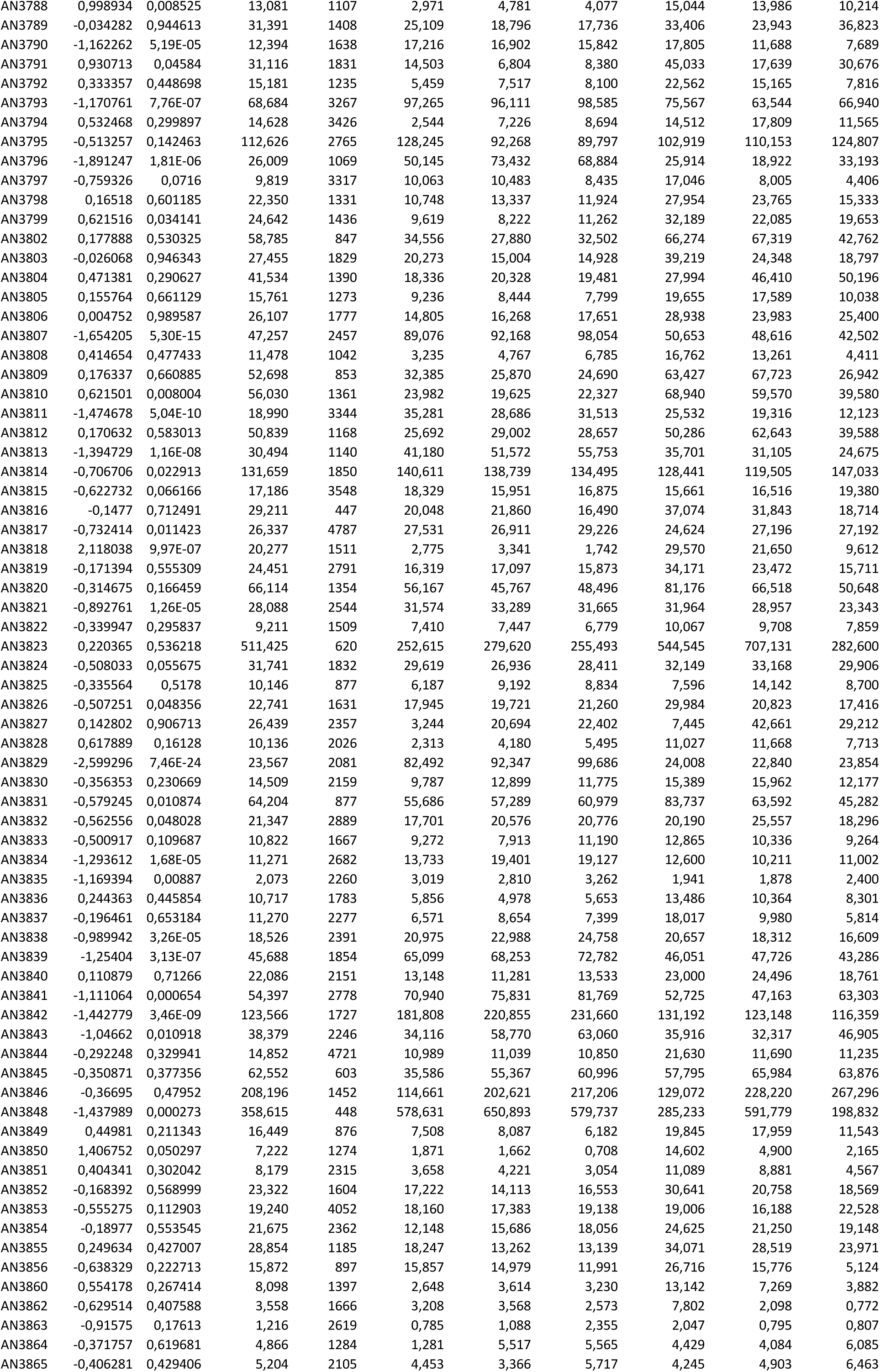
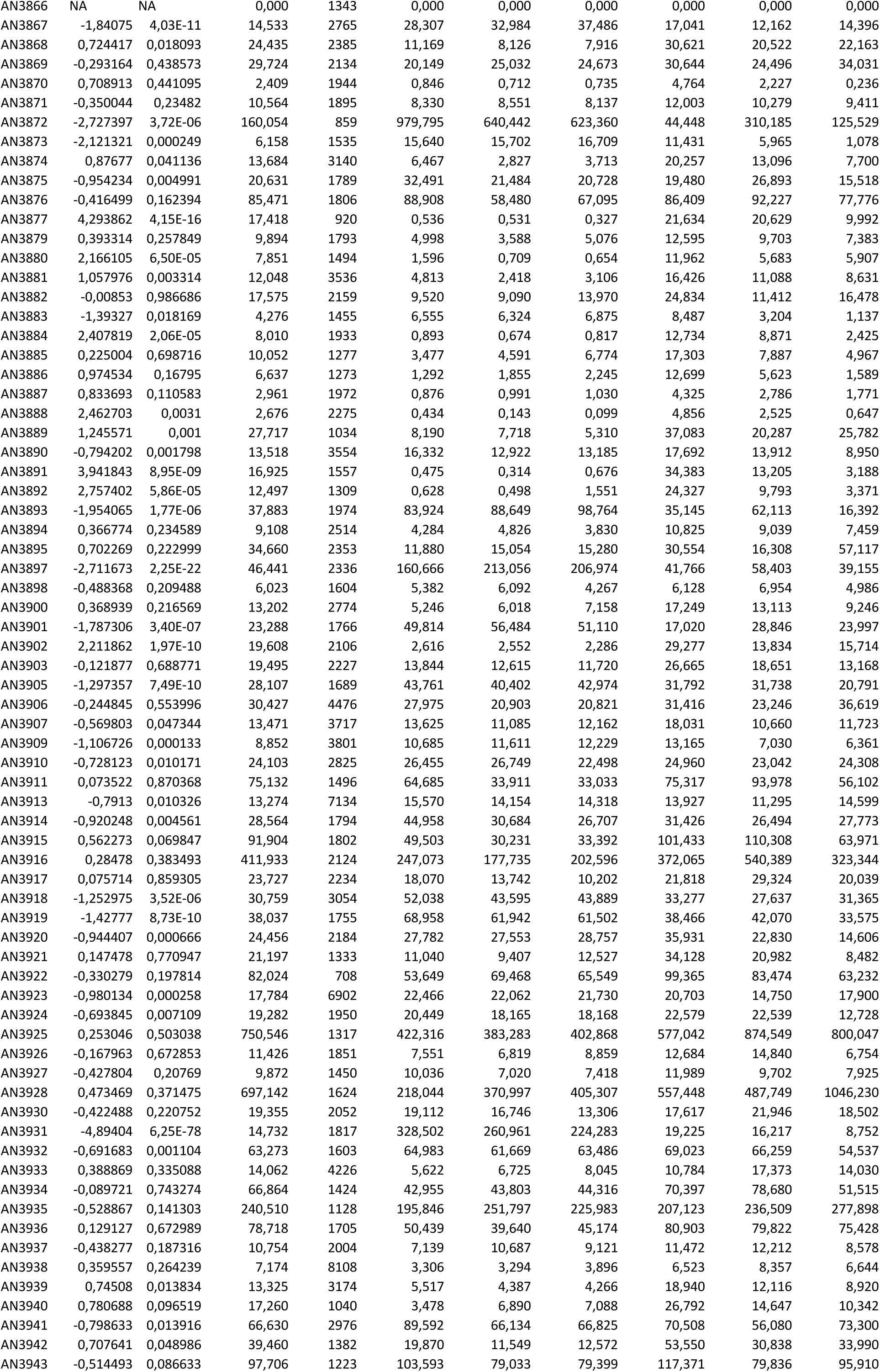
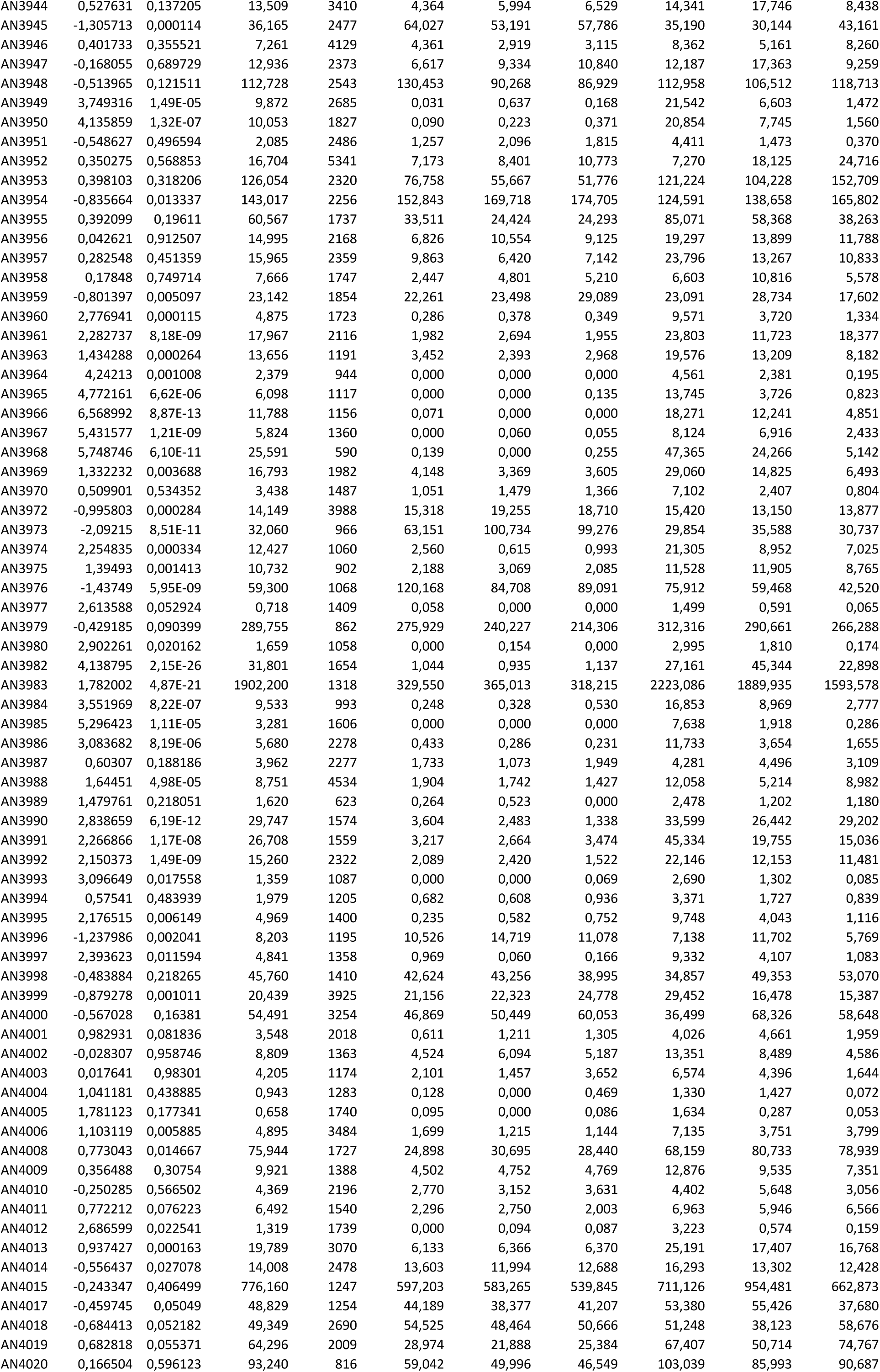
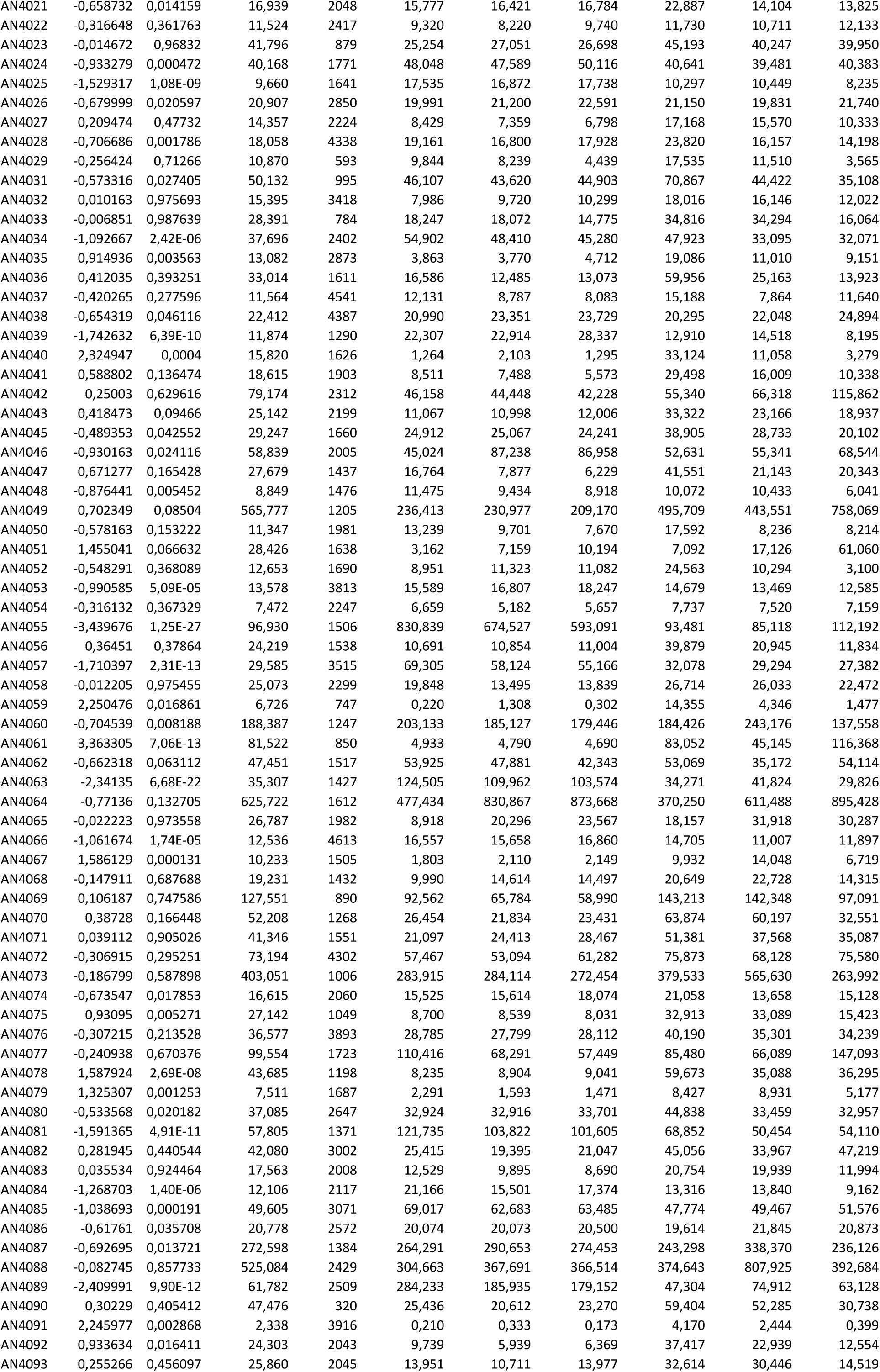
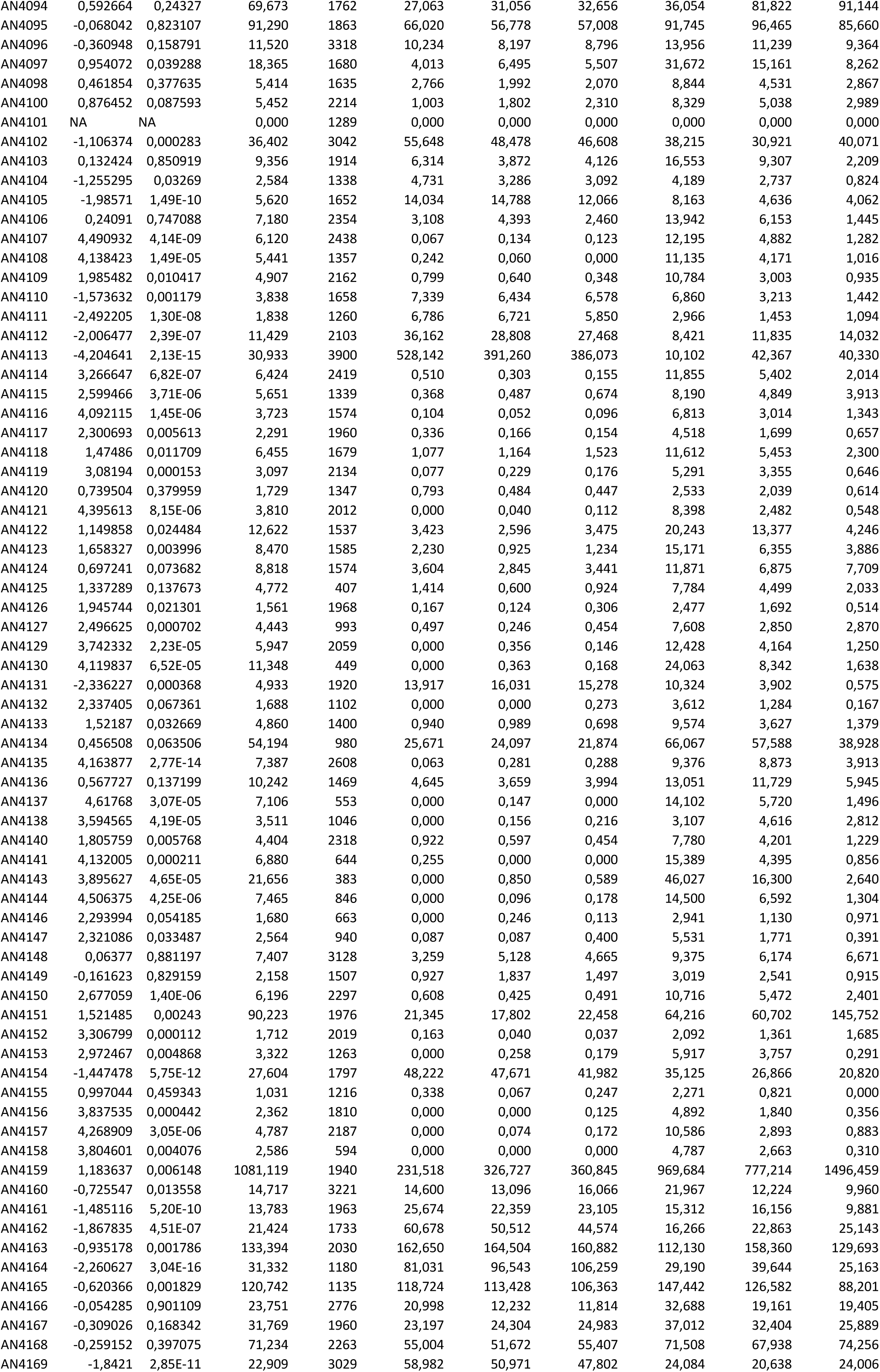
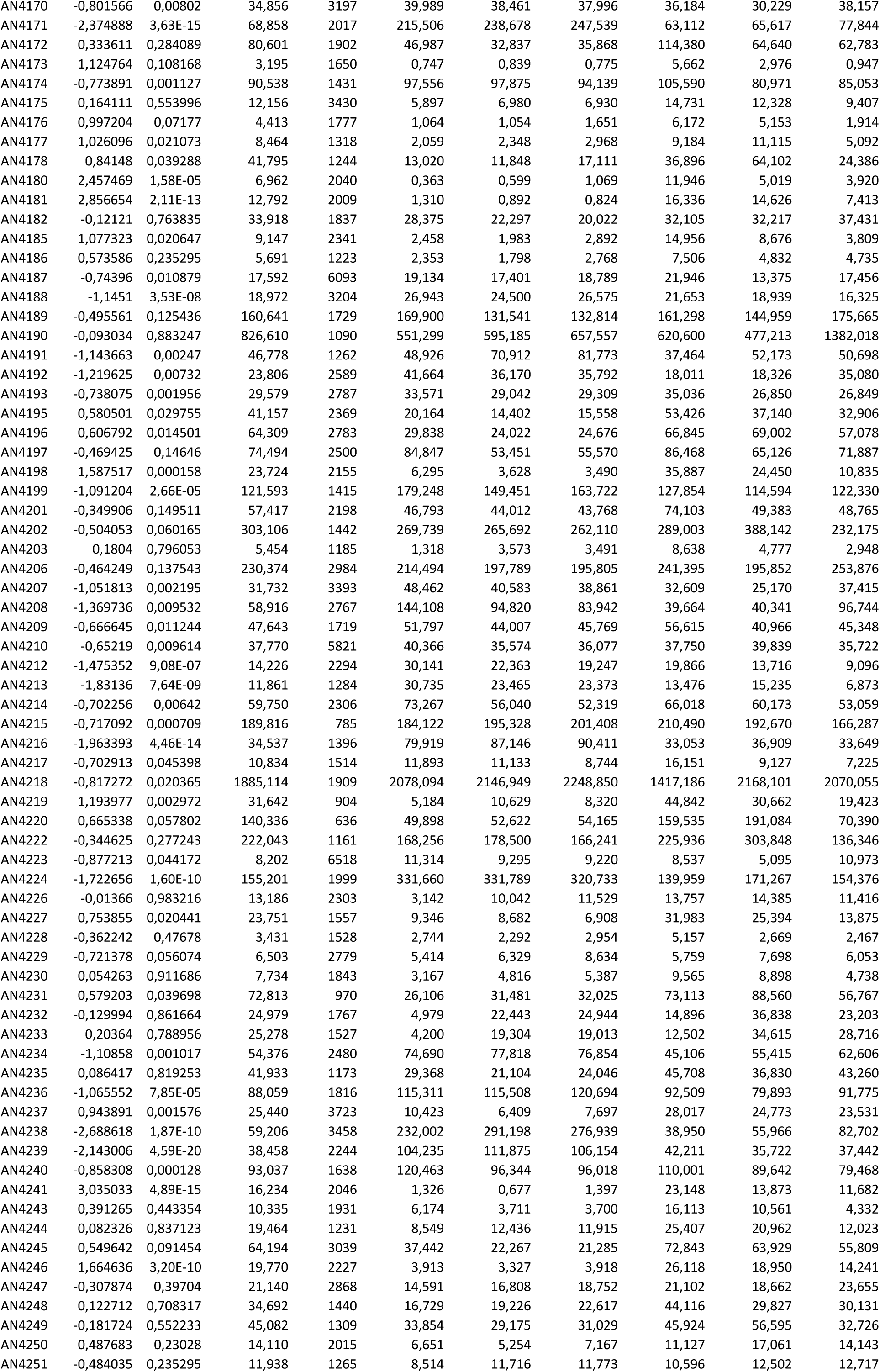
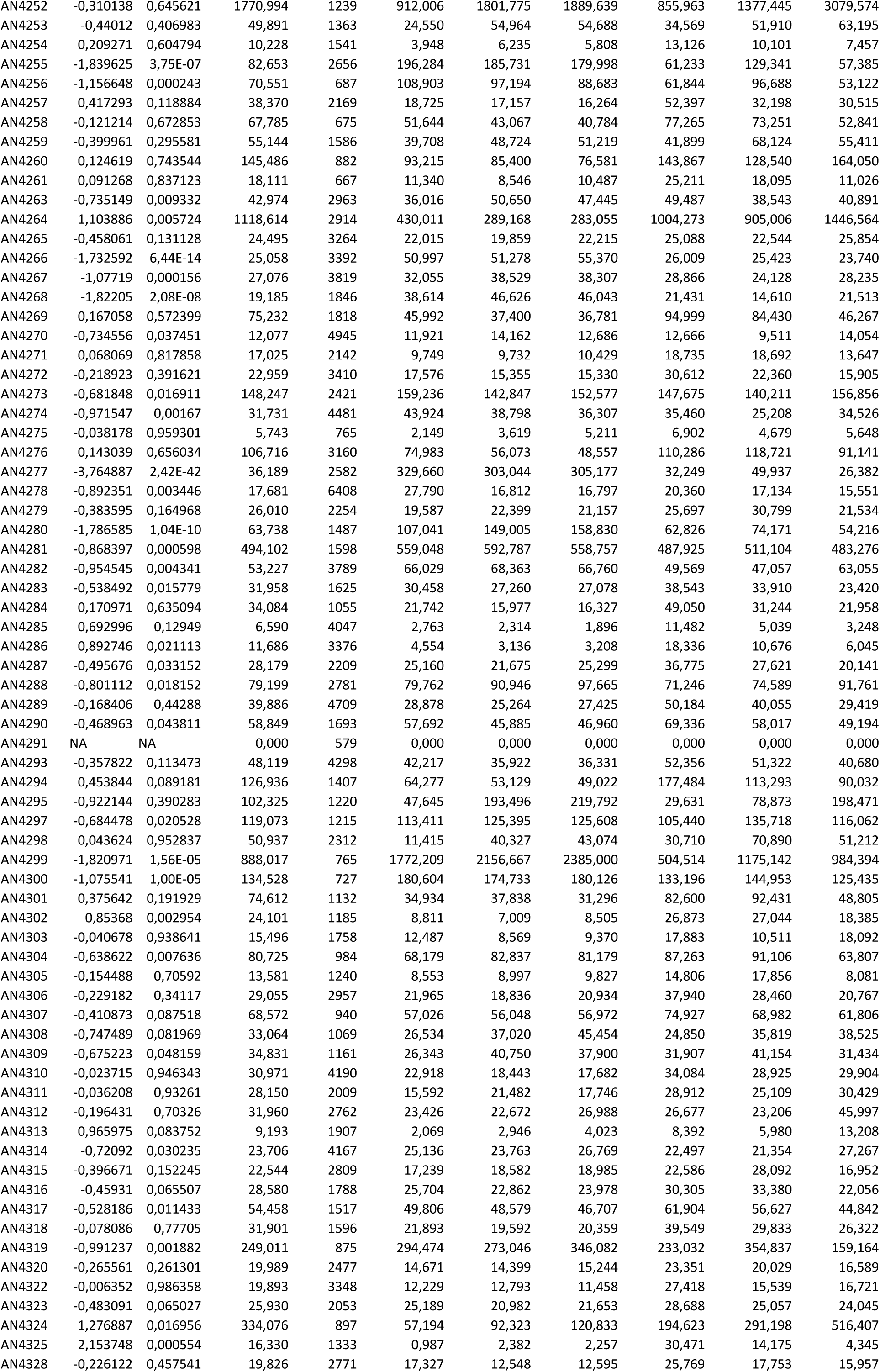
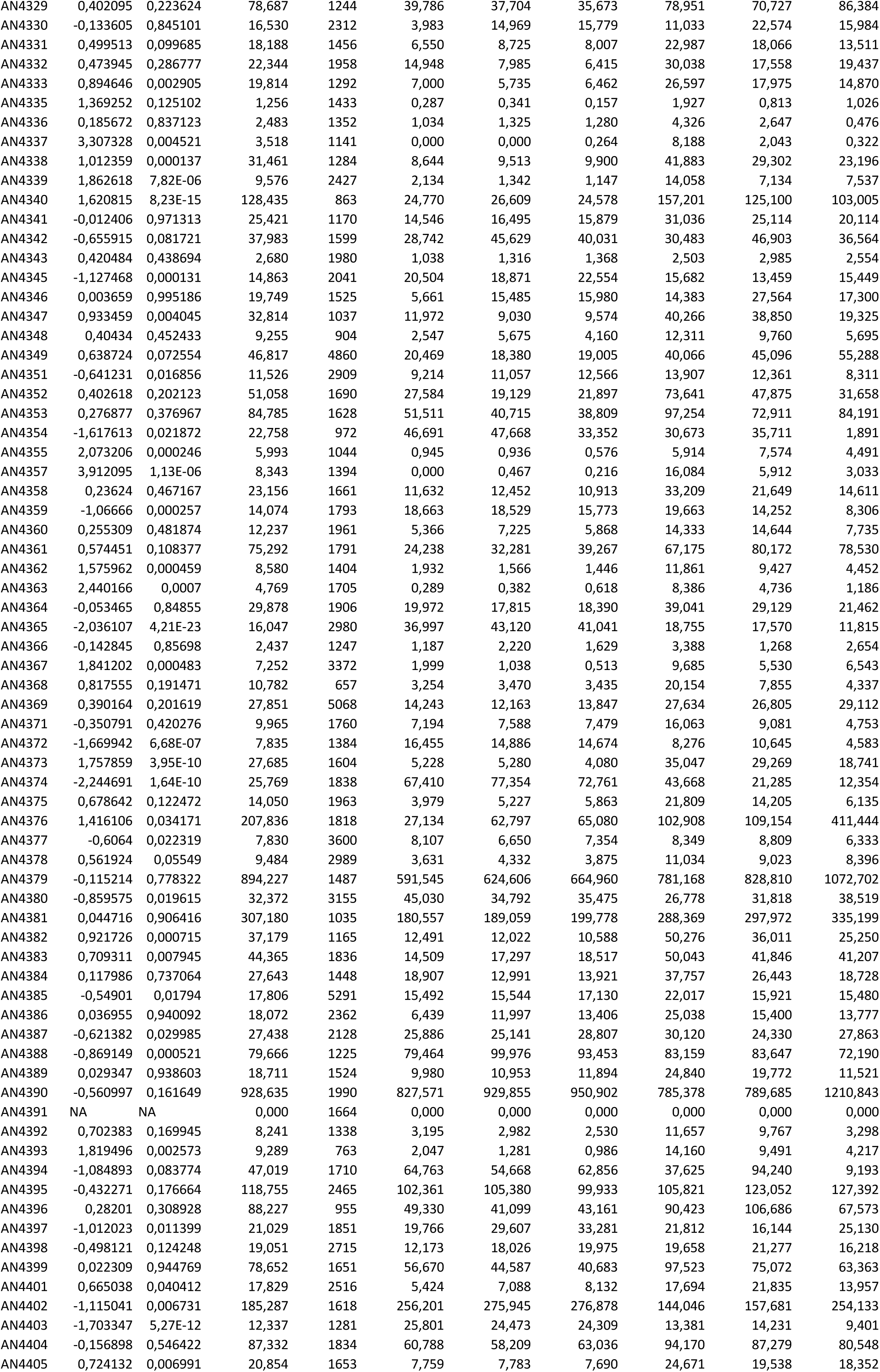
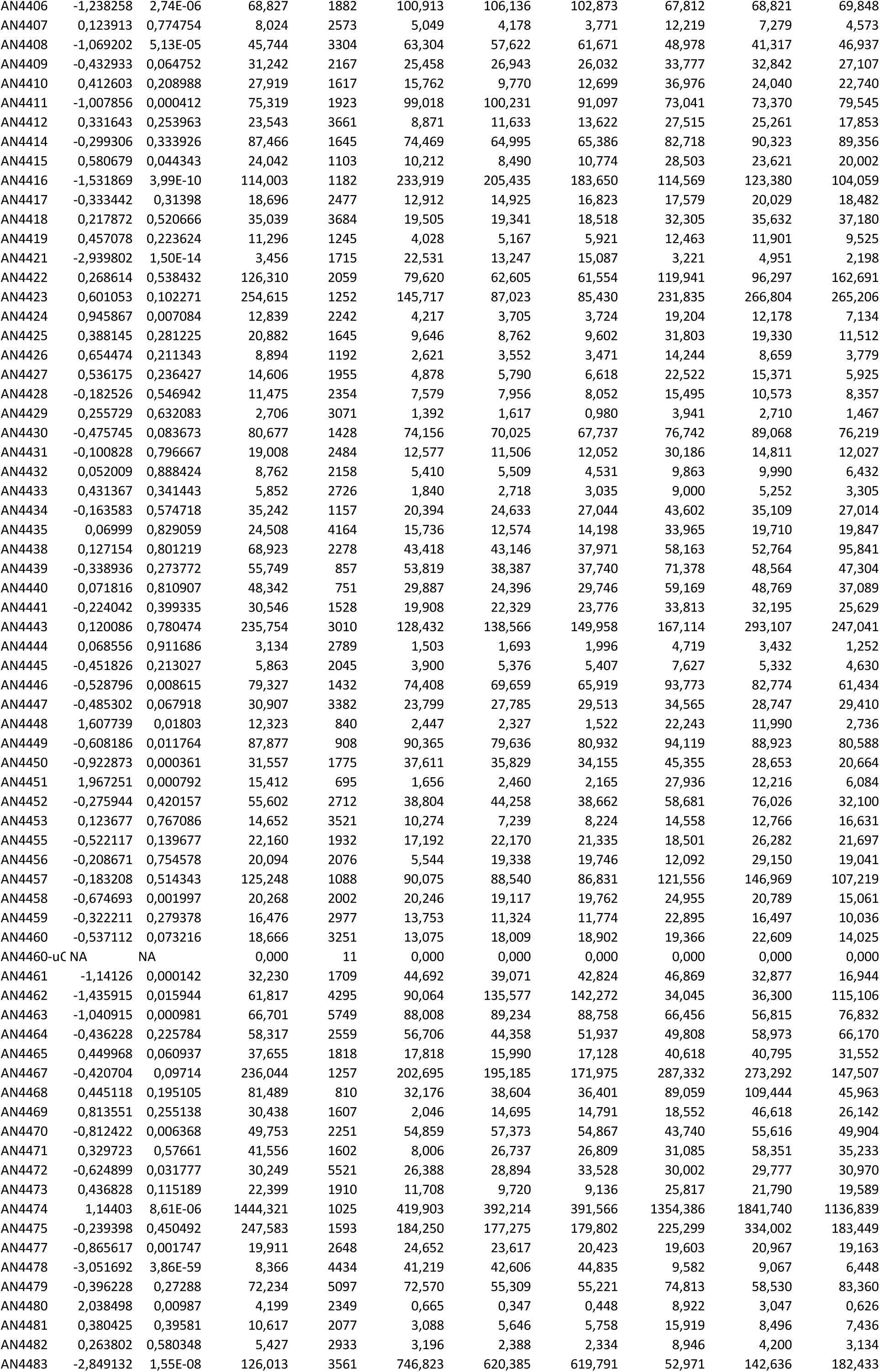
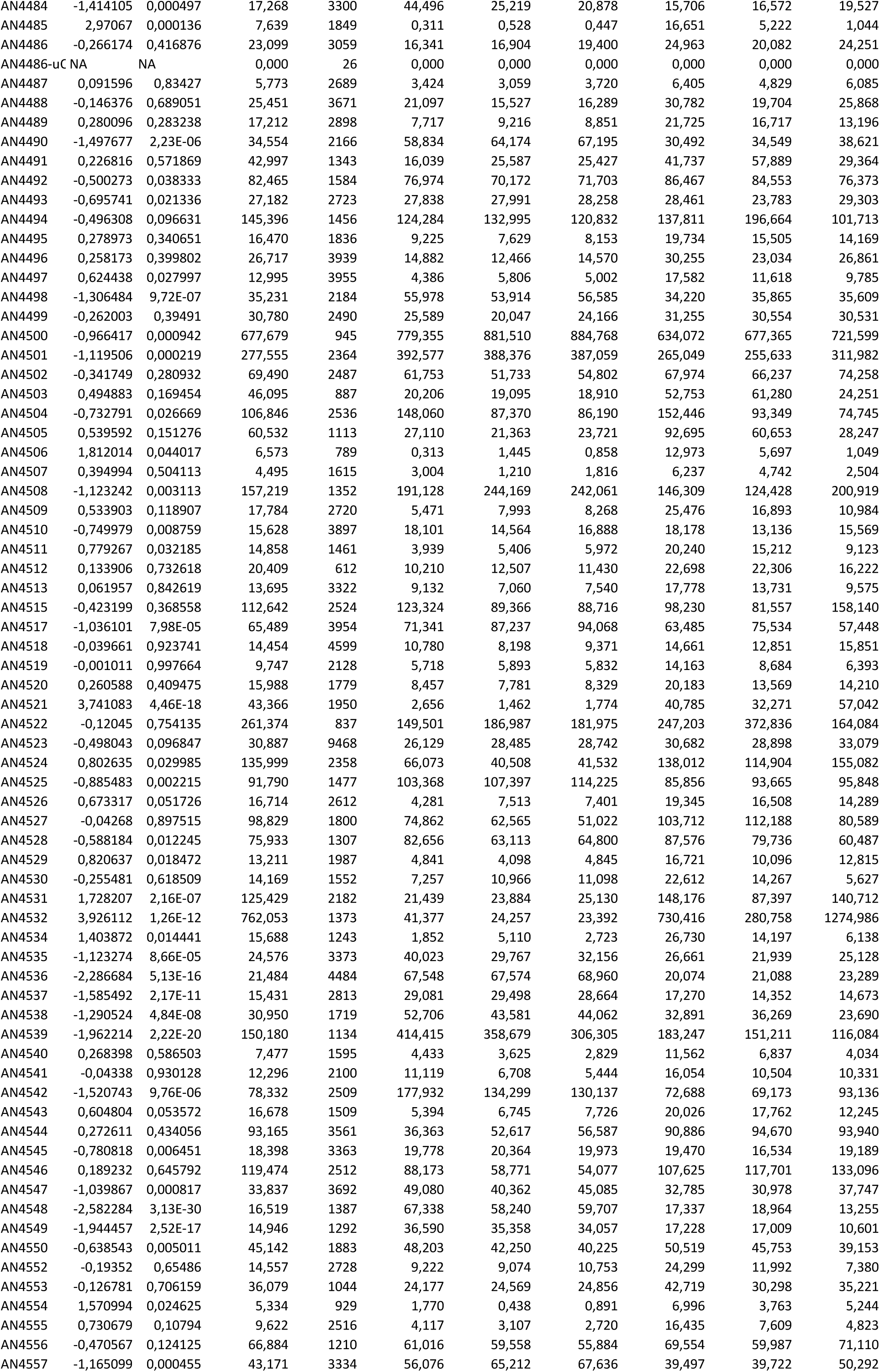
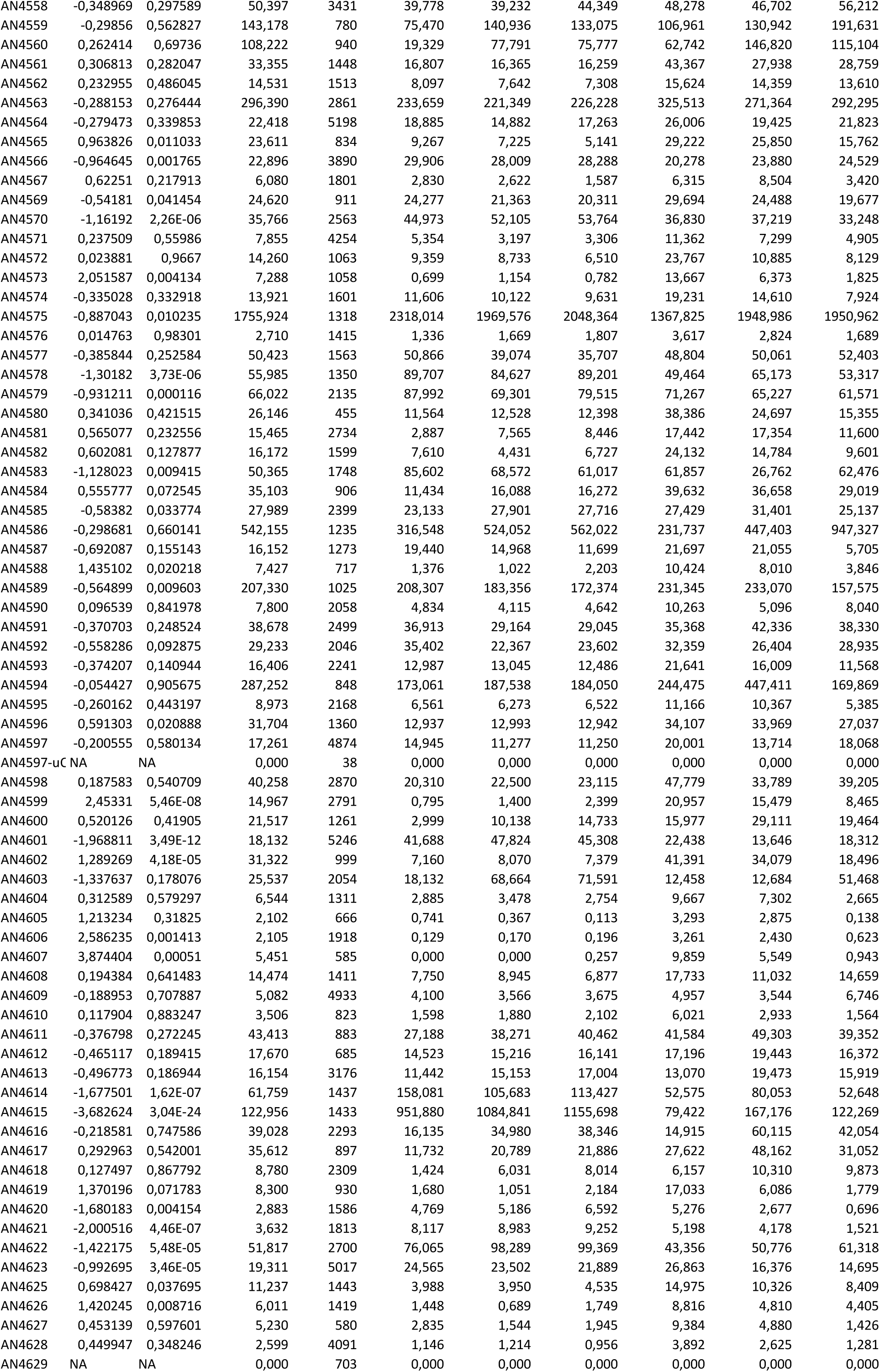
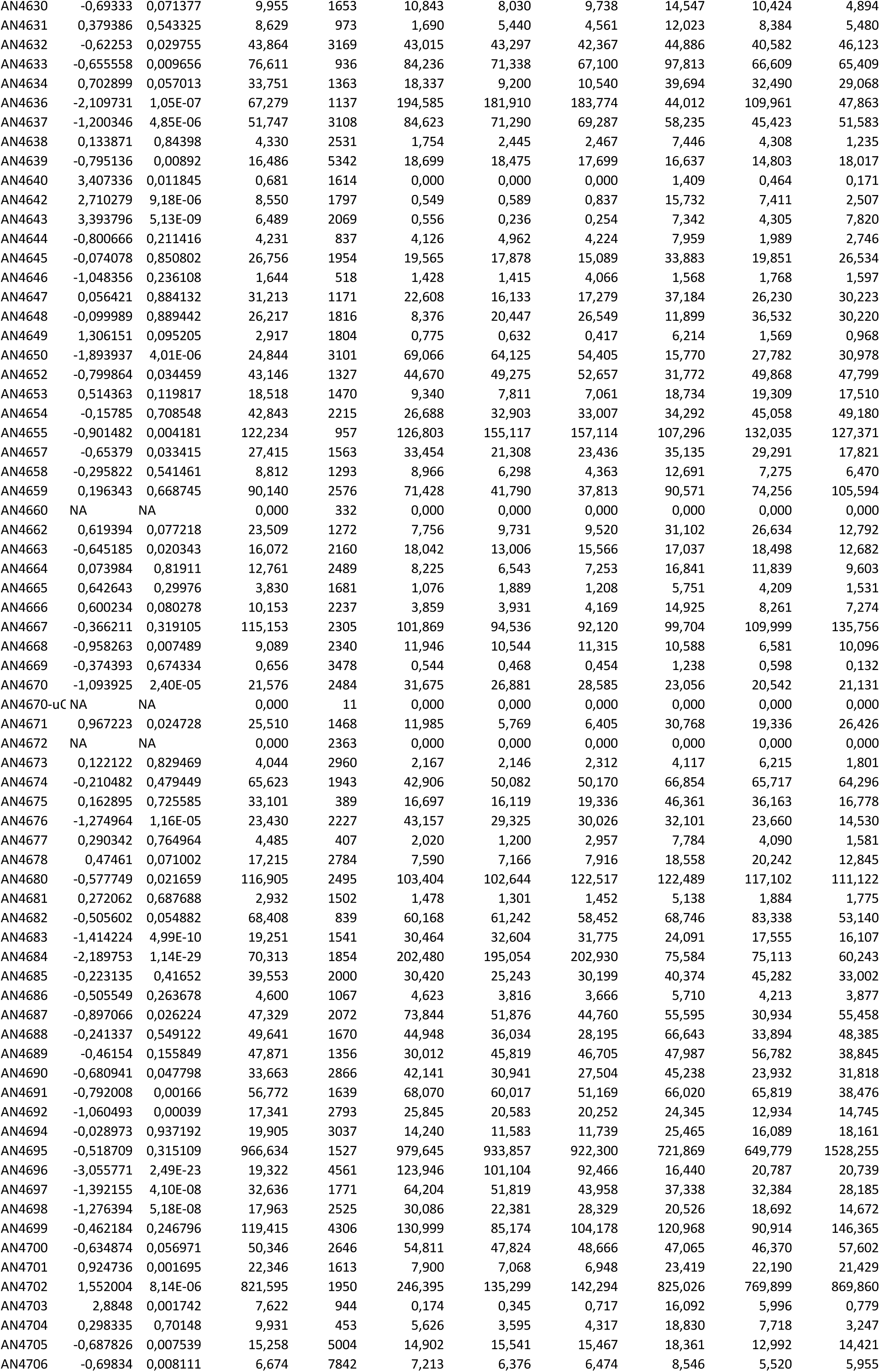
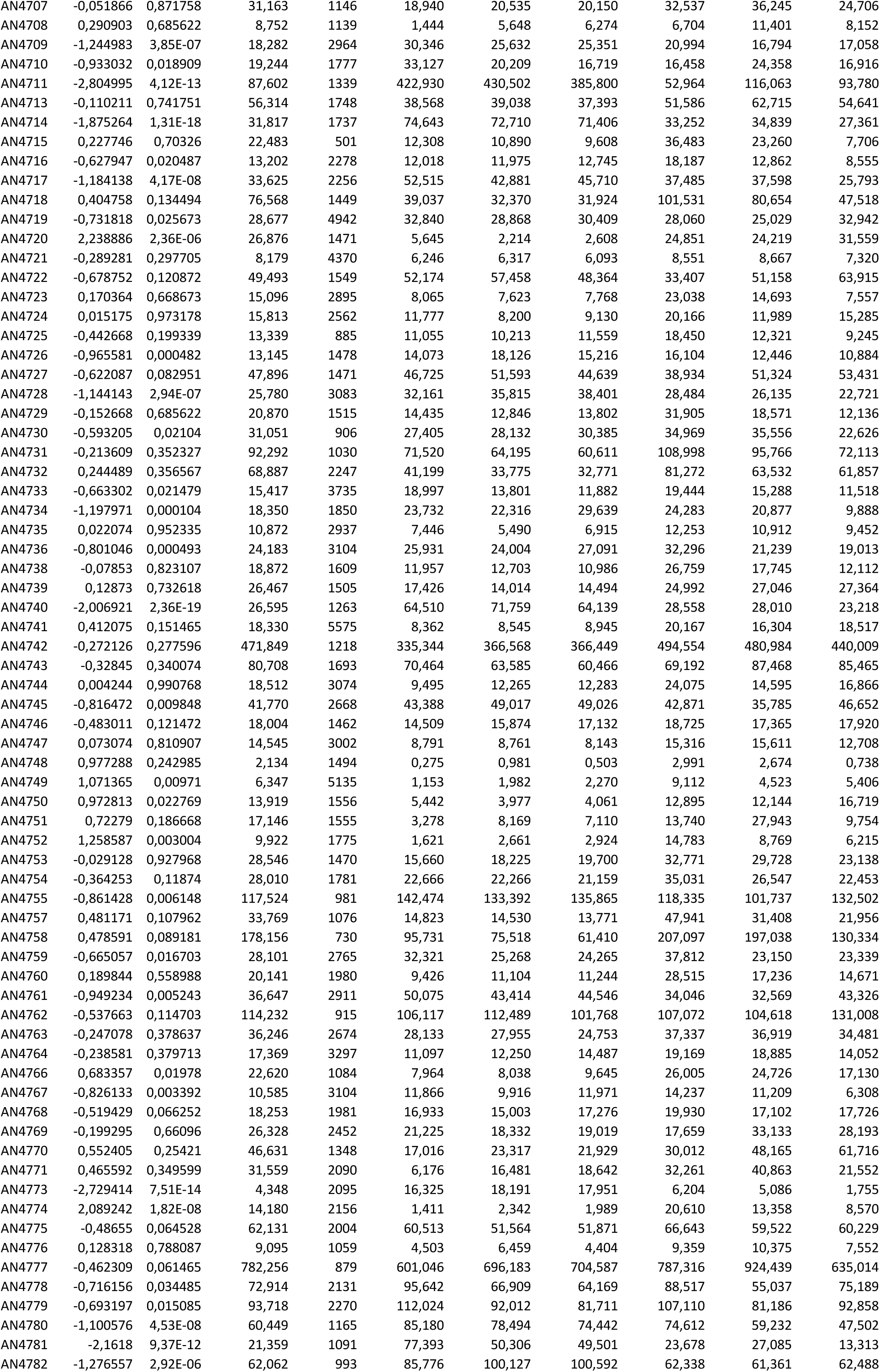
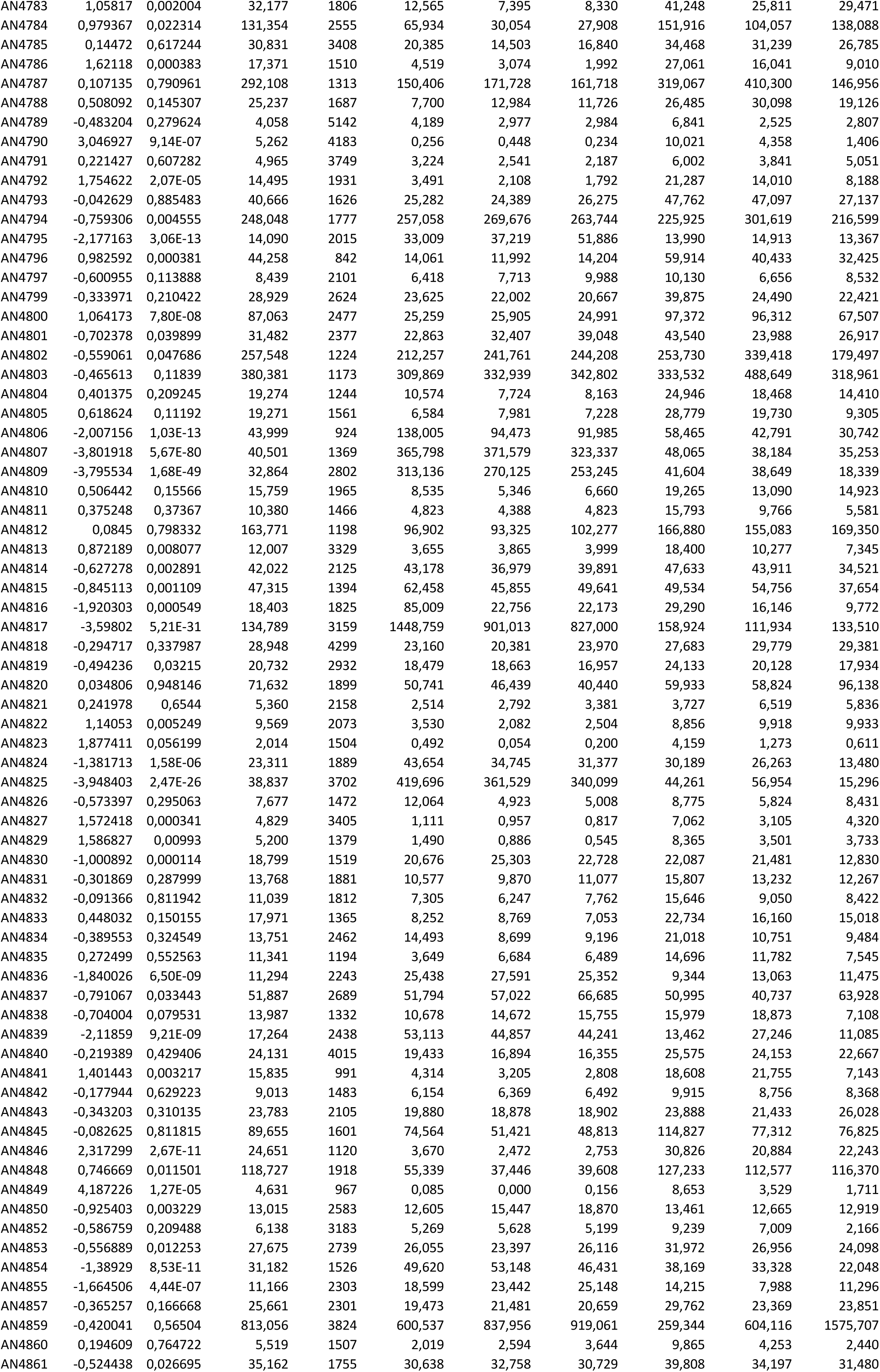
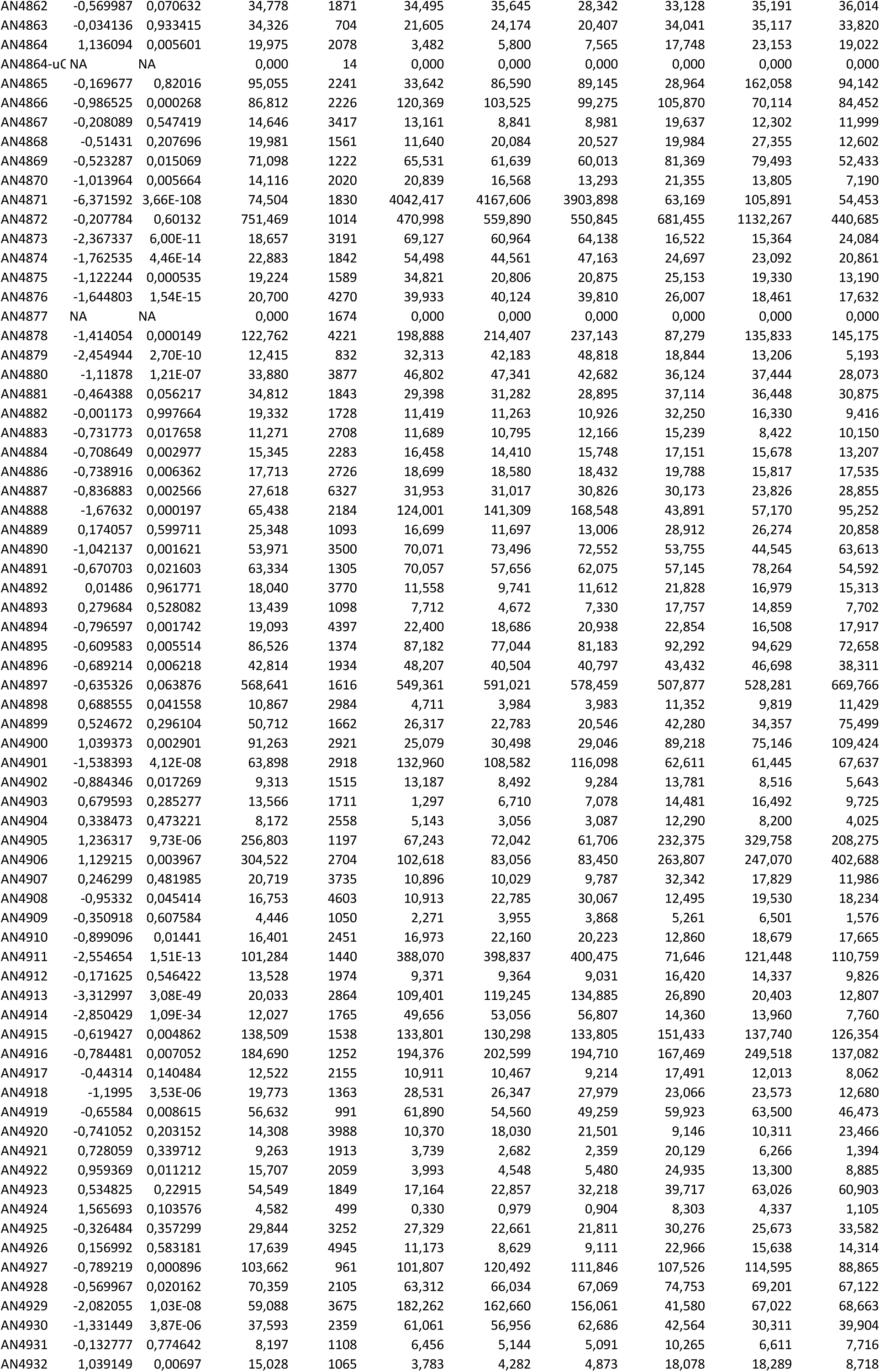
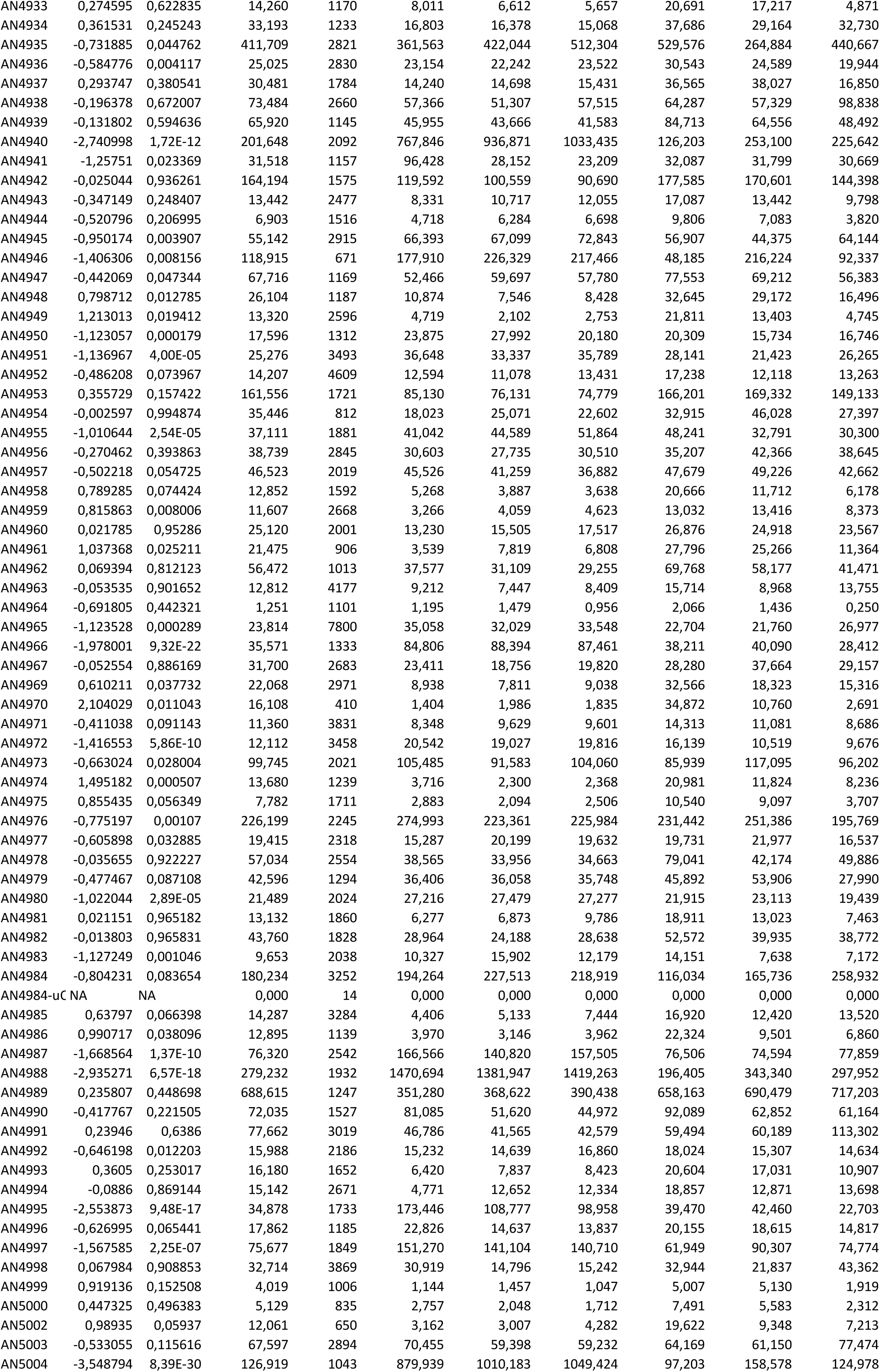
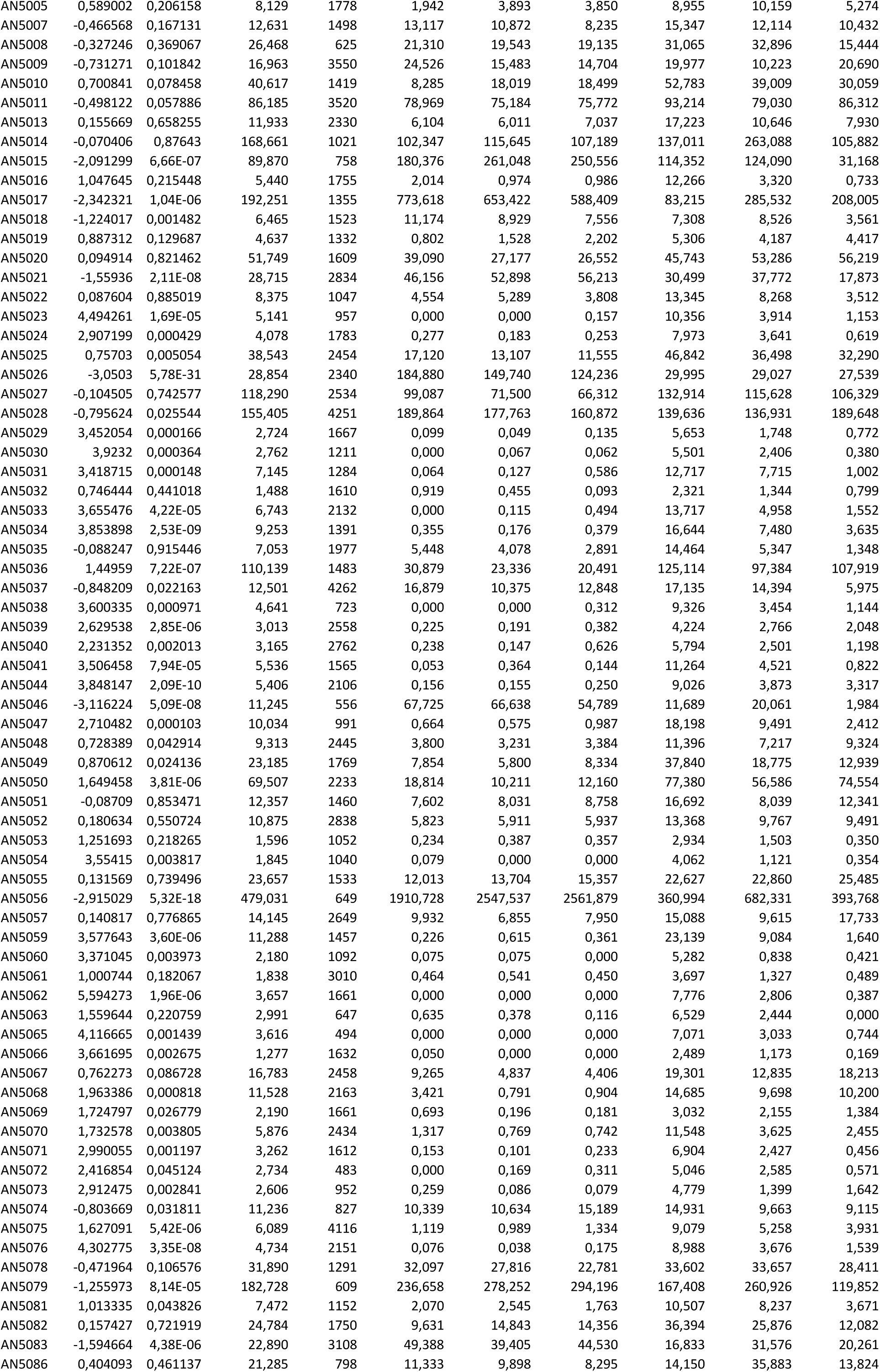
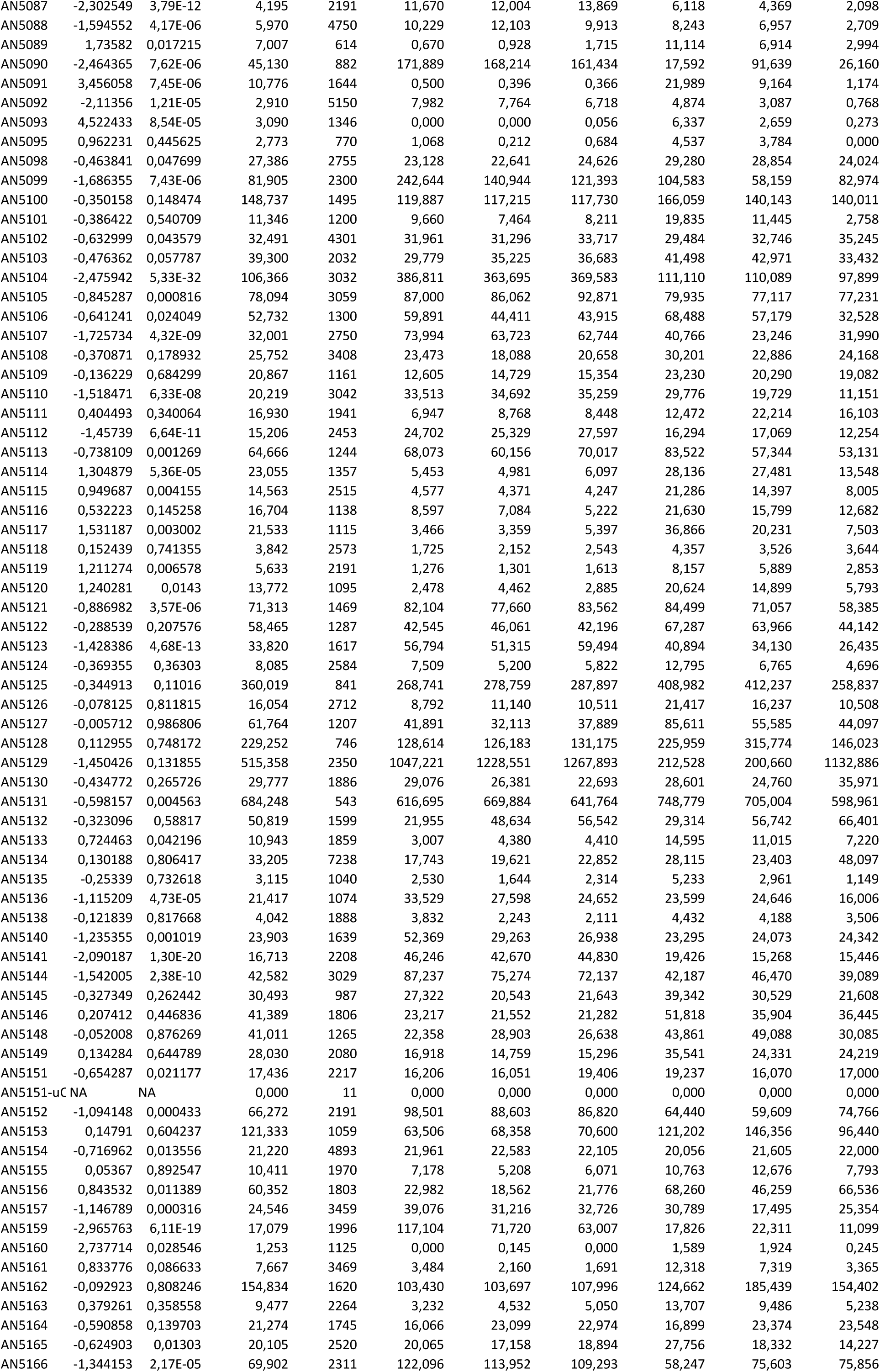
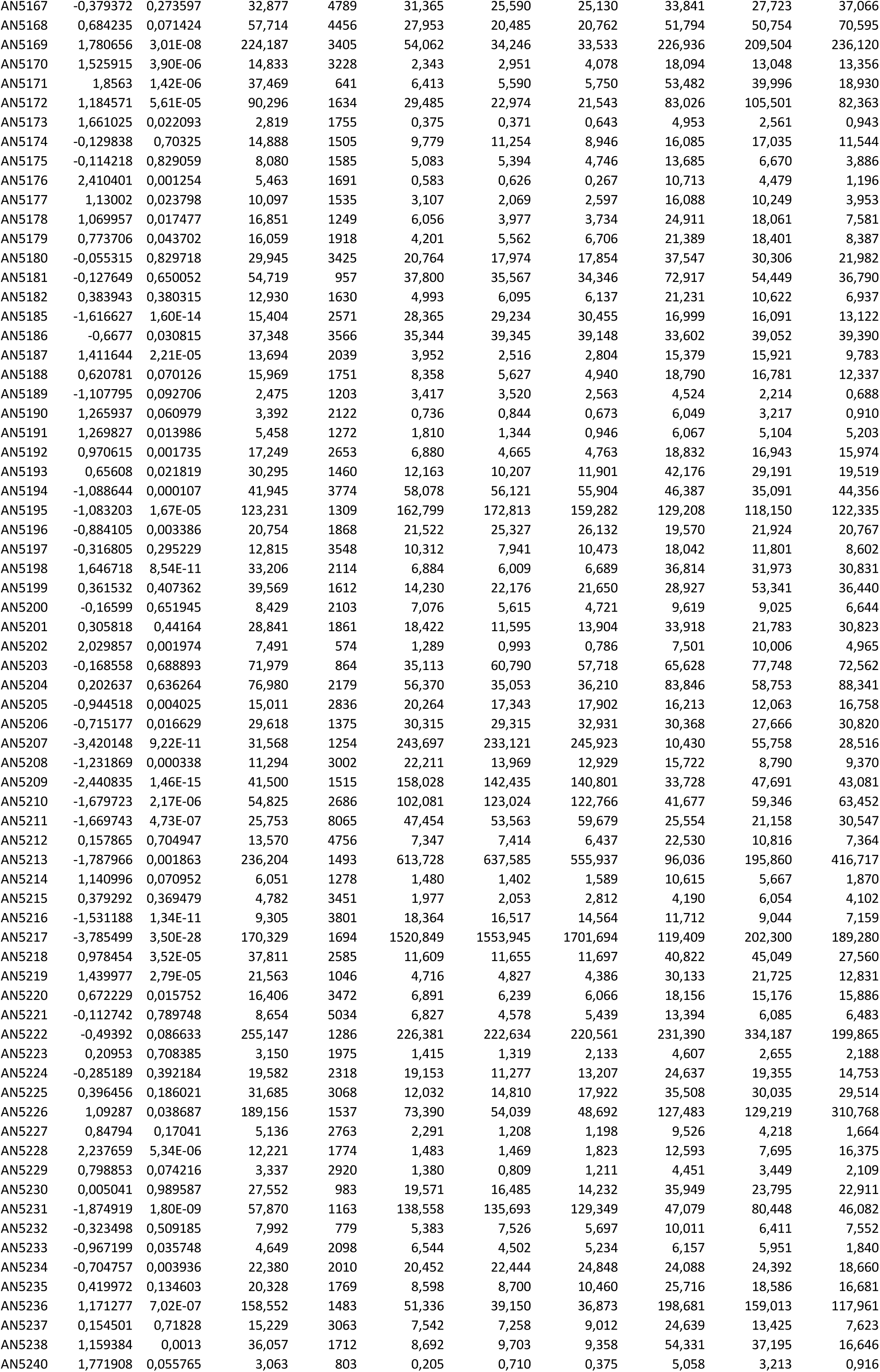
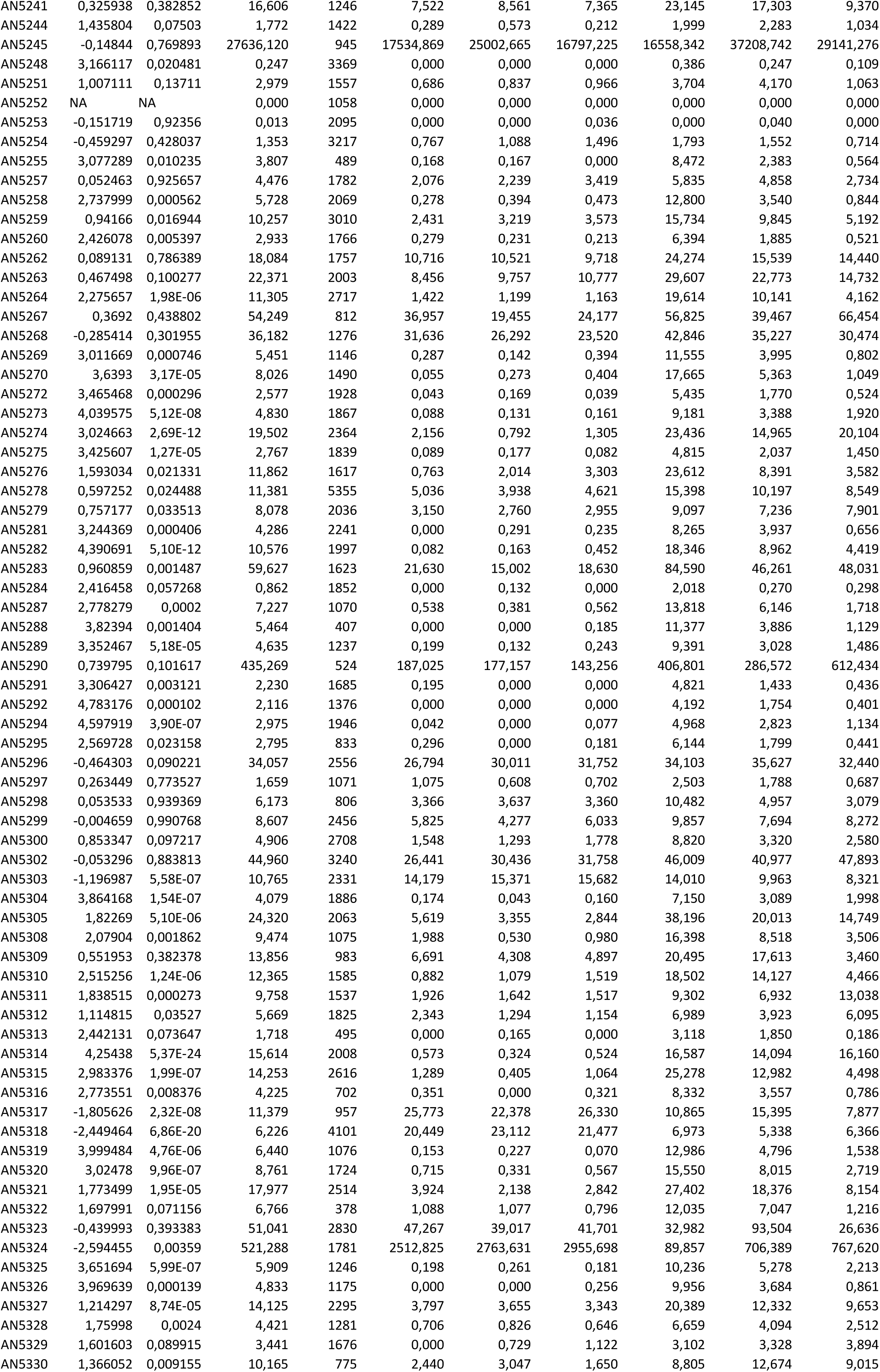
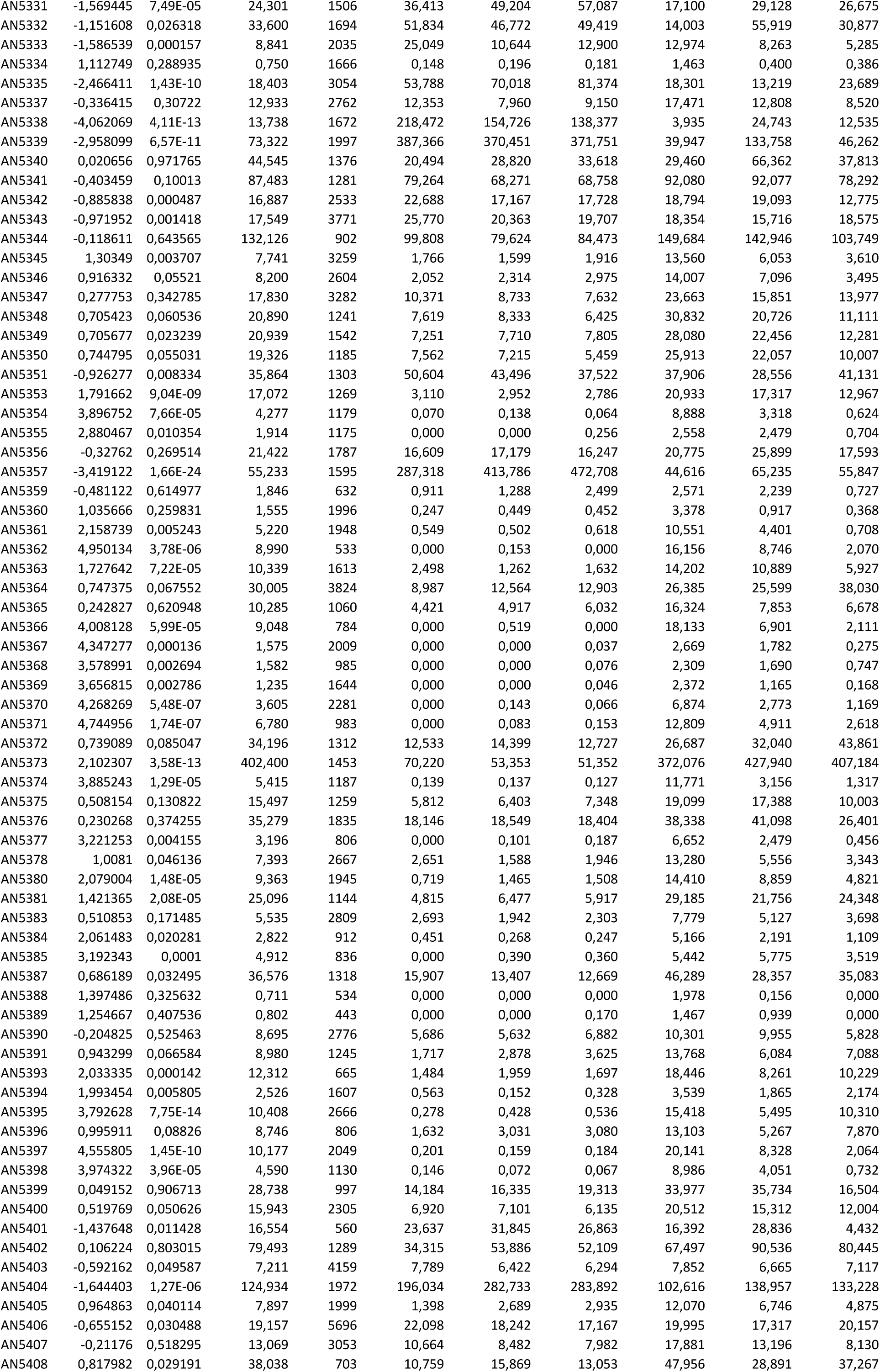
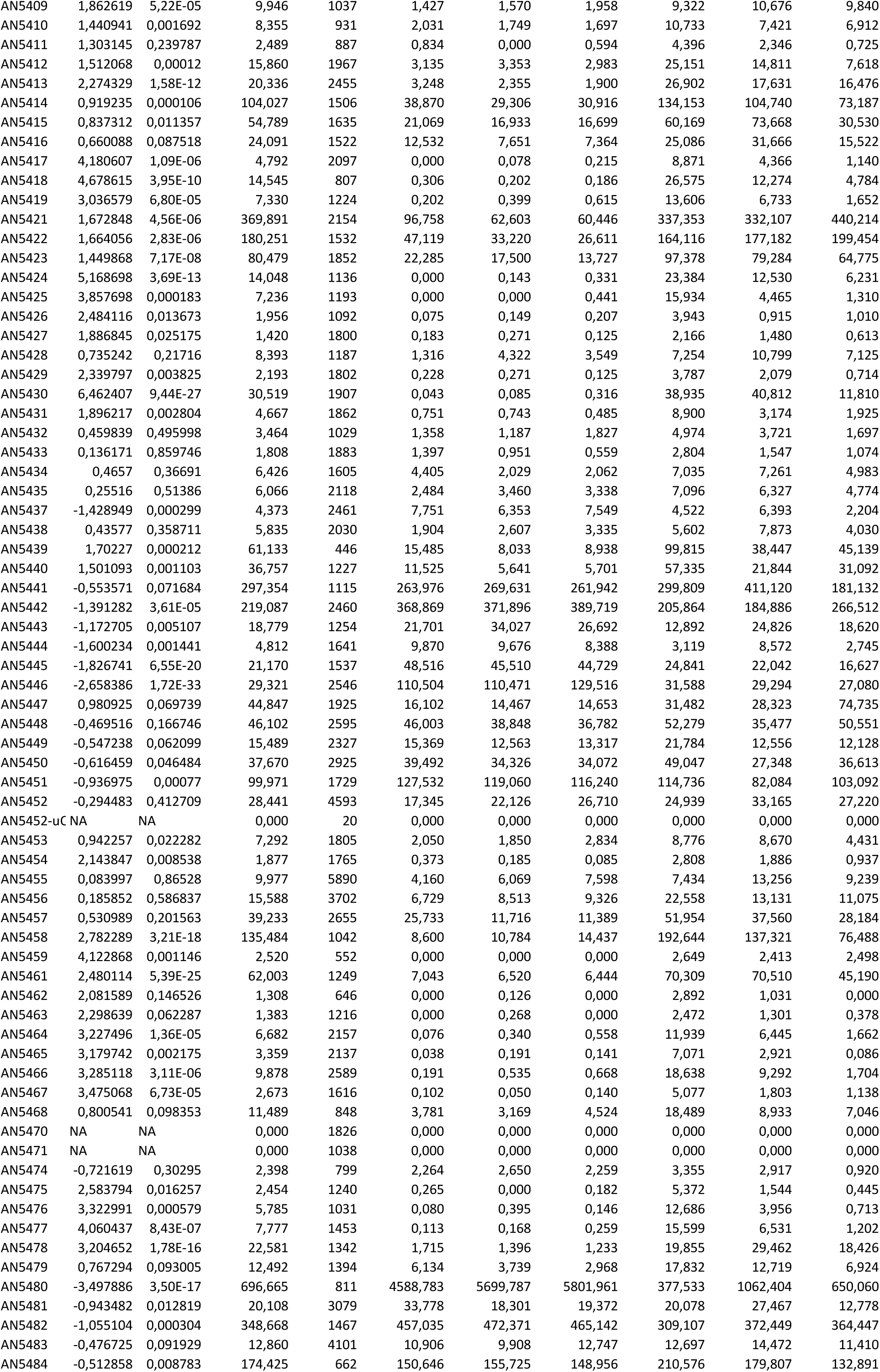
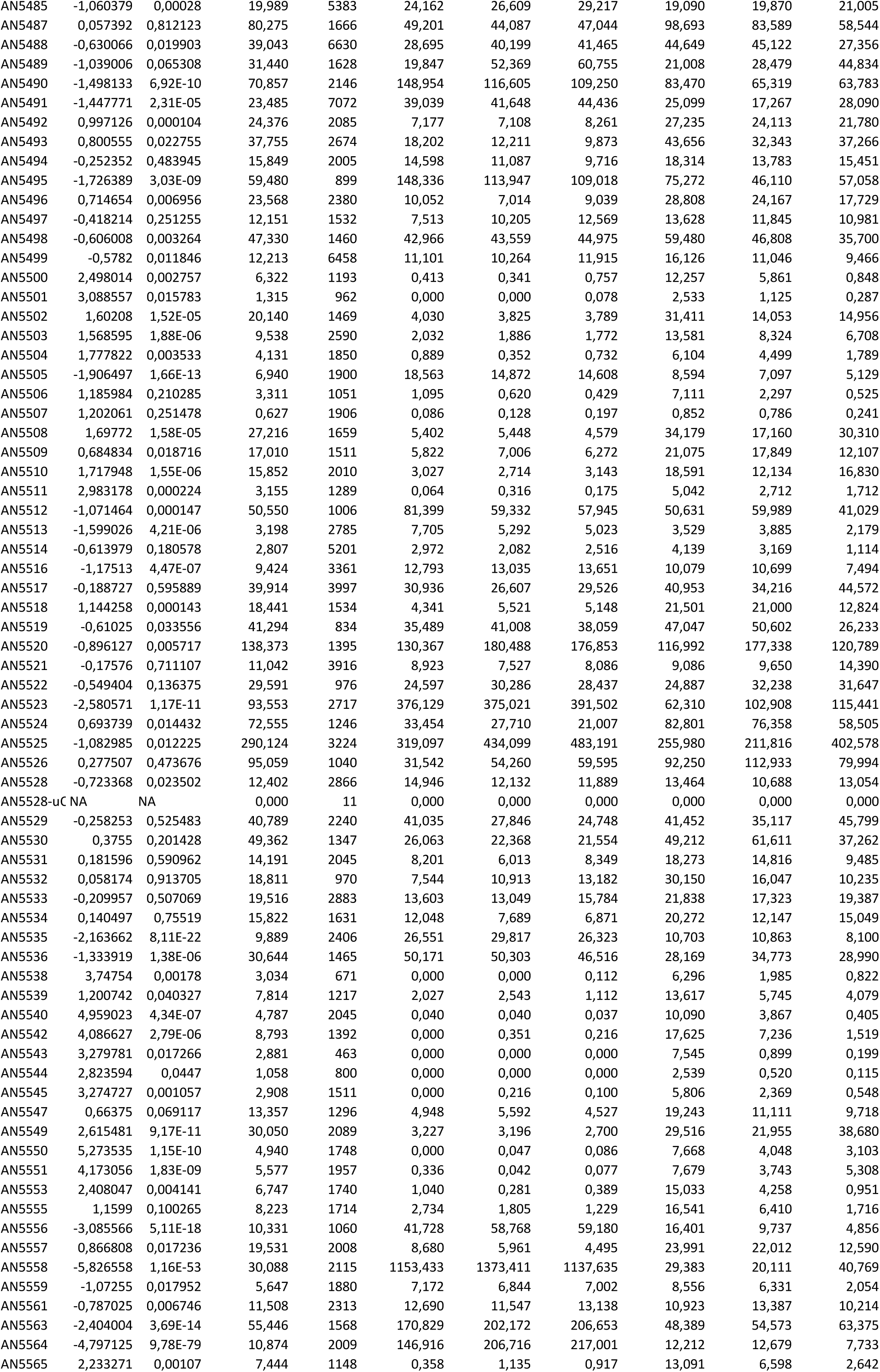
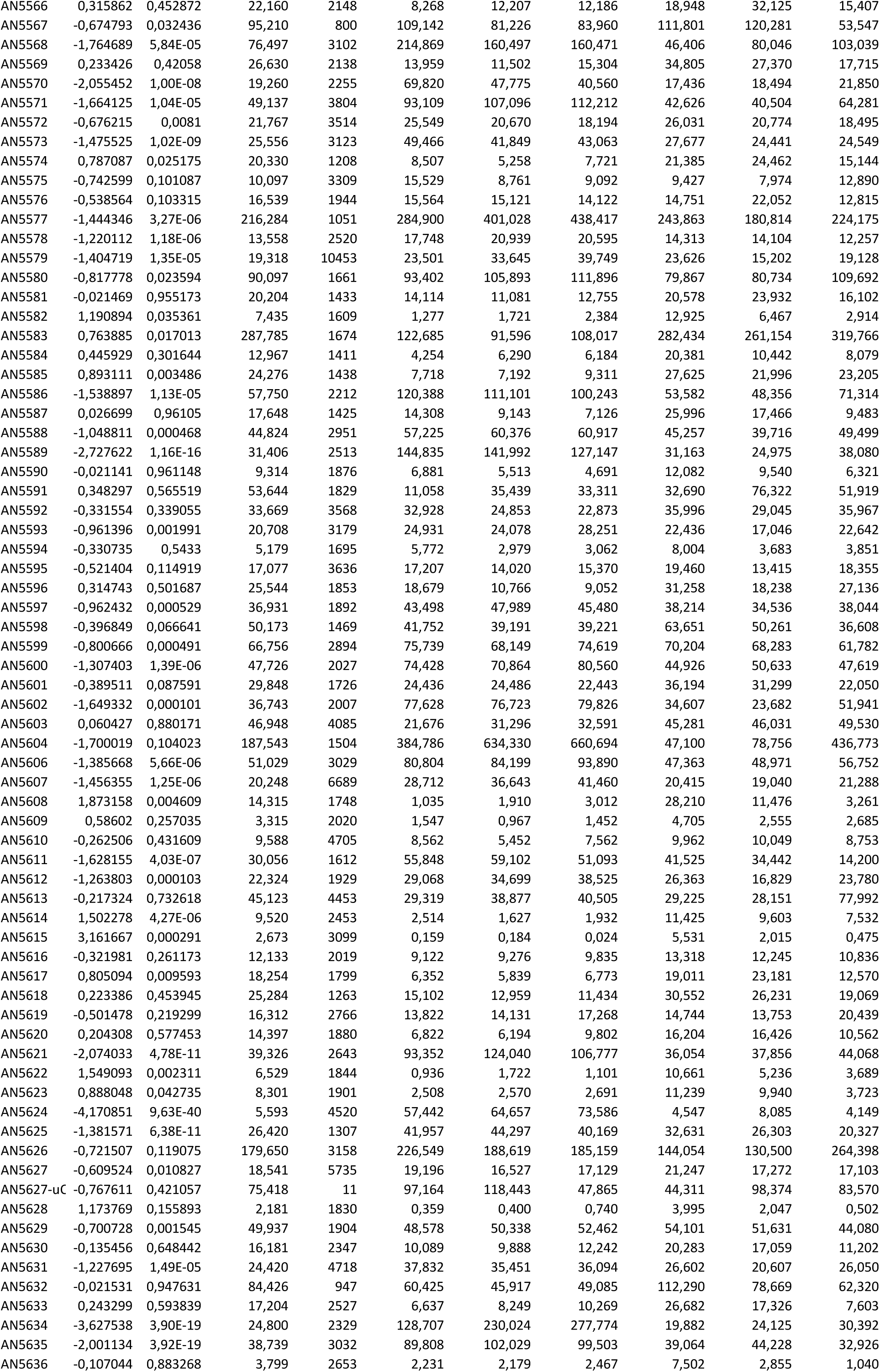
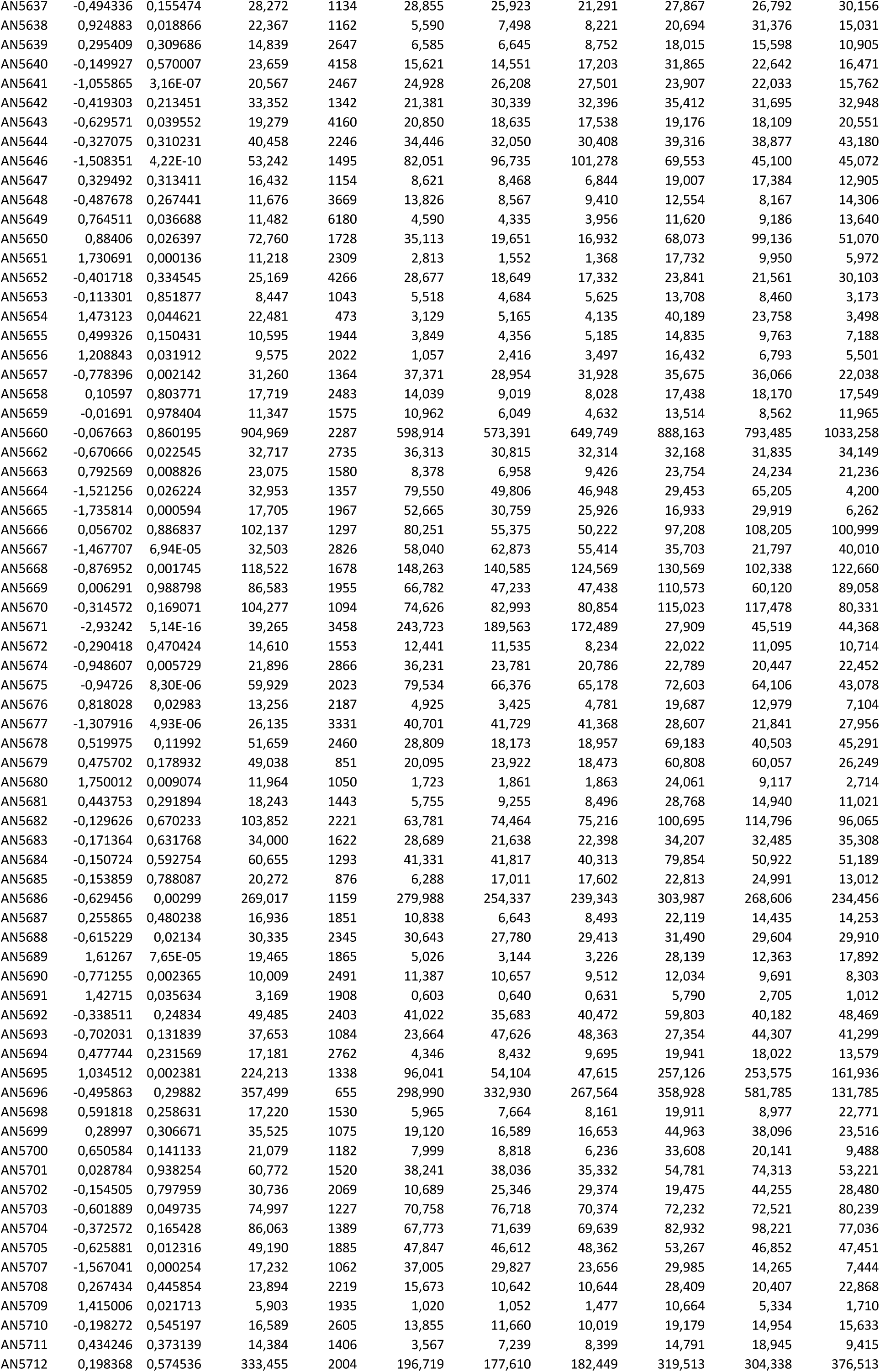
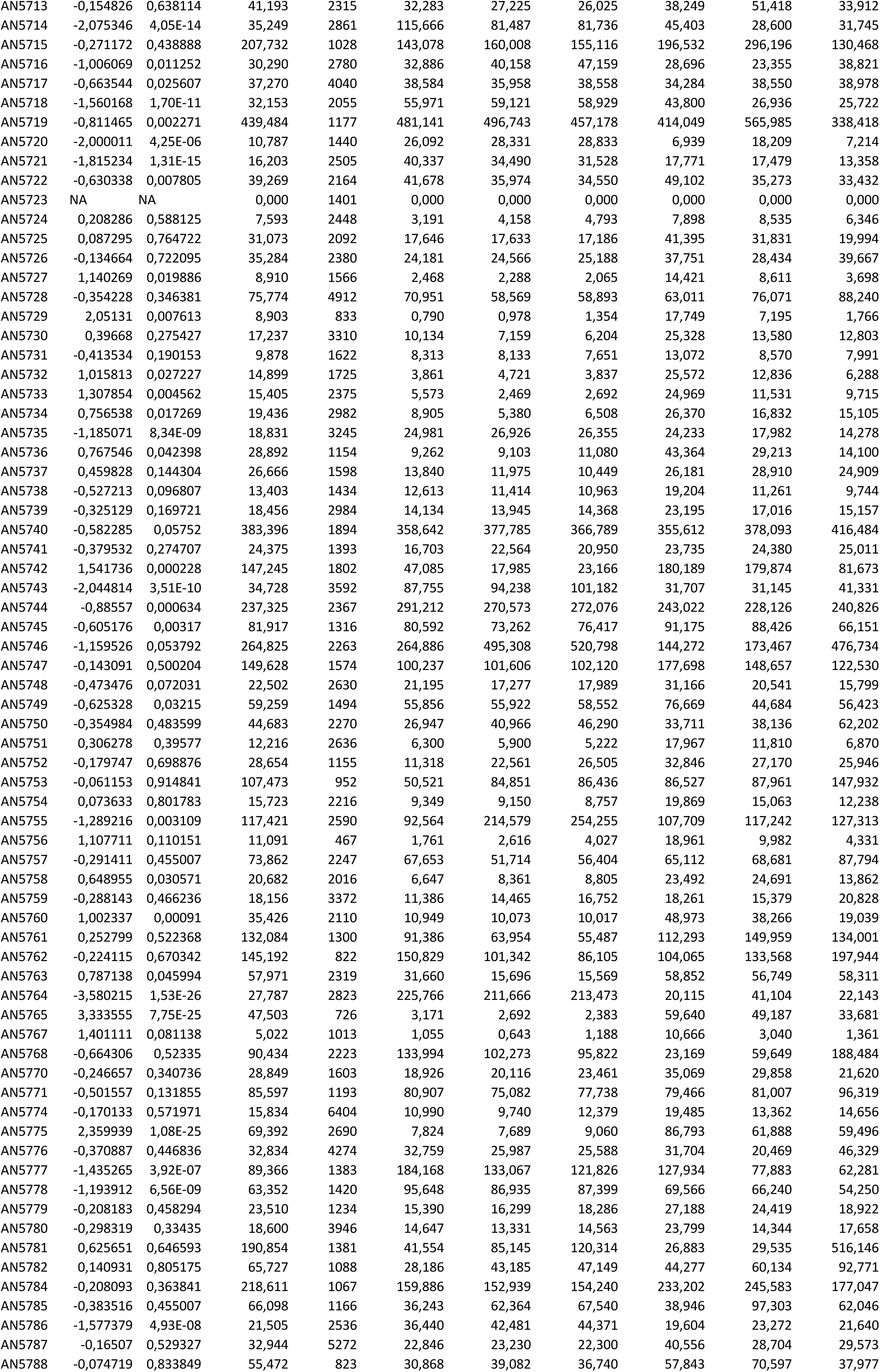
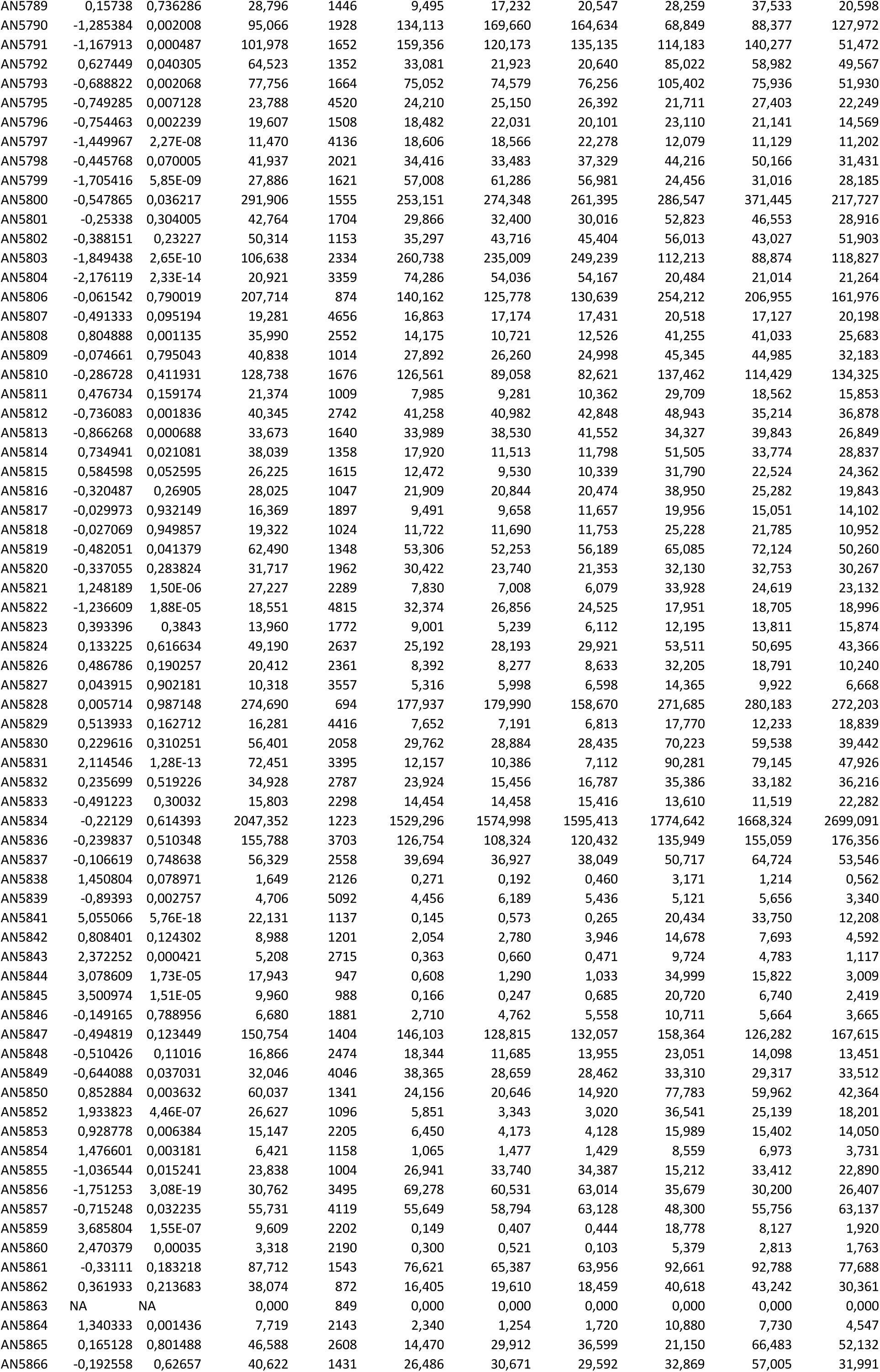
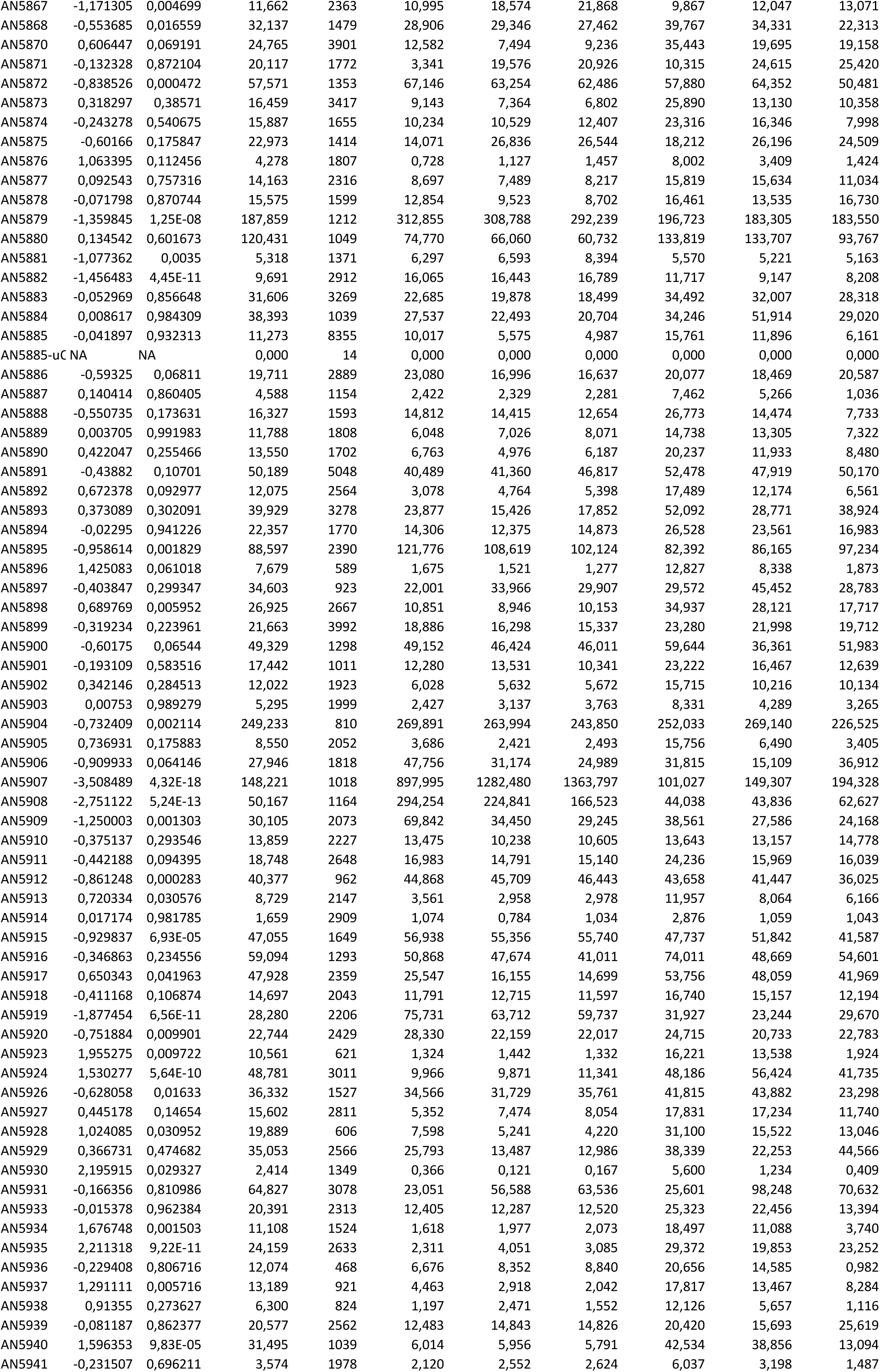
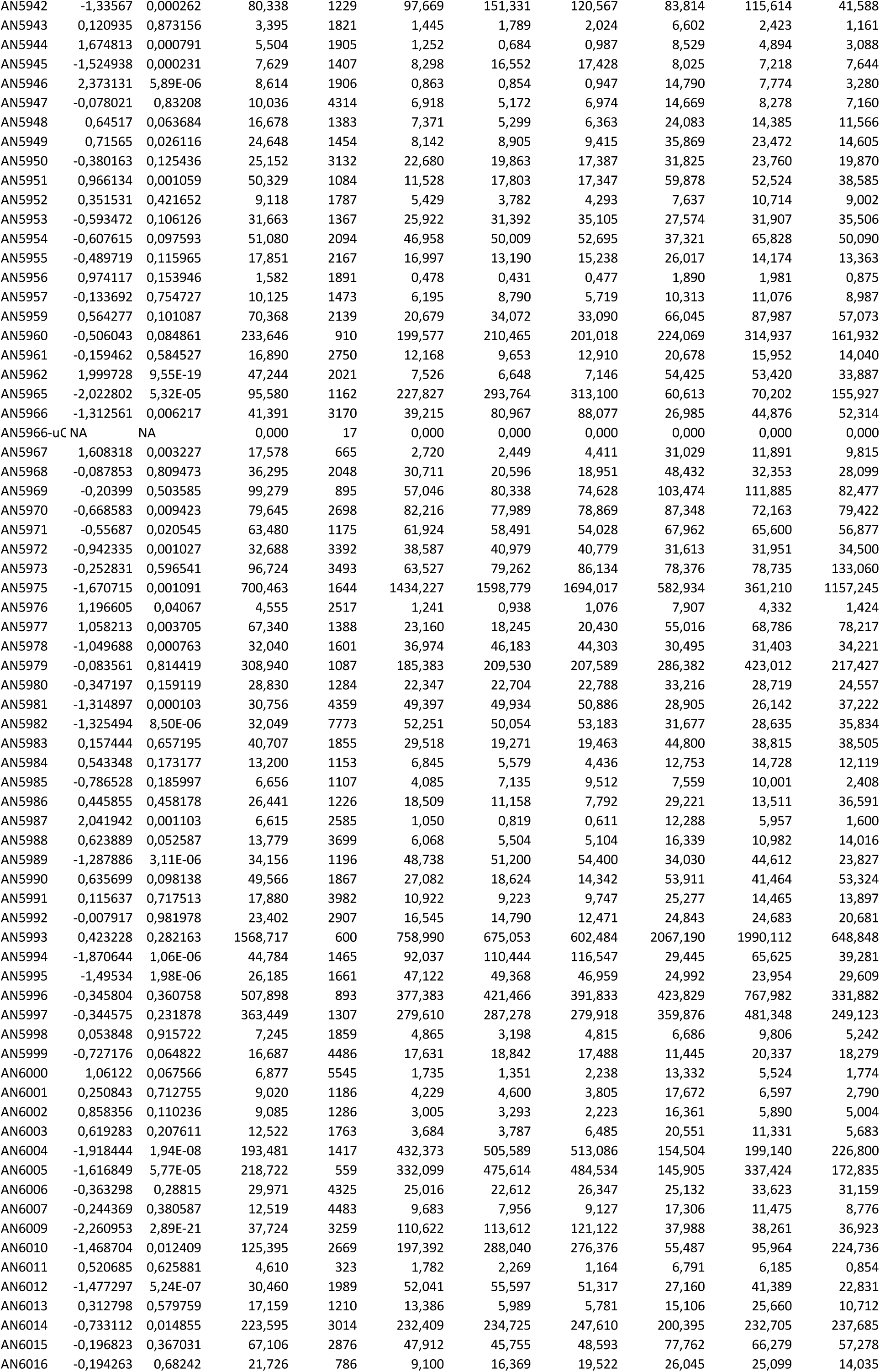
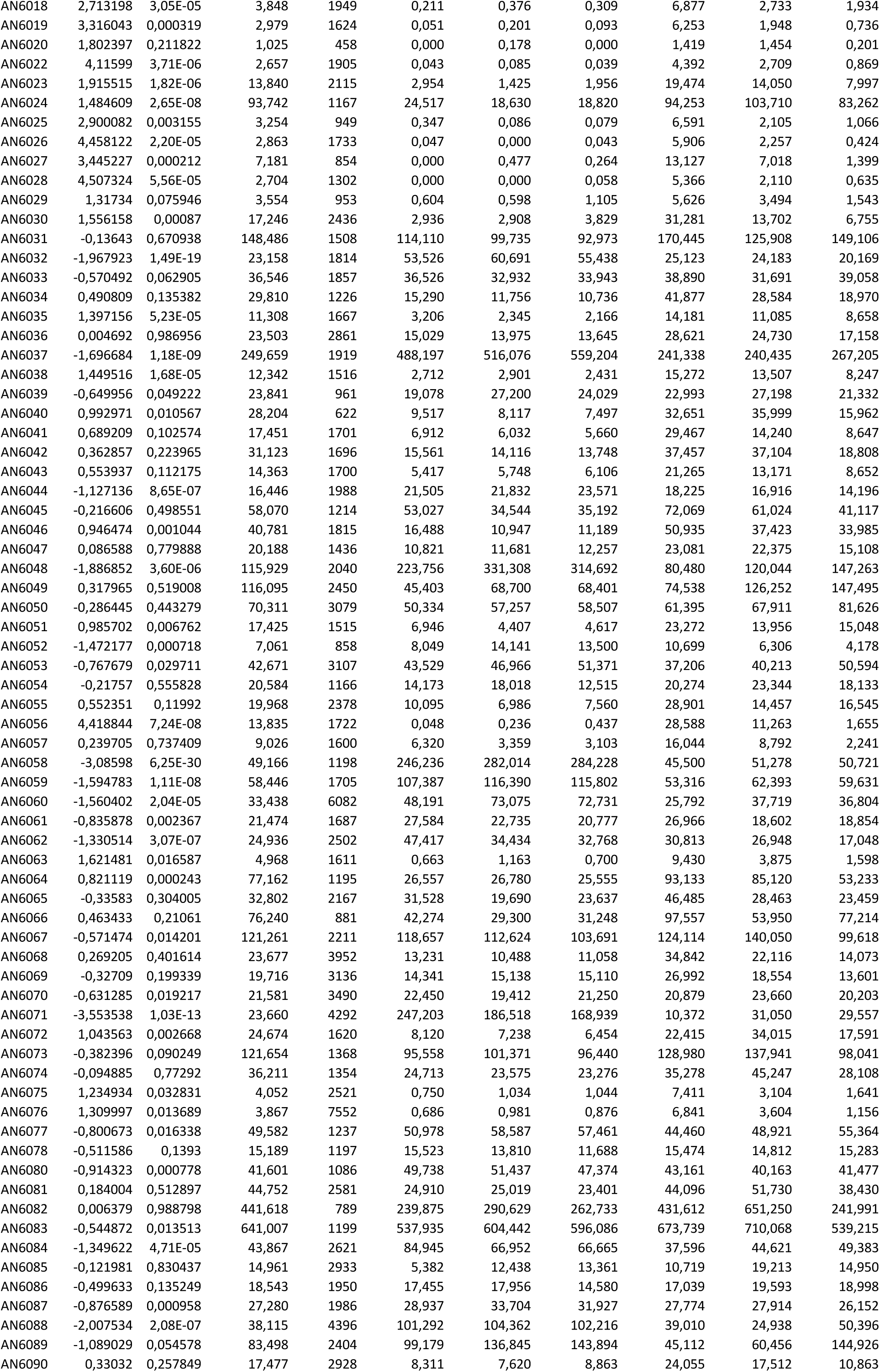
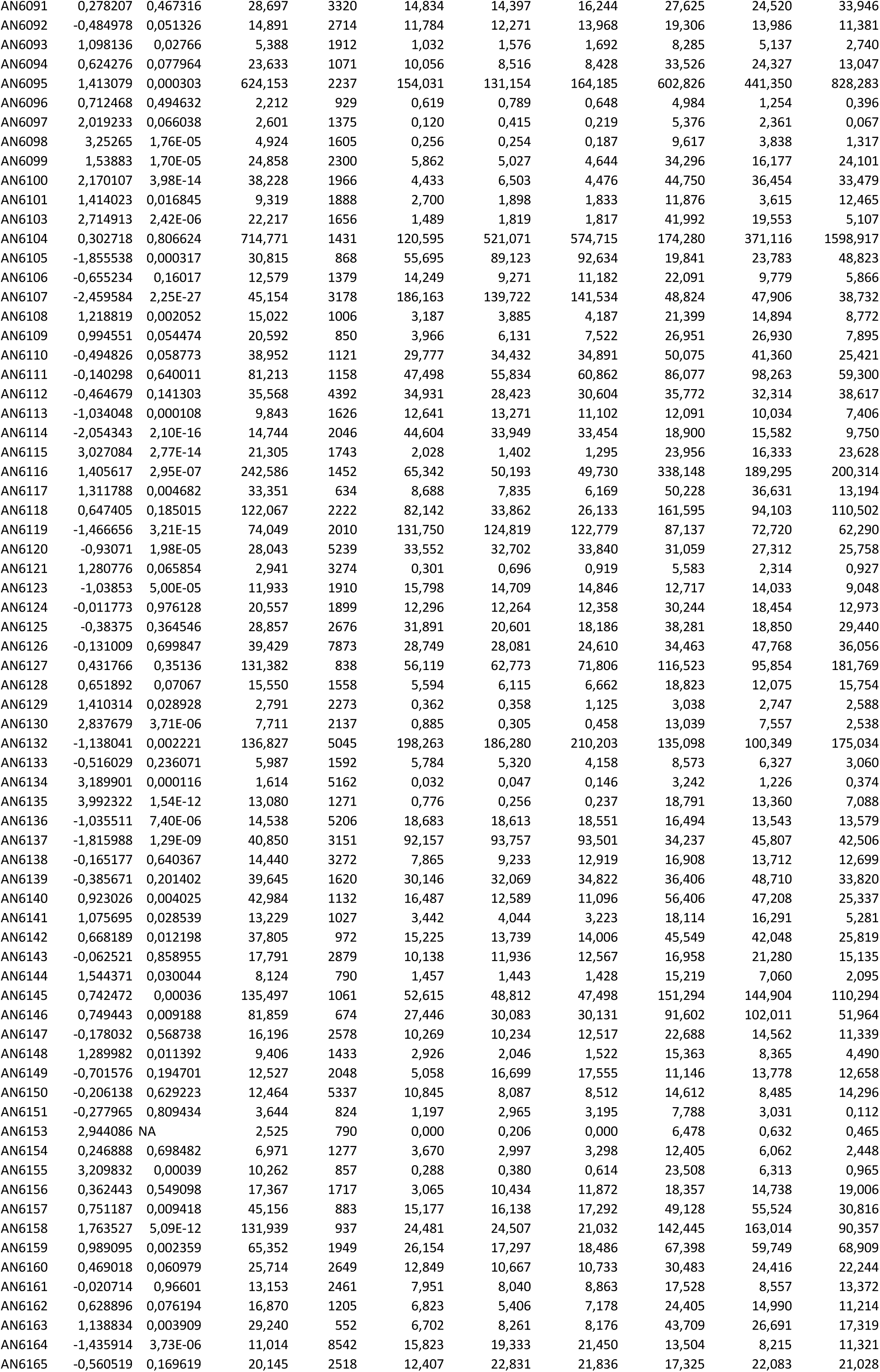
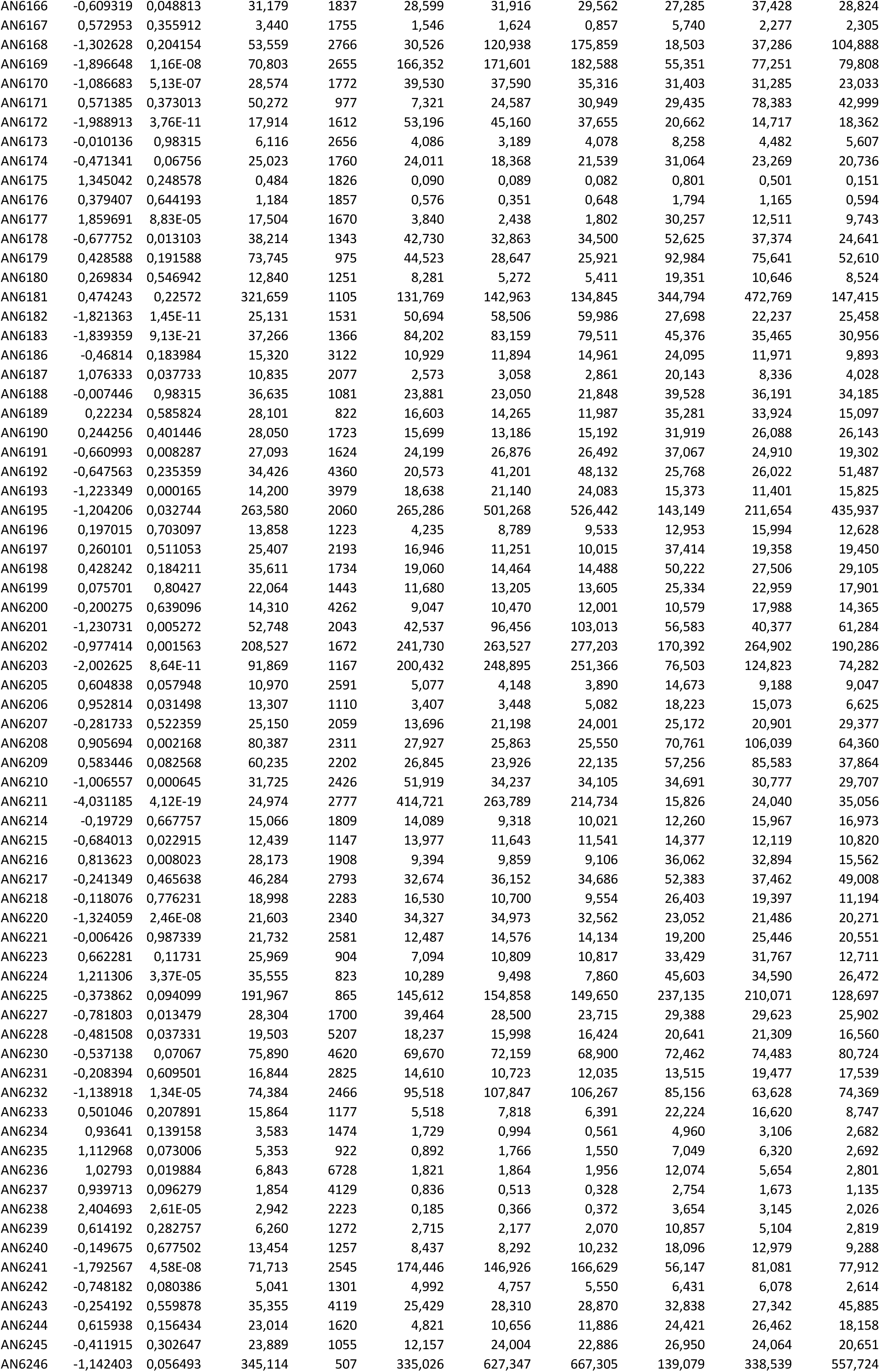
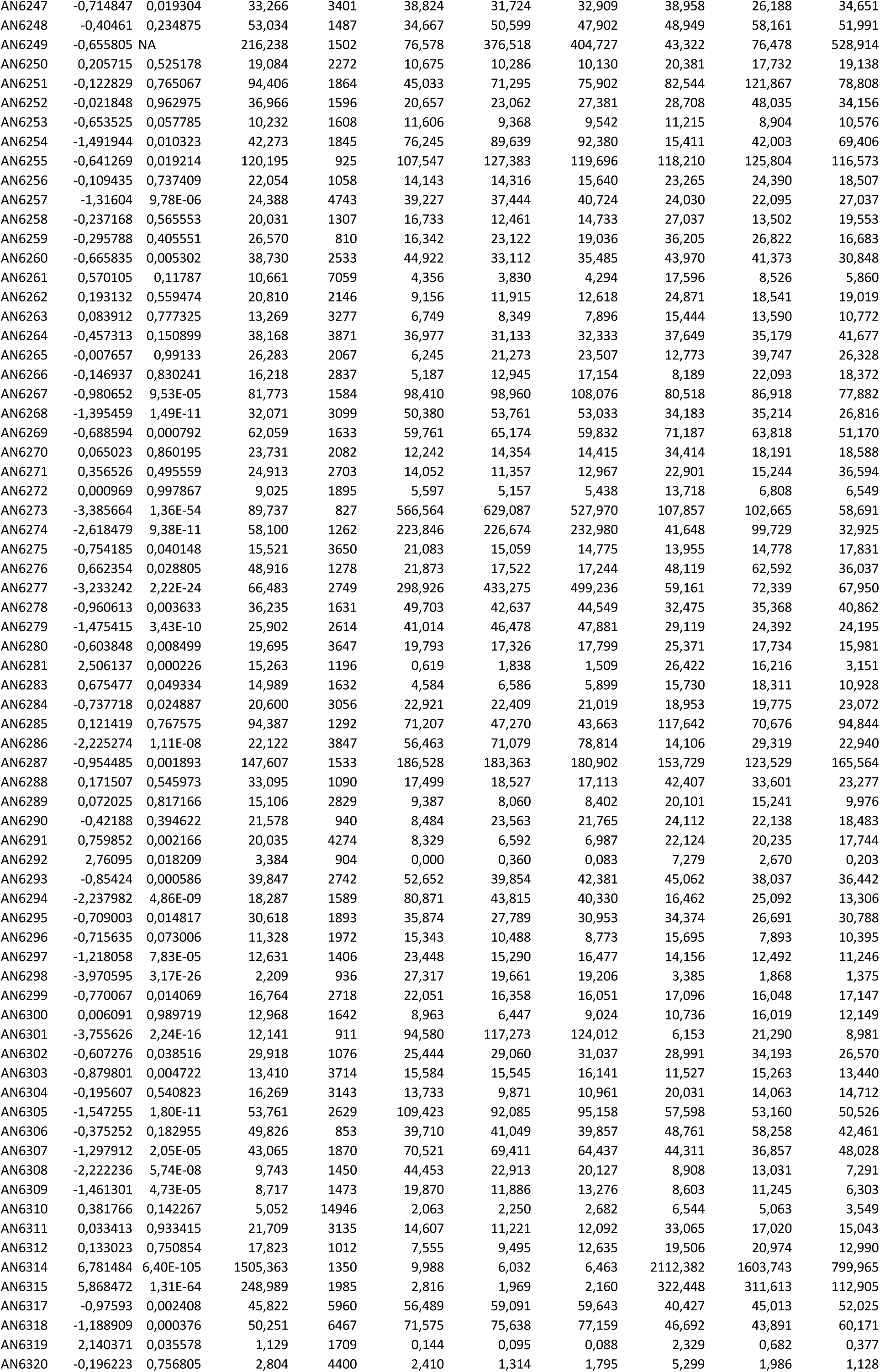
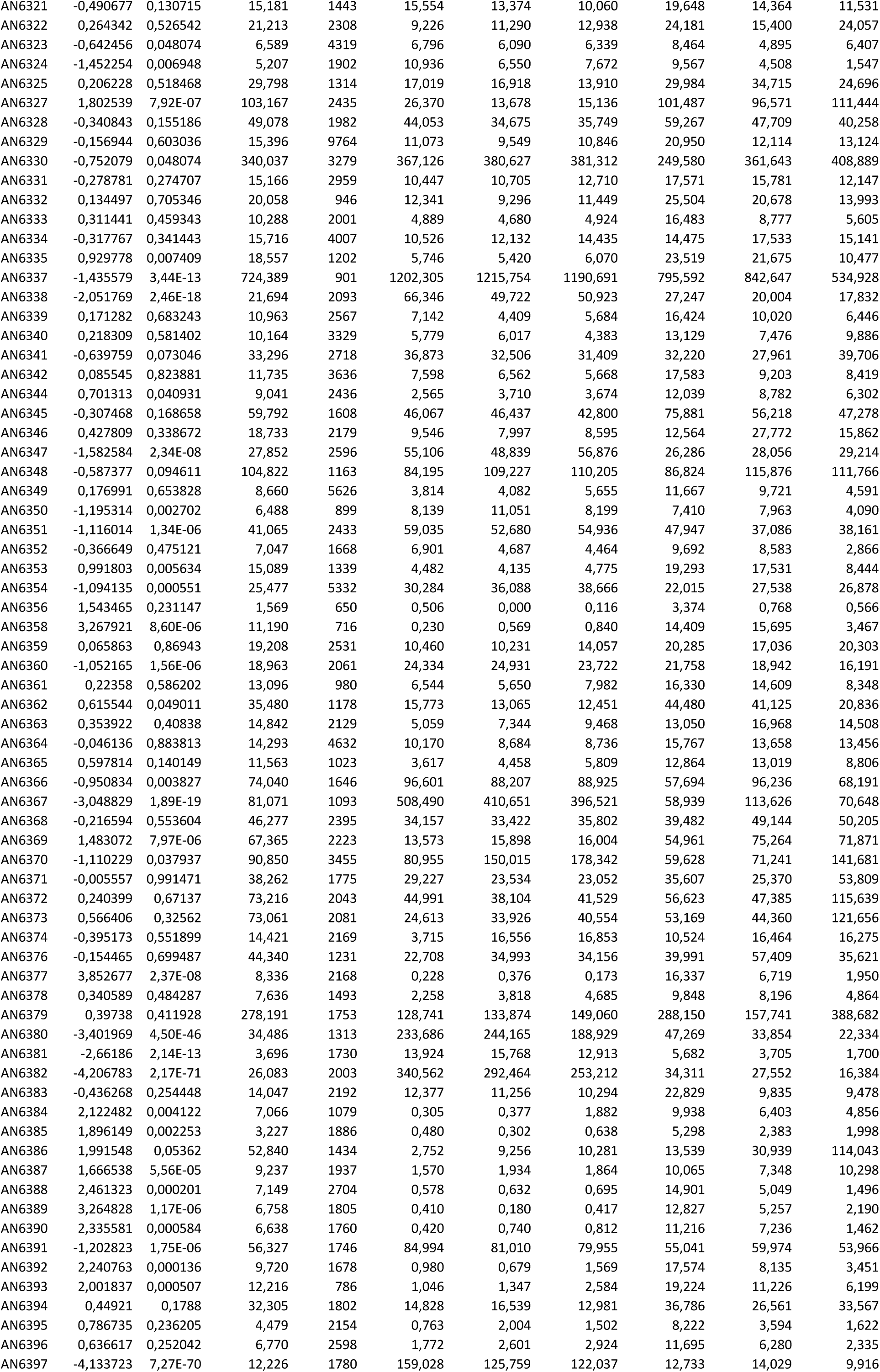
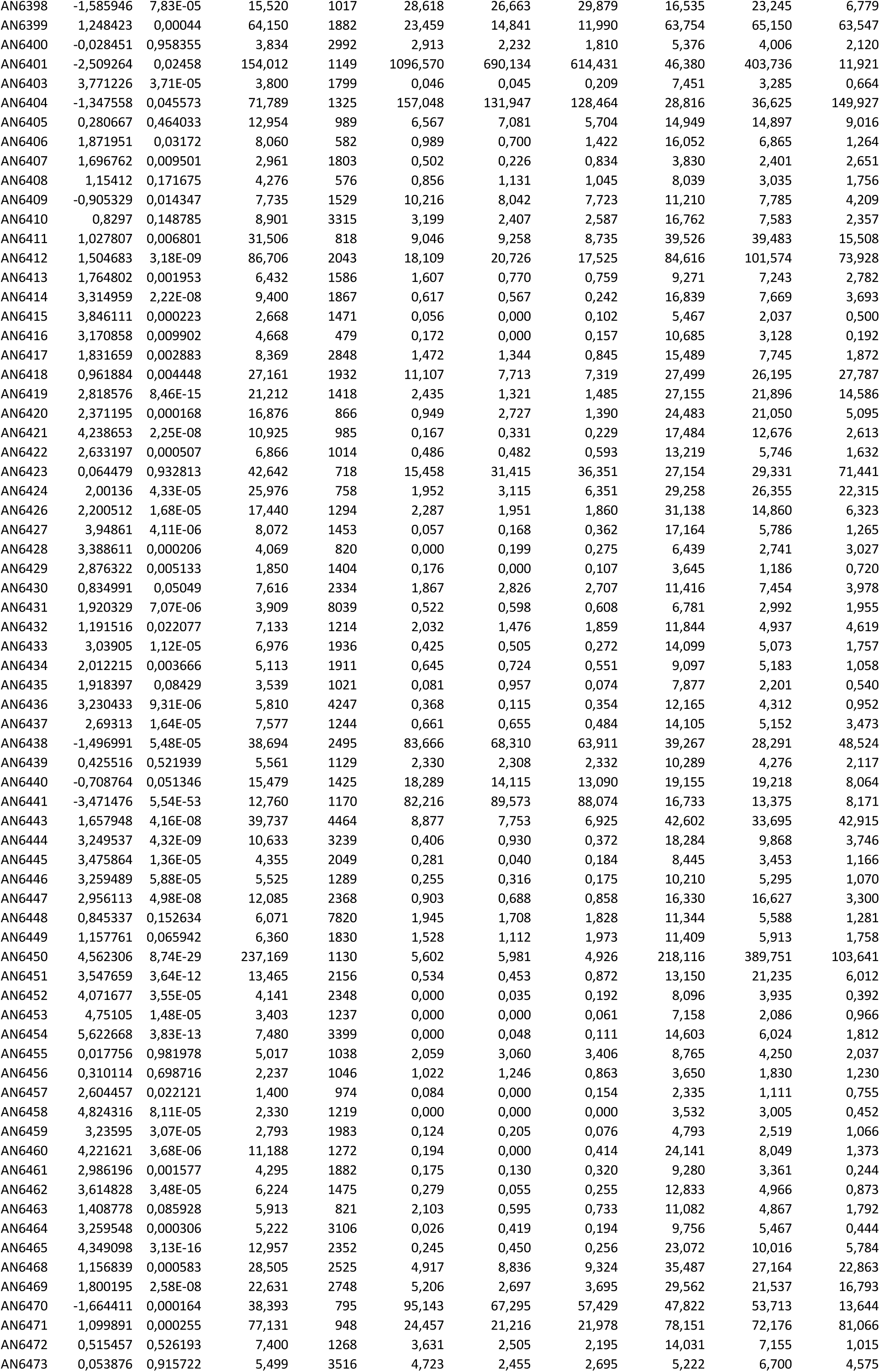
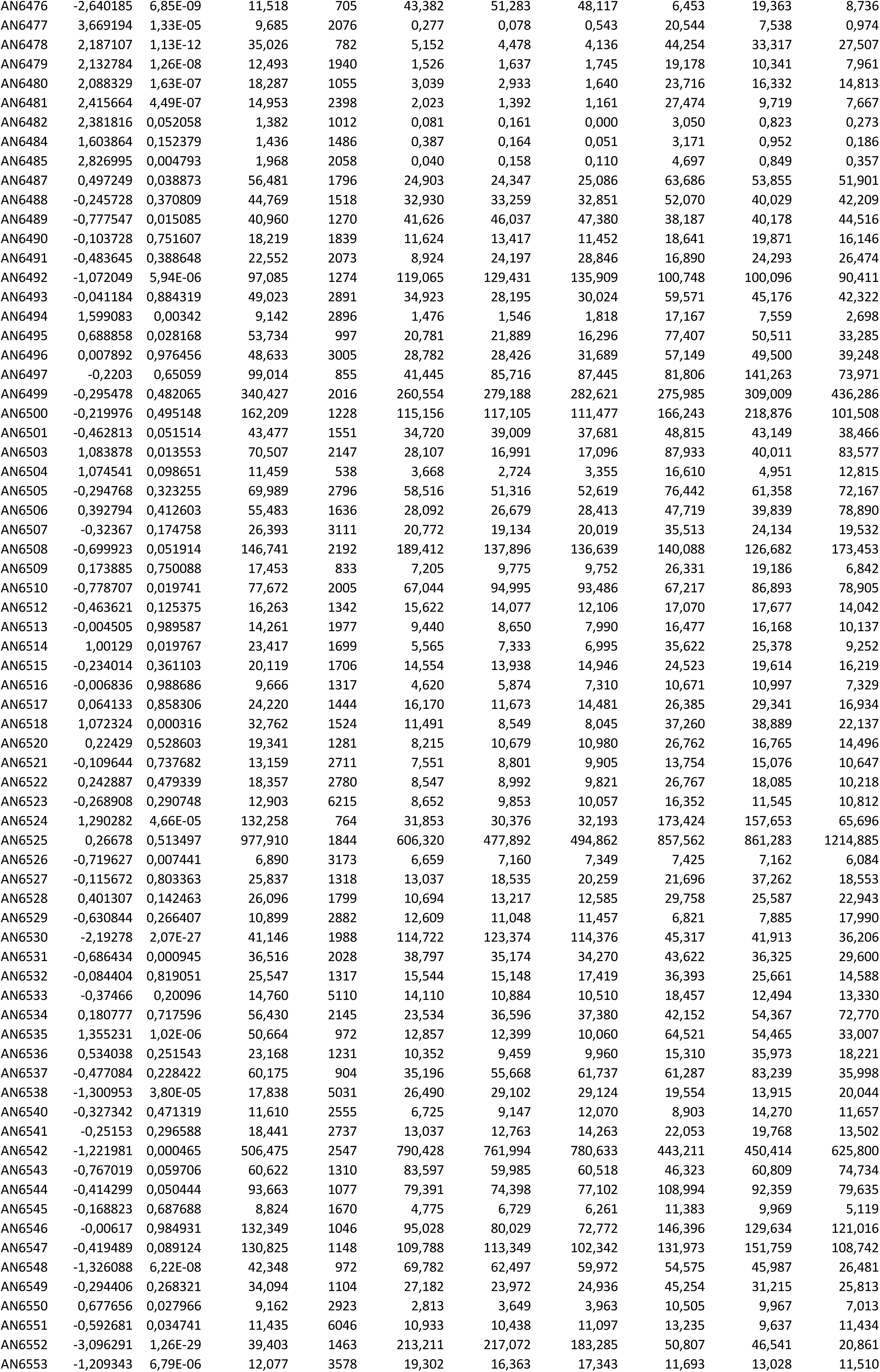
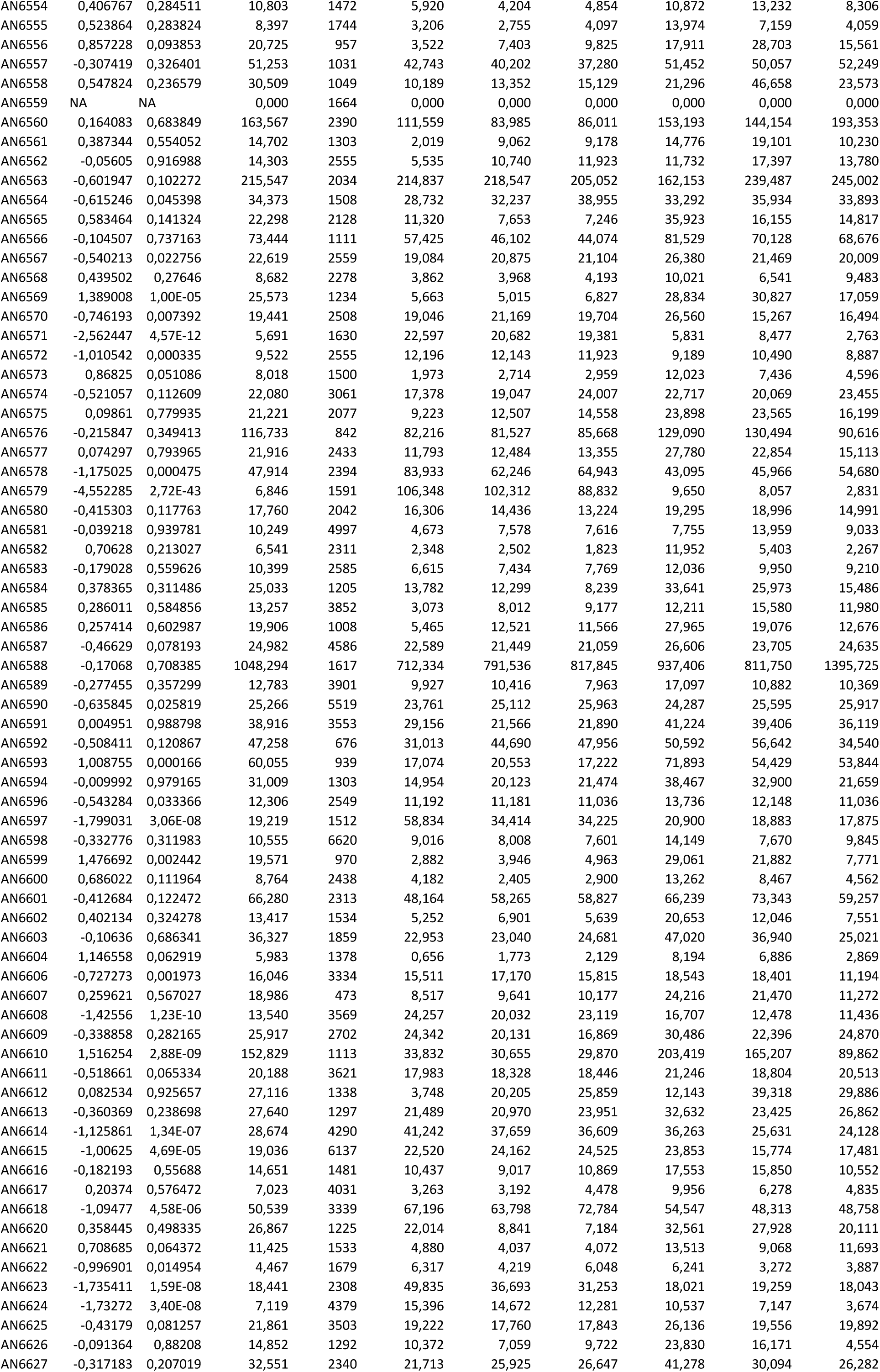
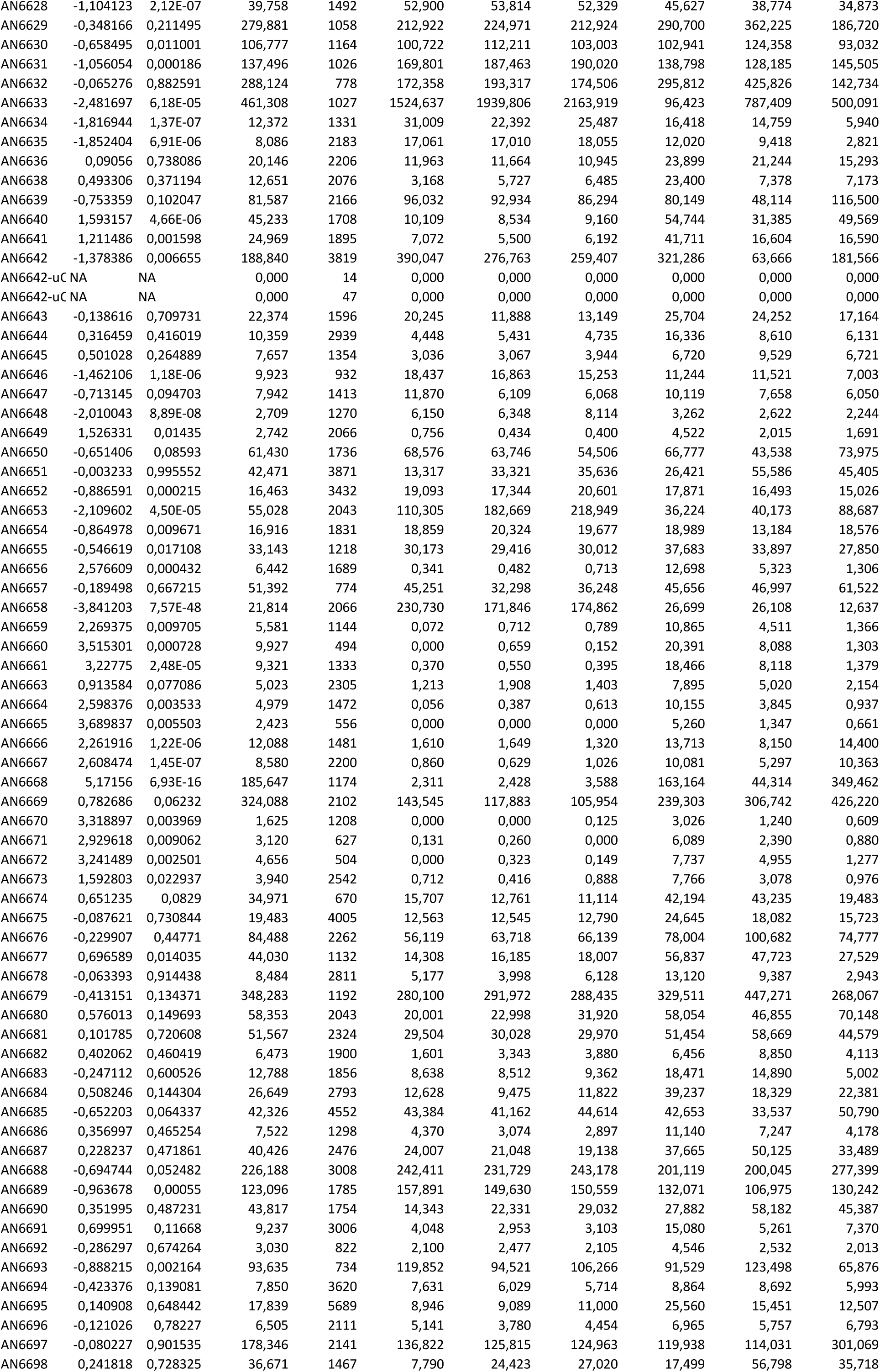
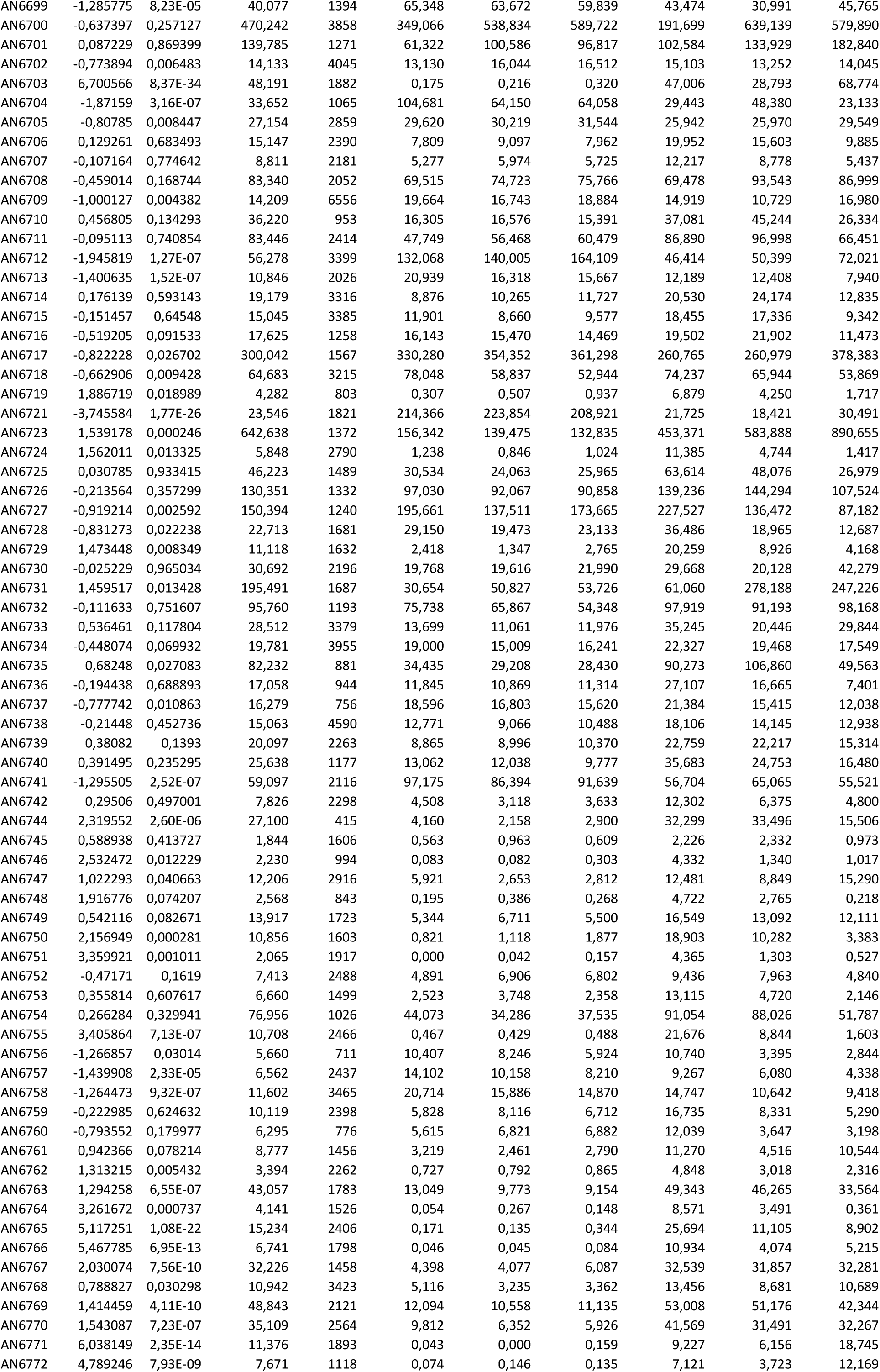
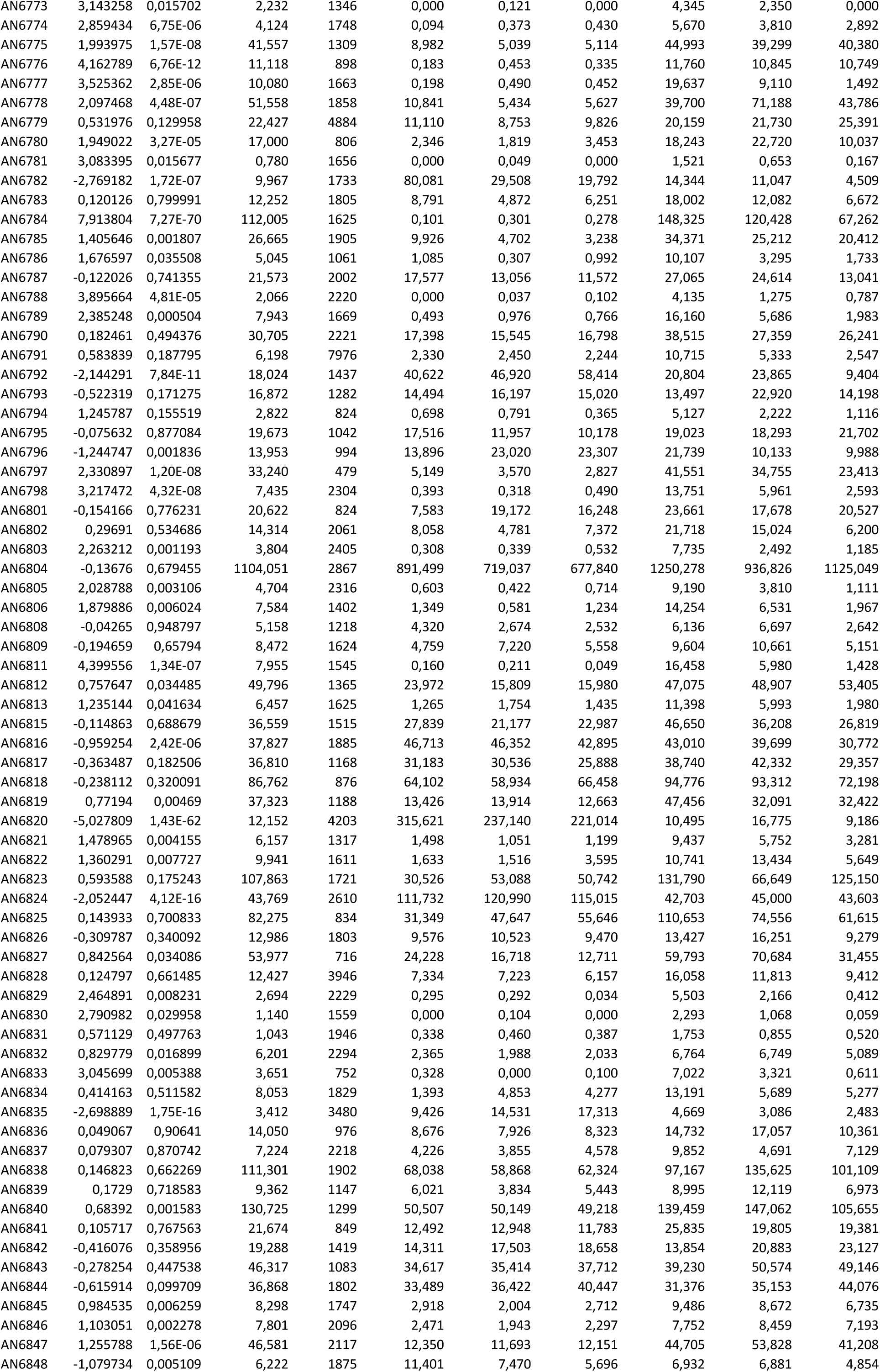
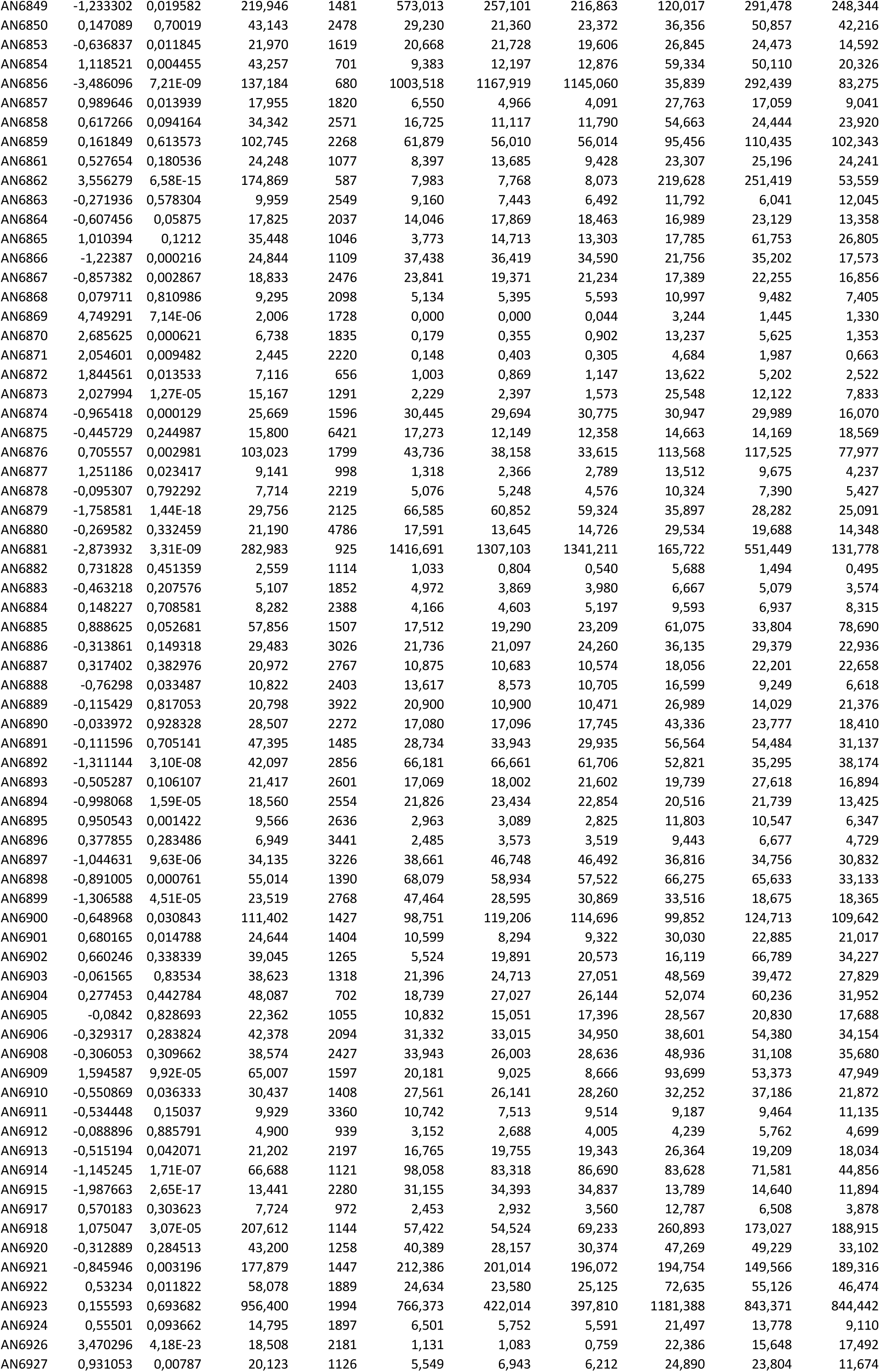
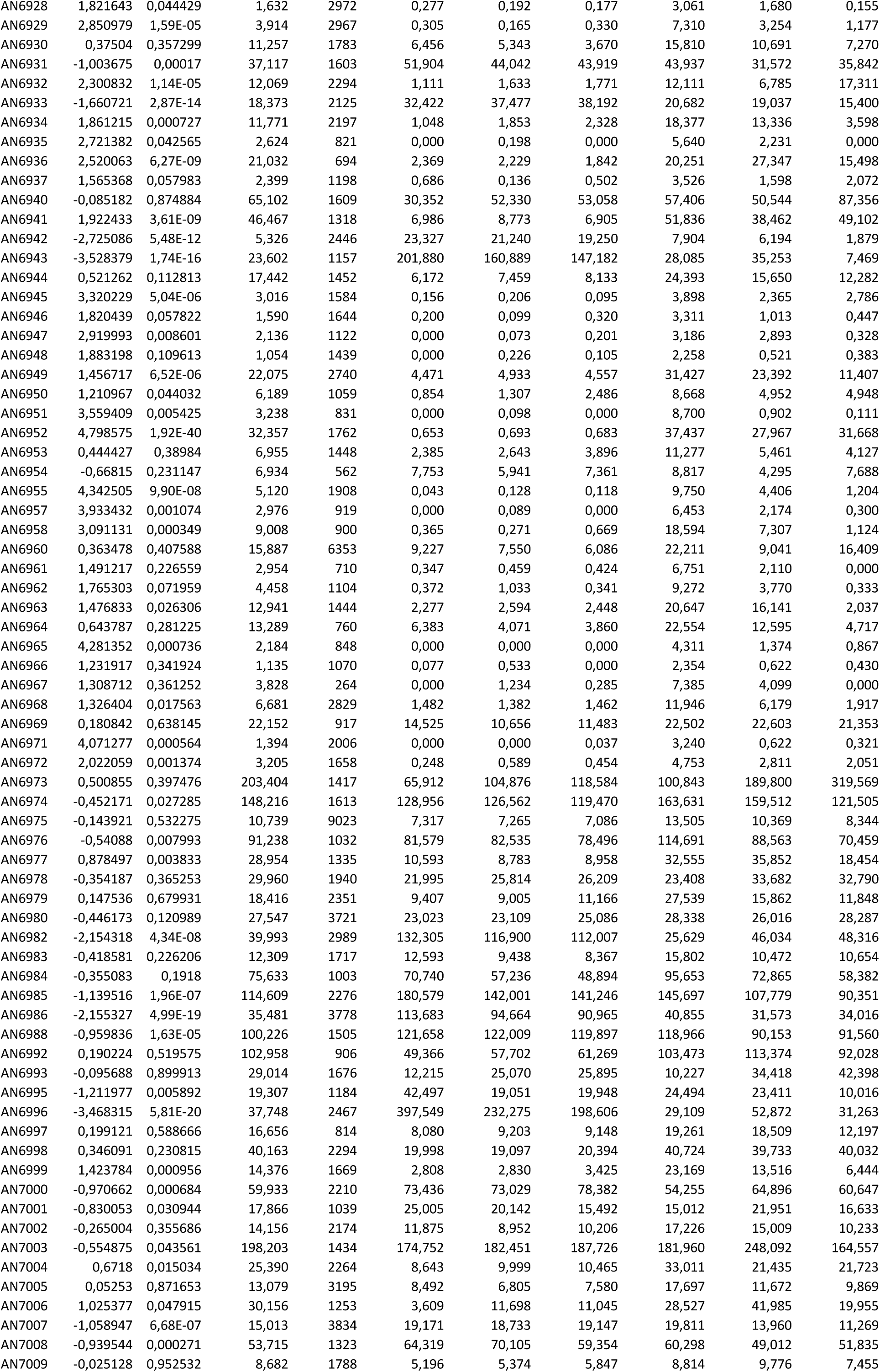
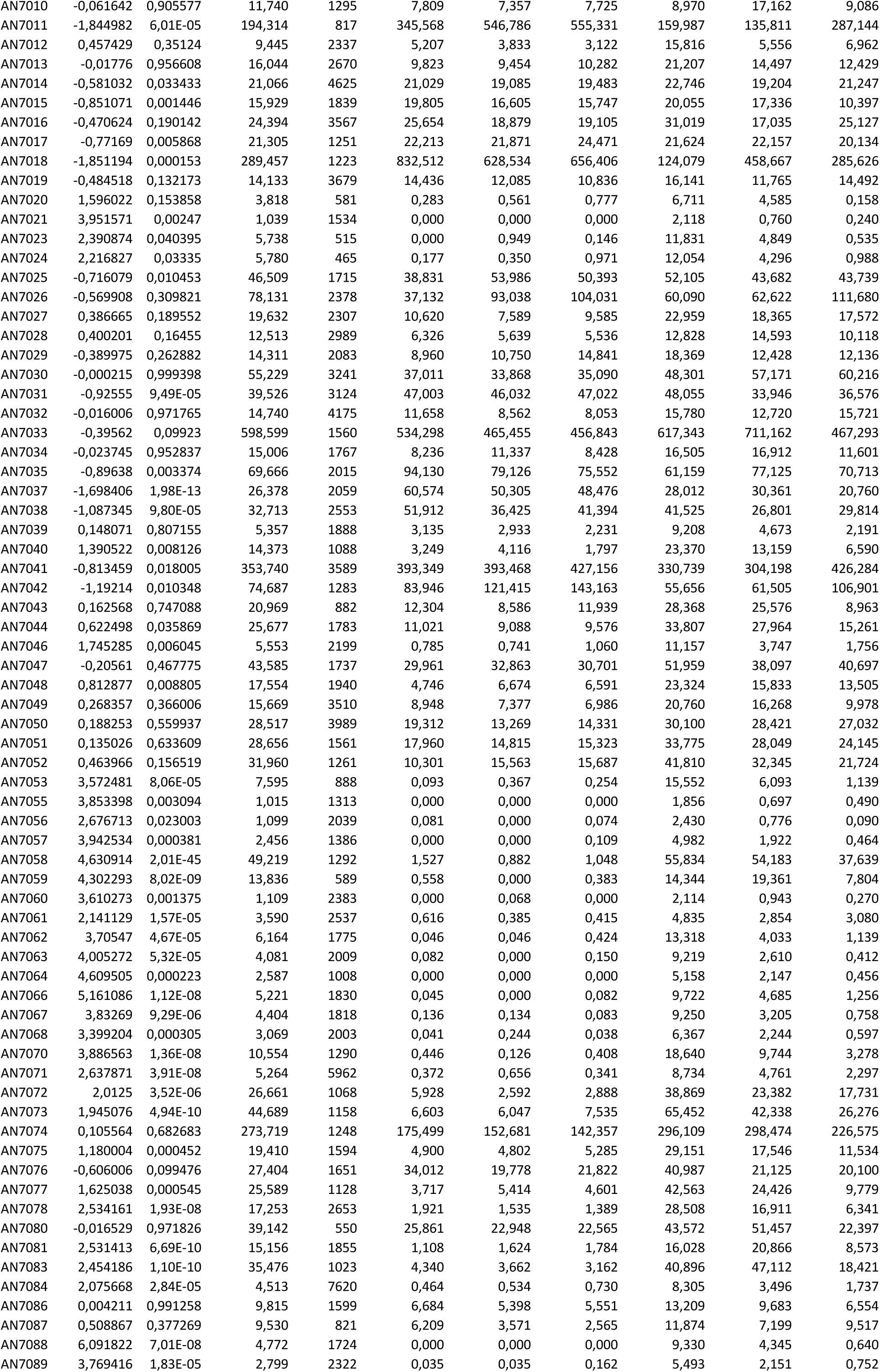
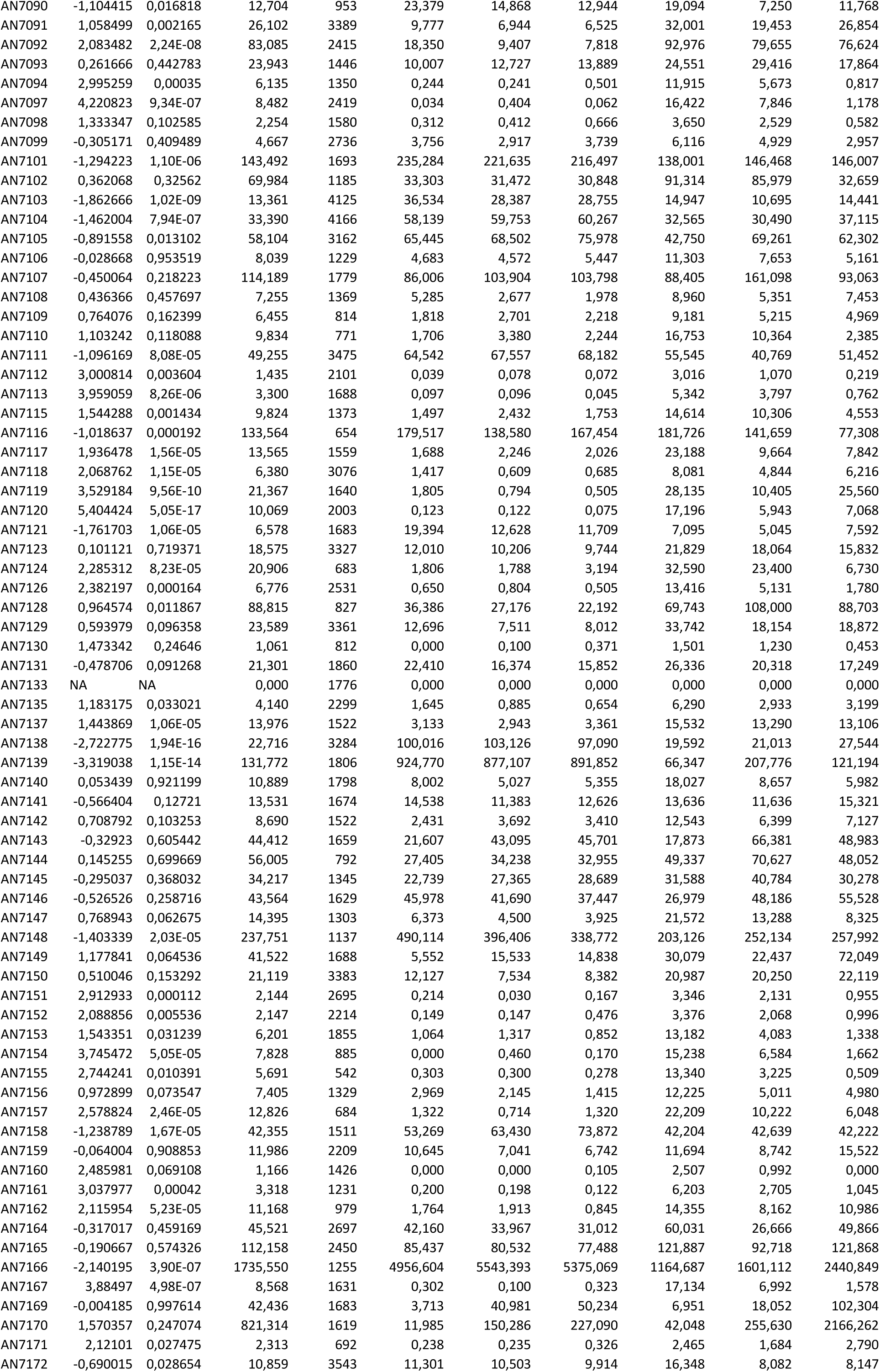
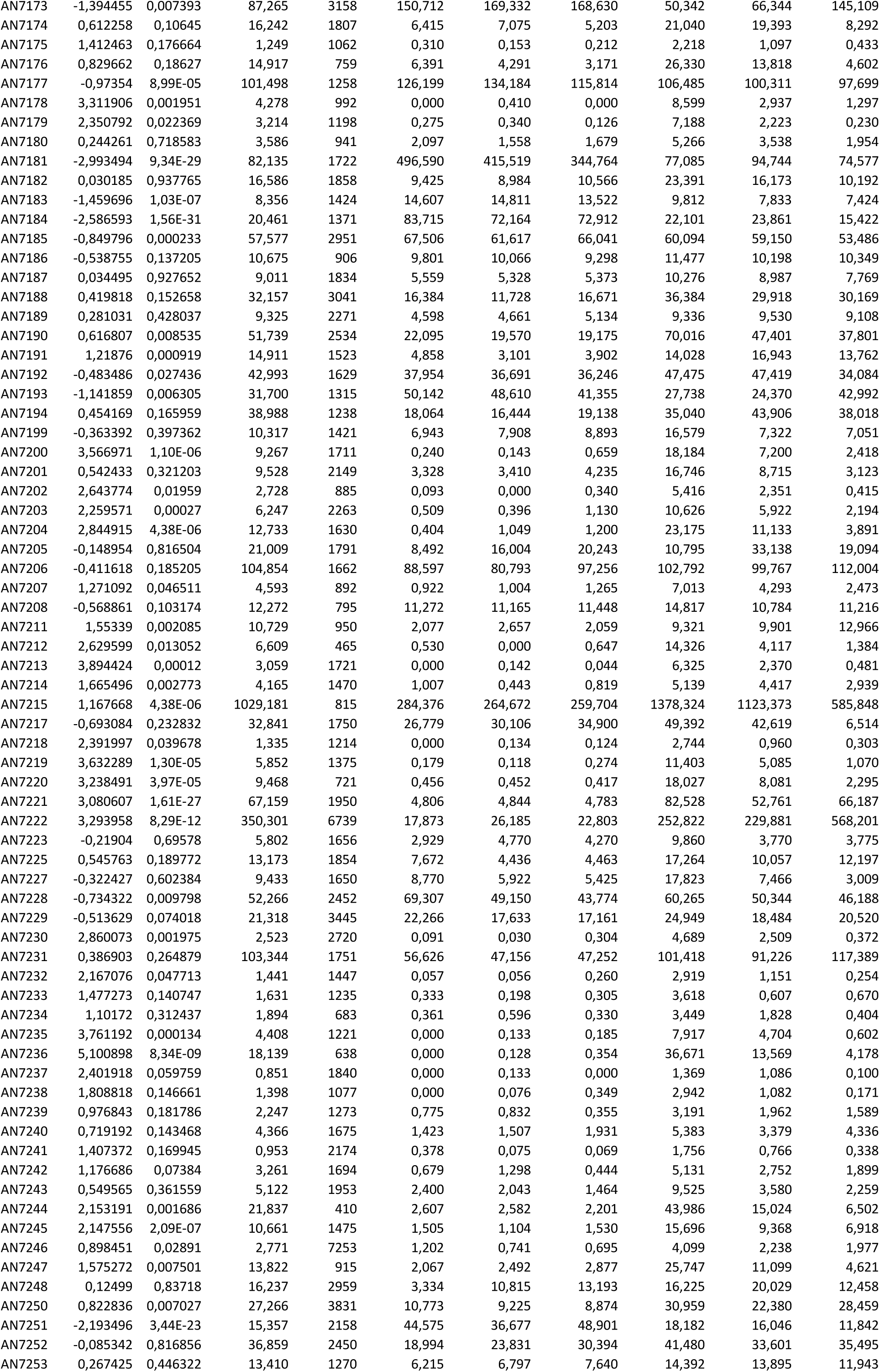
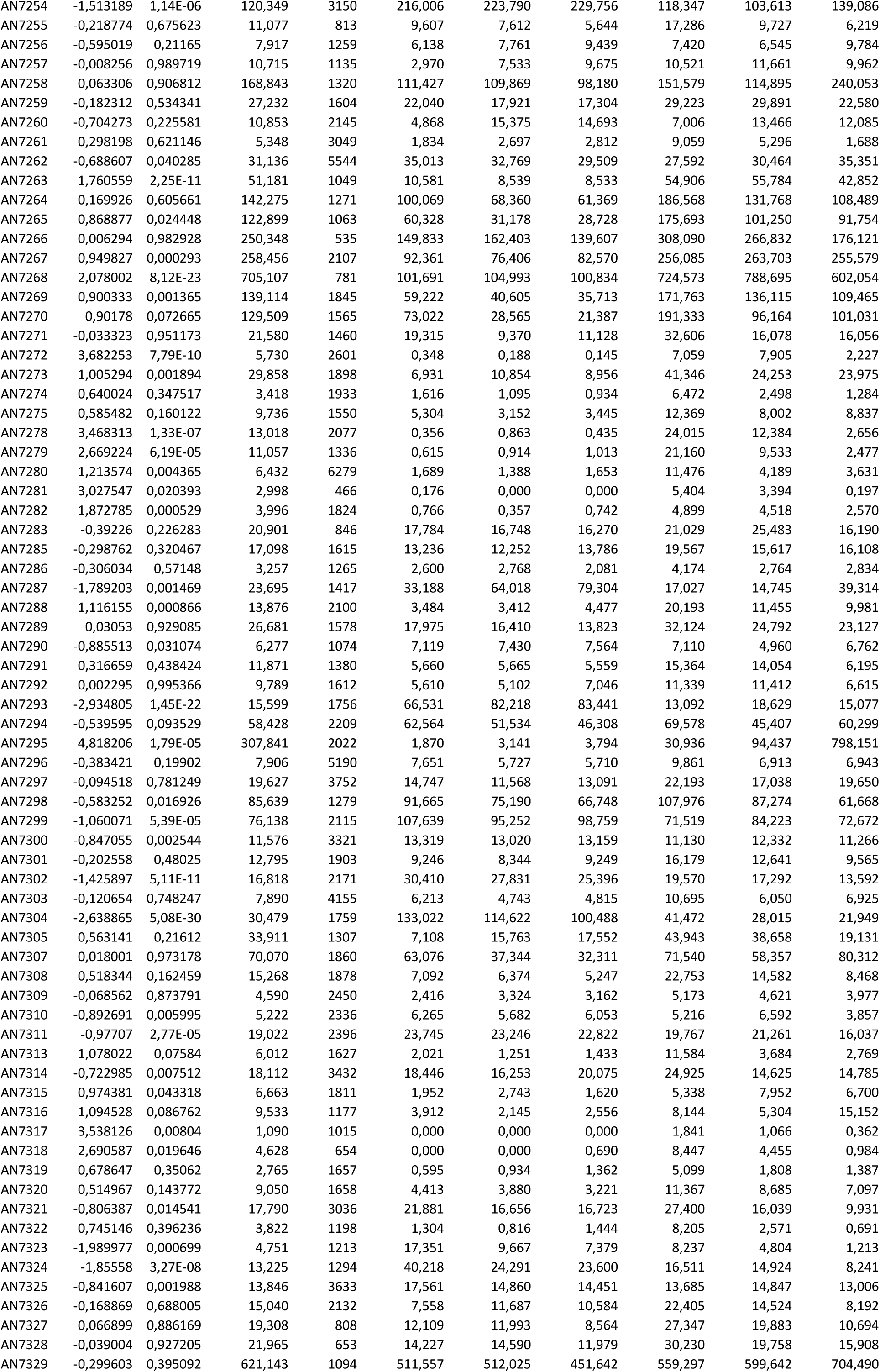
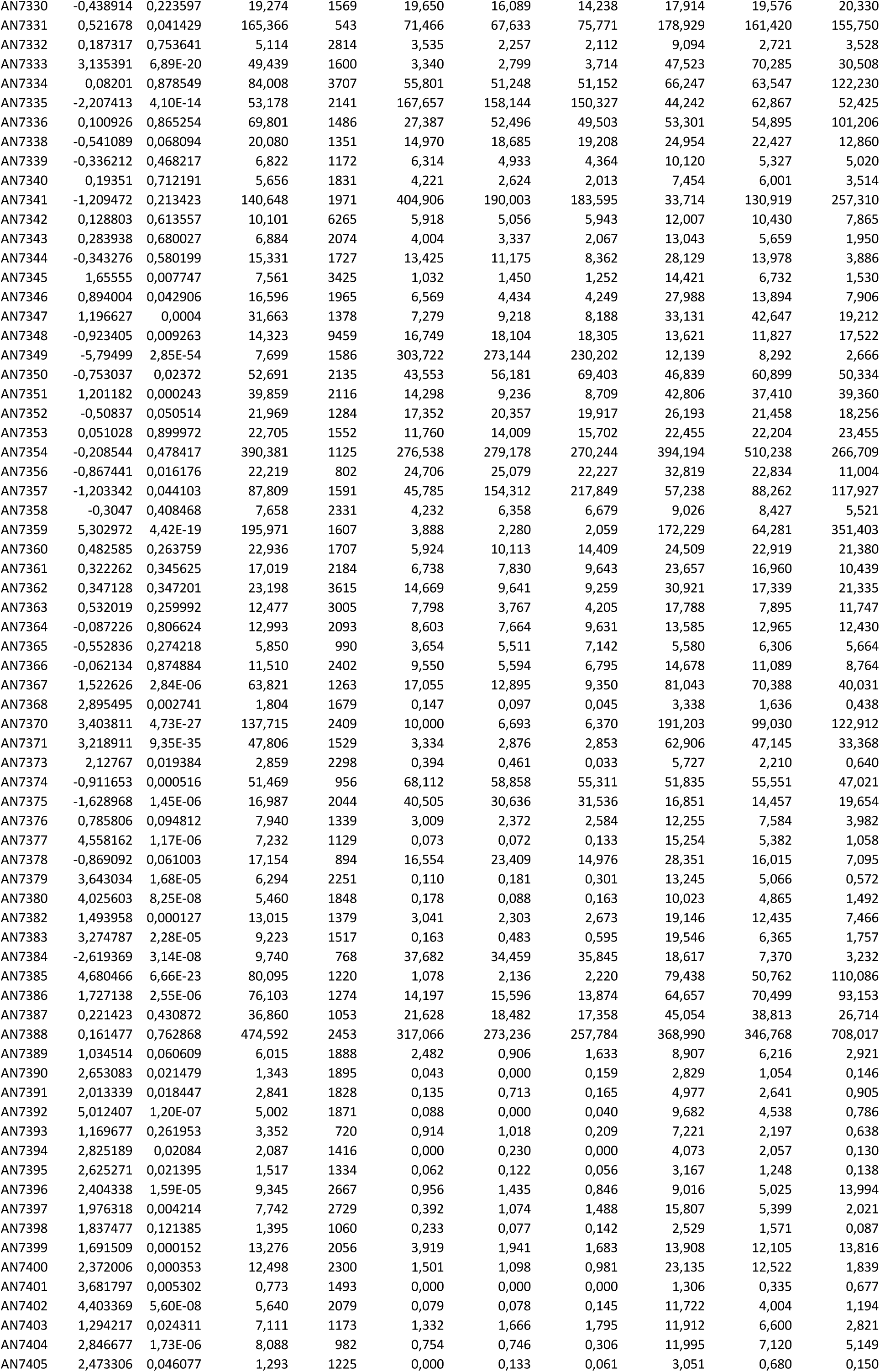
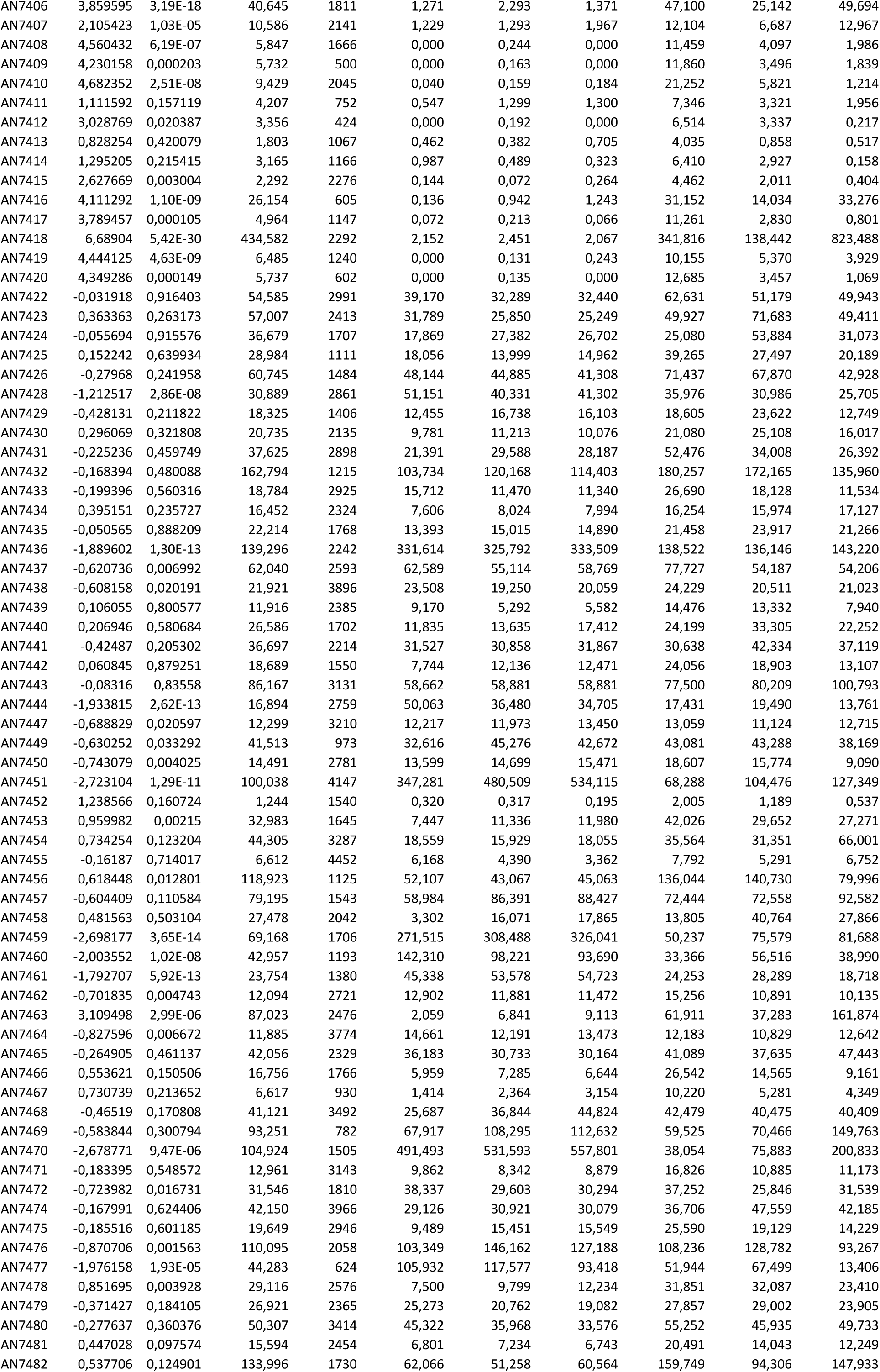
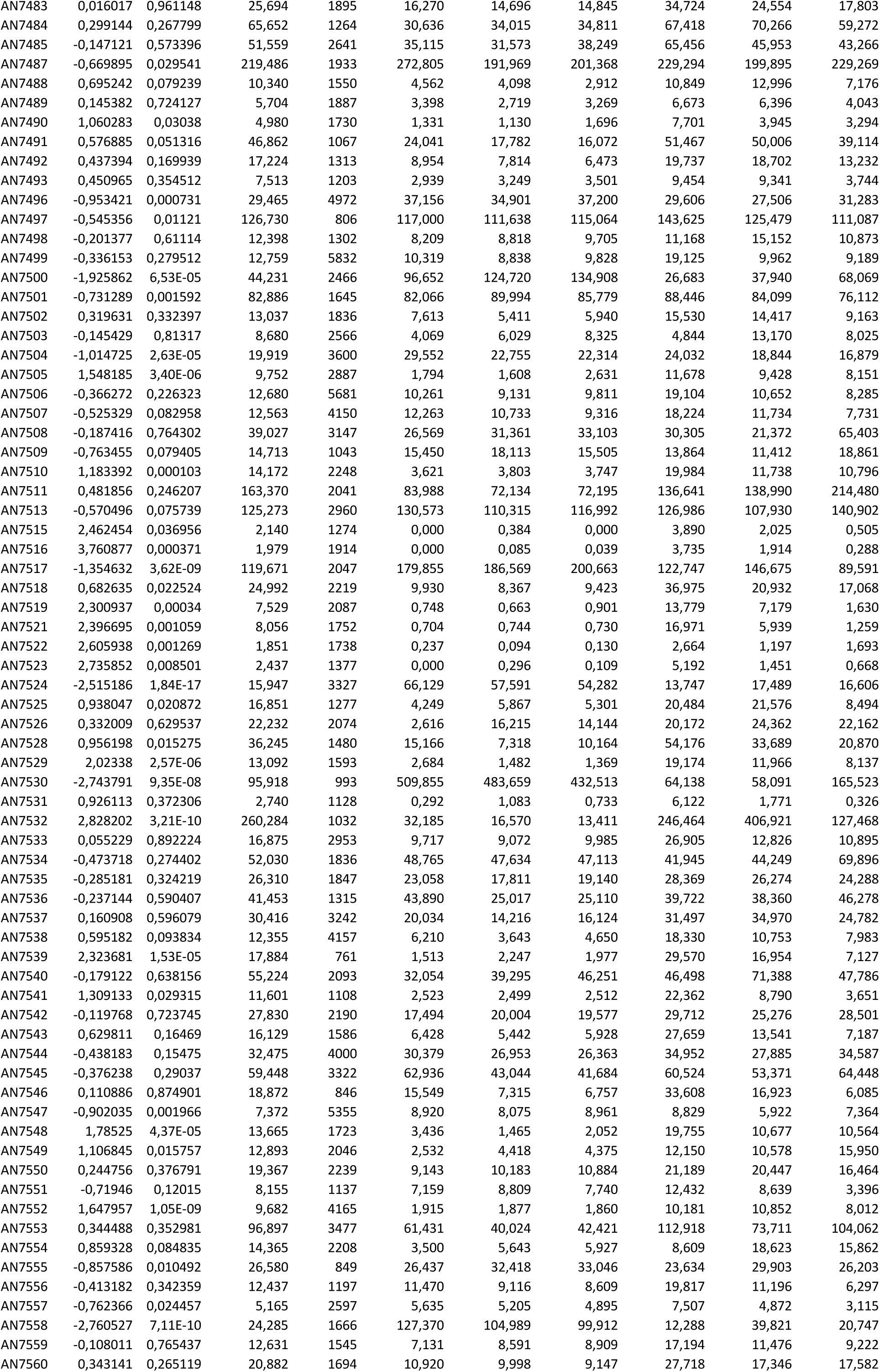
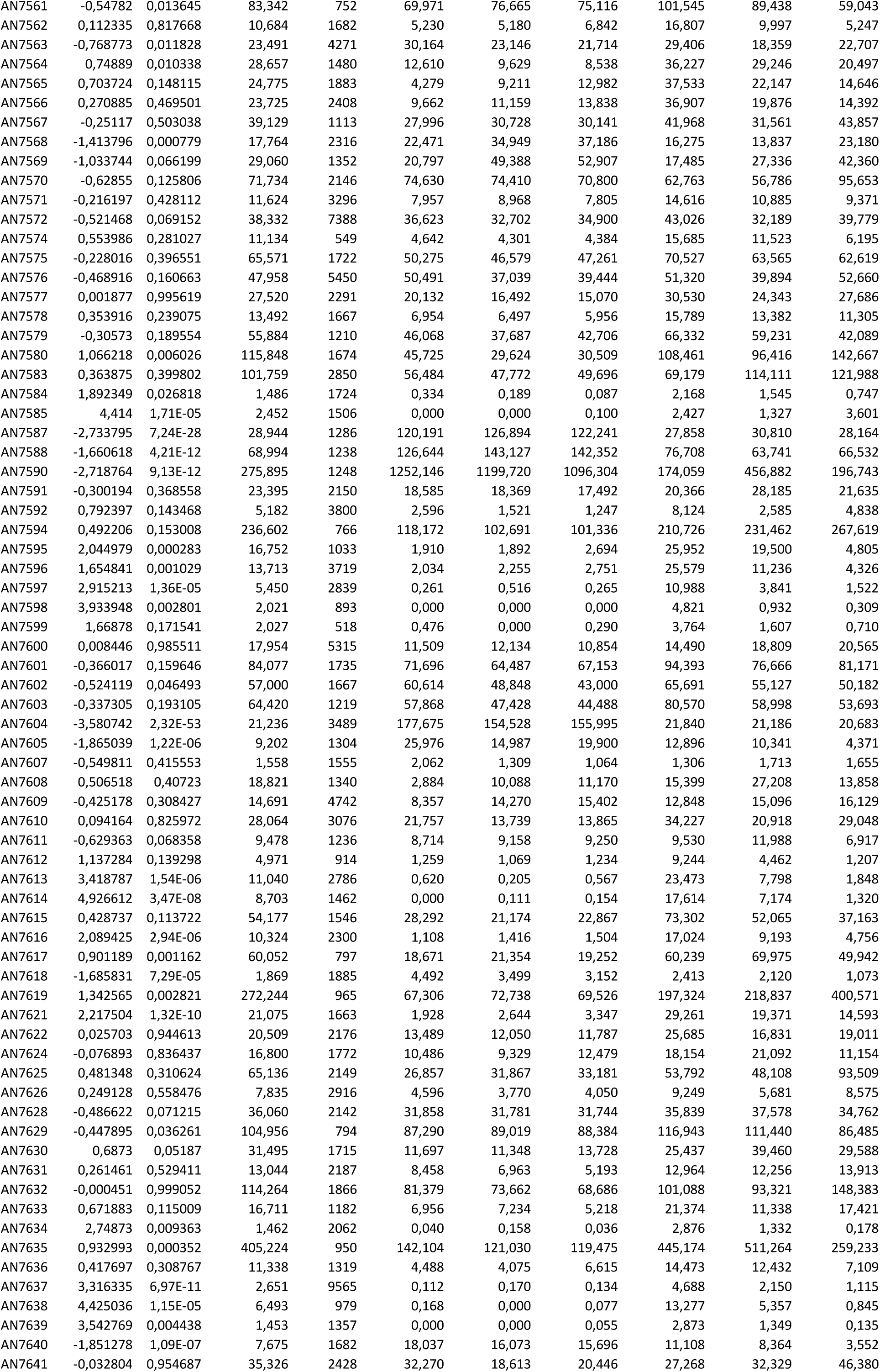
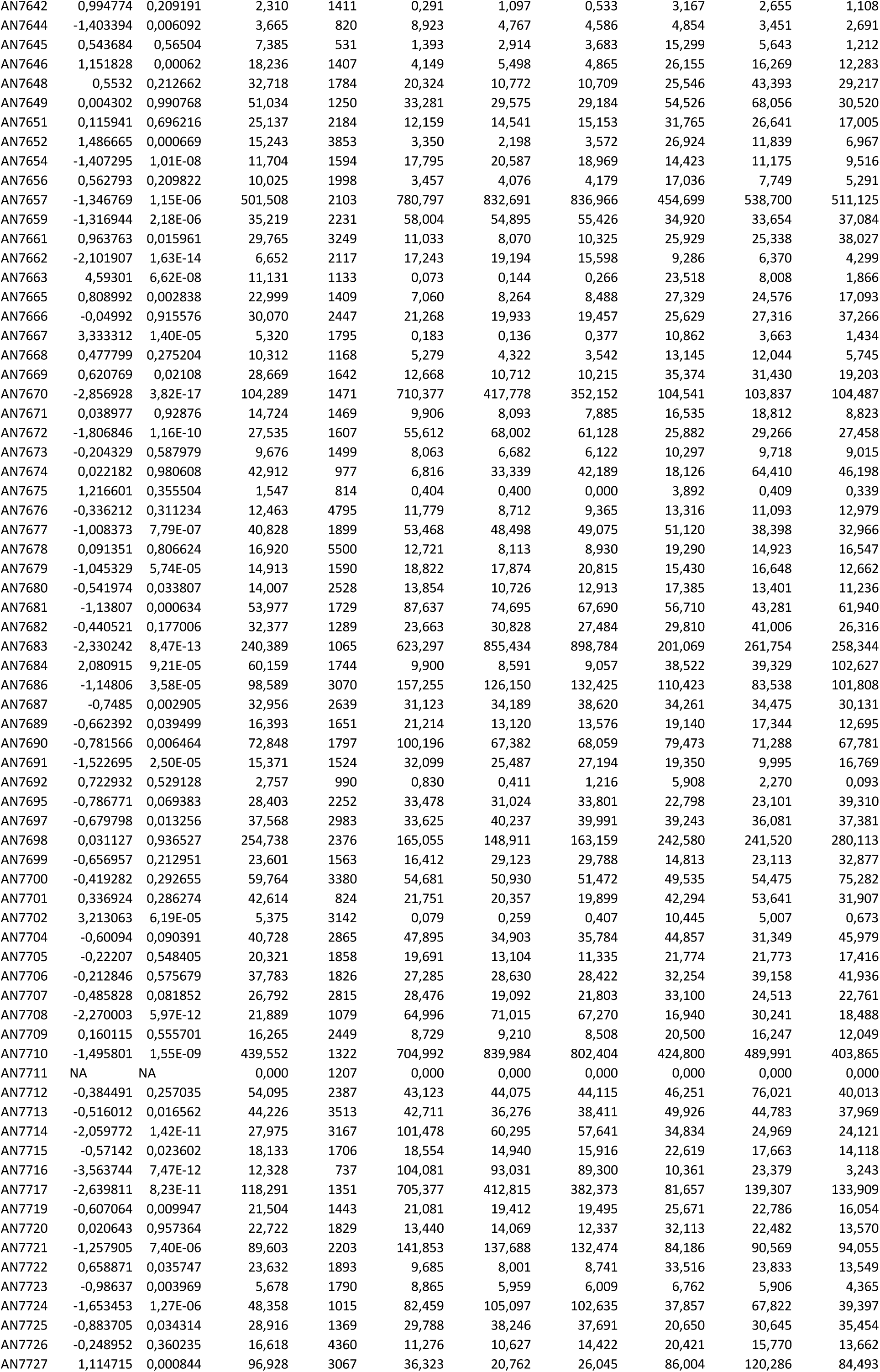
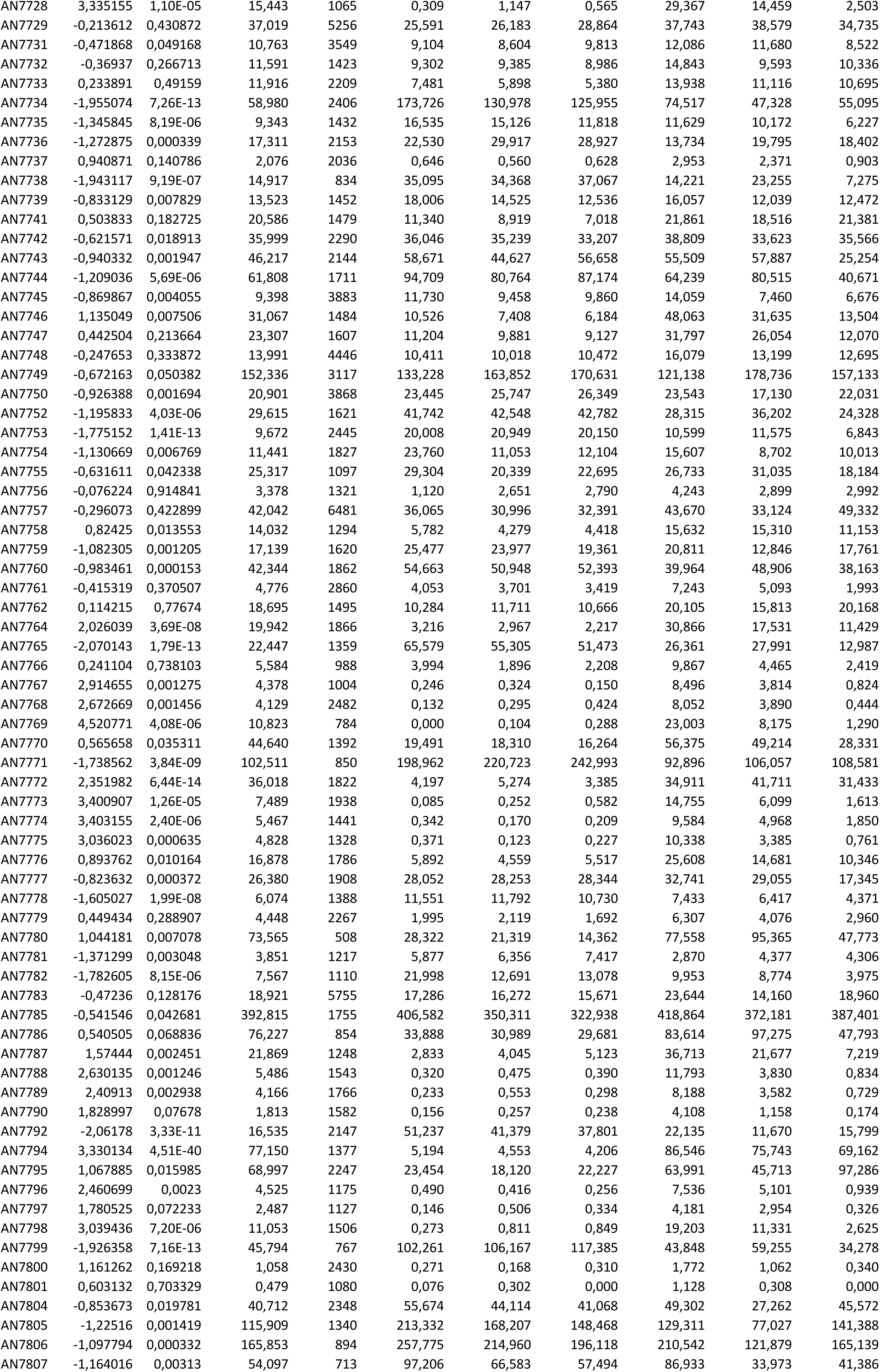
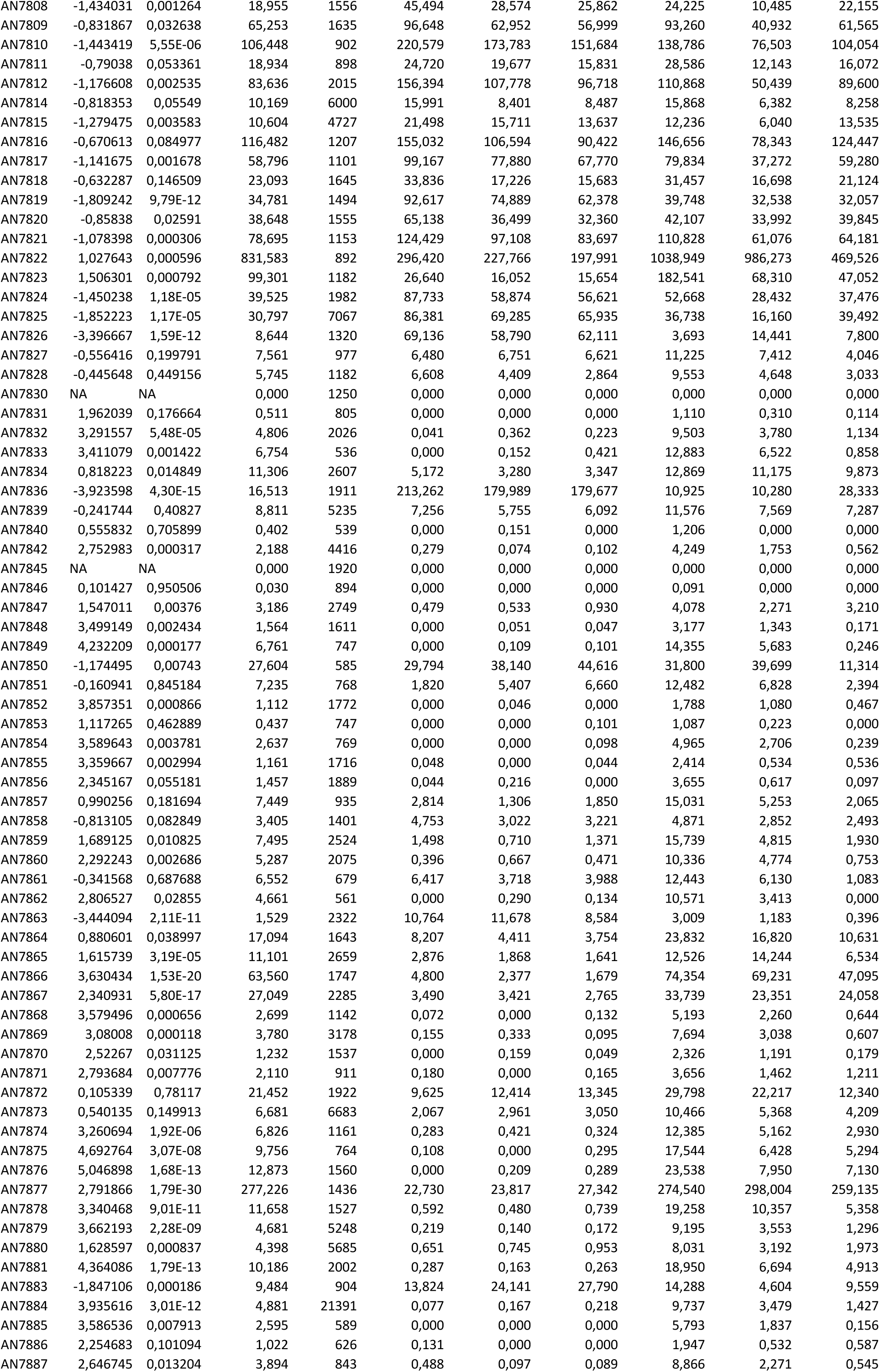
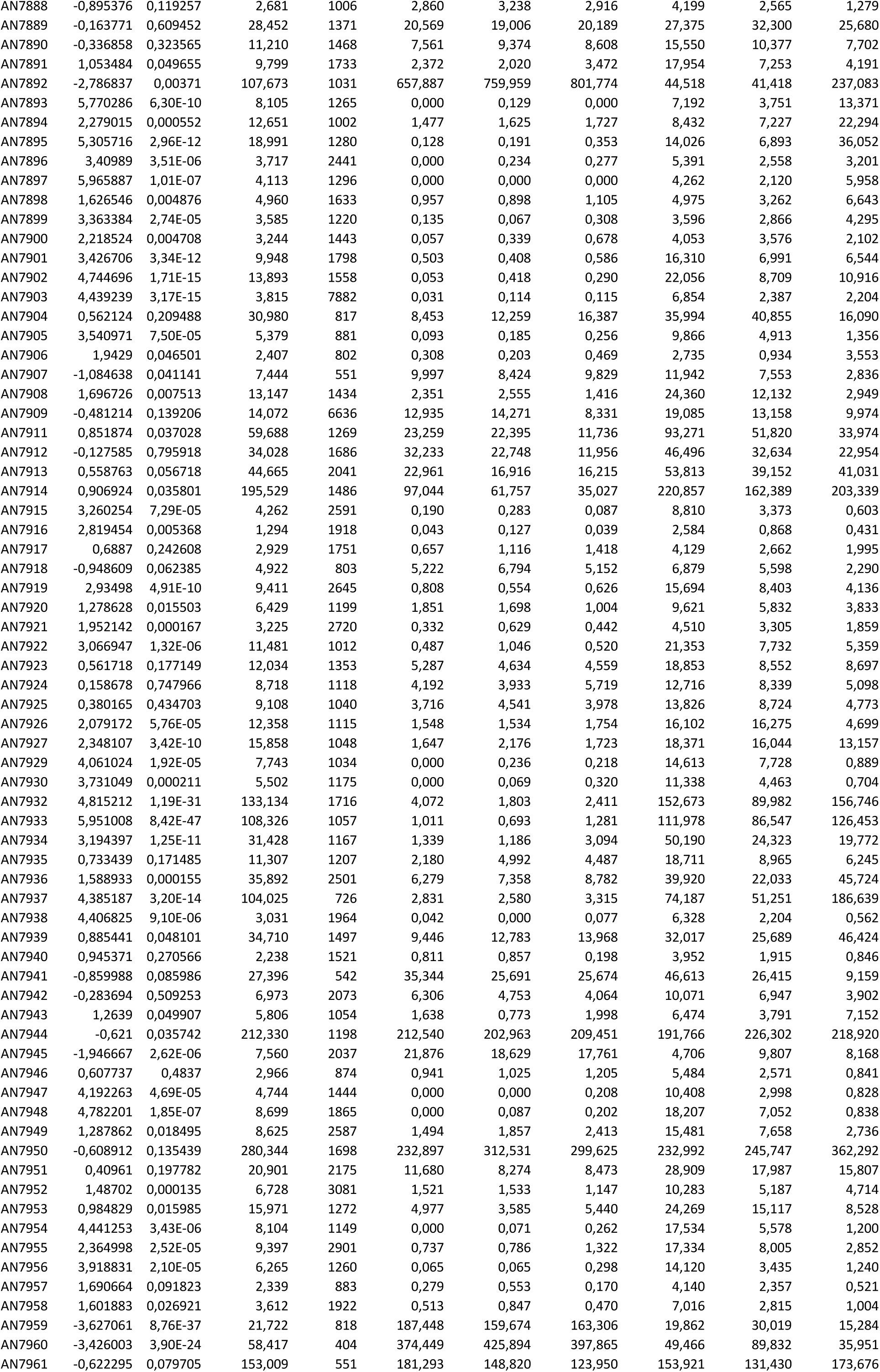
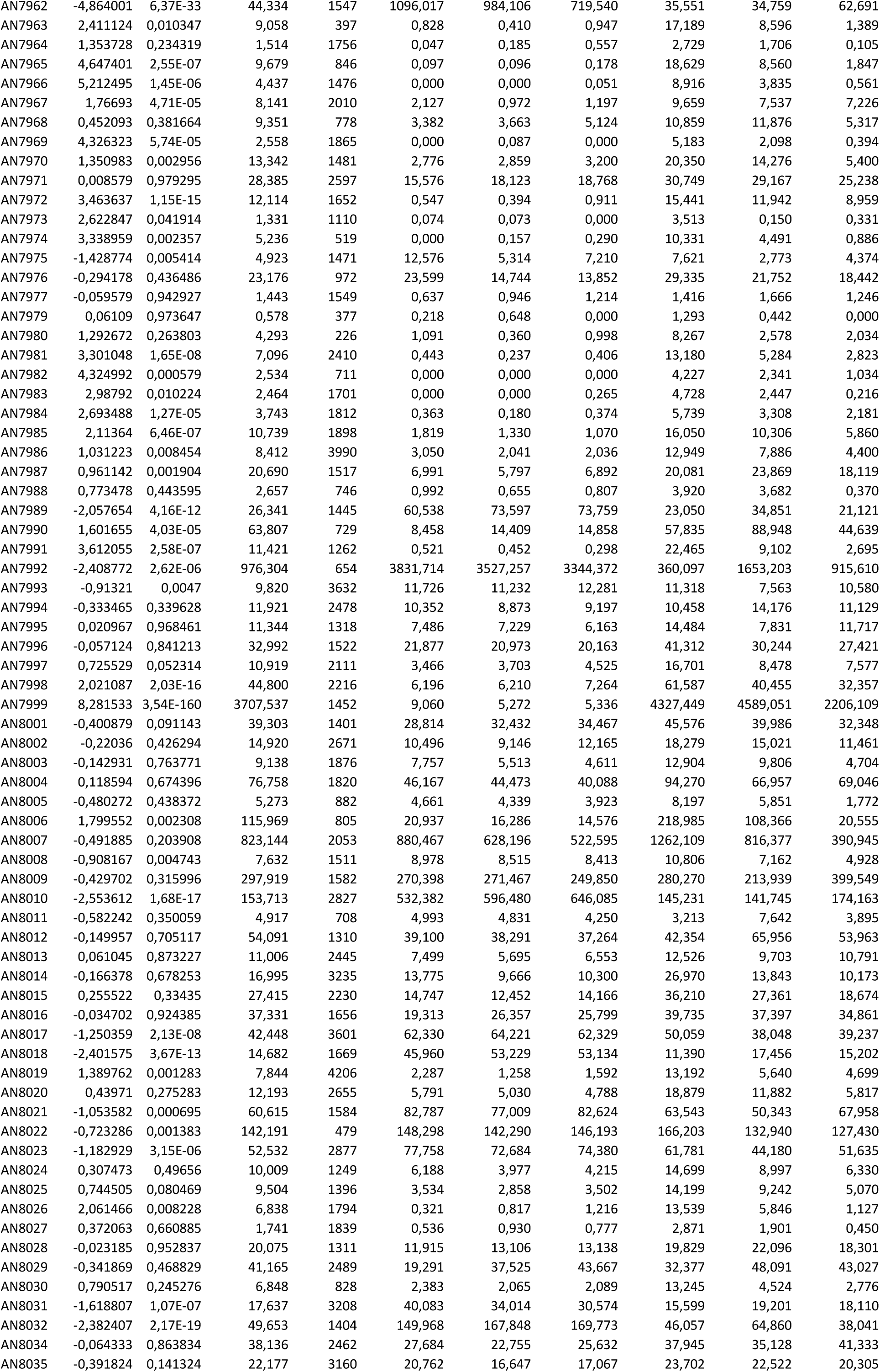
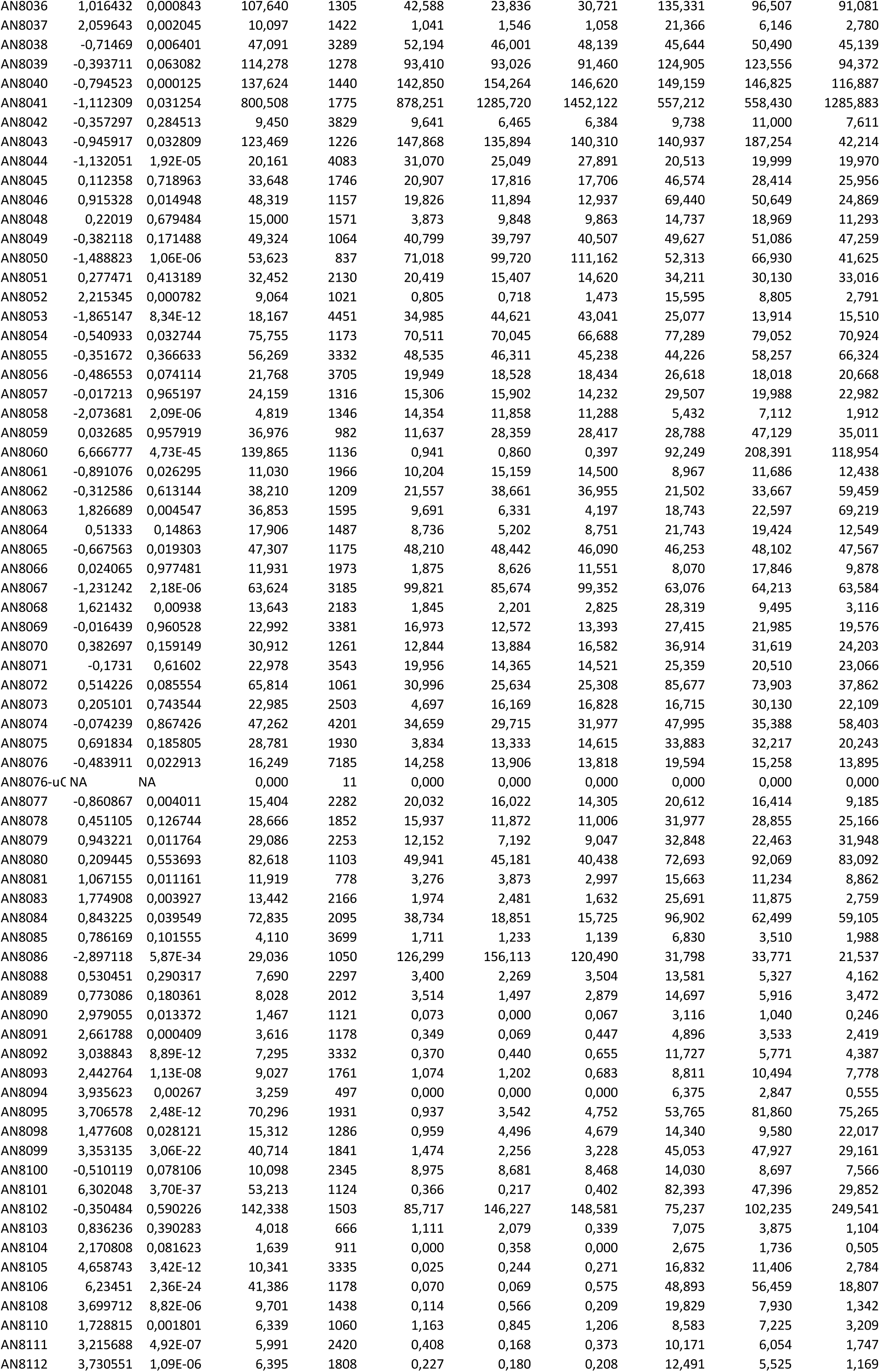
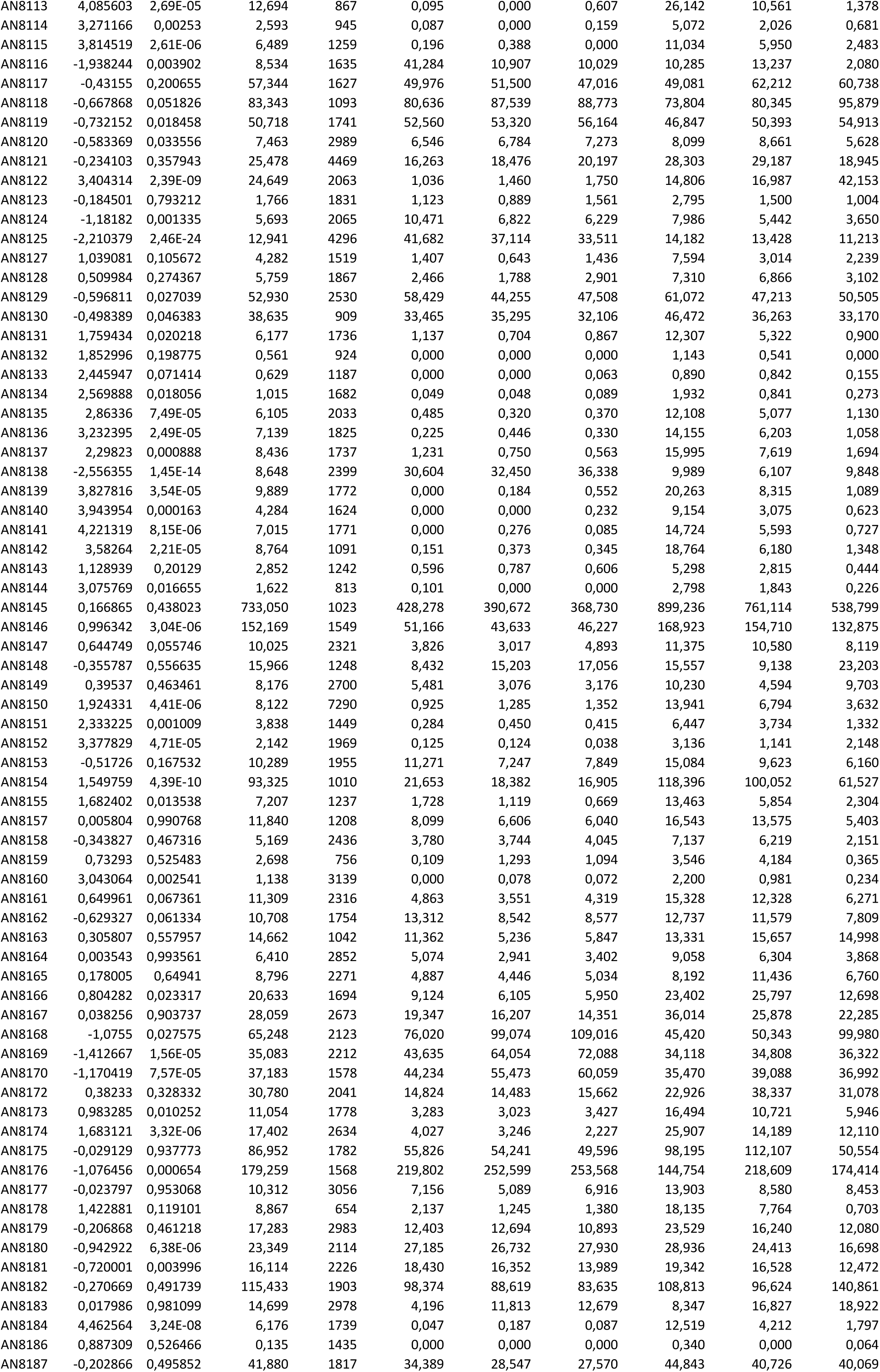
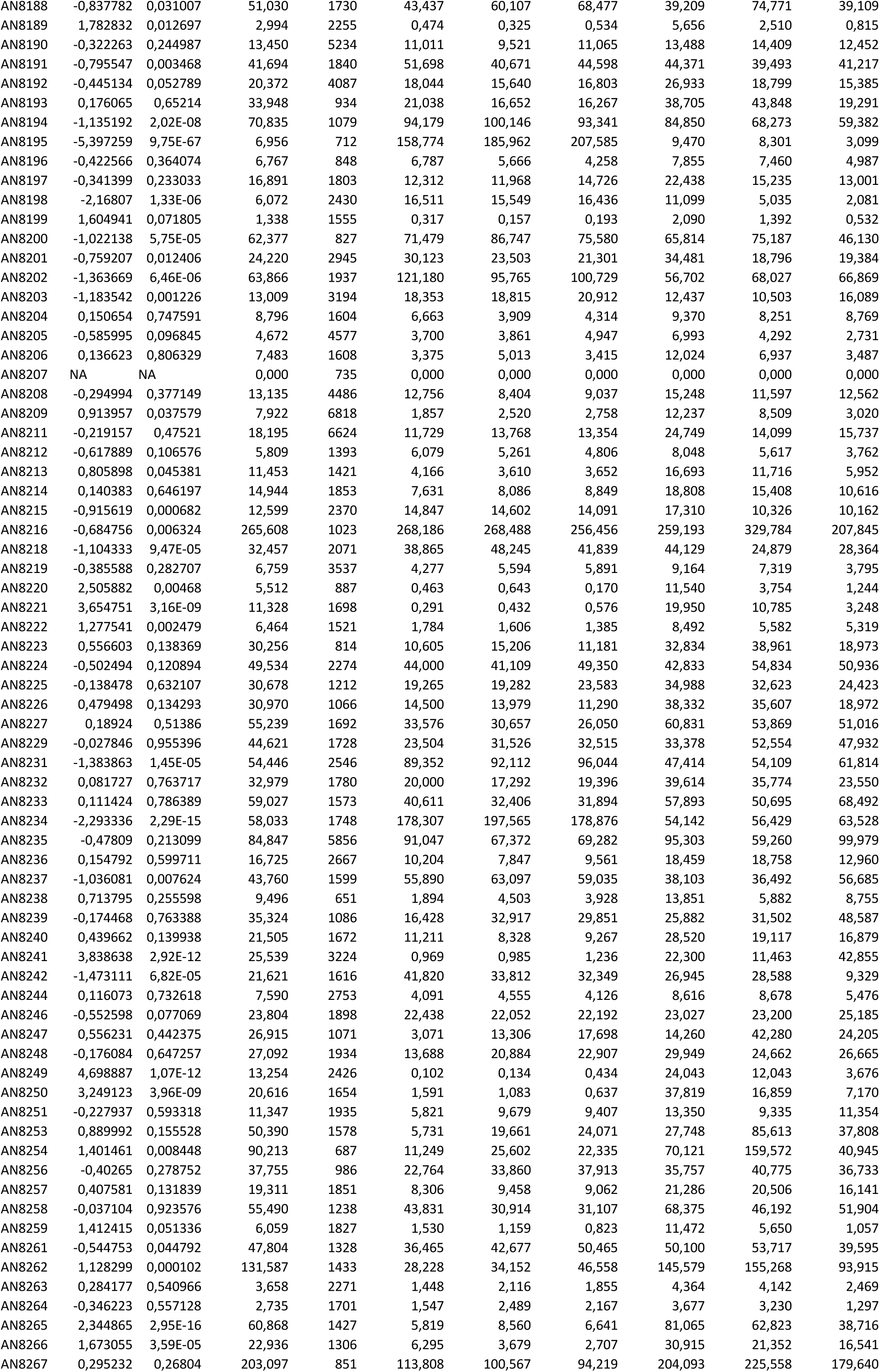
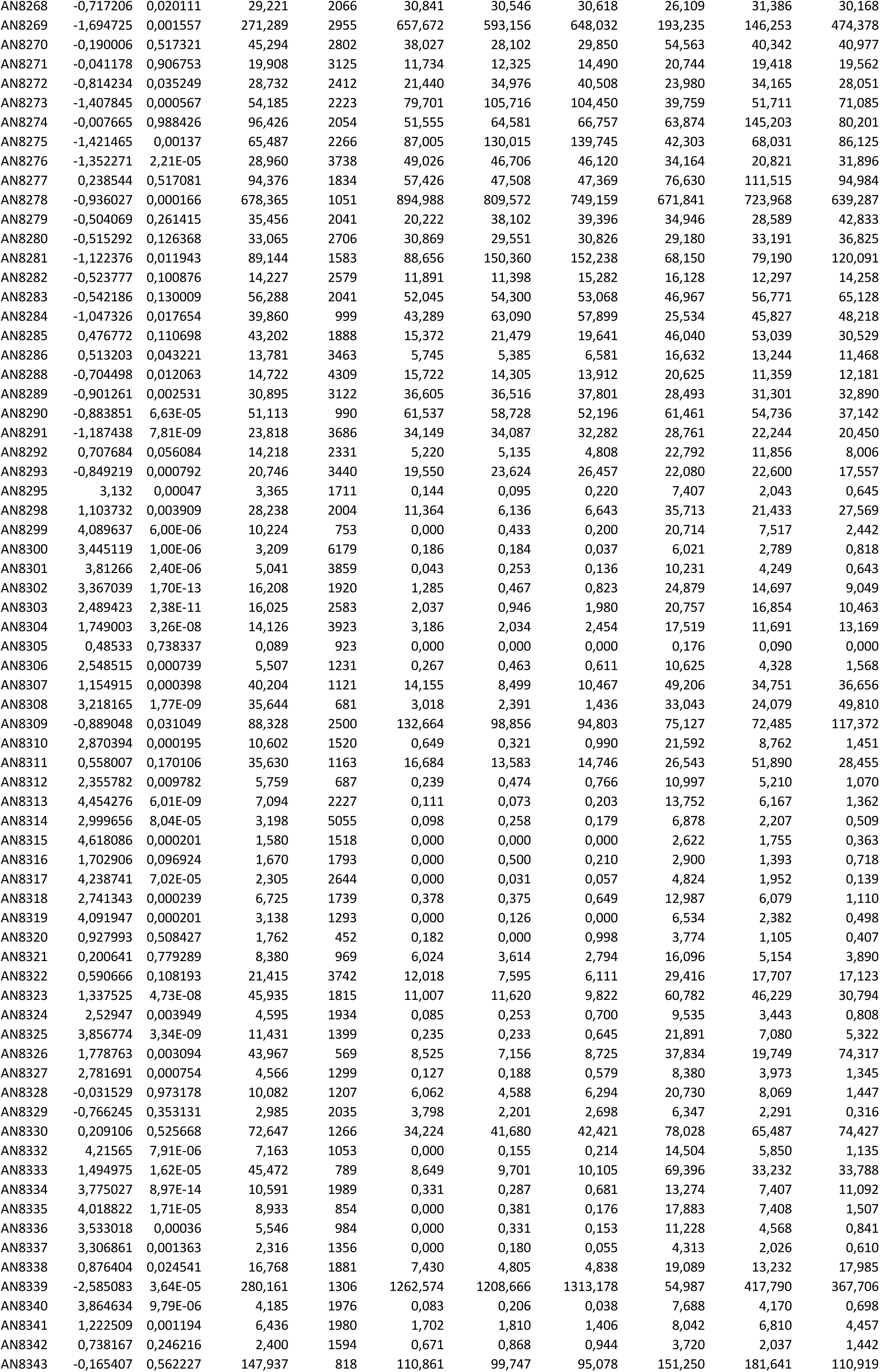
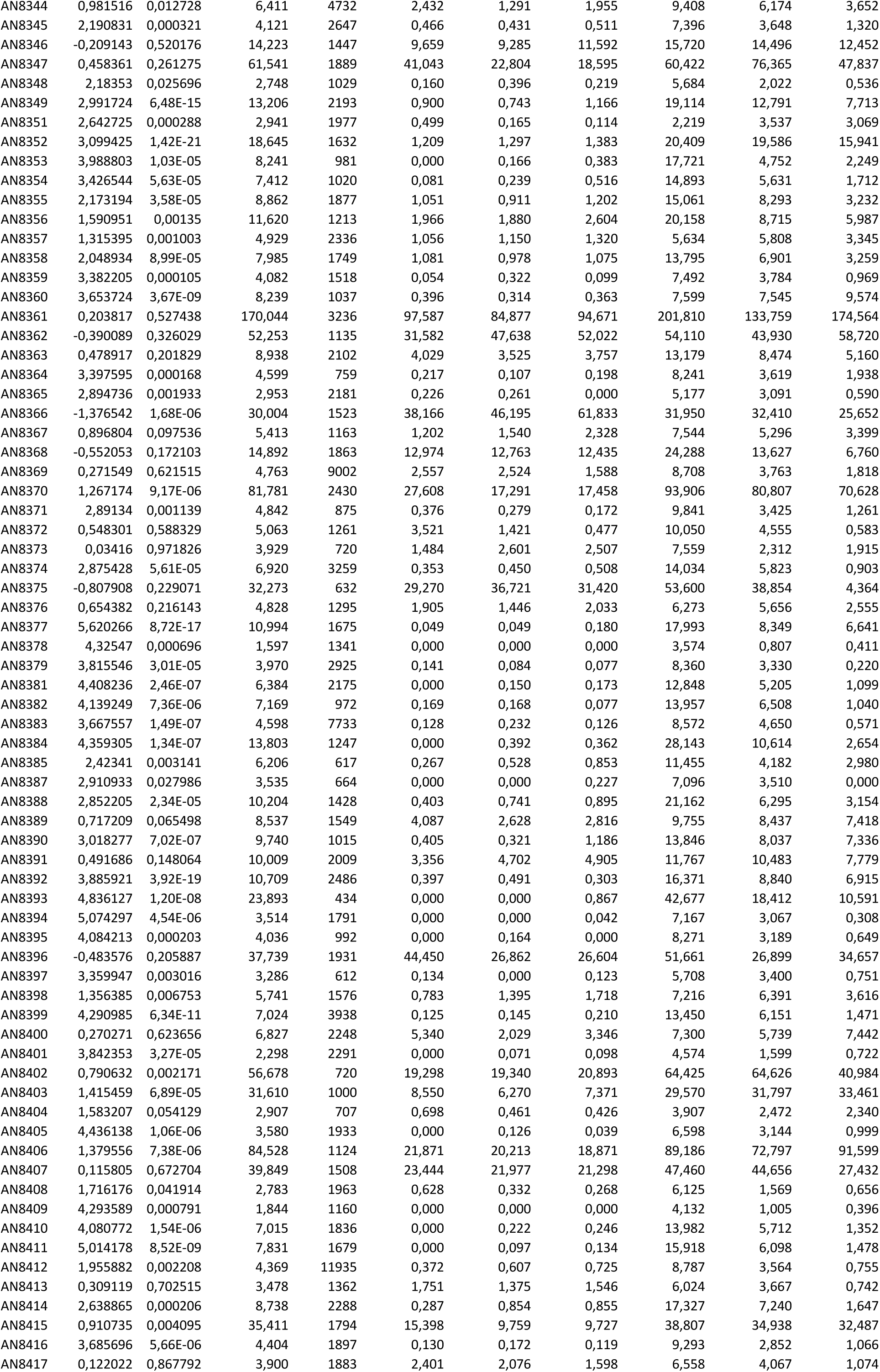
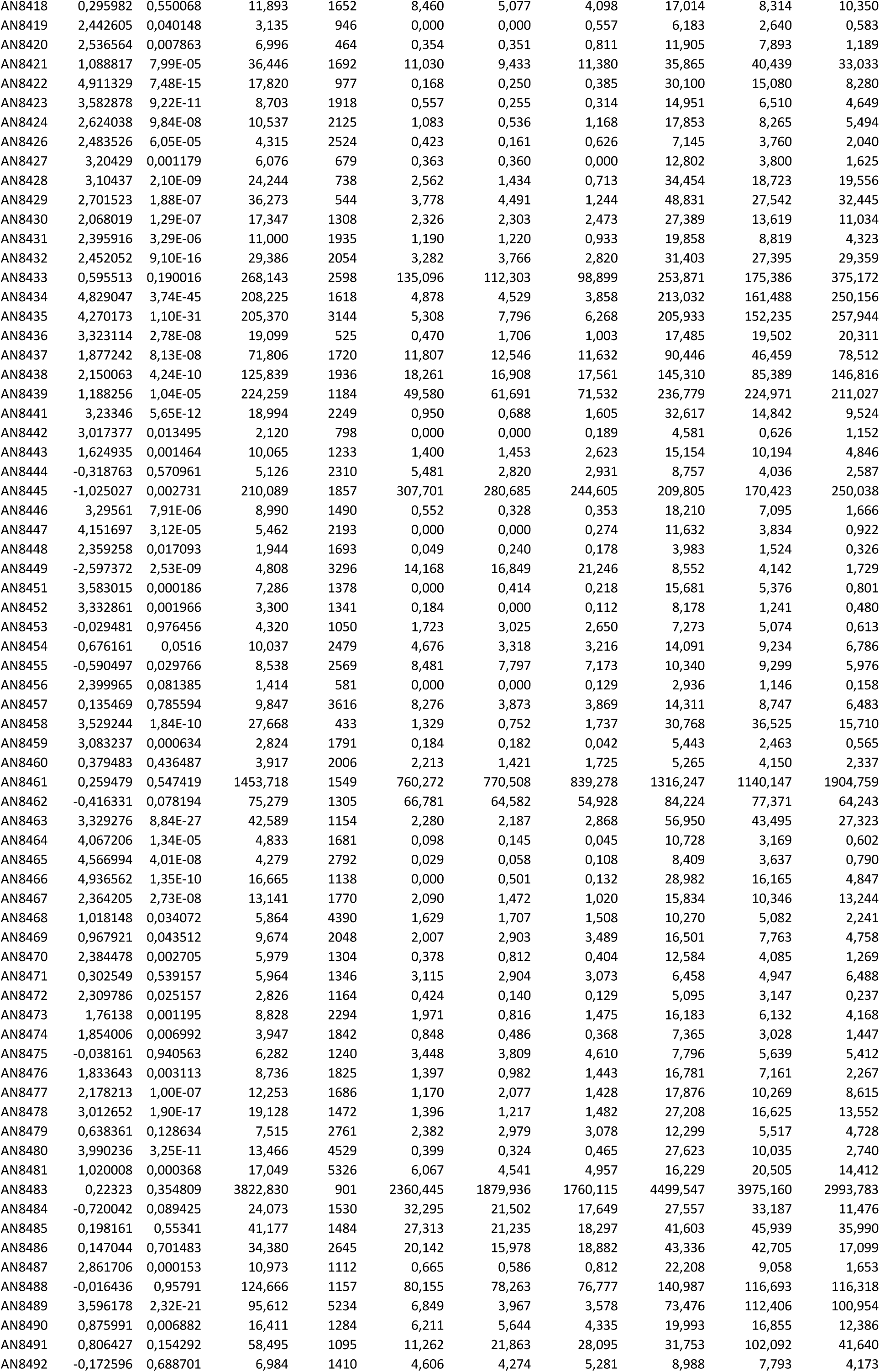
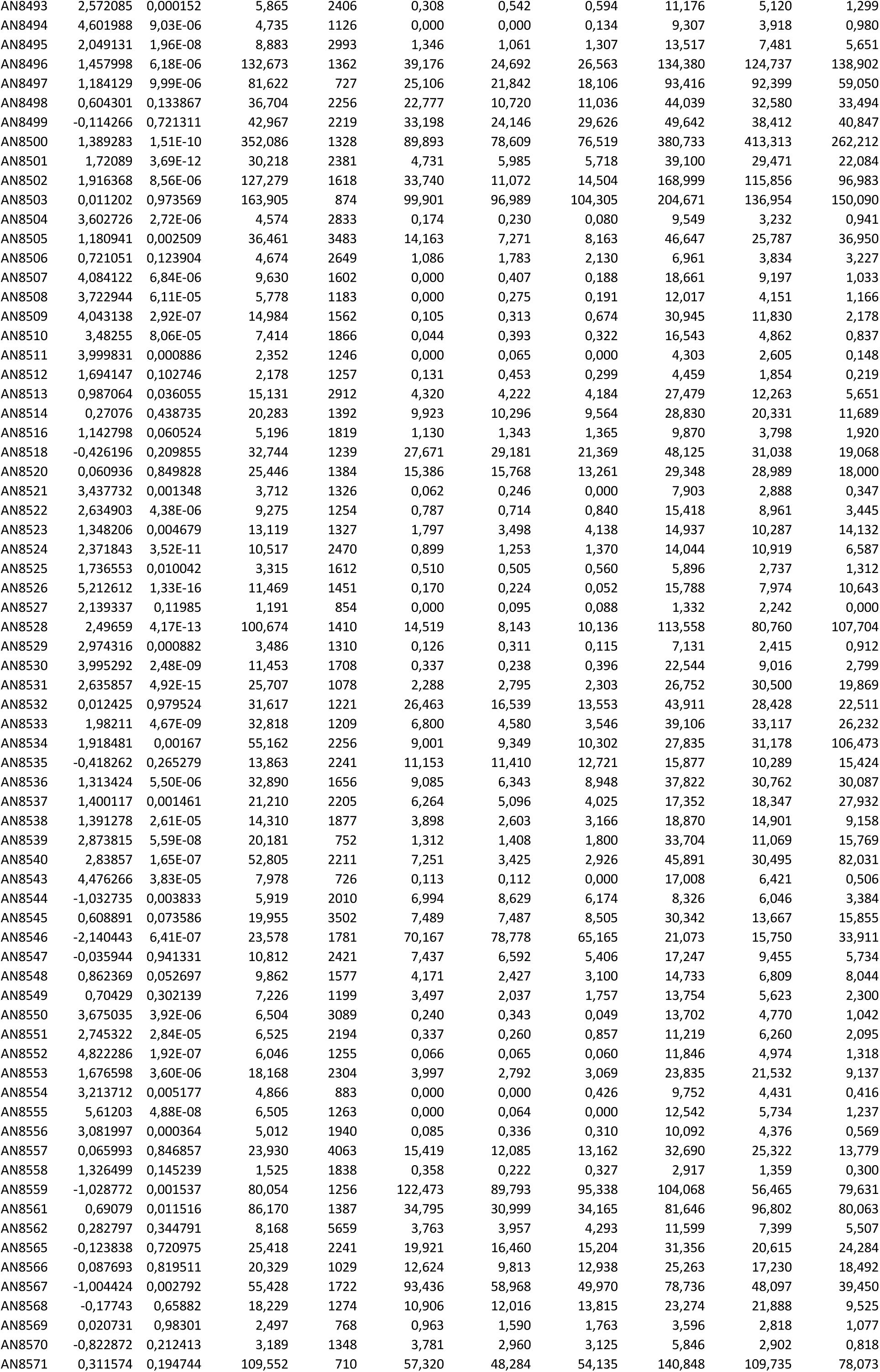
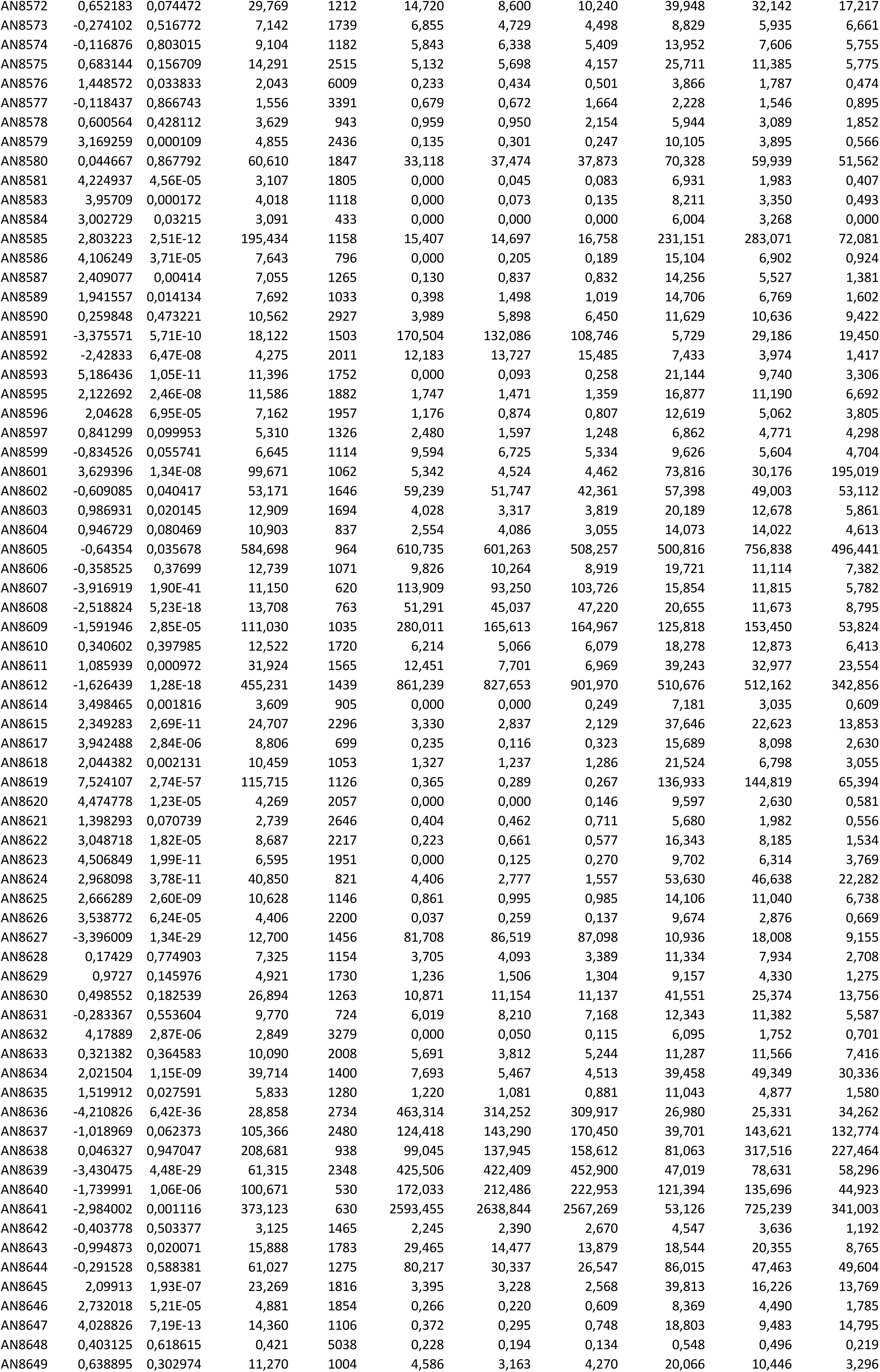
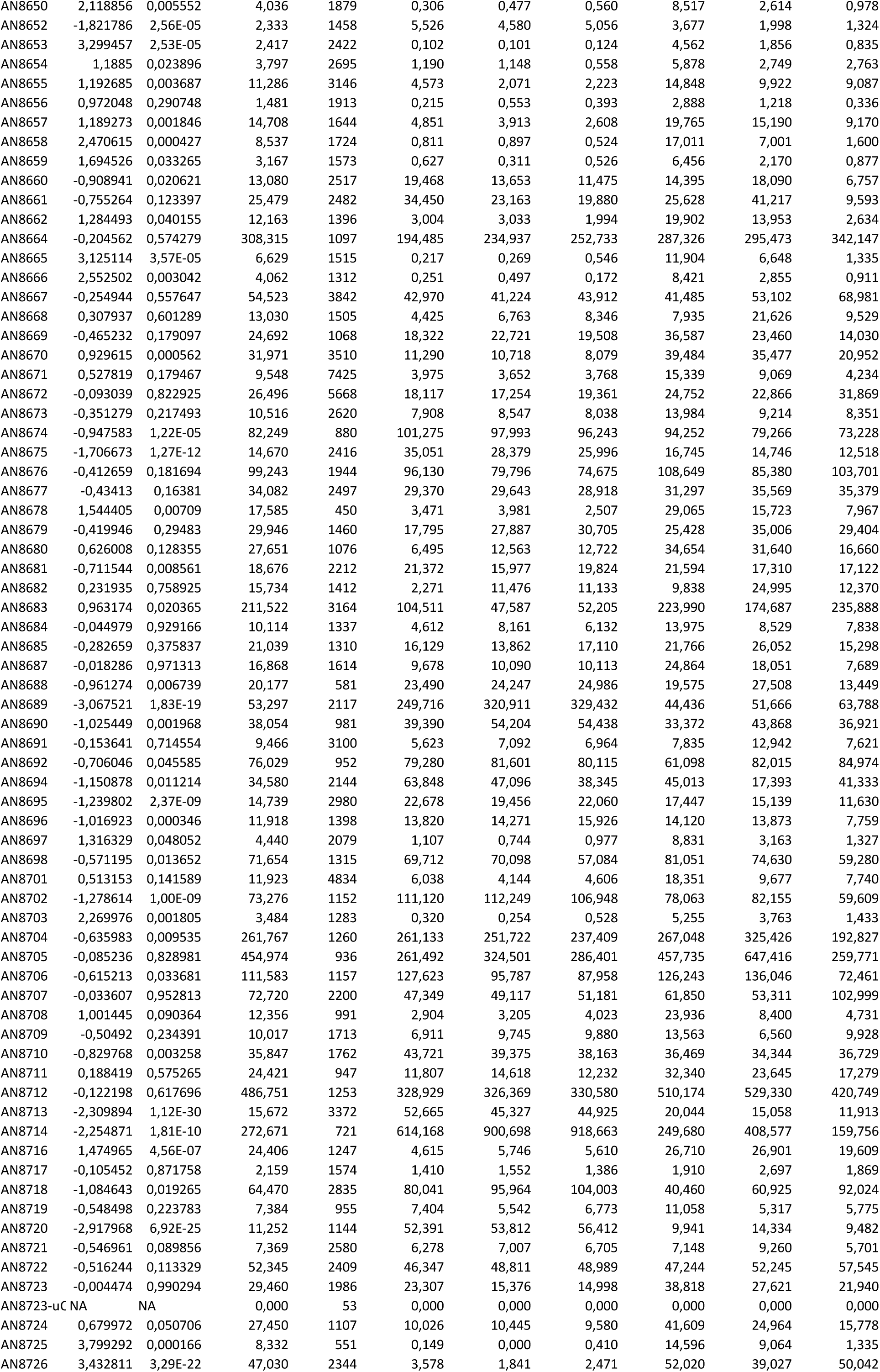
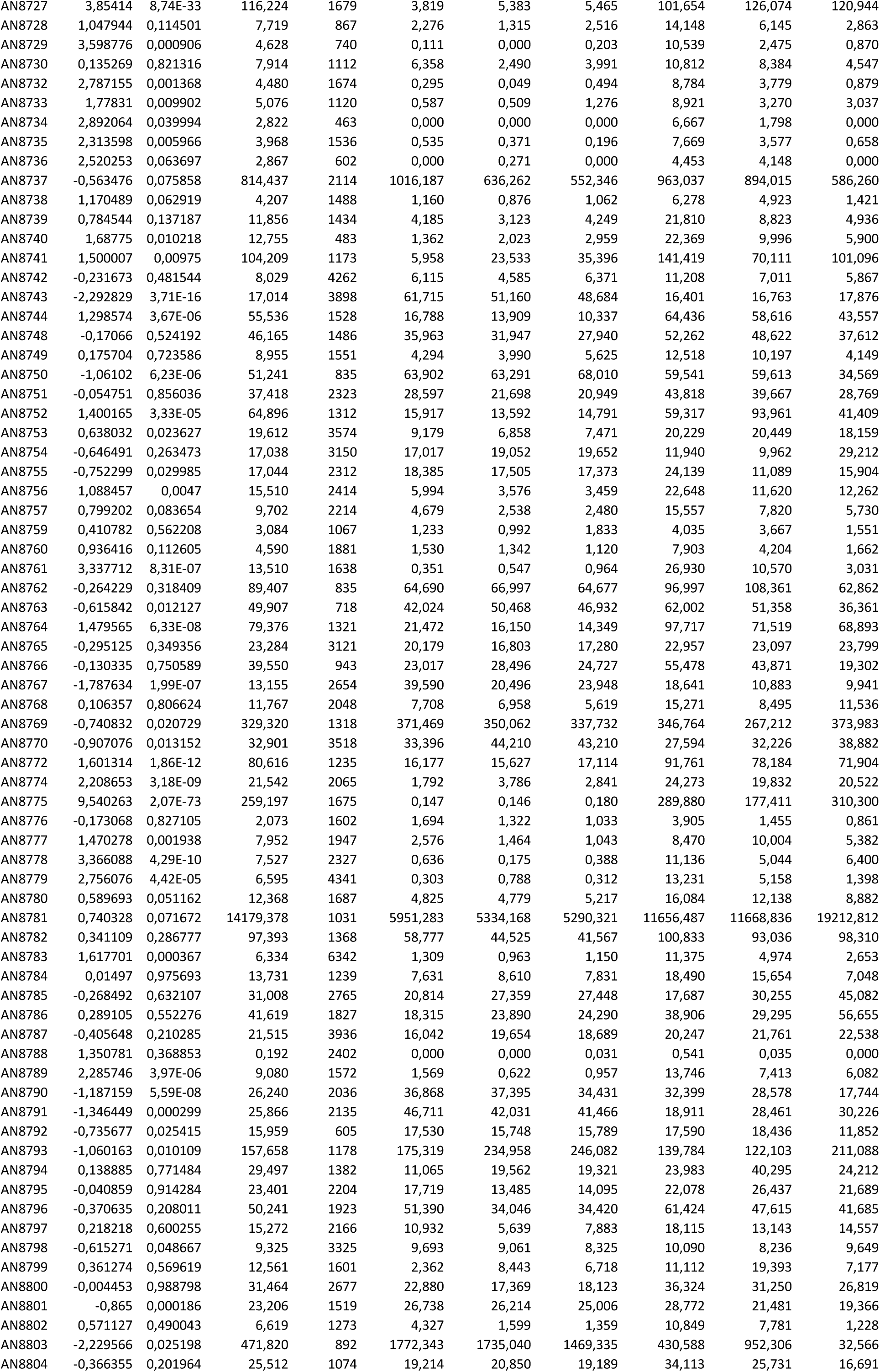
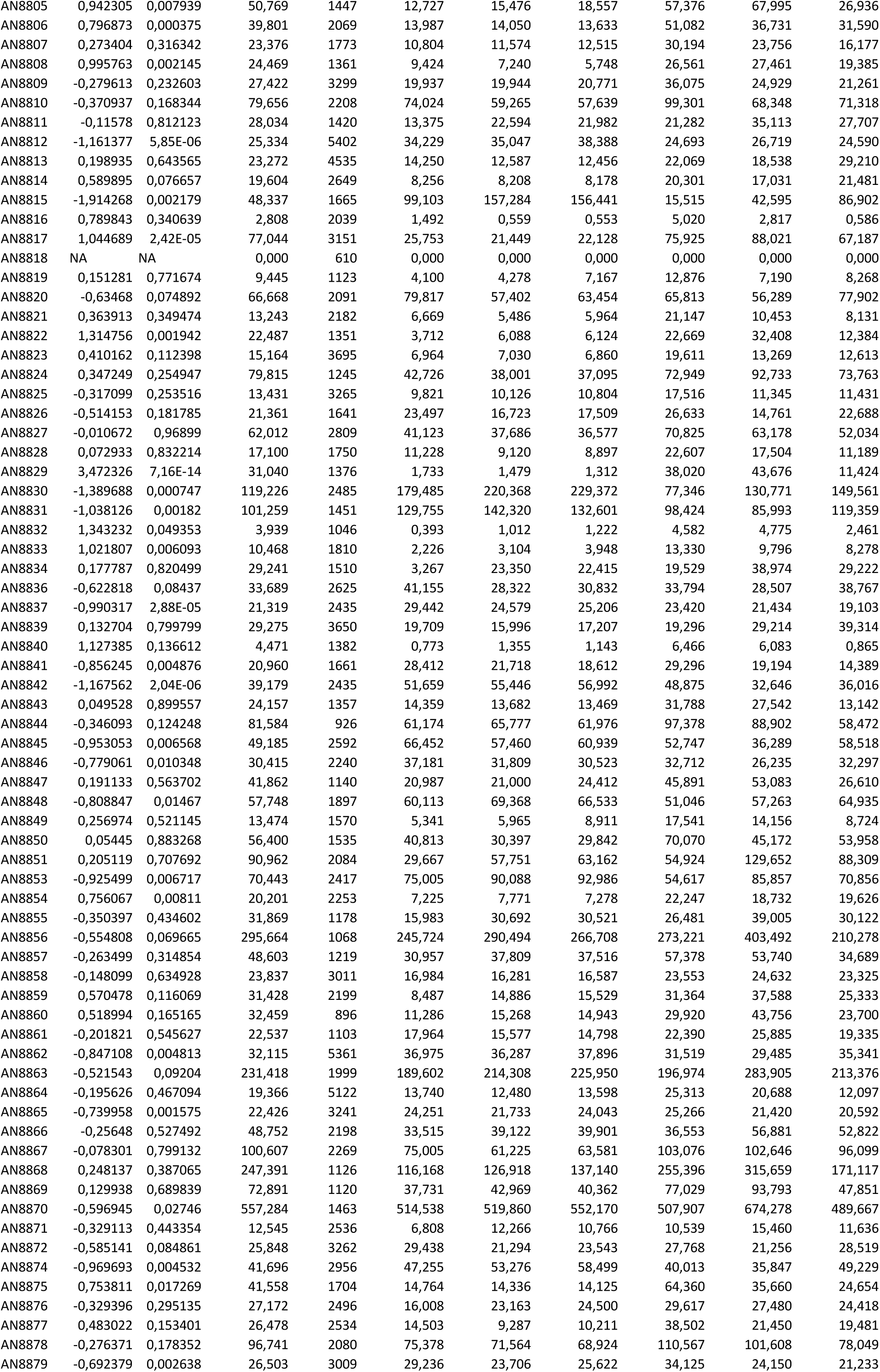
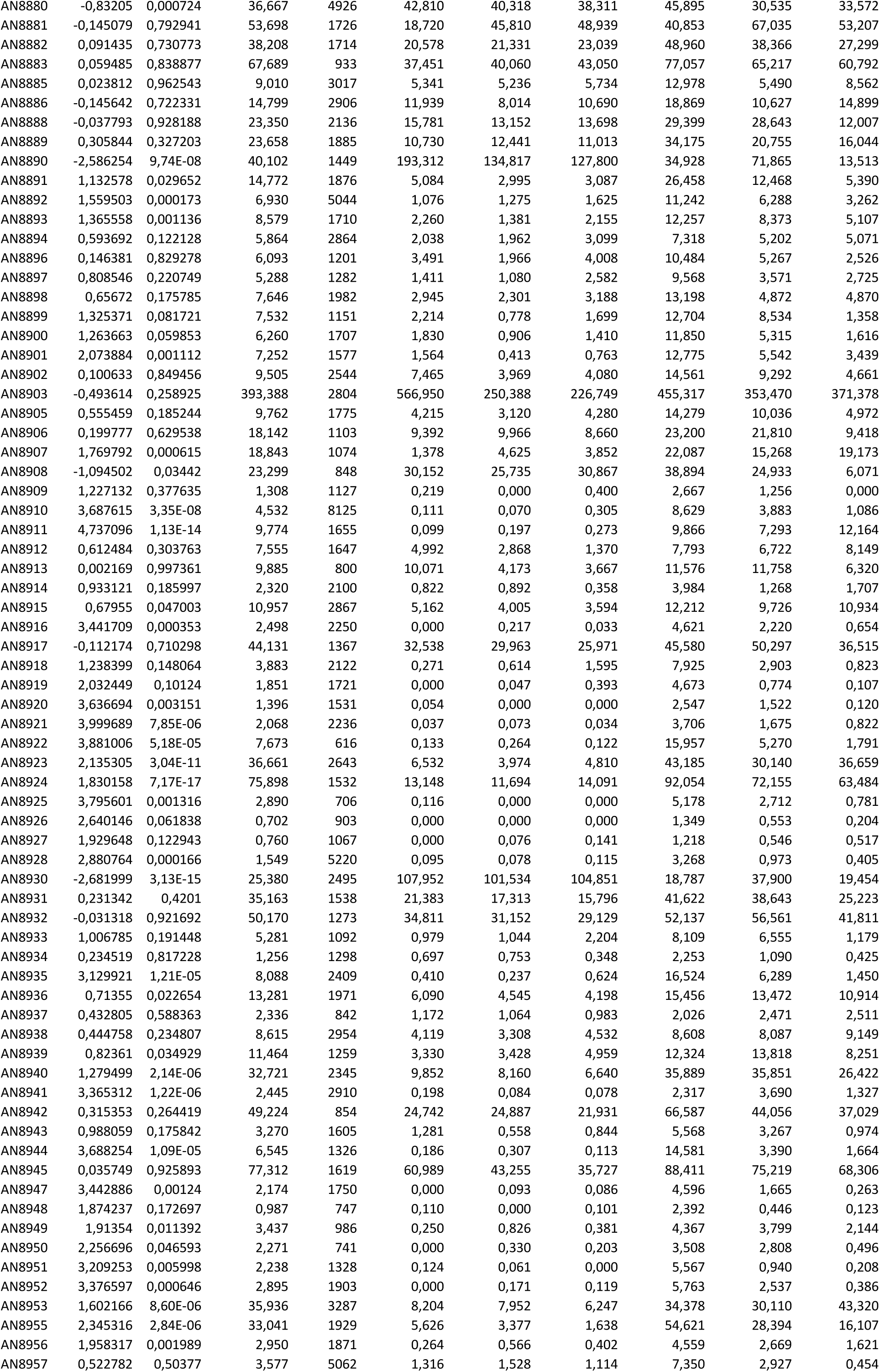
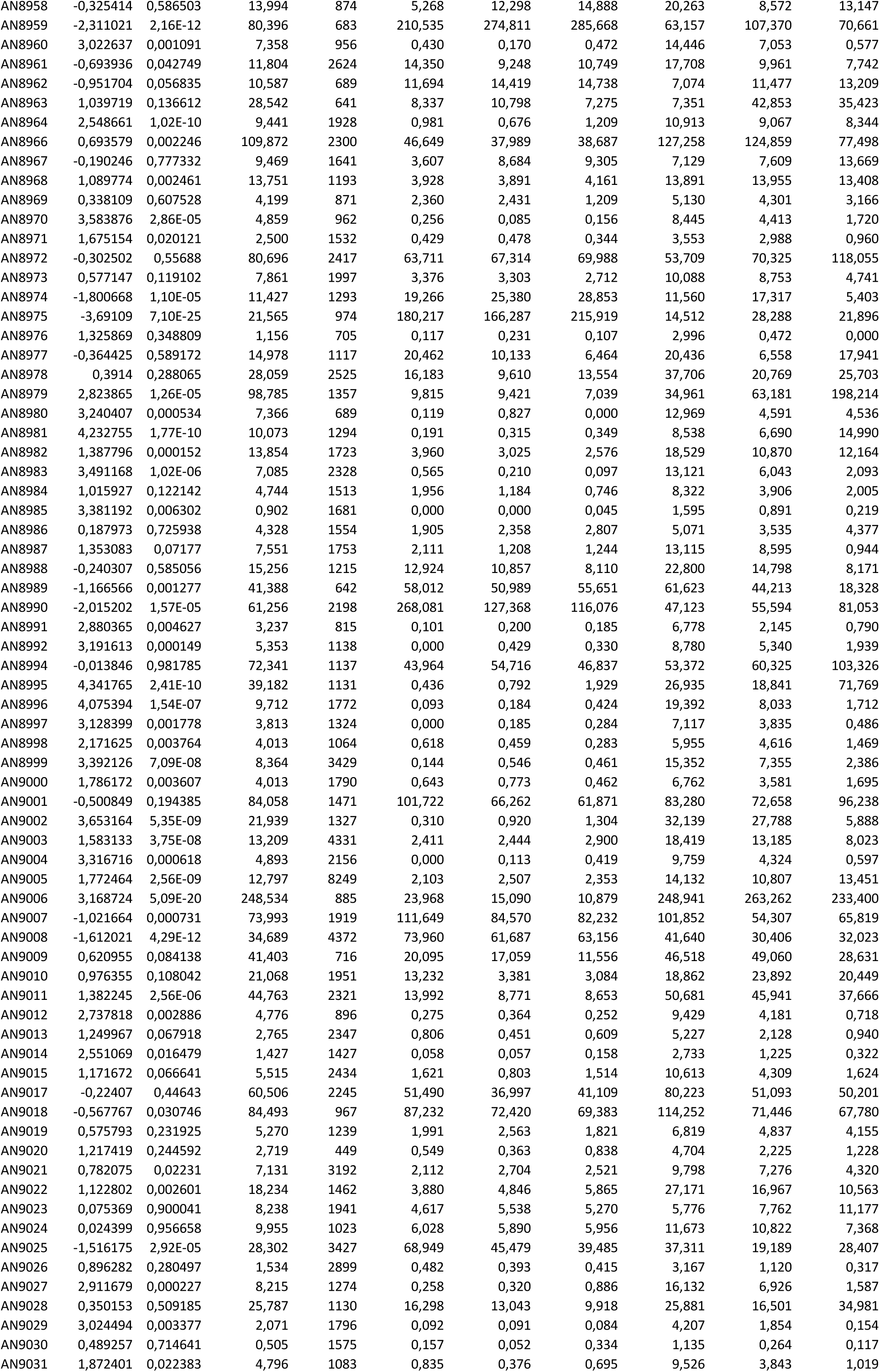
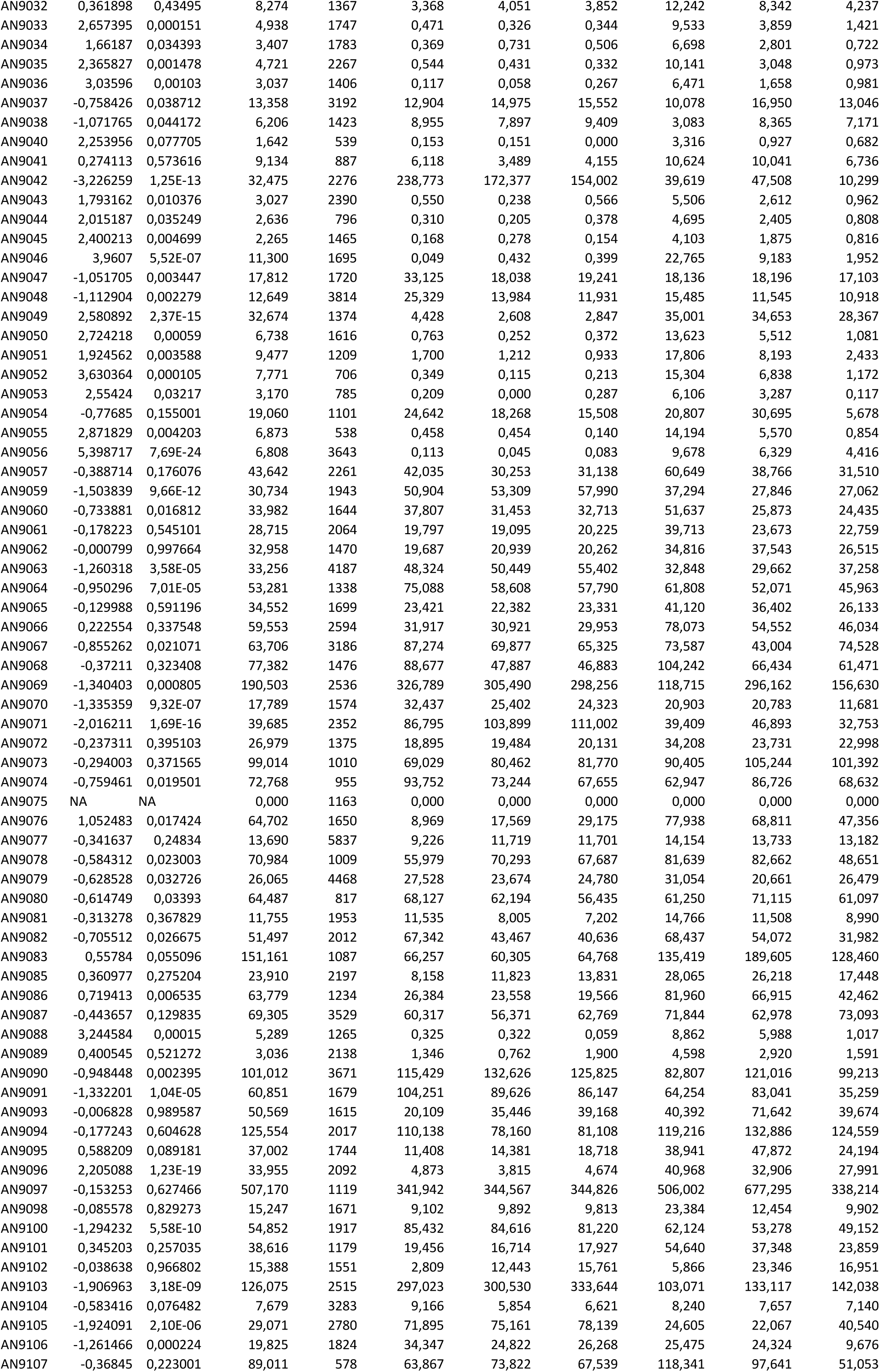
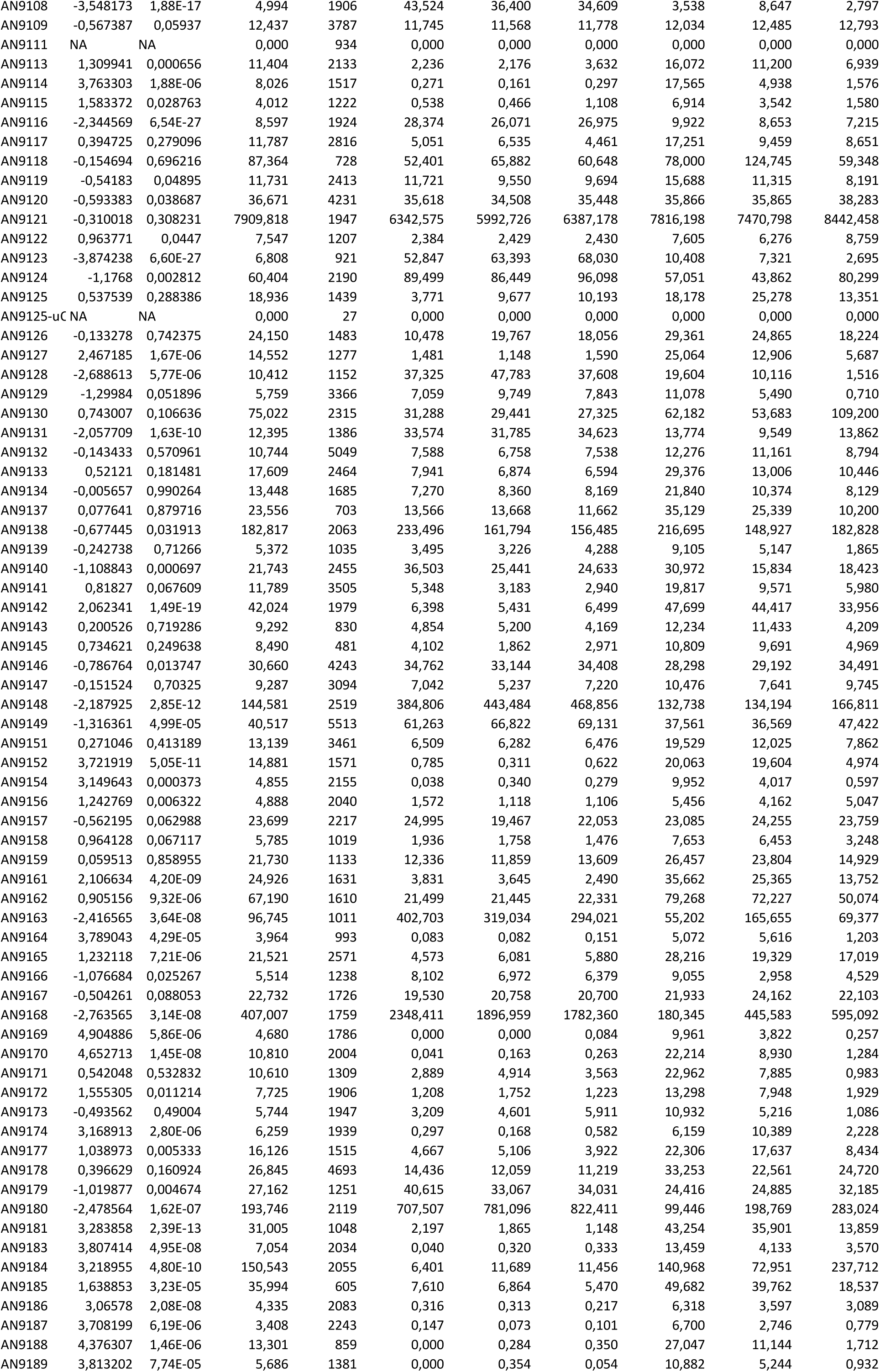
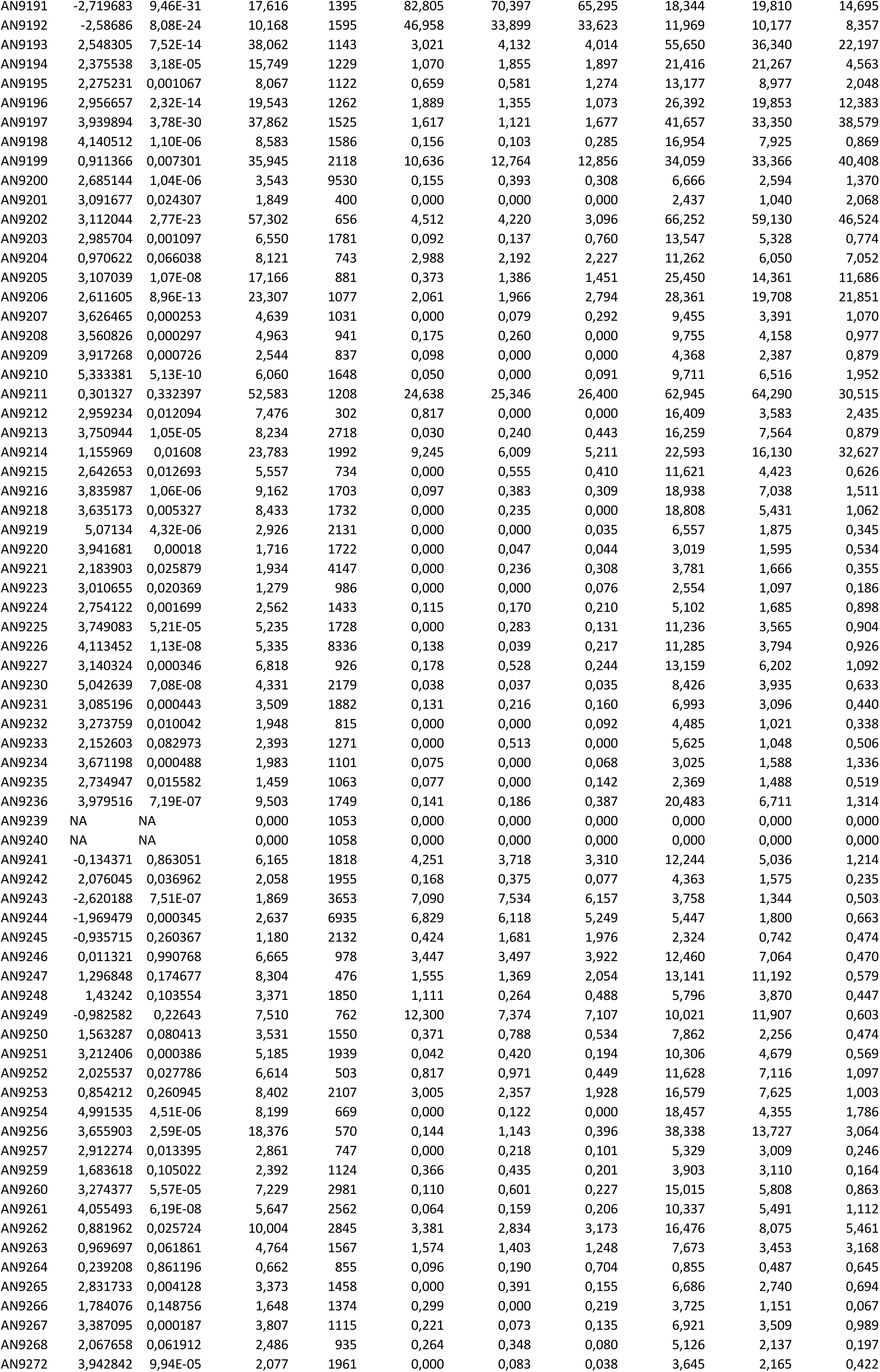
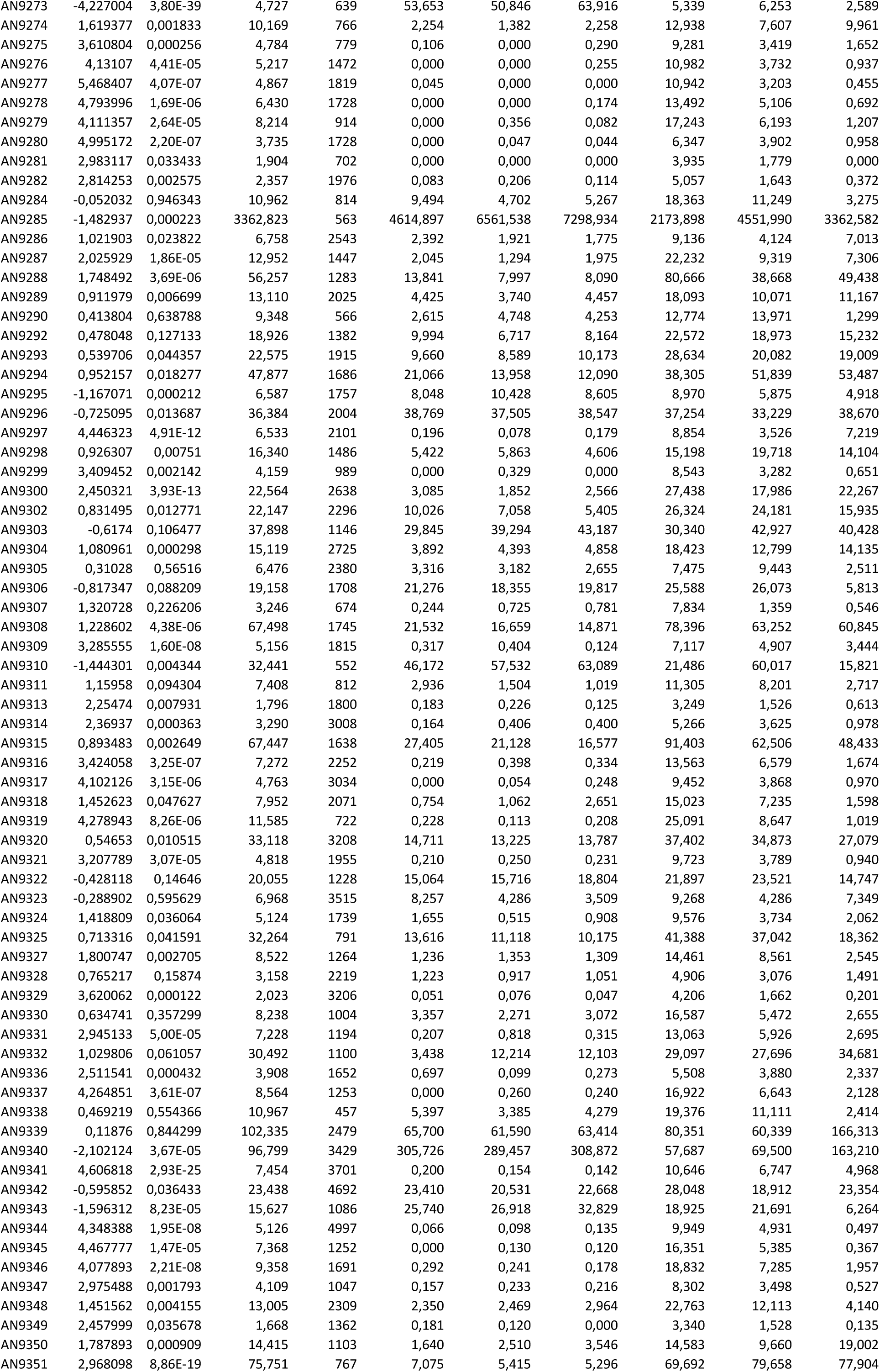
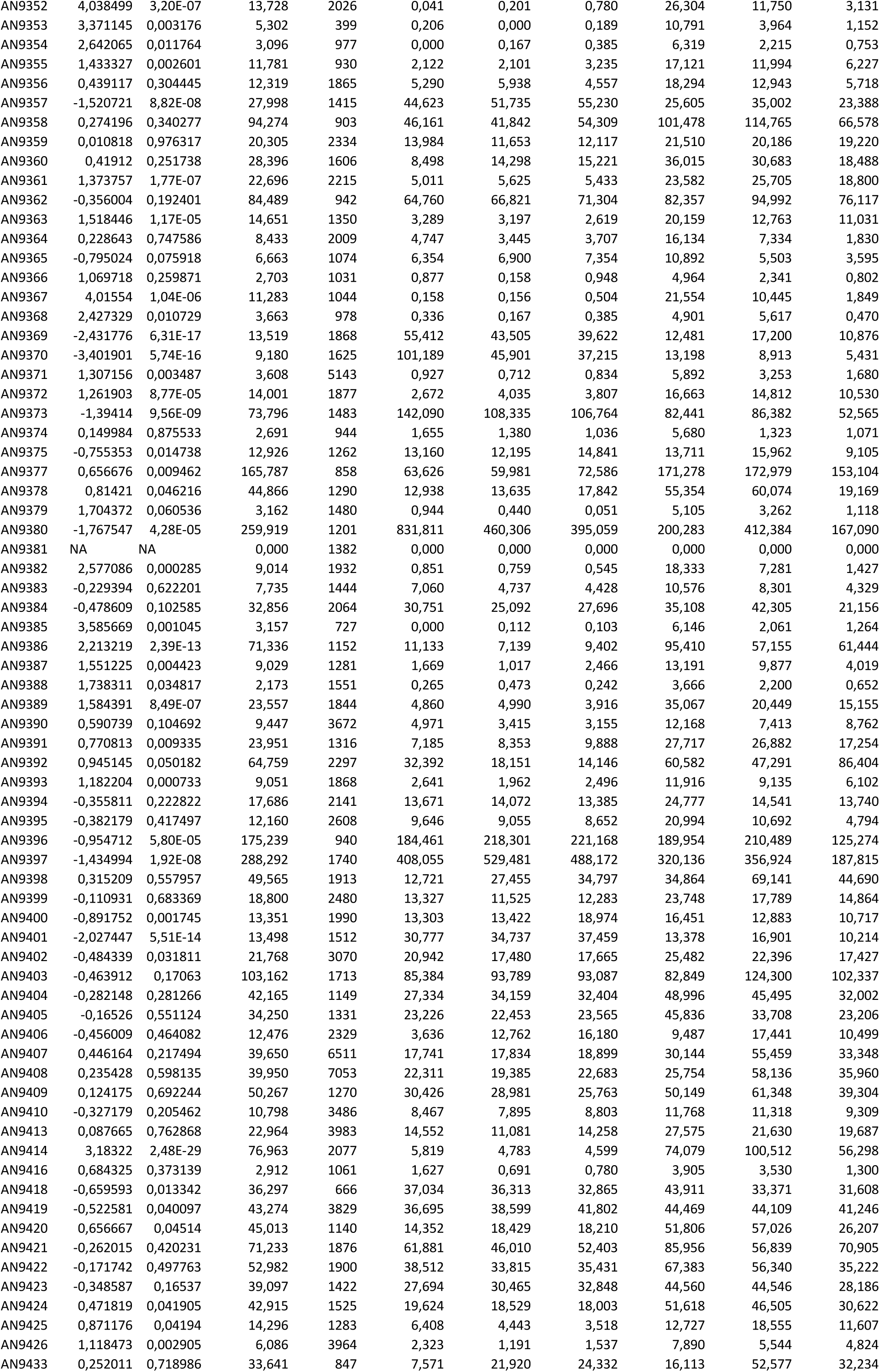
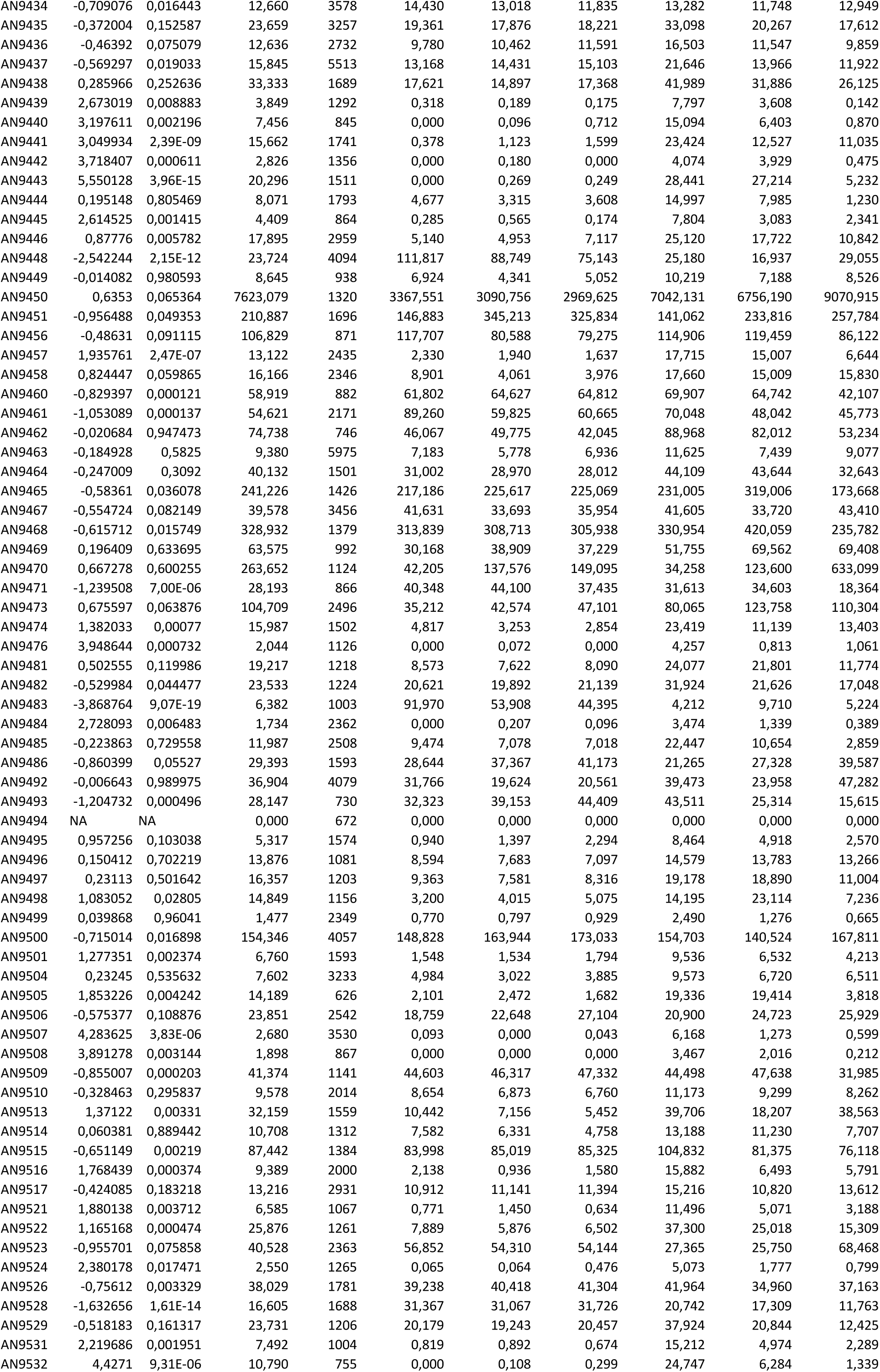
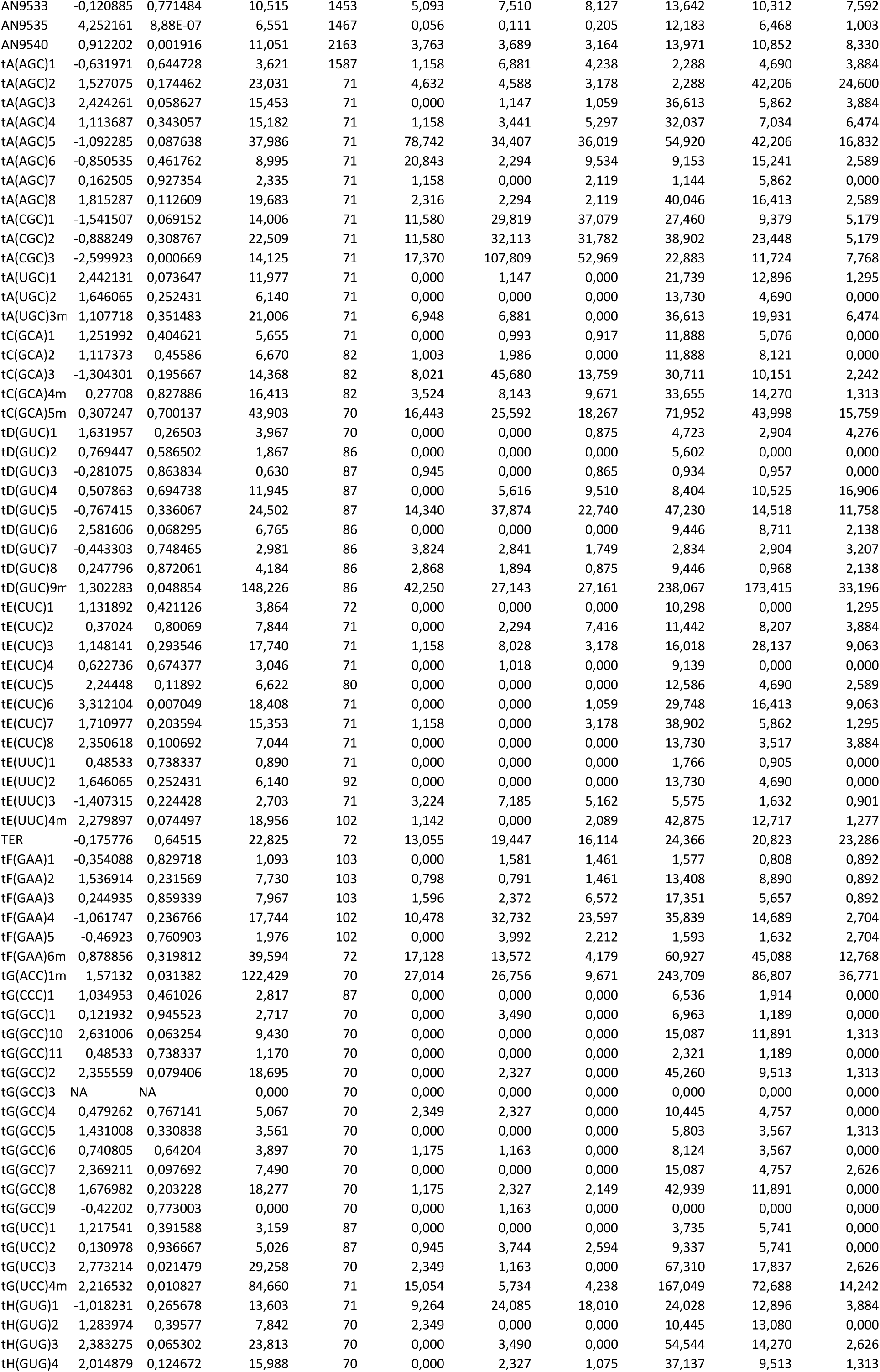
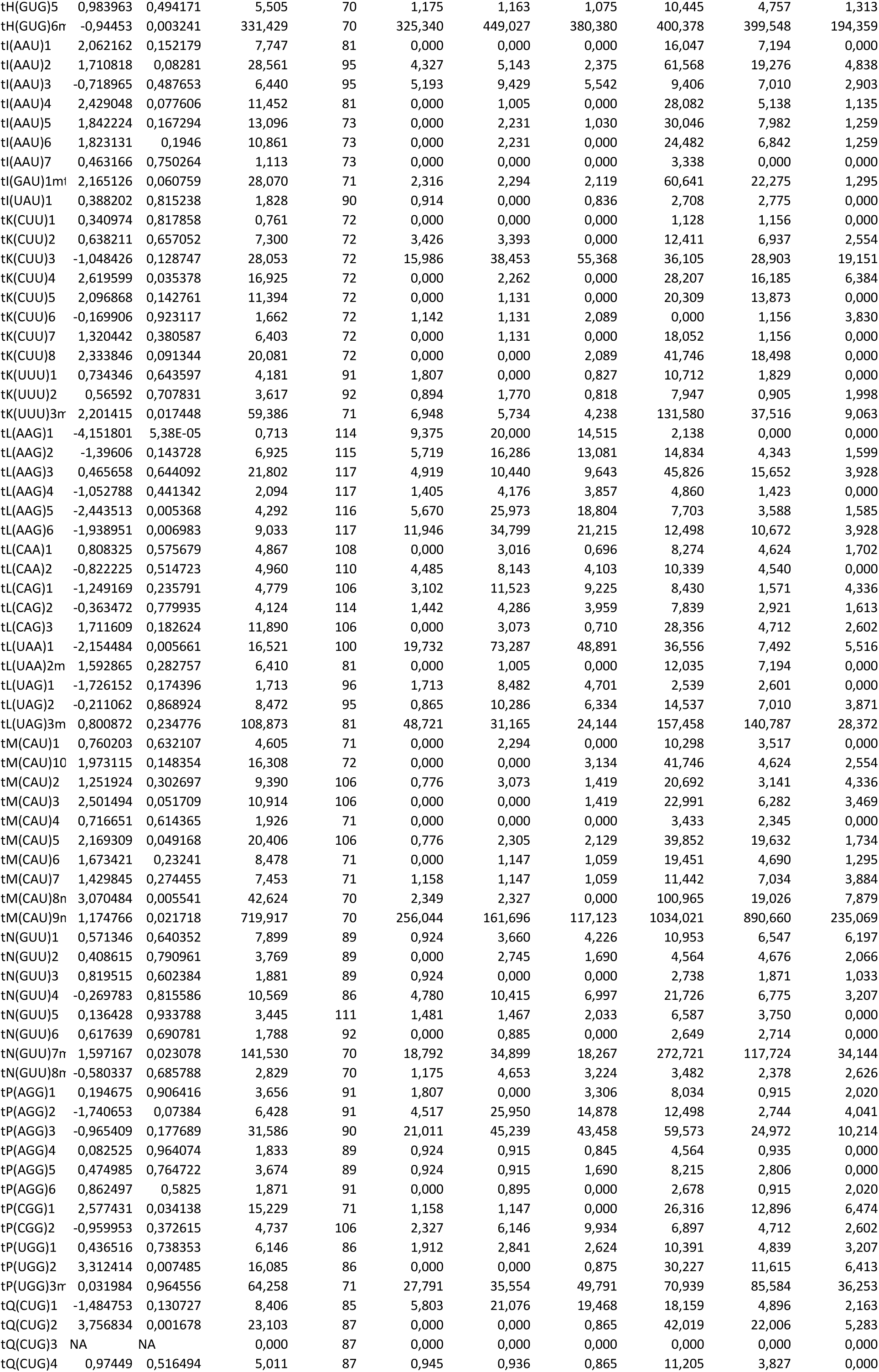
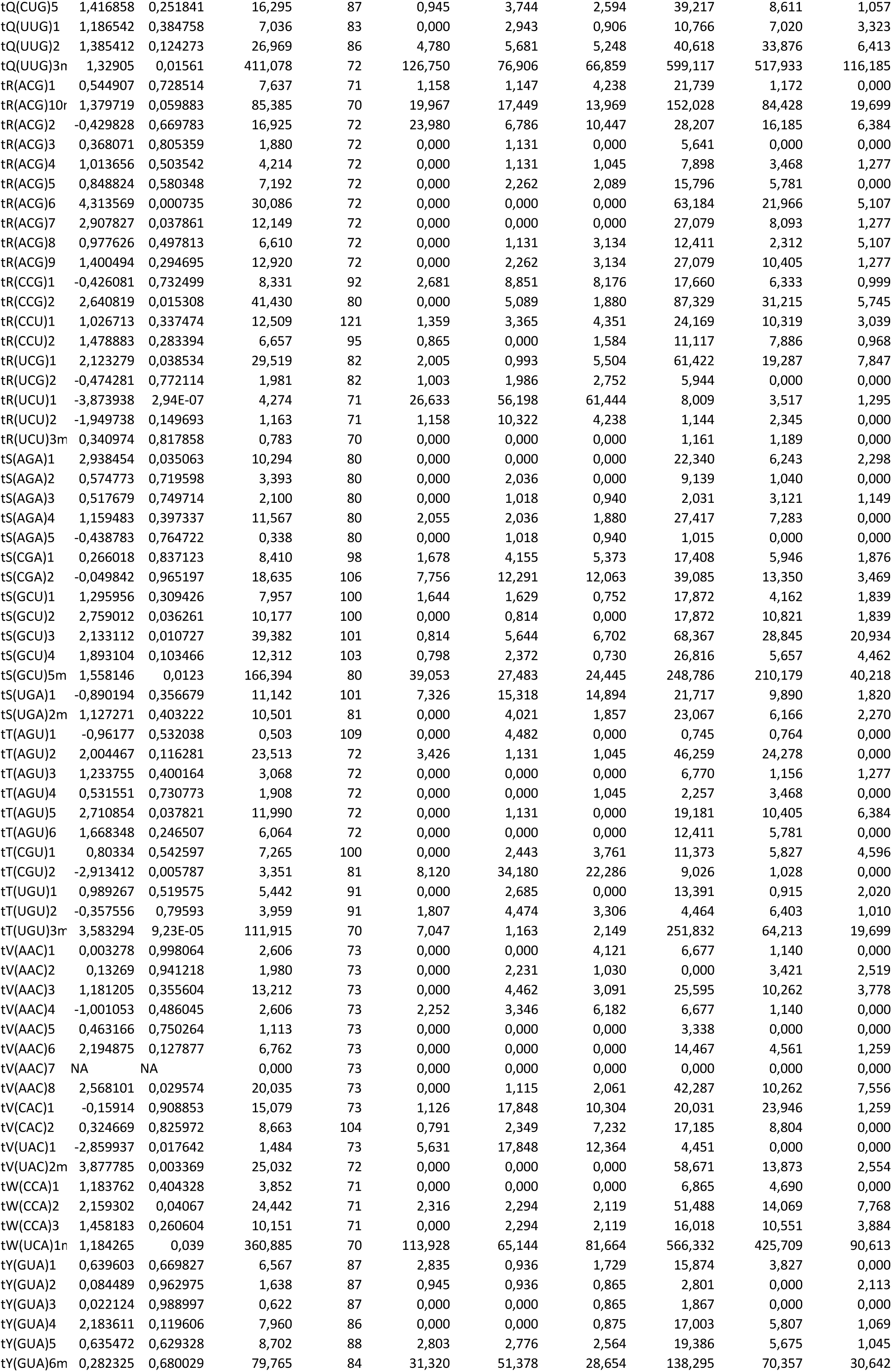
Aspergillus nidulans RNA-seq gene expression data.

**Table S6.**
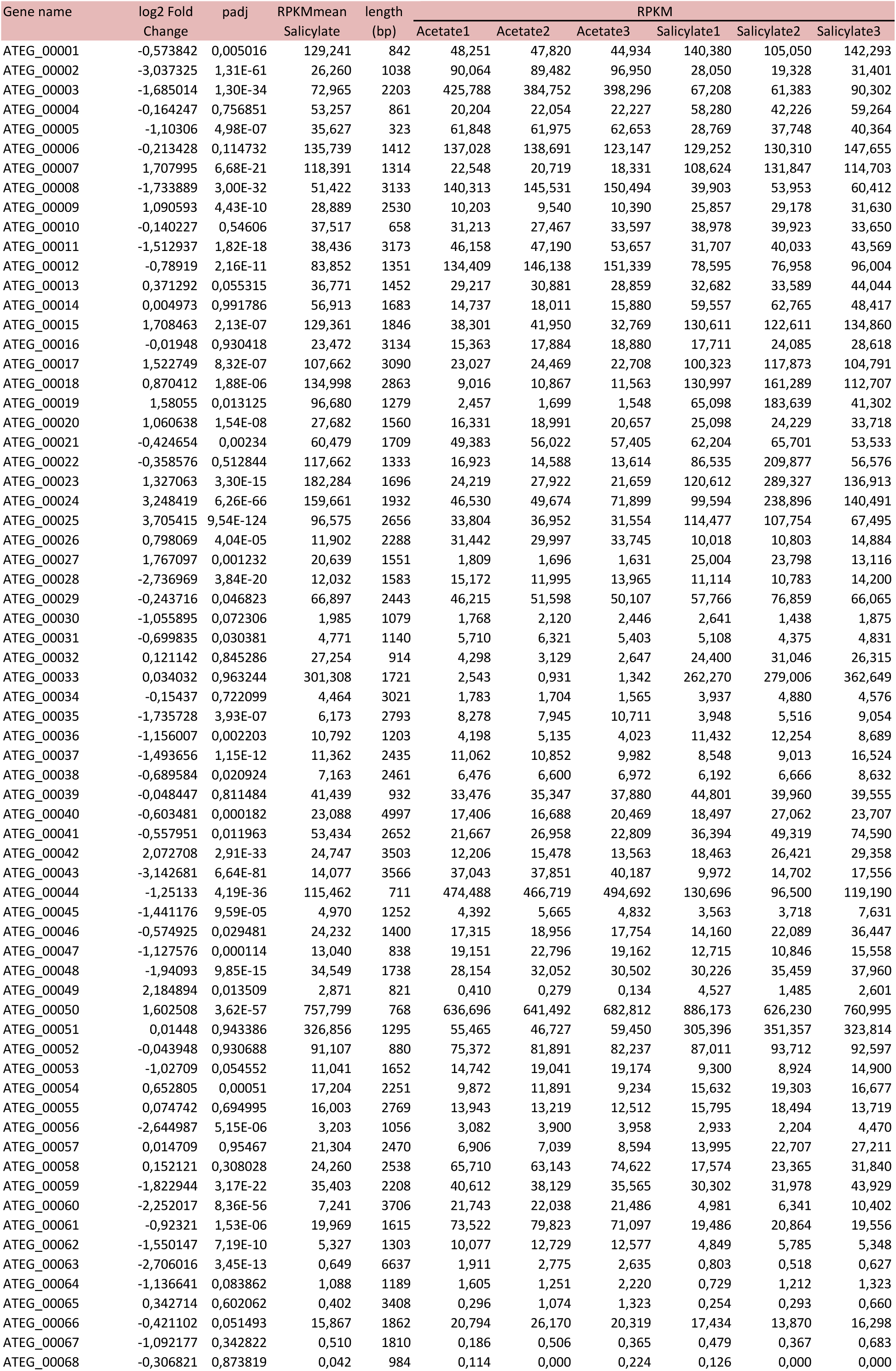
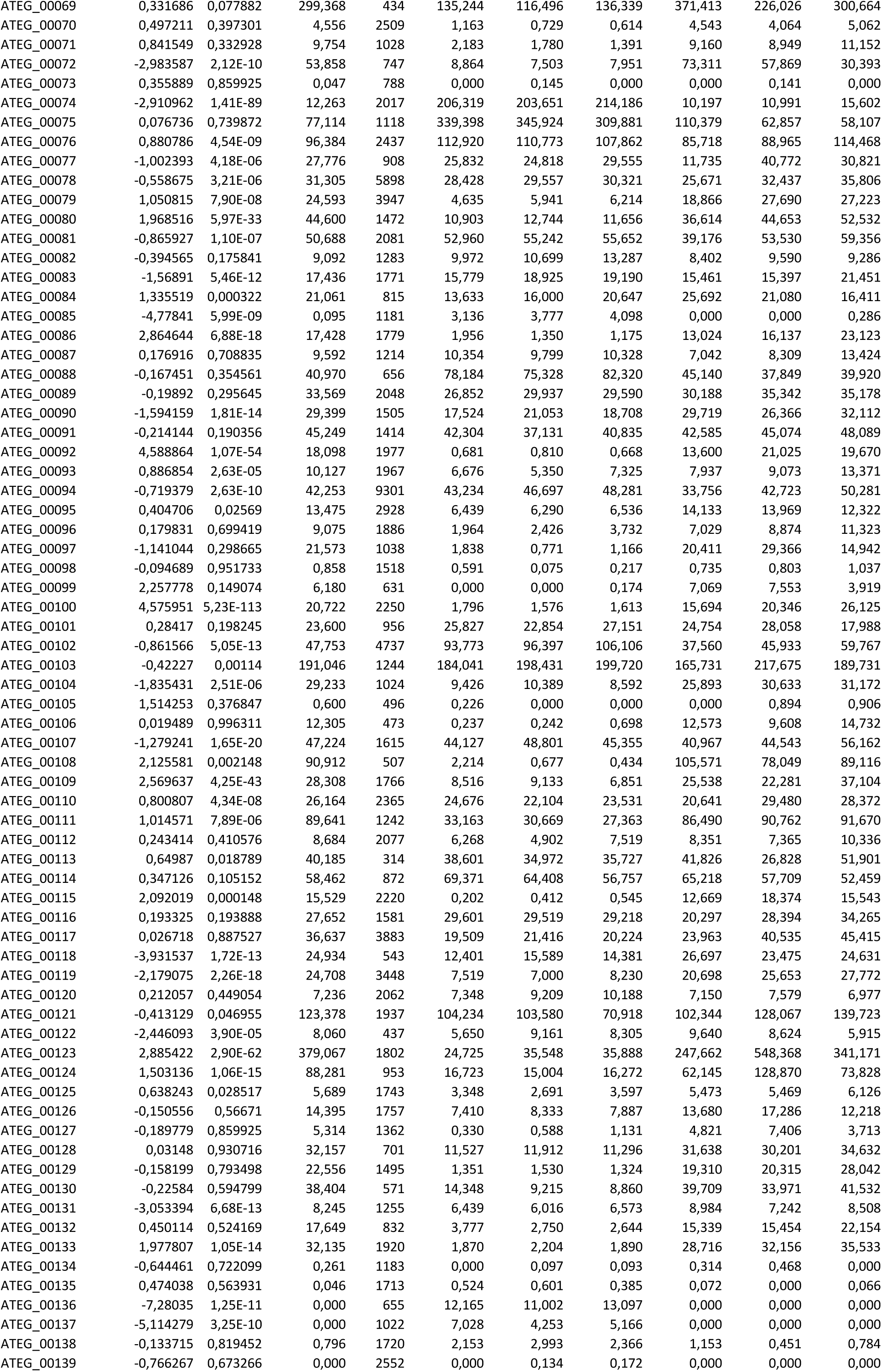
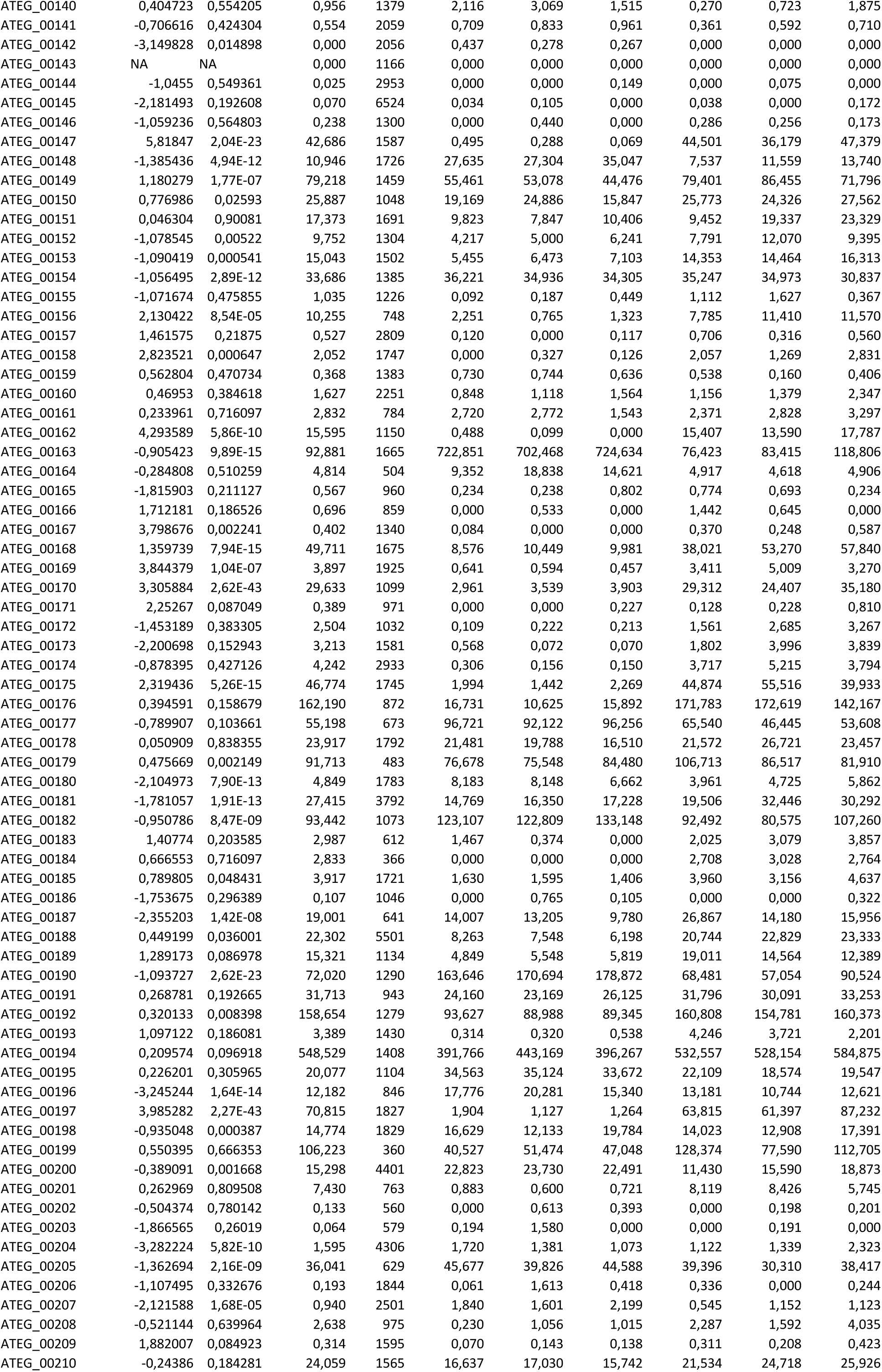
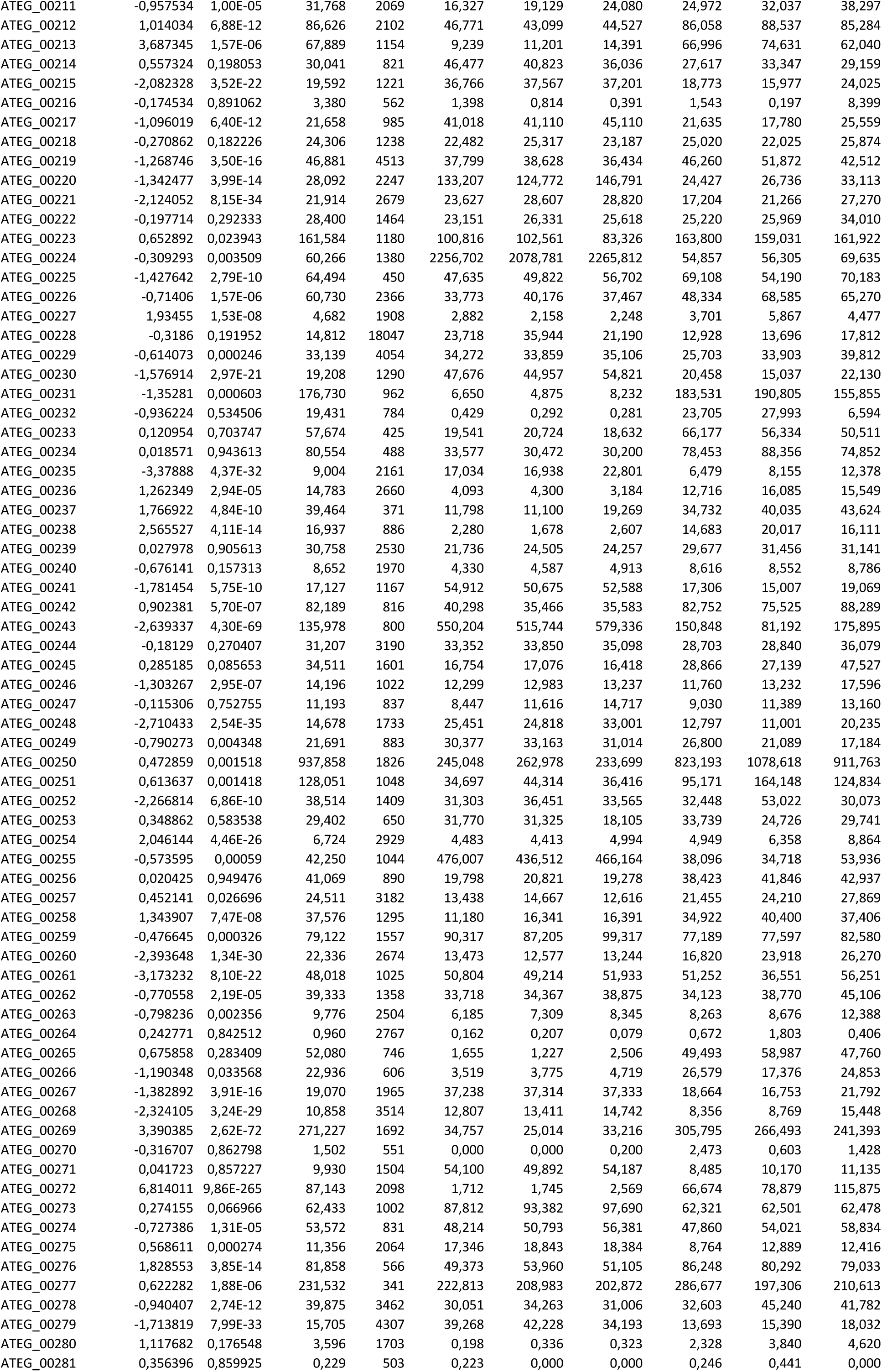
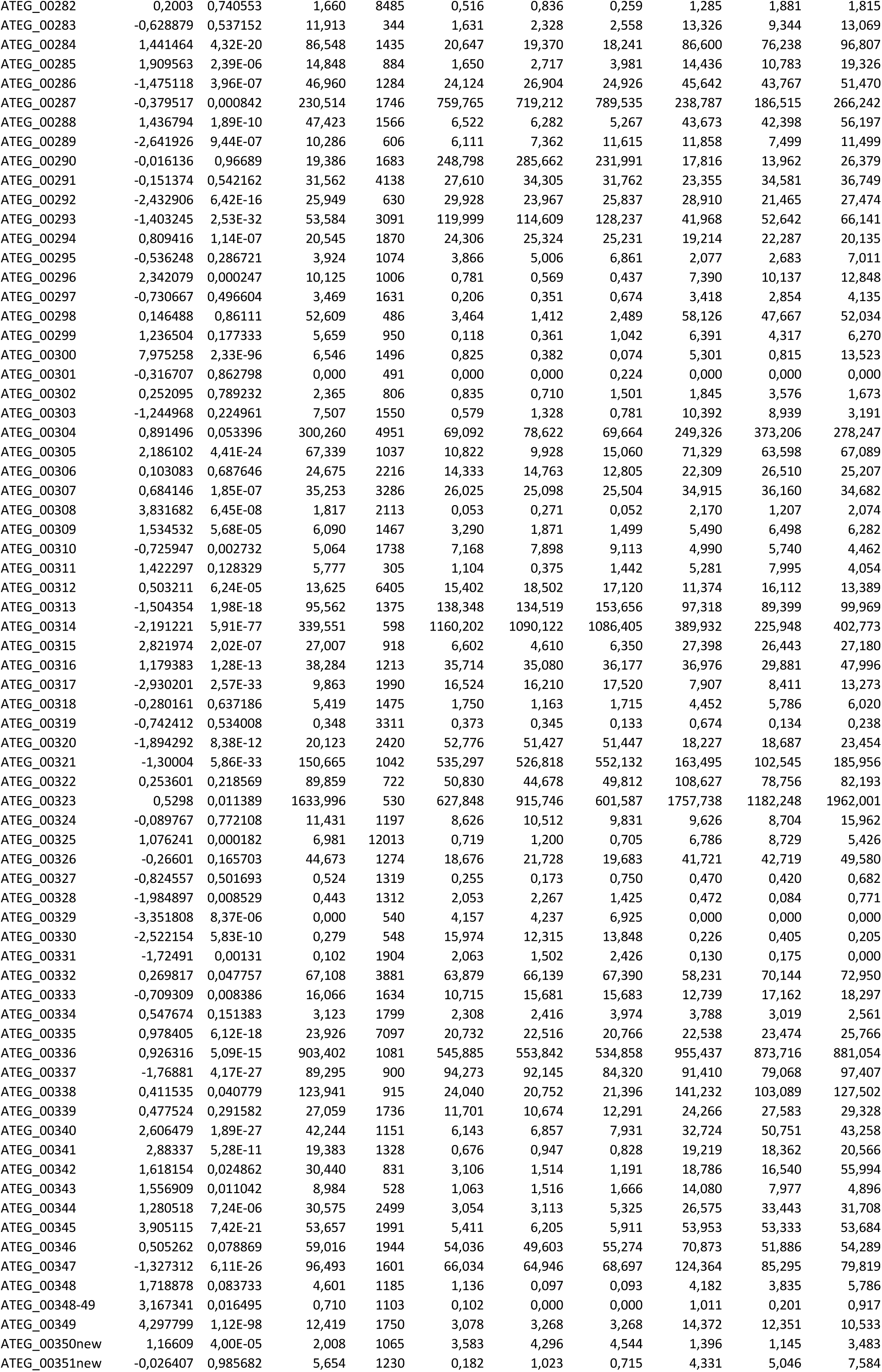
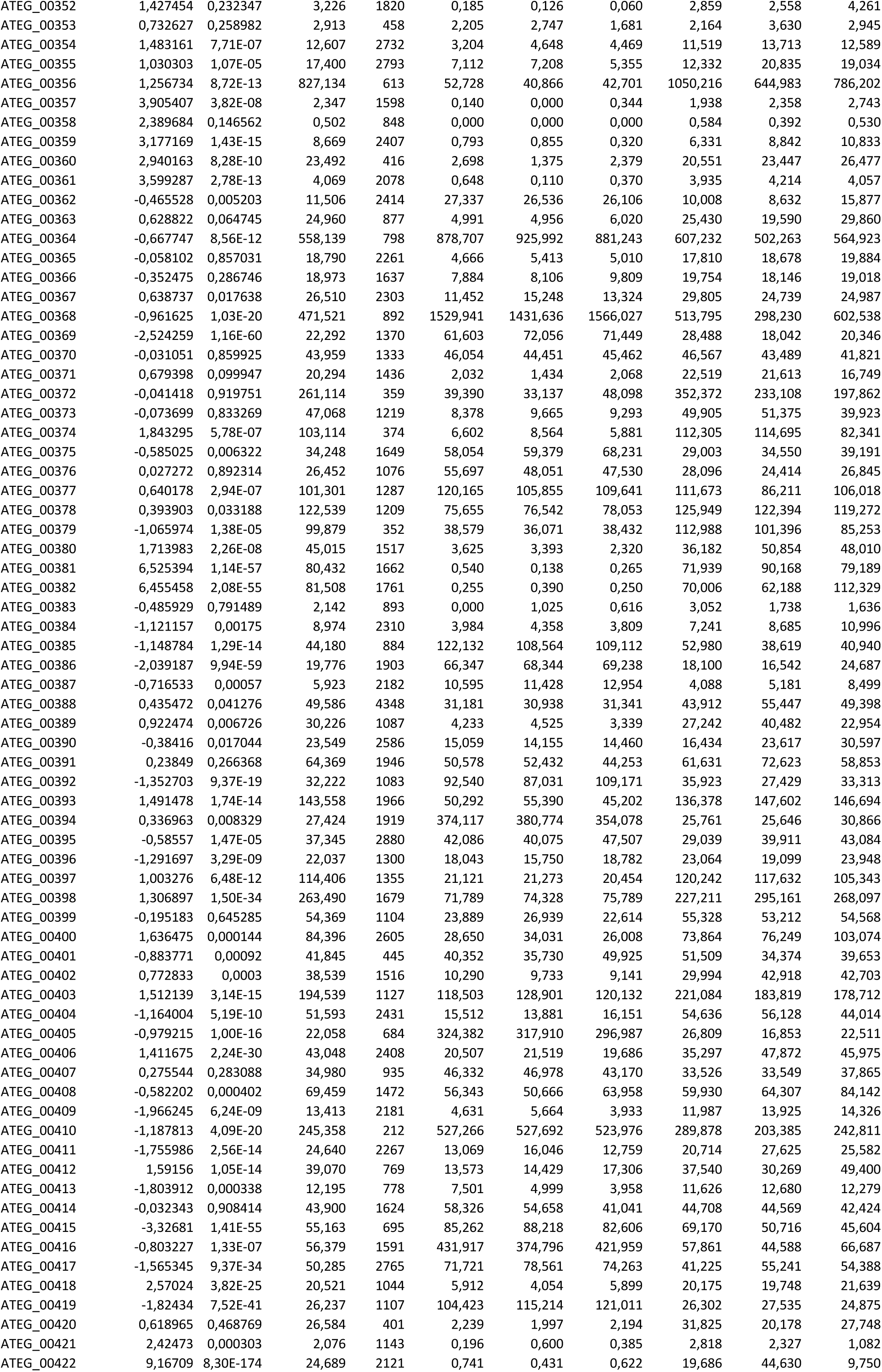
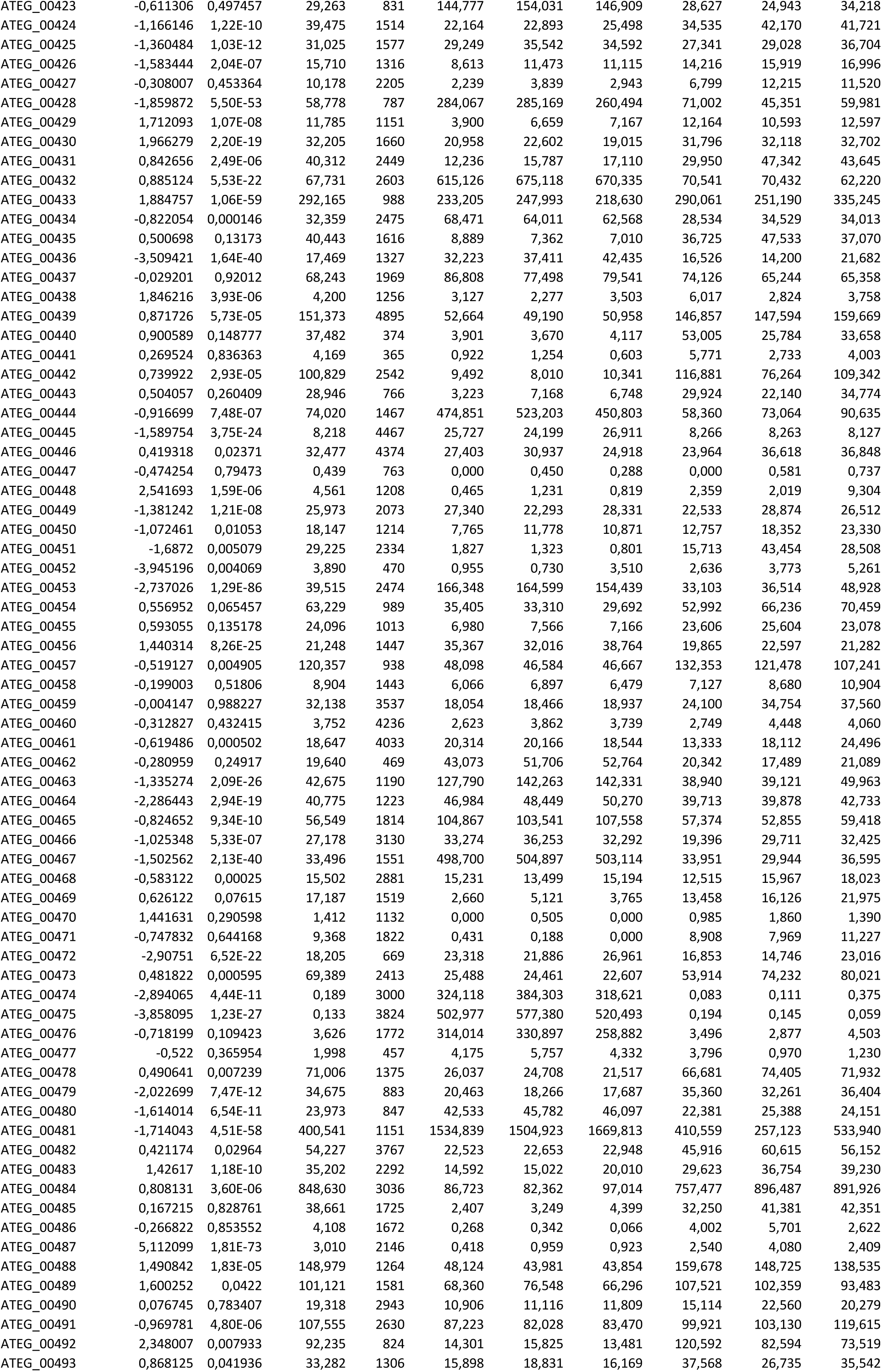
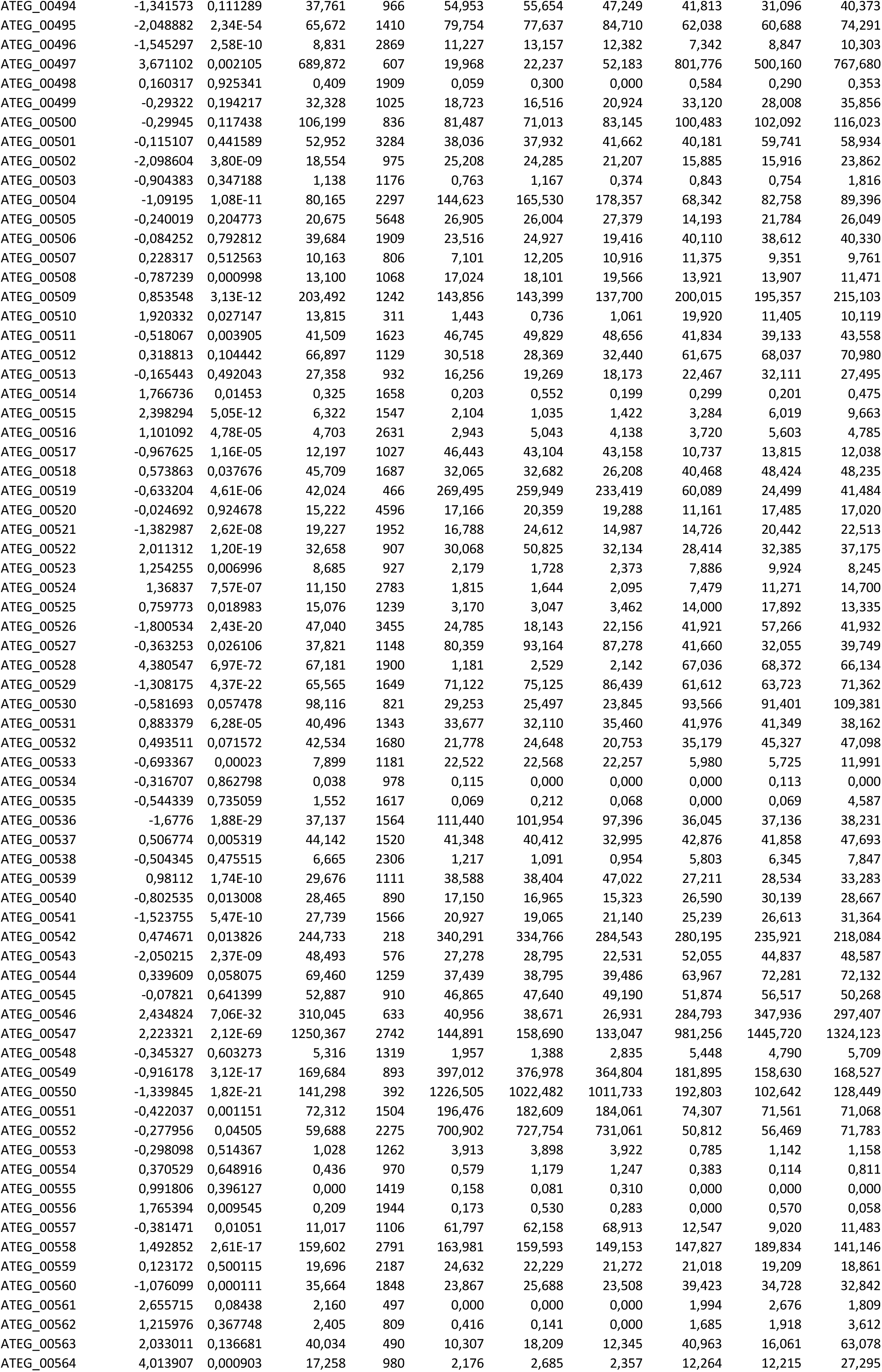
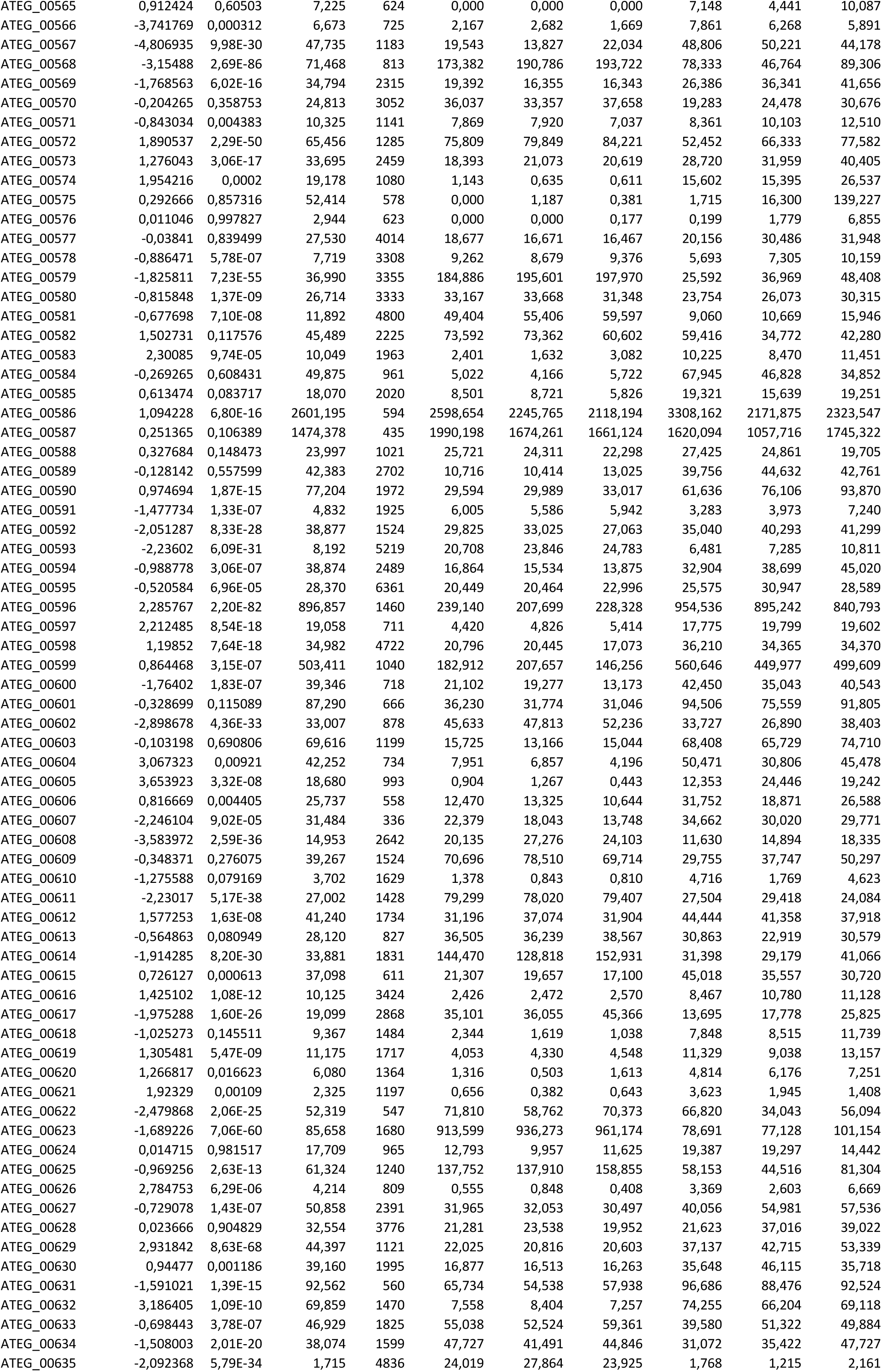
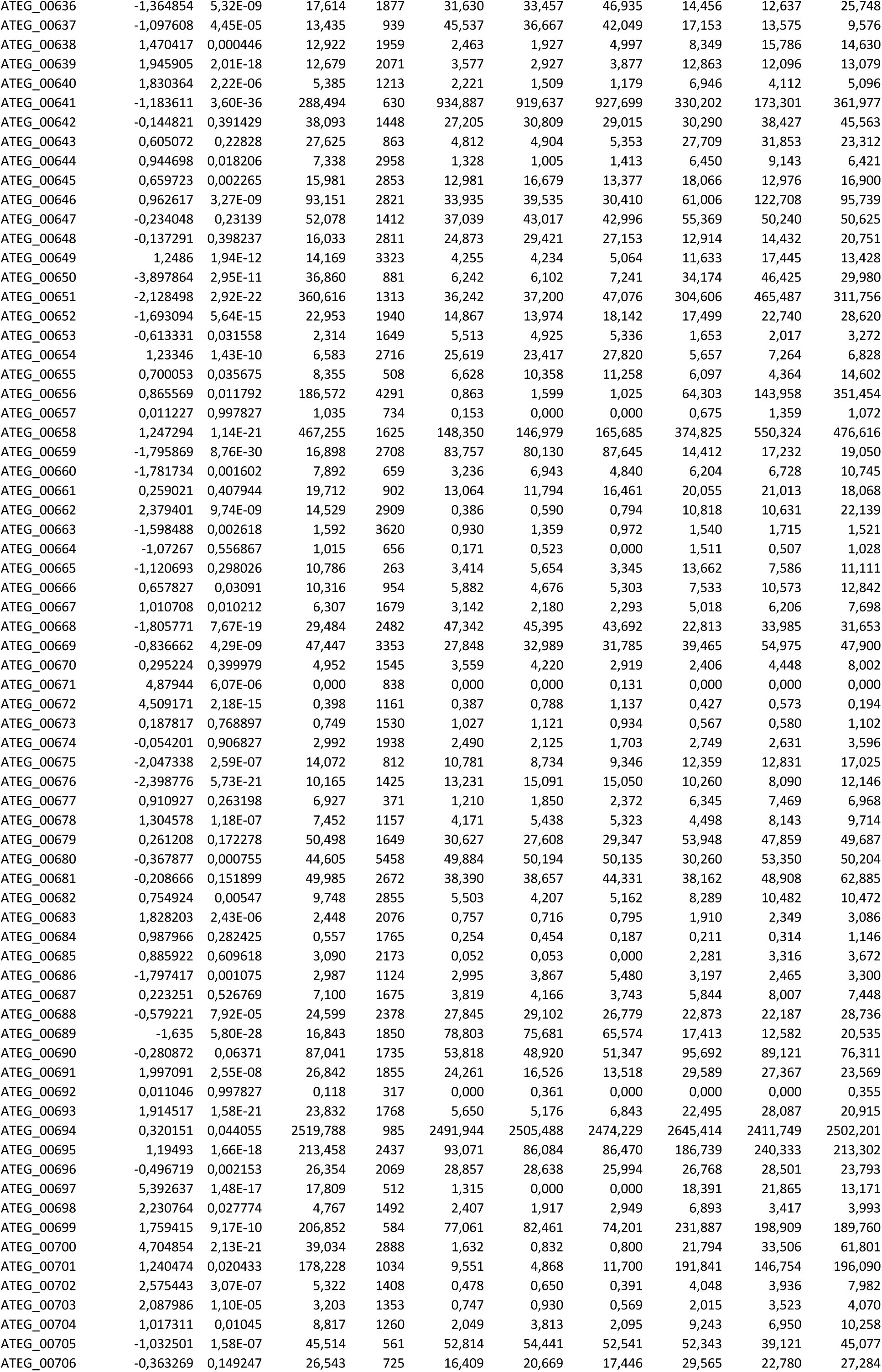
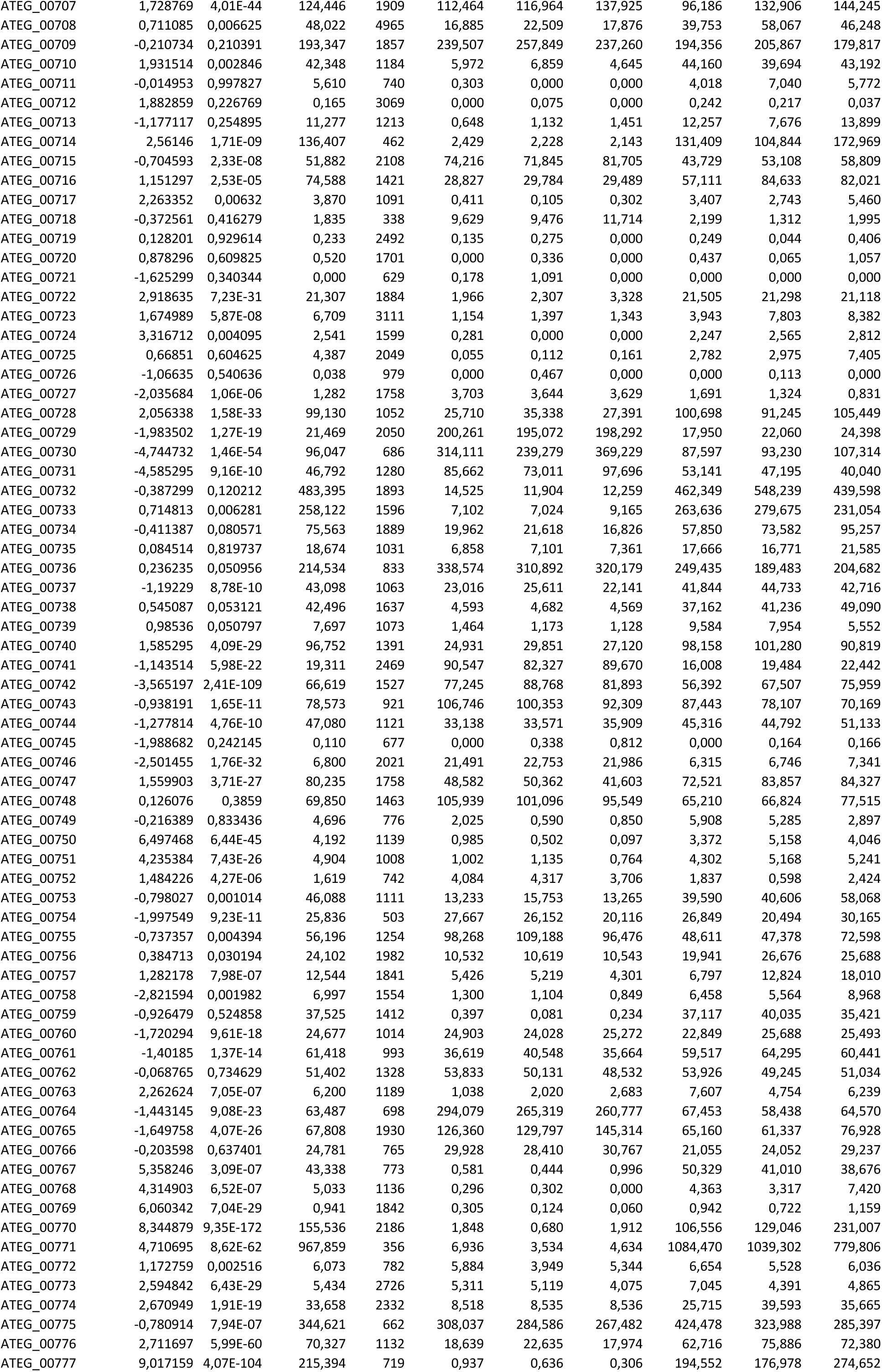
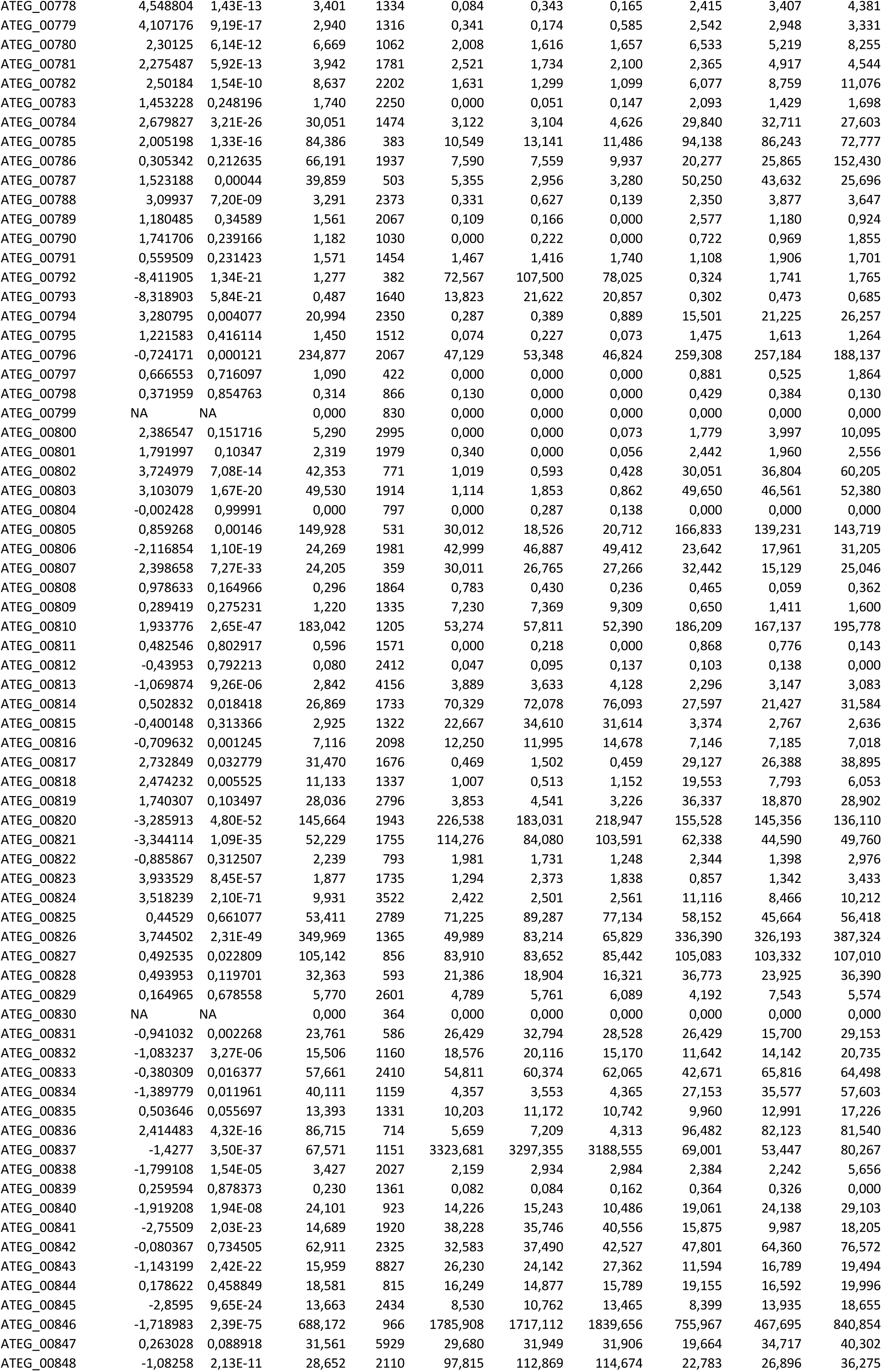
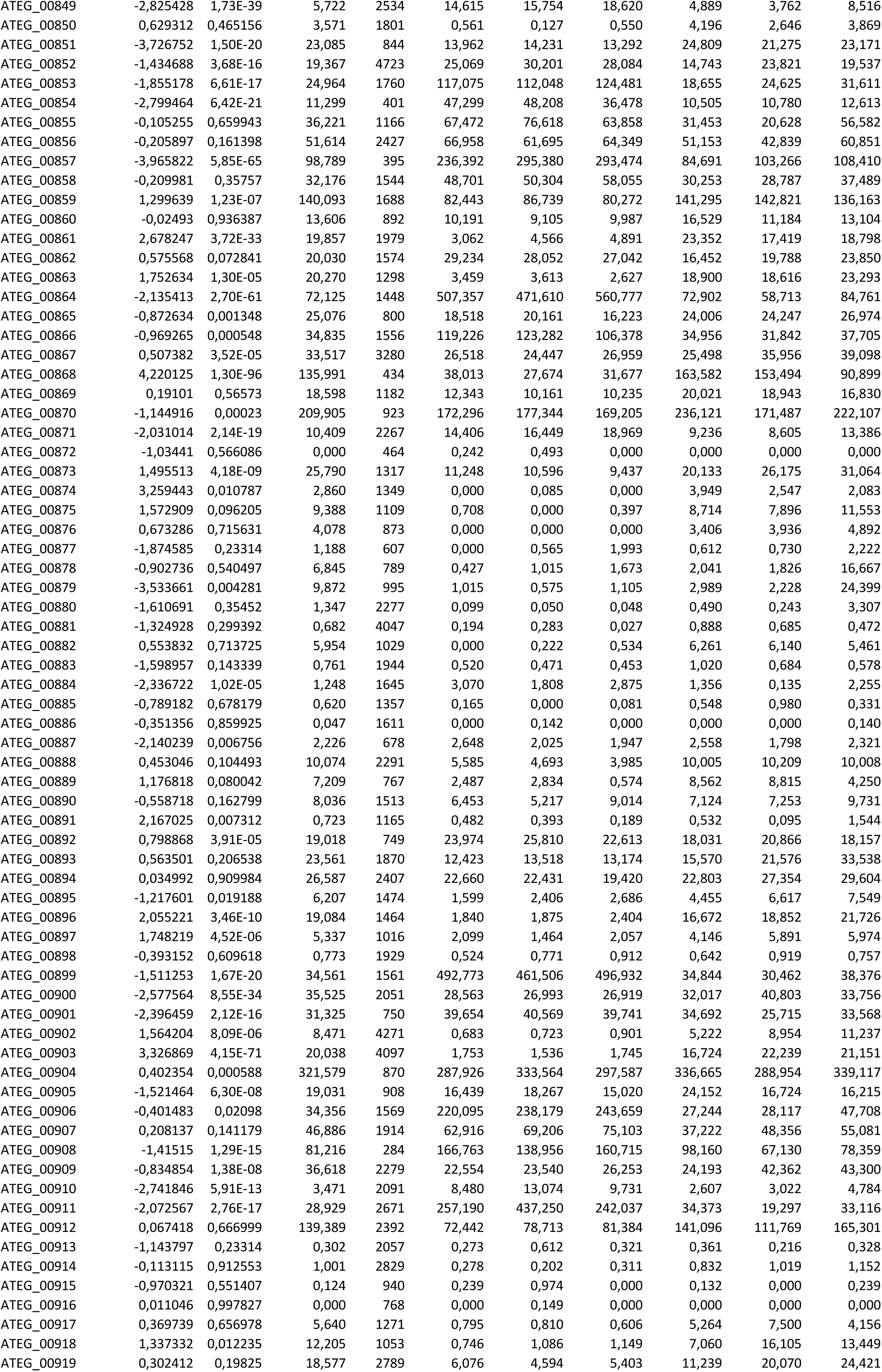
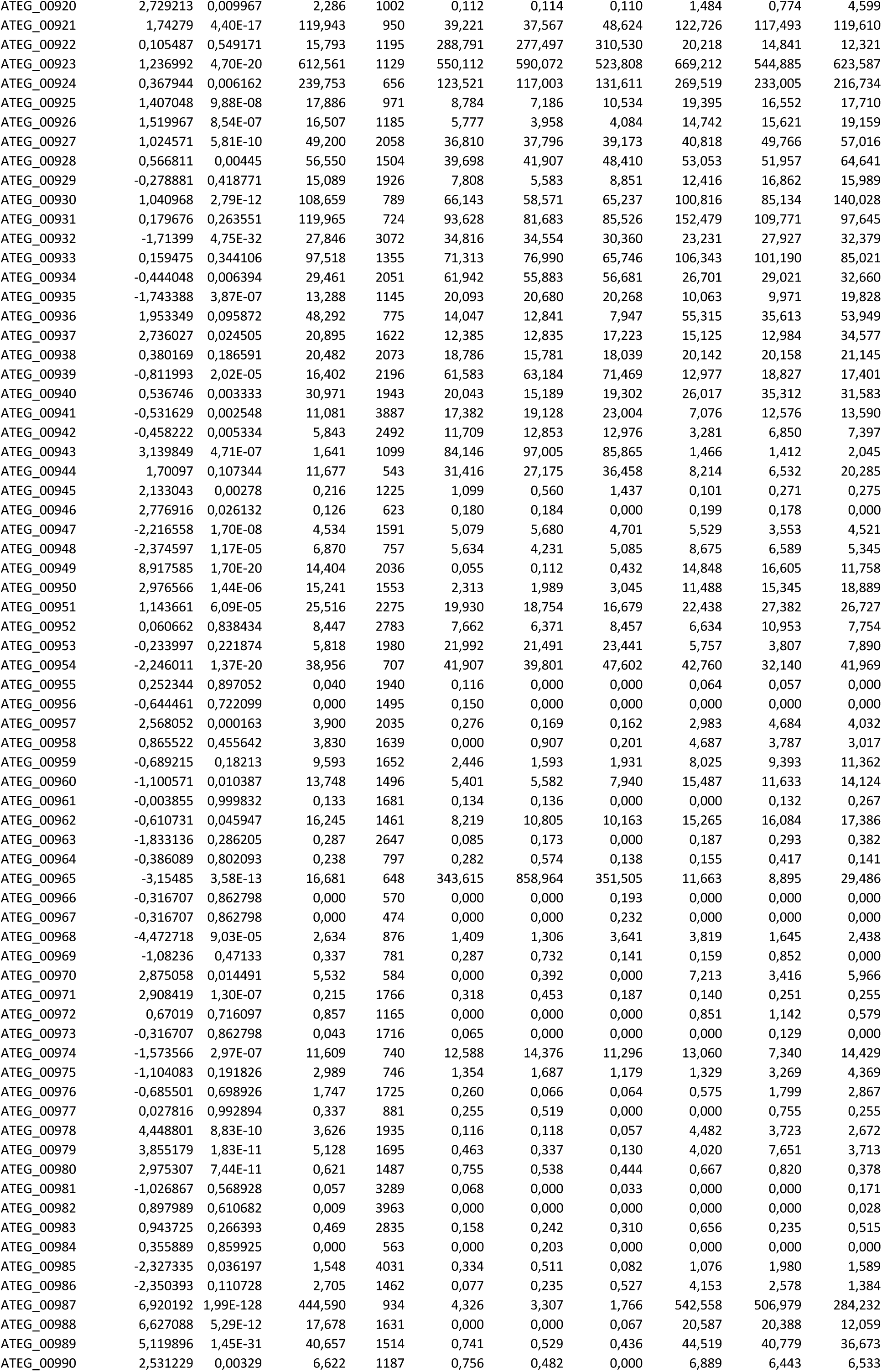
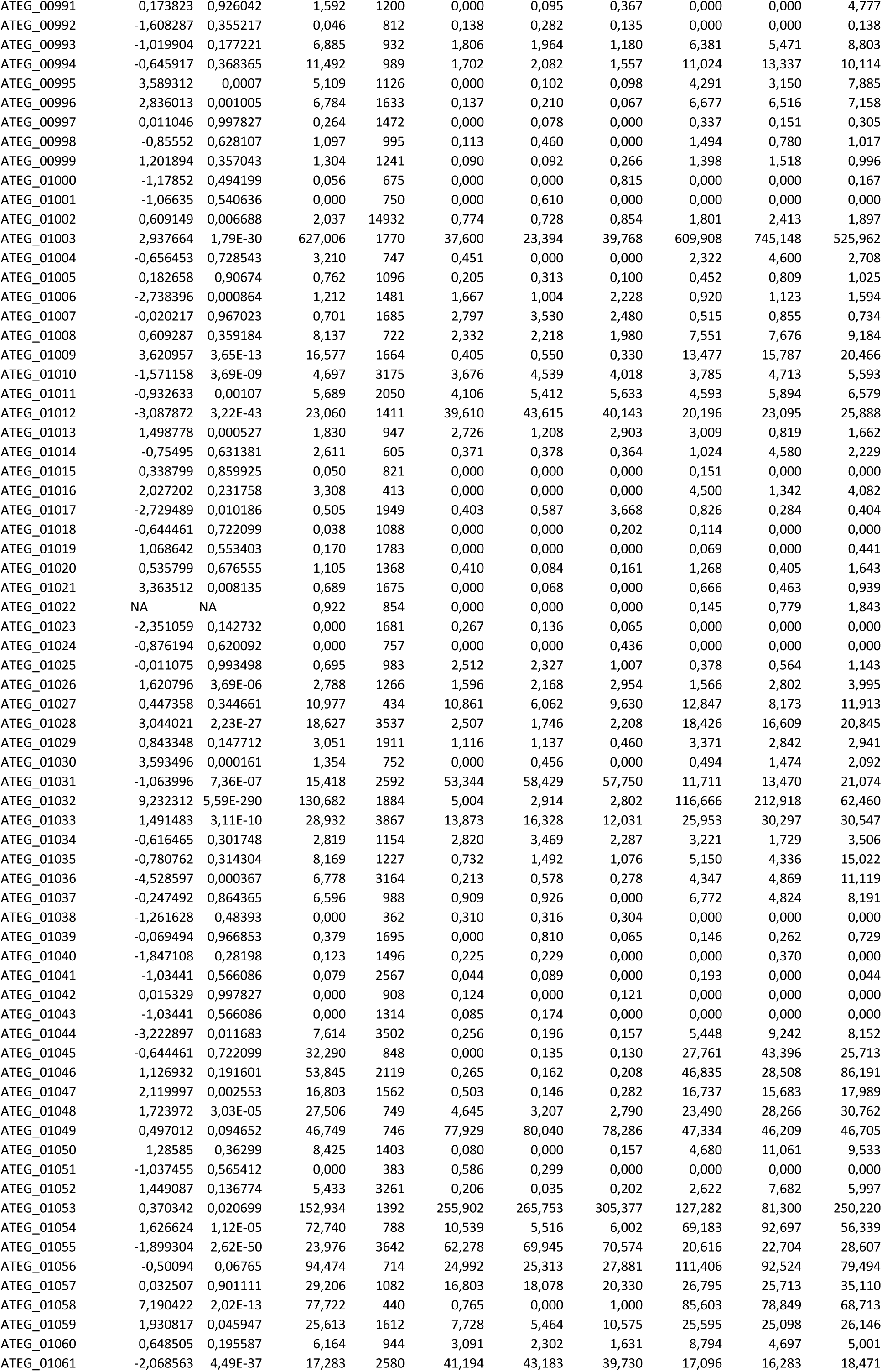
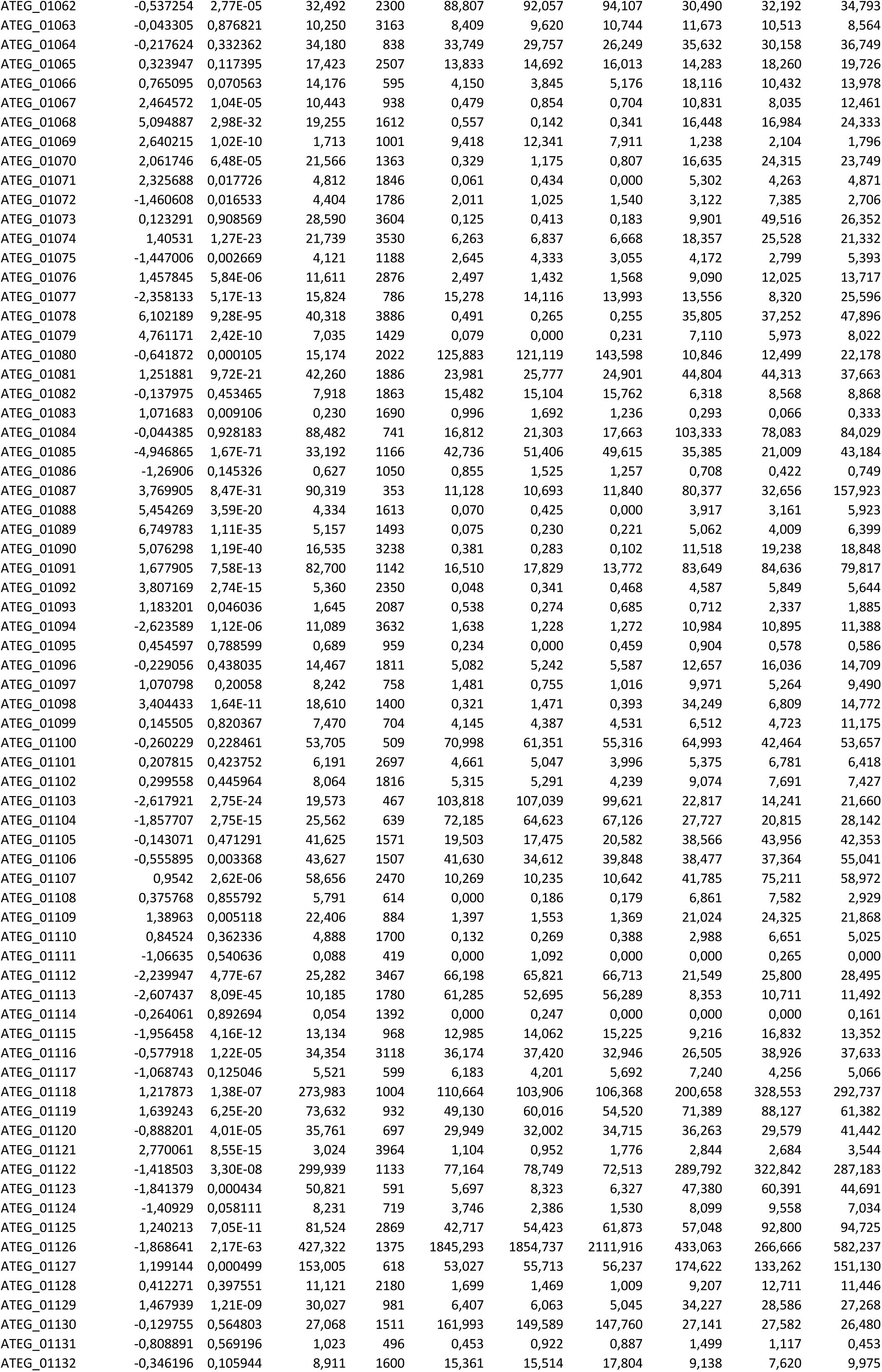
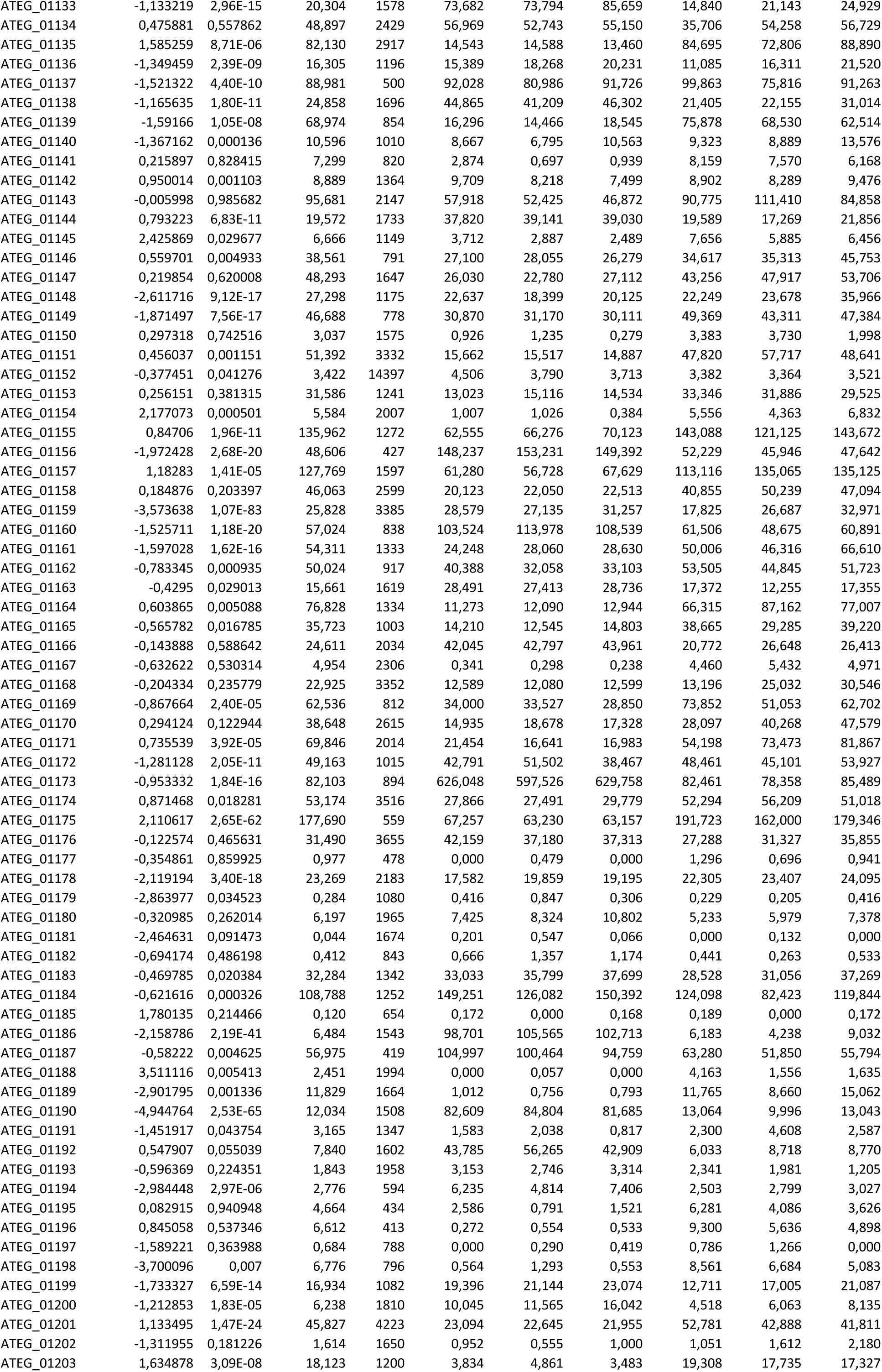
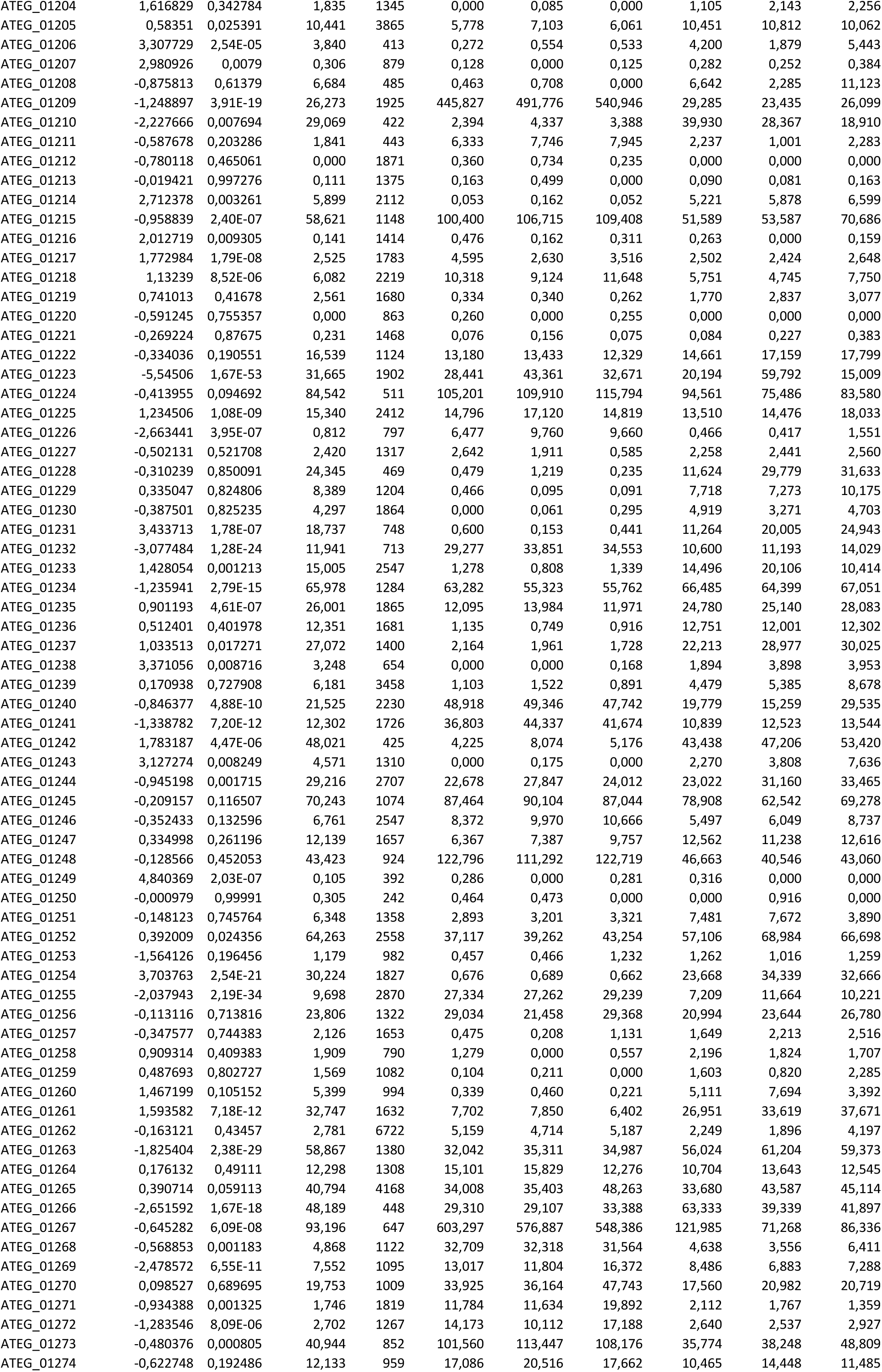
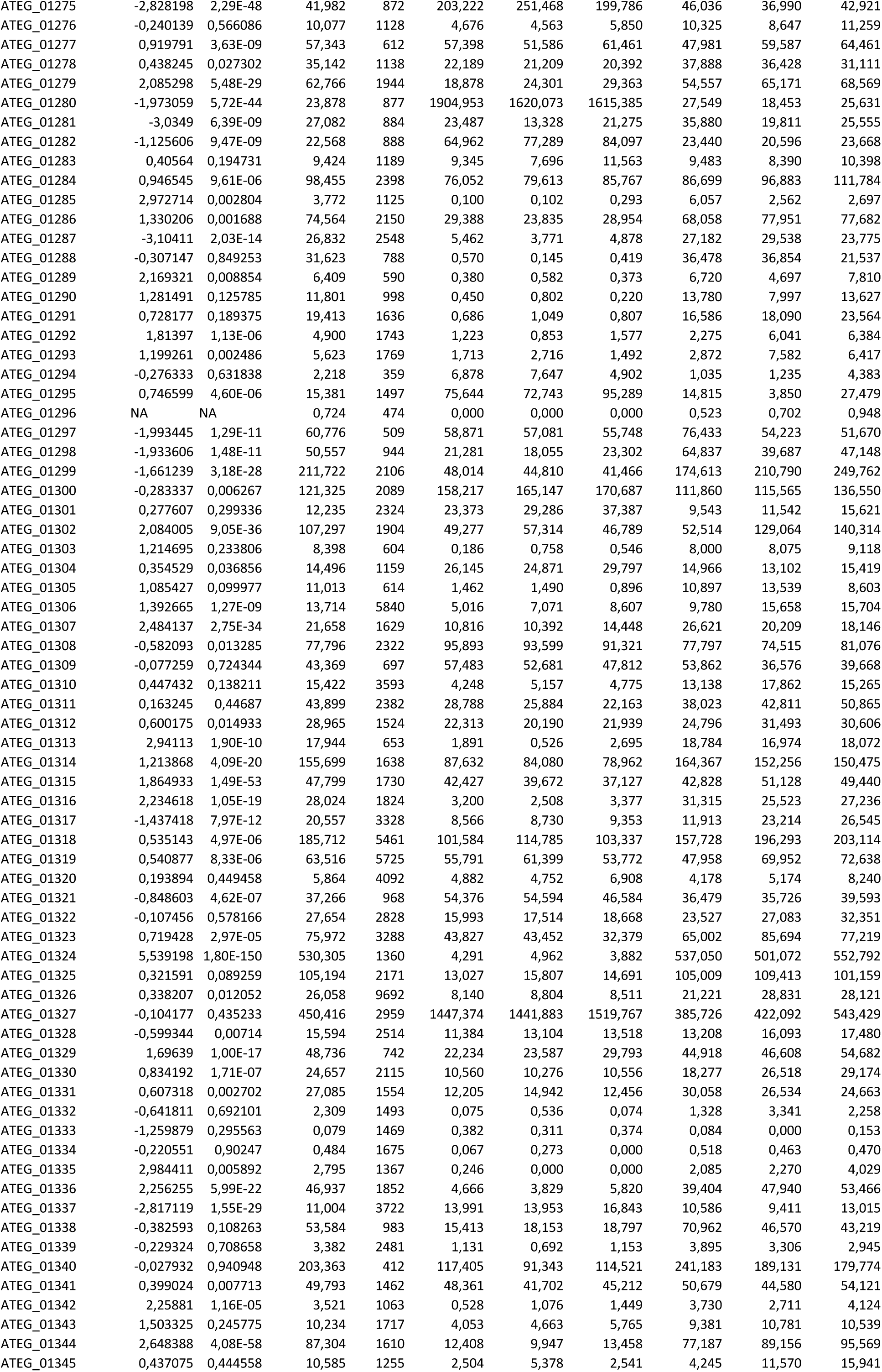
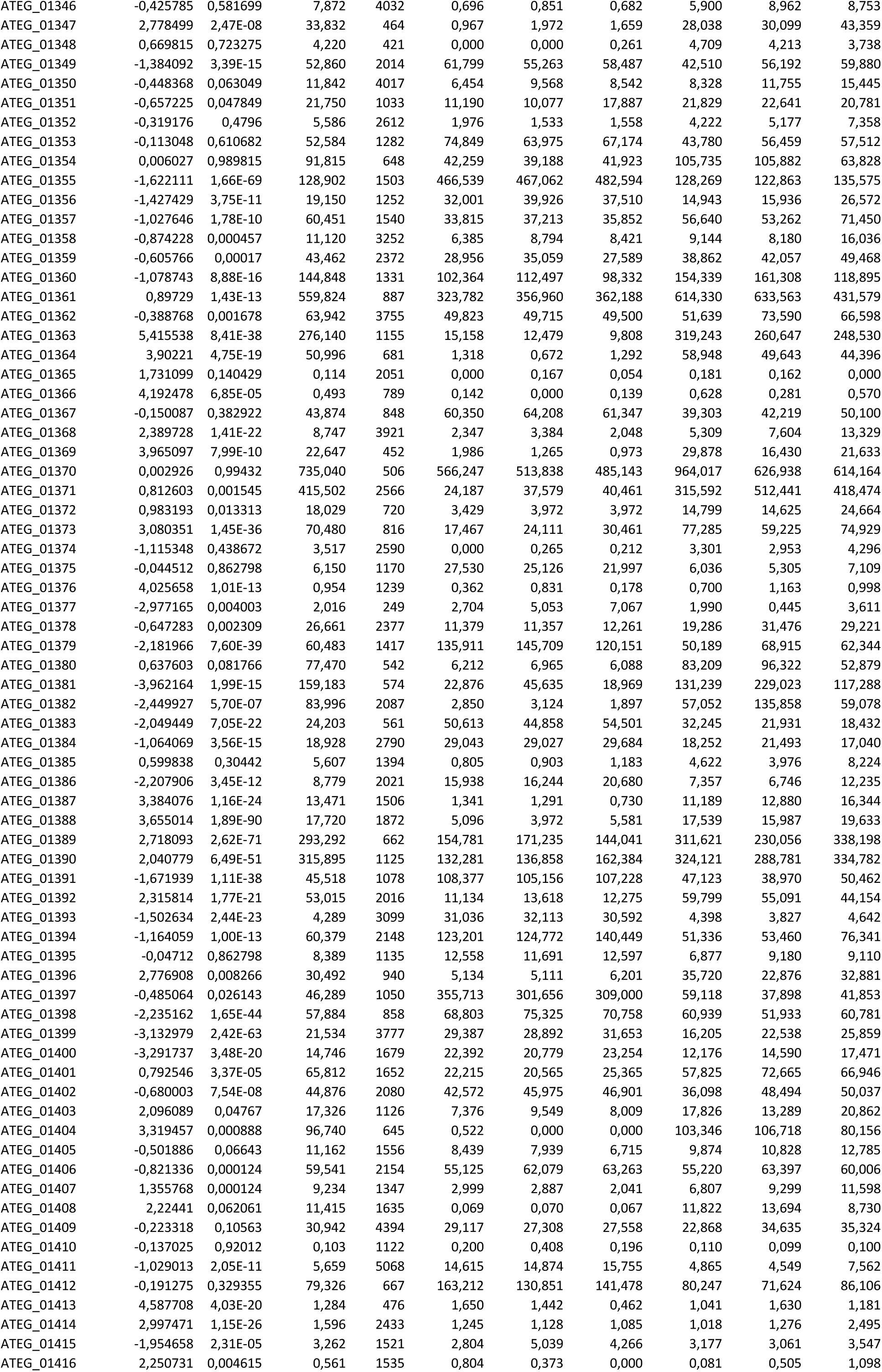
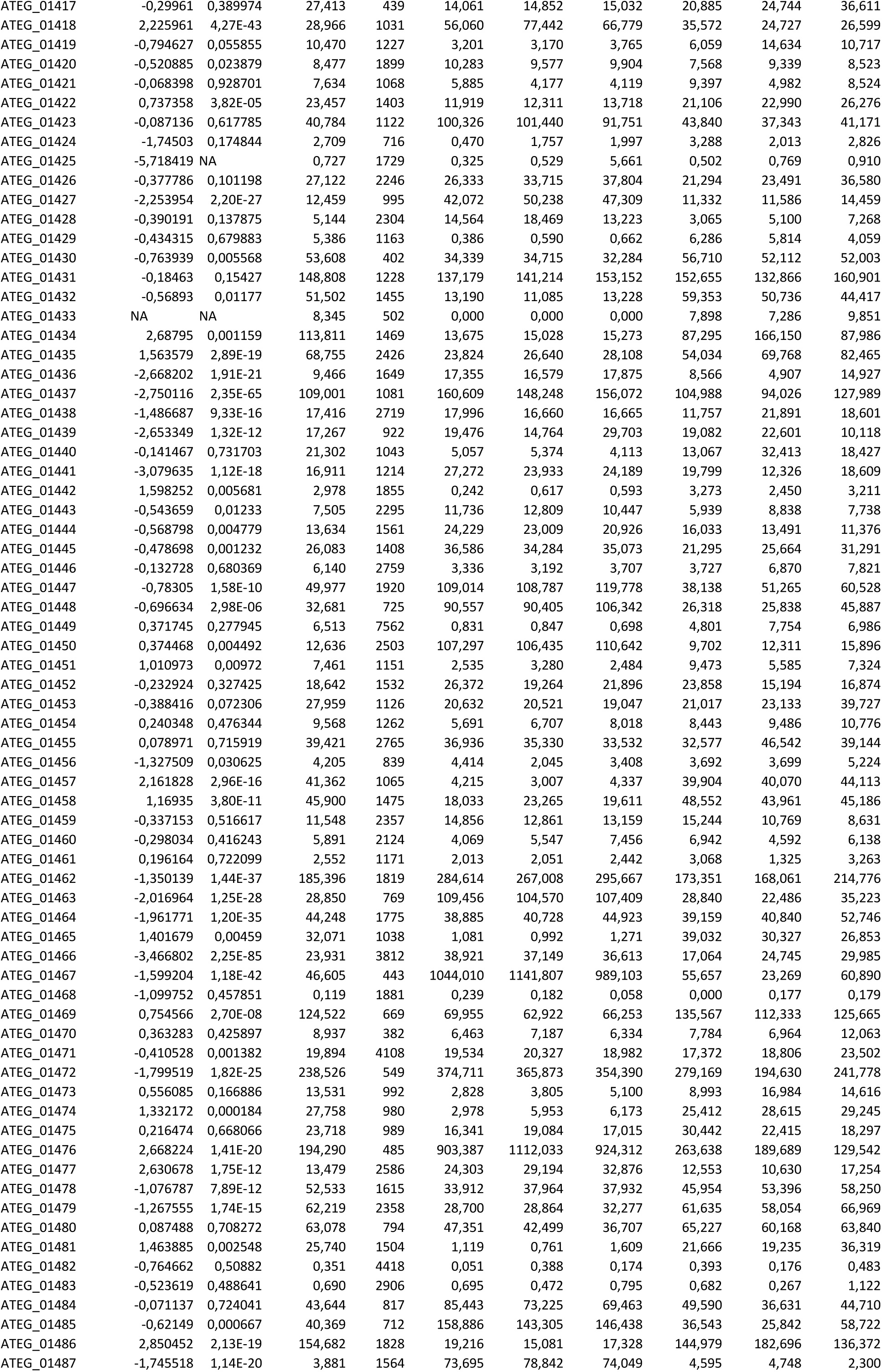
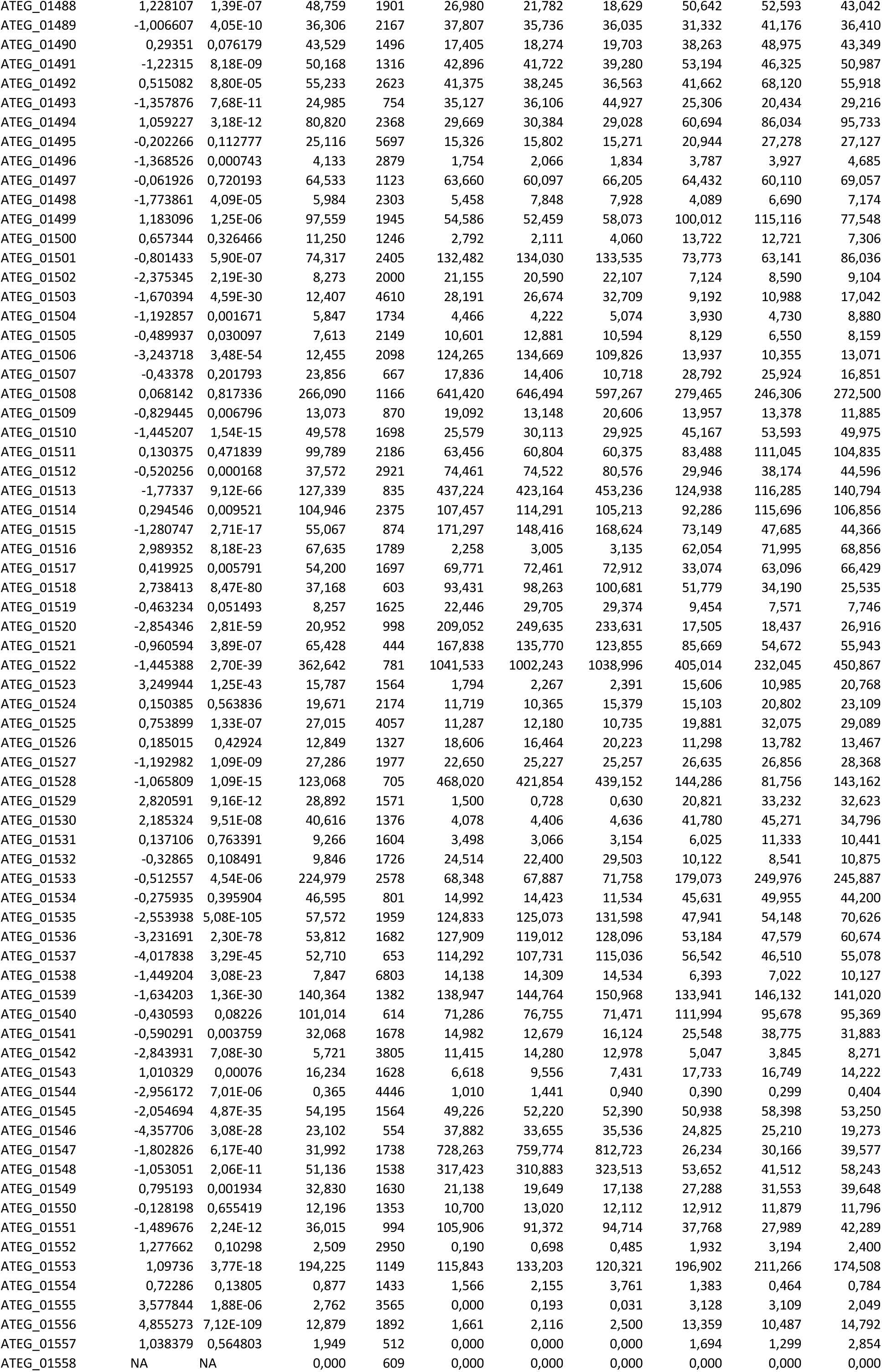
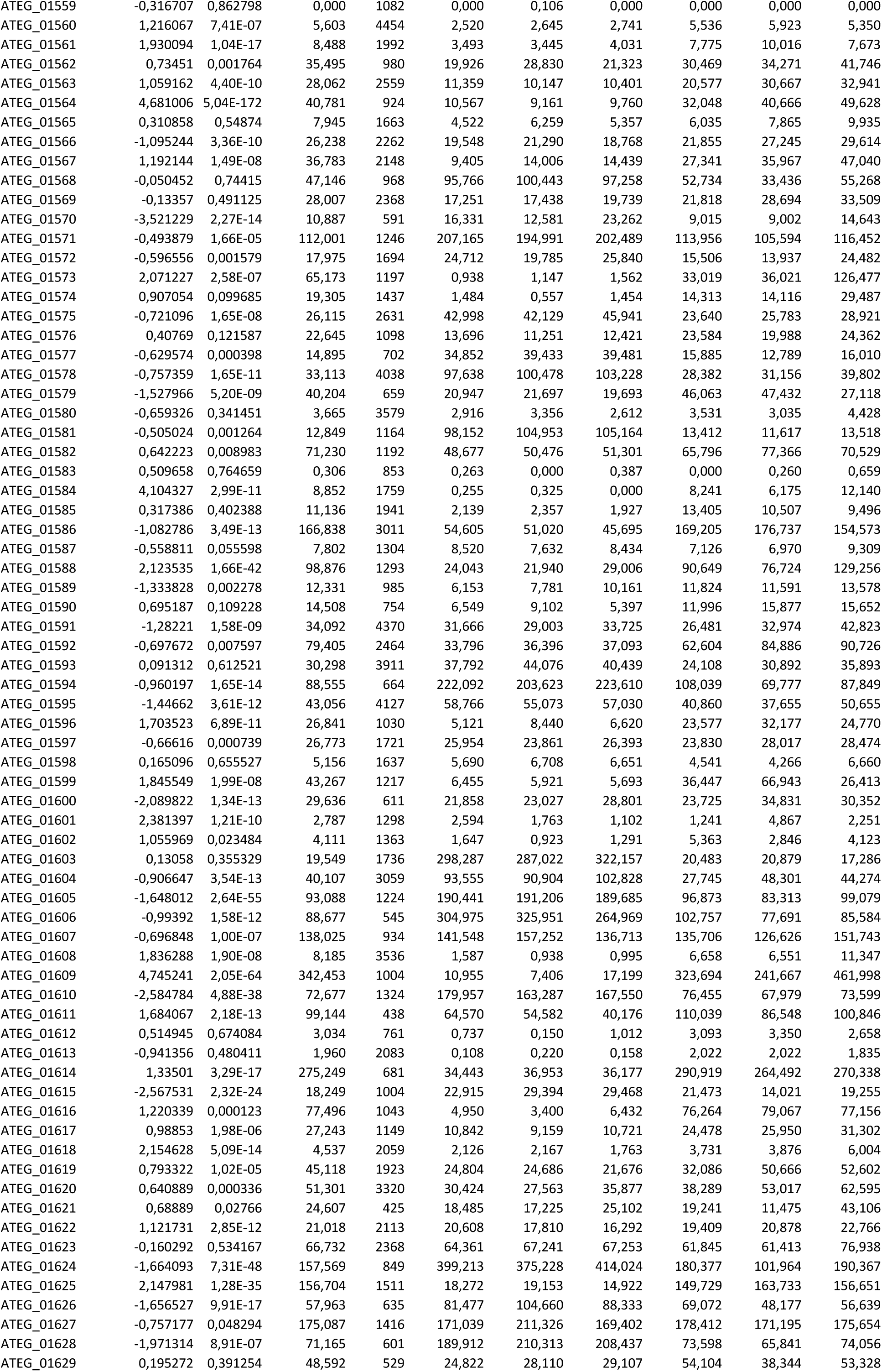
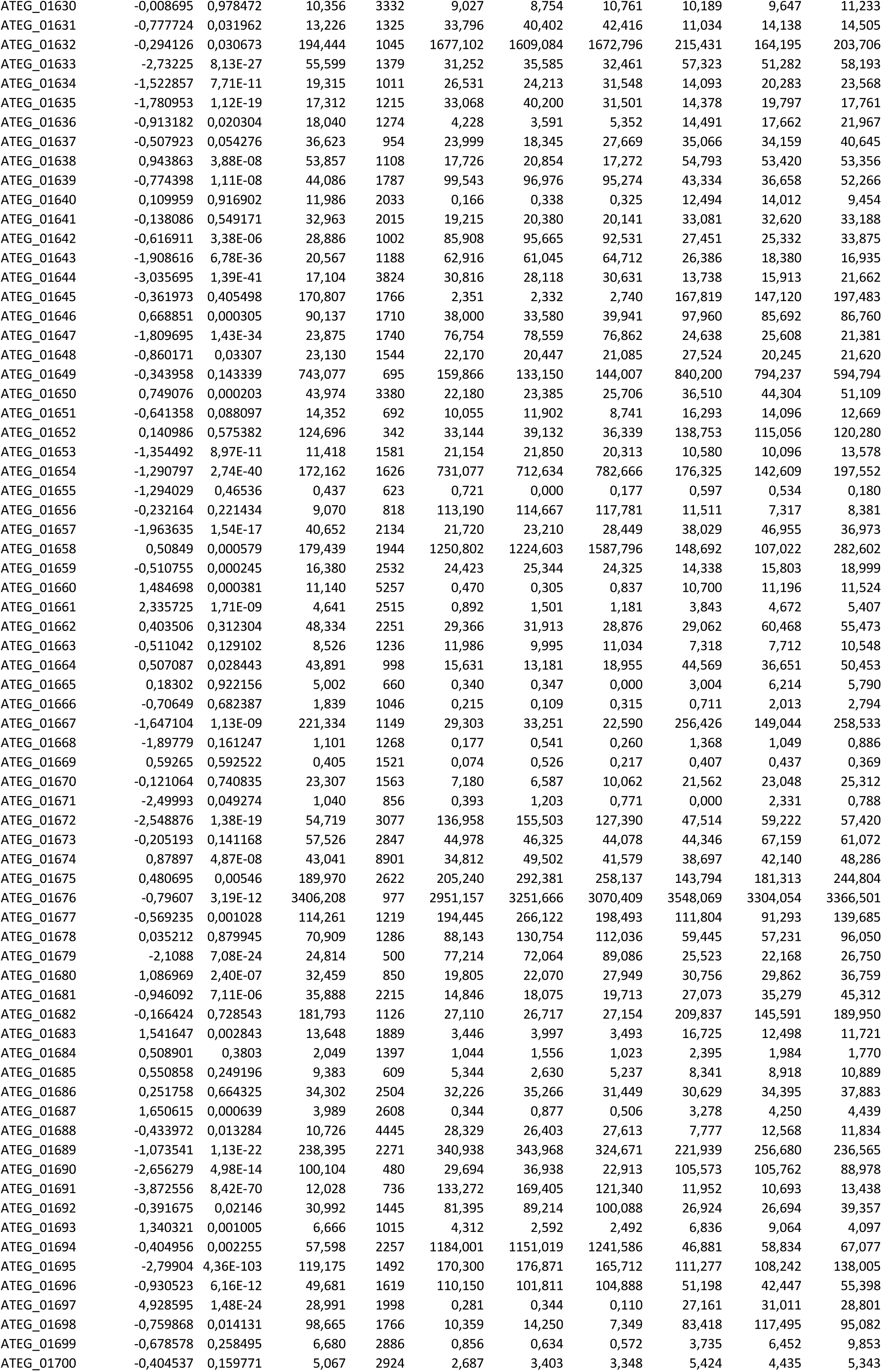
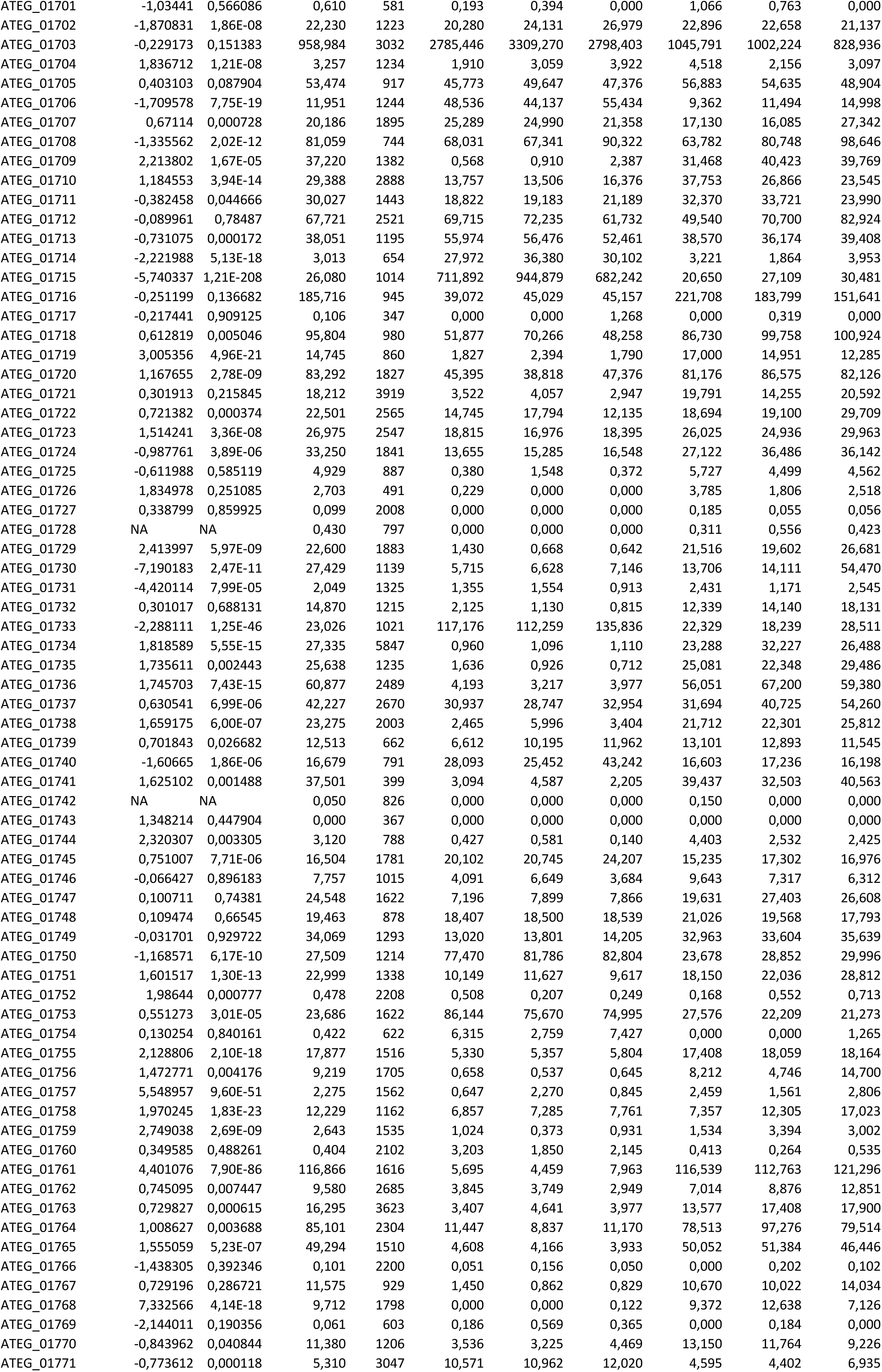
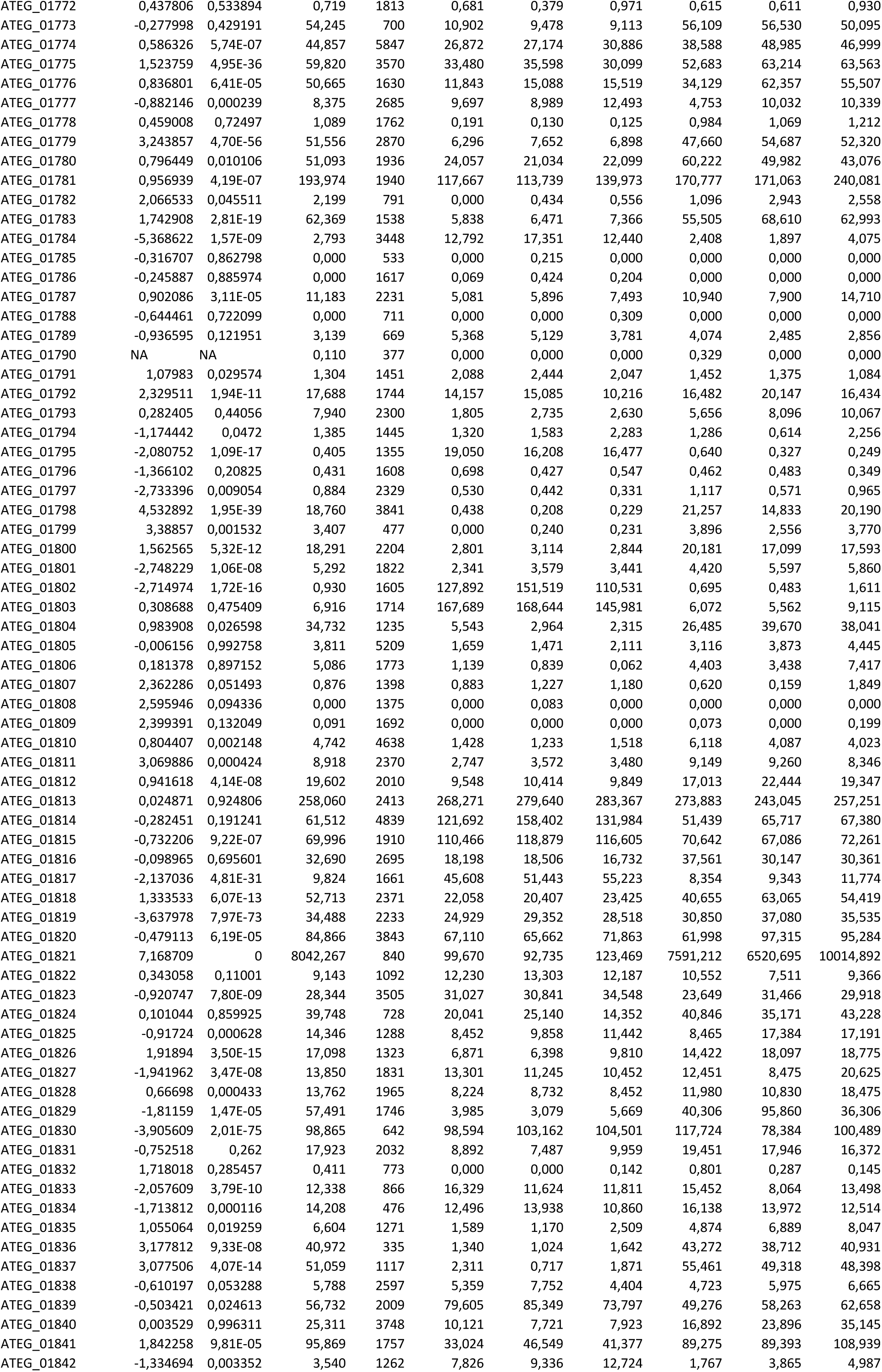
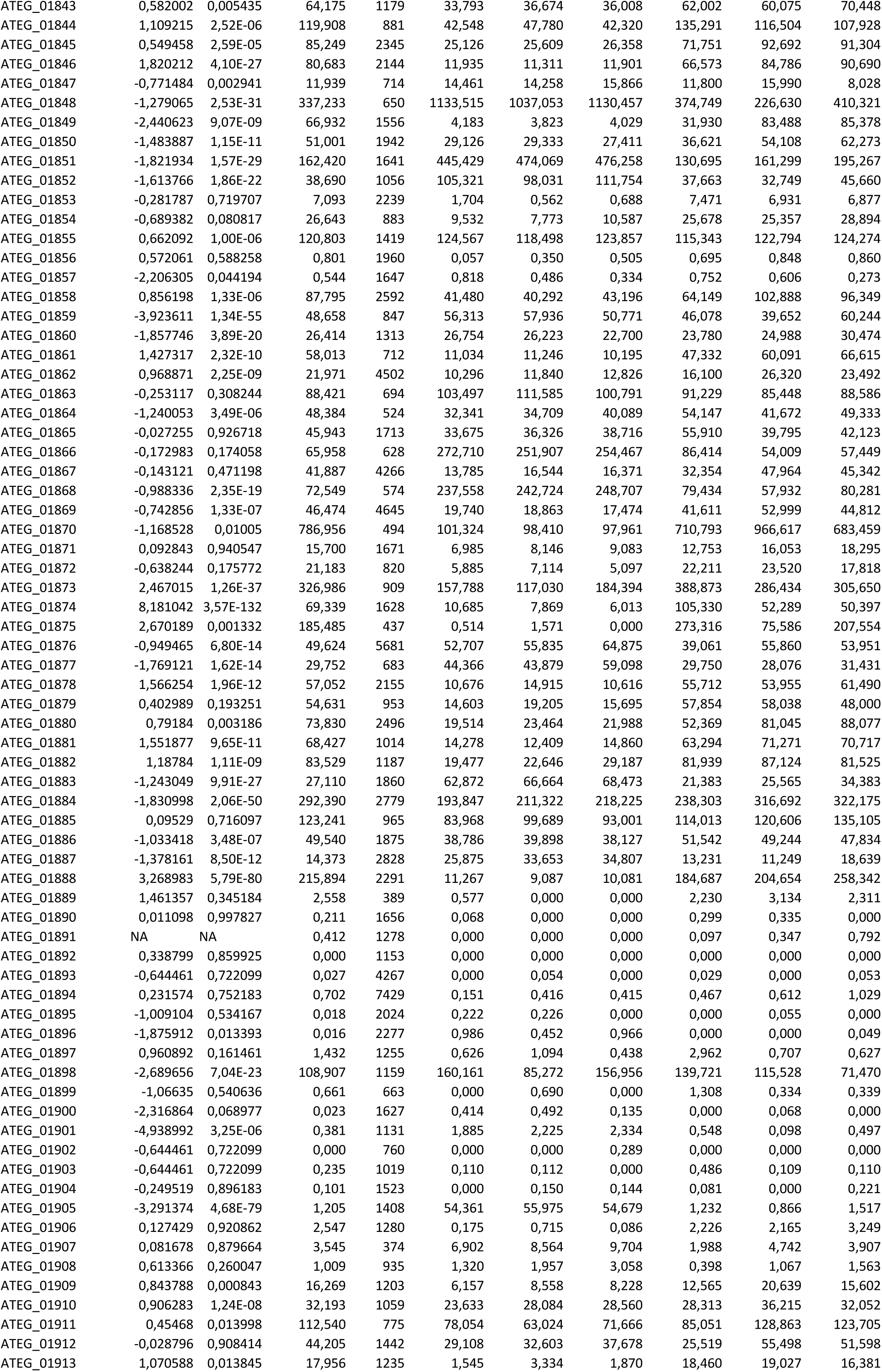
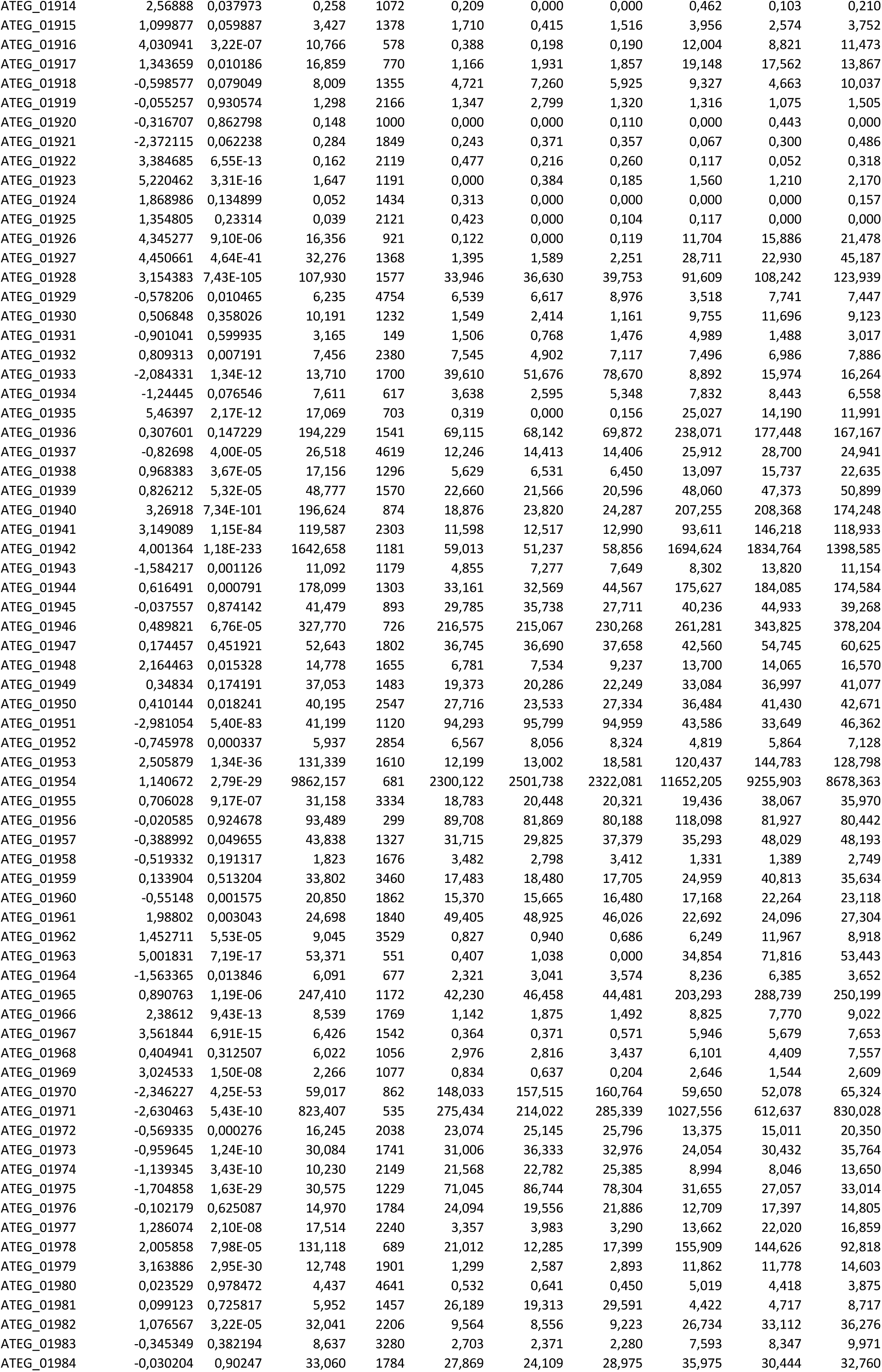
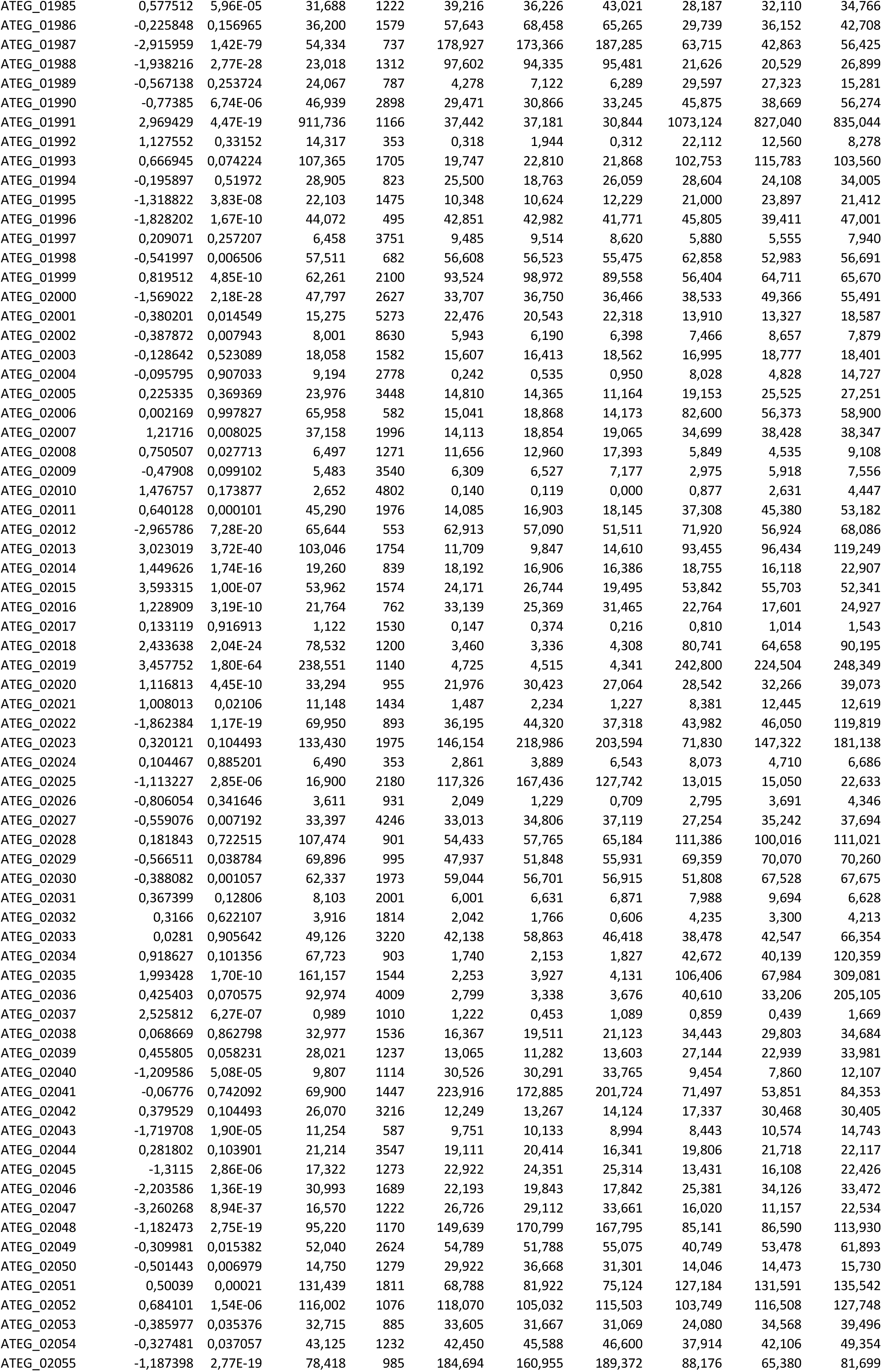
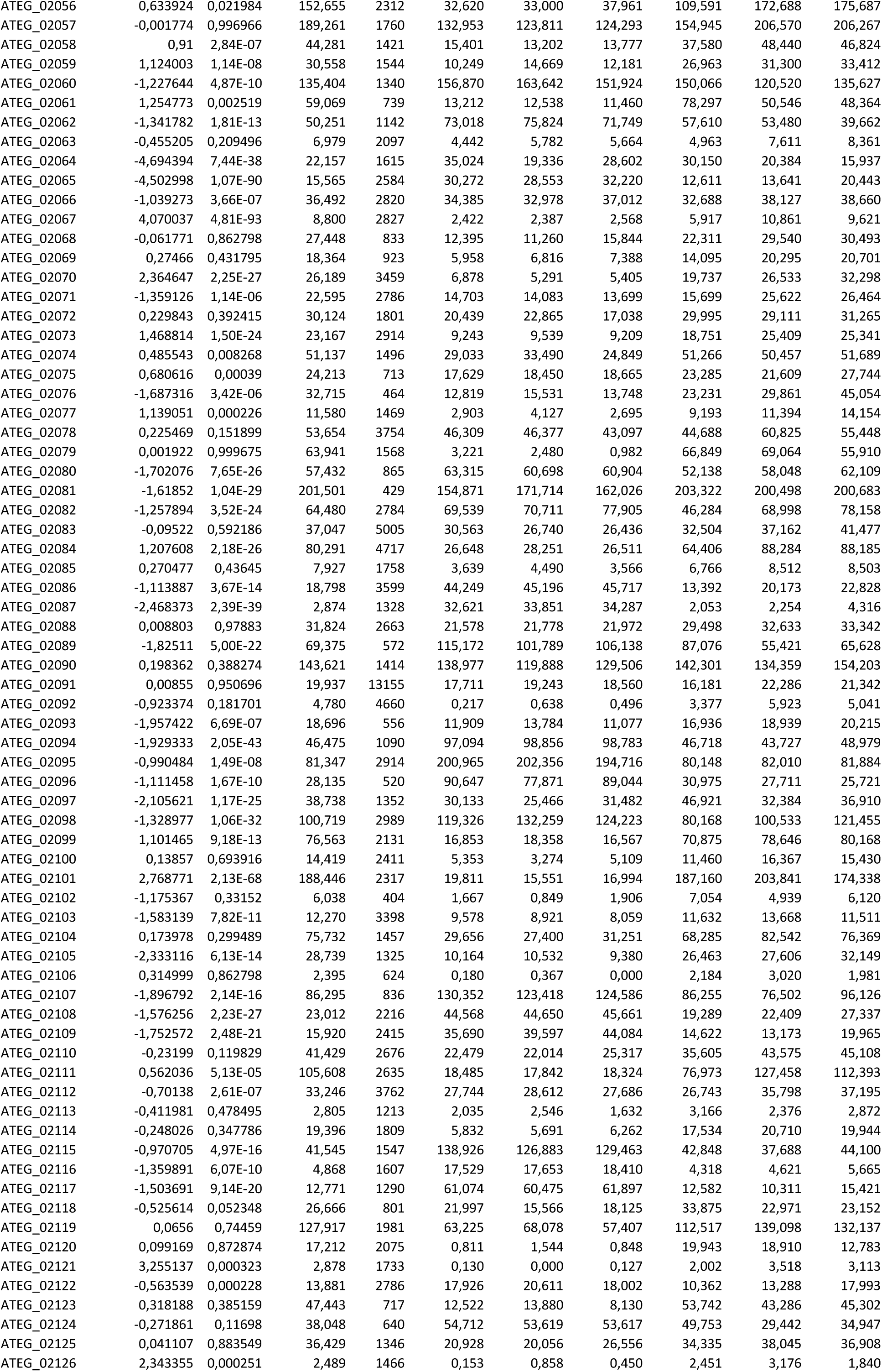
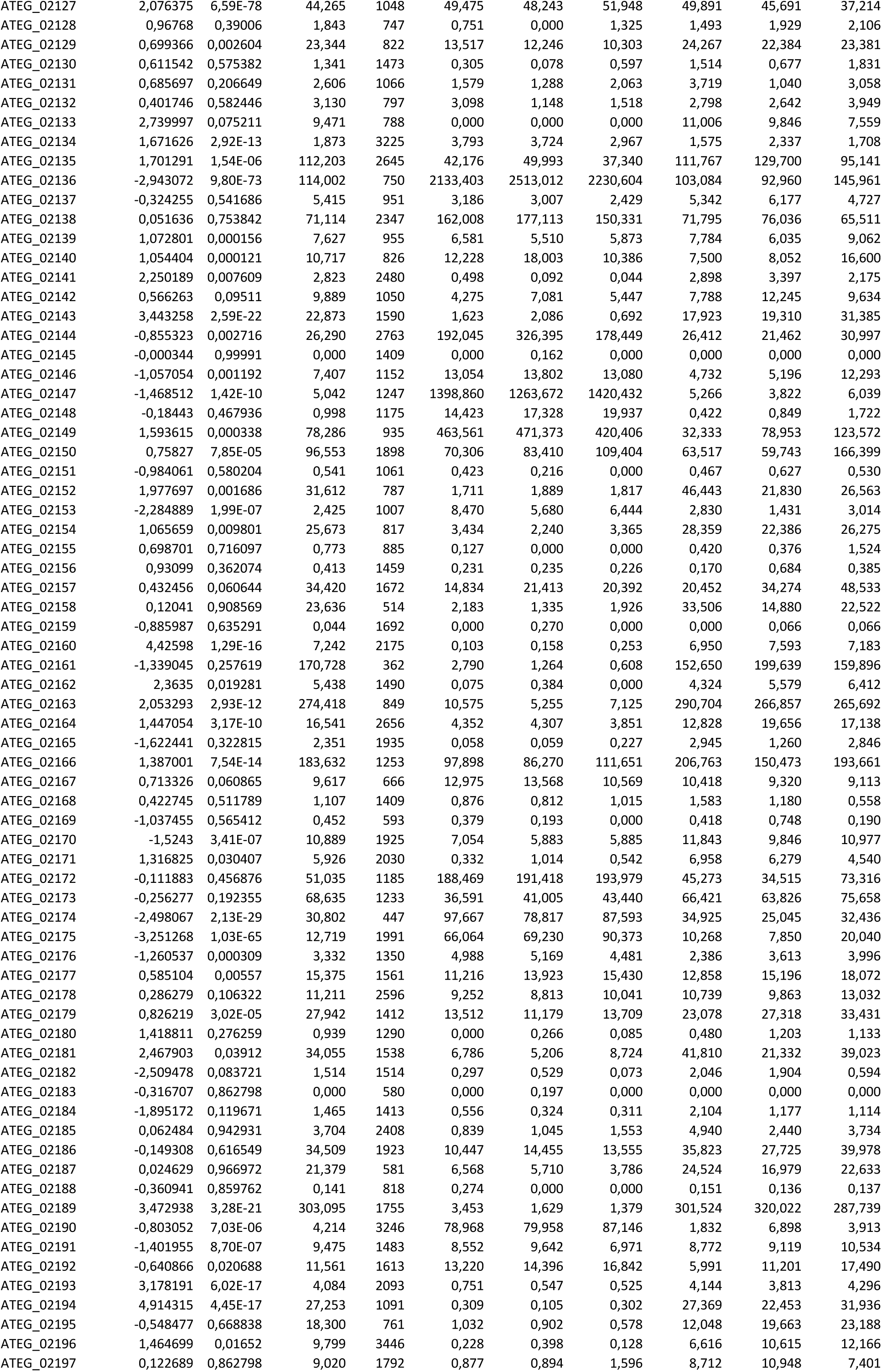
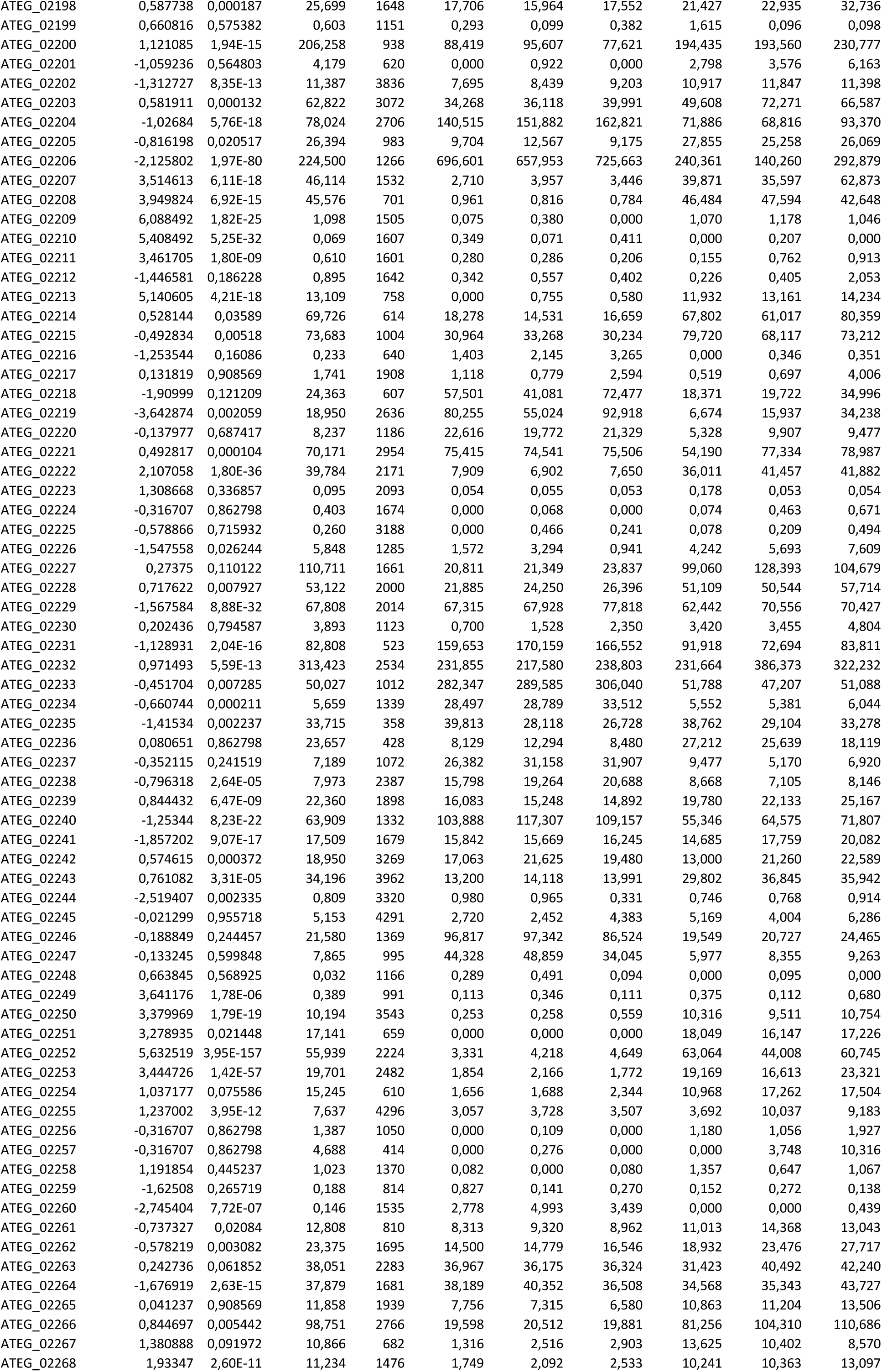
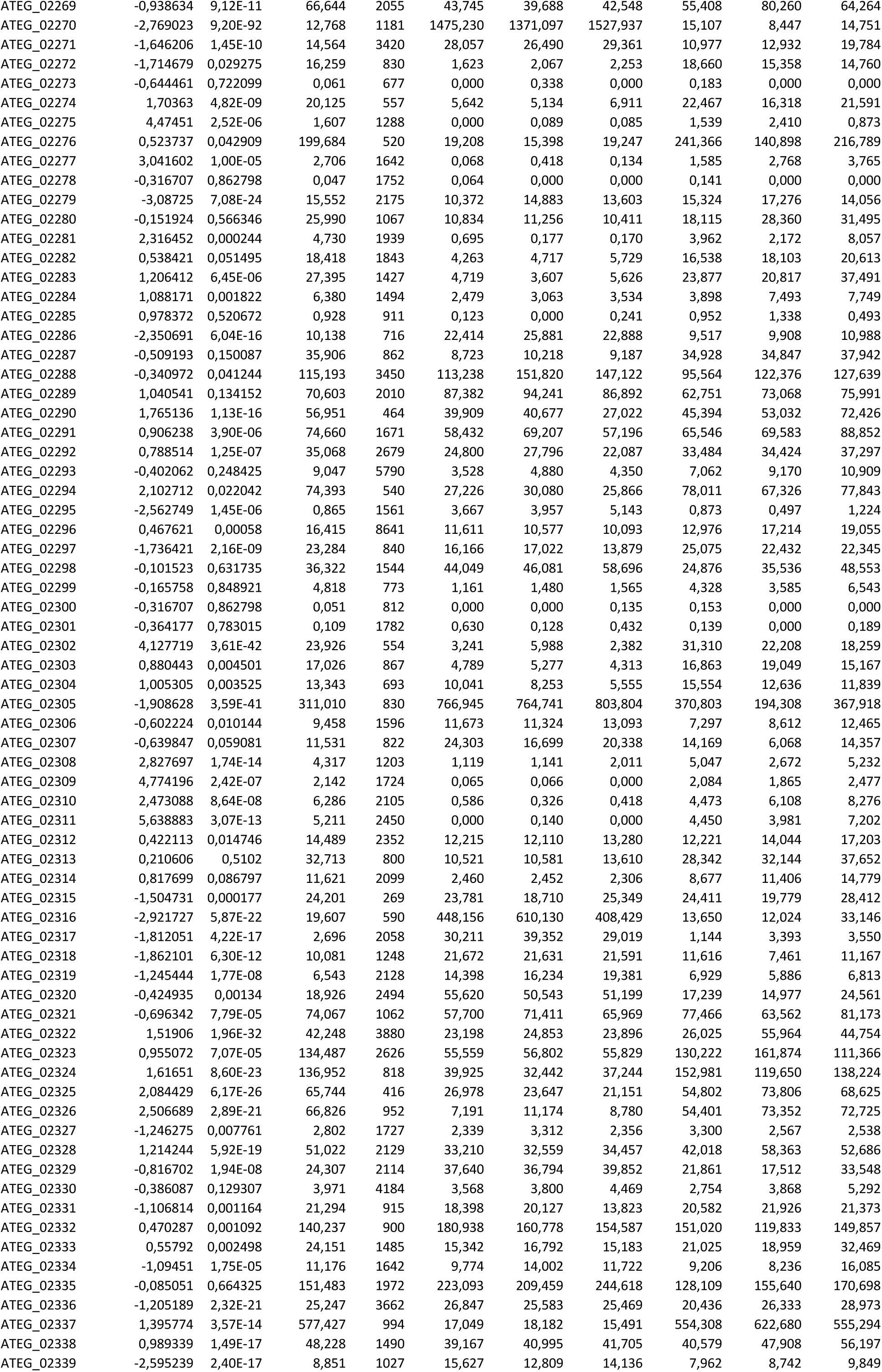
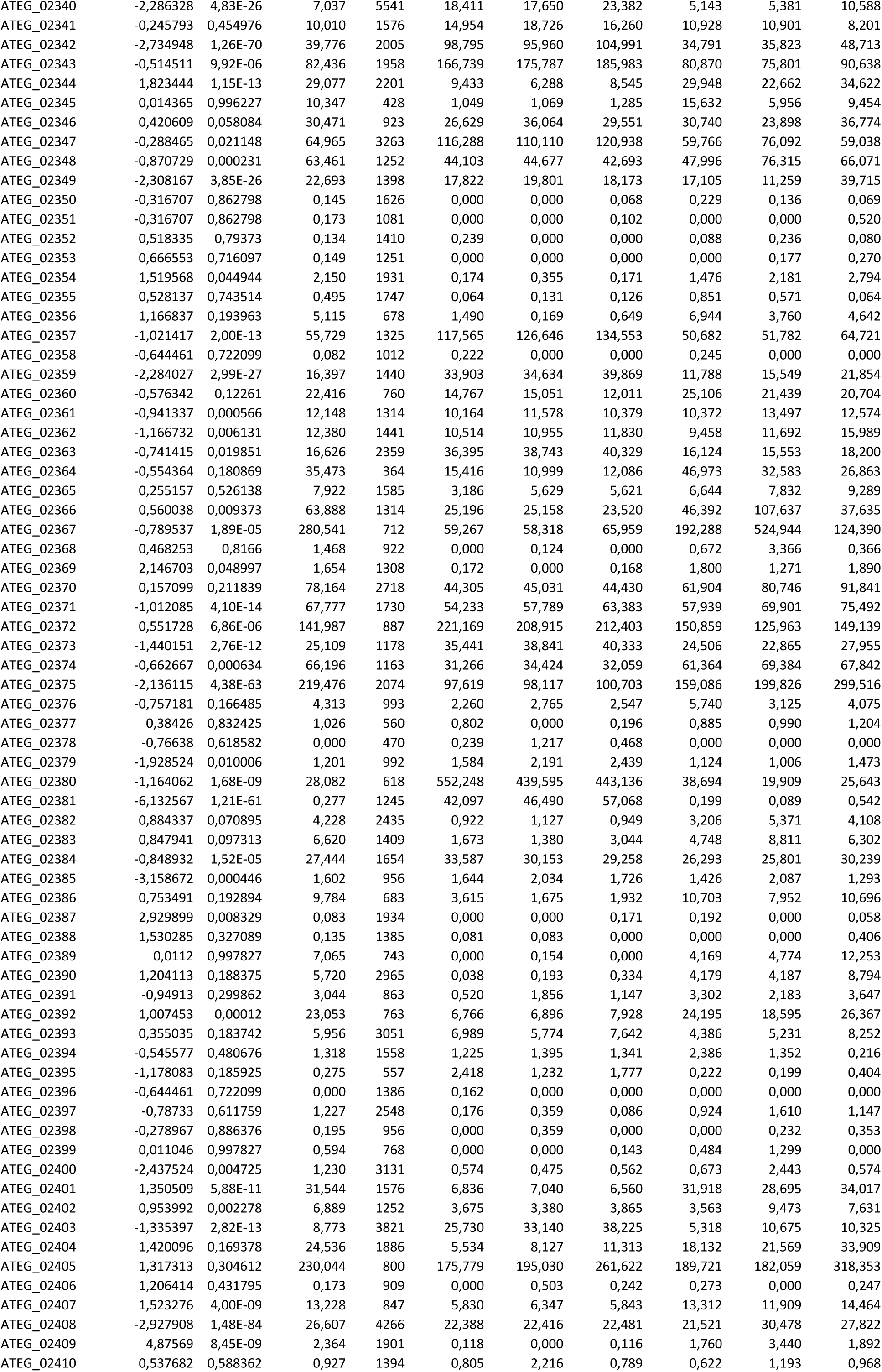
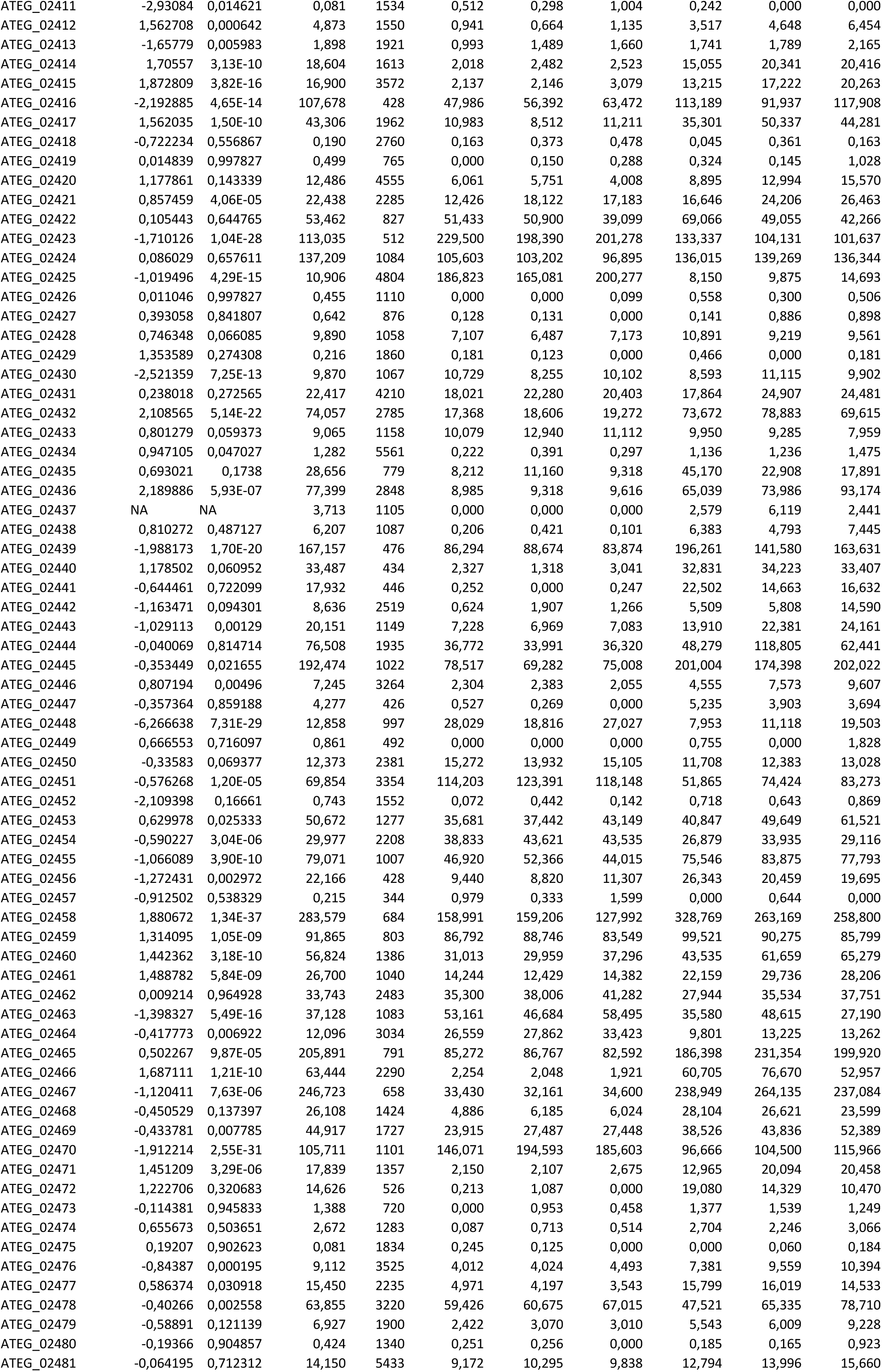
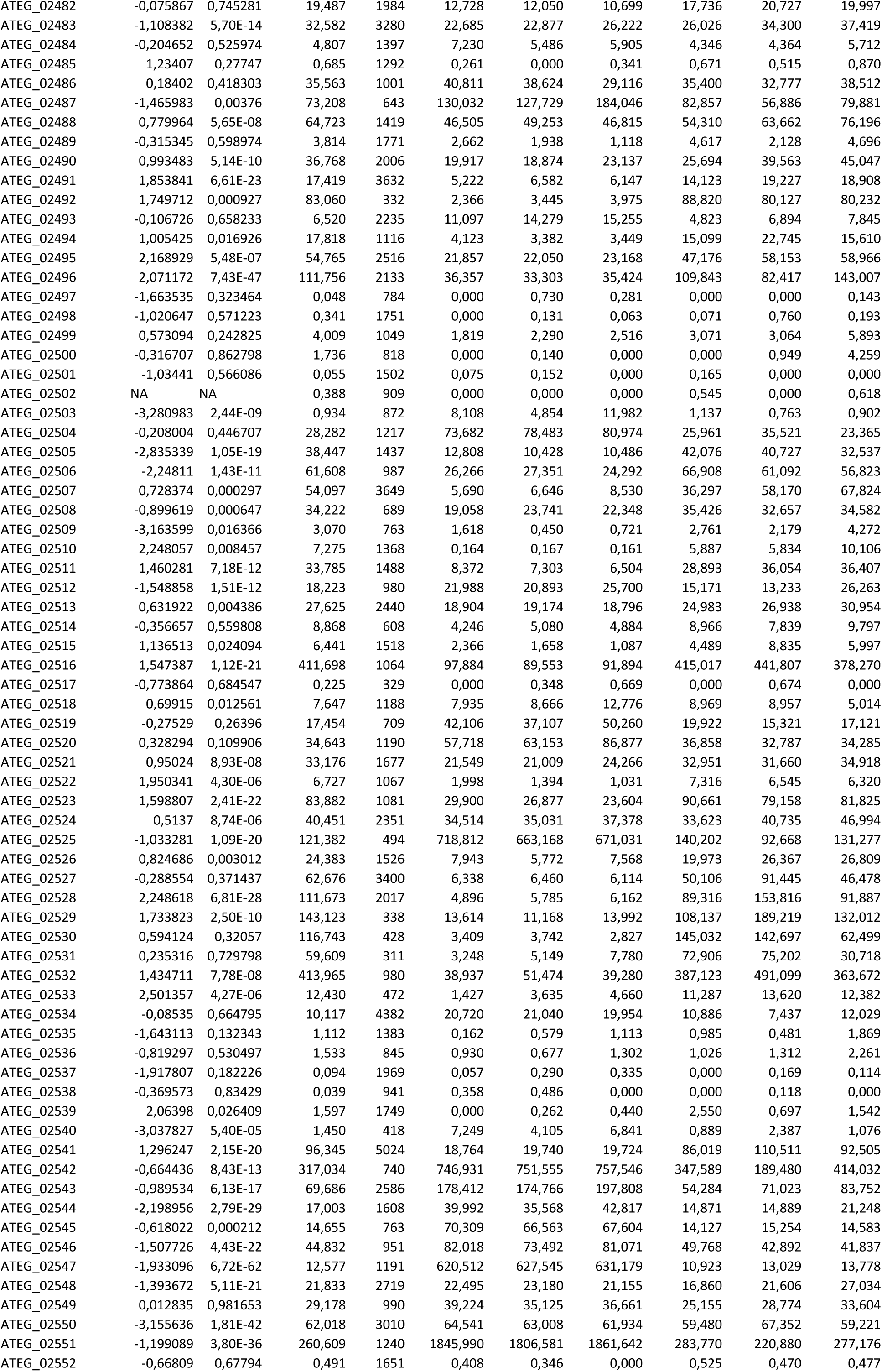
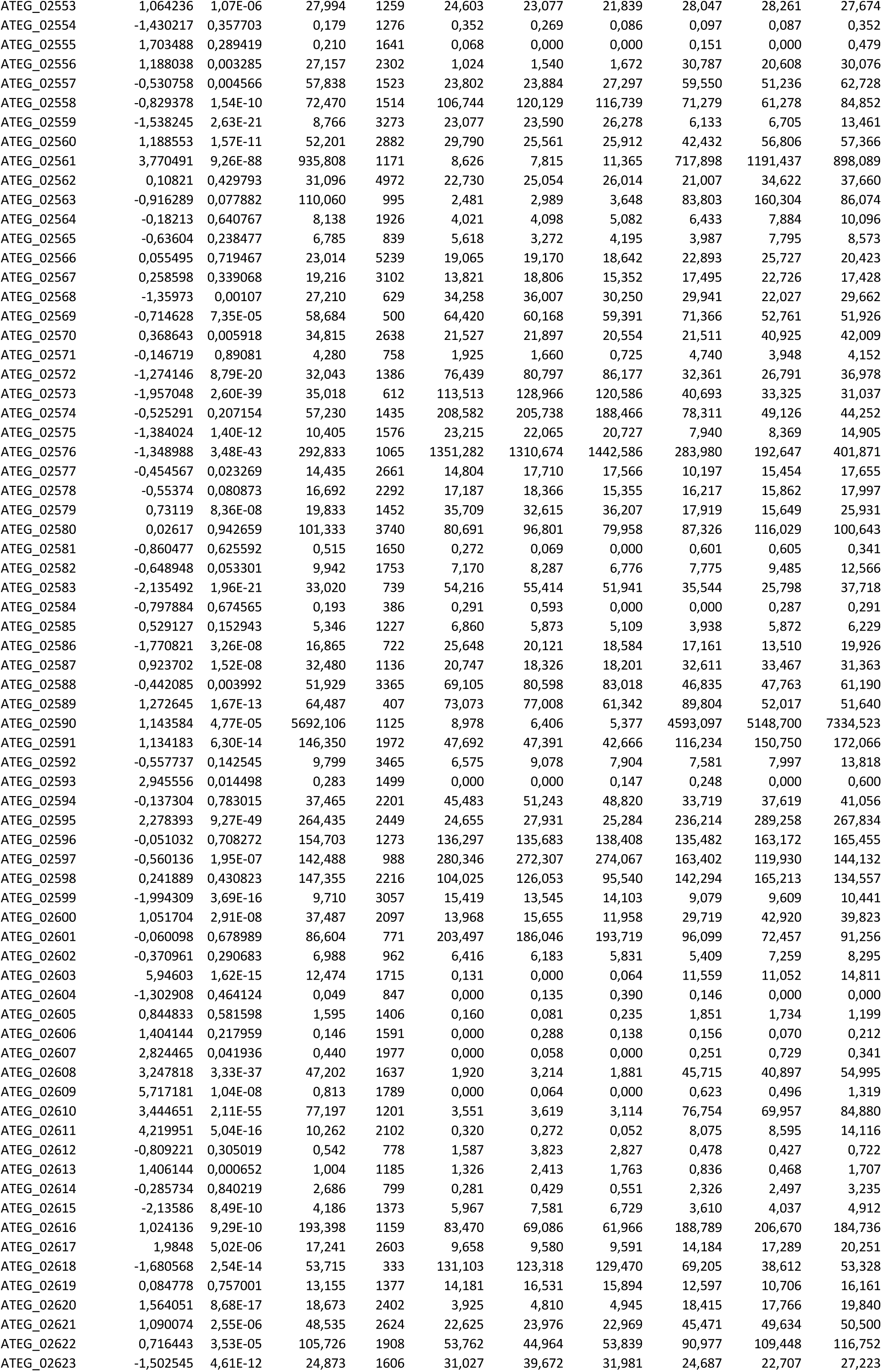
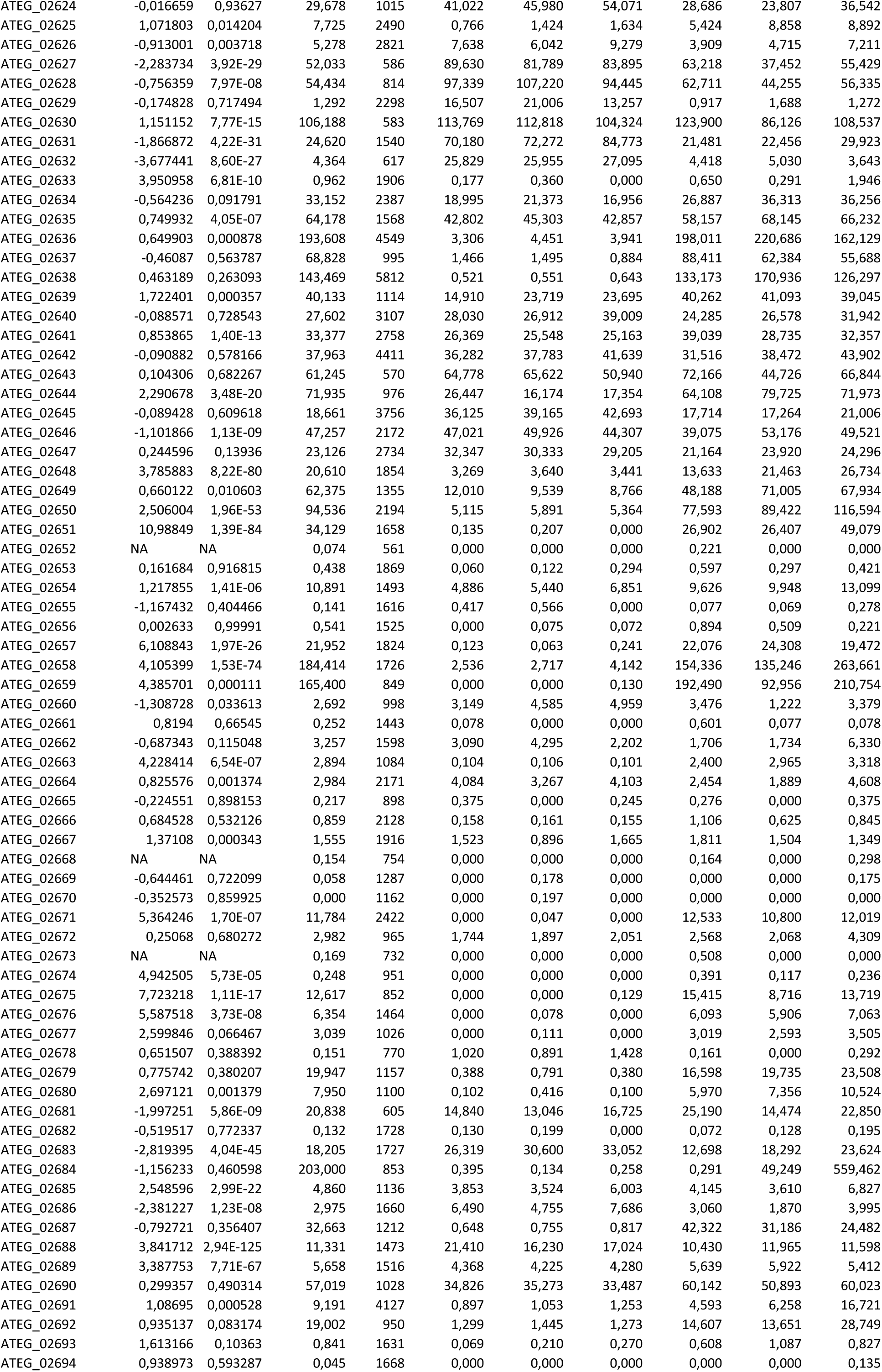
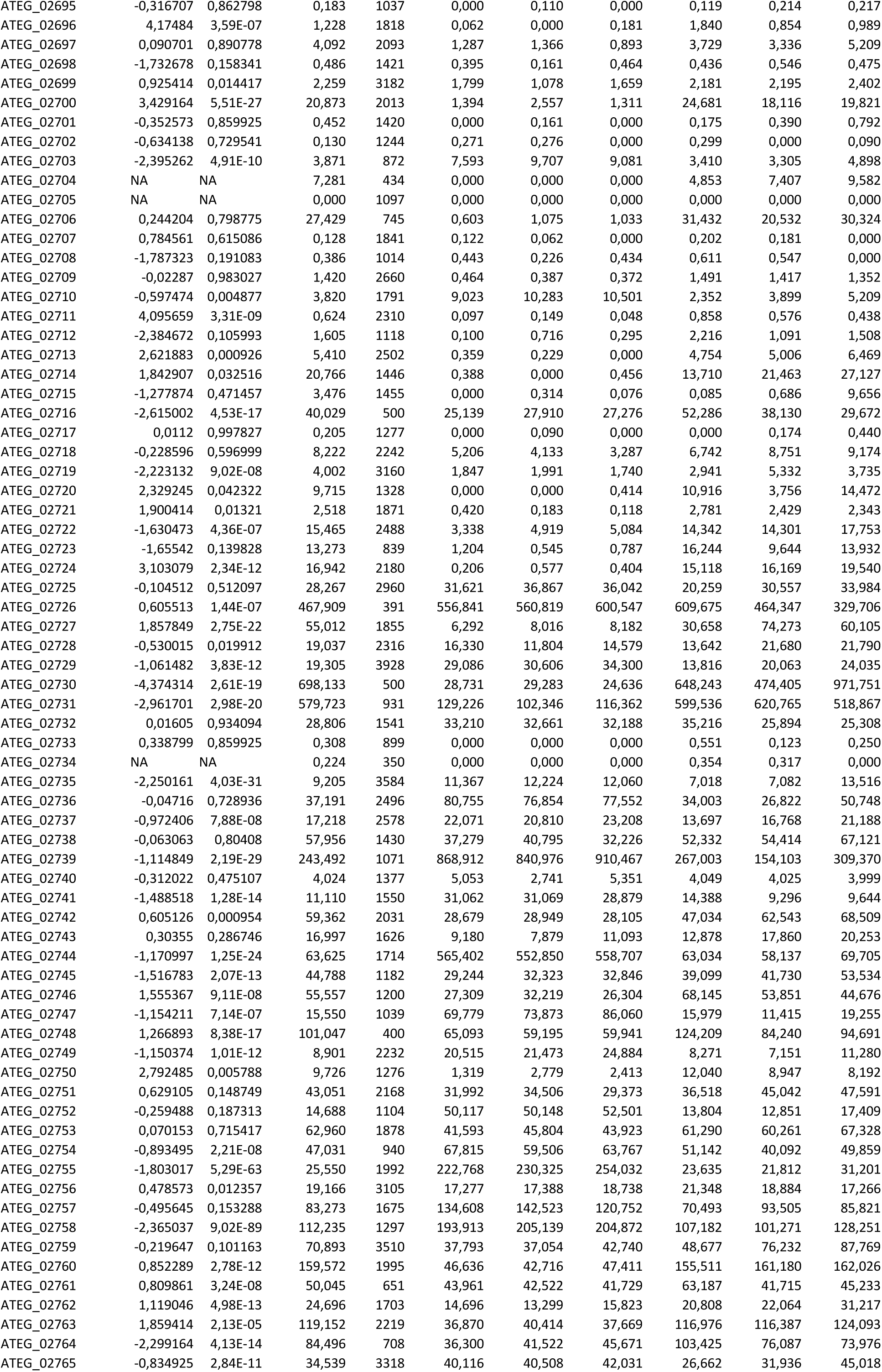
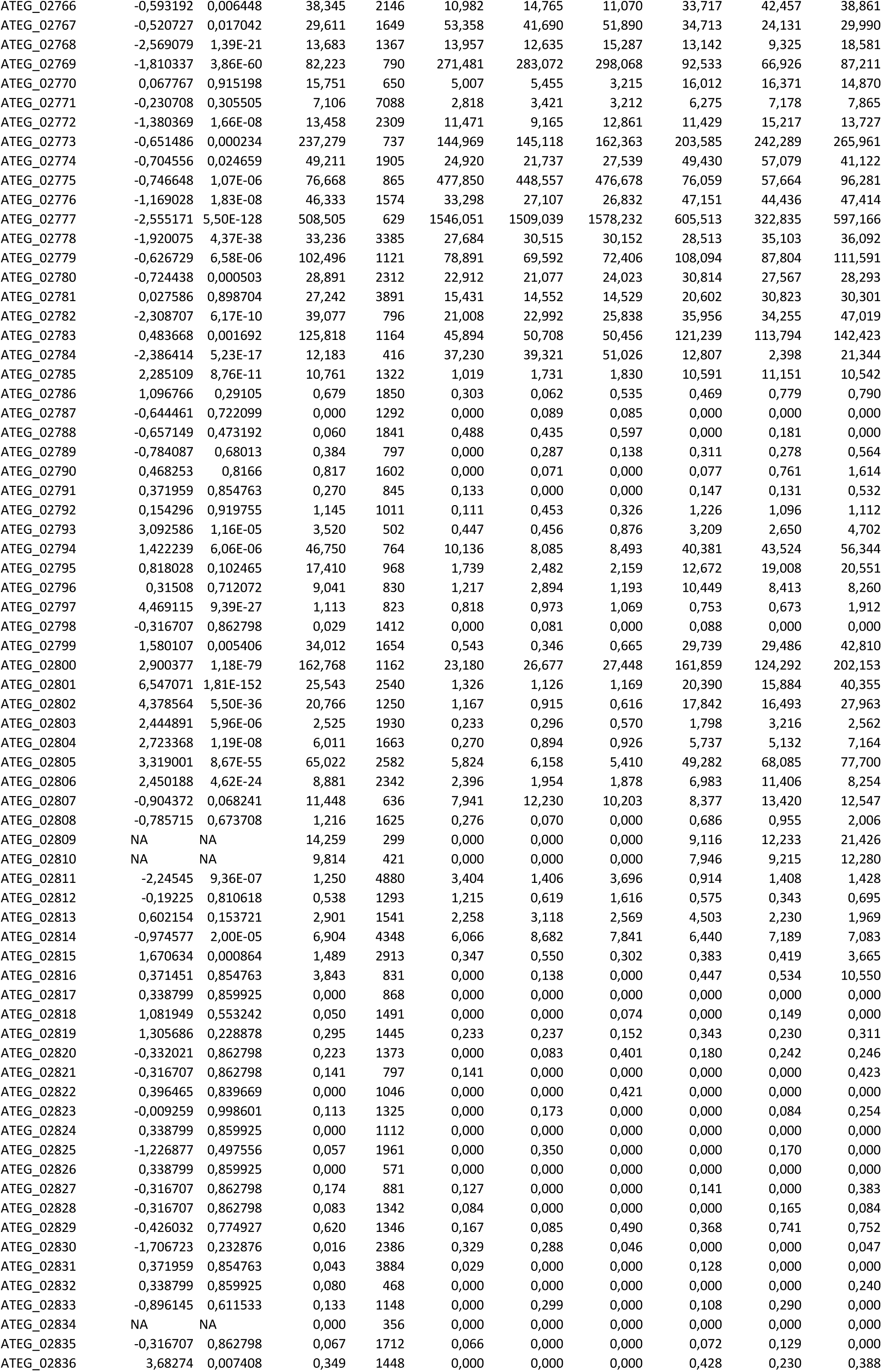
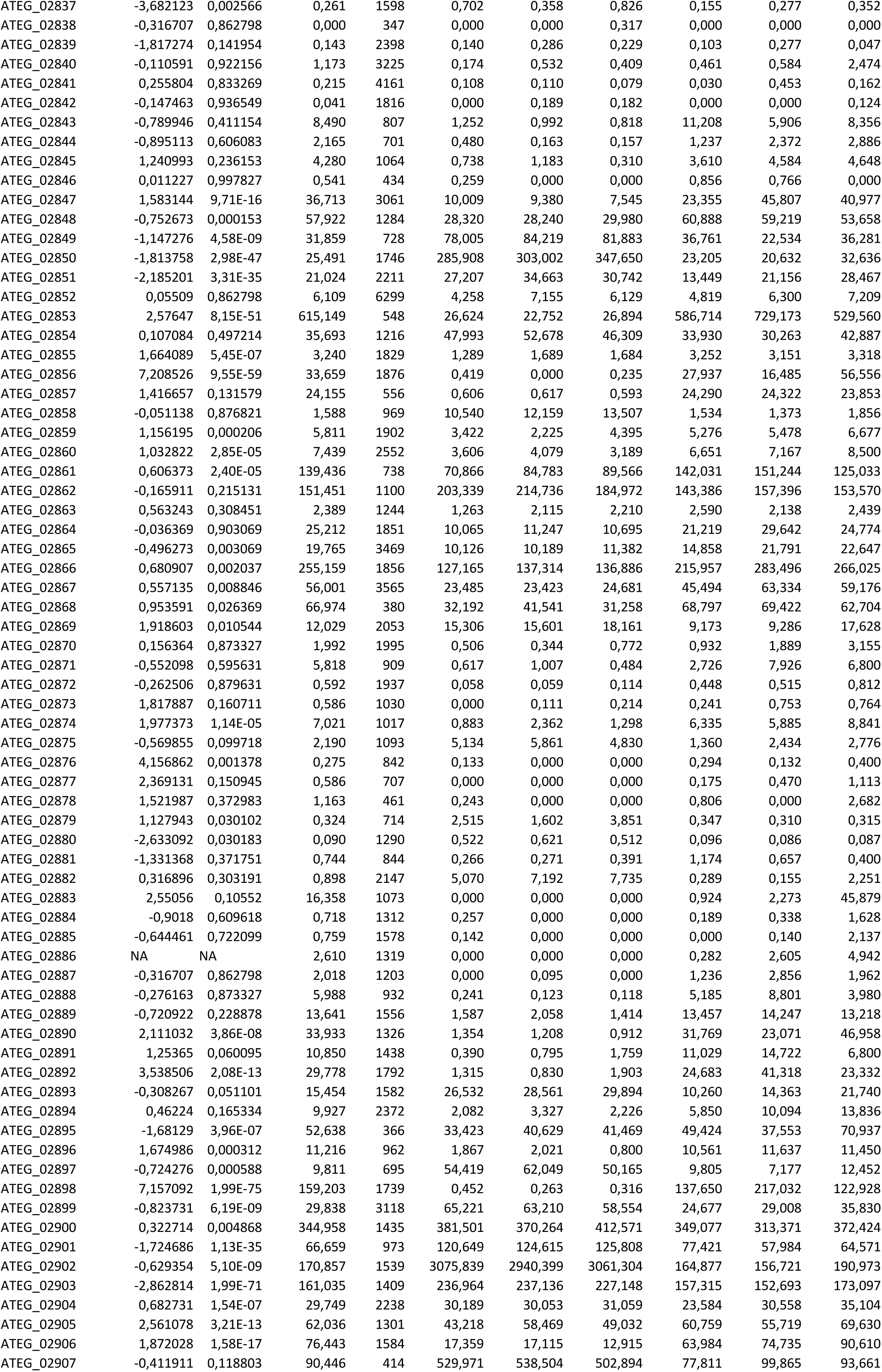
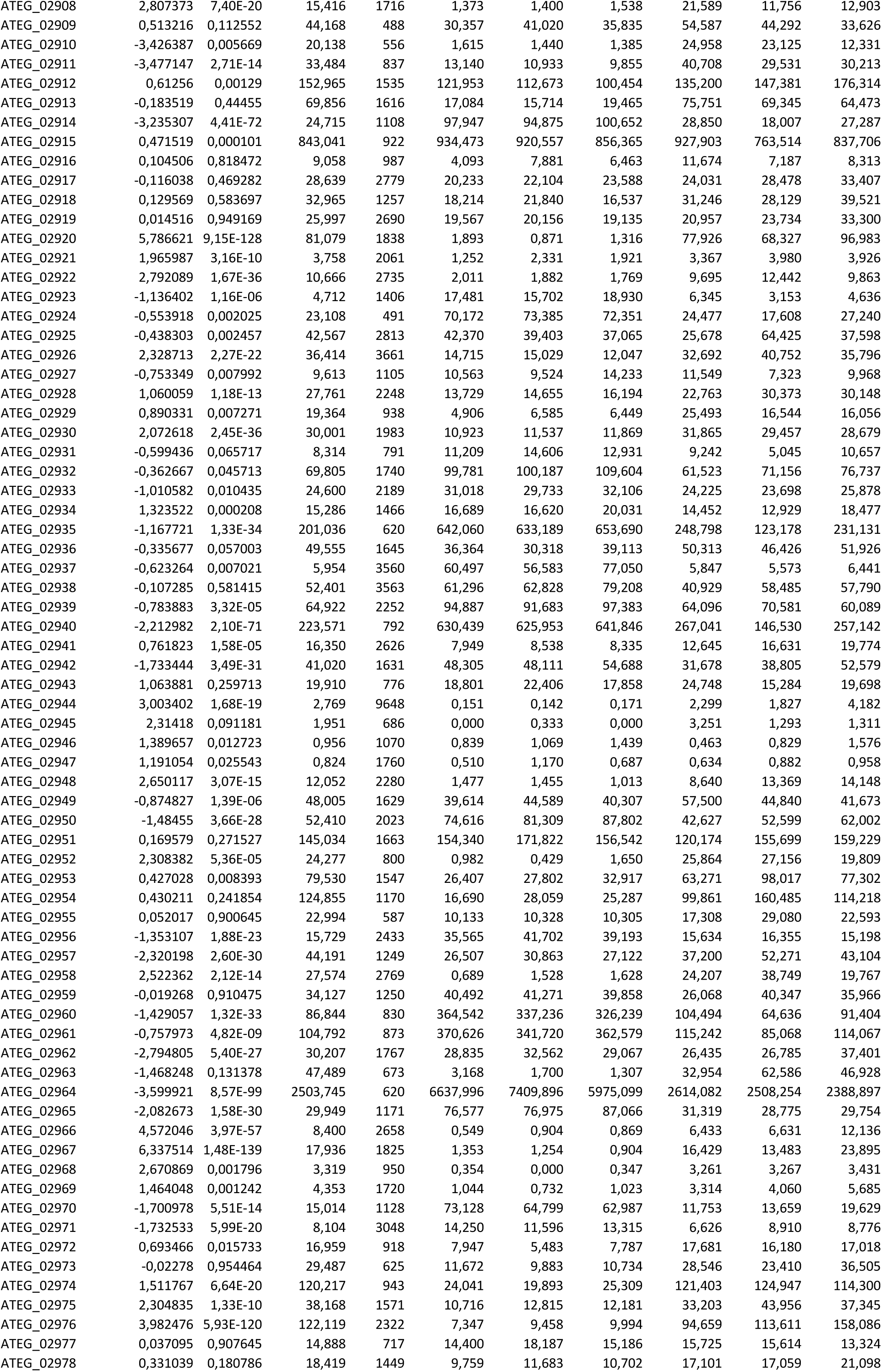
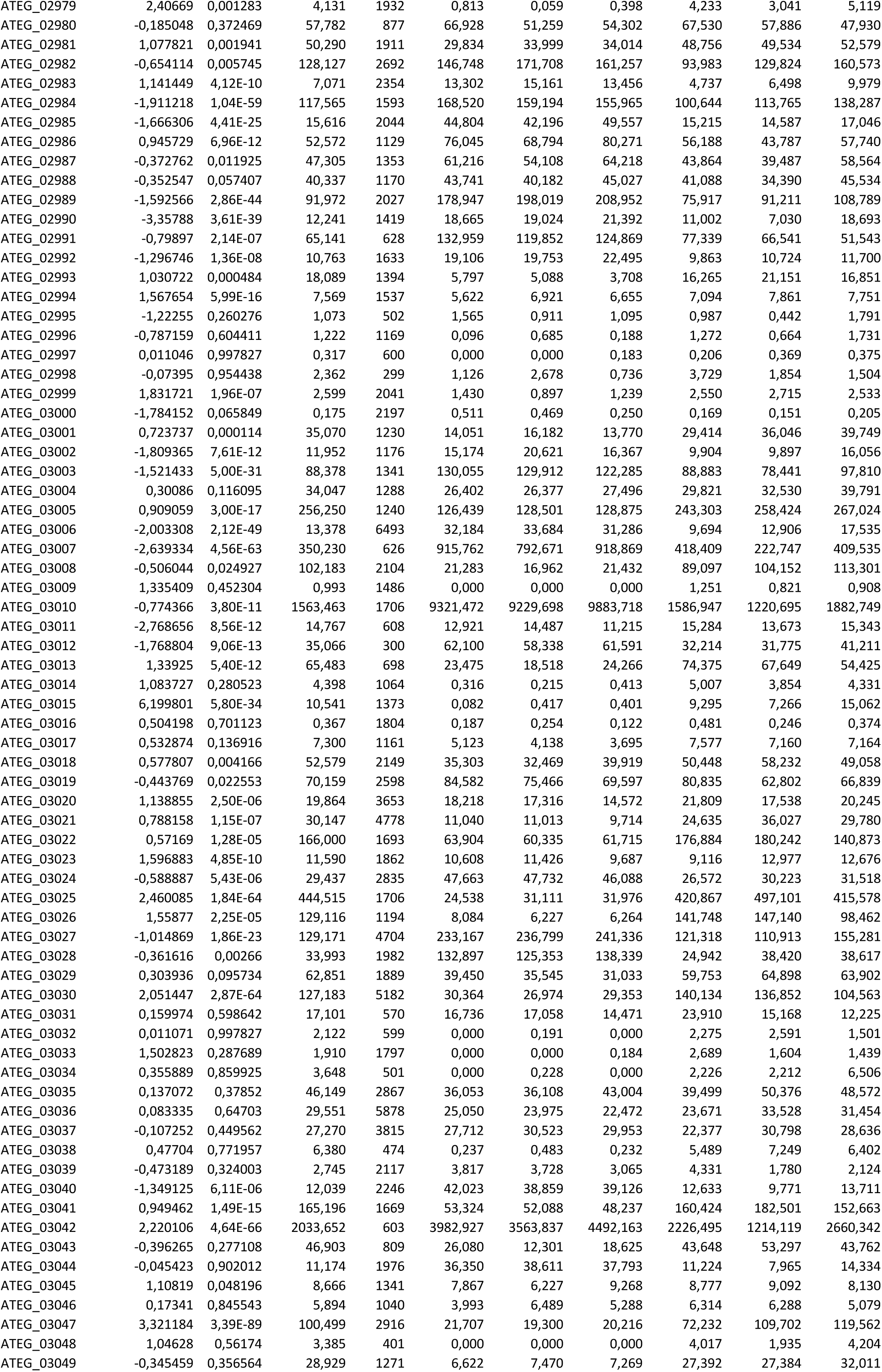
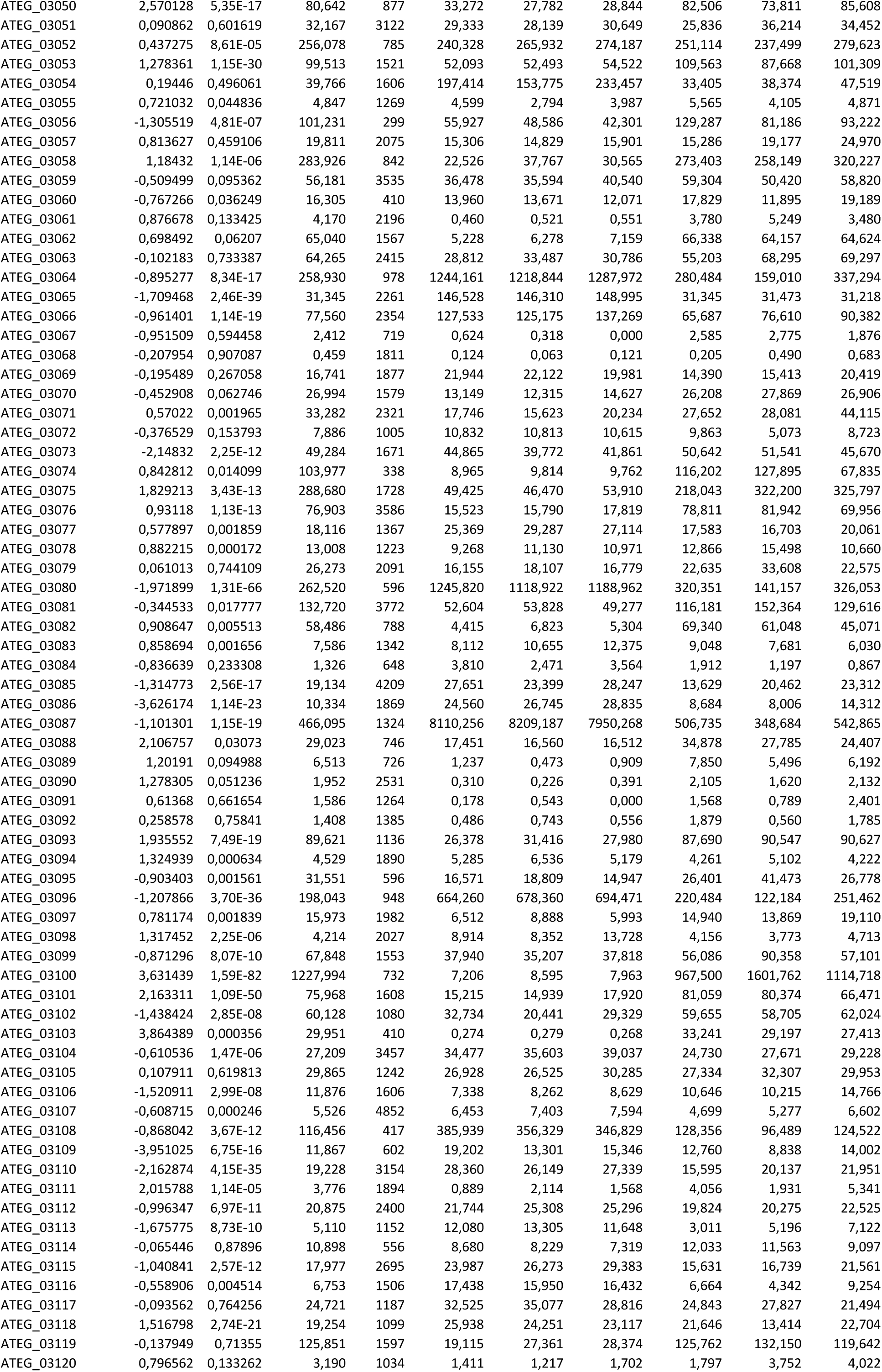
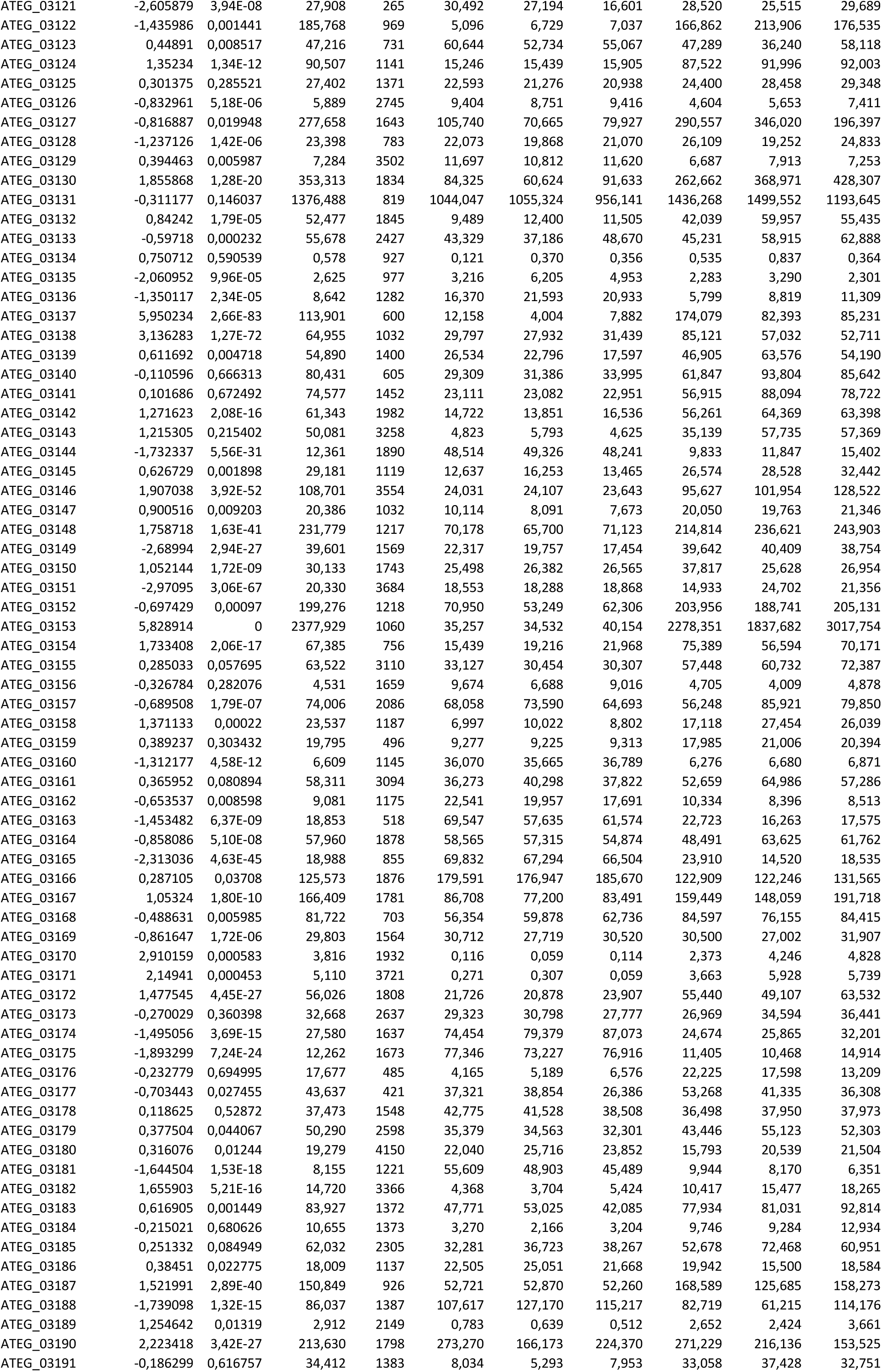
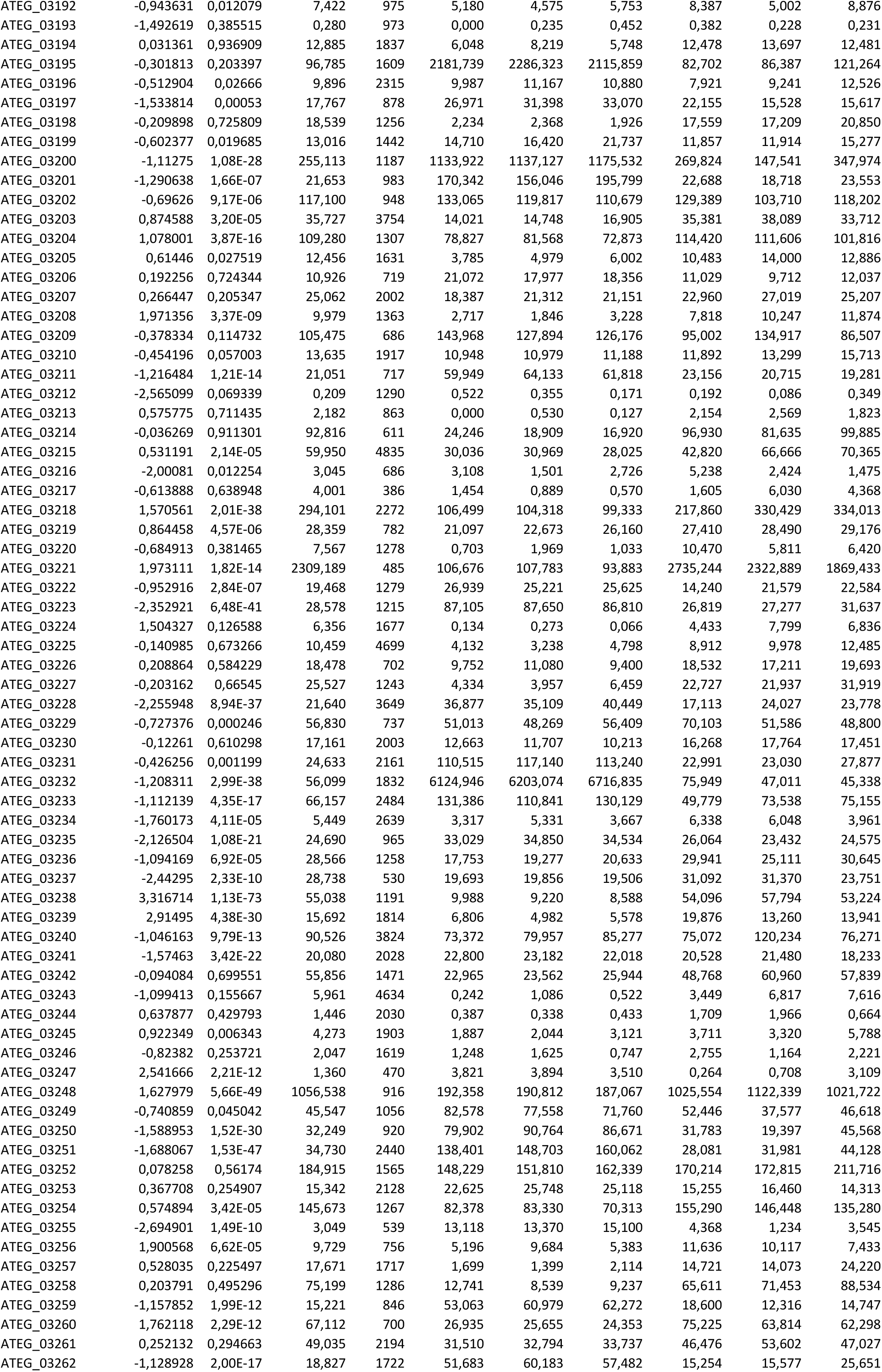
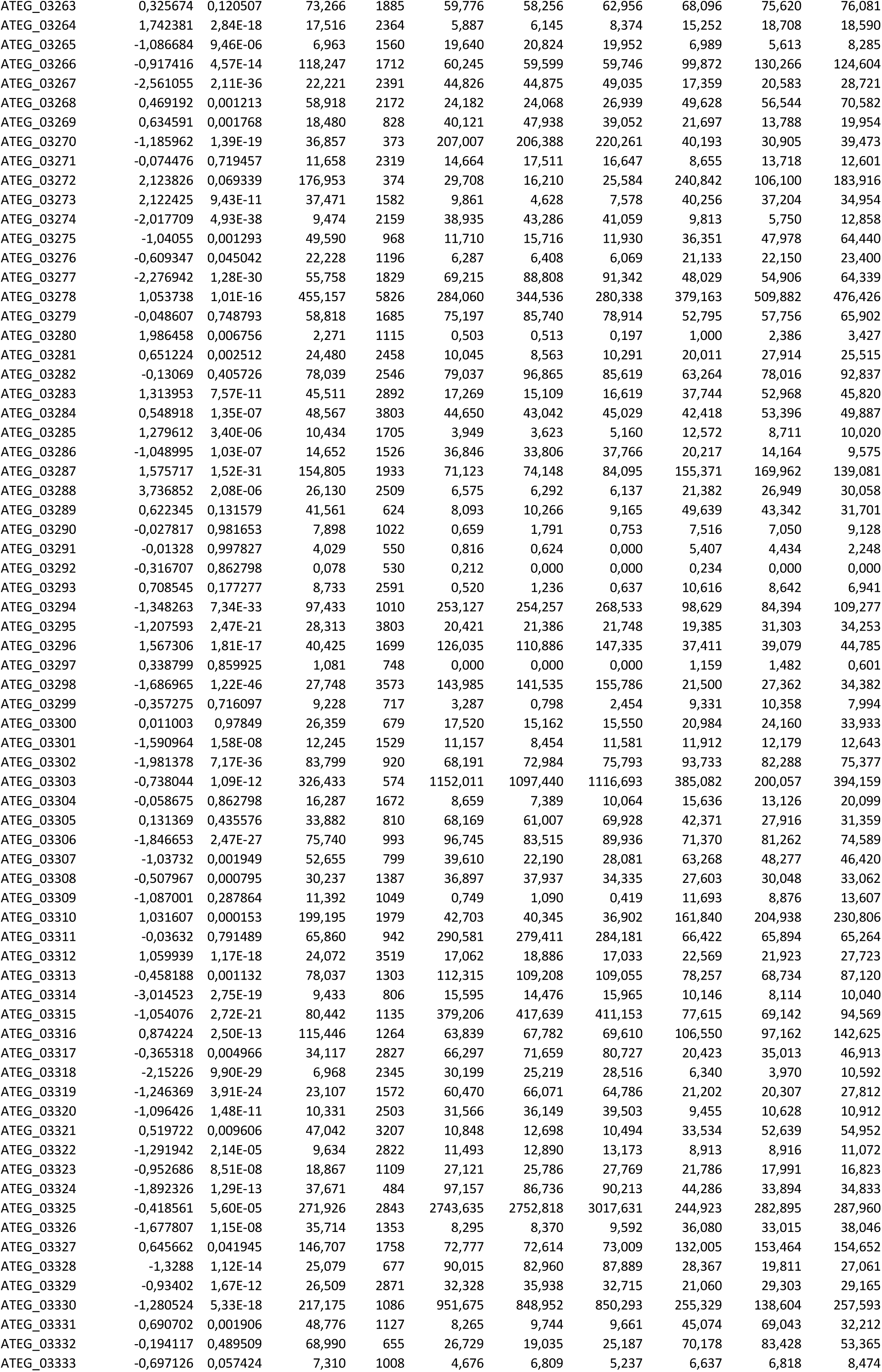
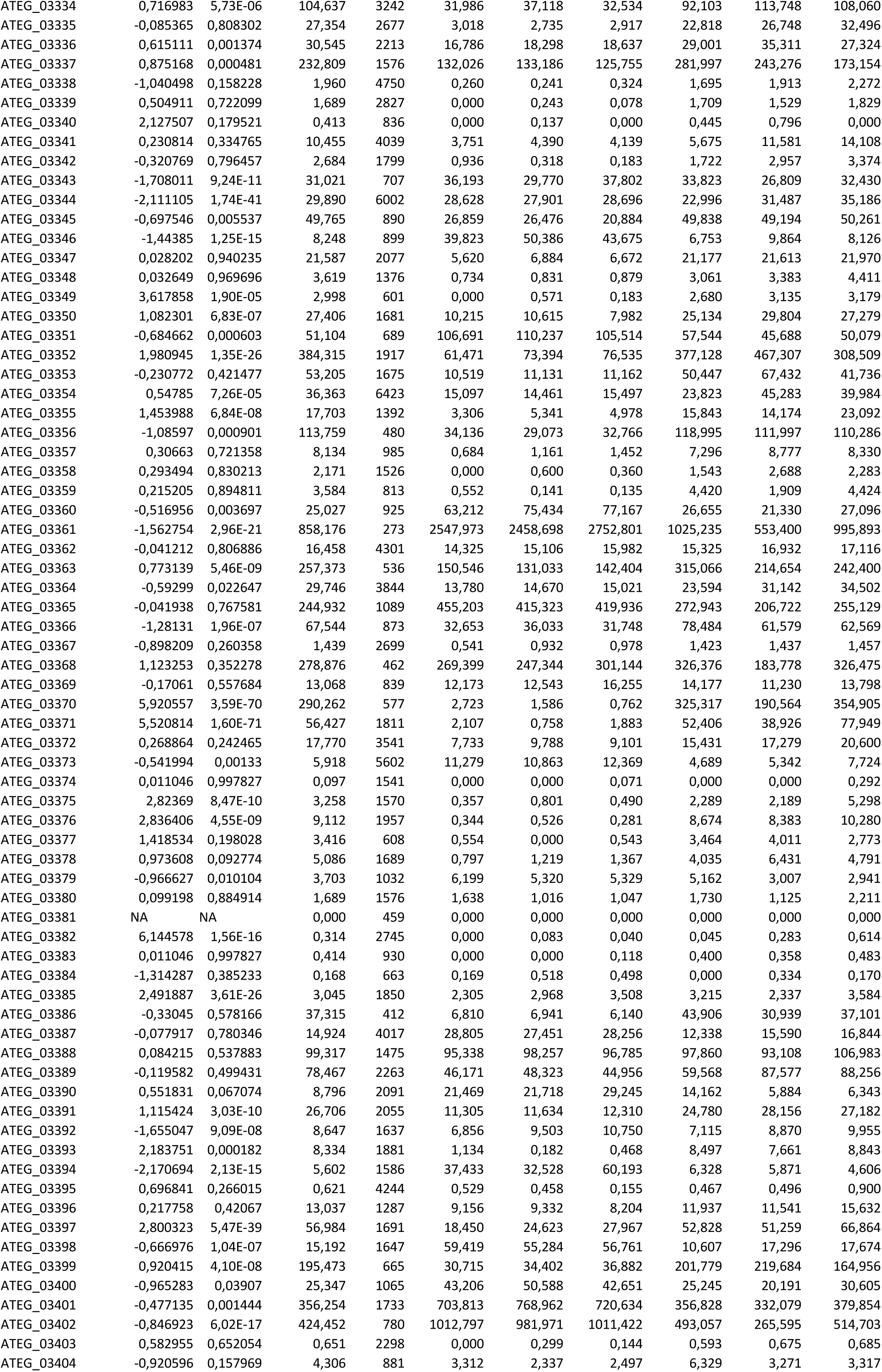
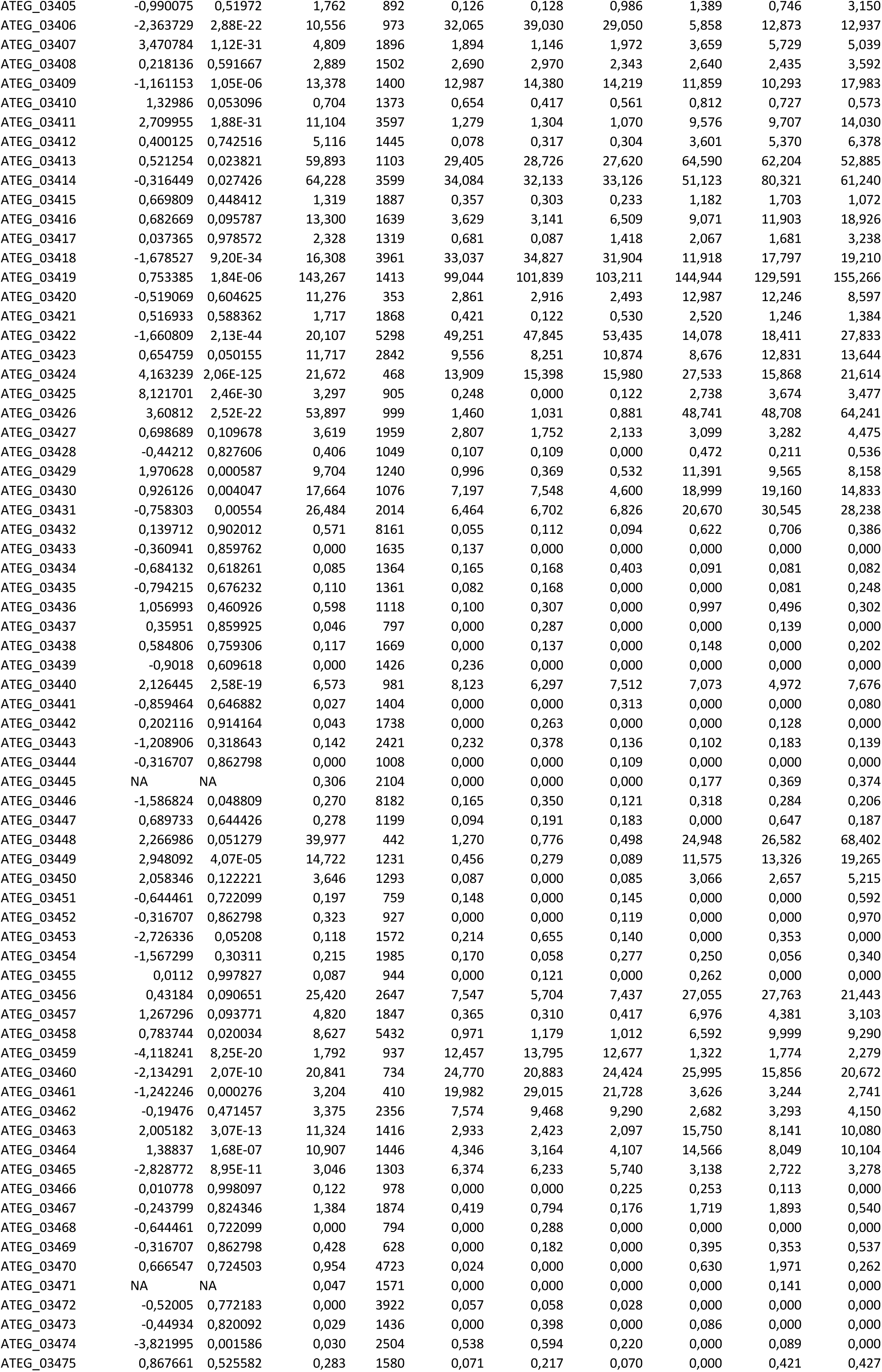
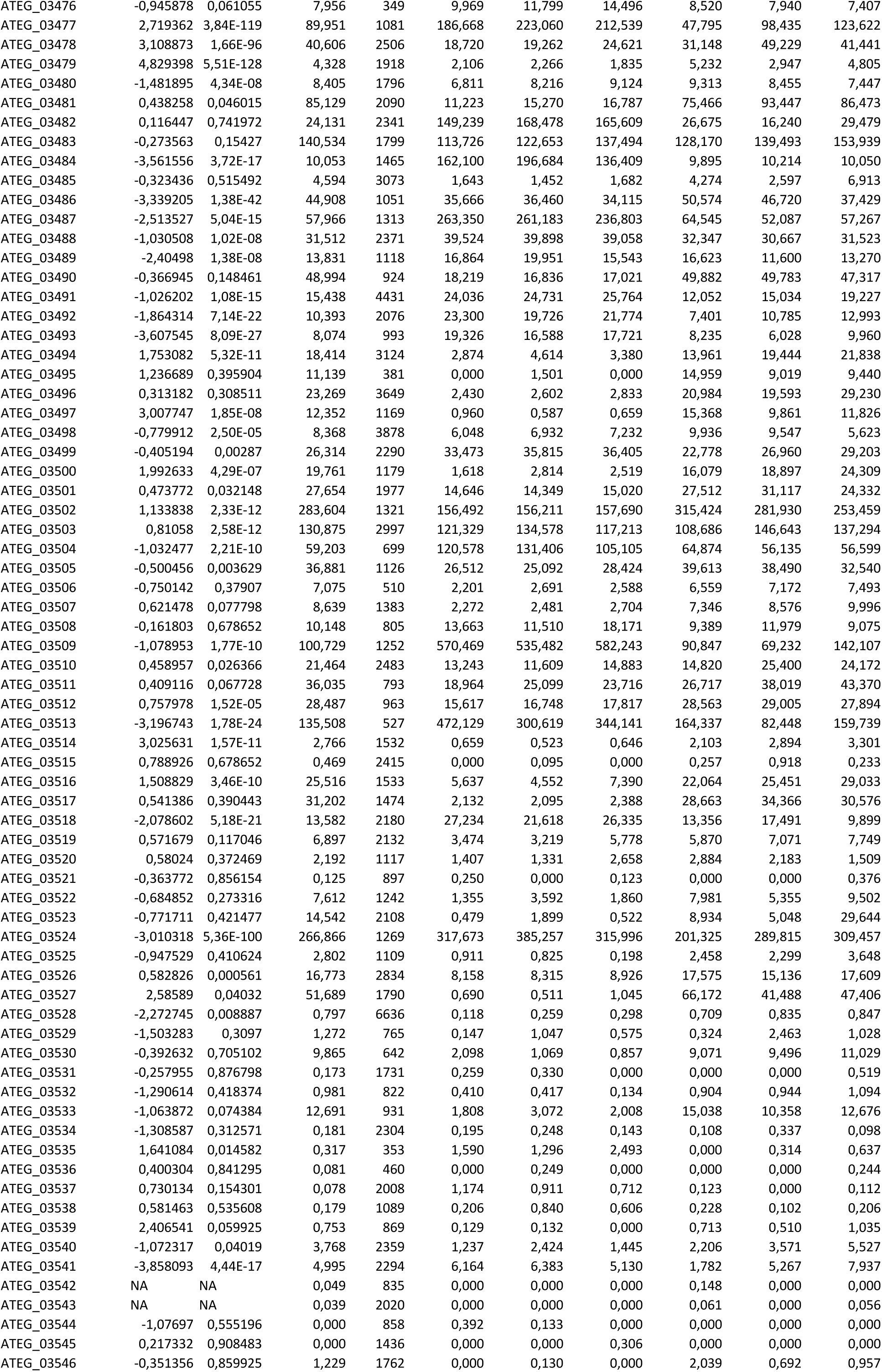
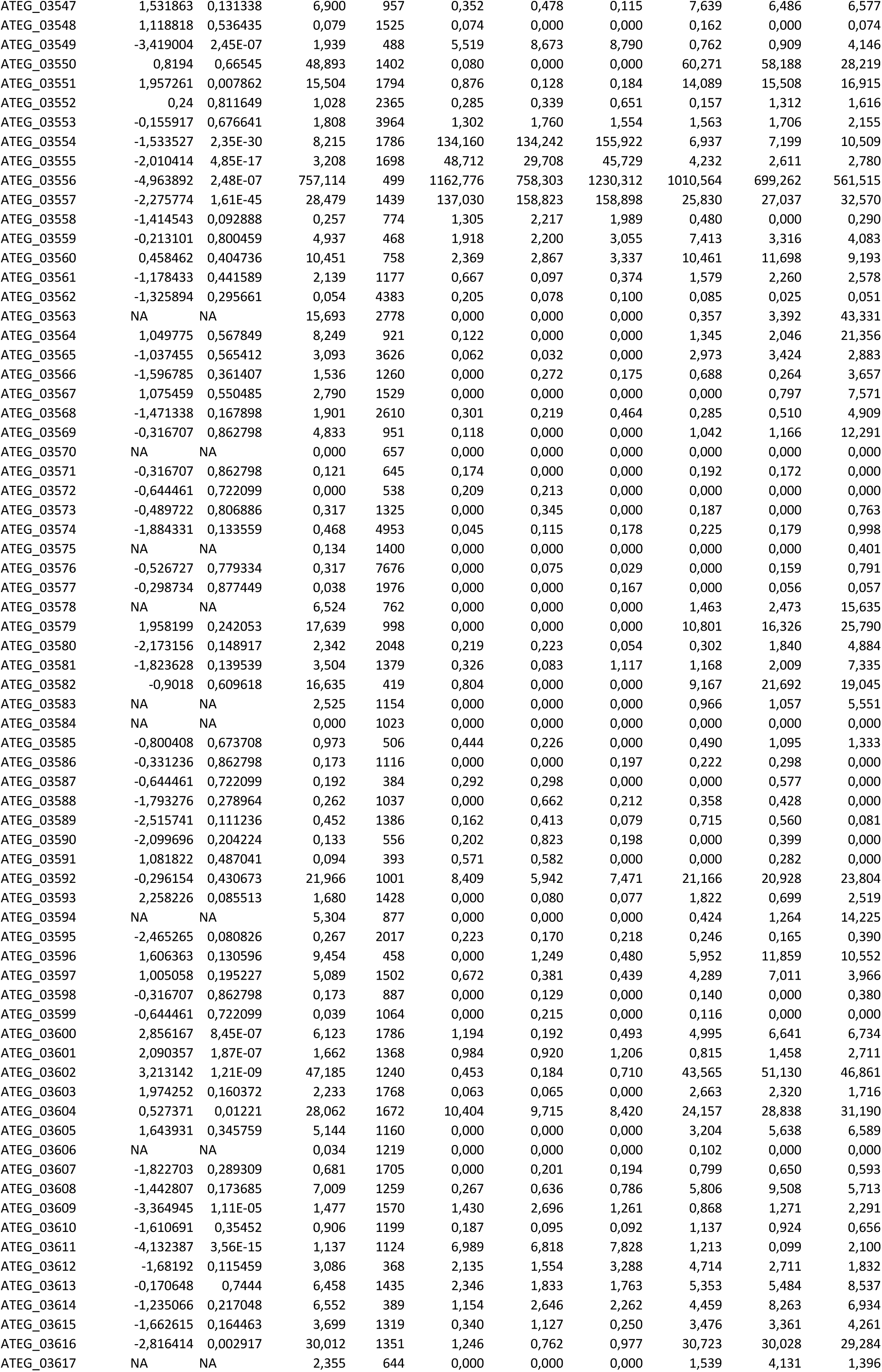
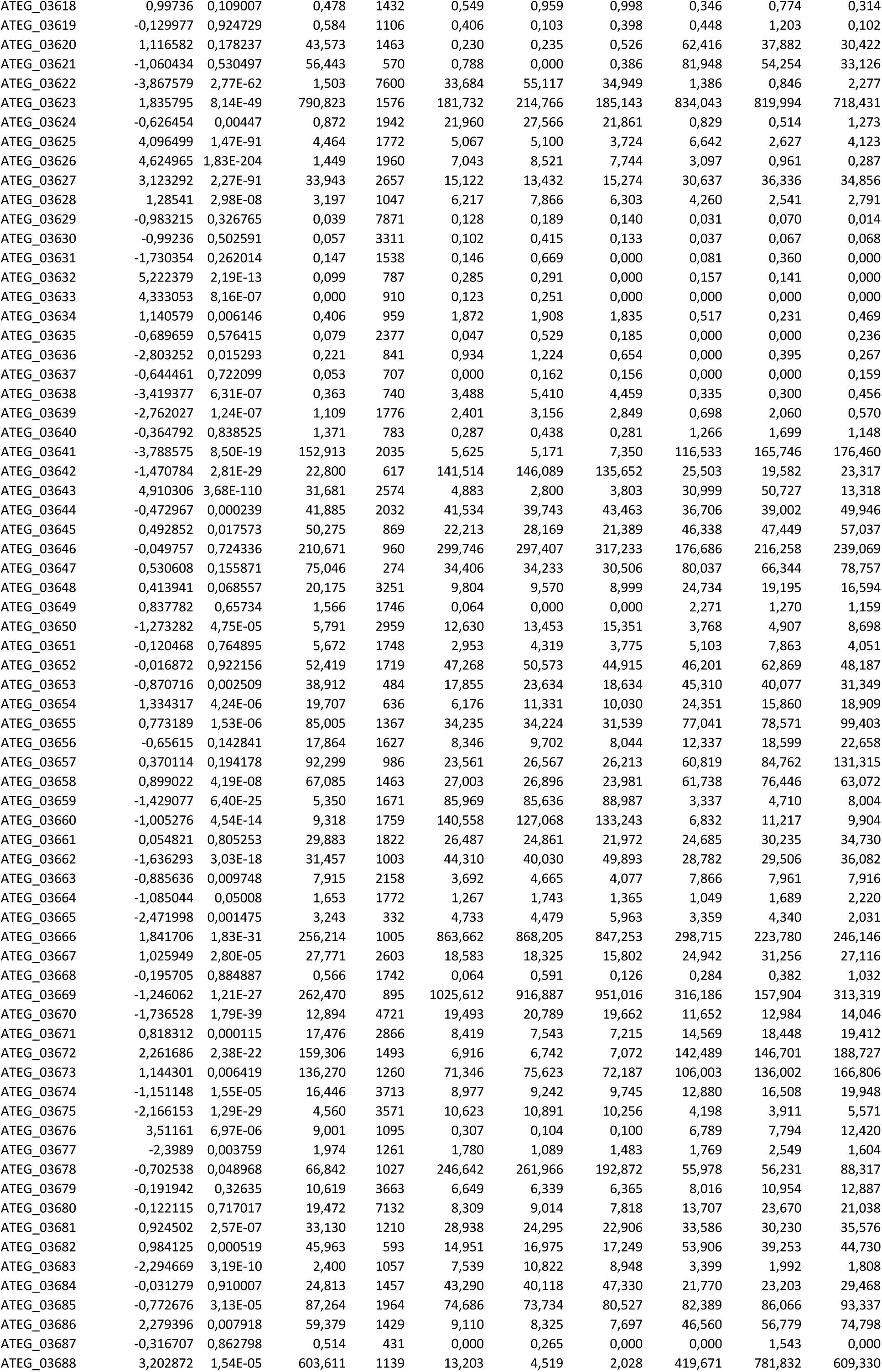
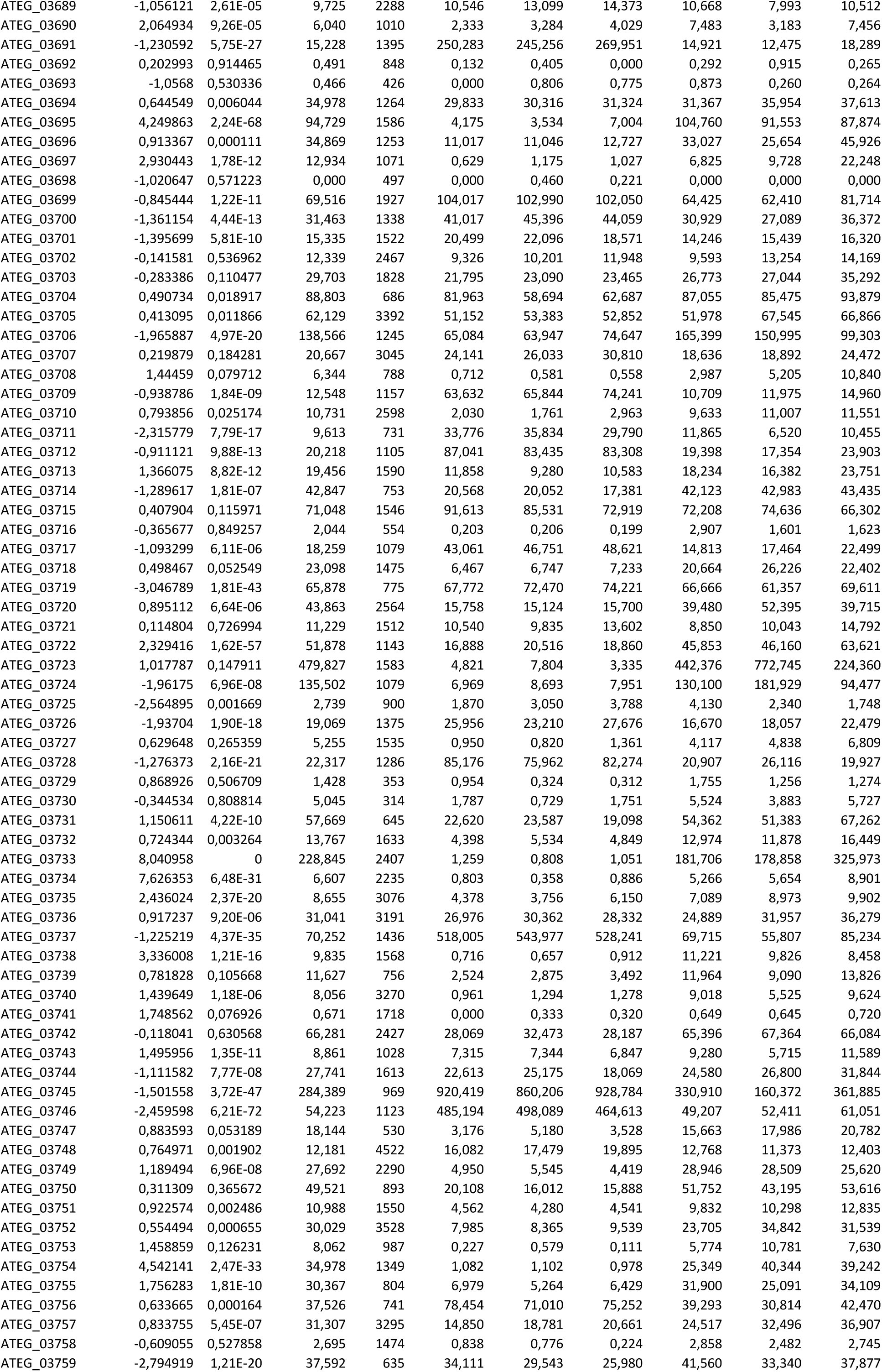
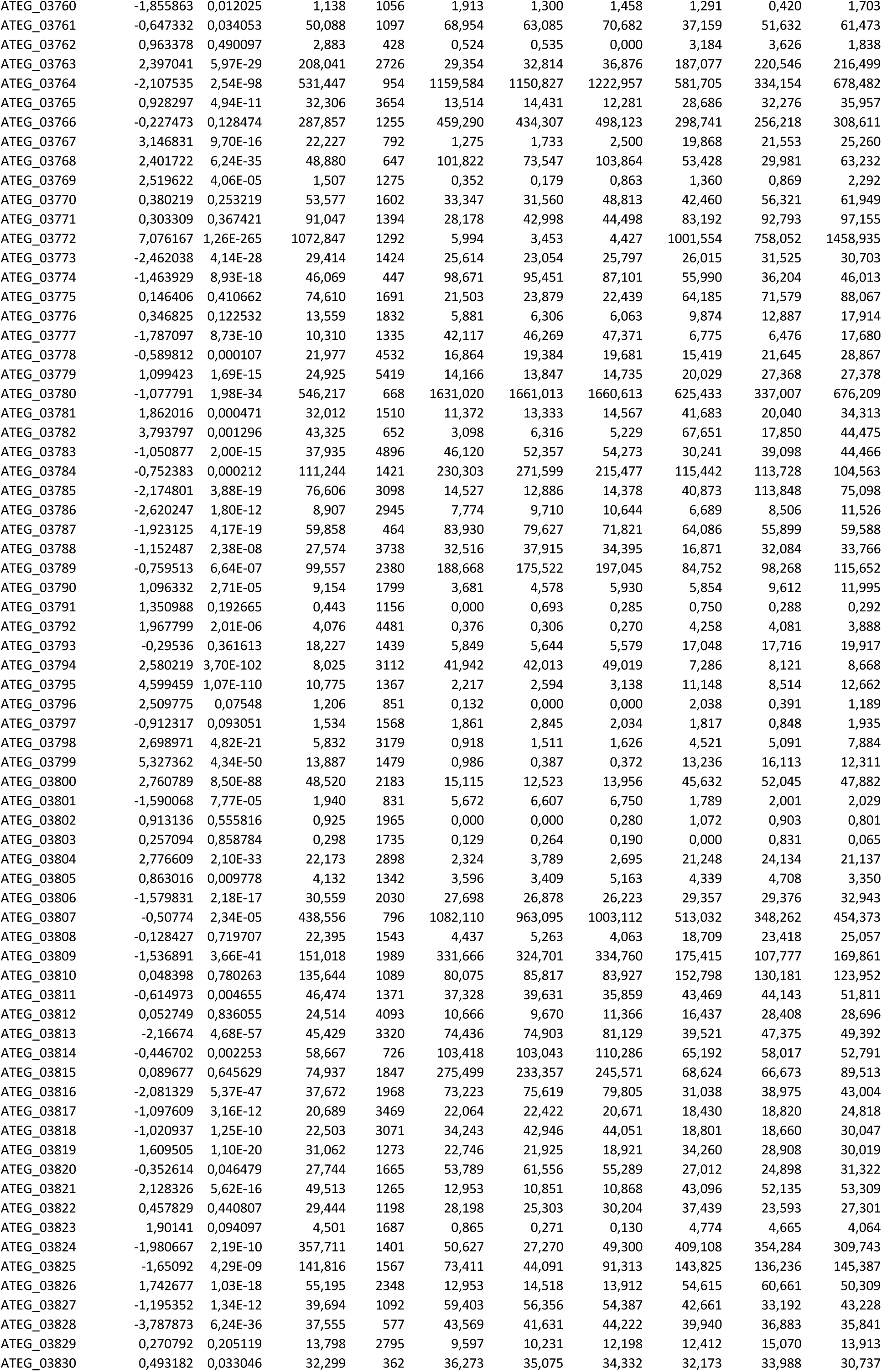
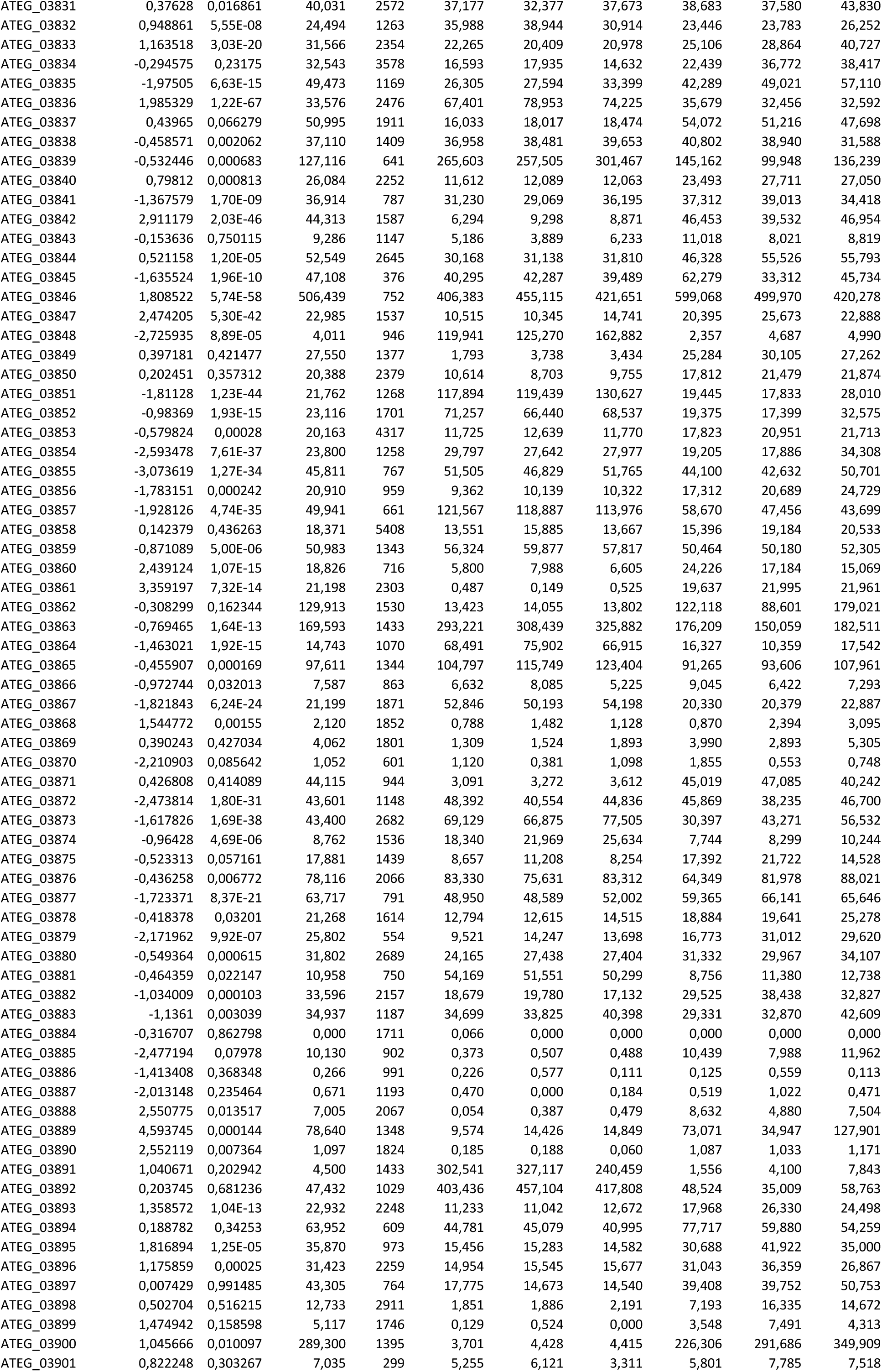
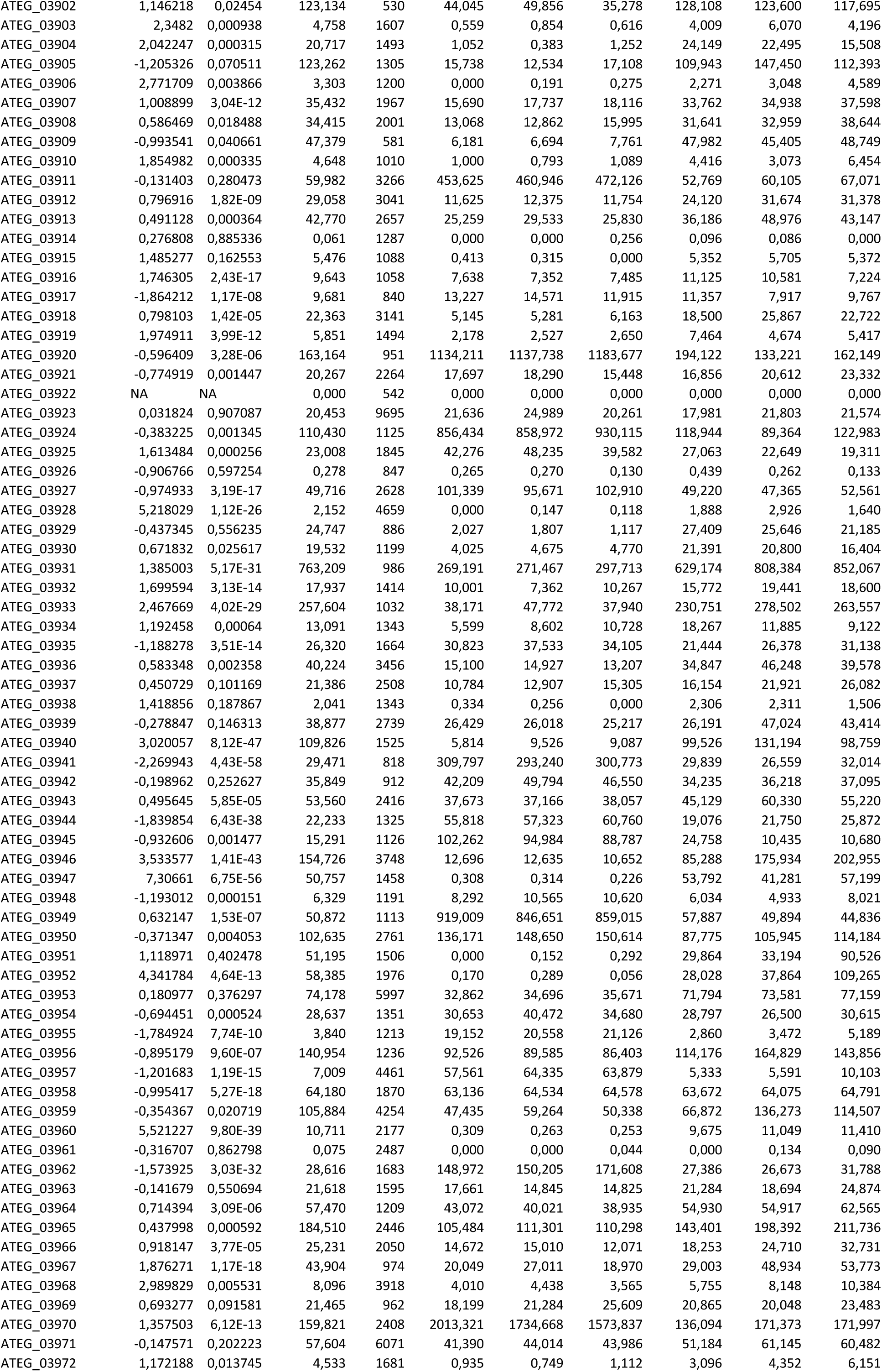
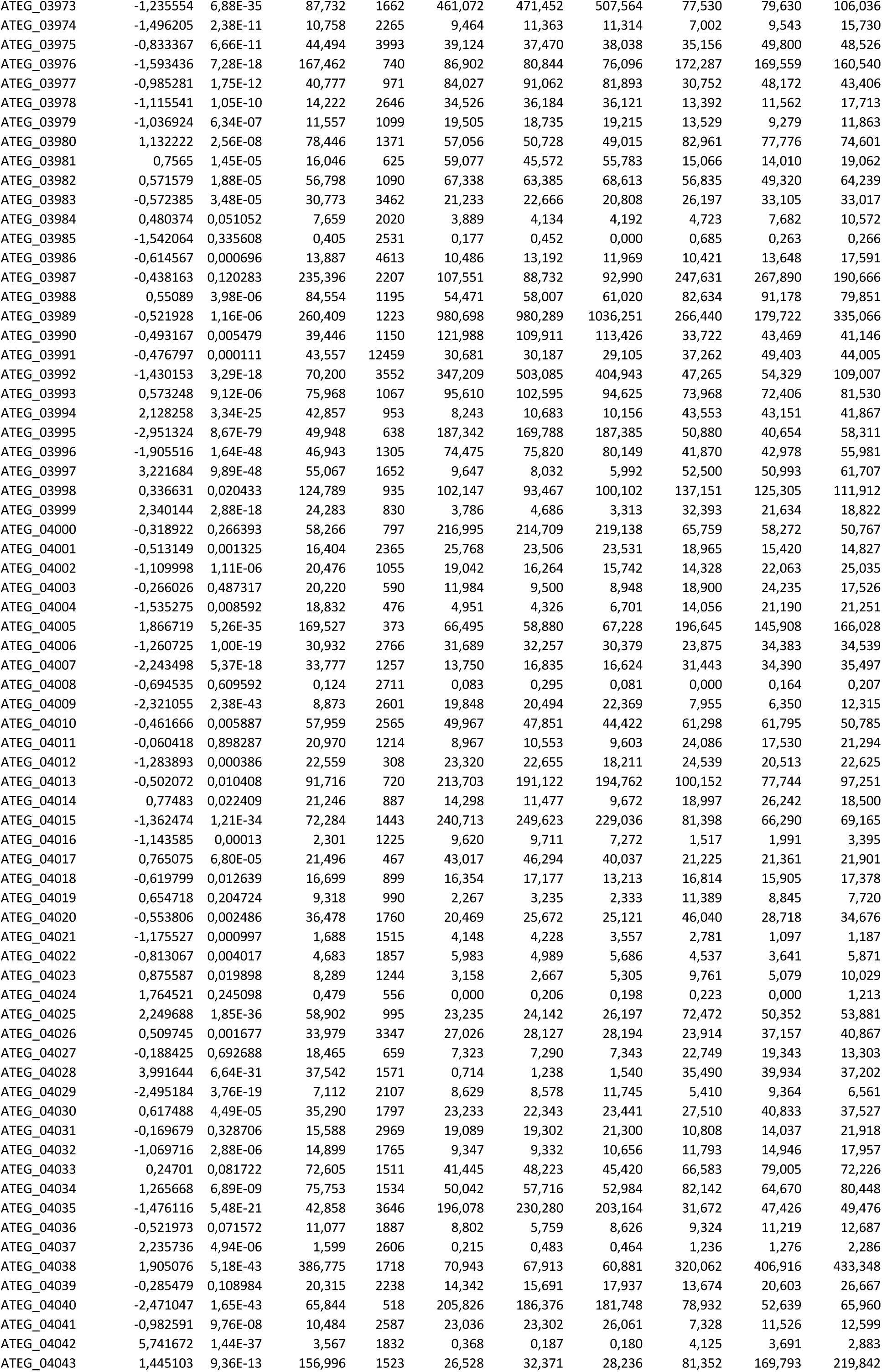
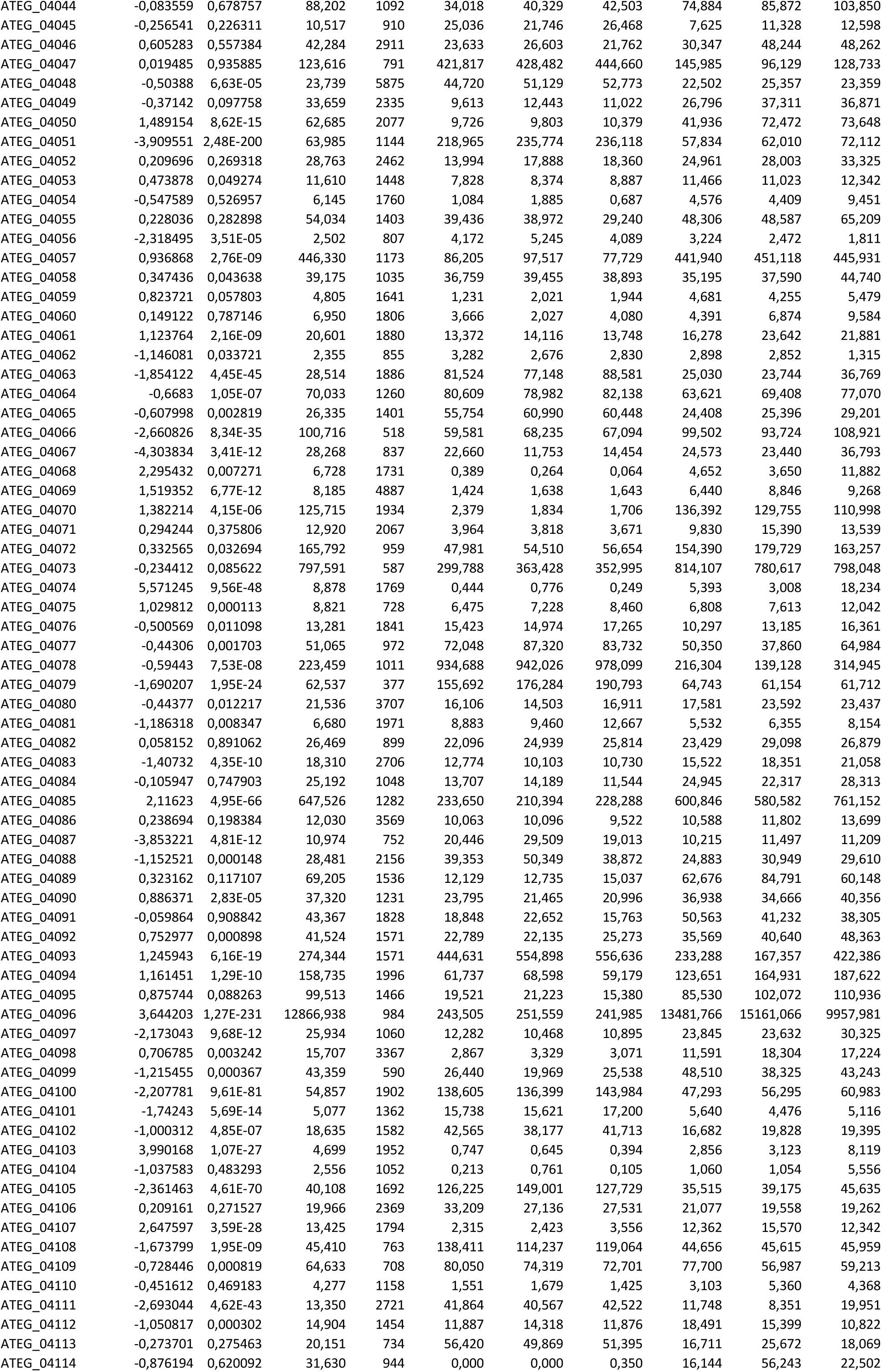
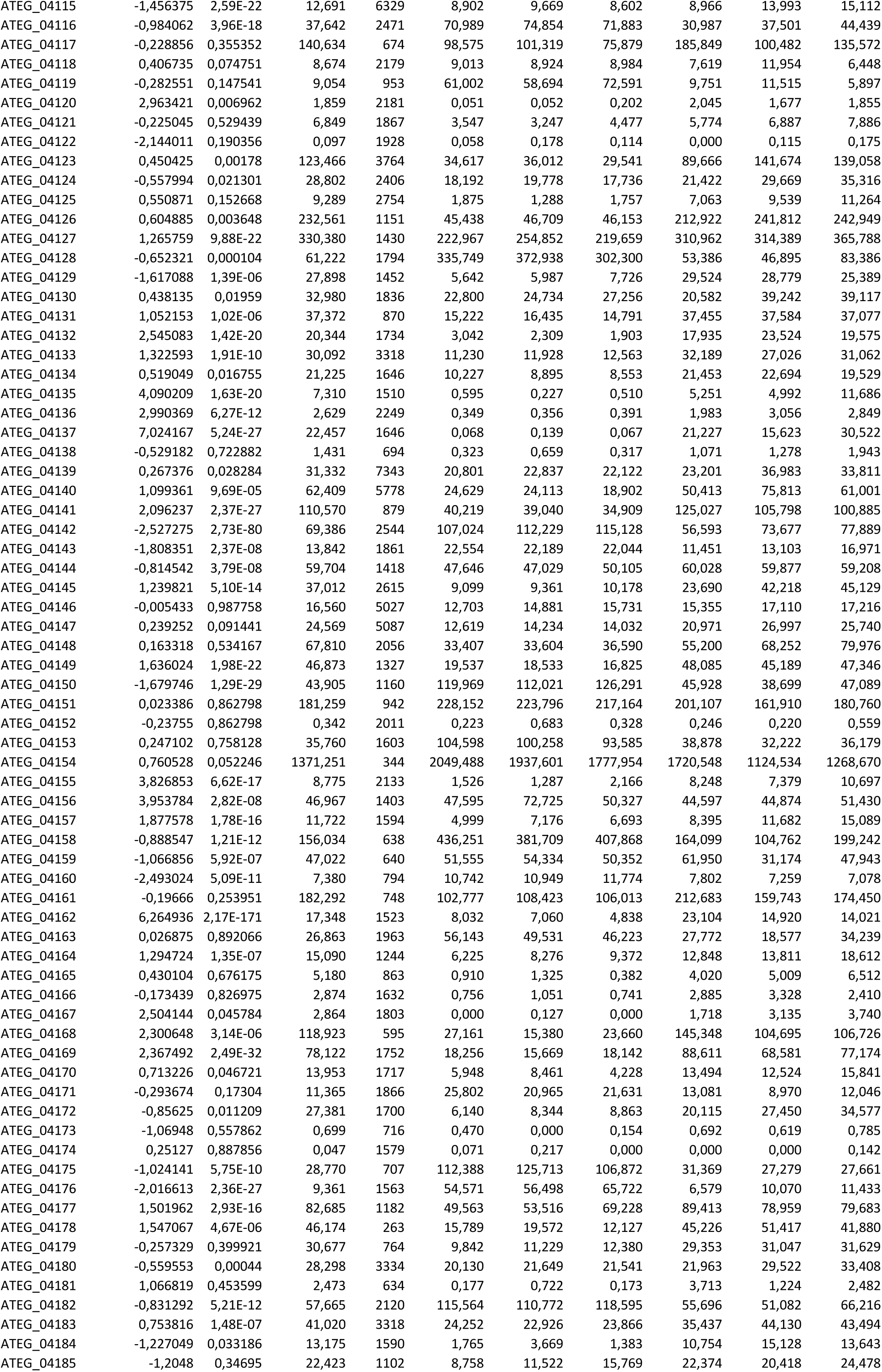
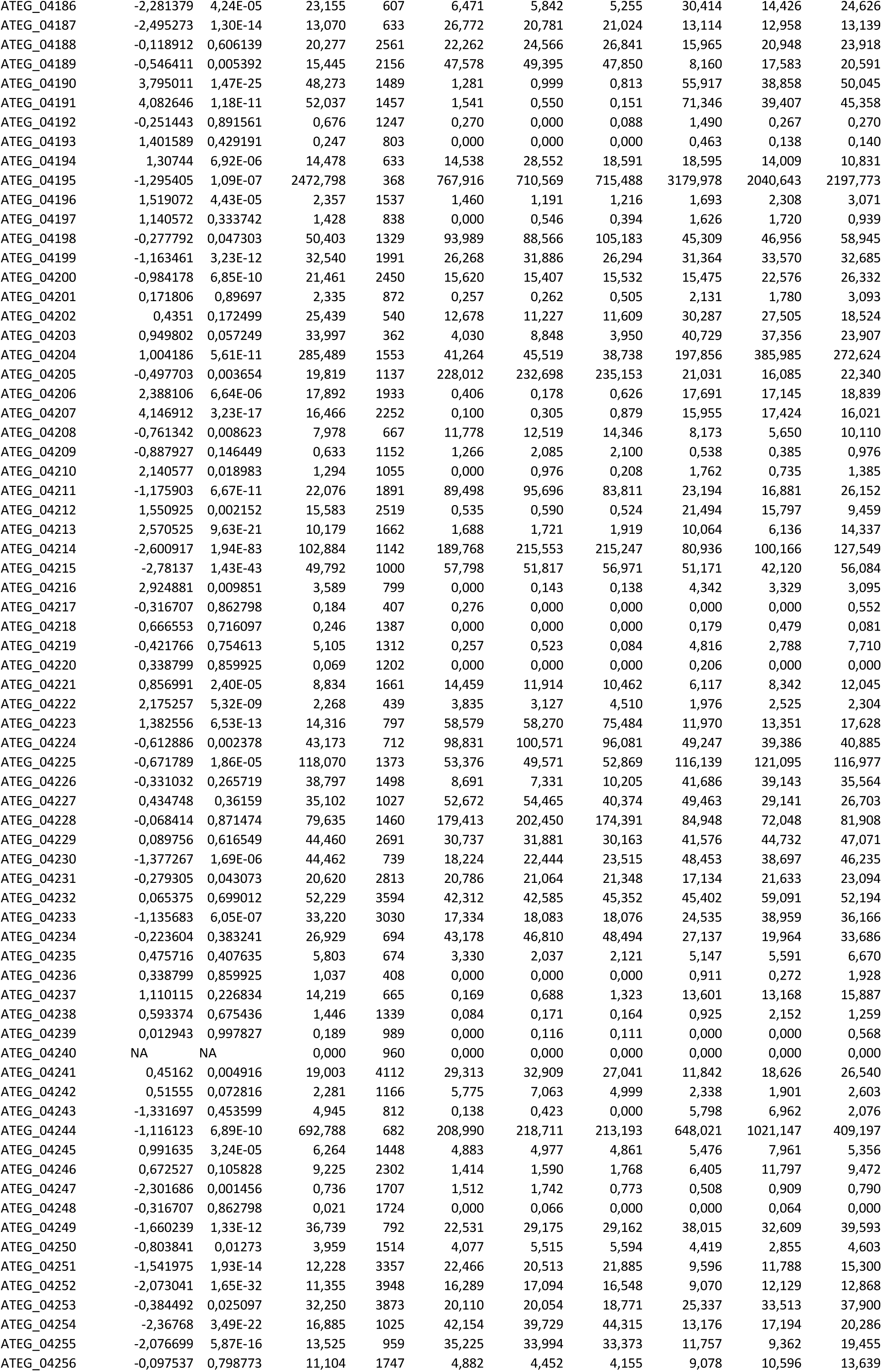
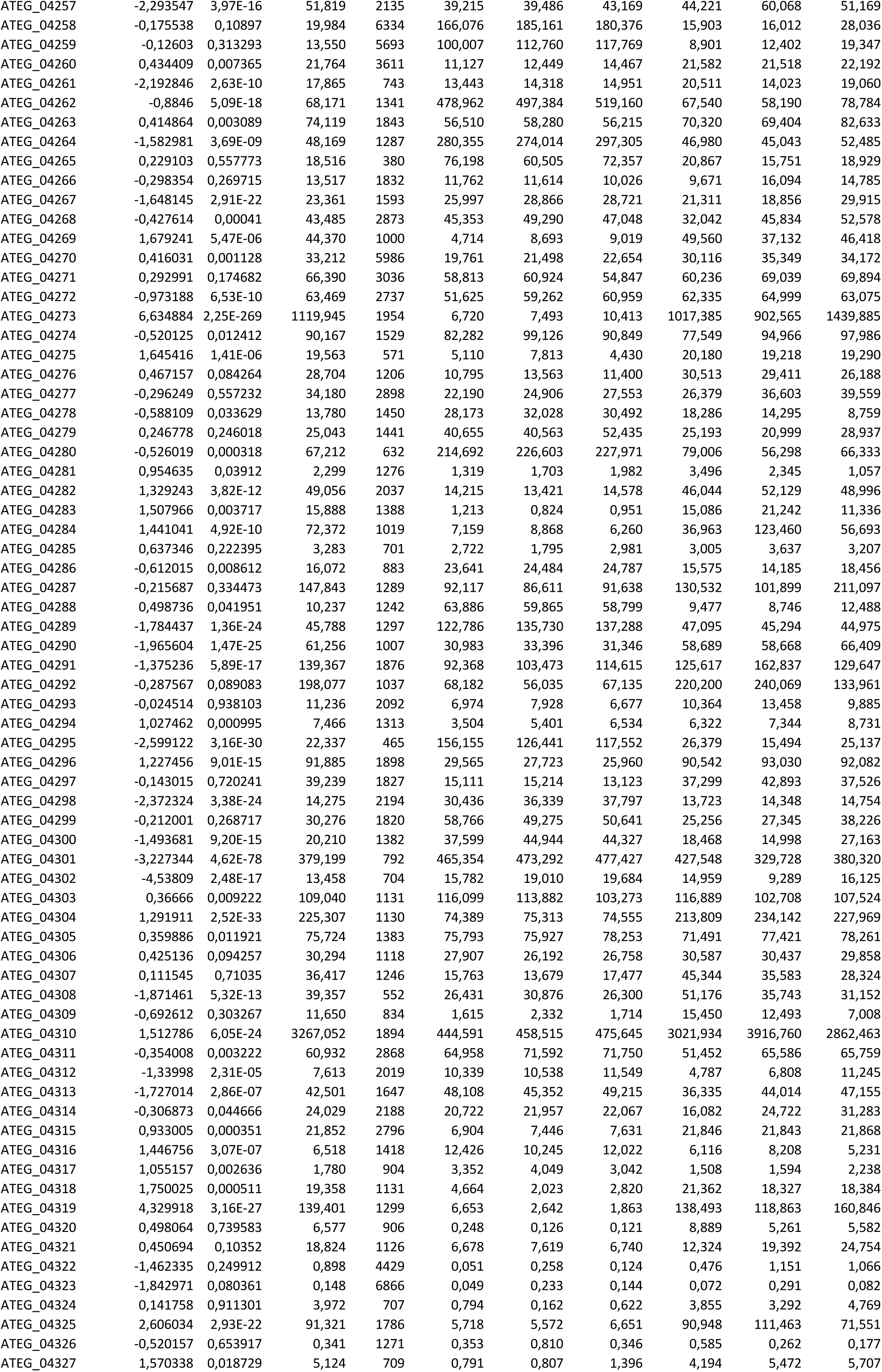
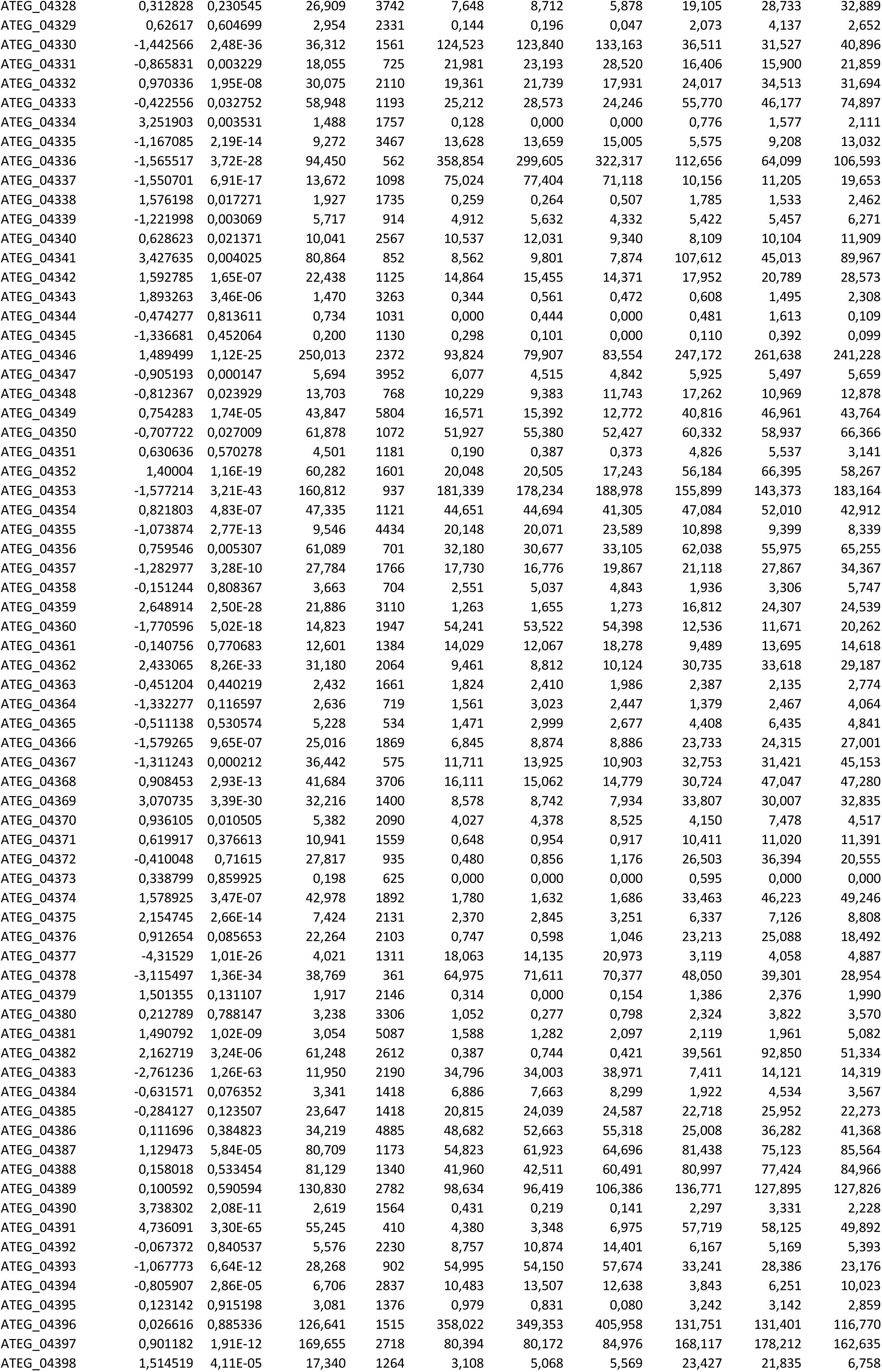
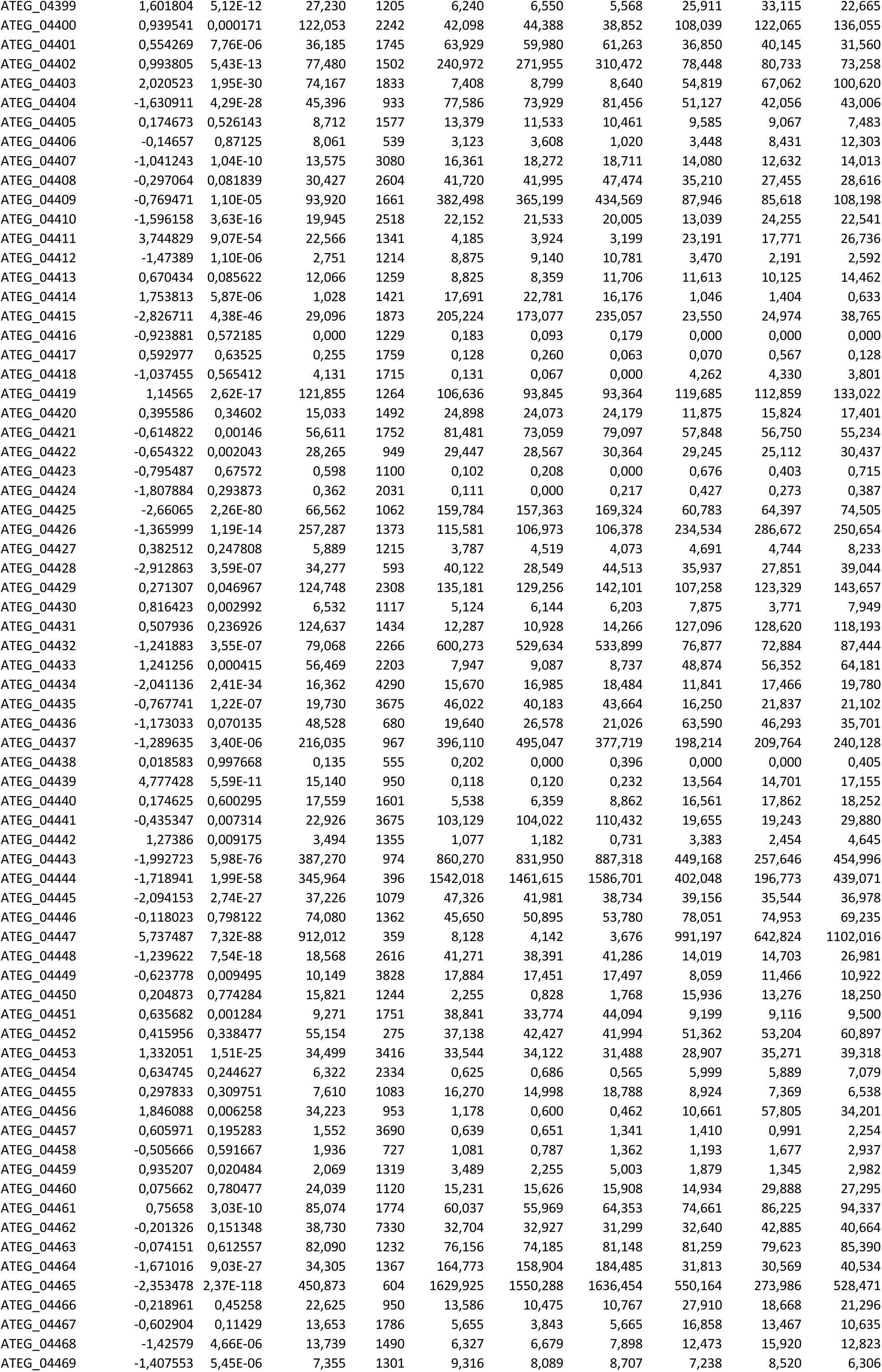
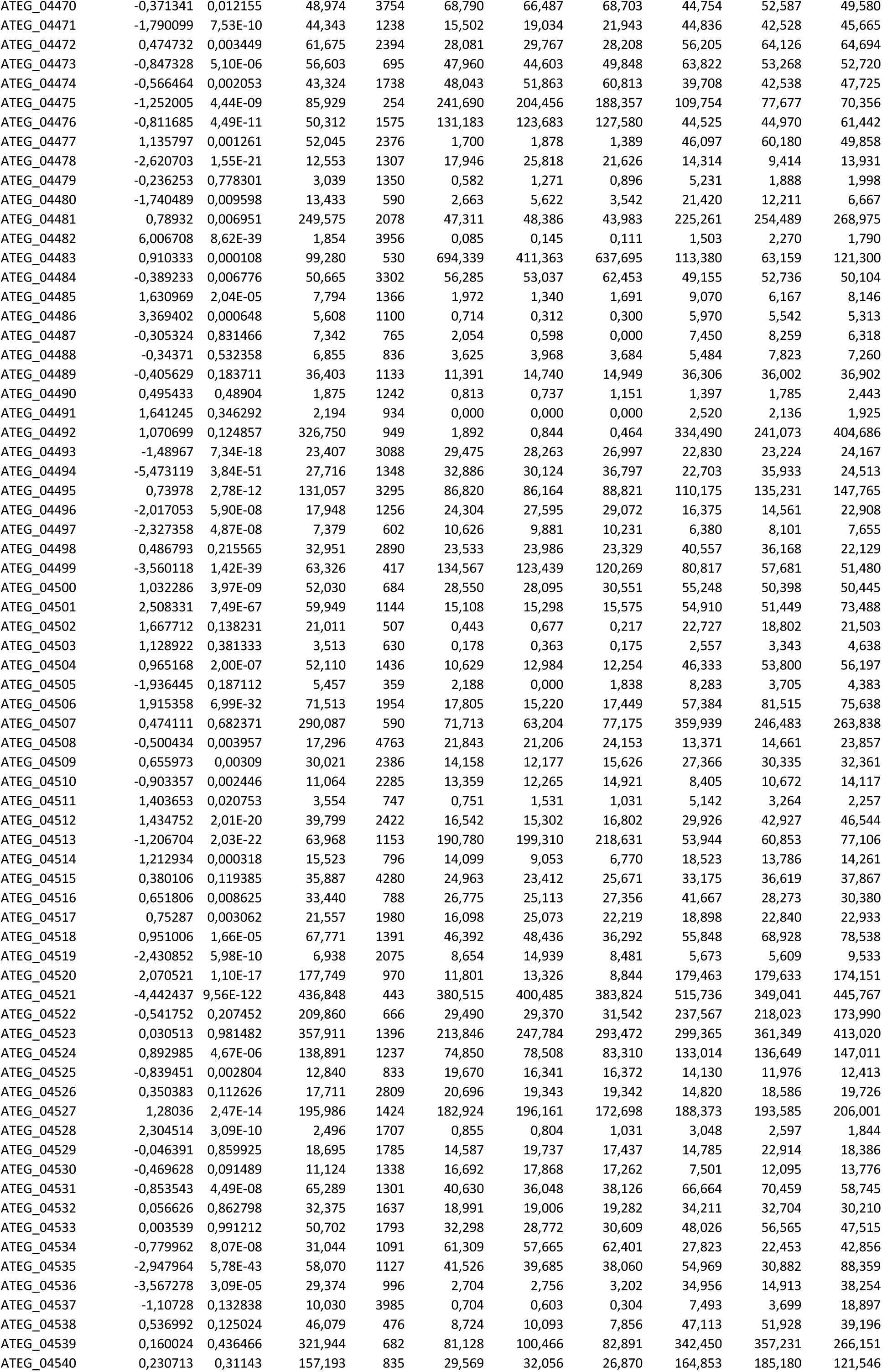
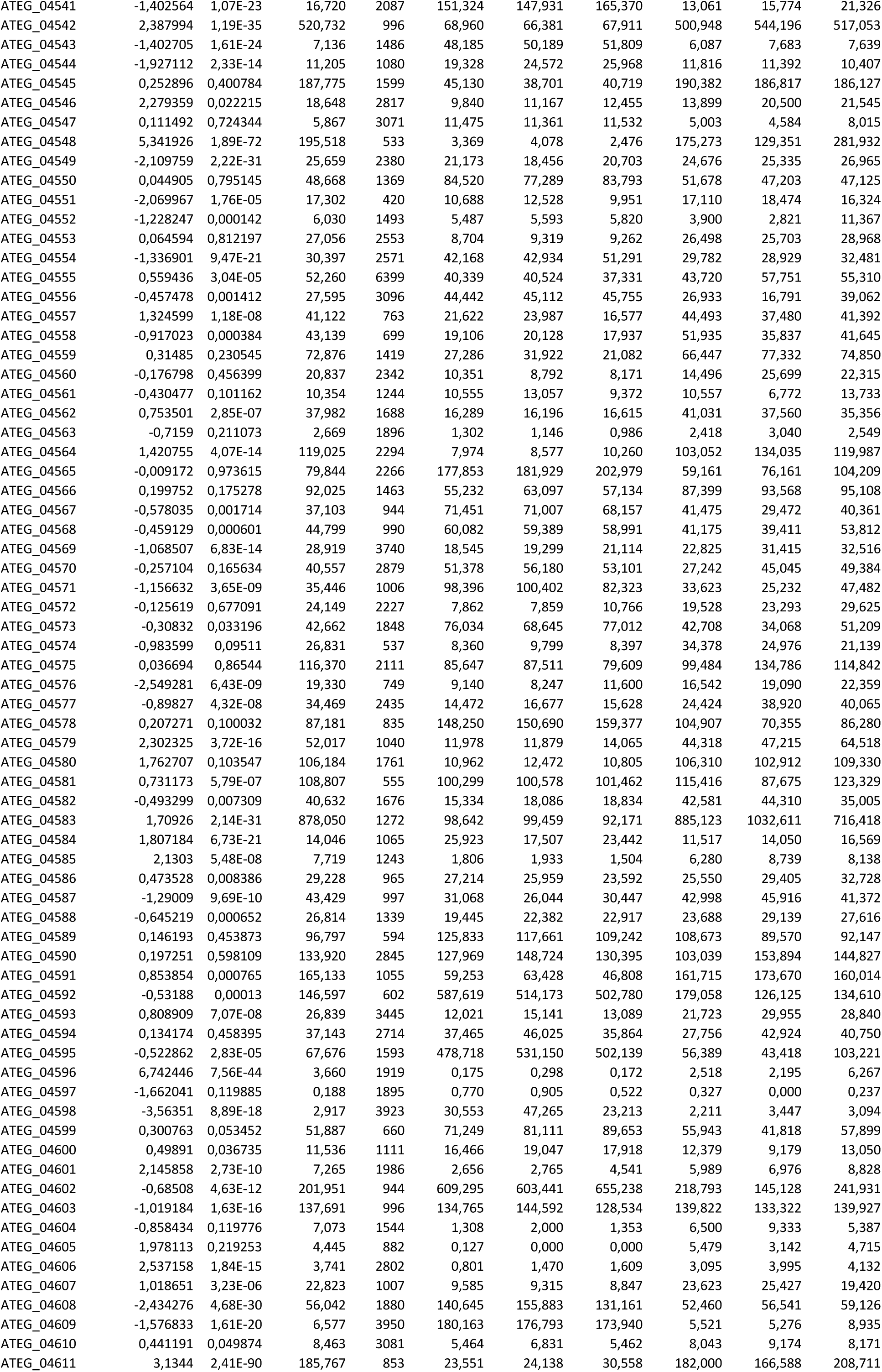
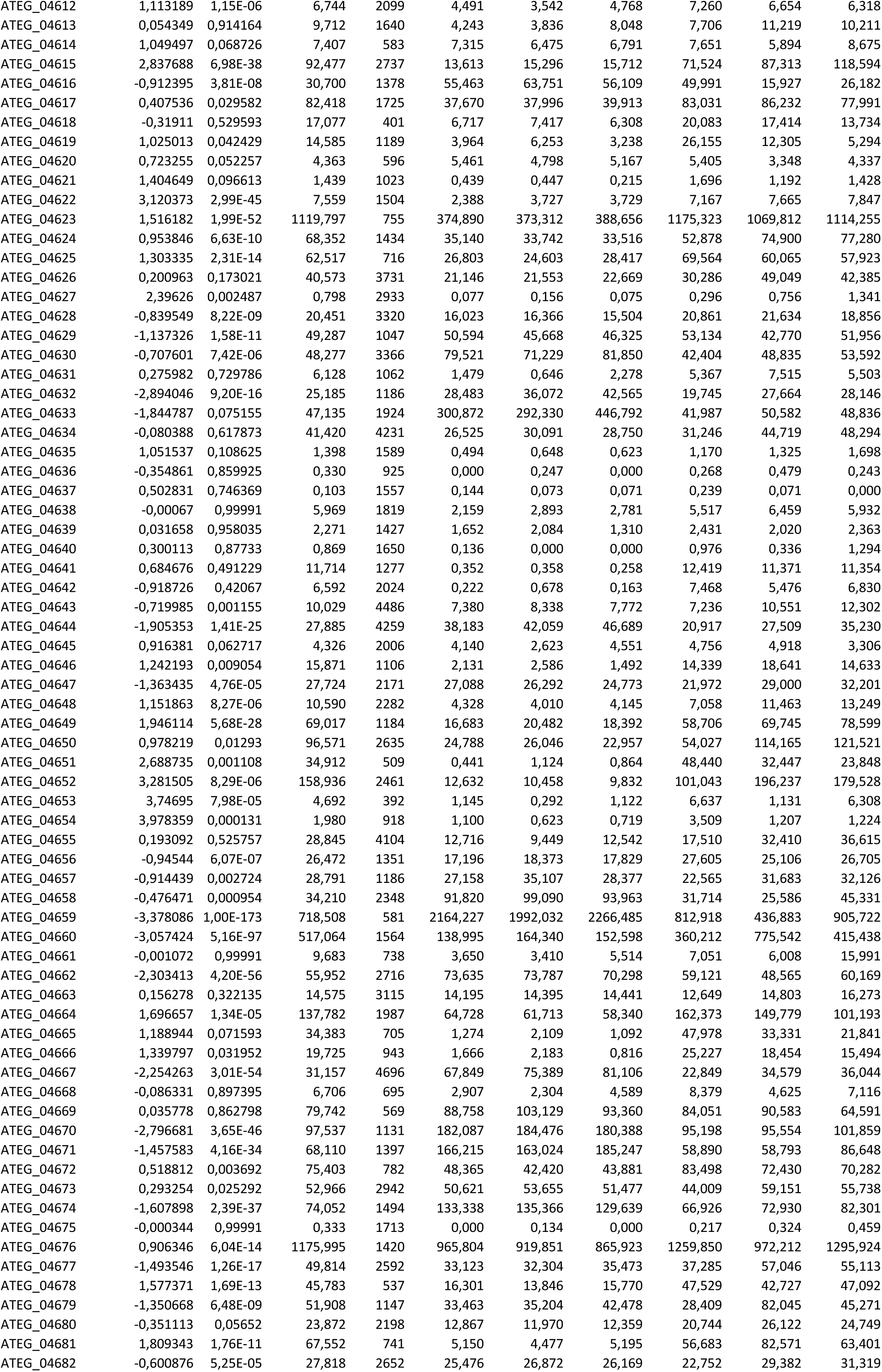
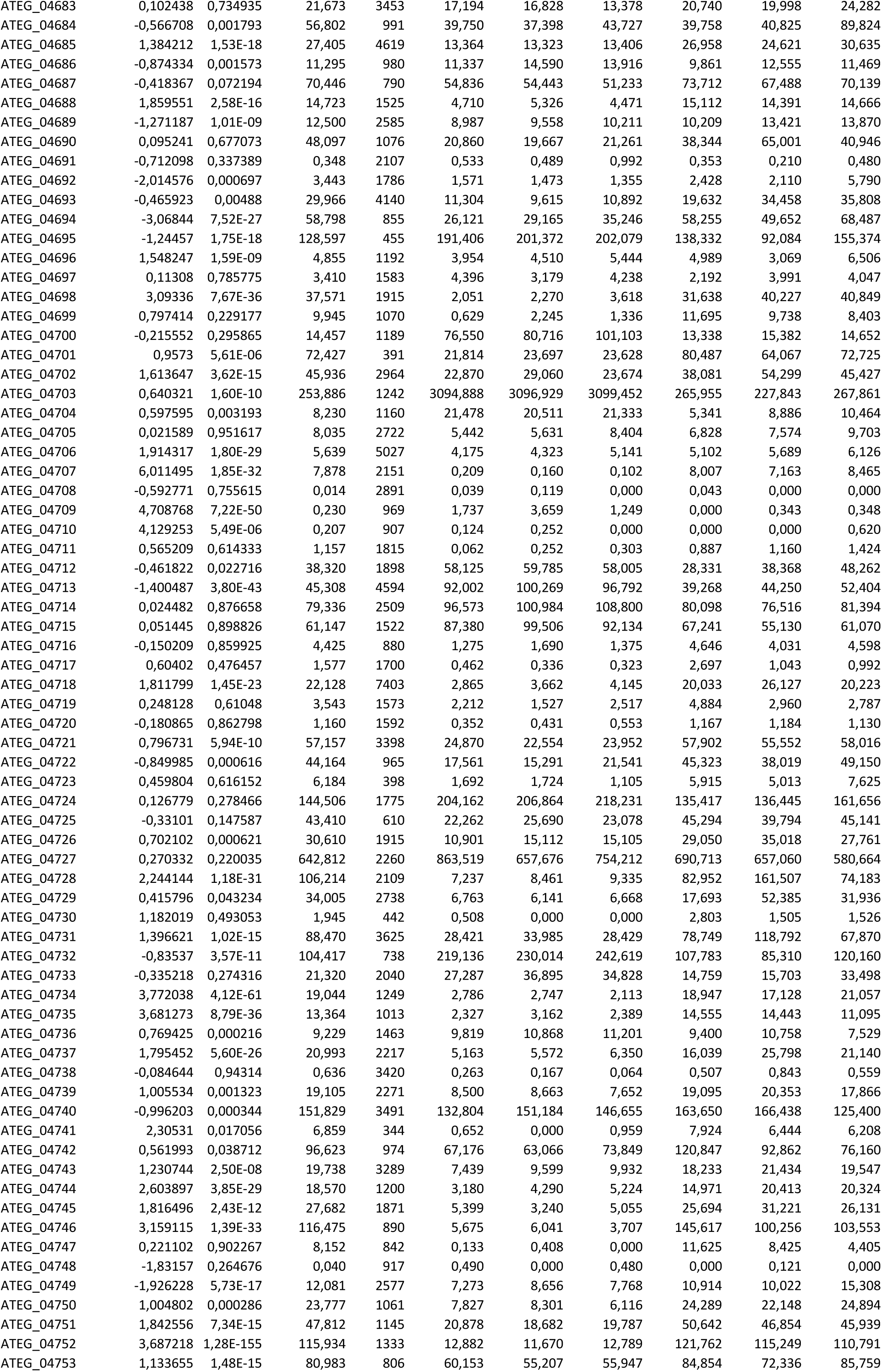
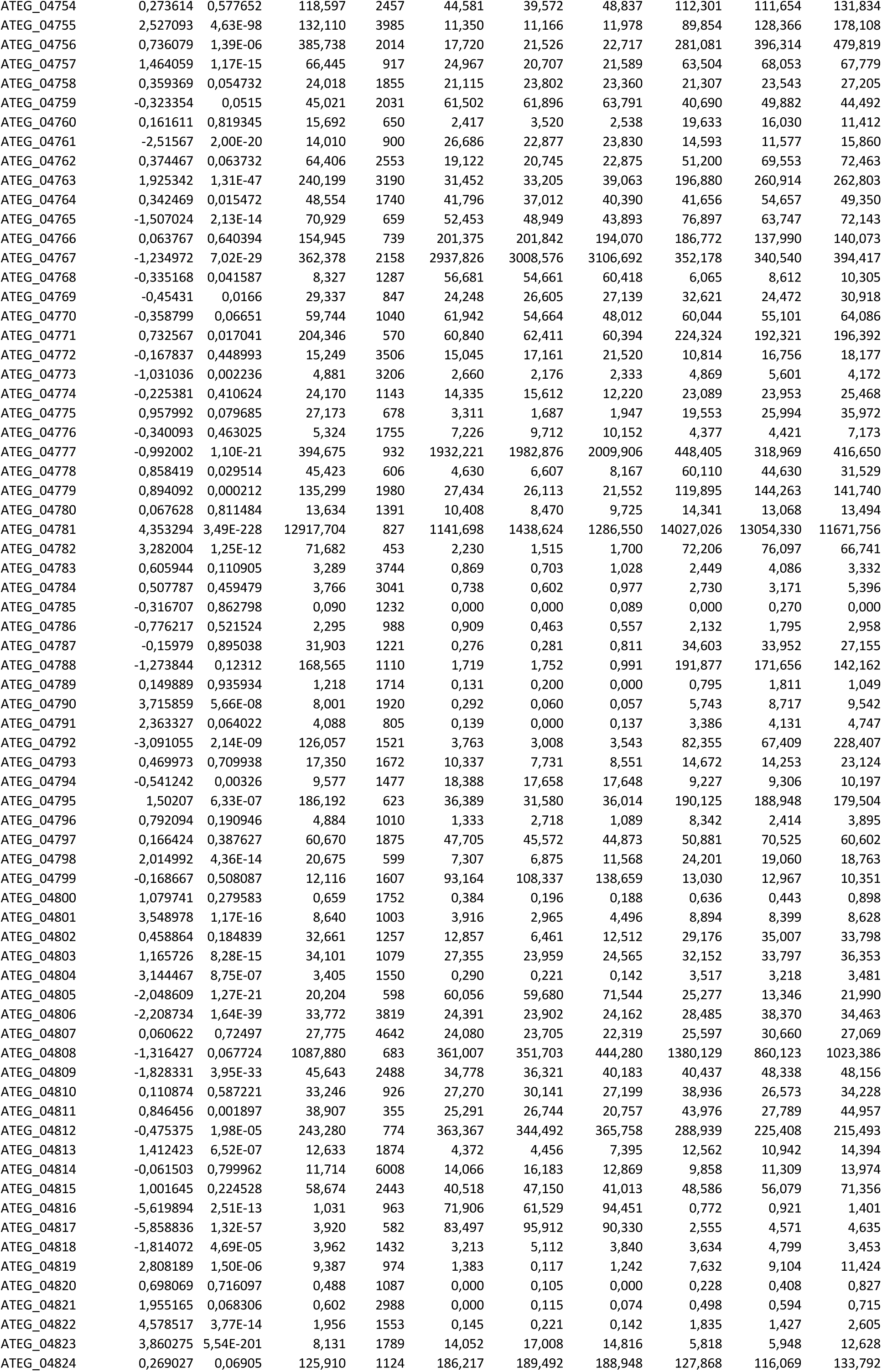
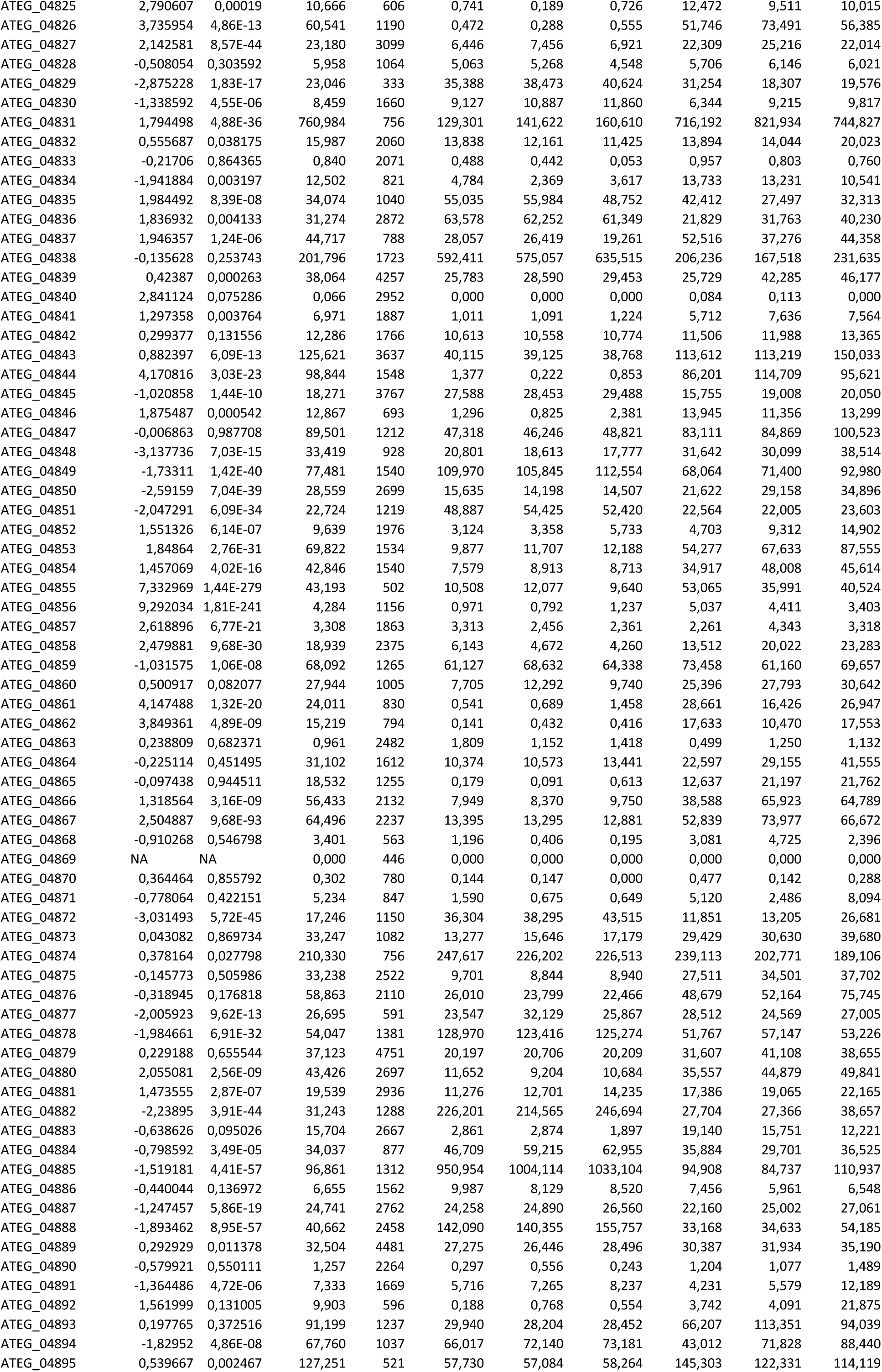
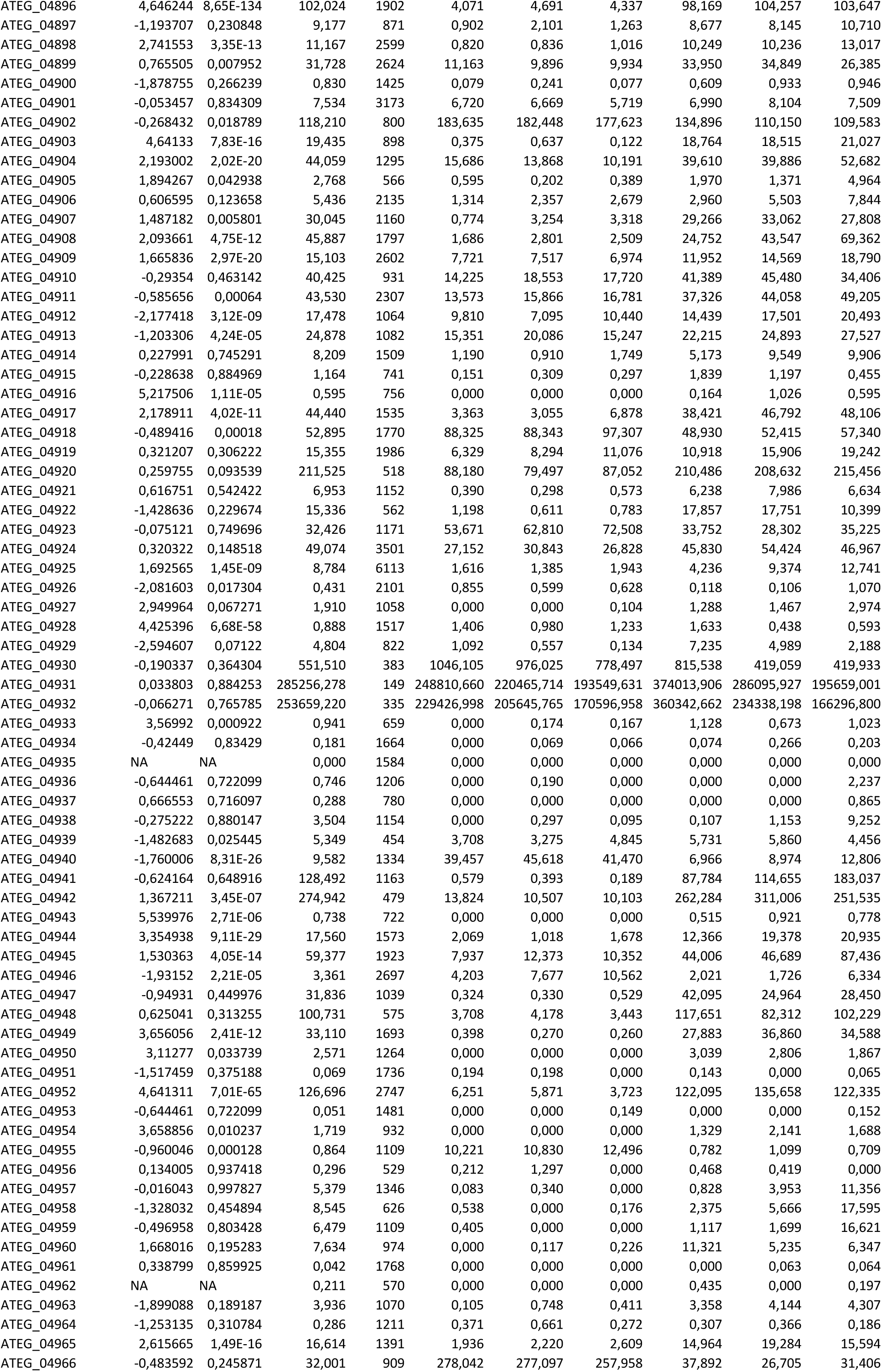
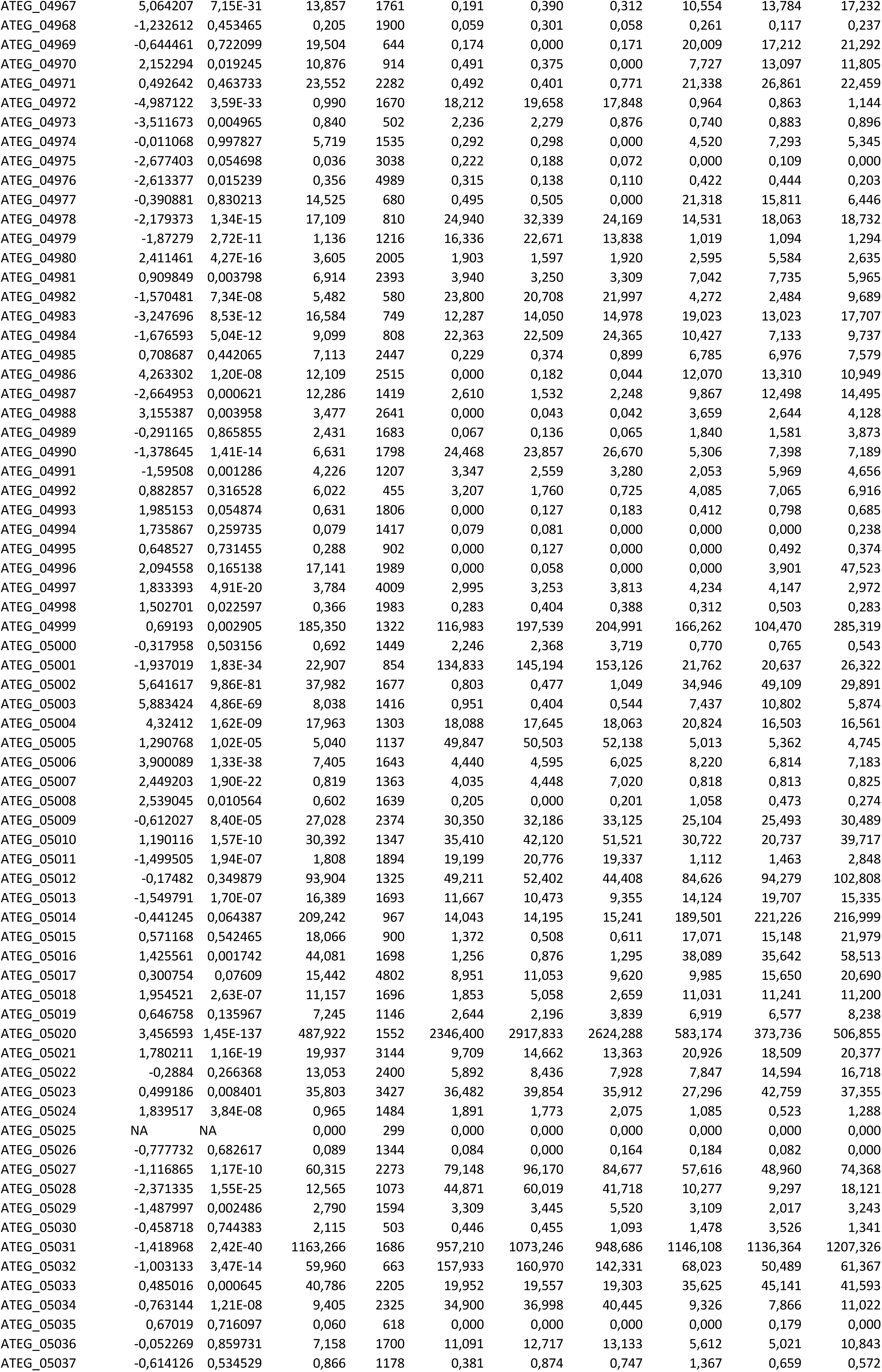
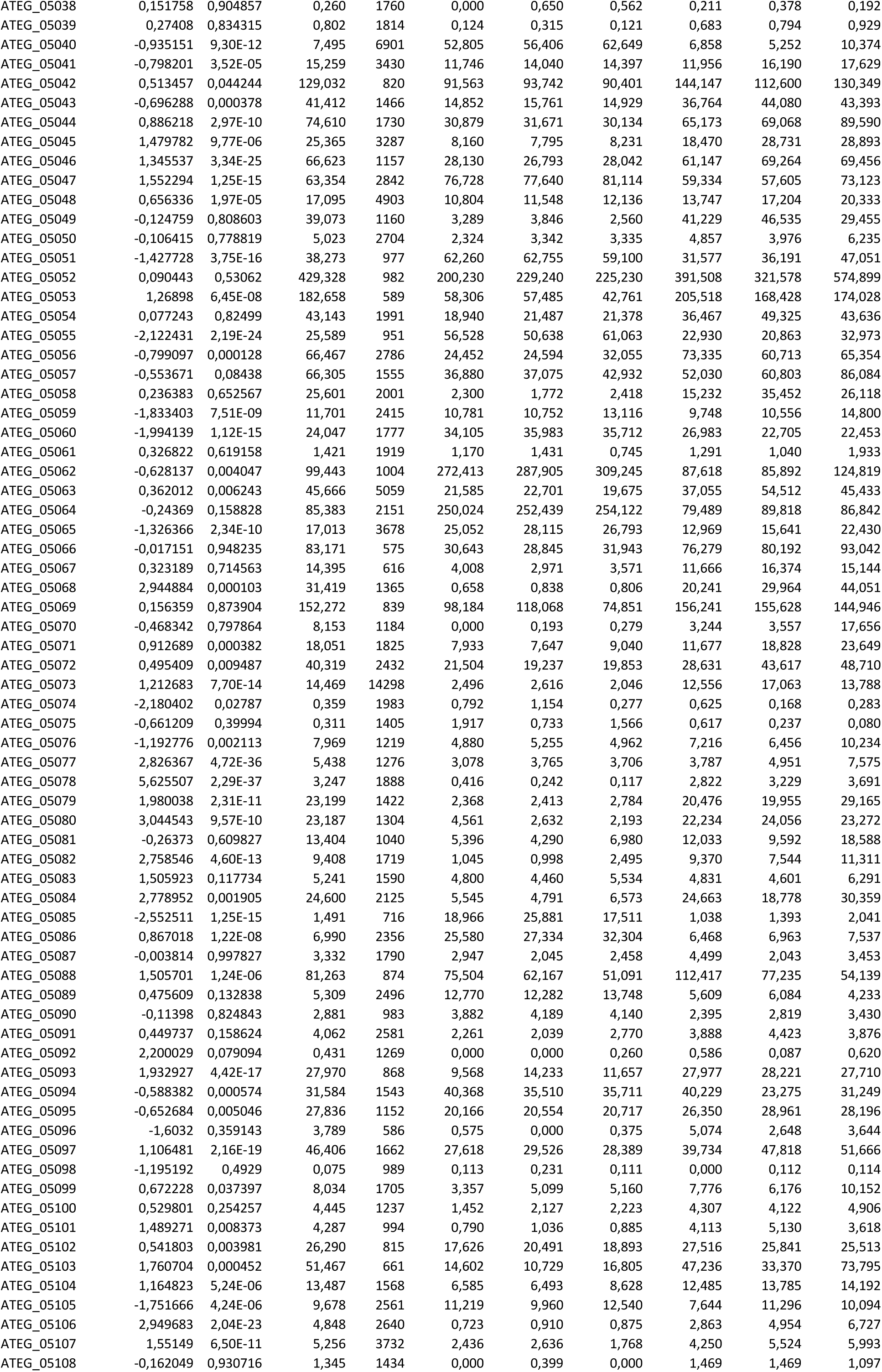
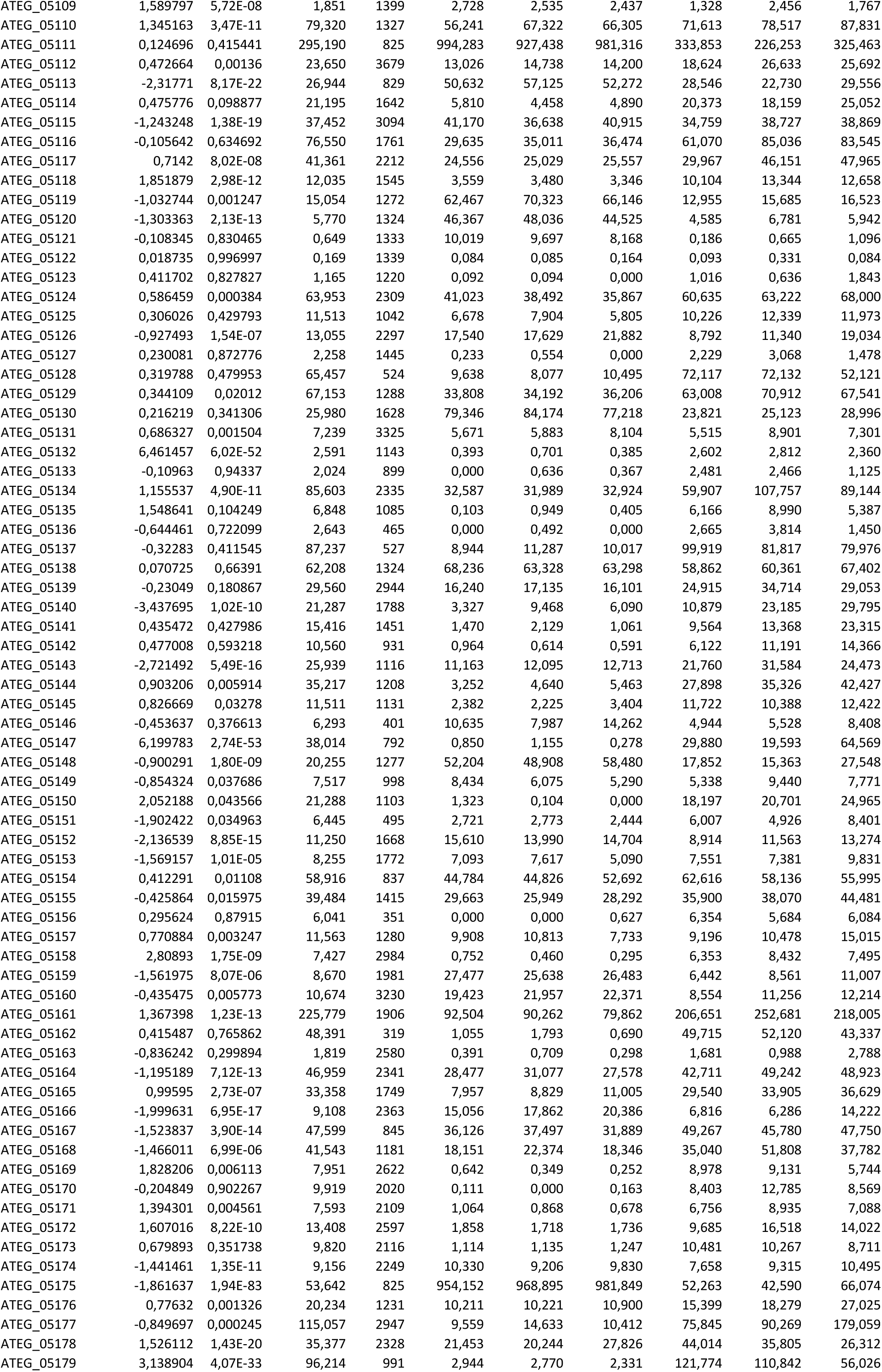
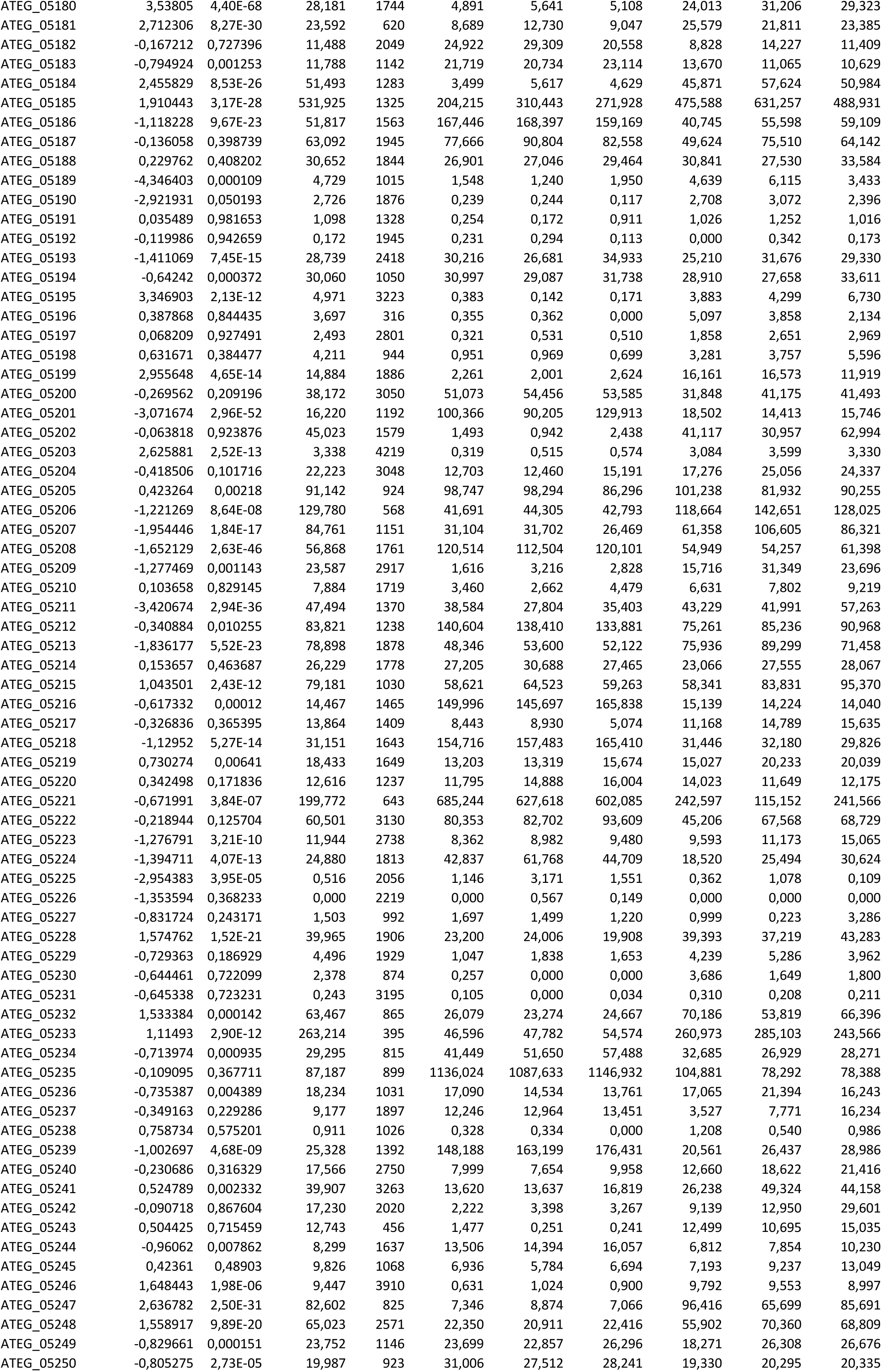
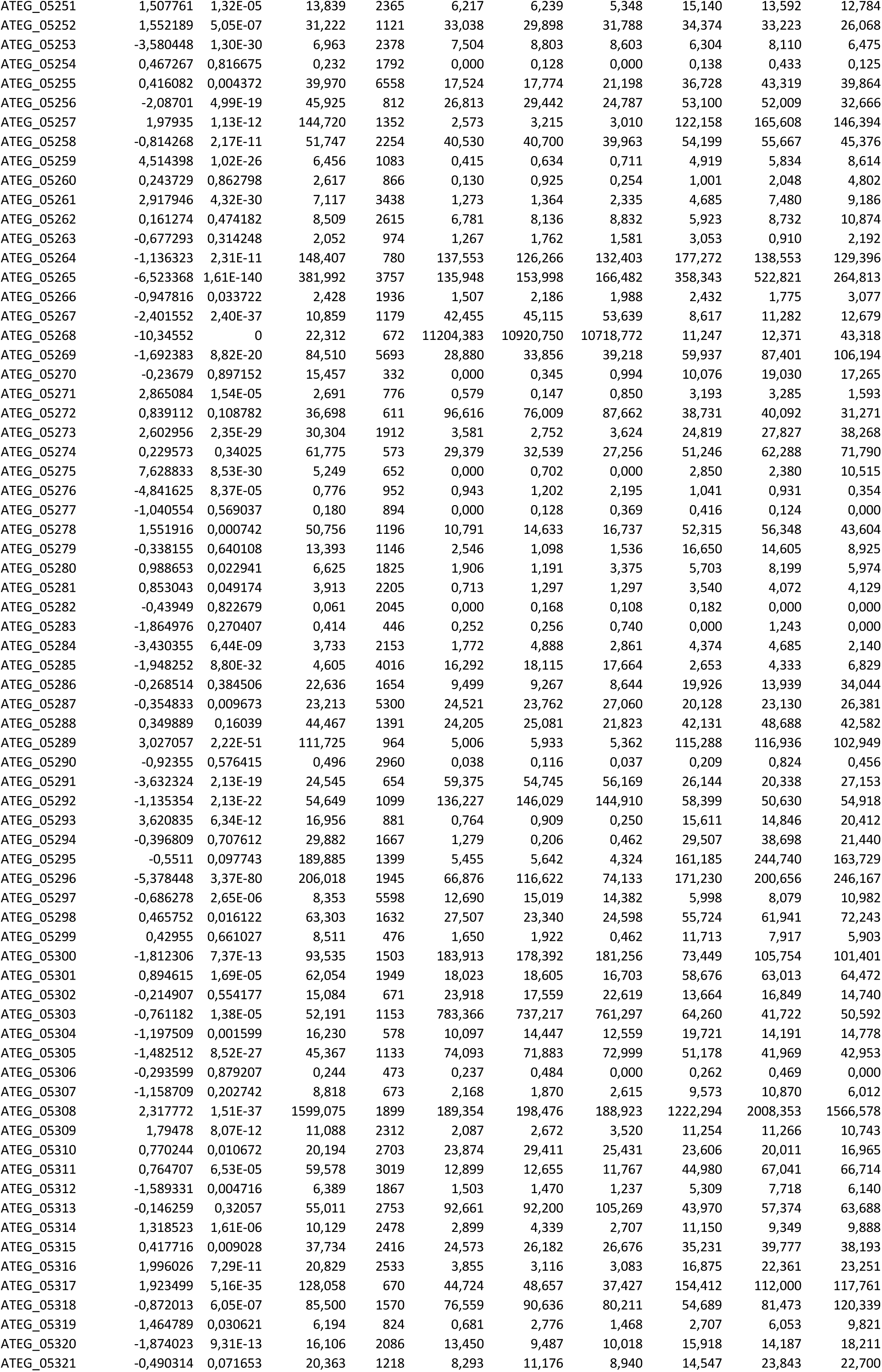
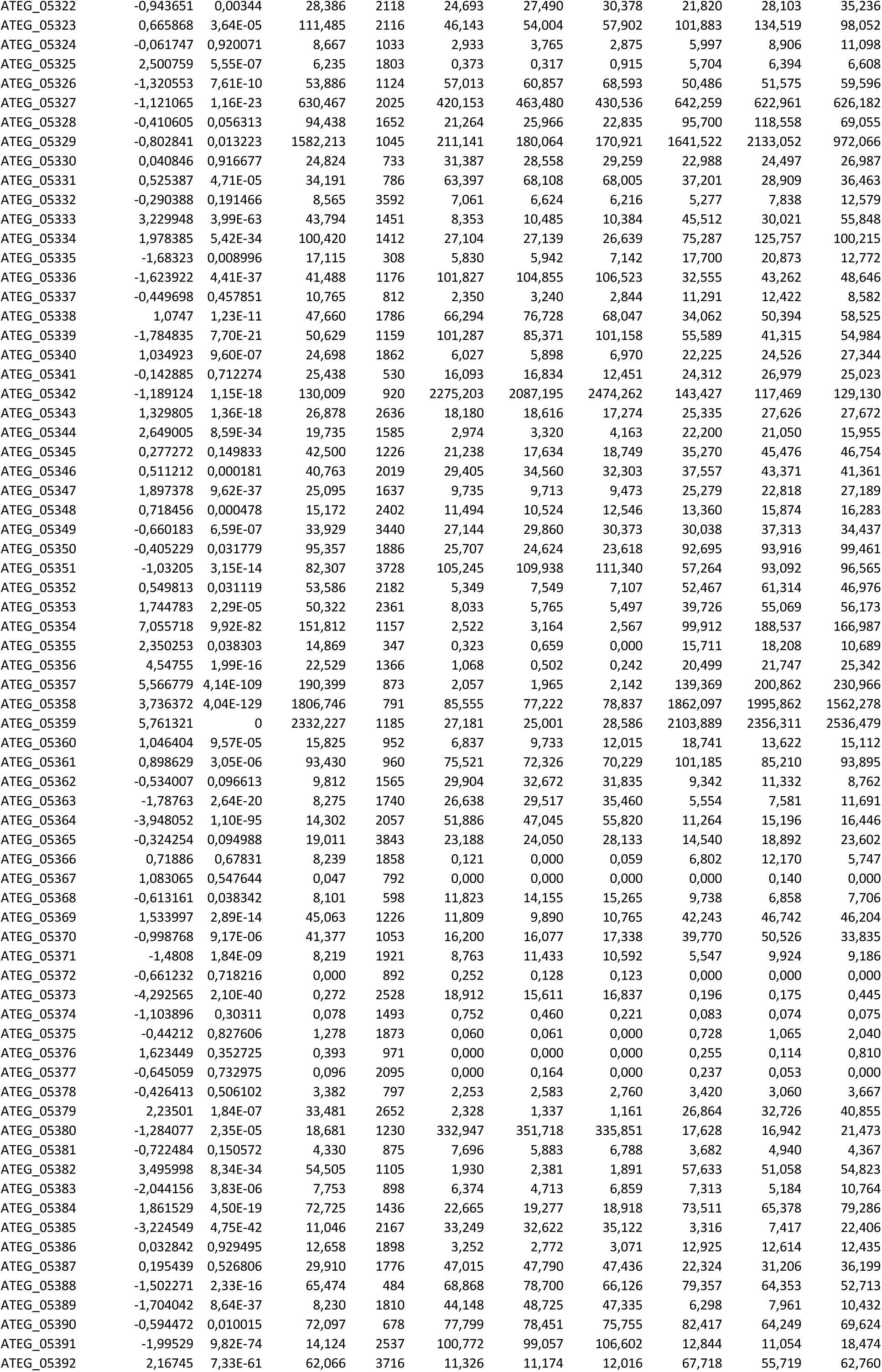
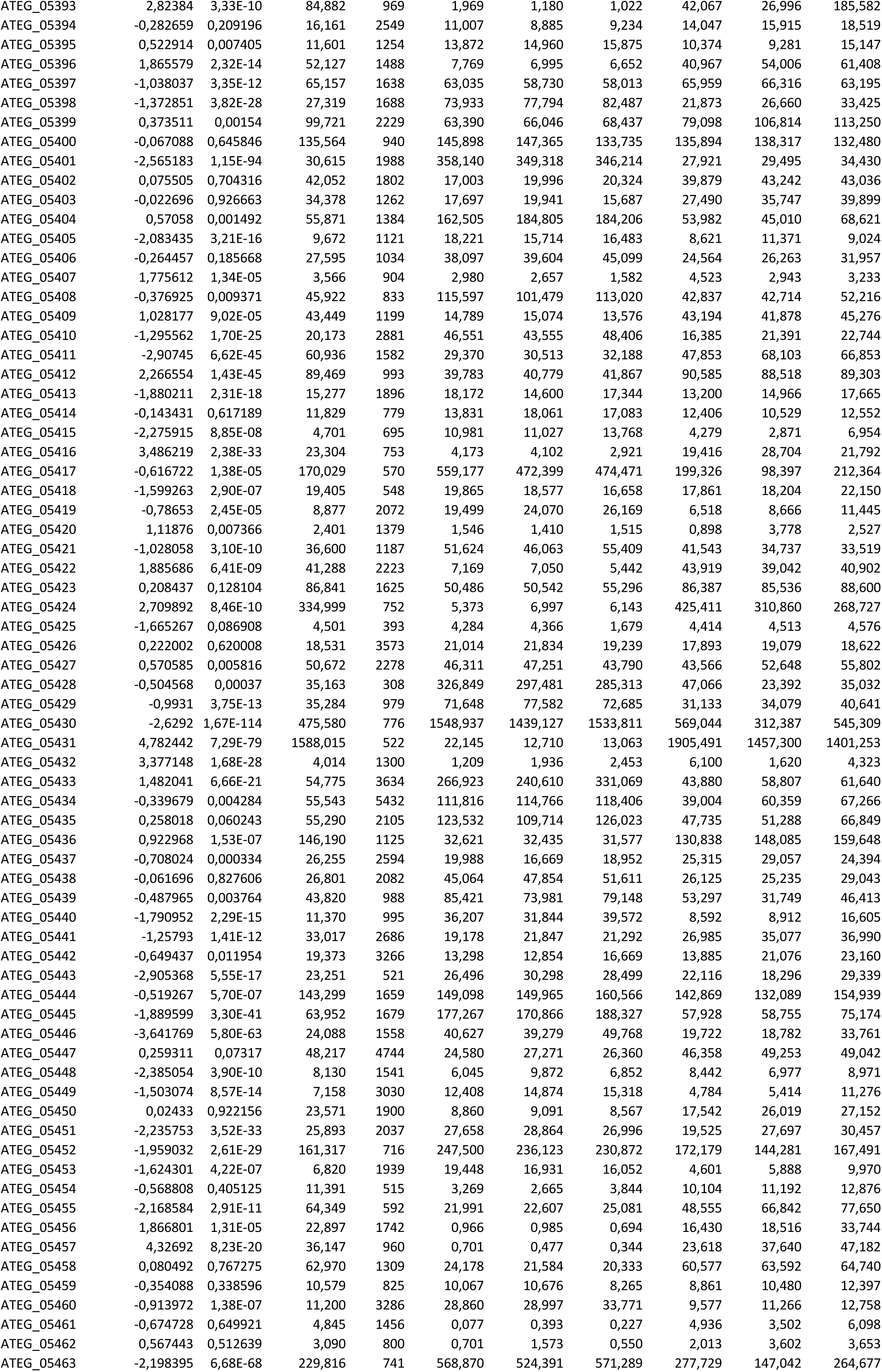
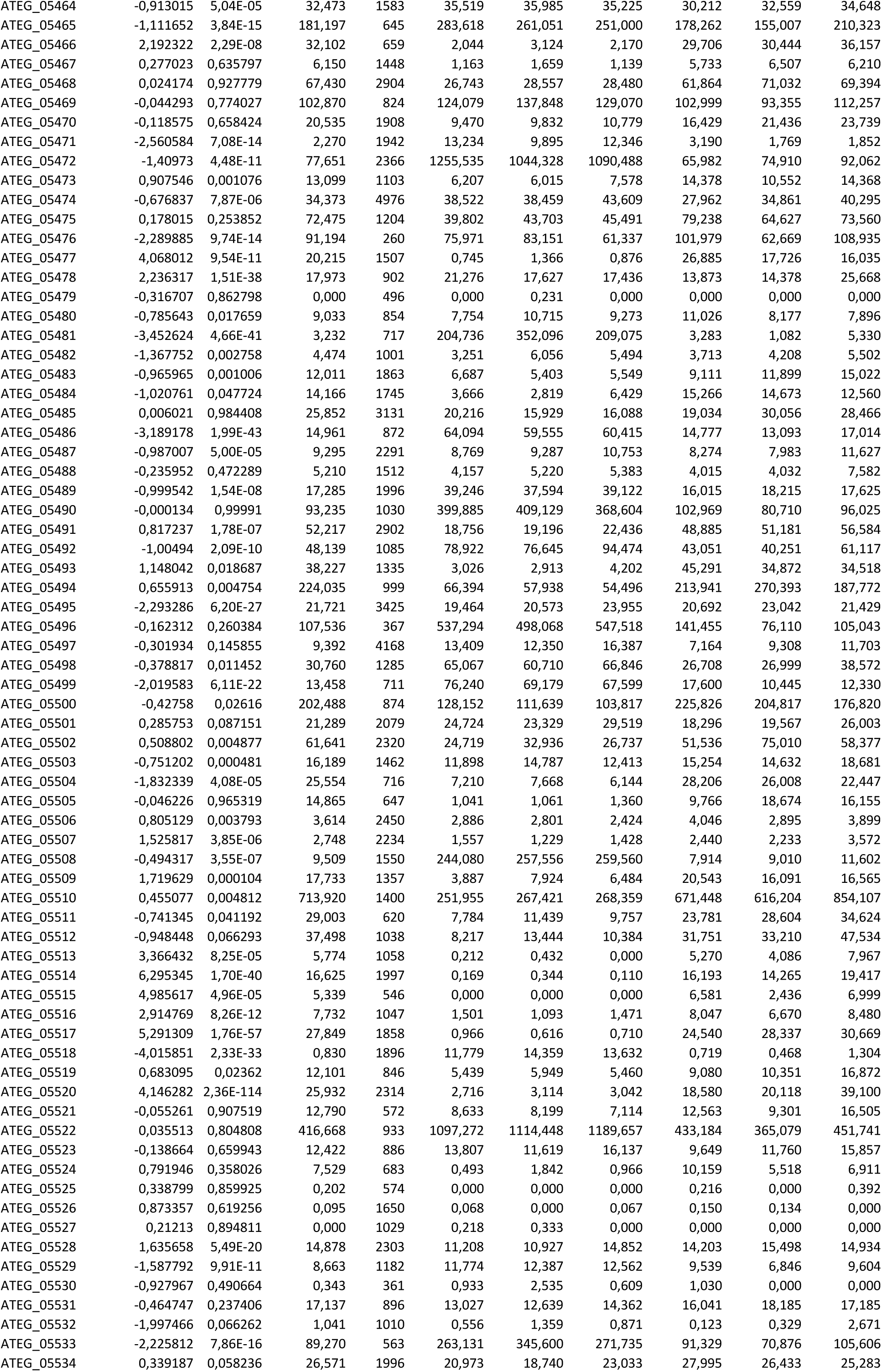
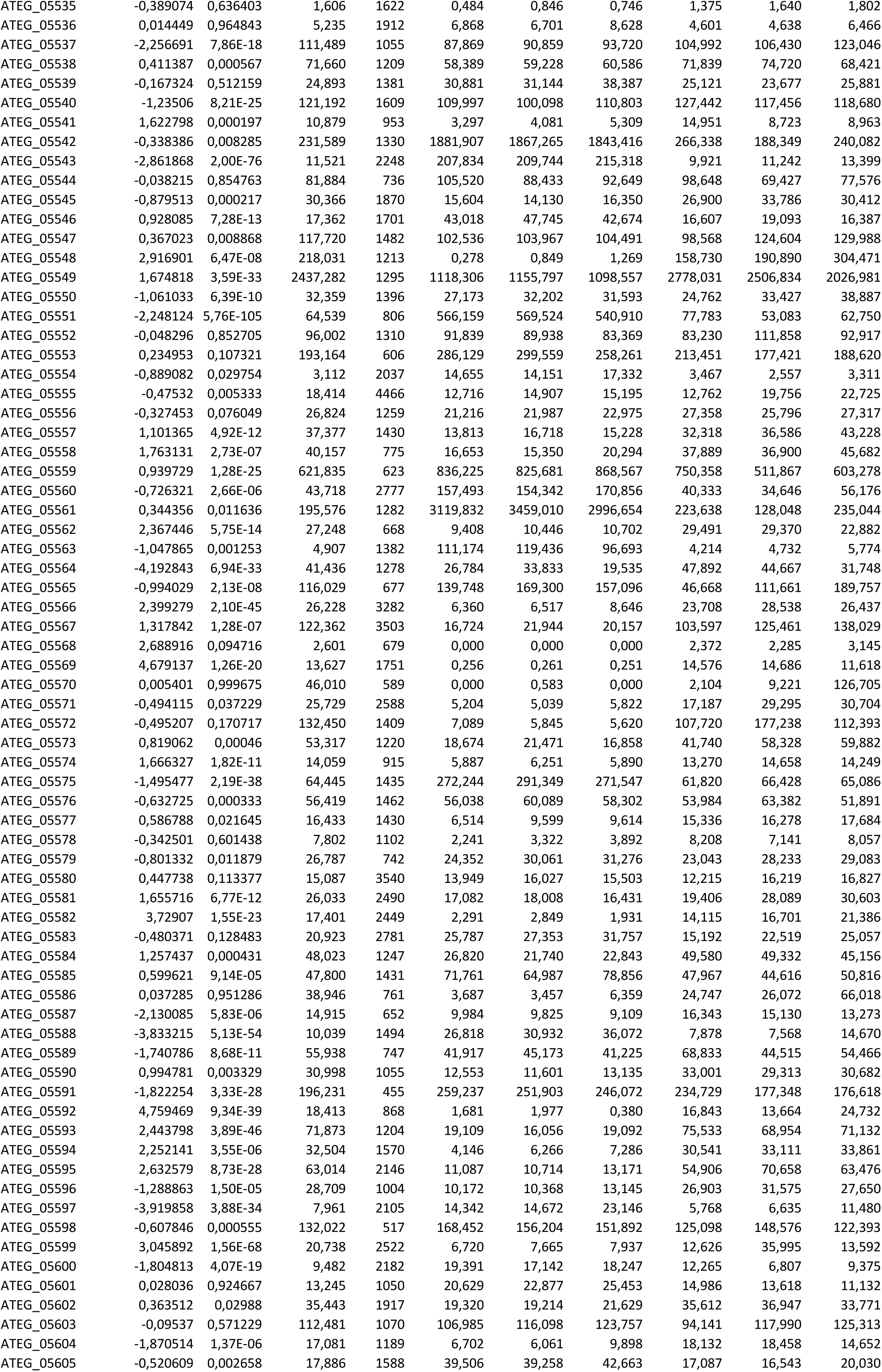
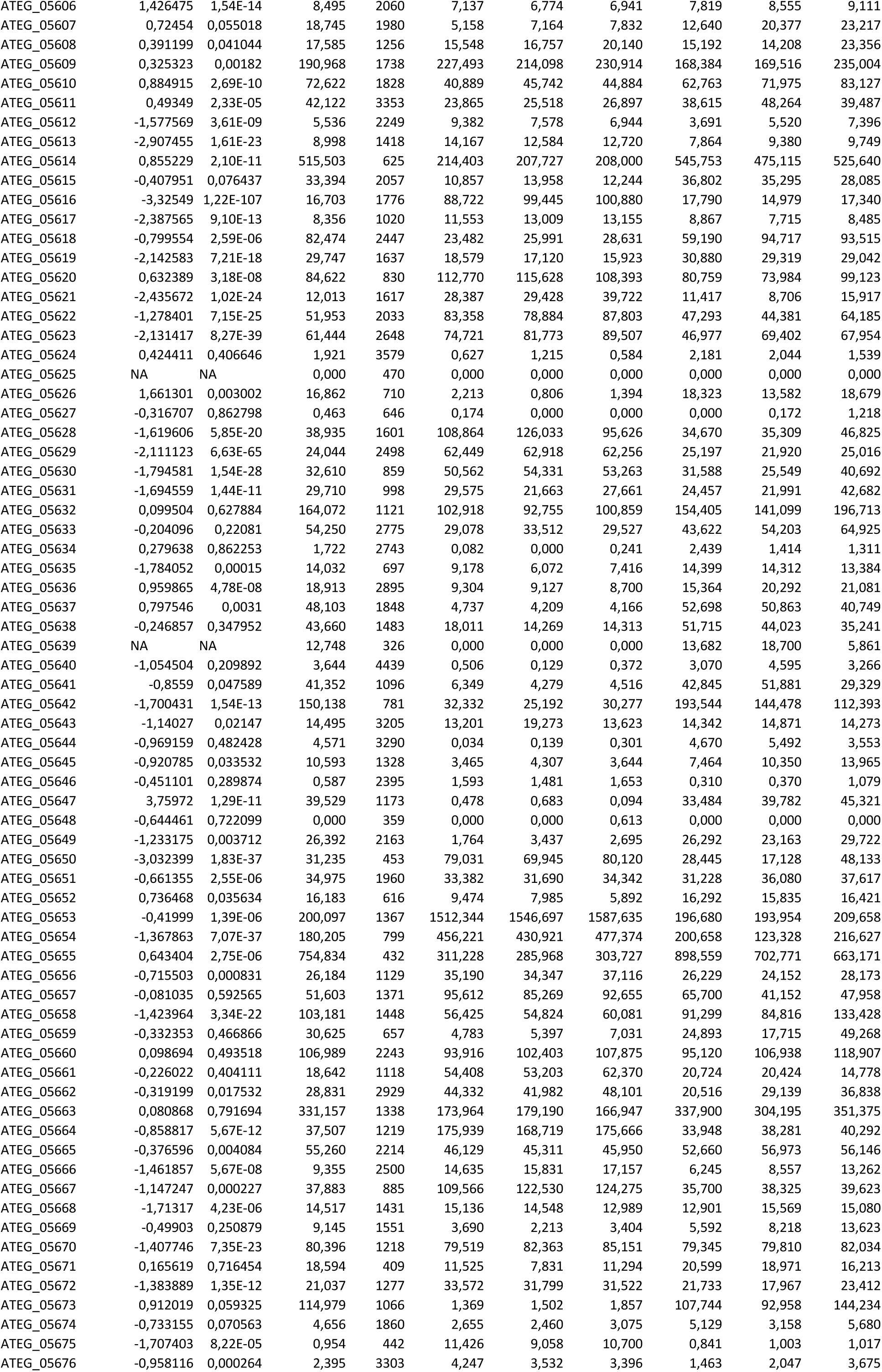
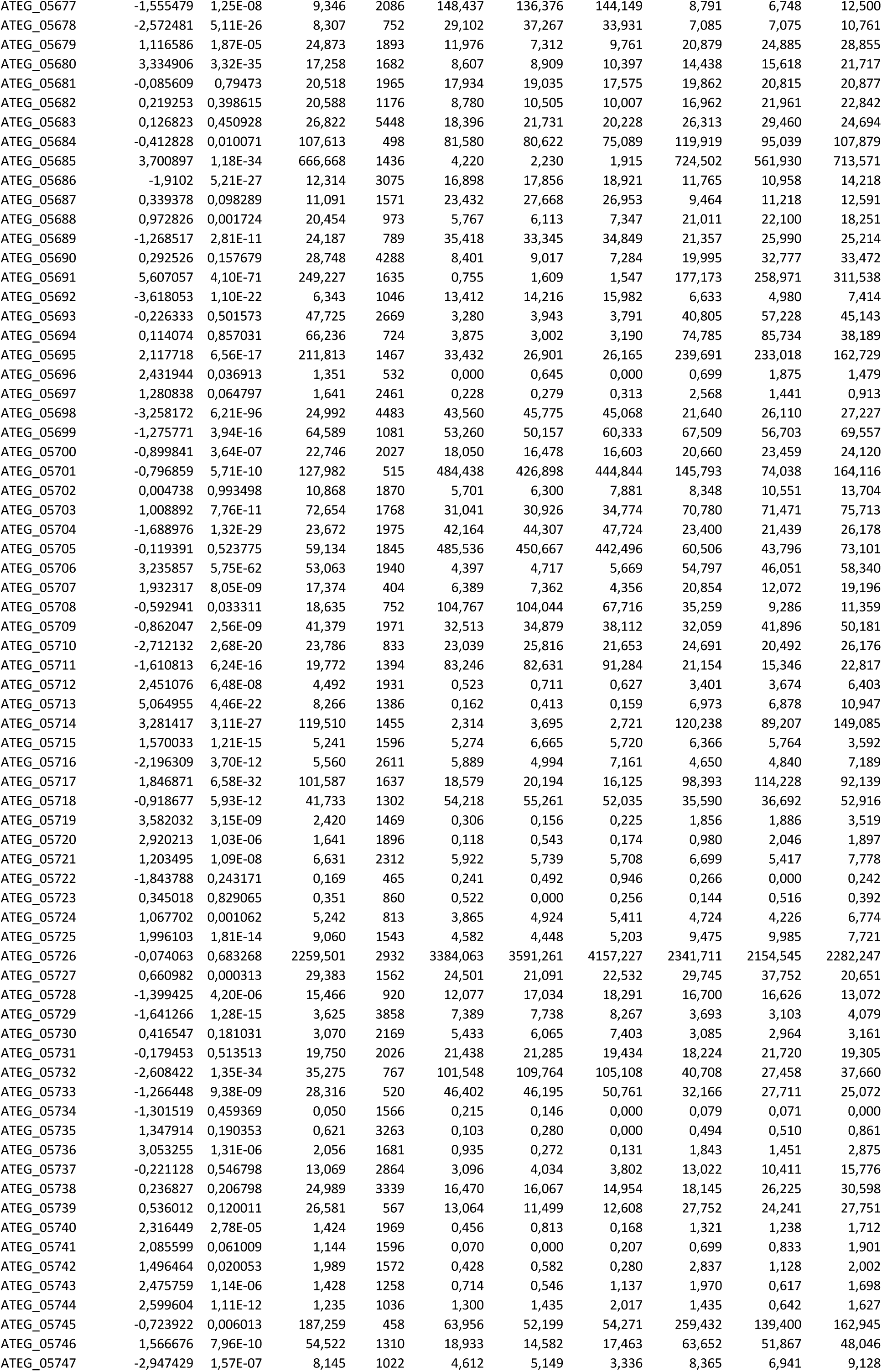
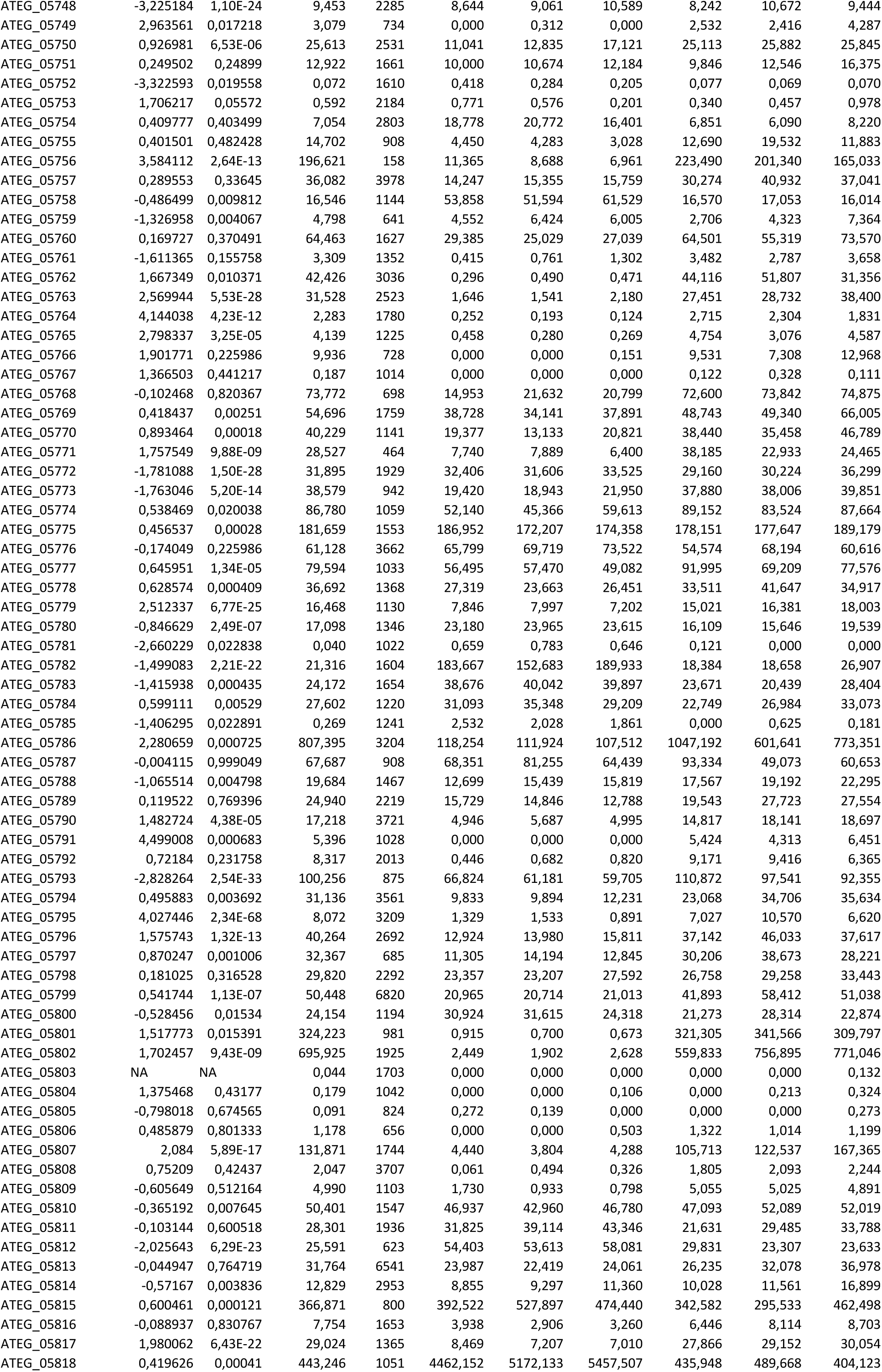
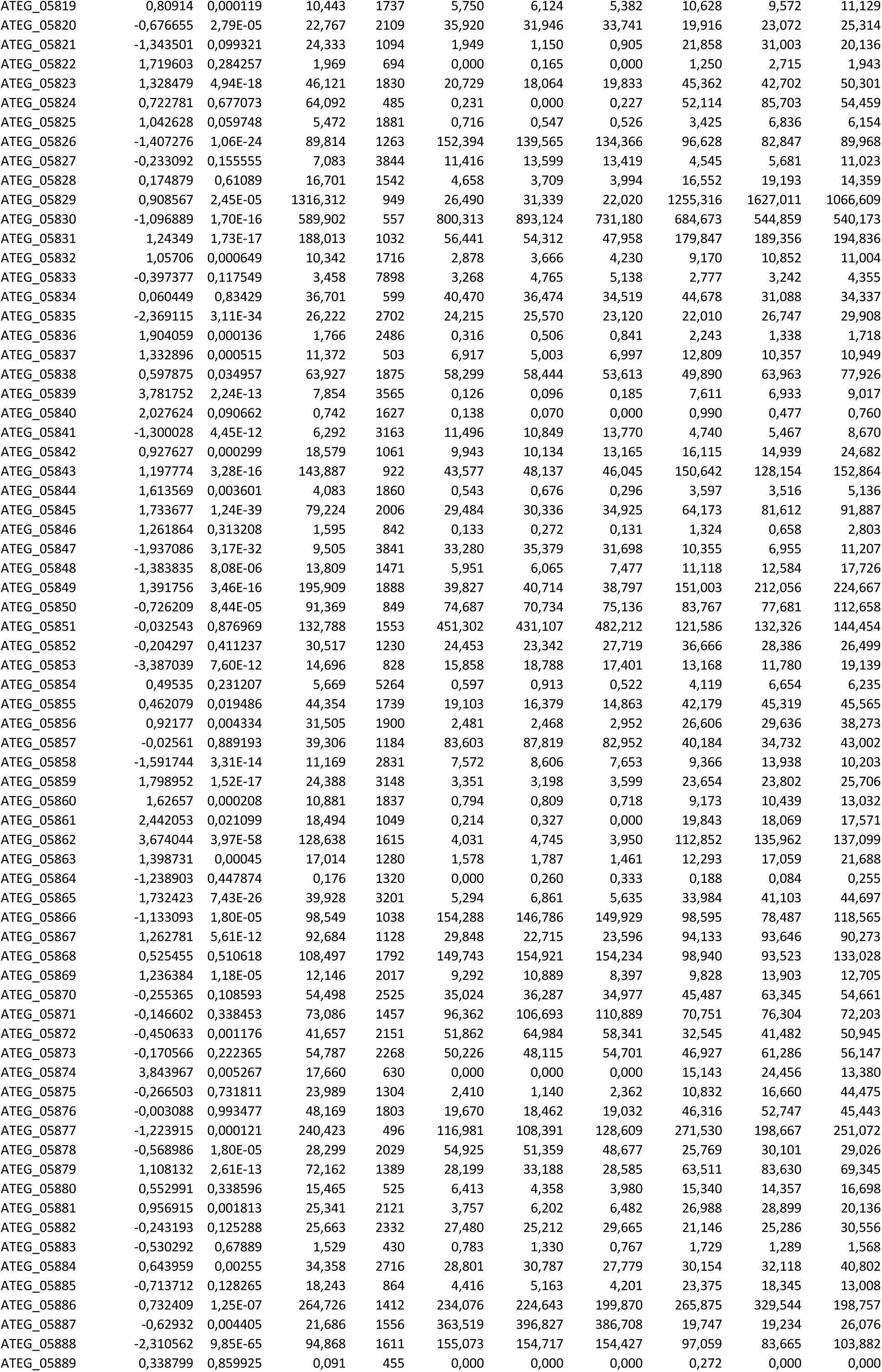
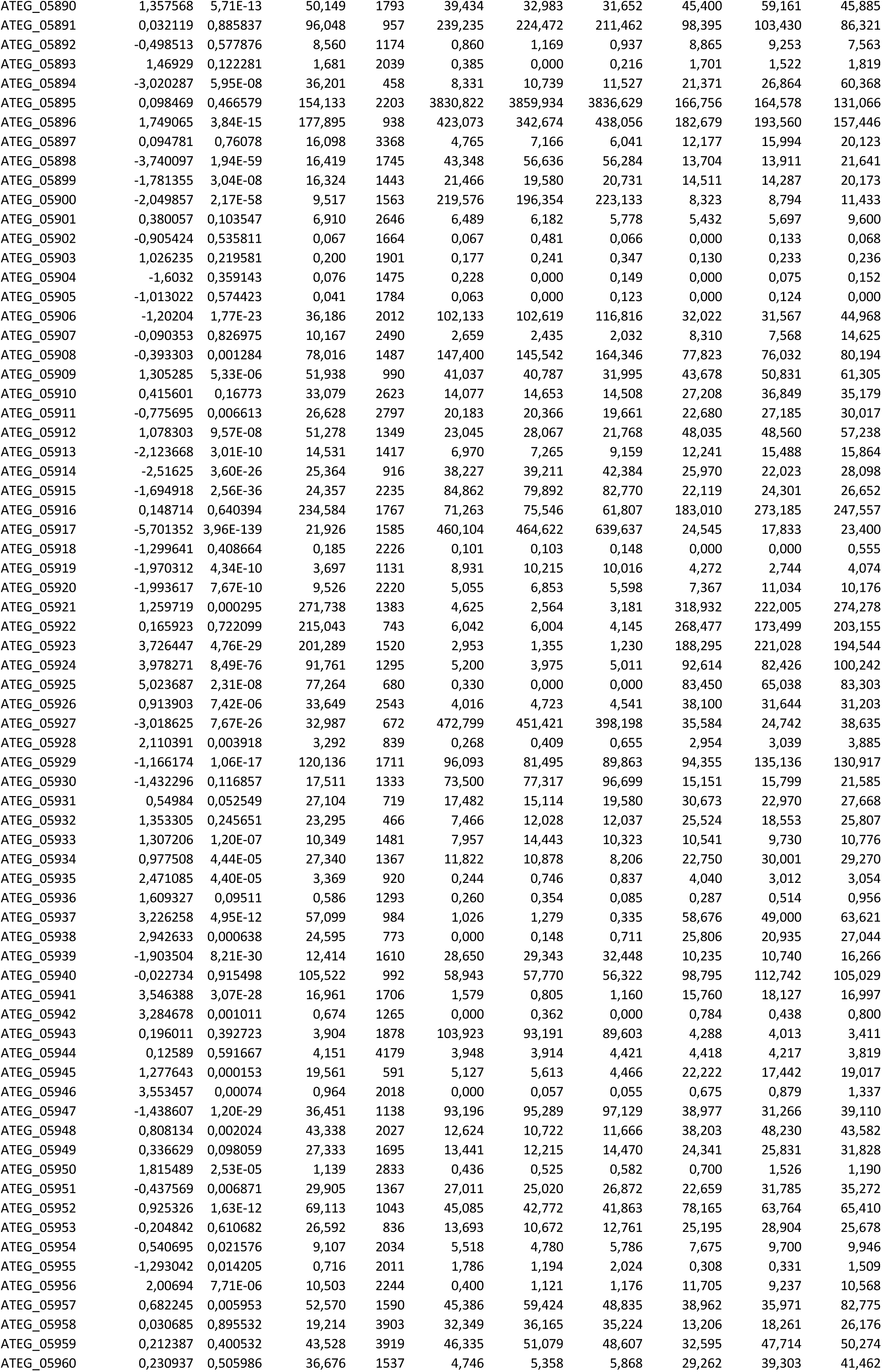
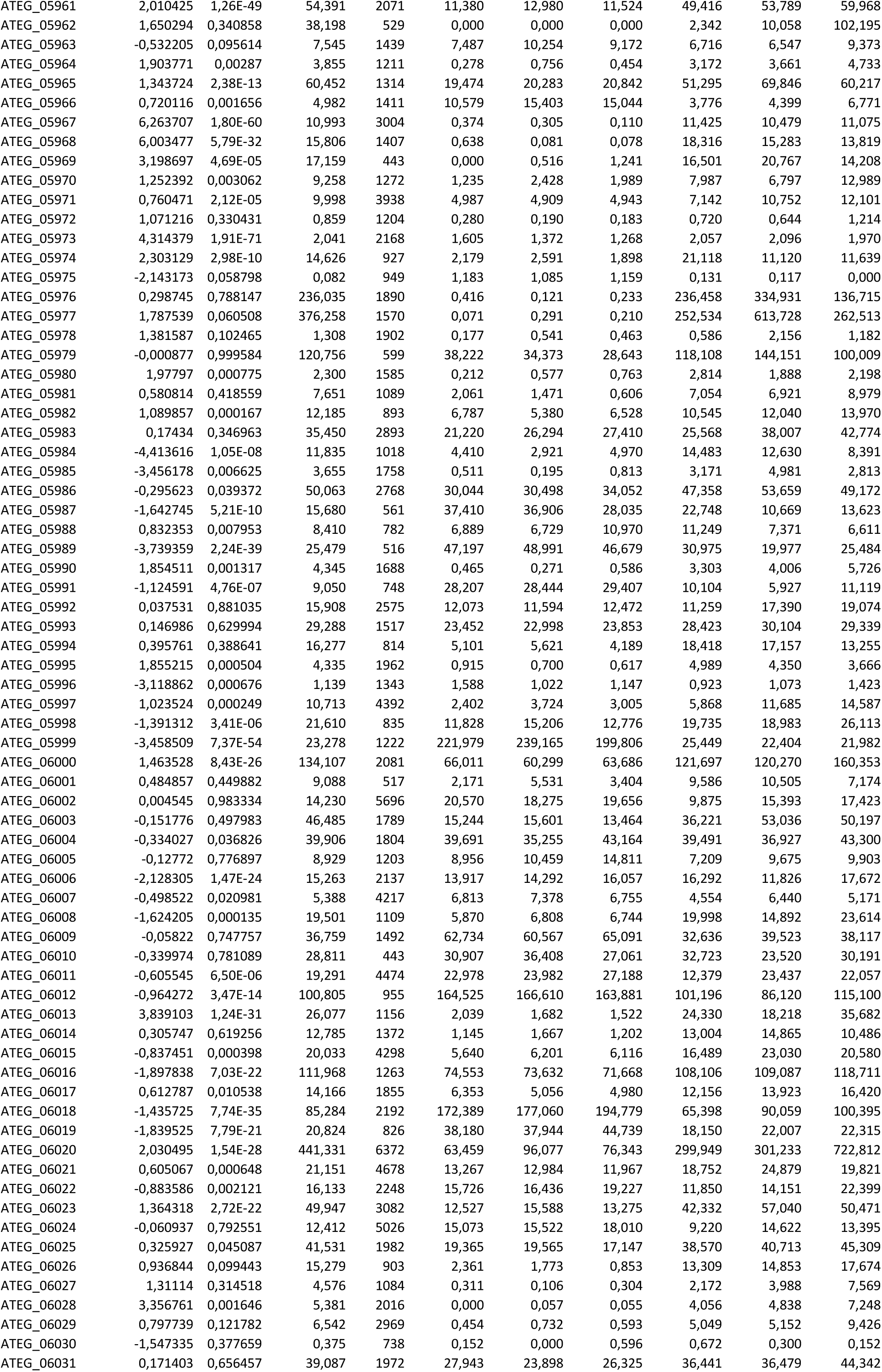
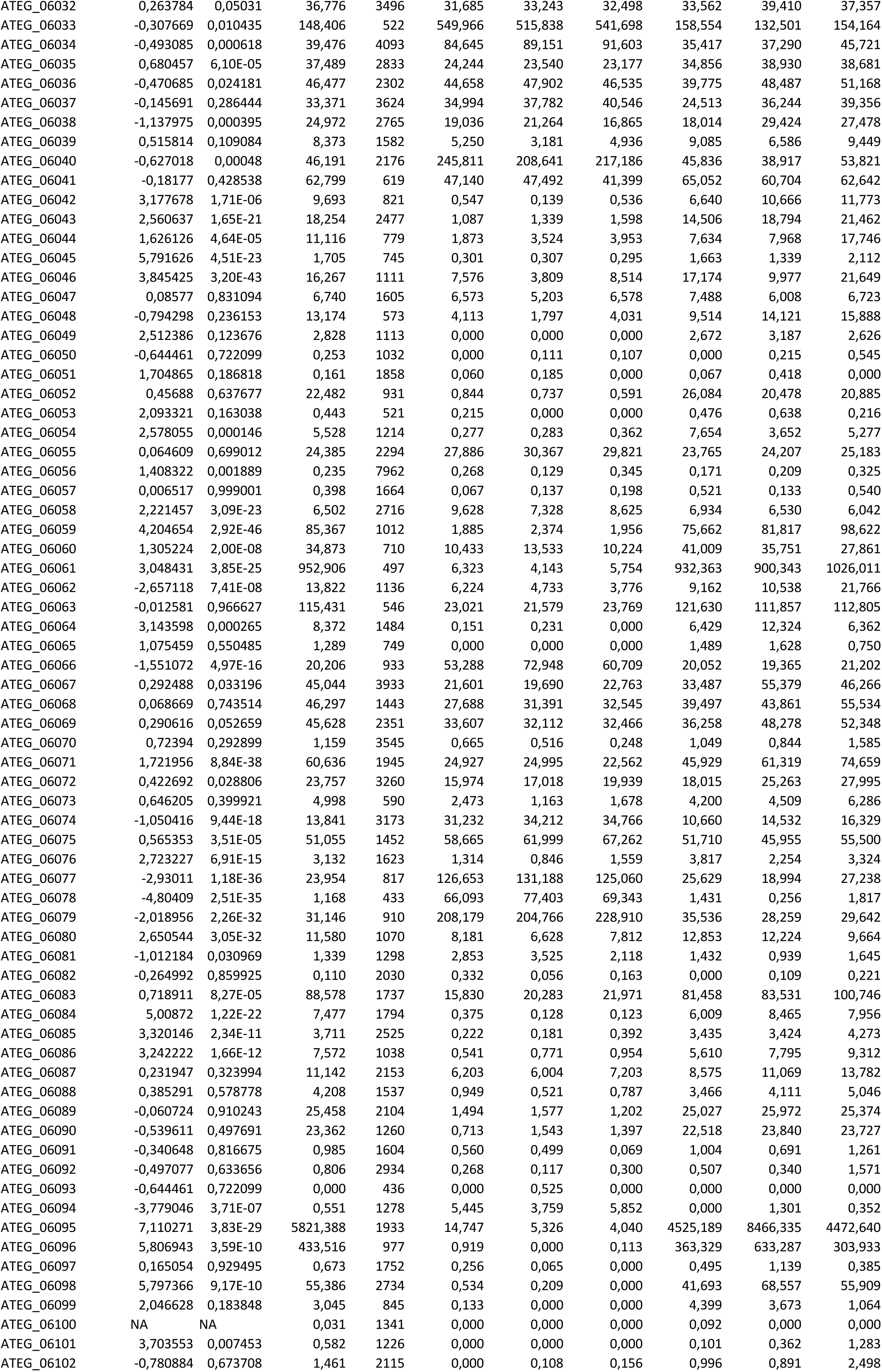
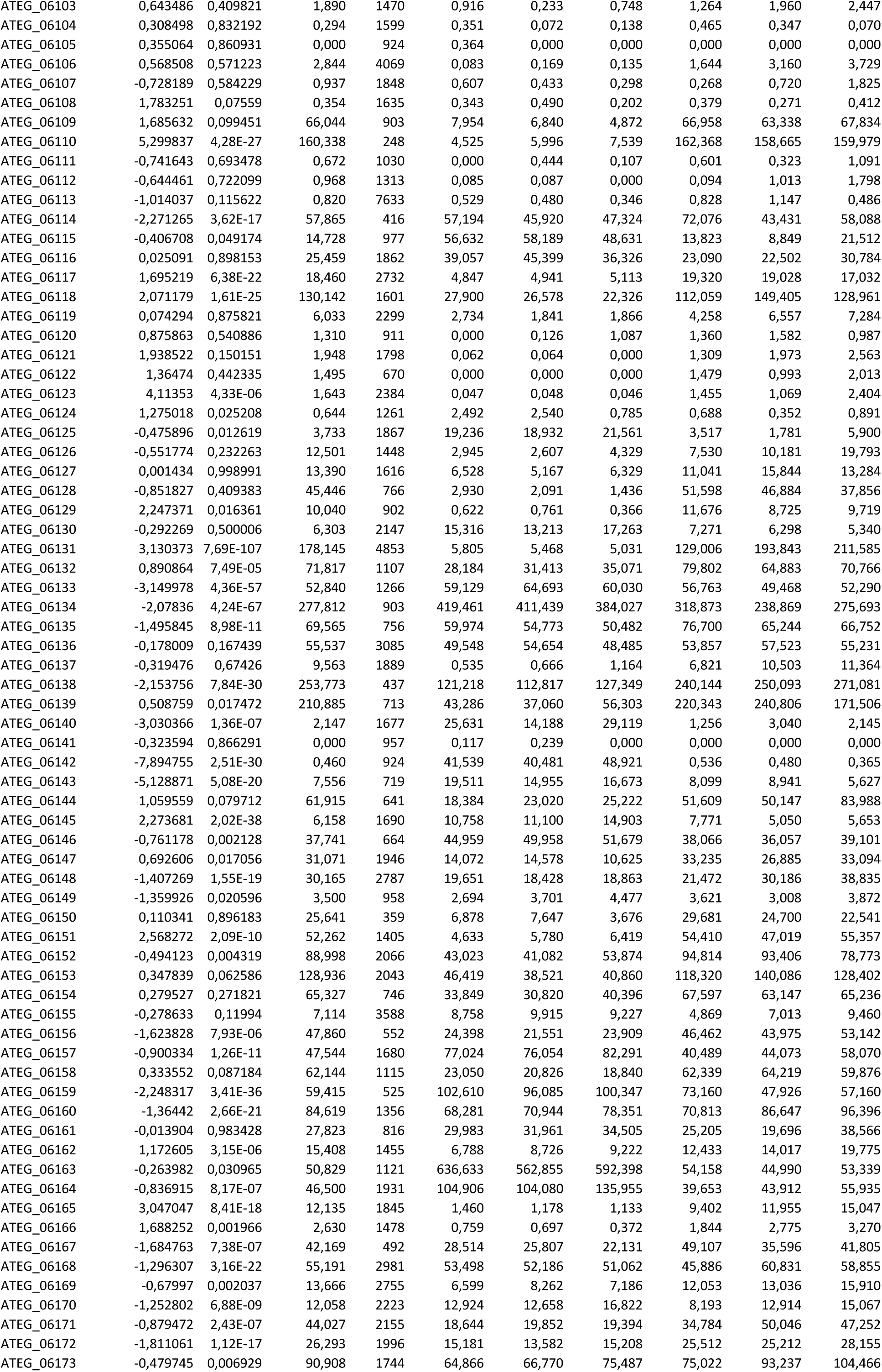
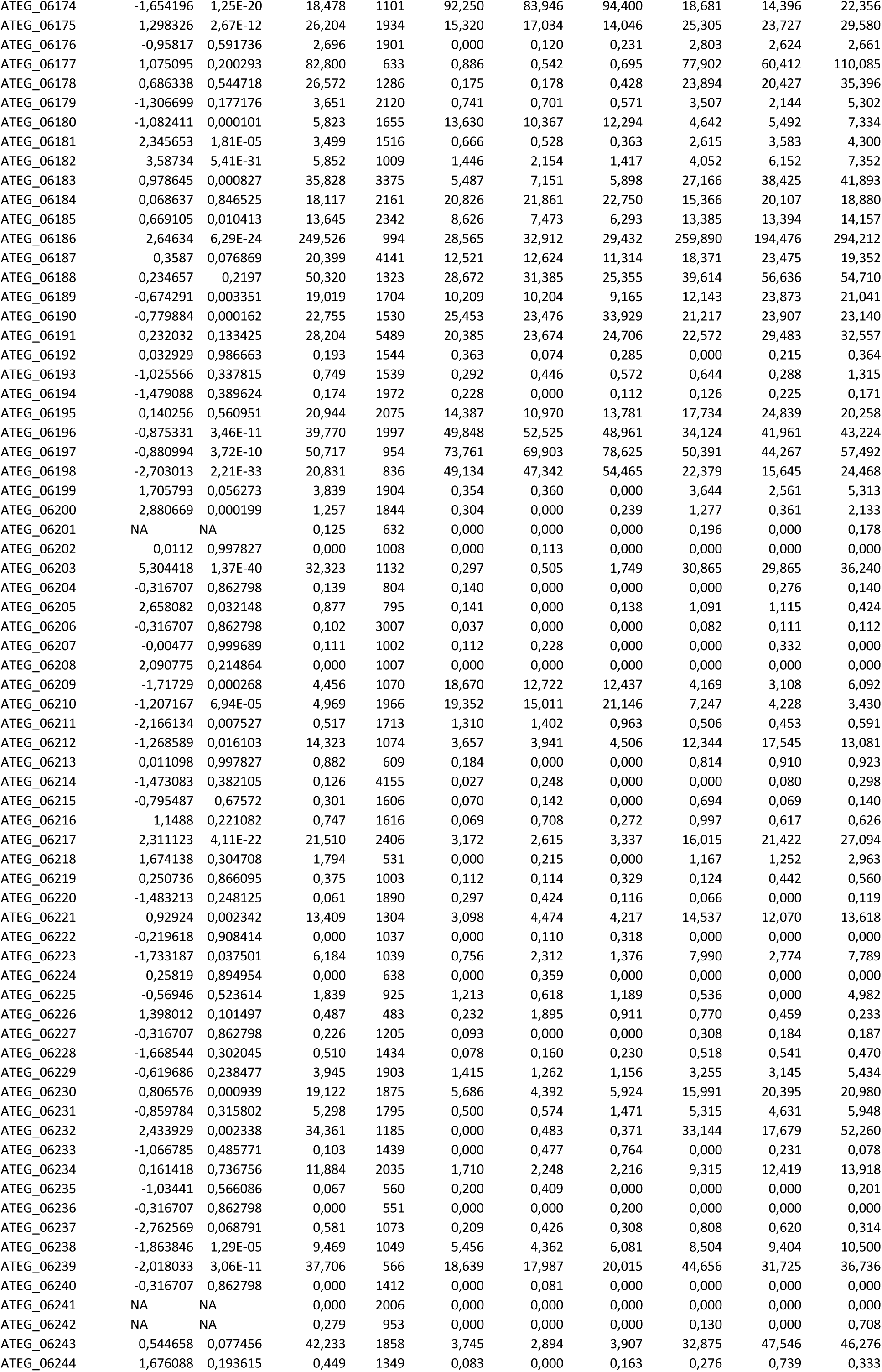
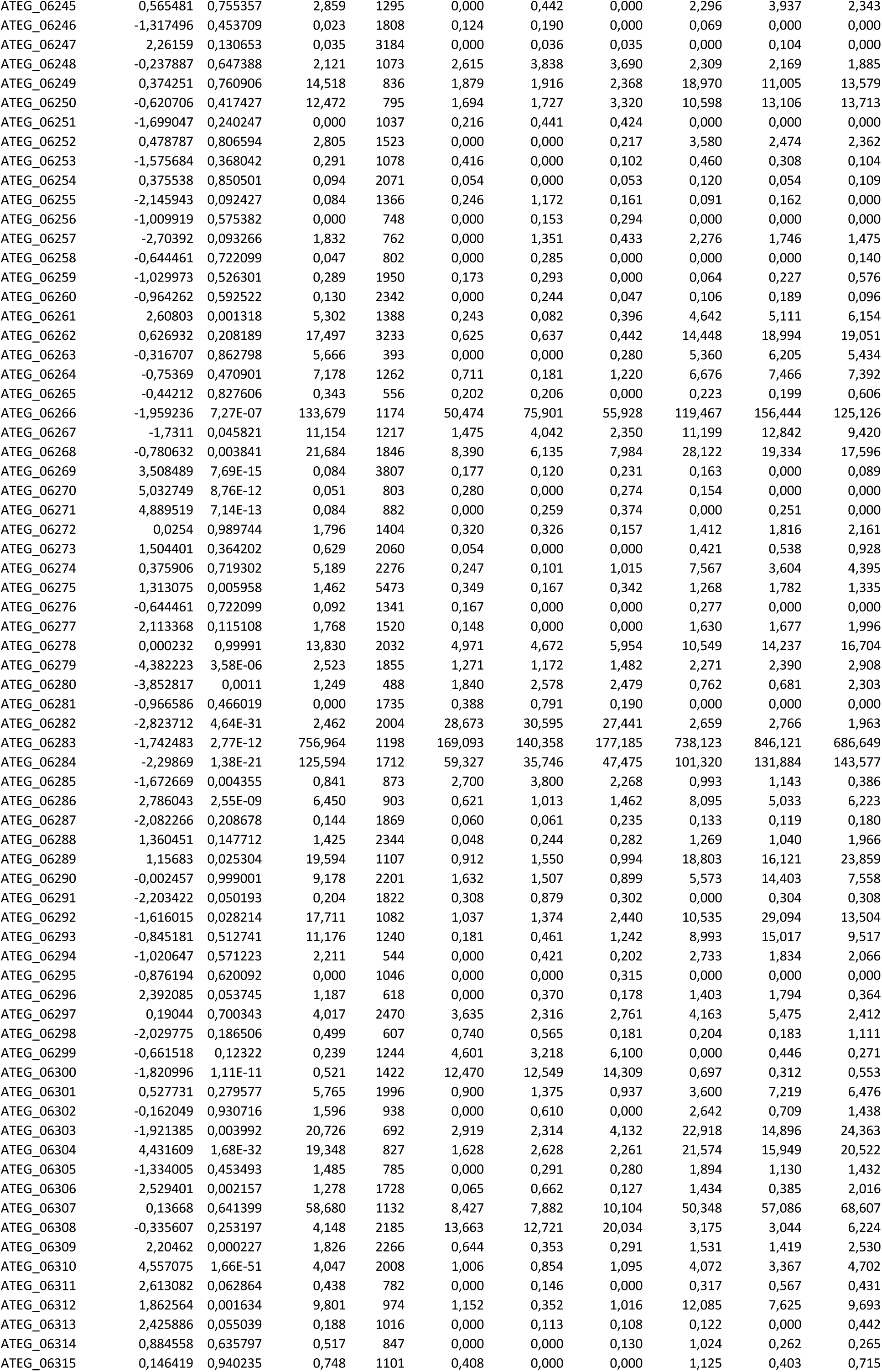
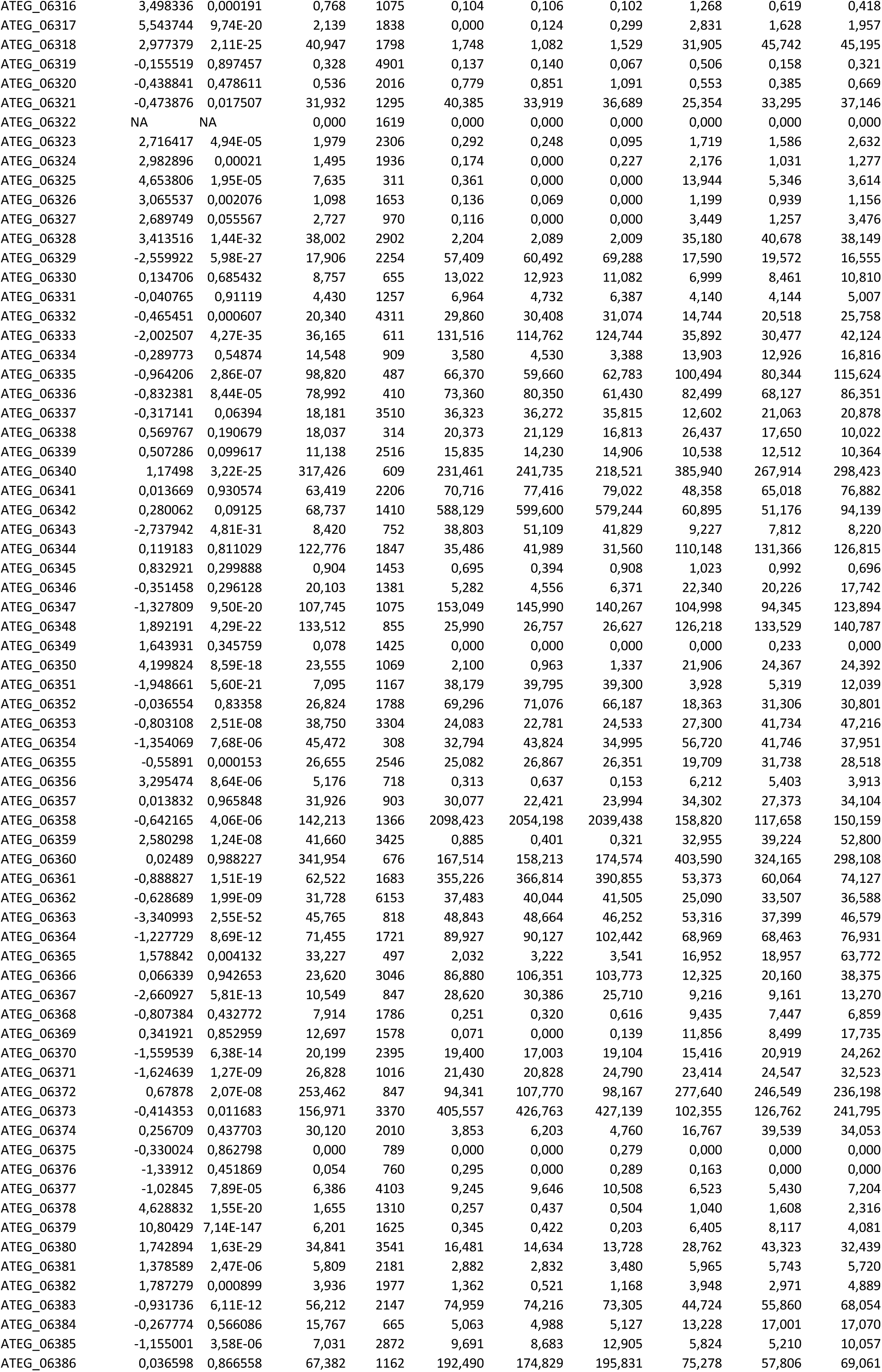
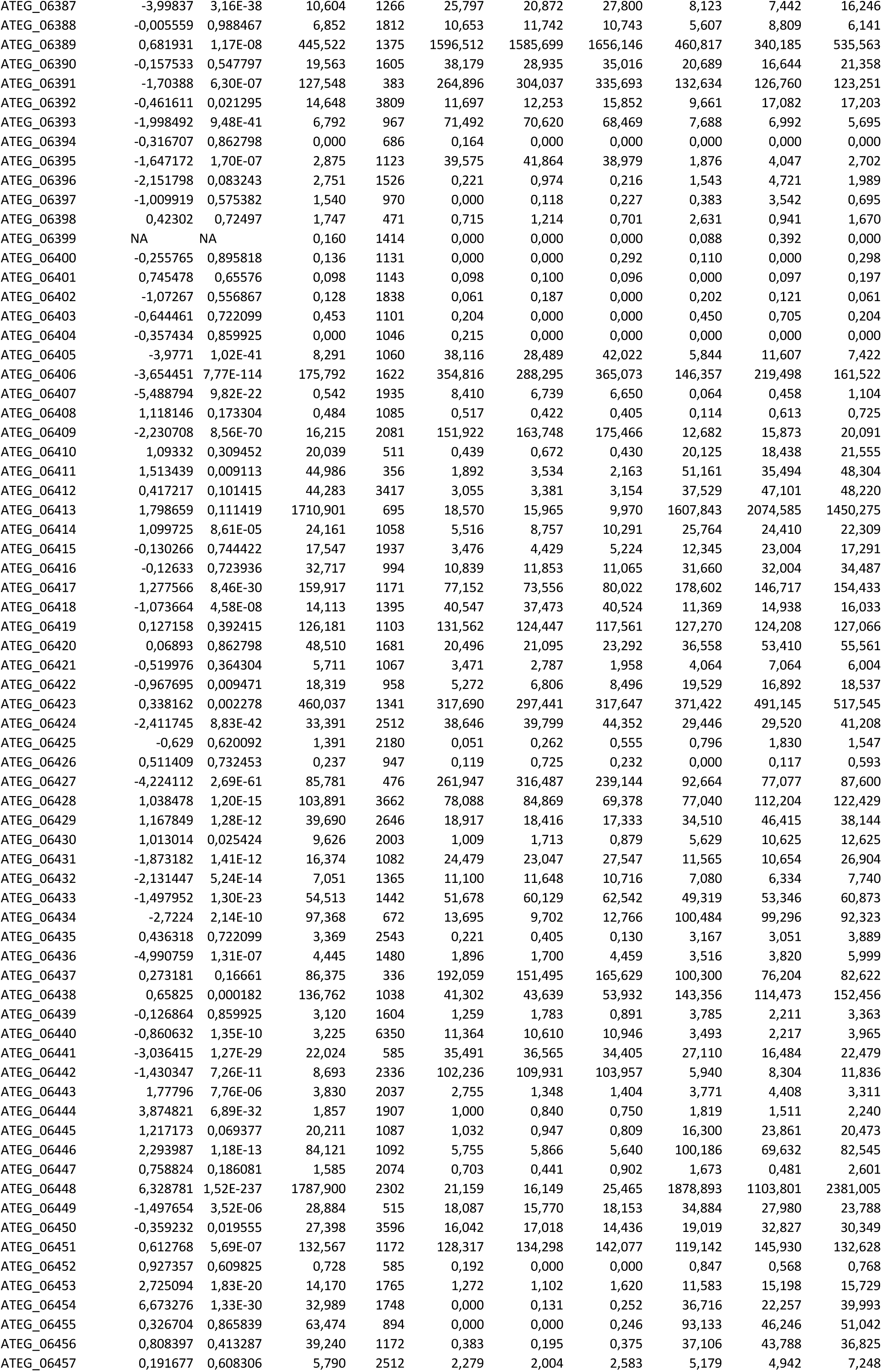
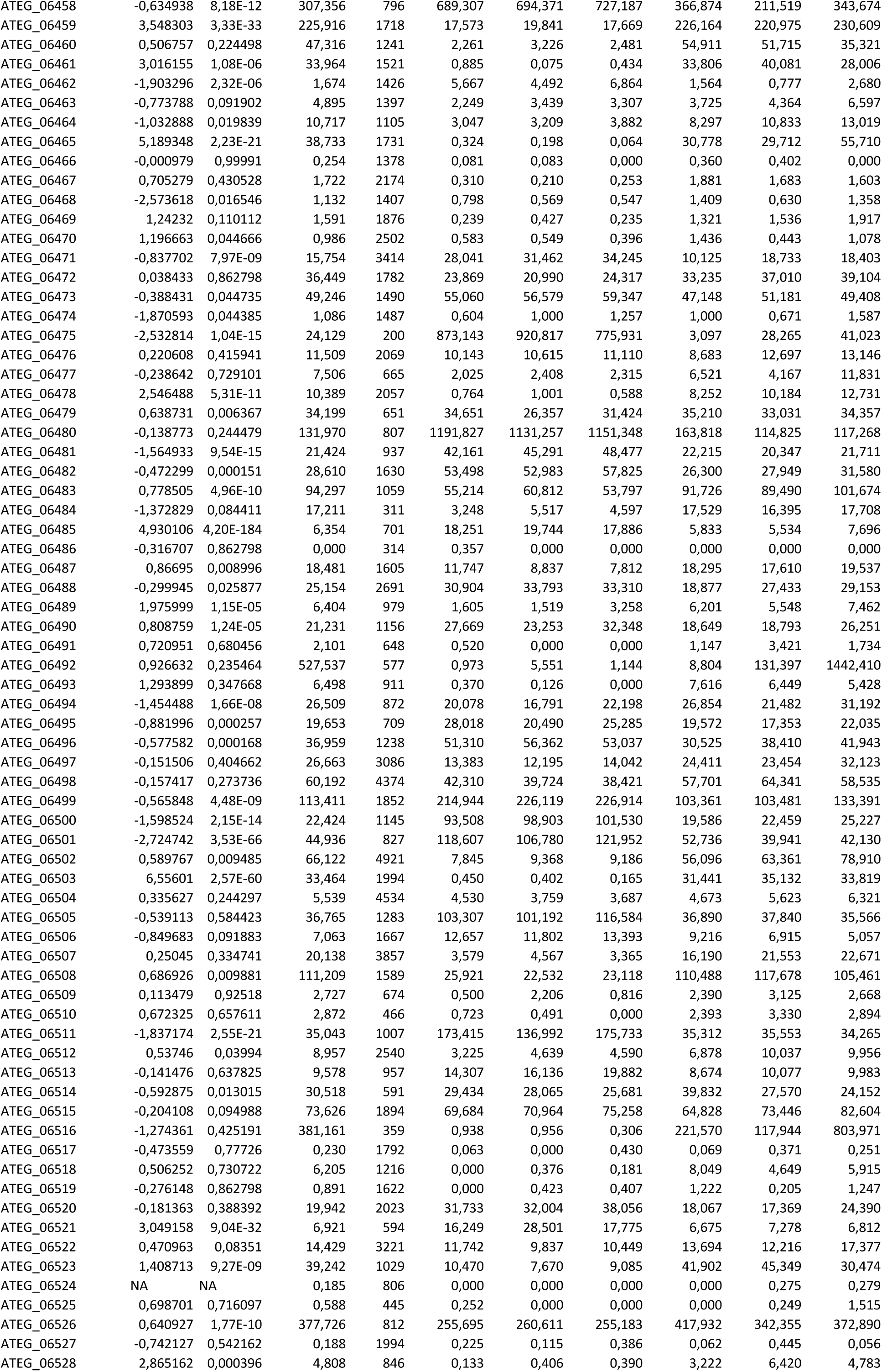
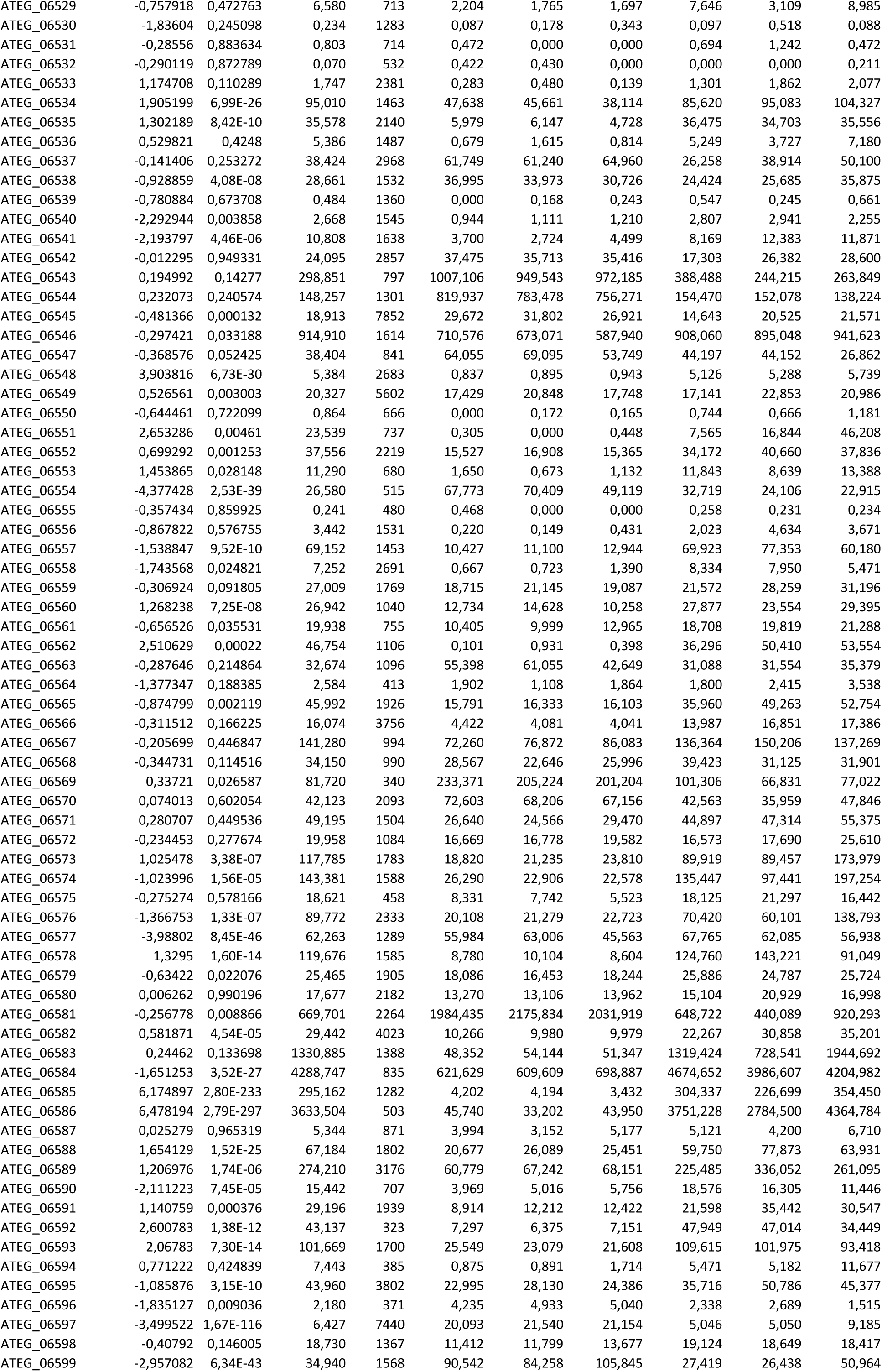
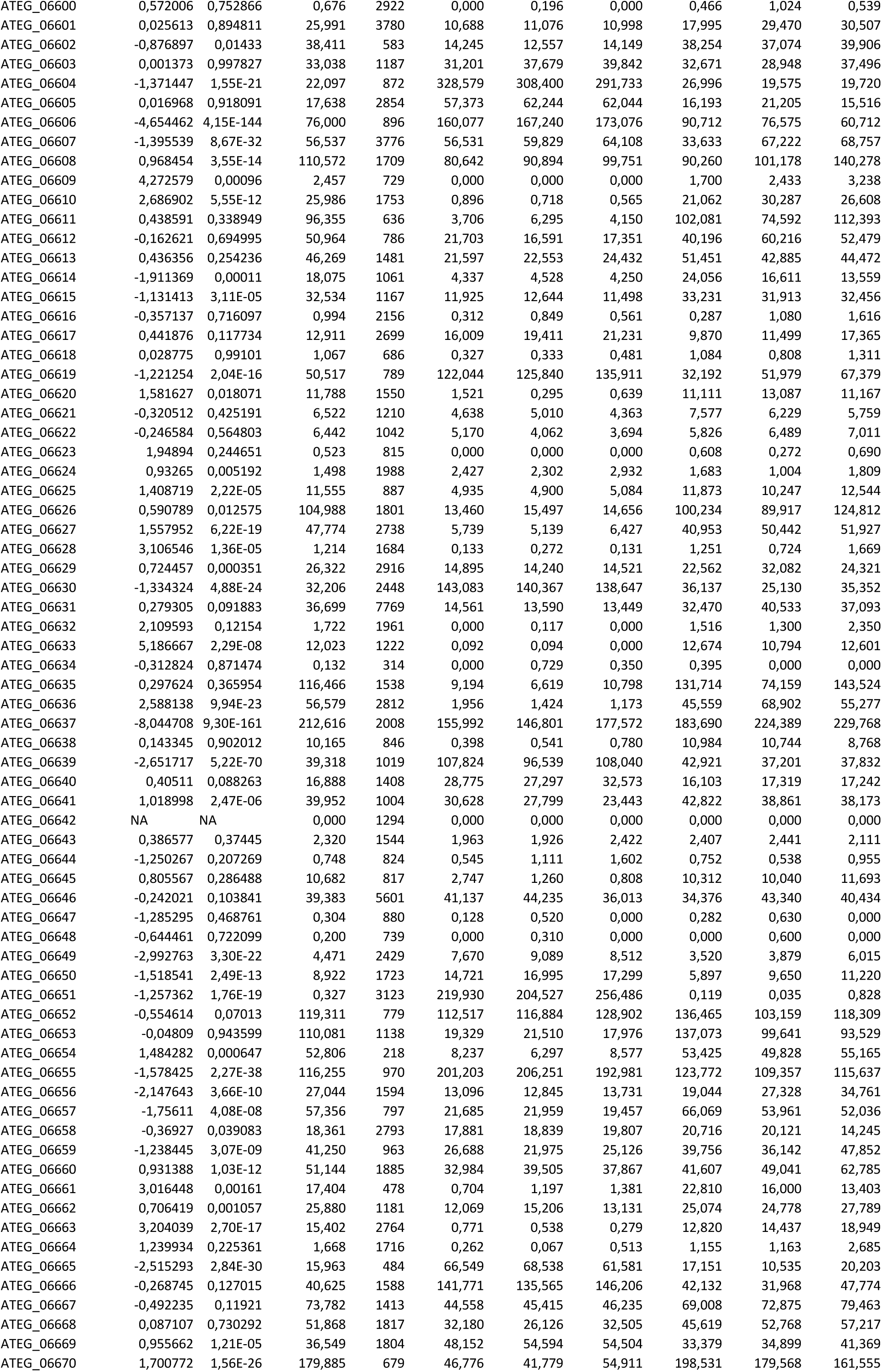
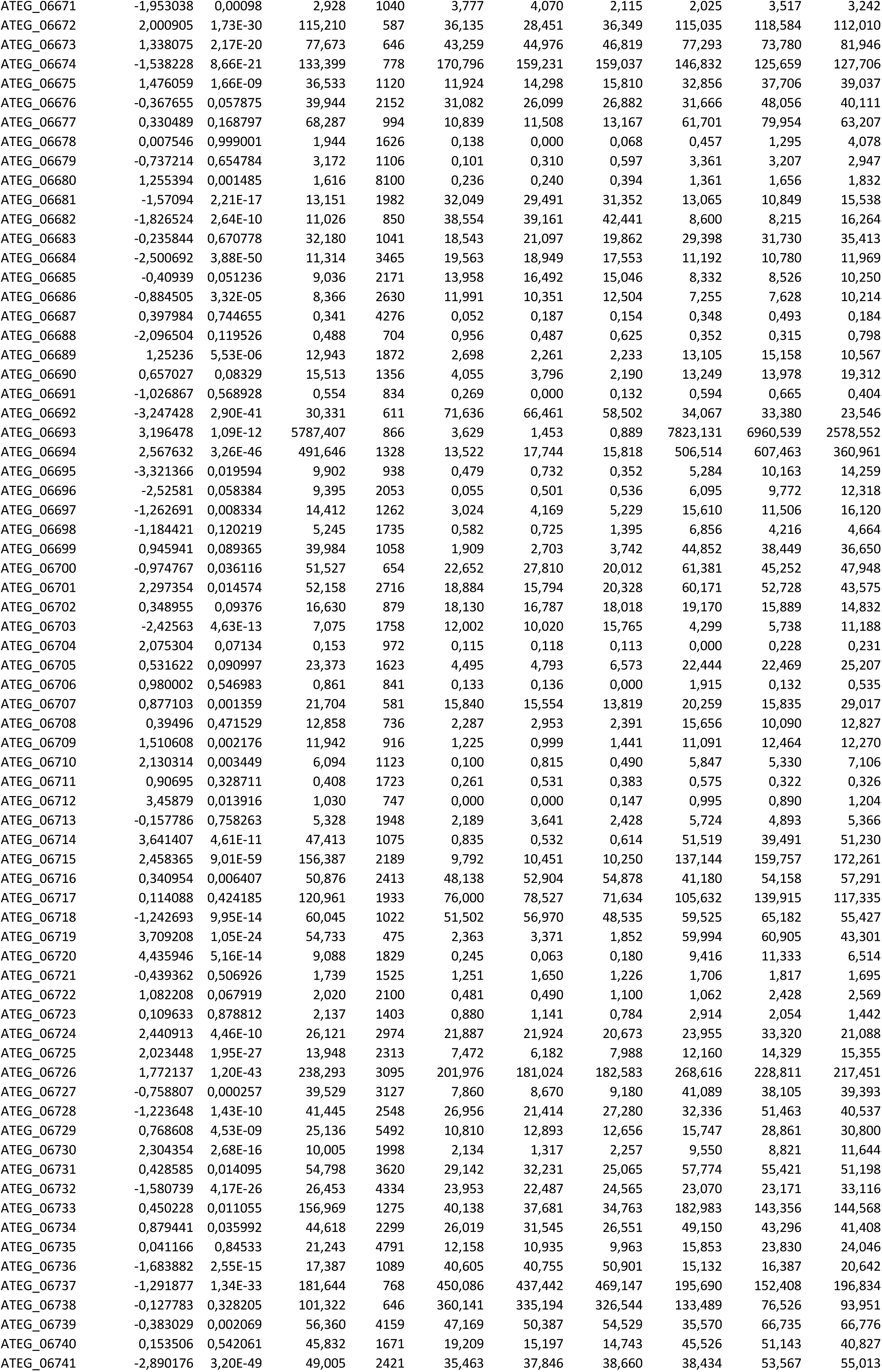
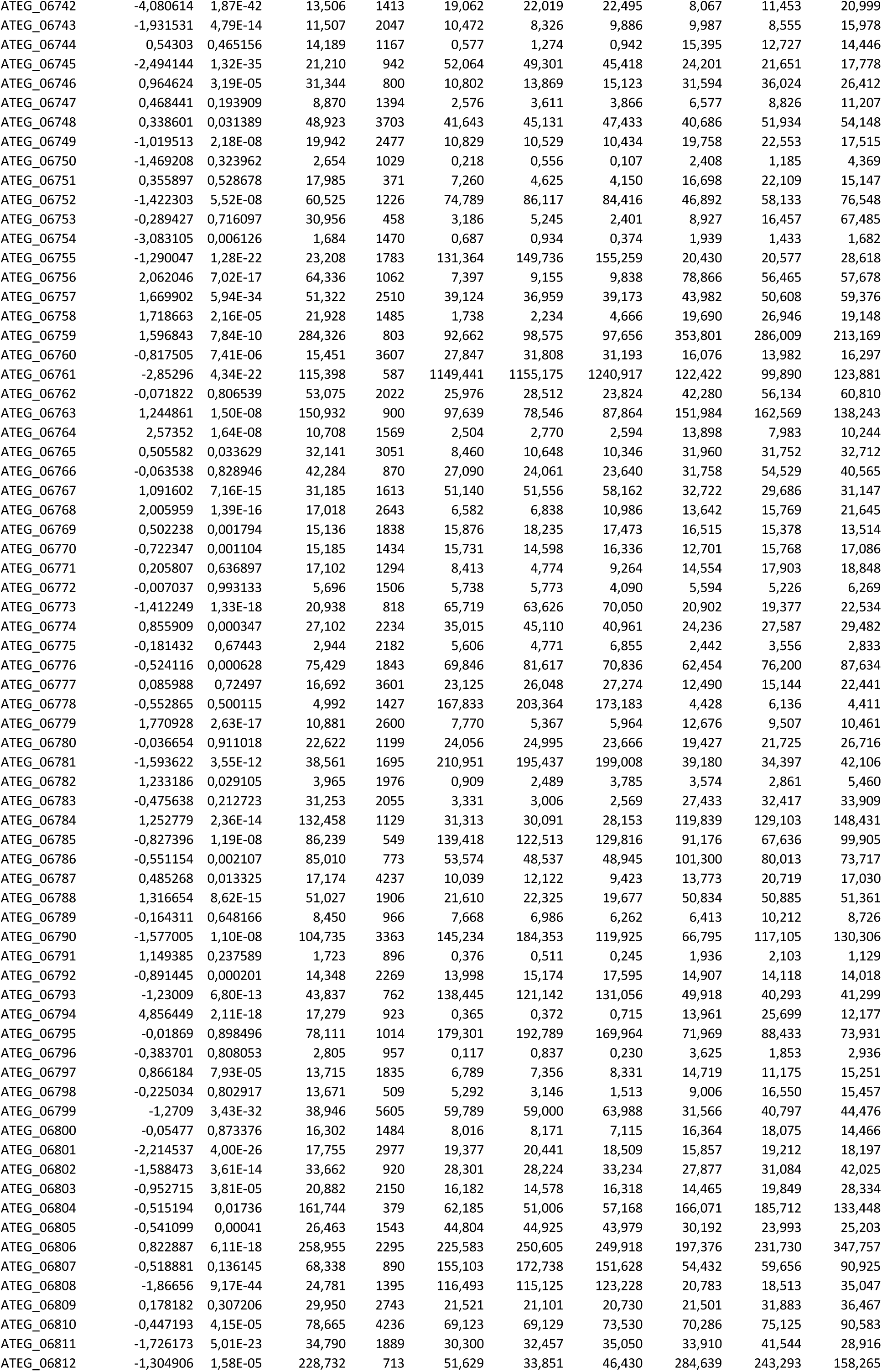
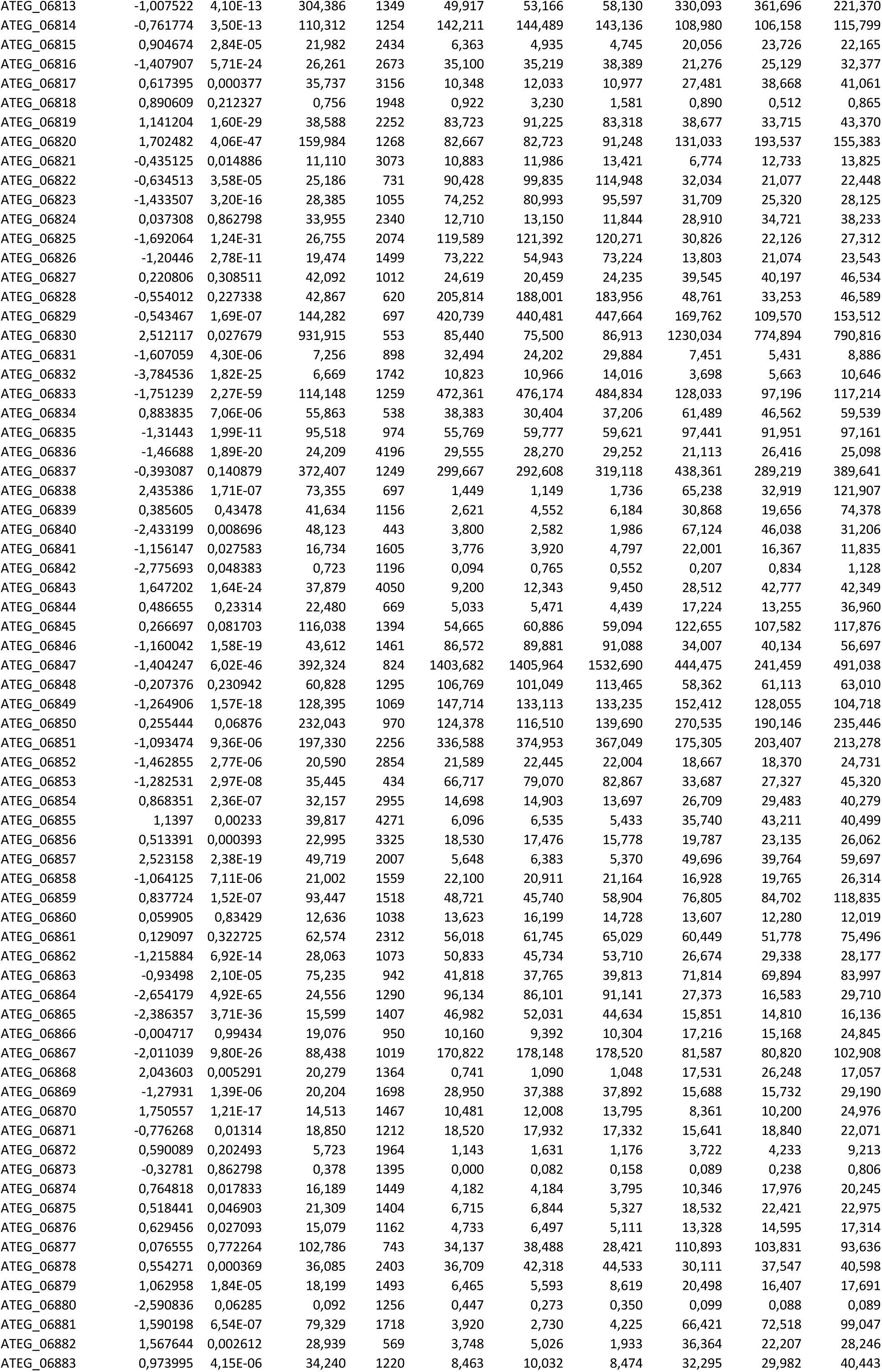
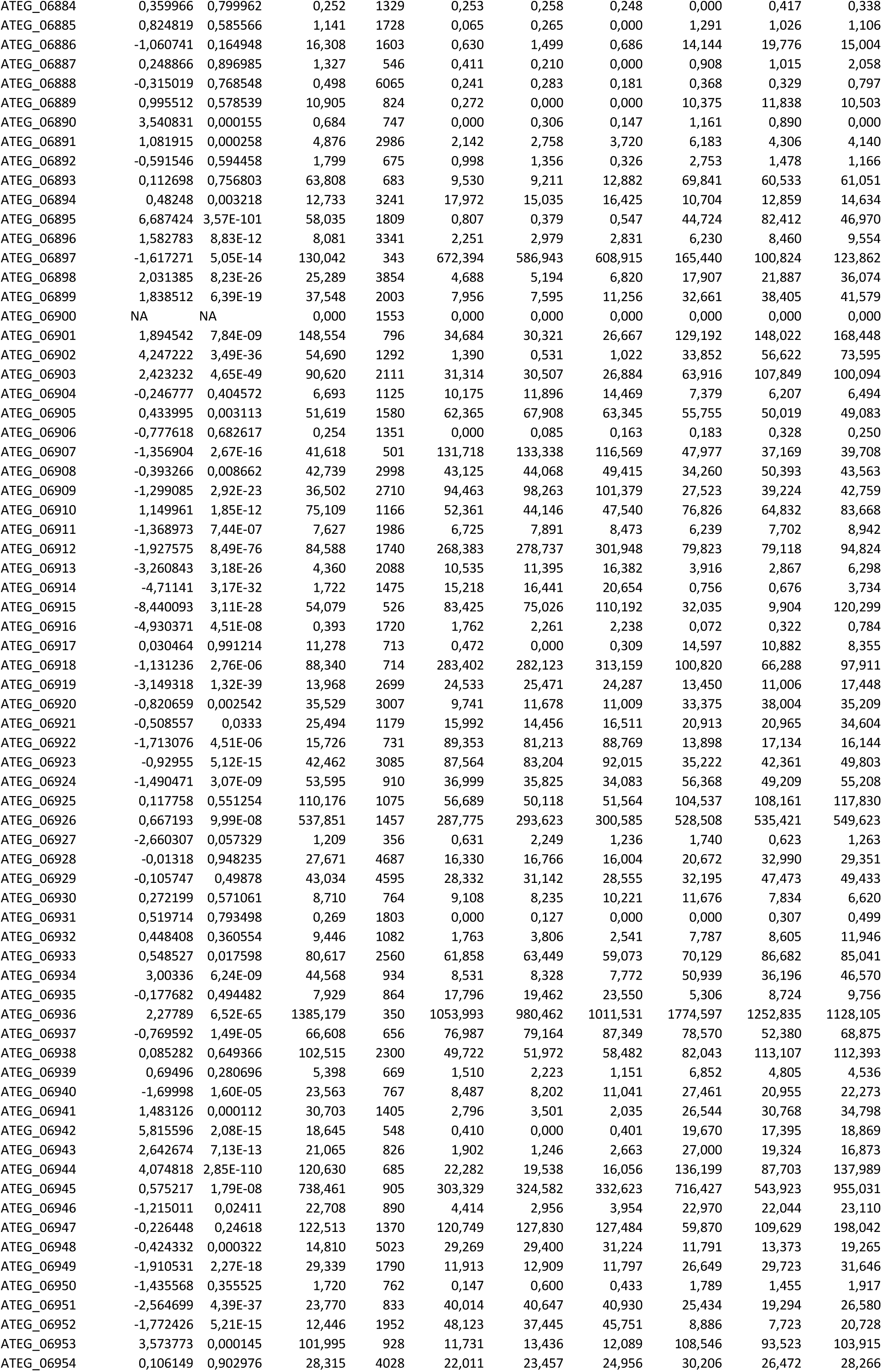
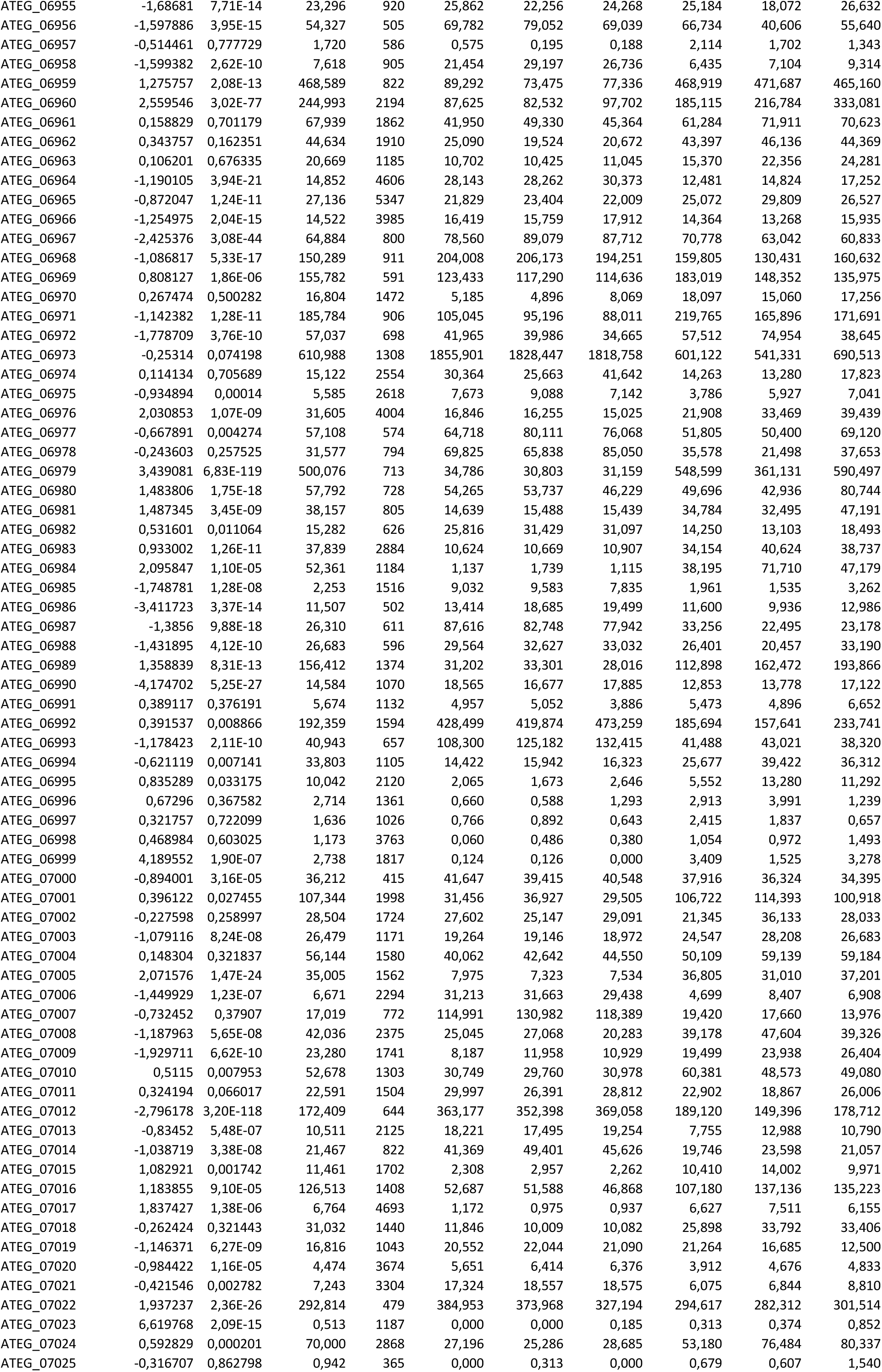
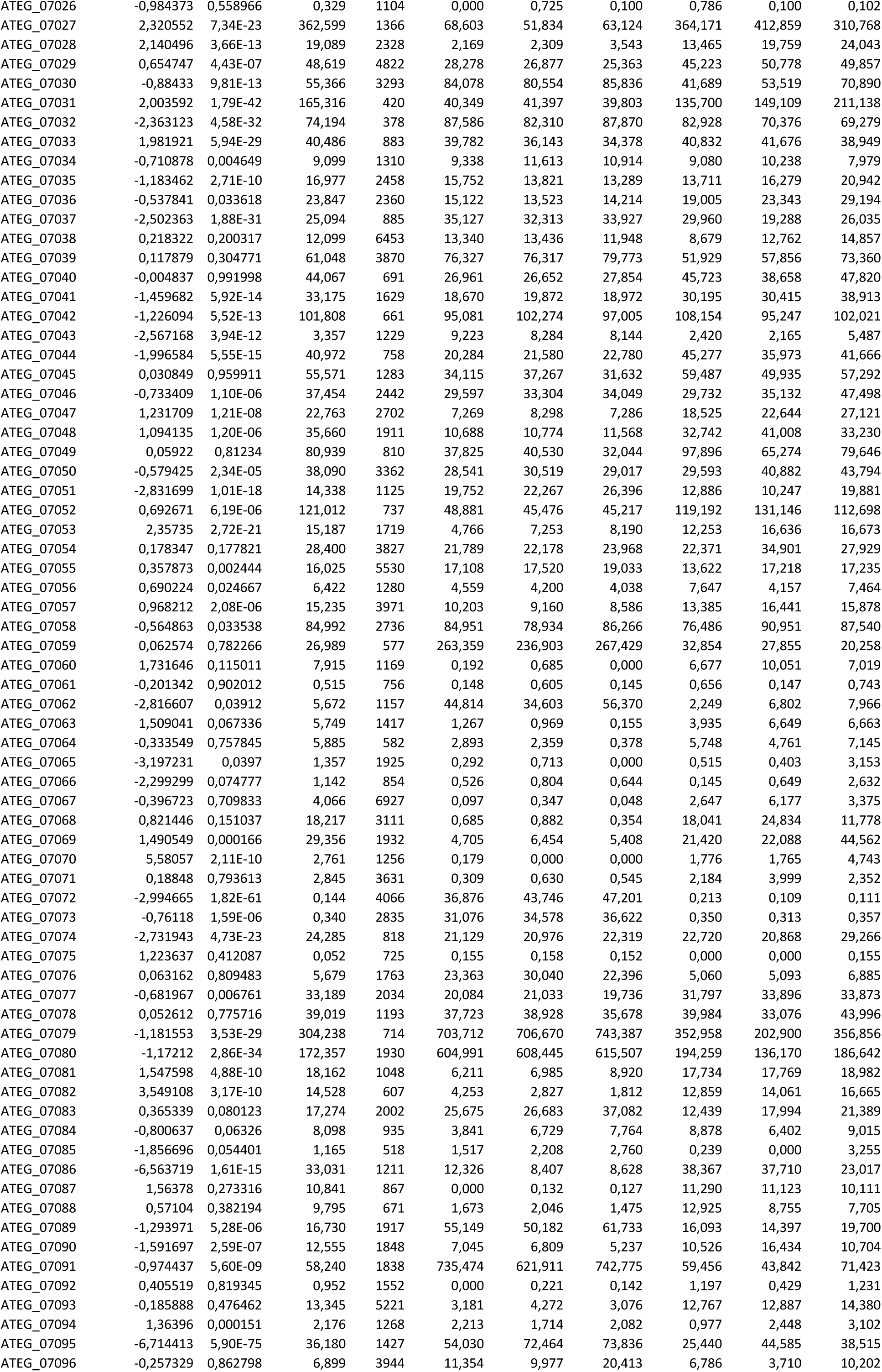
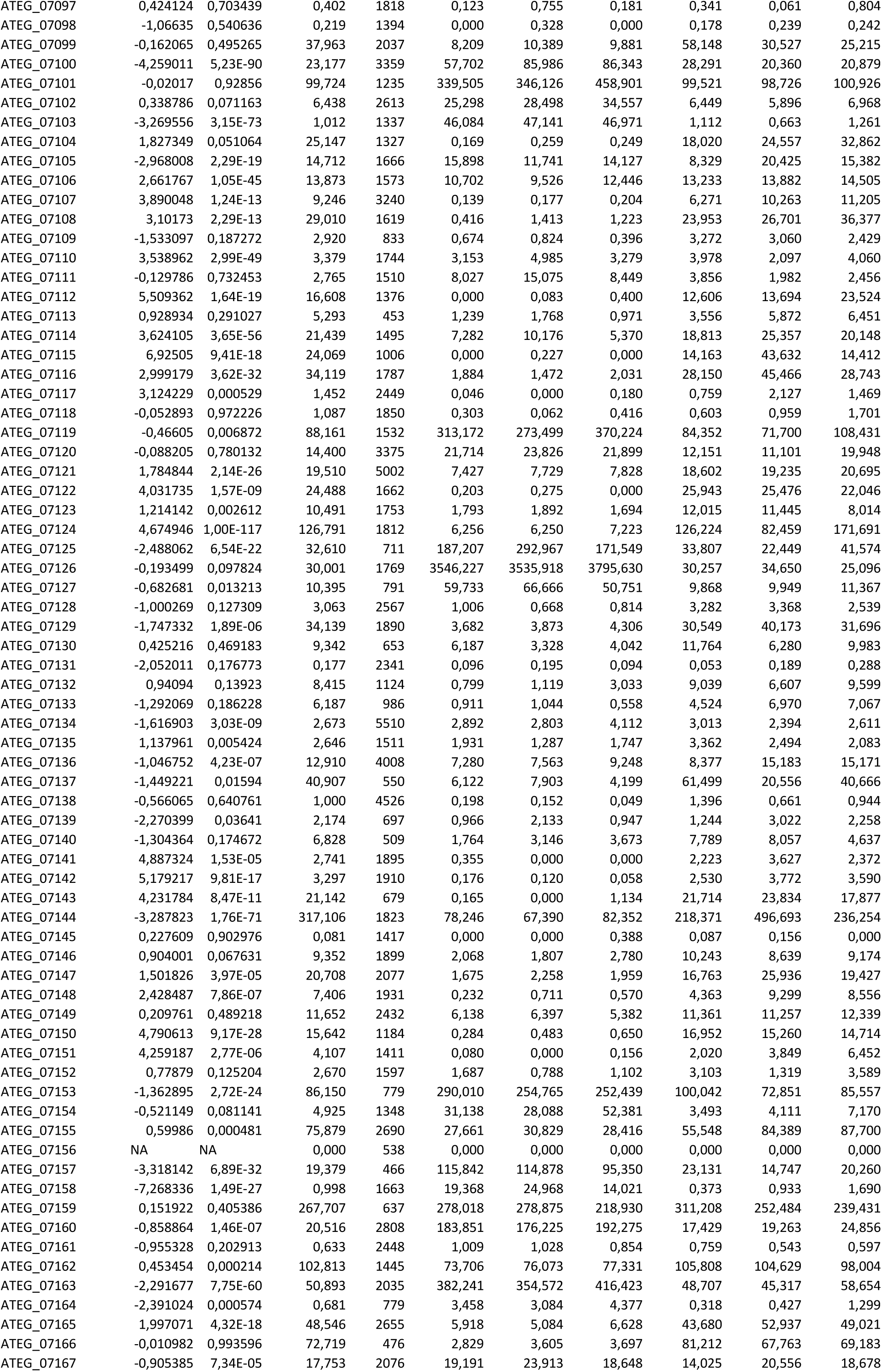
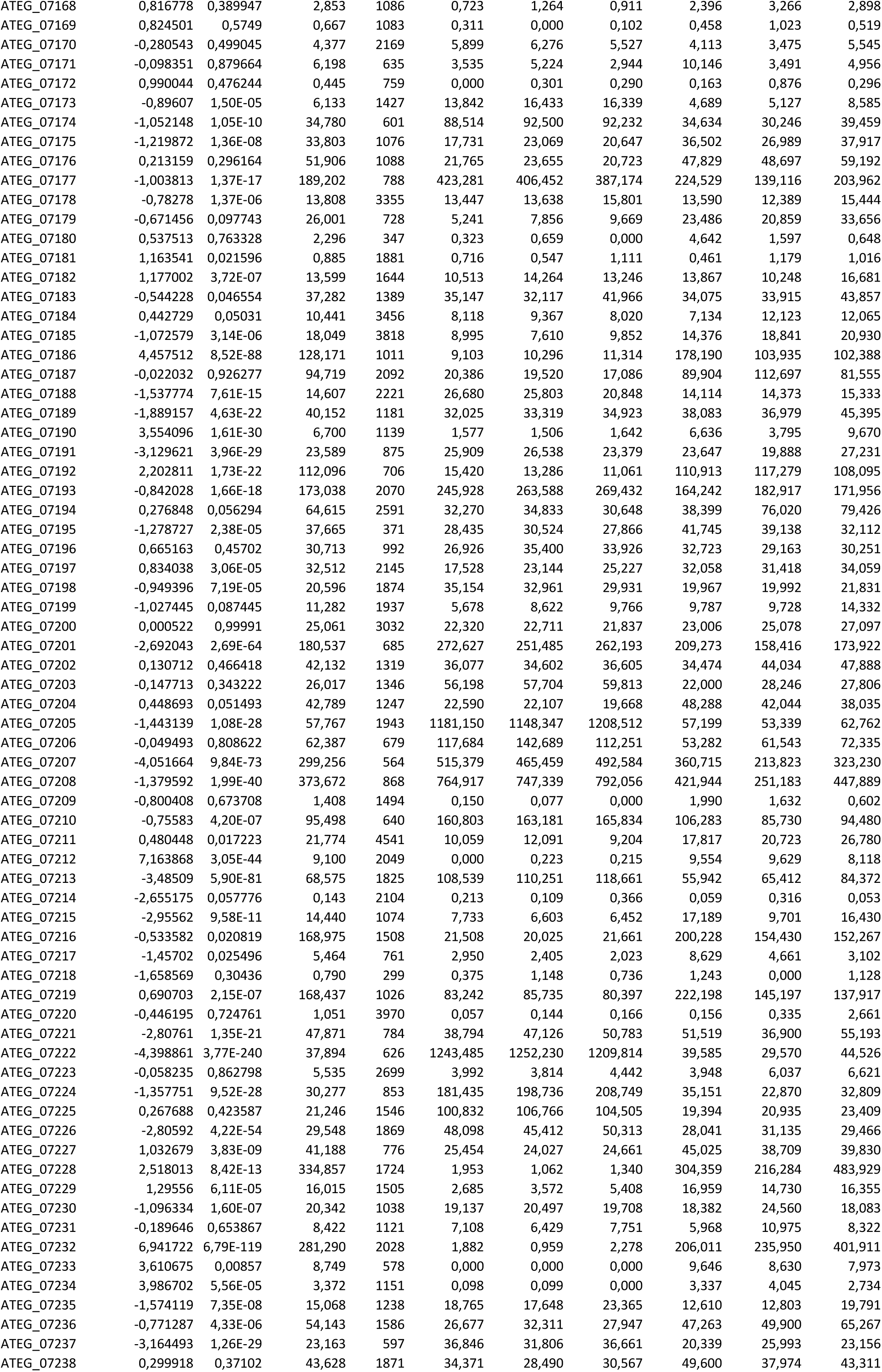
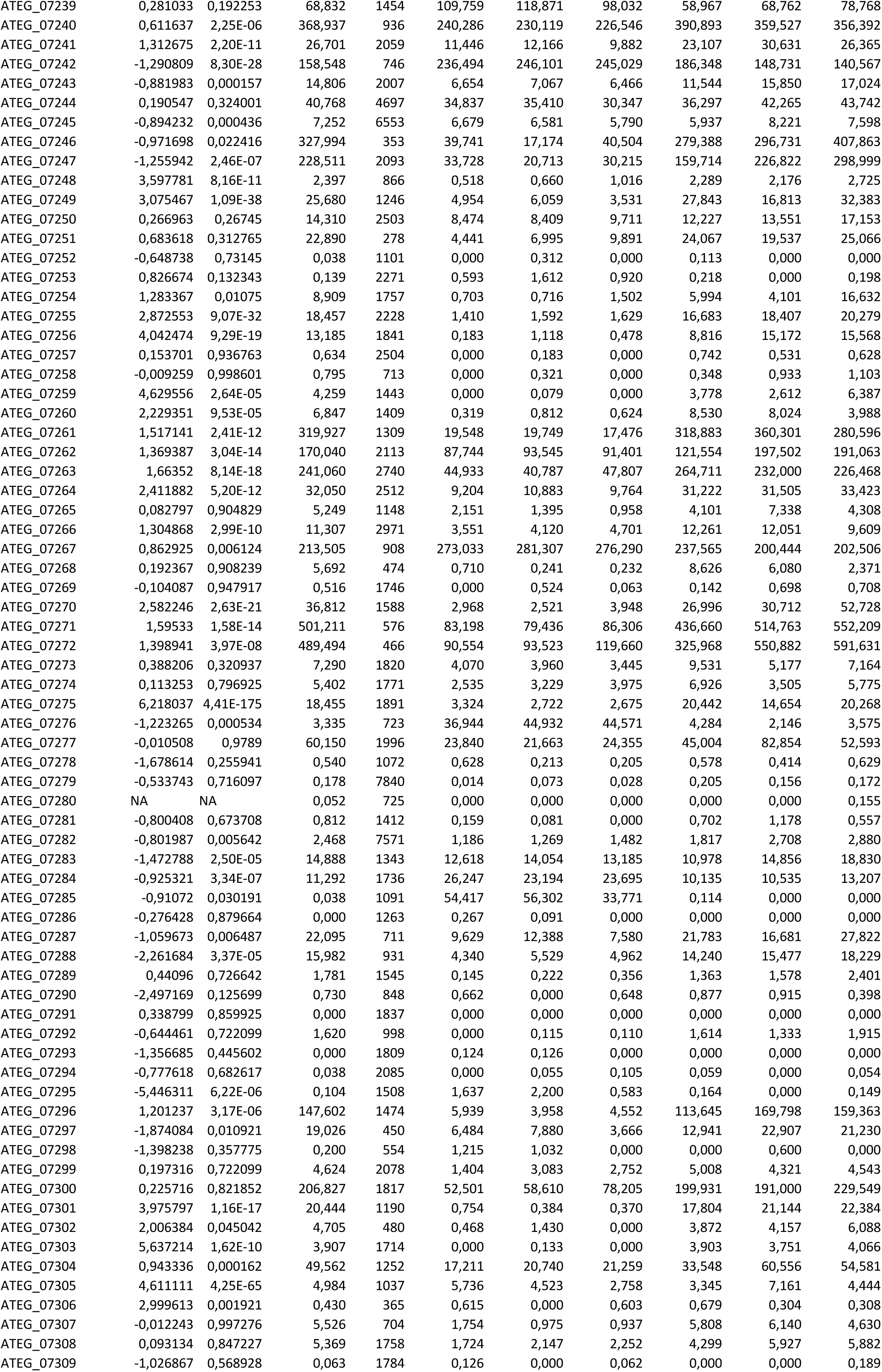
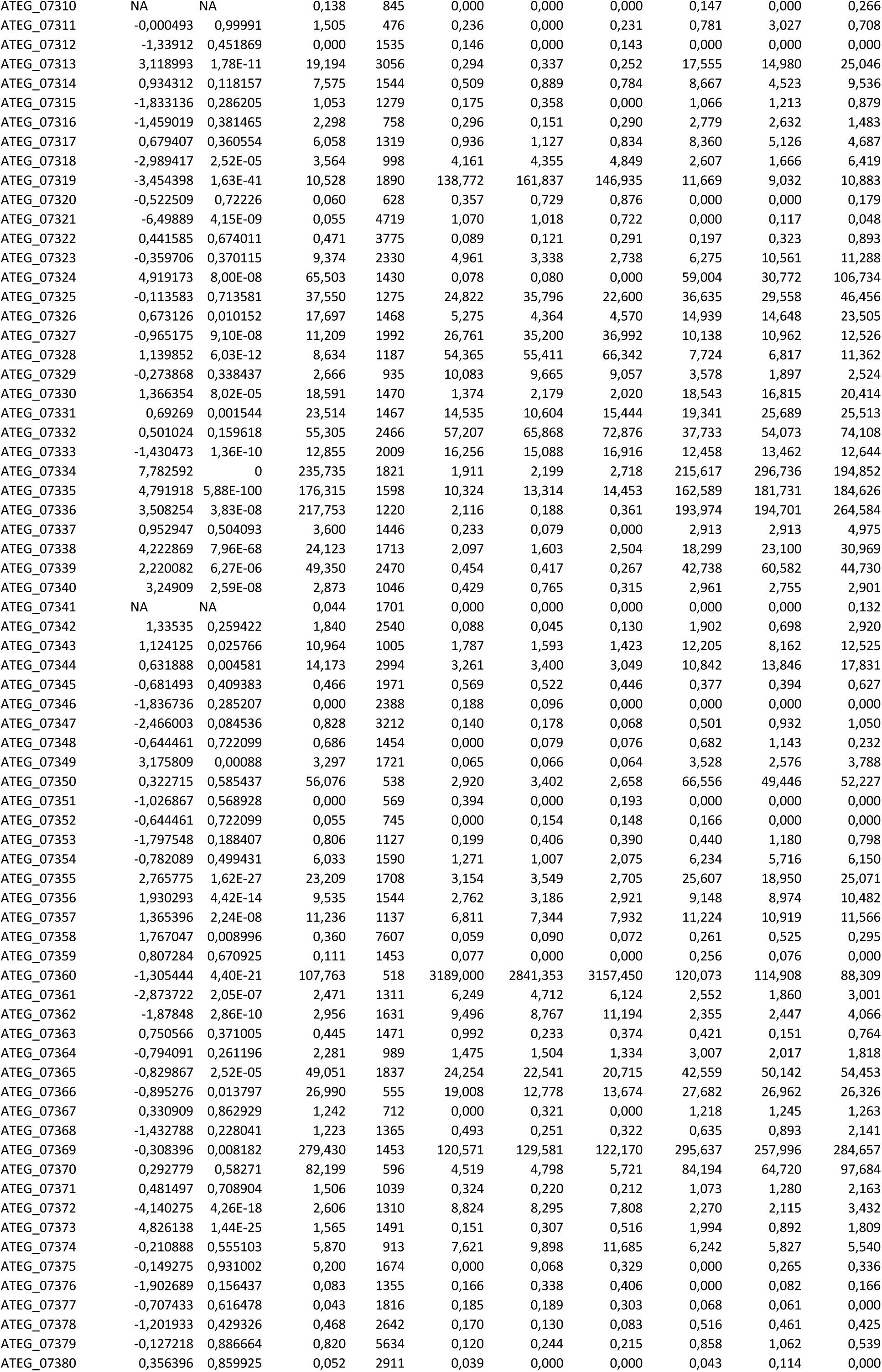
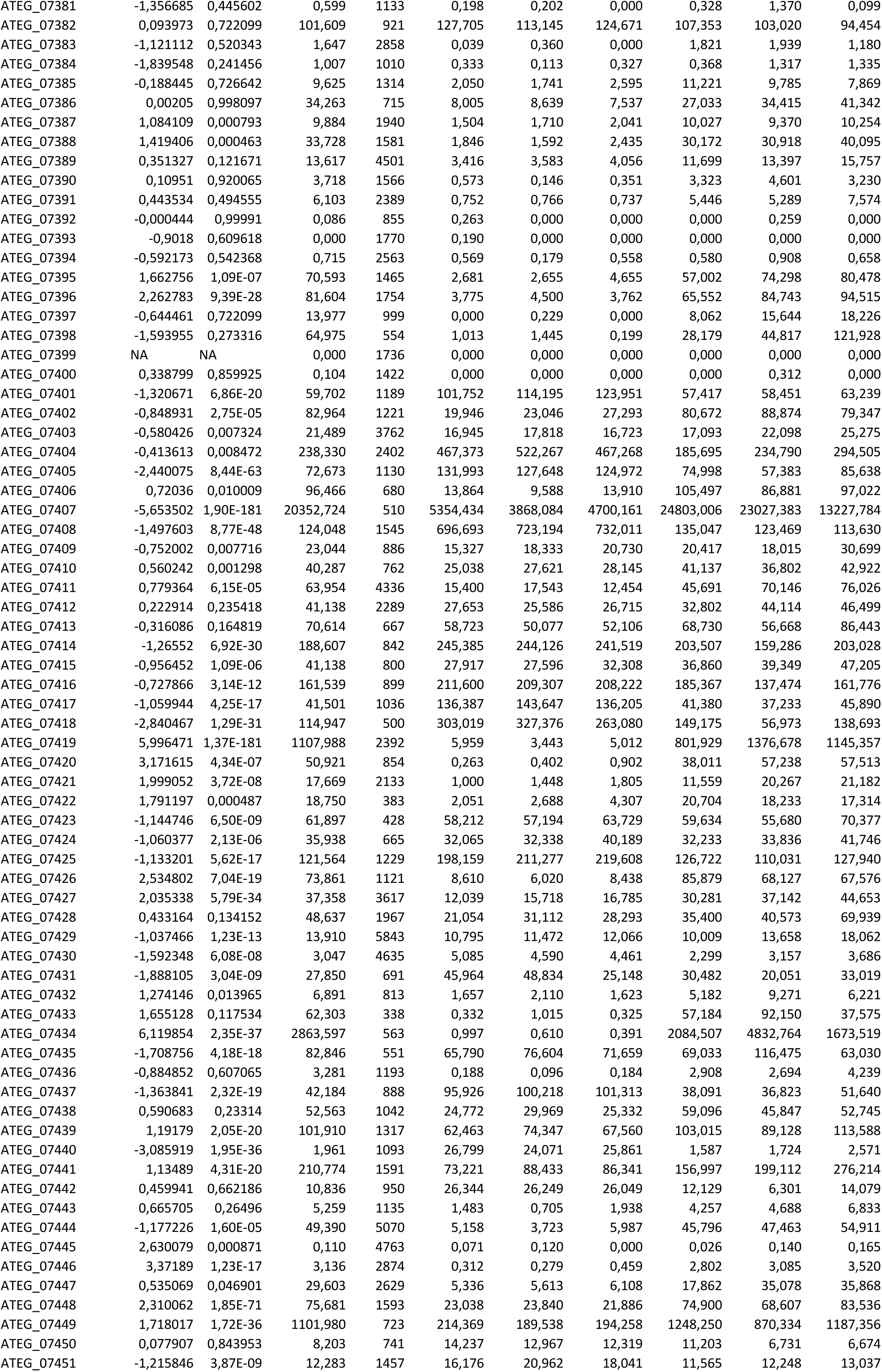
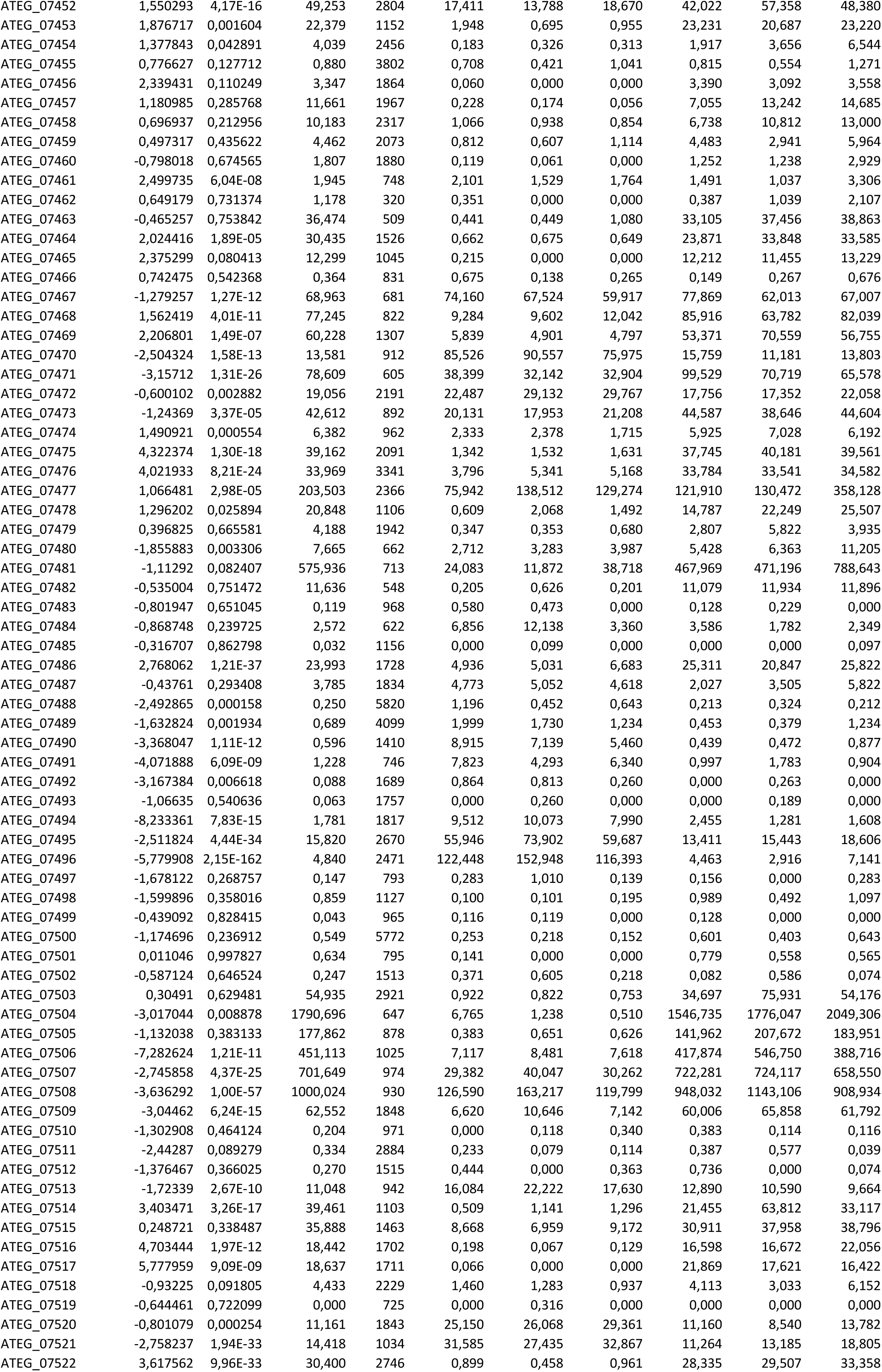
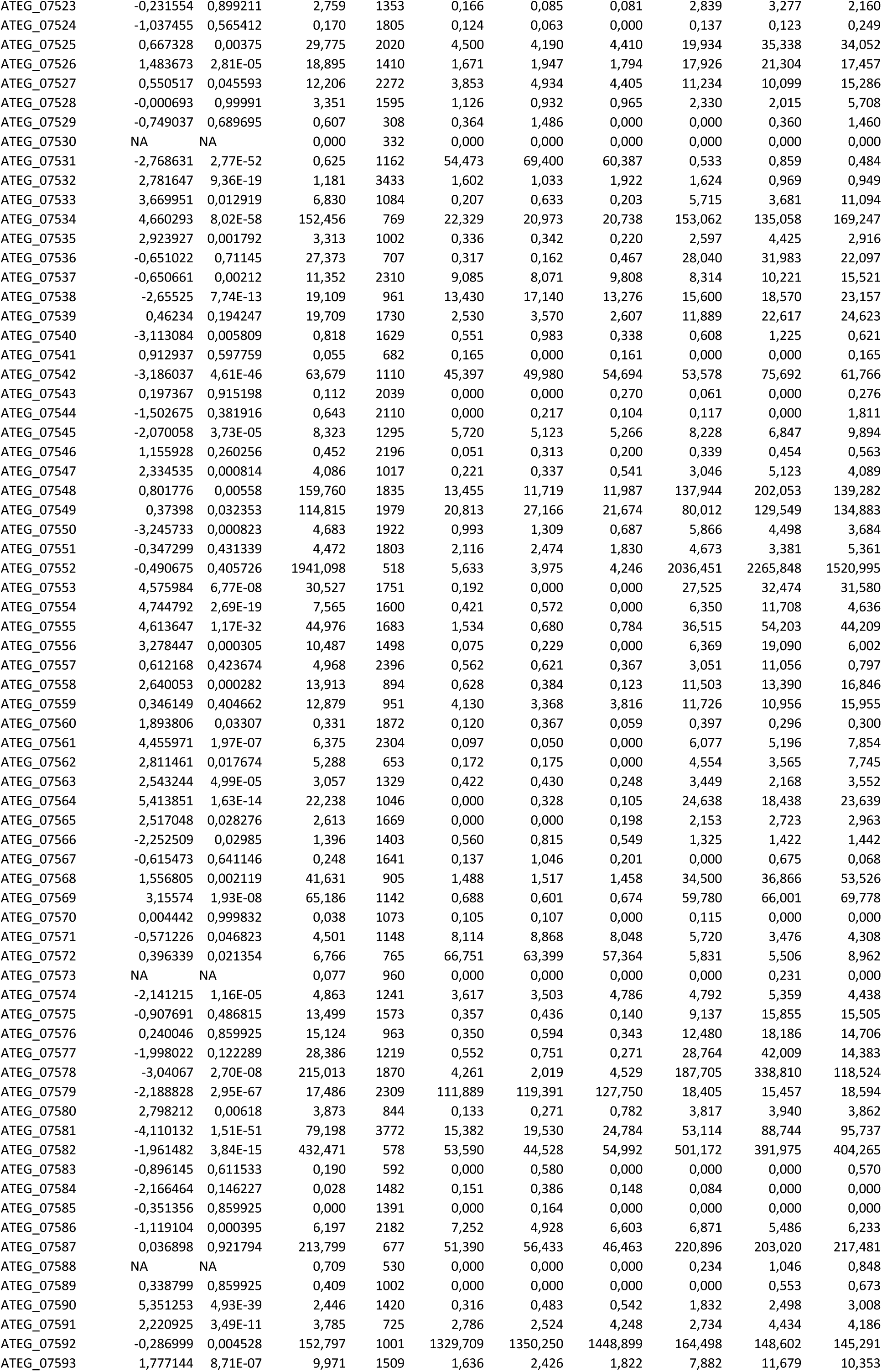
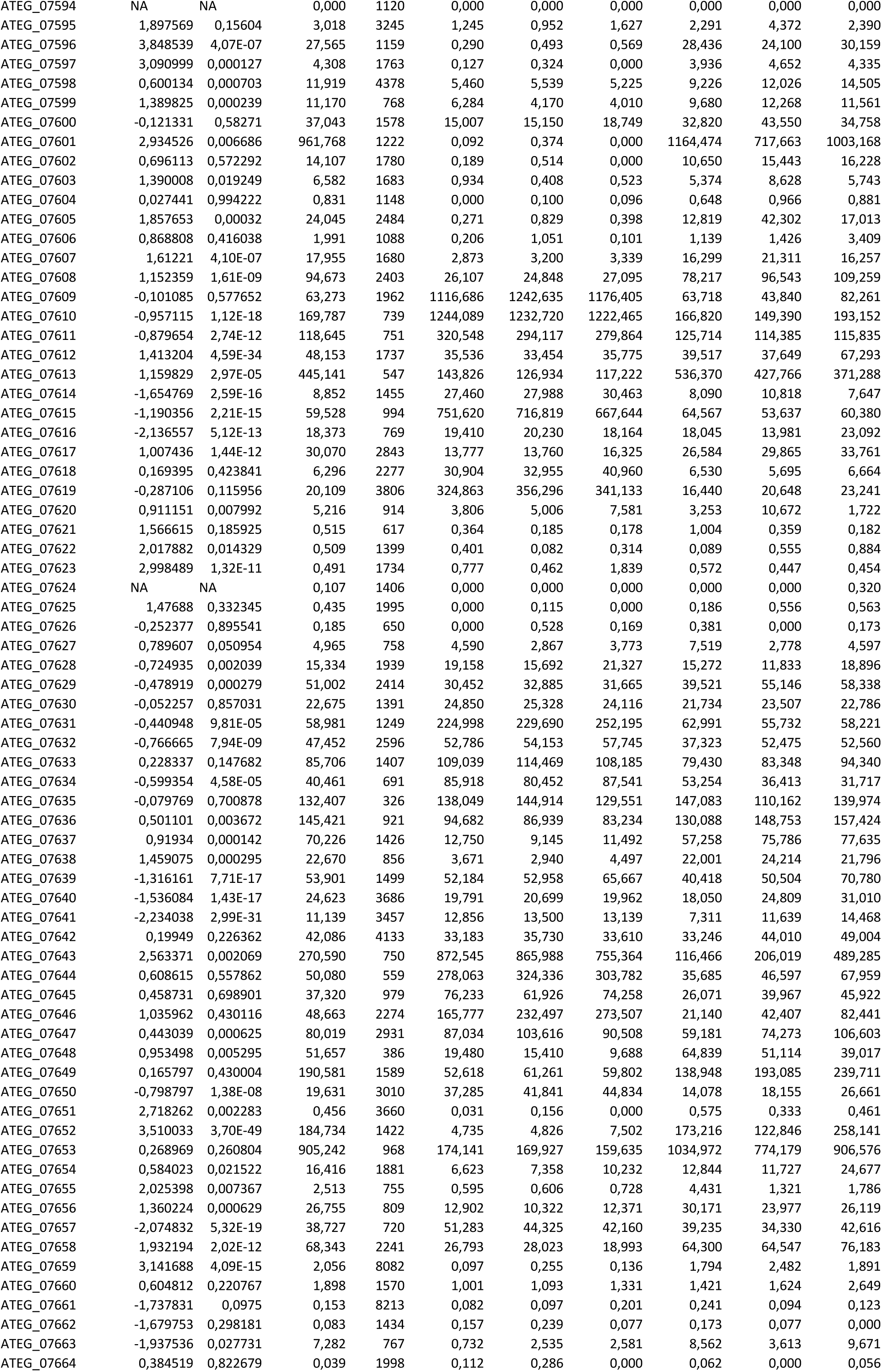
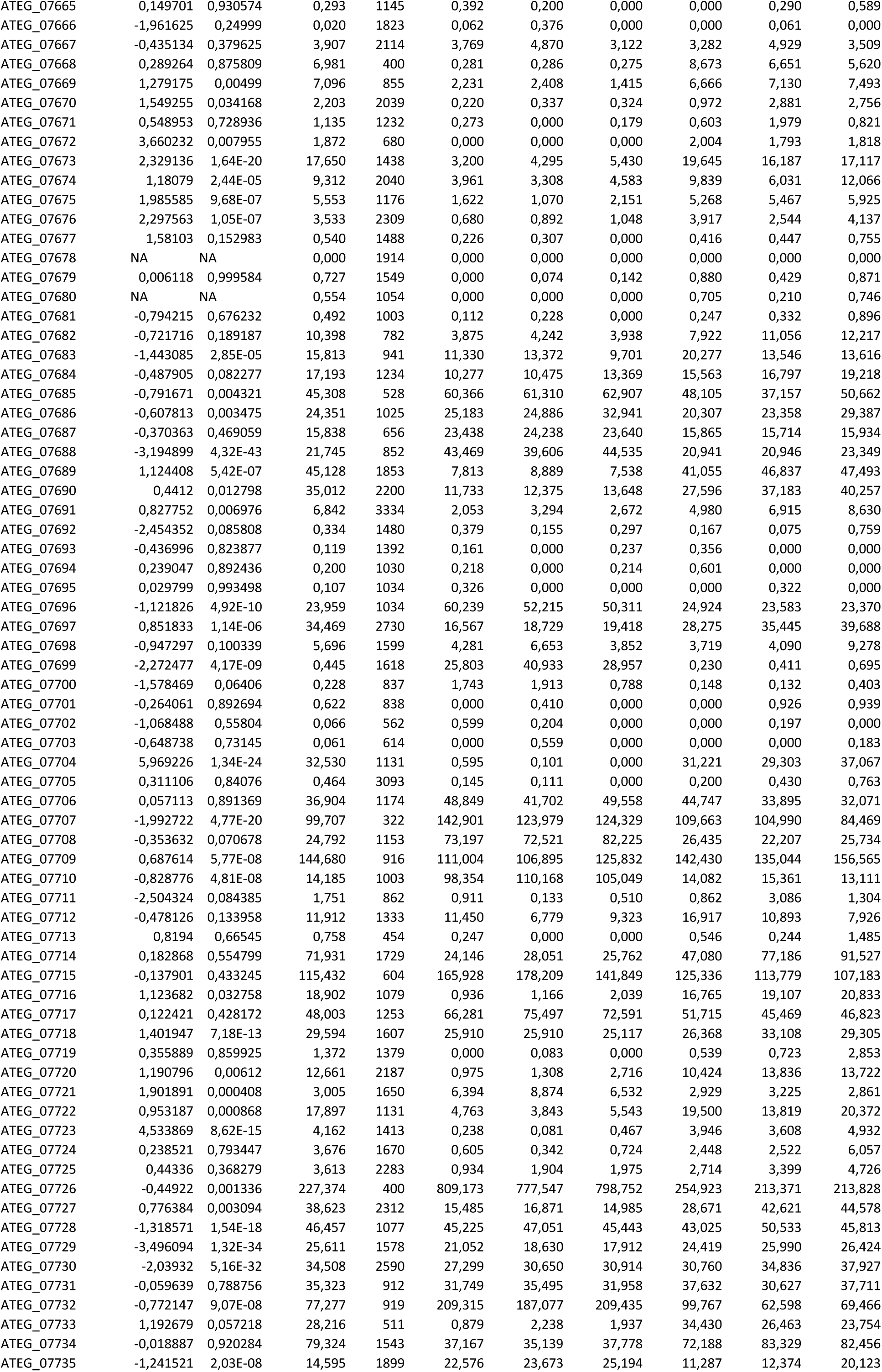
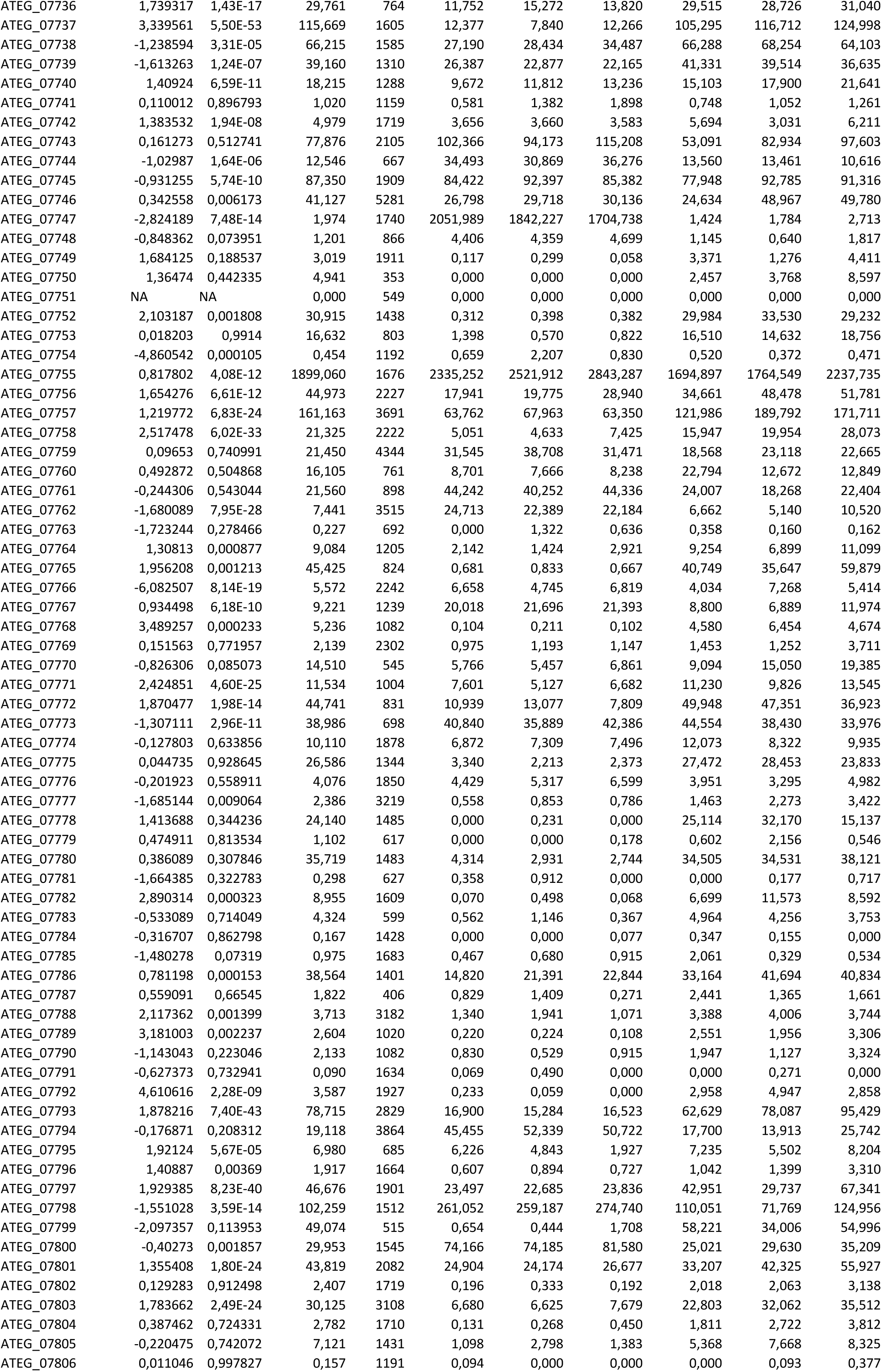
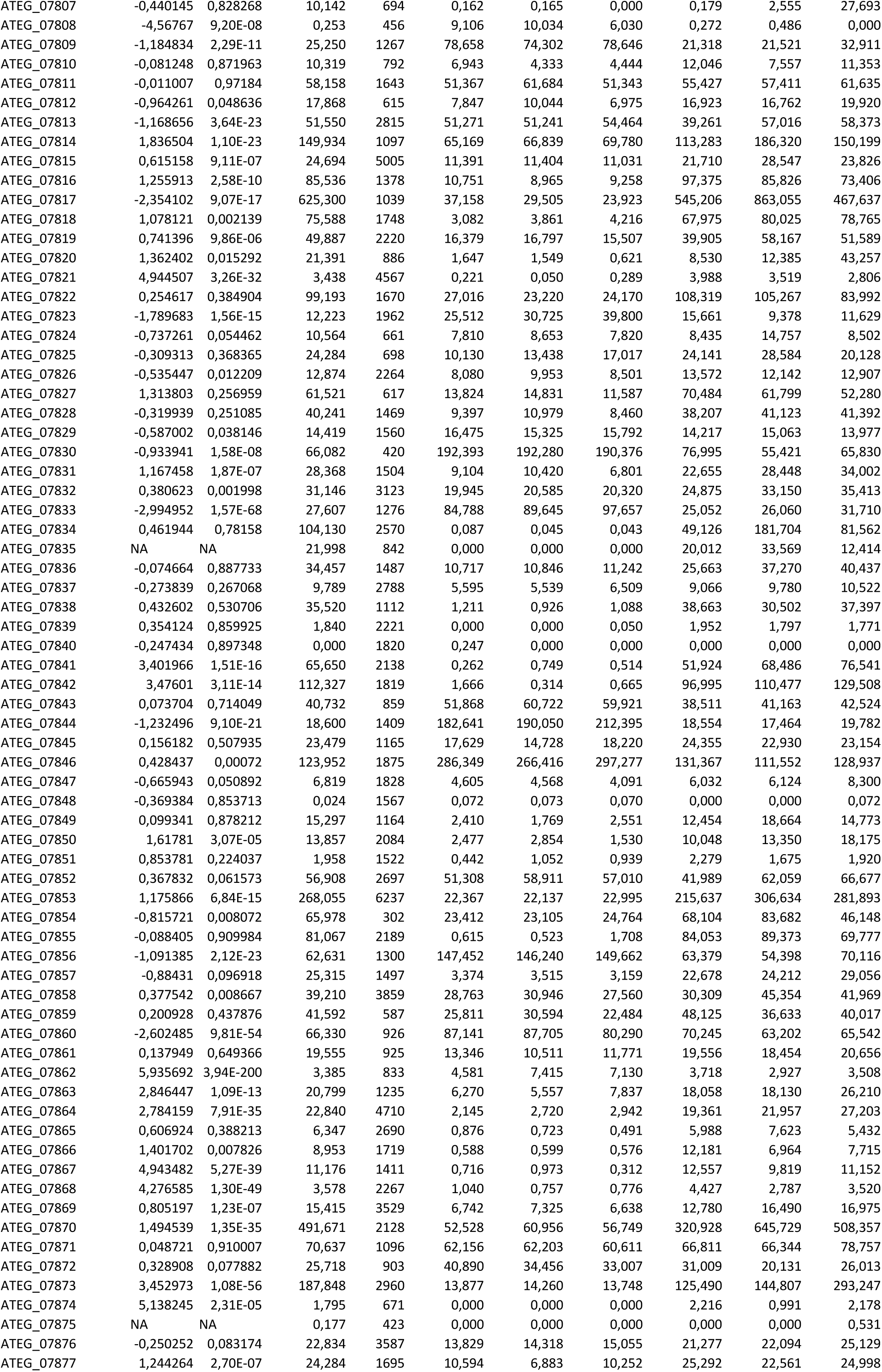
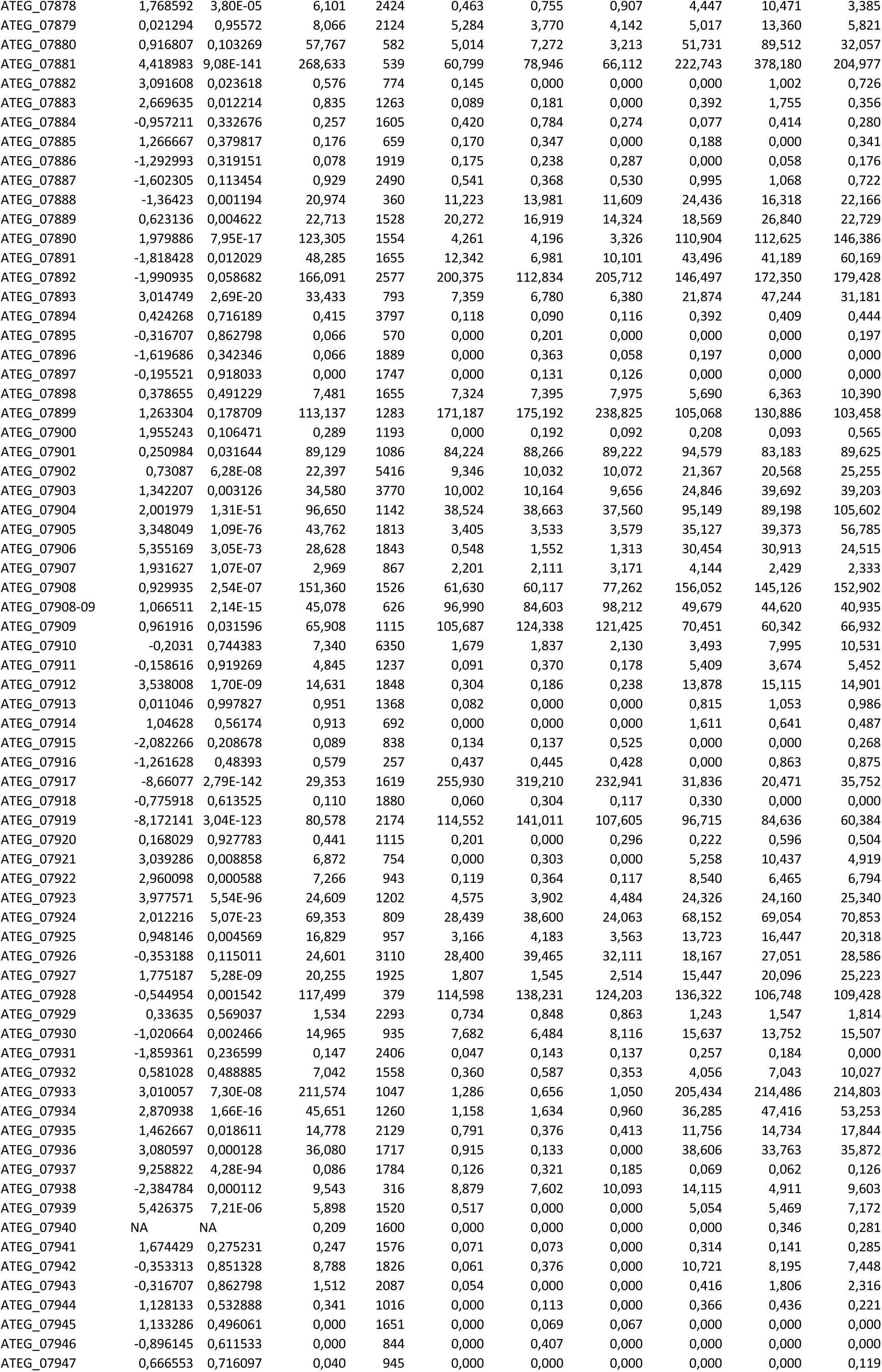
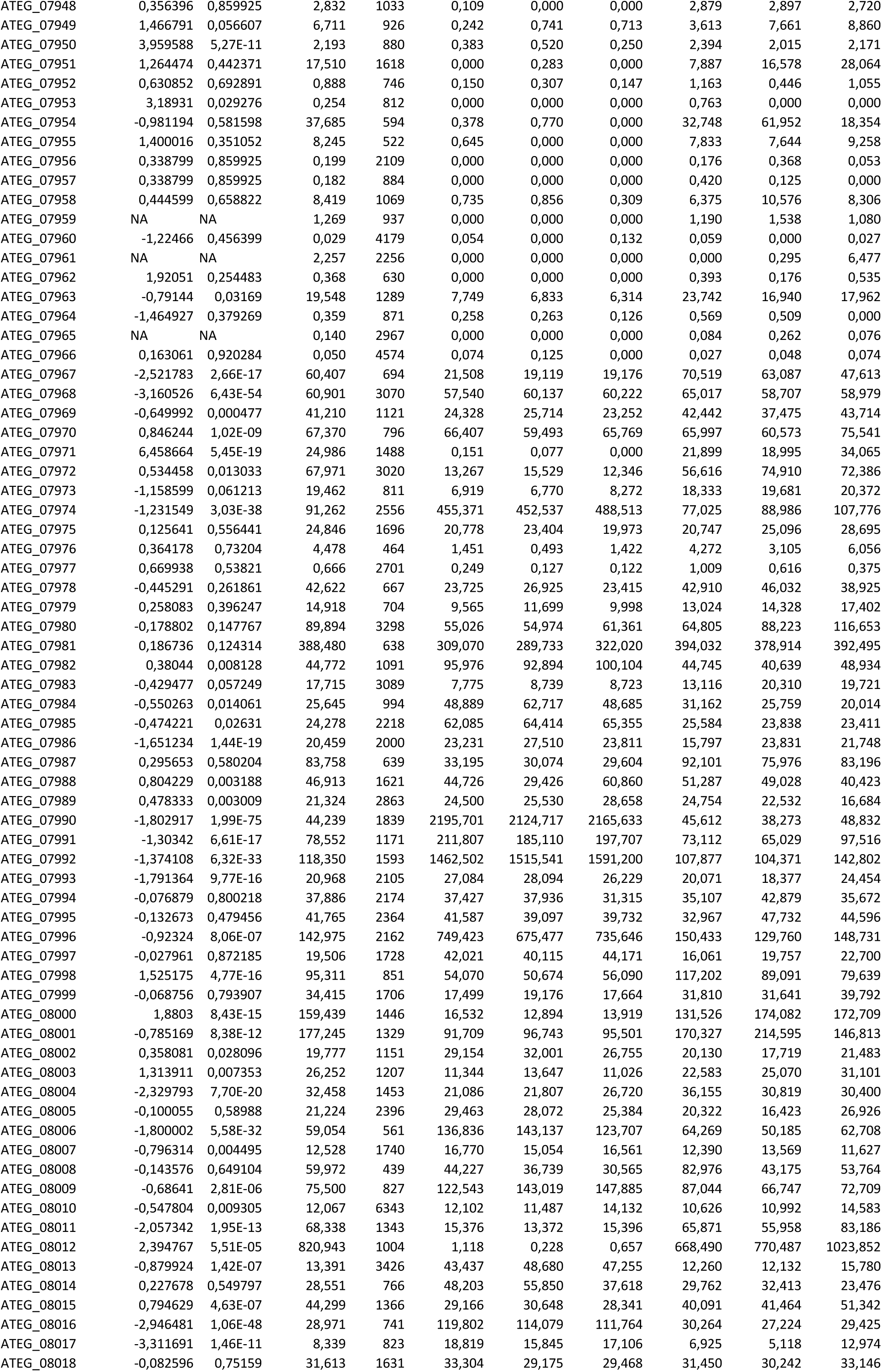
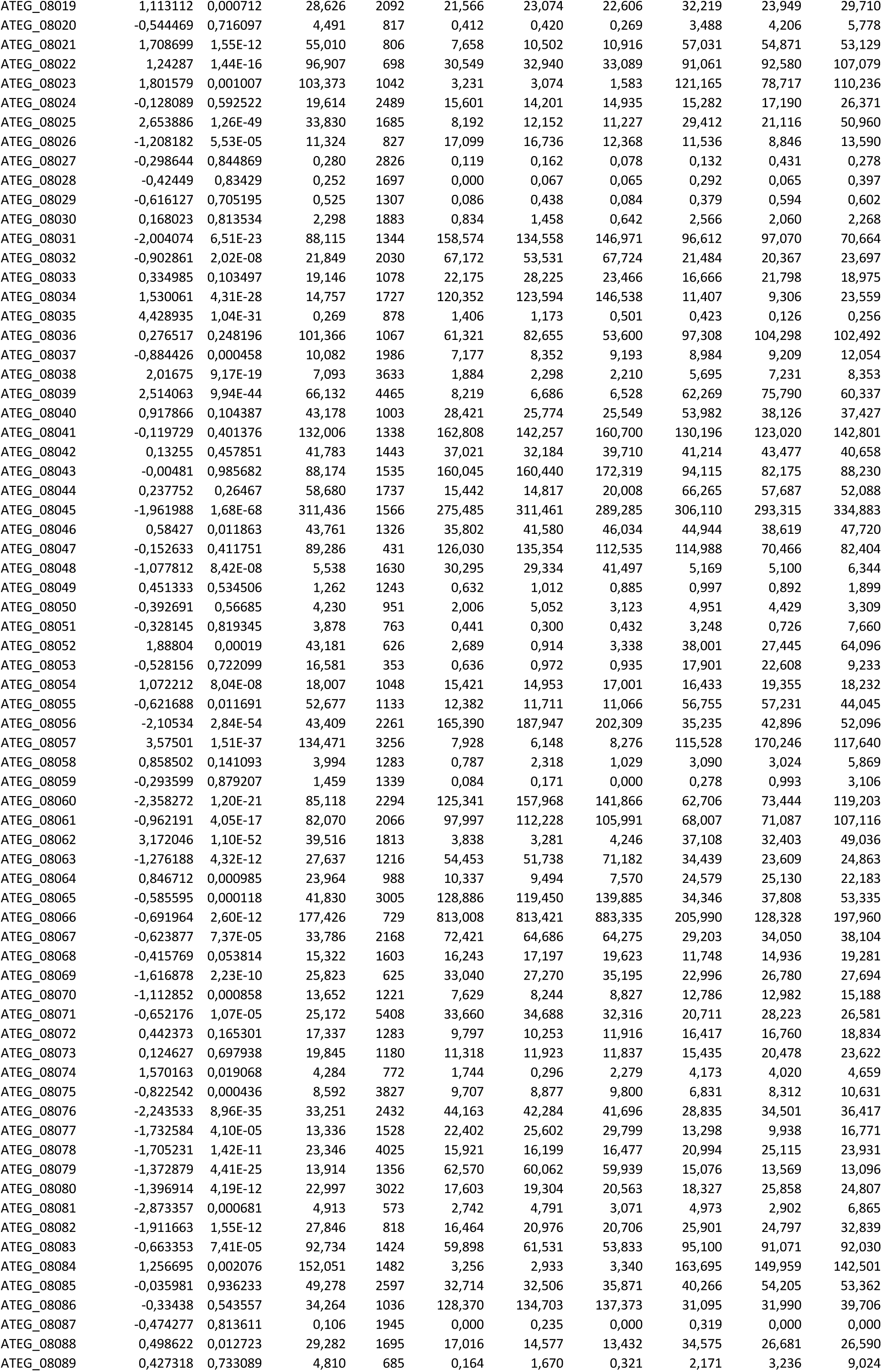
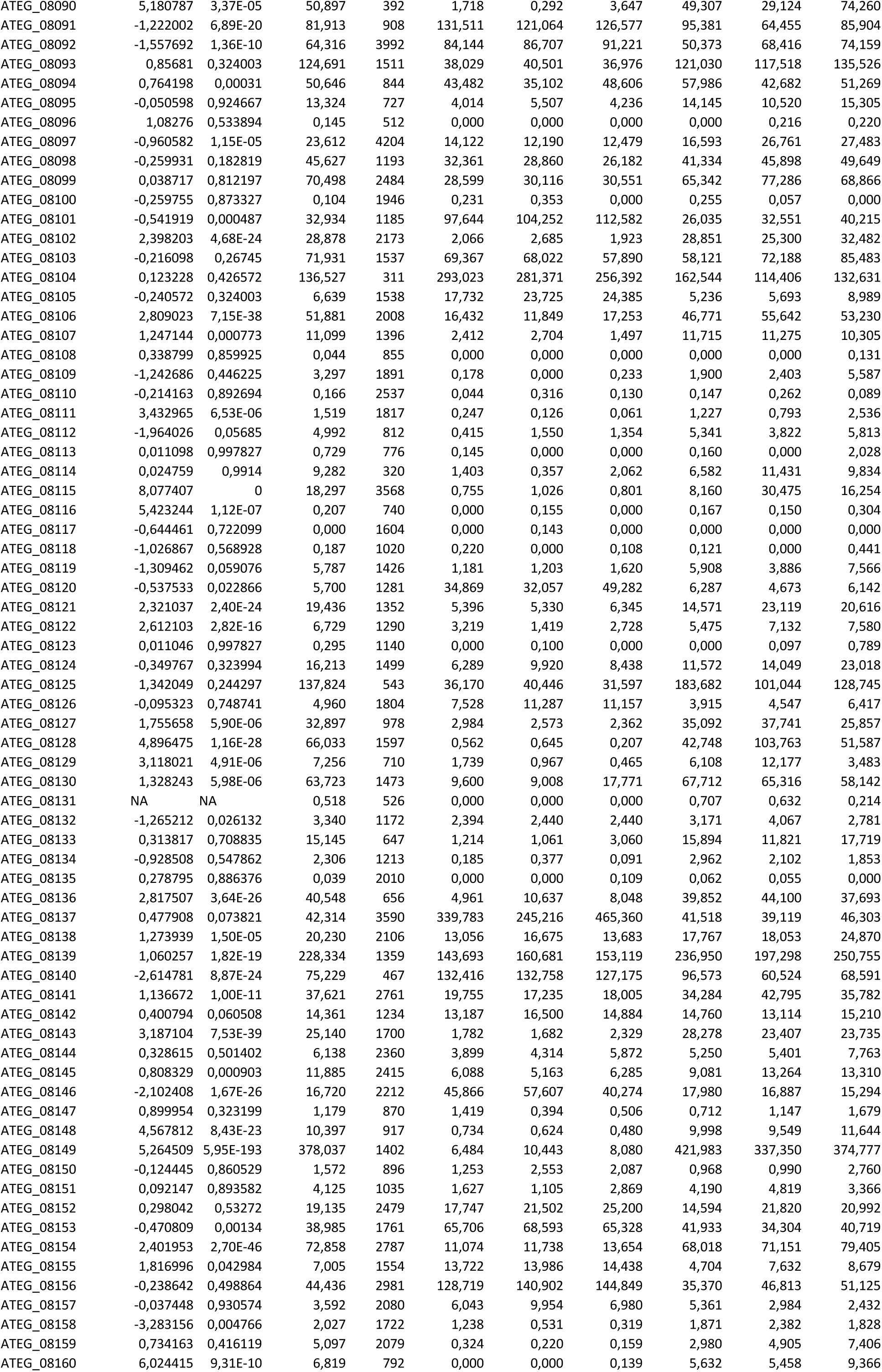
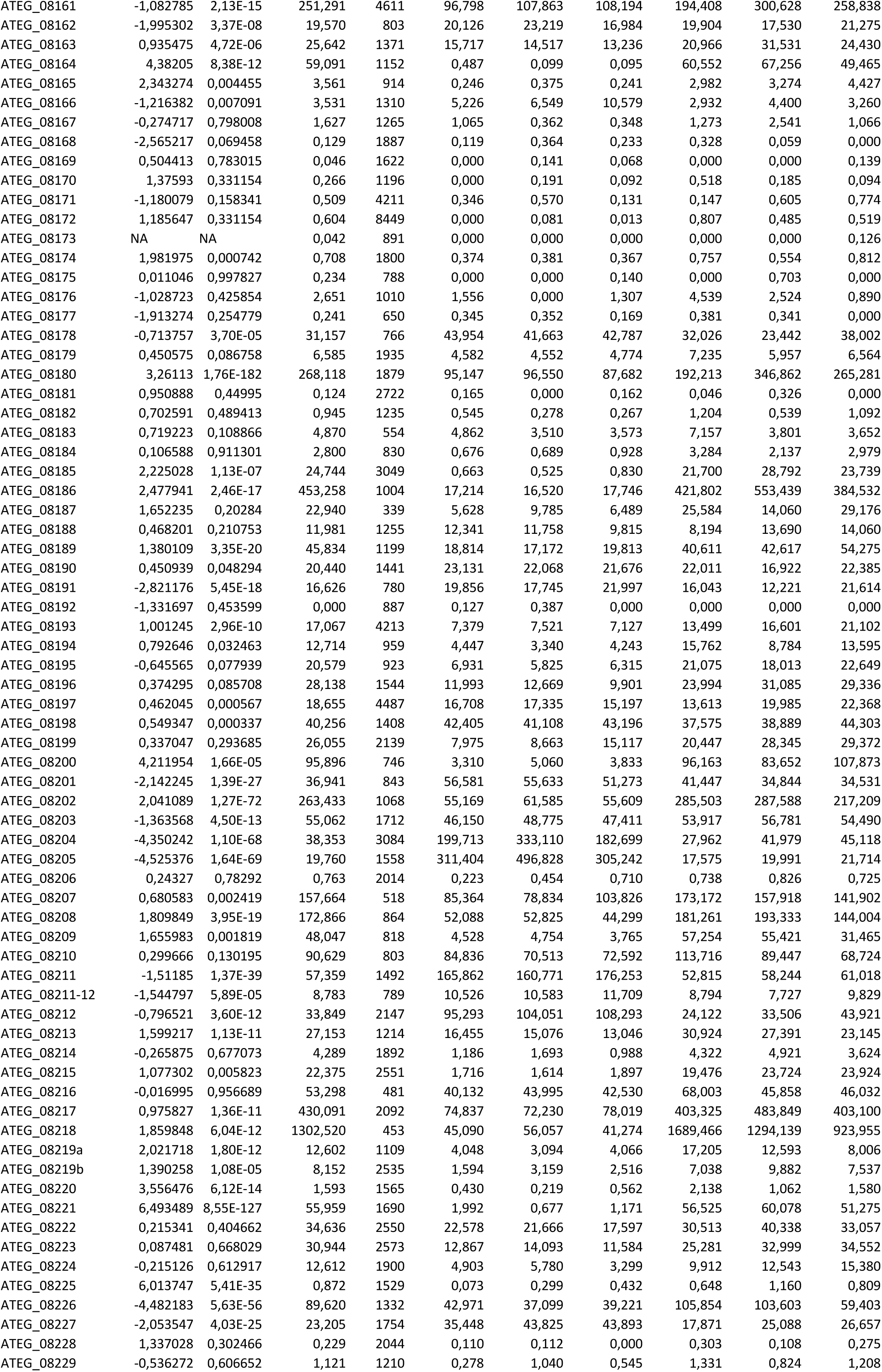
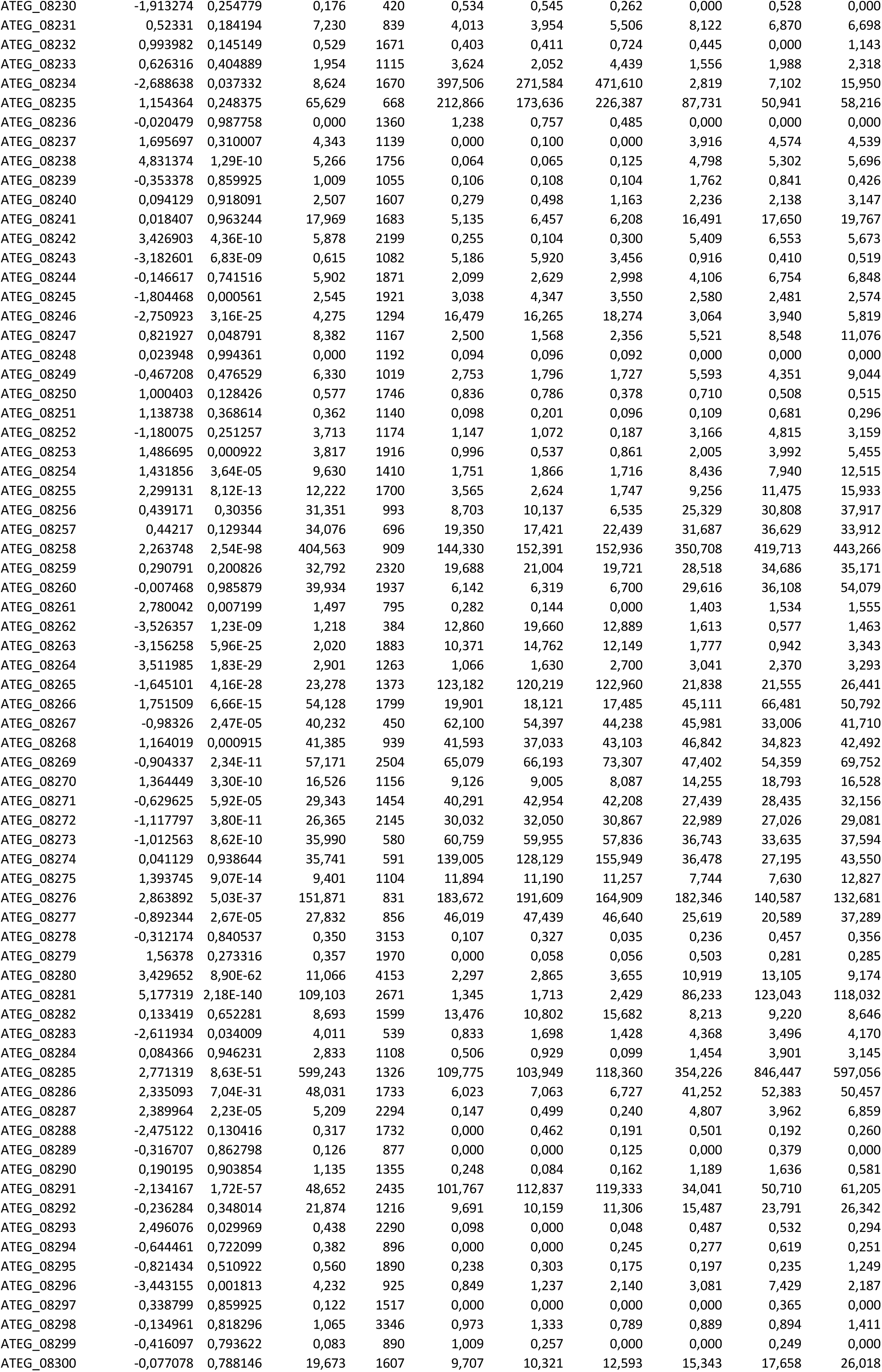
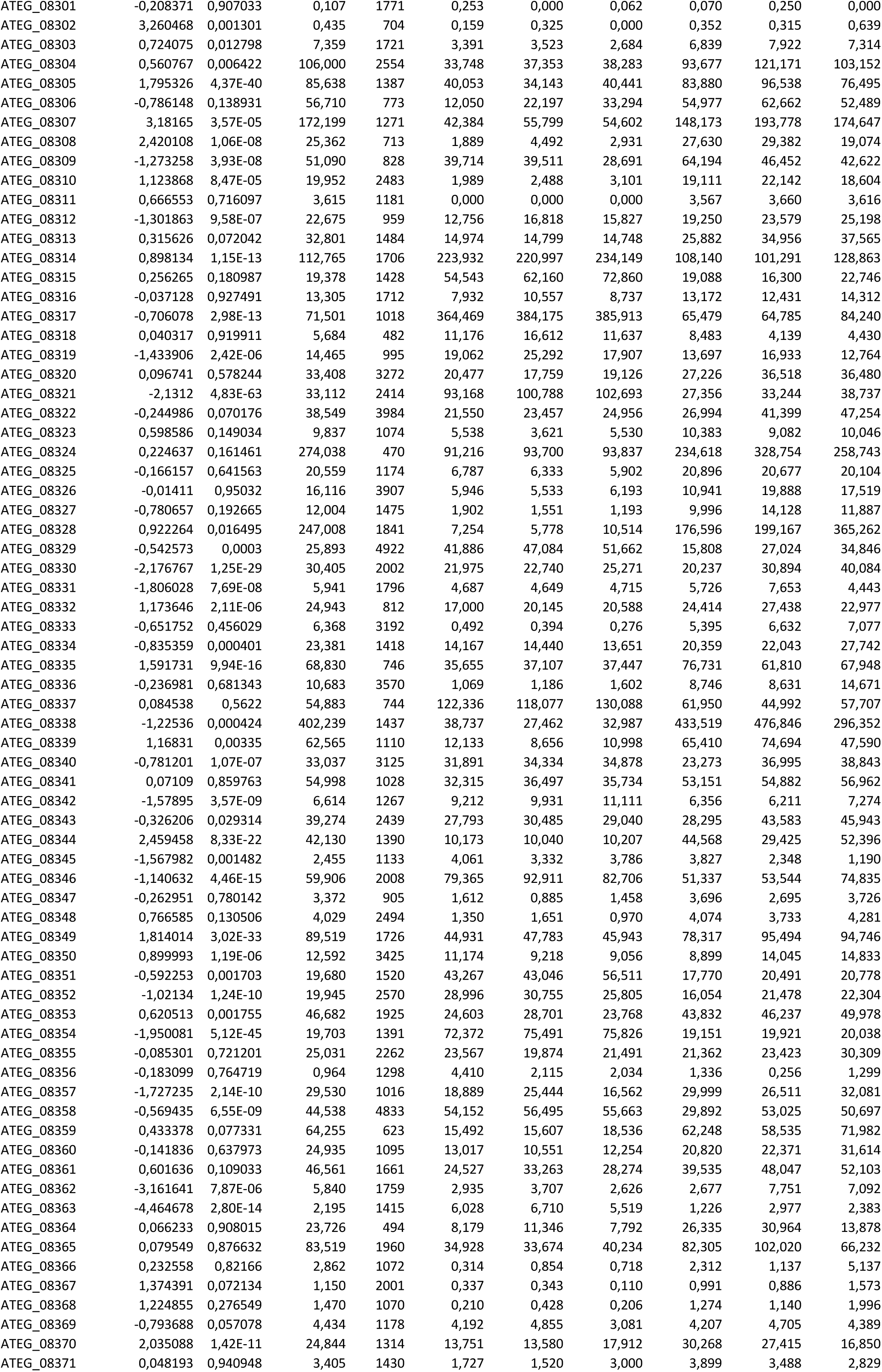
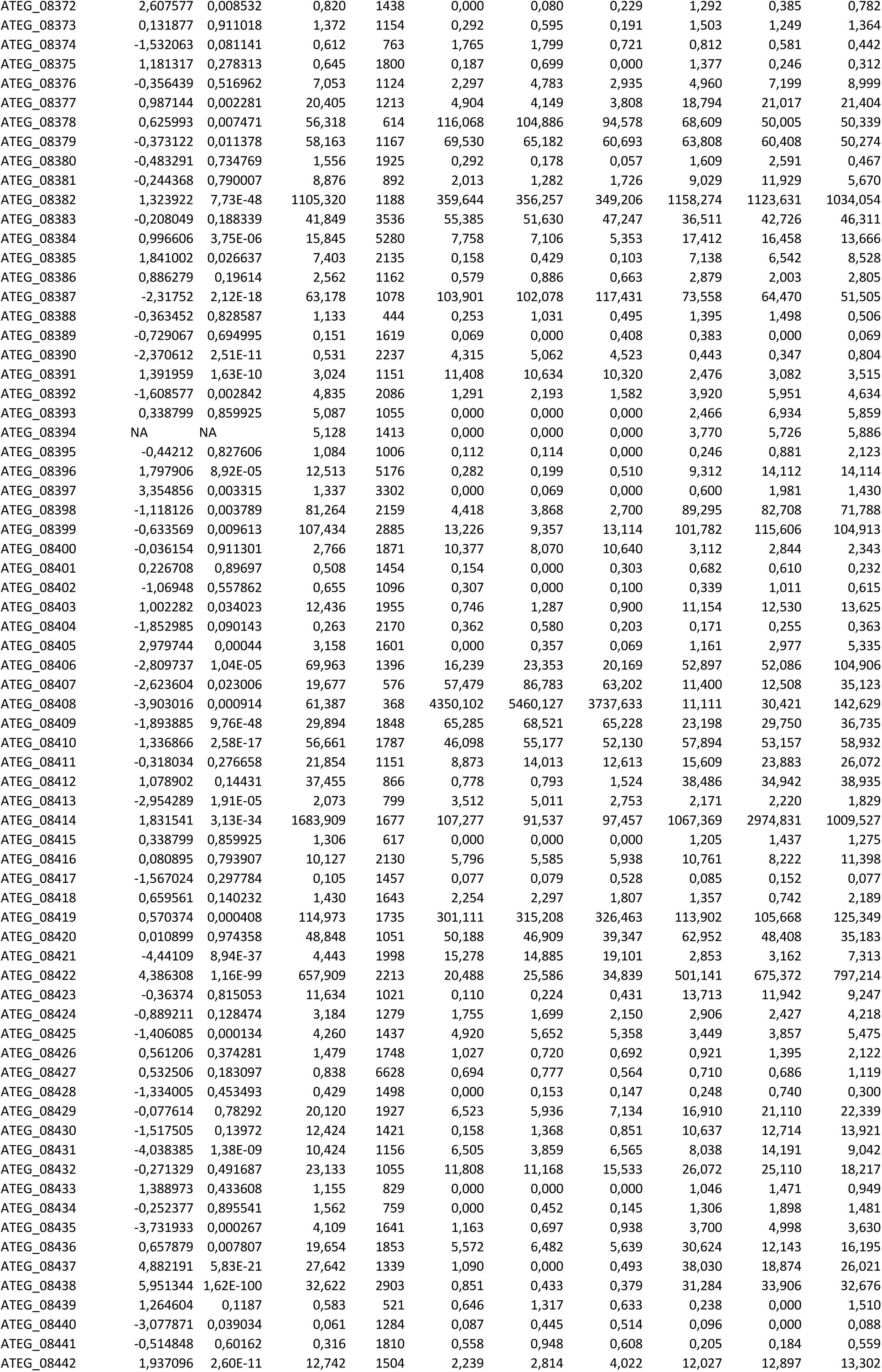
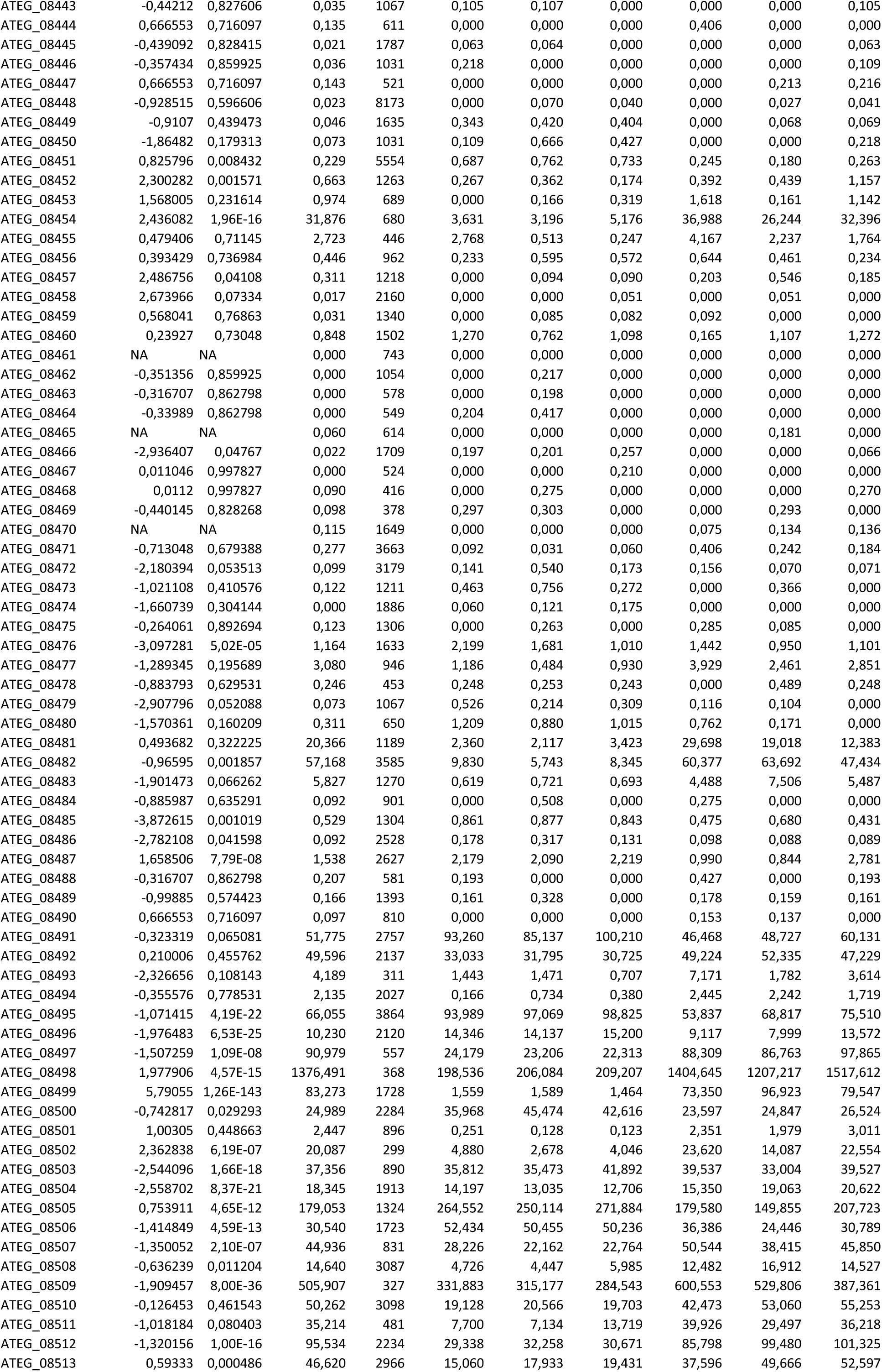
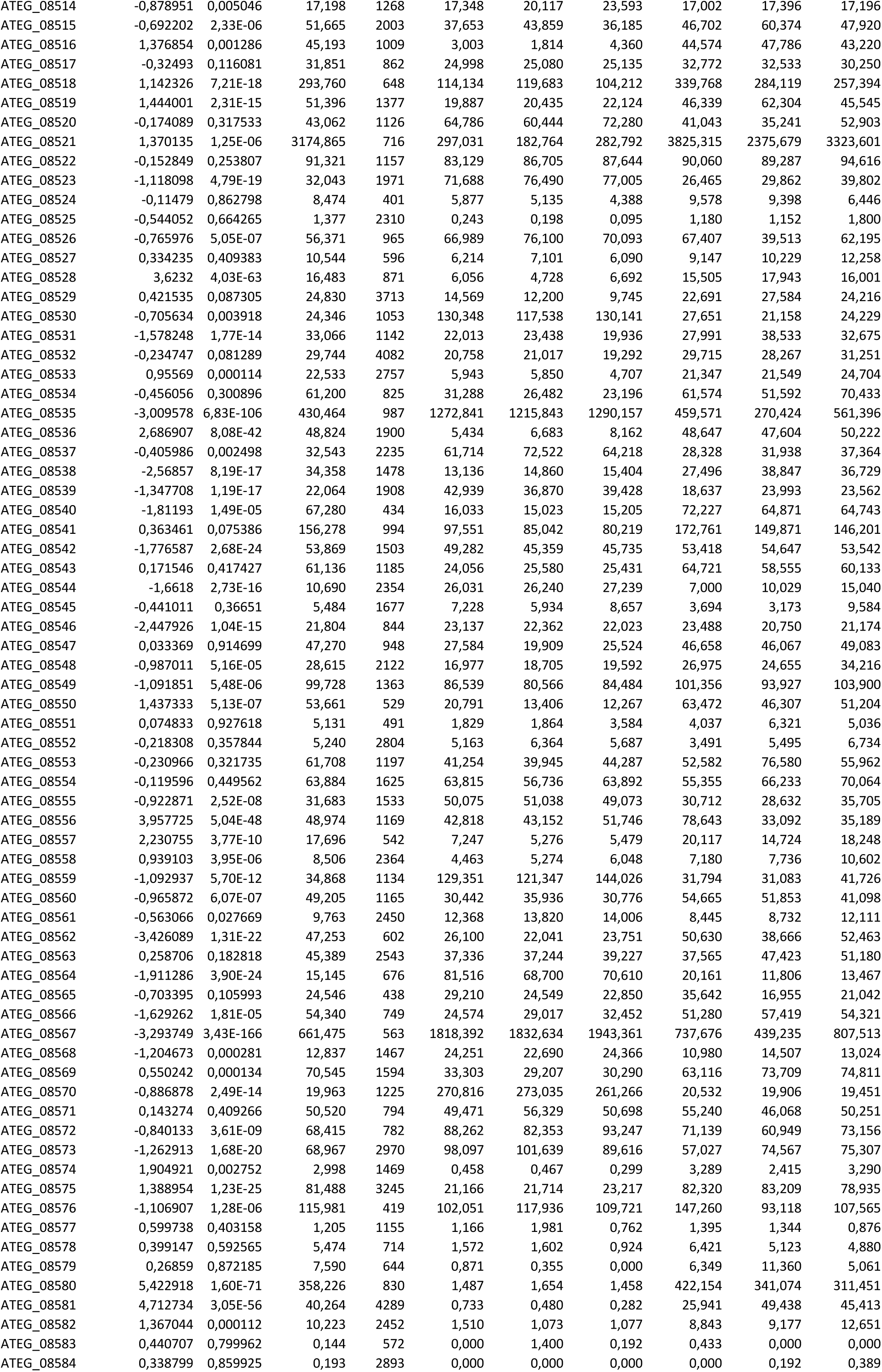
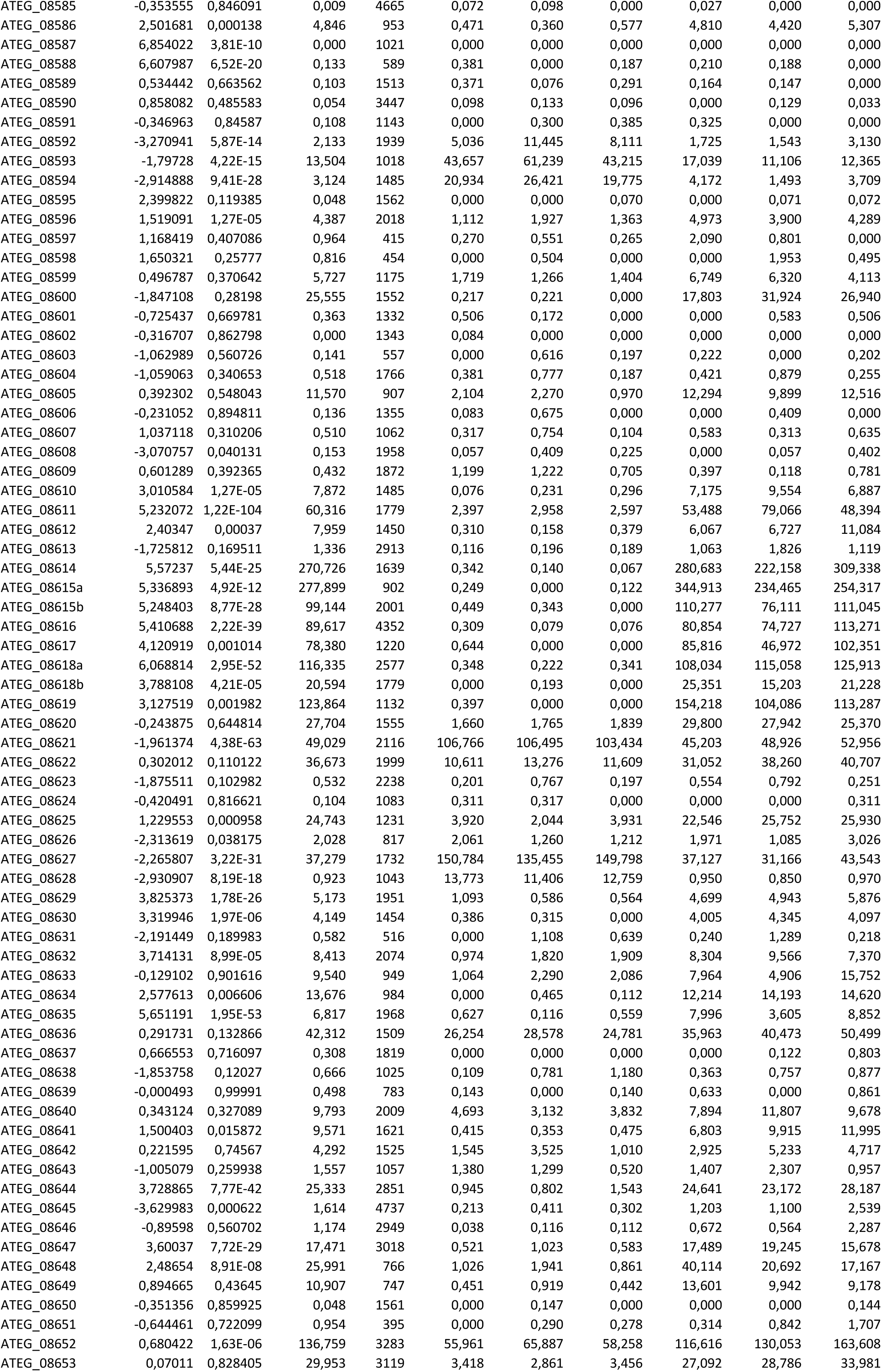
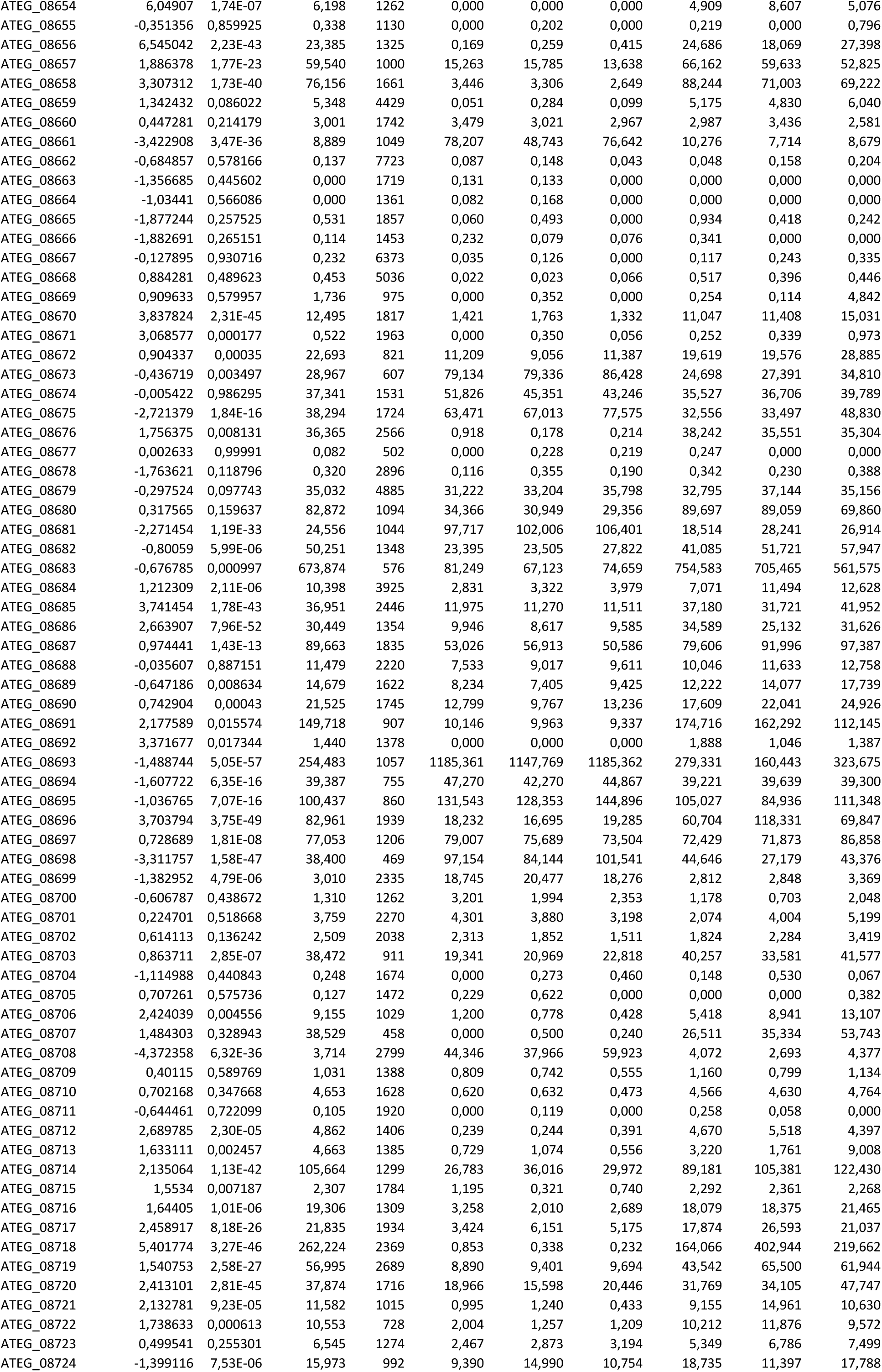
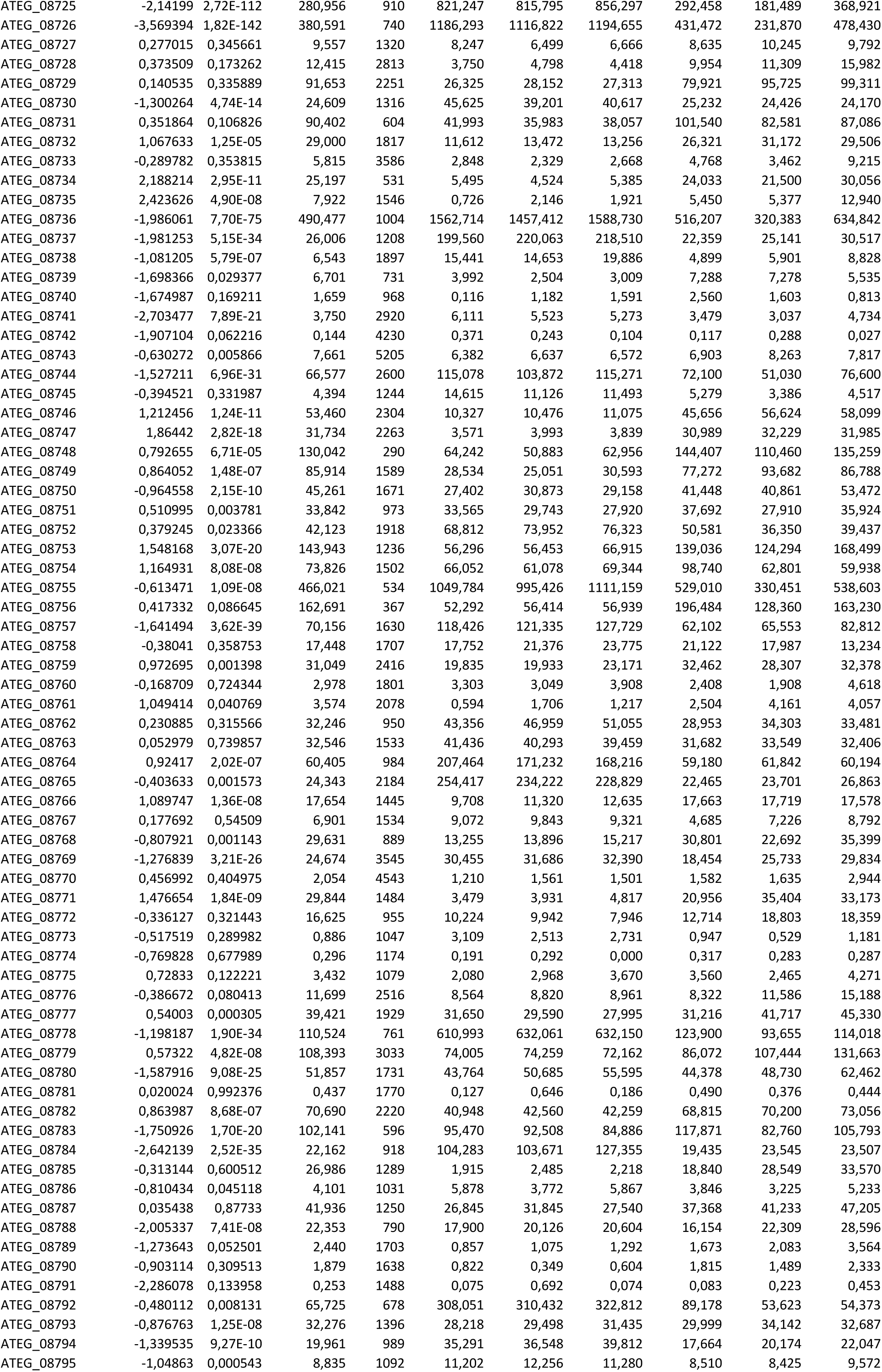
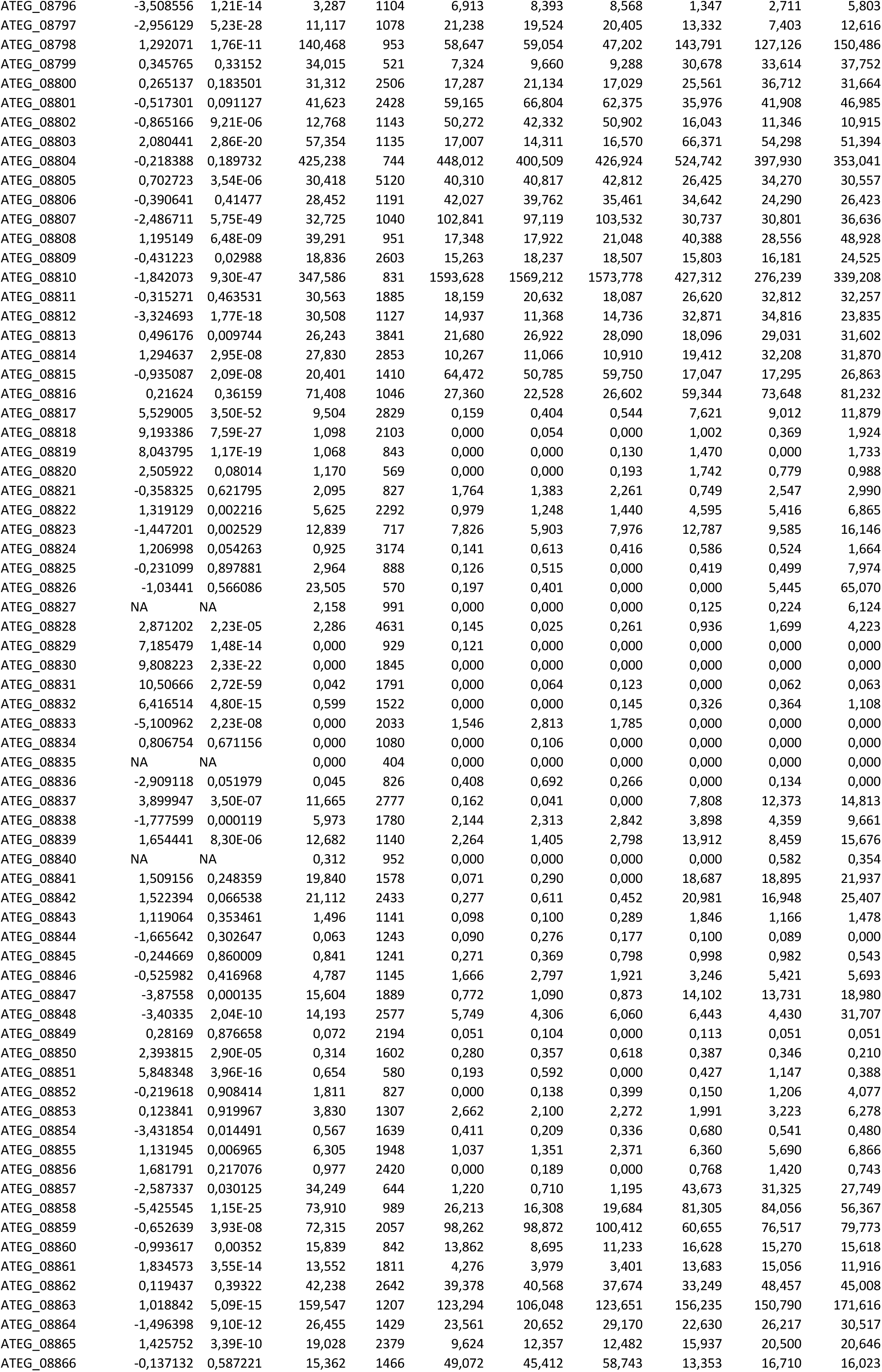
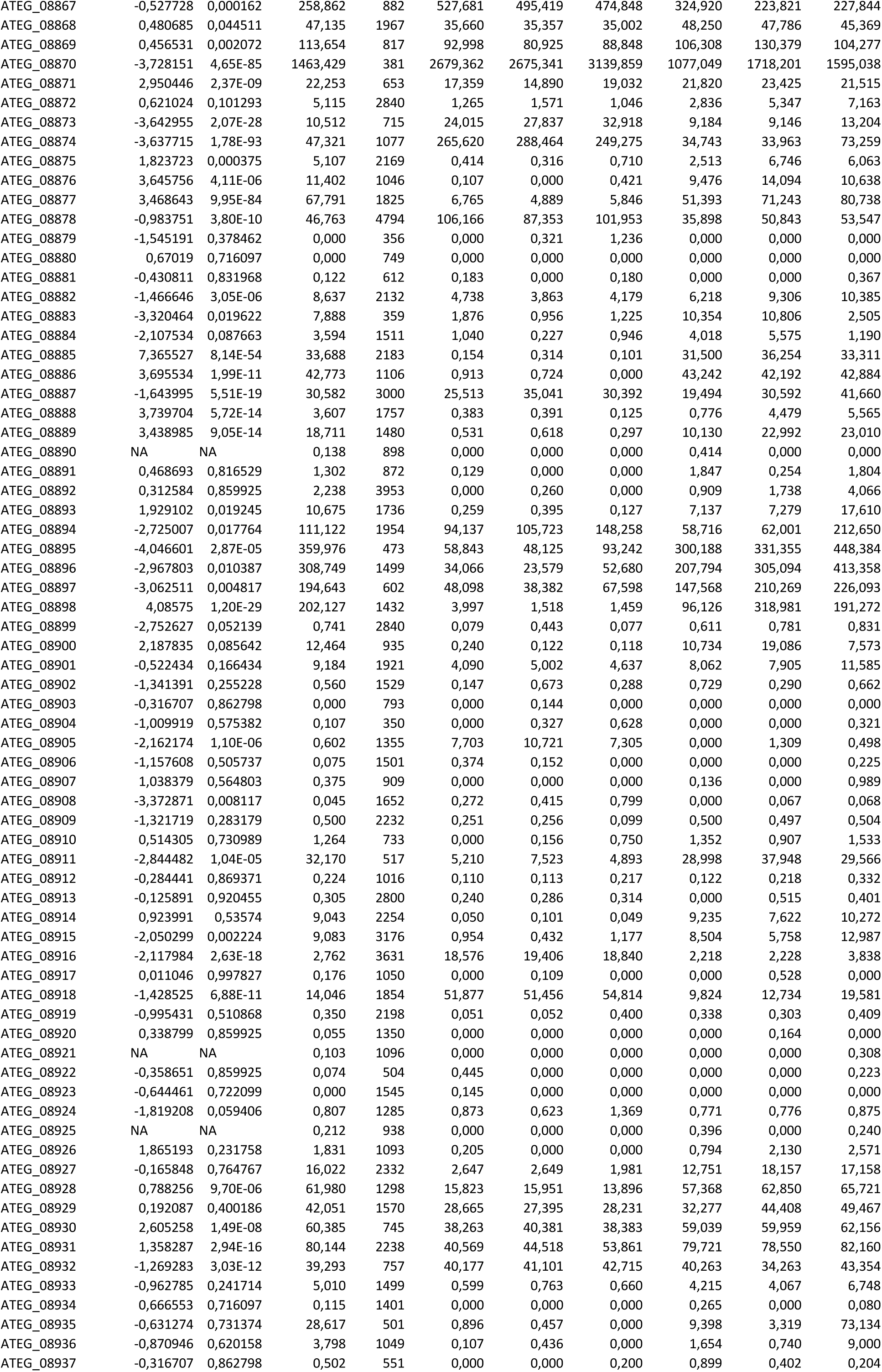
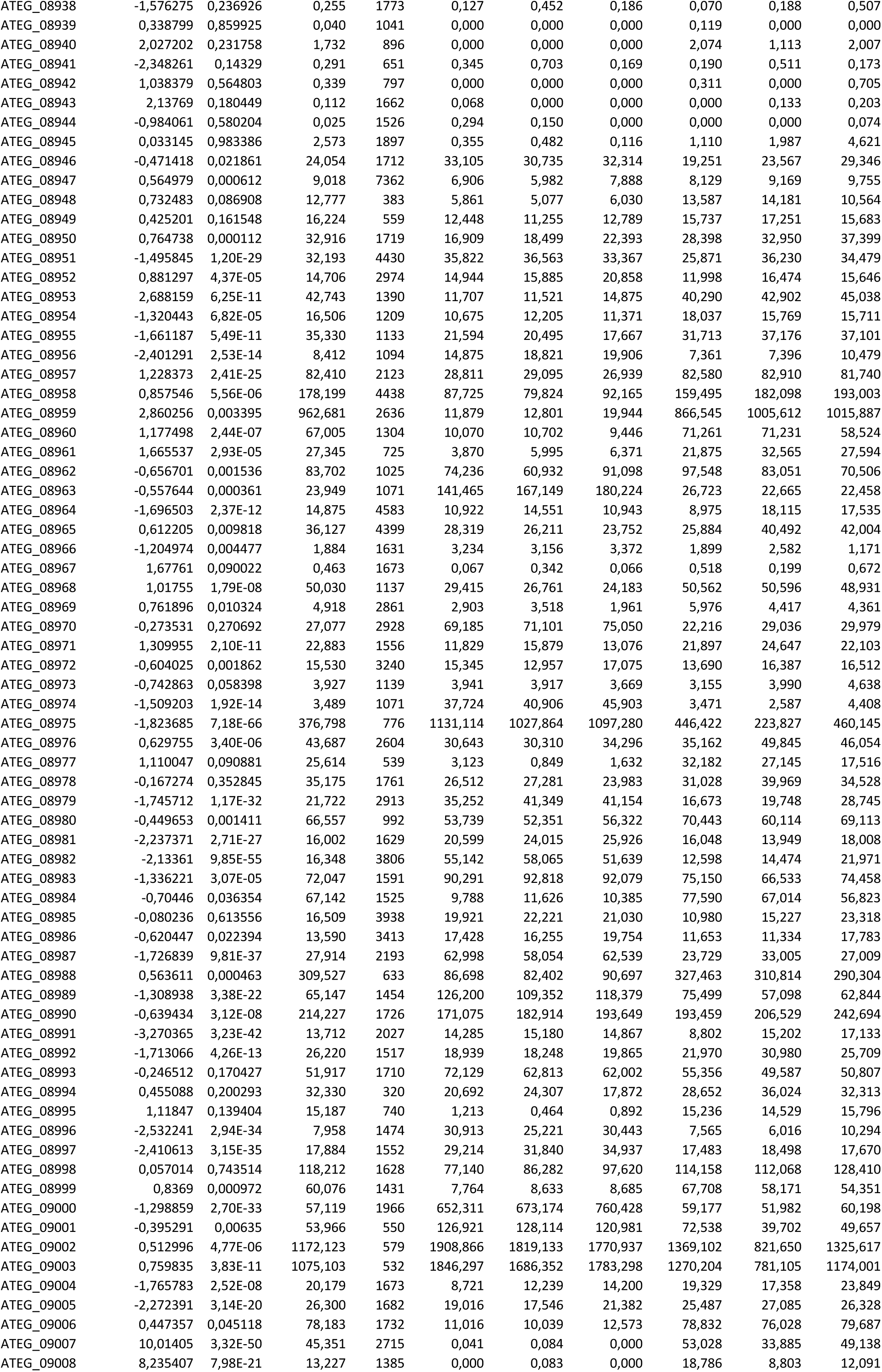
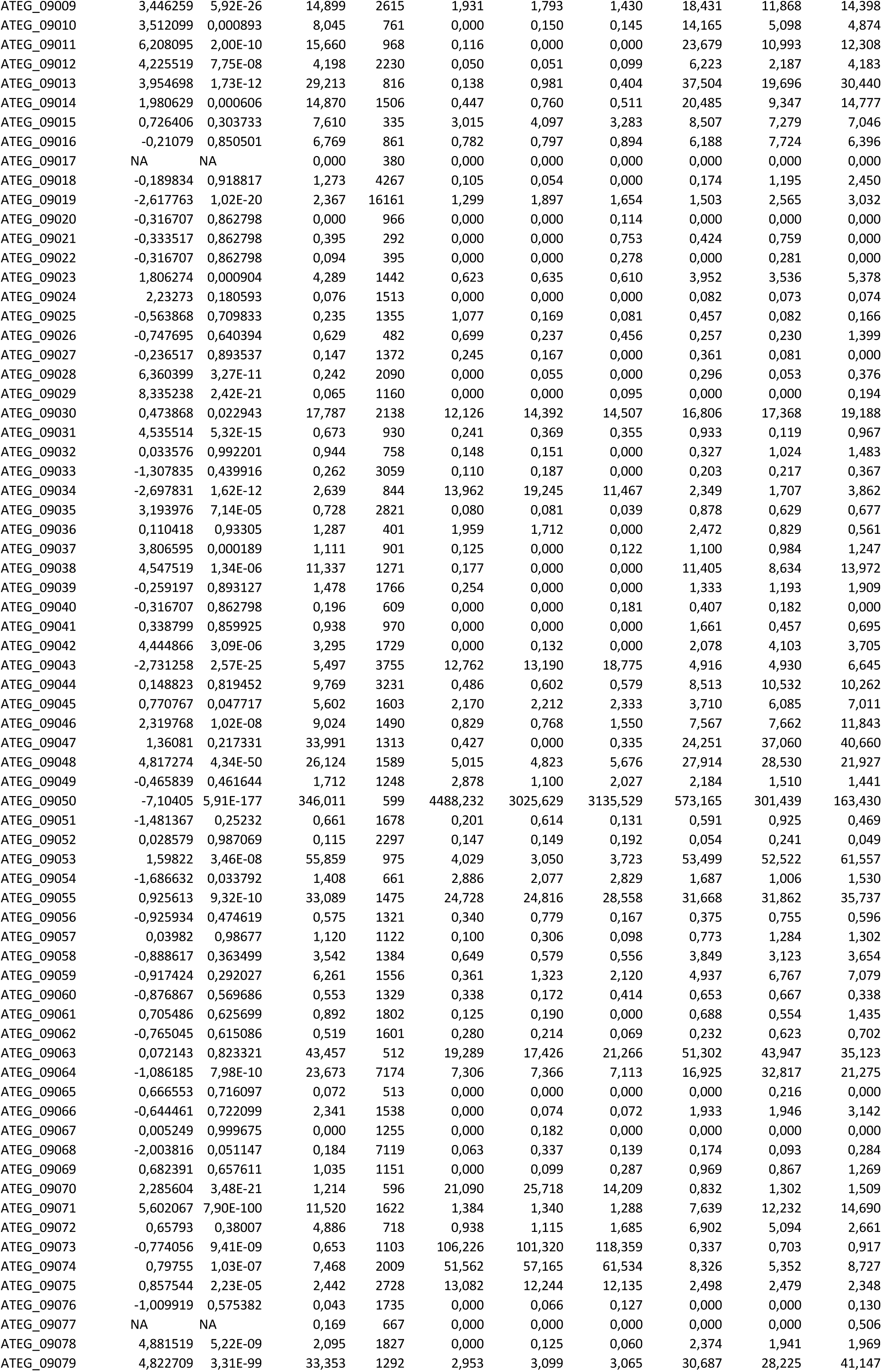
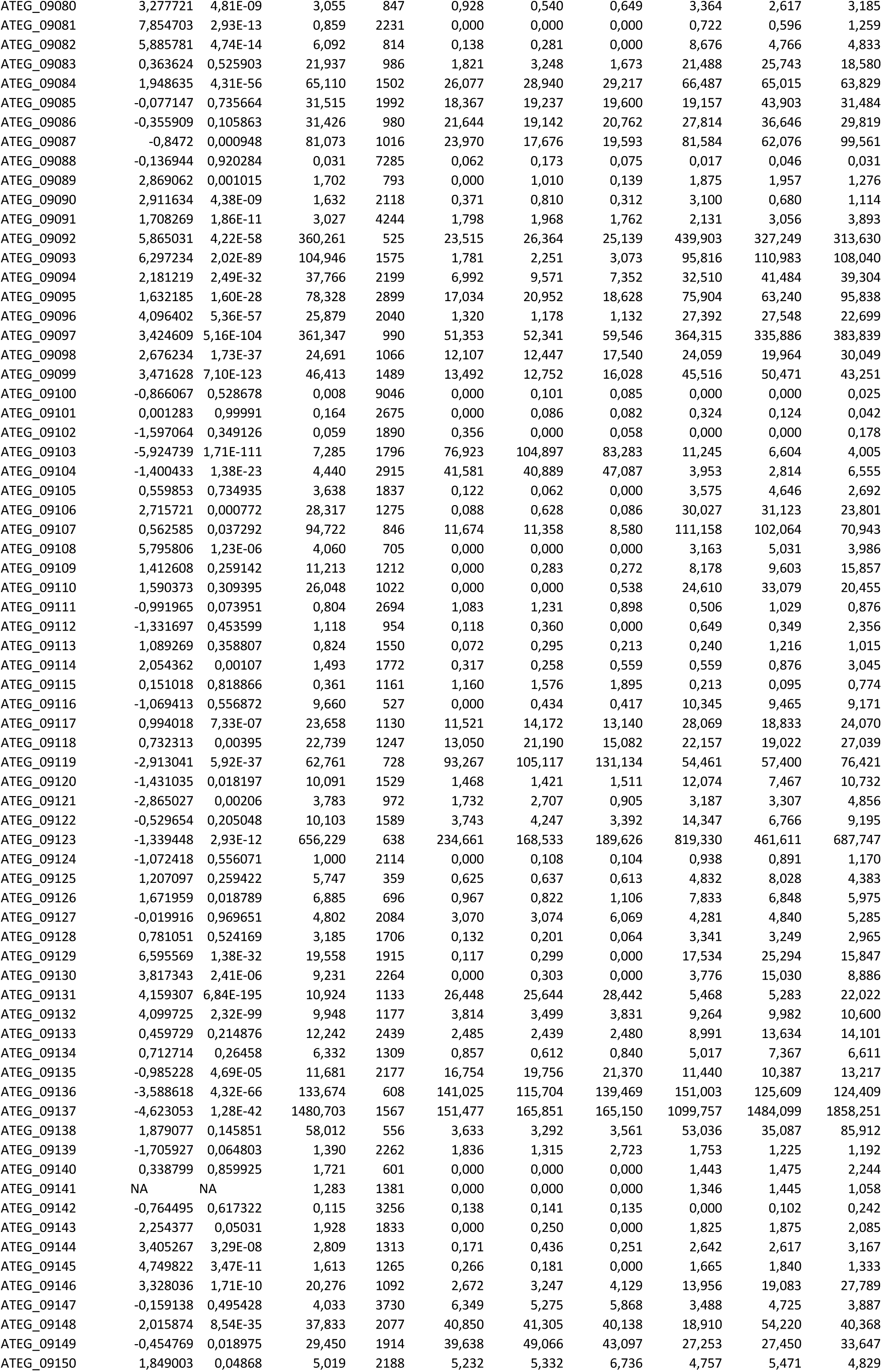
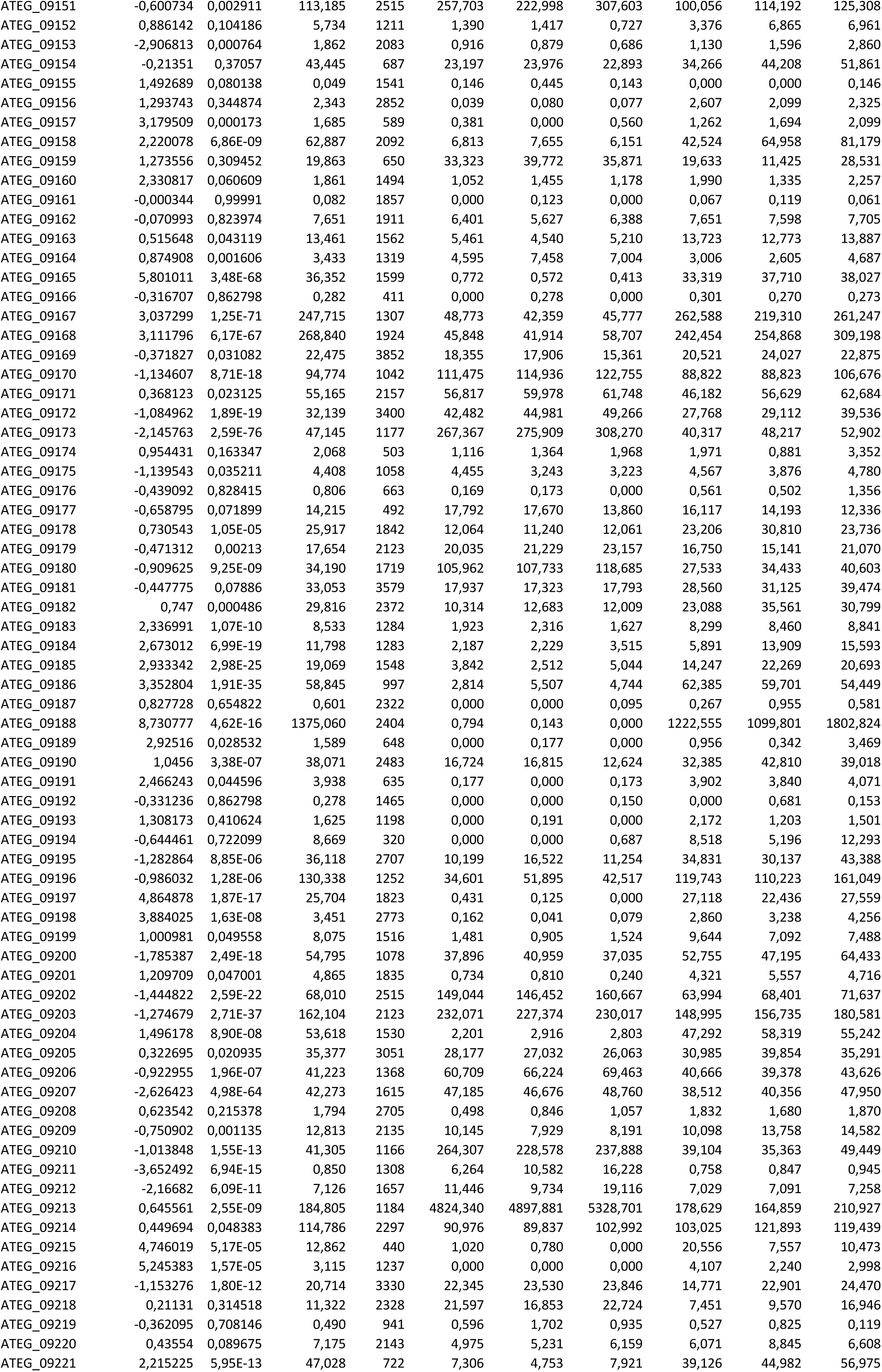
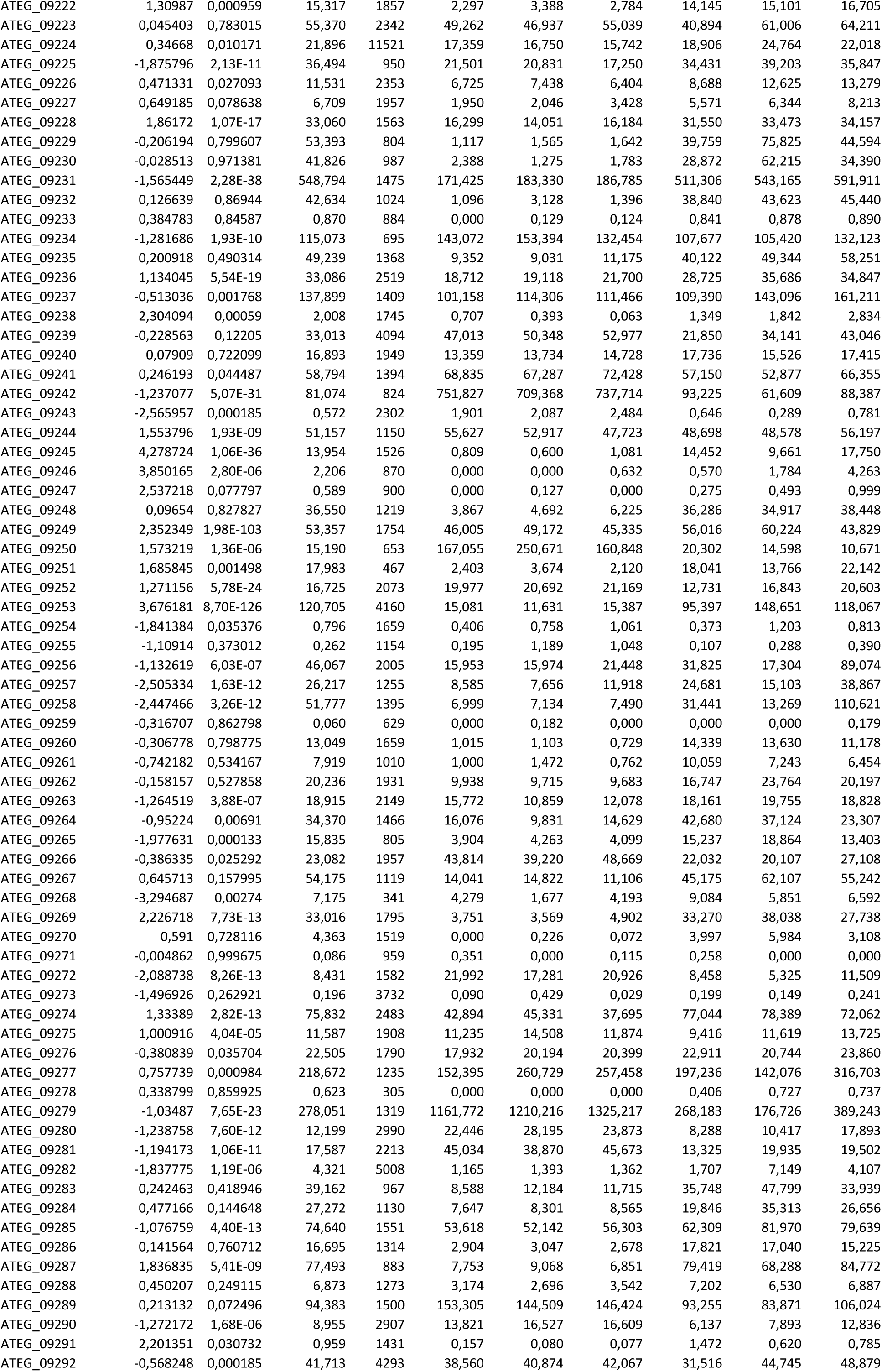
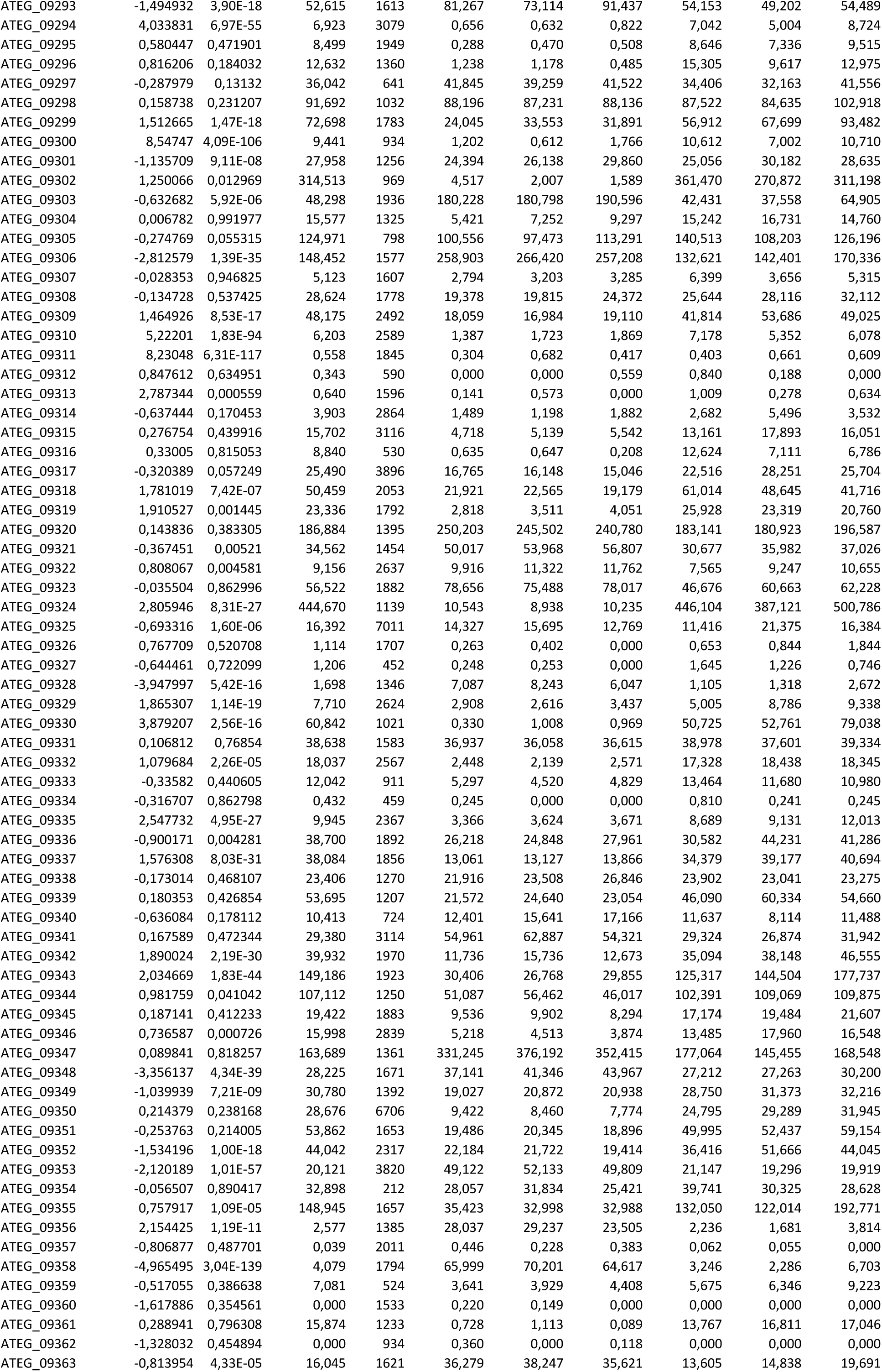
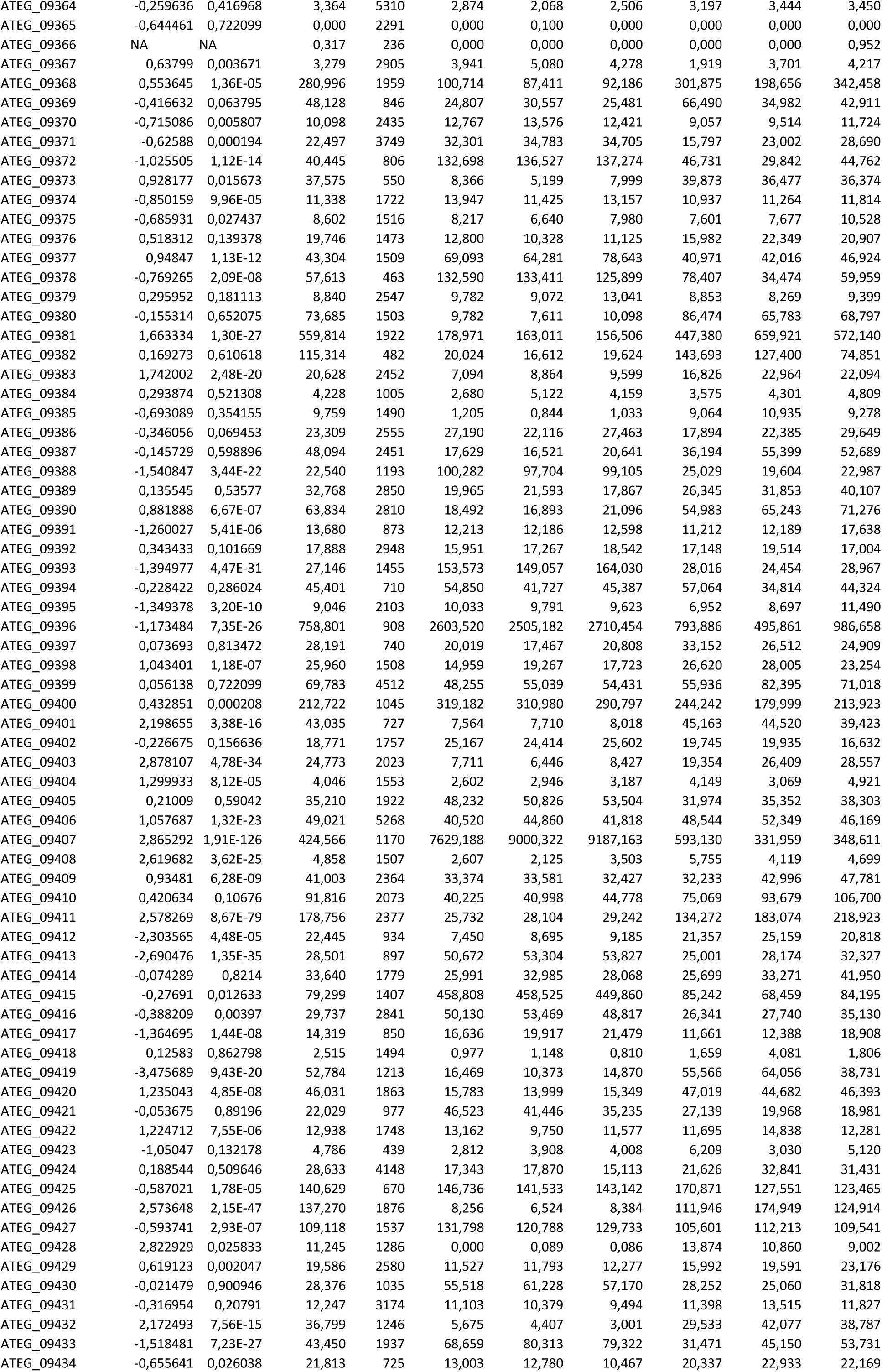
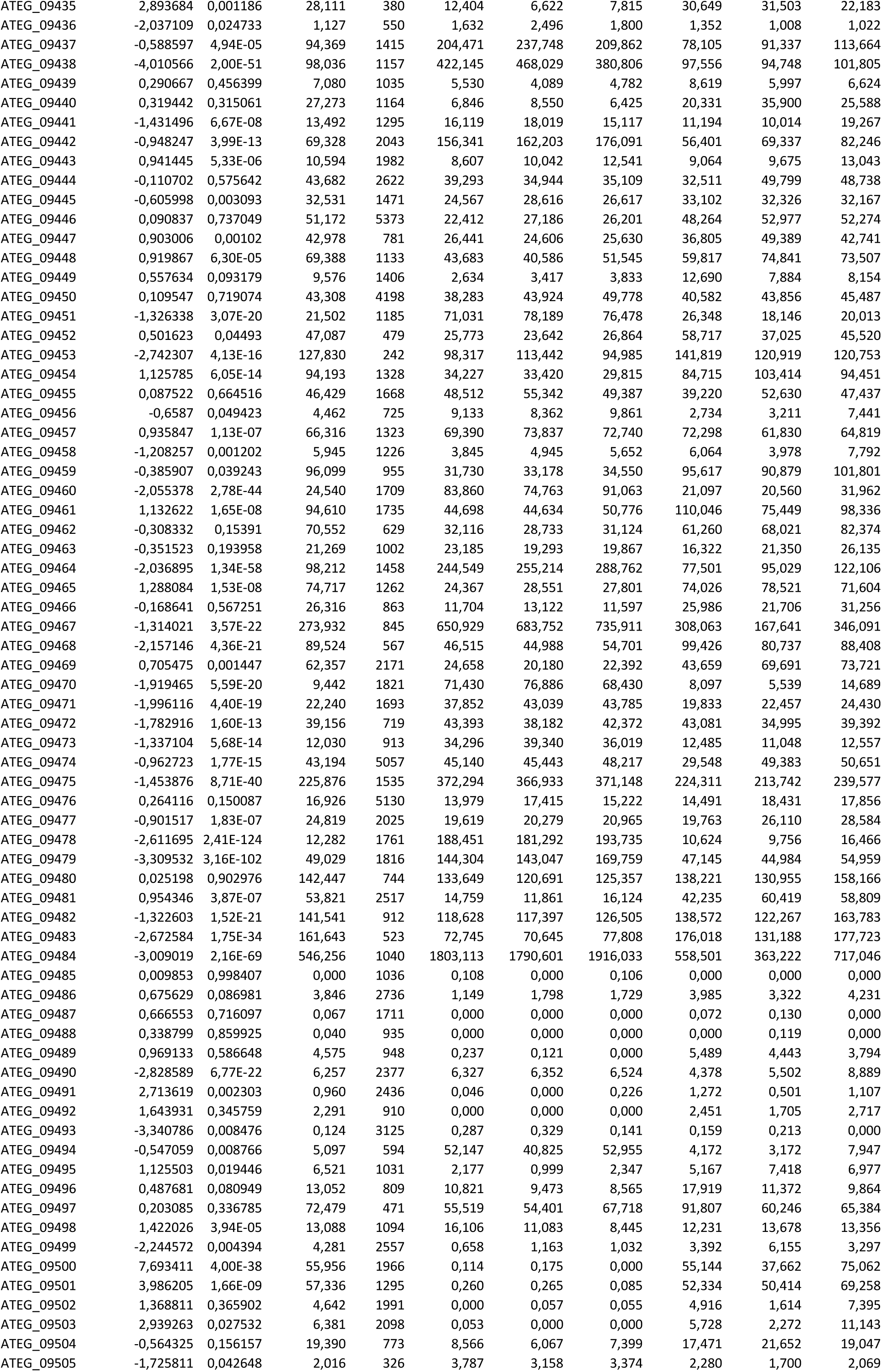
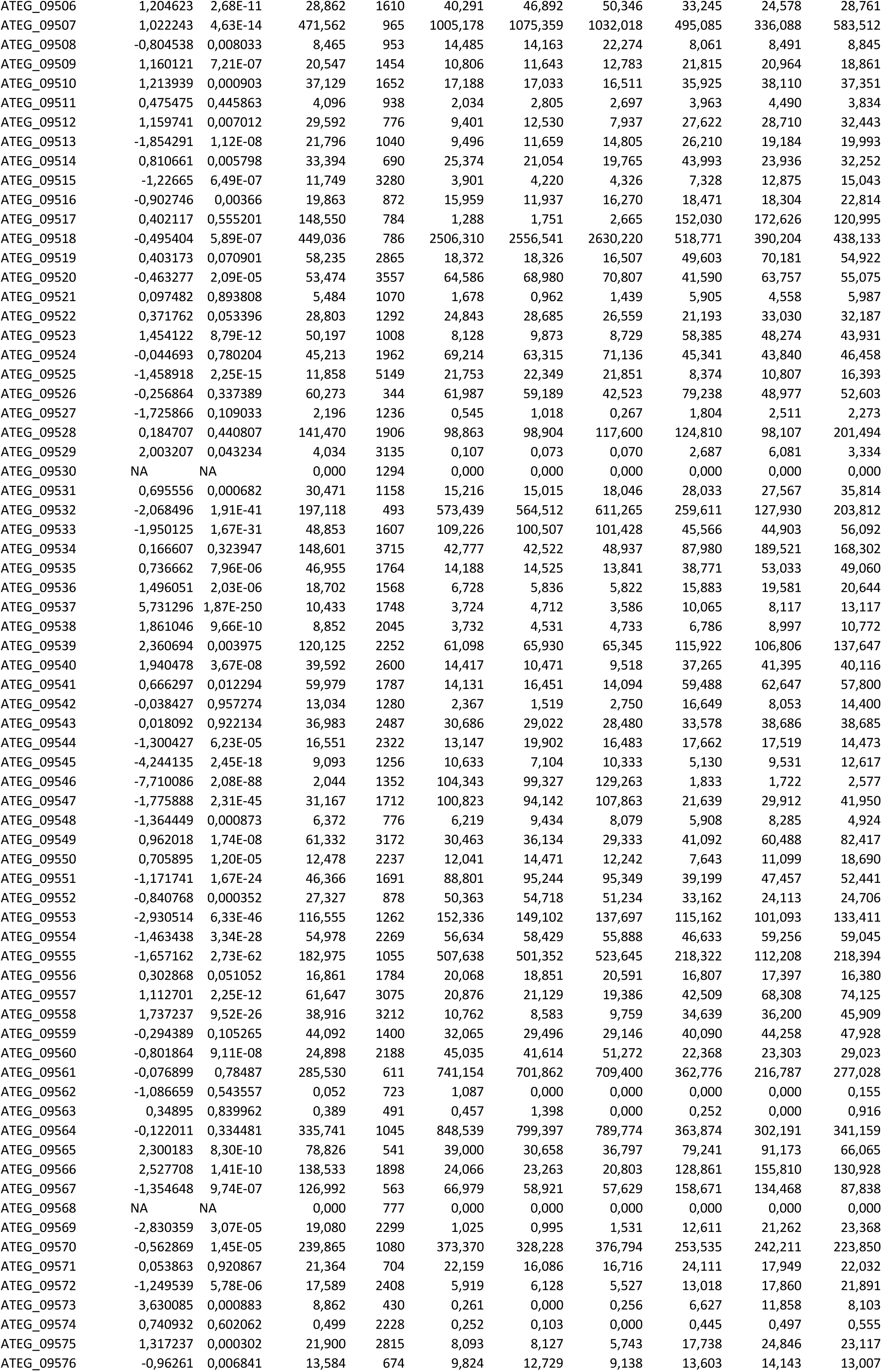
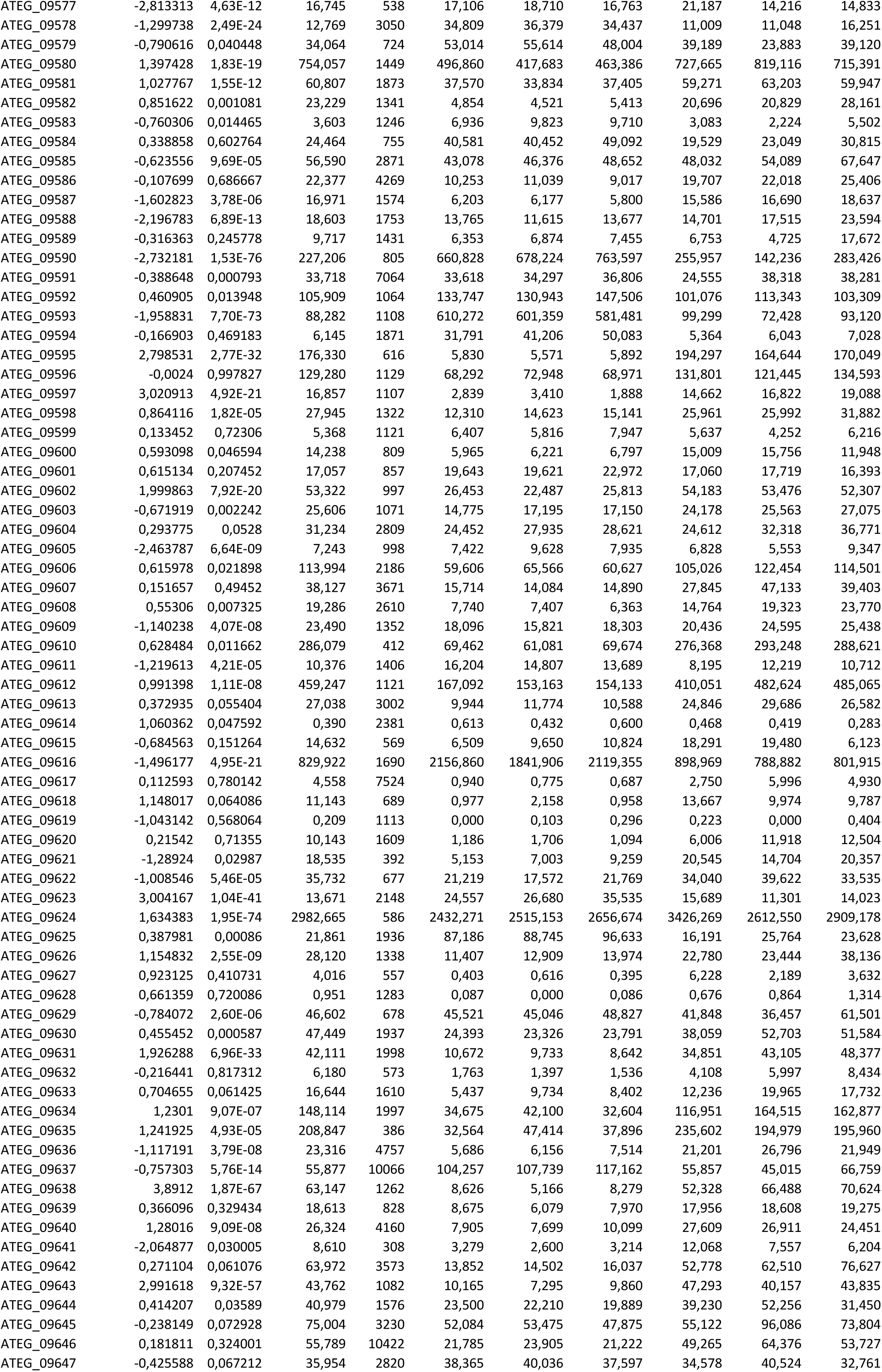
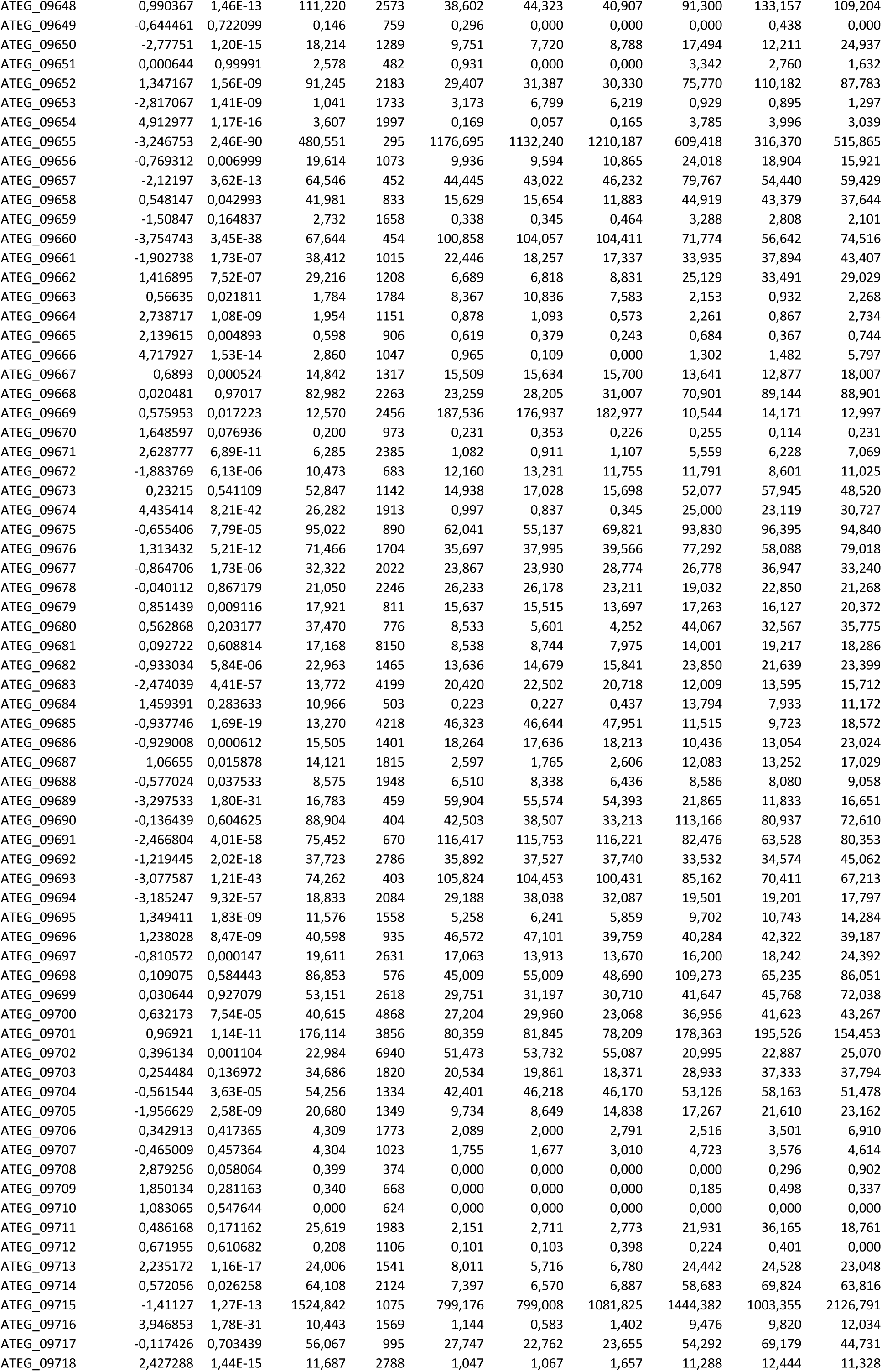
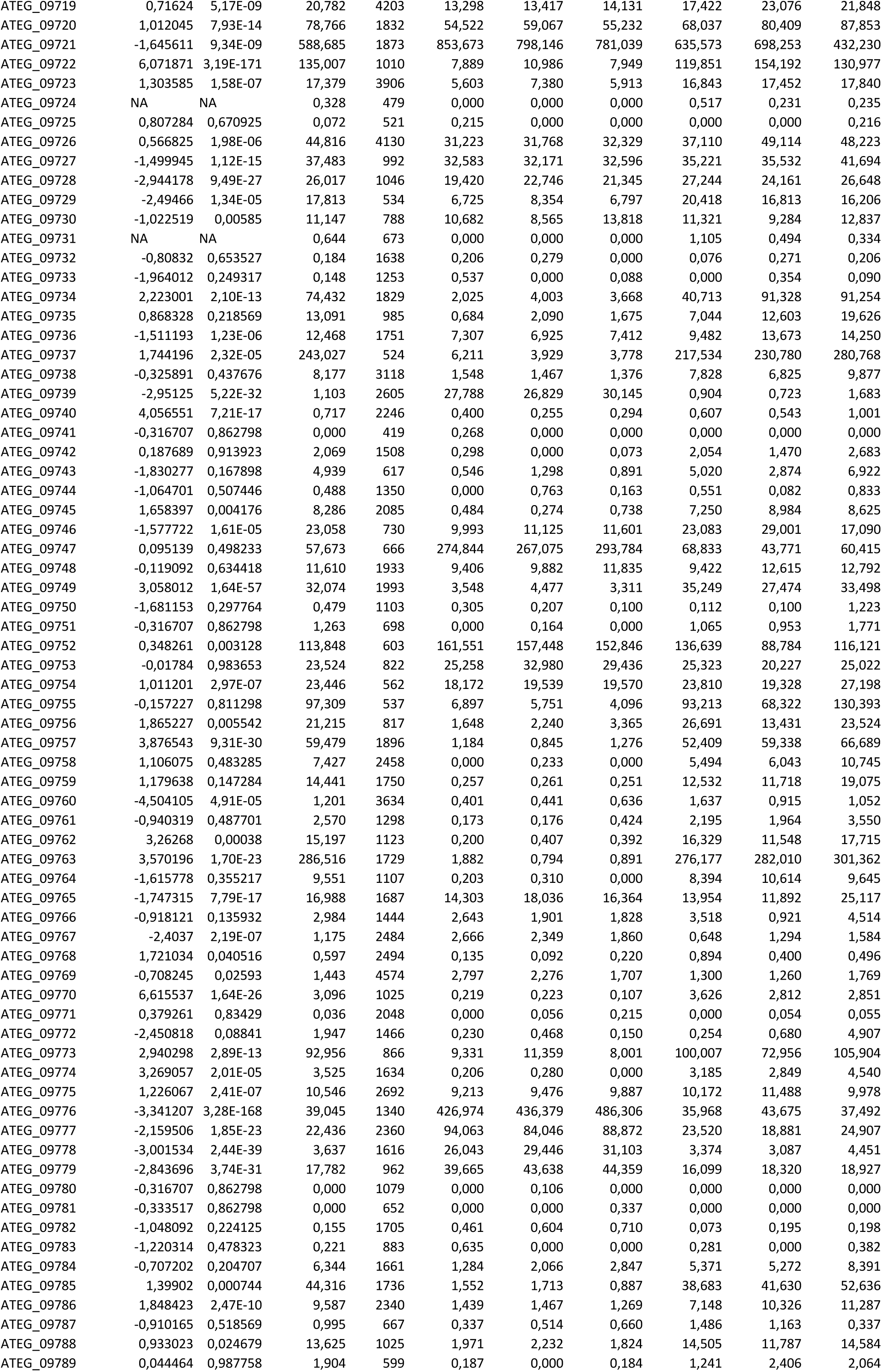
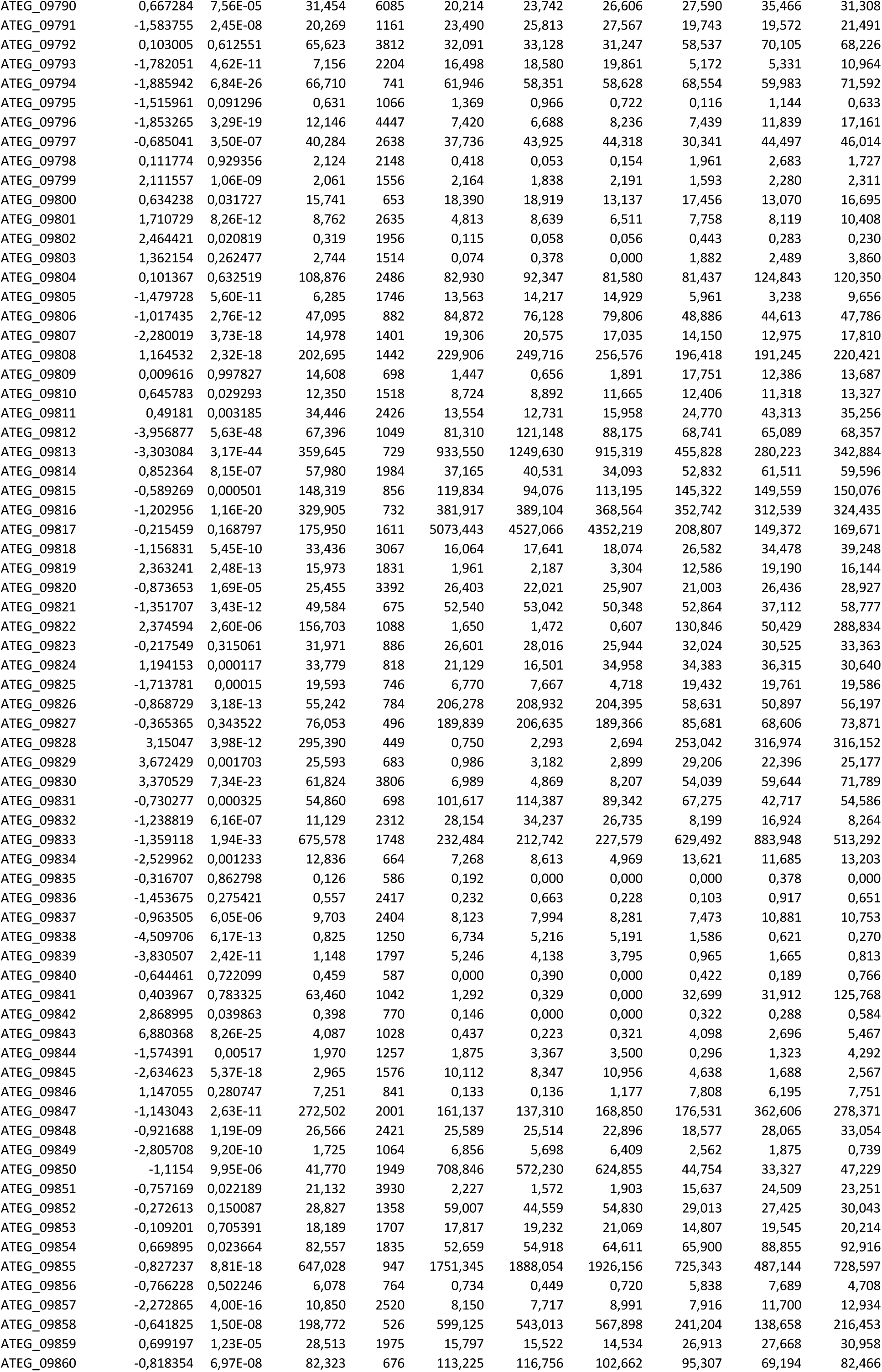
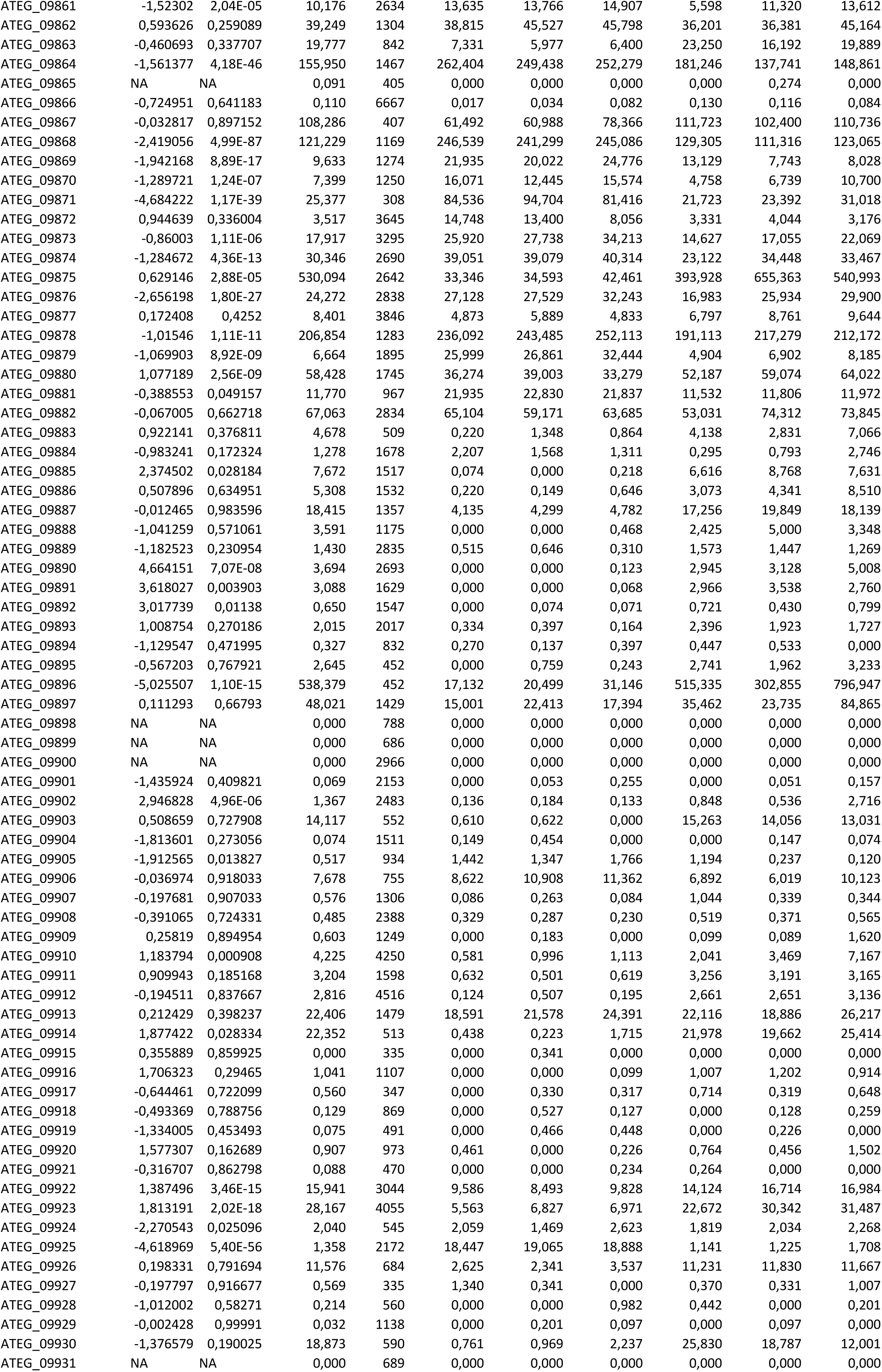
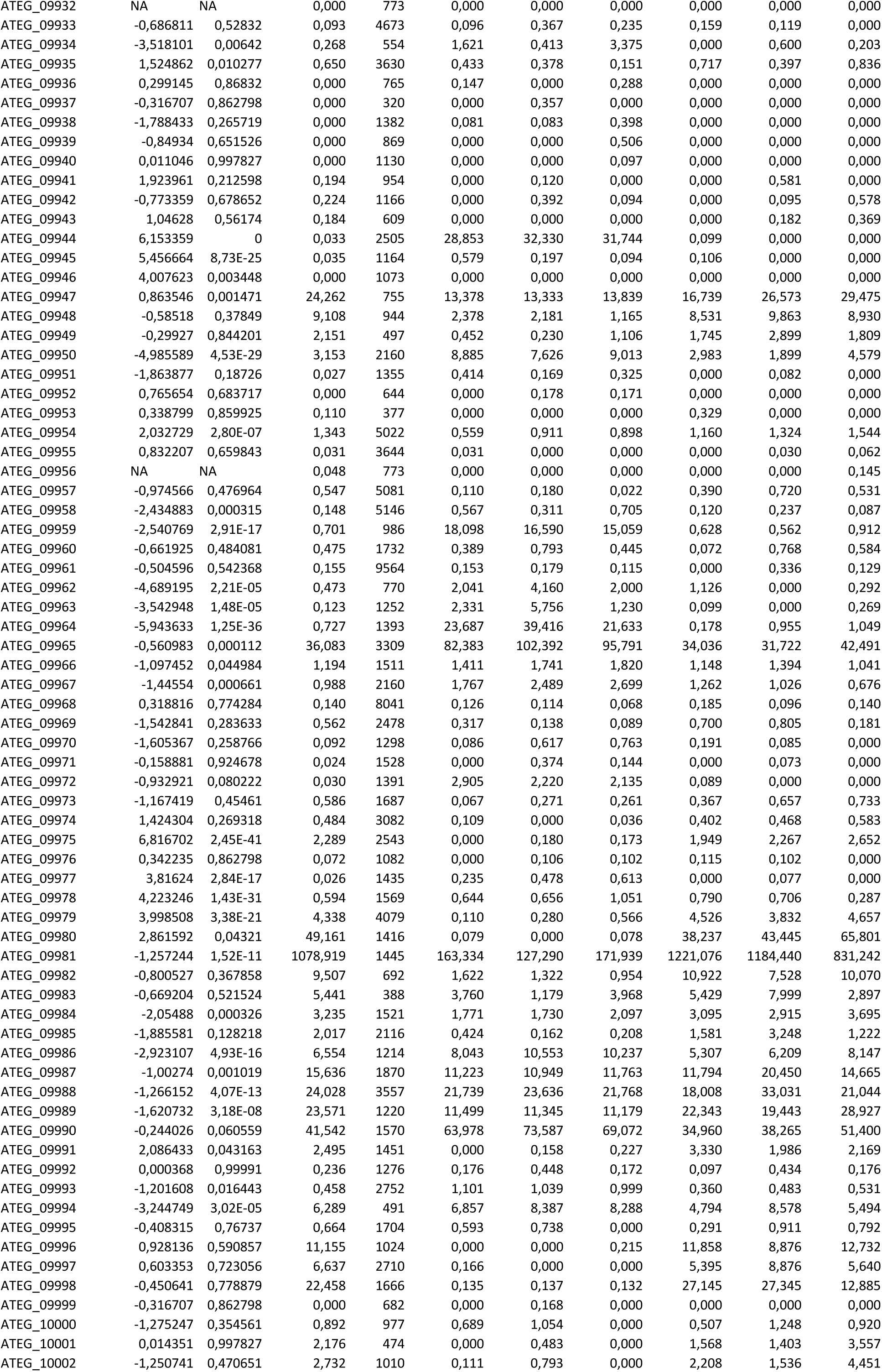
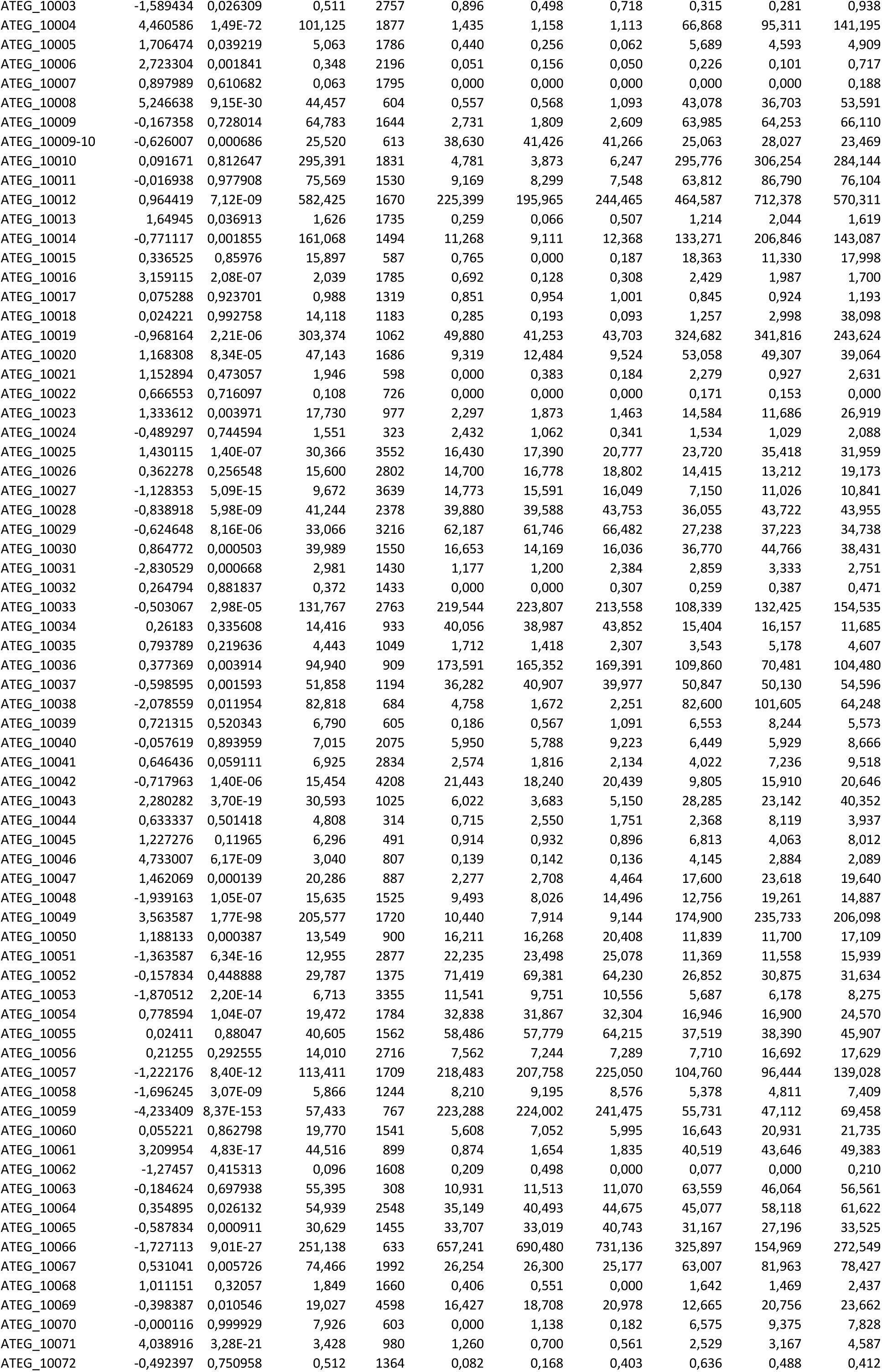
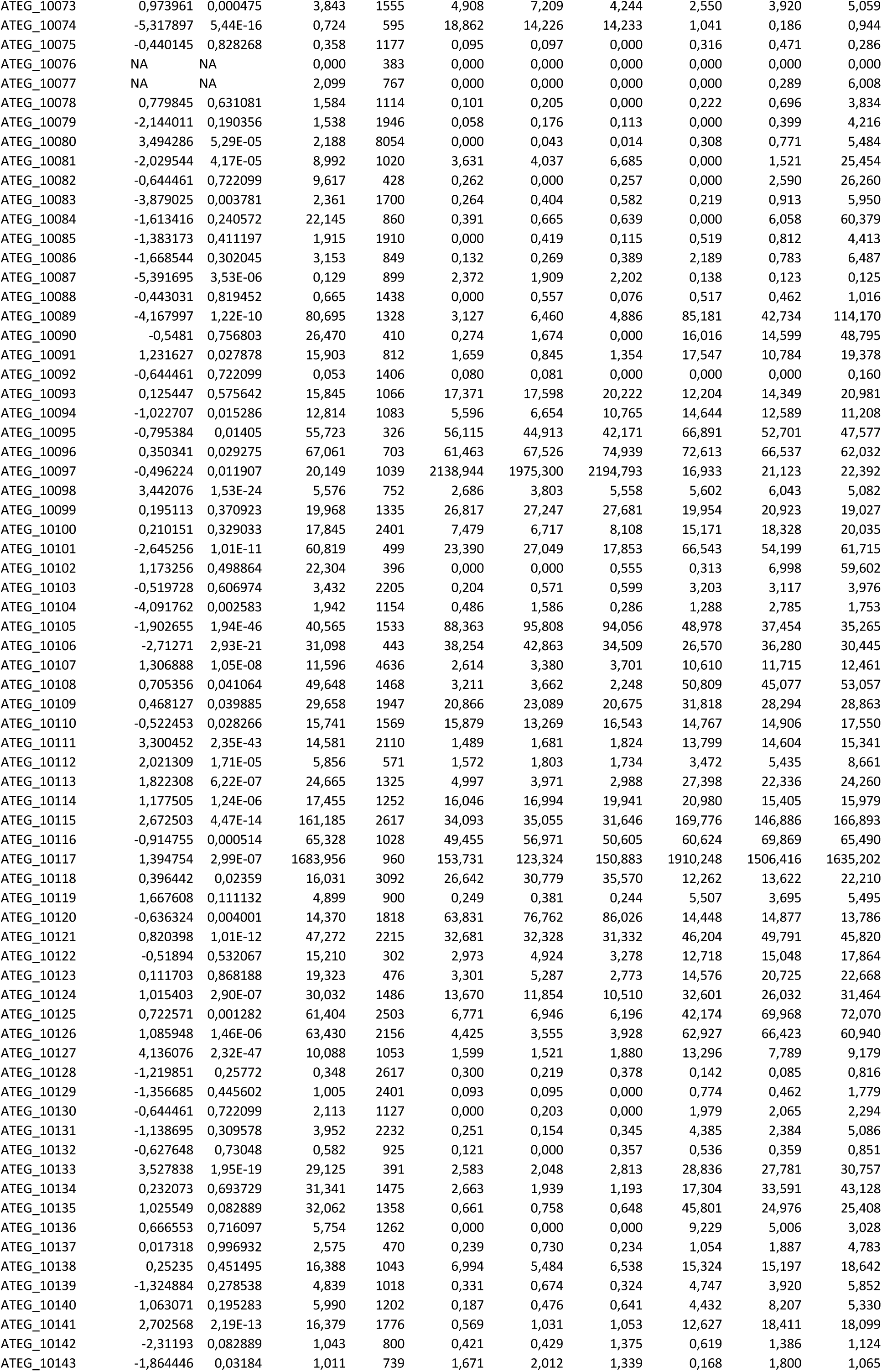
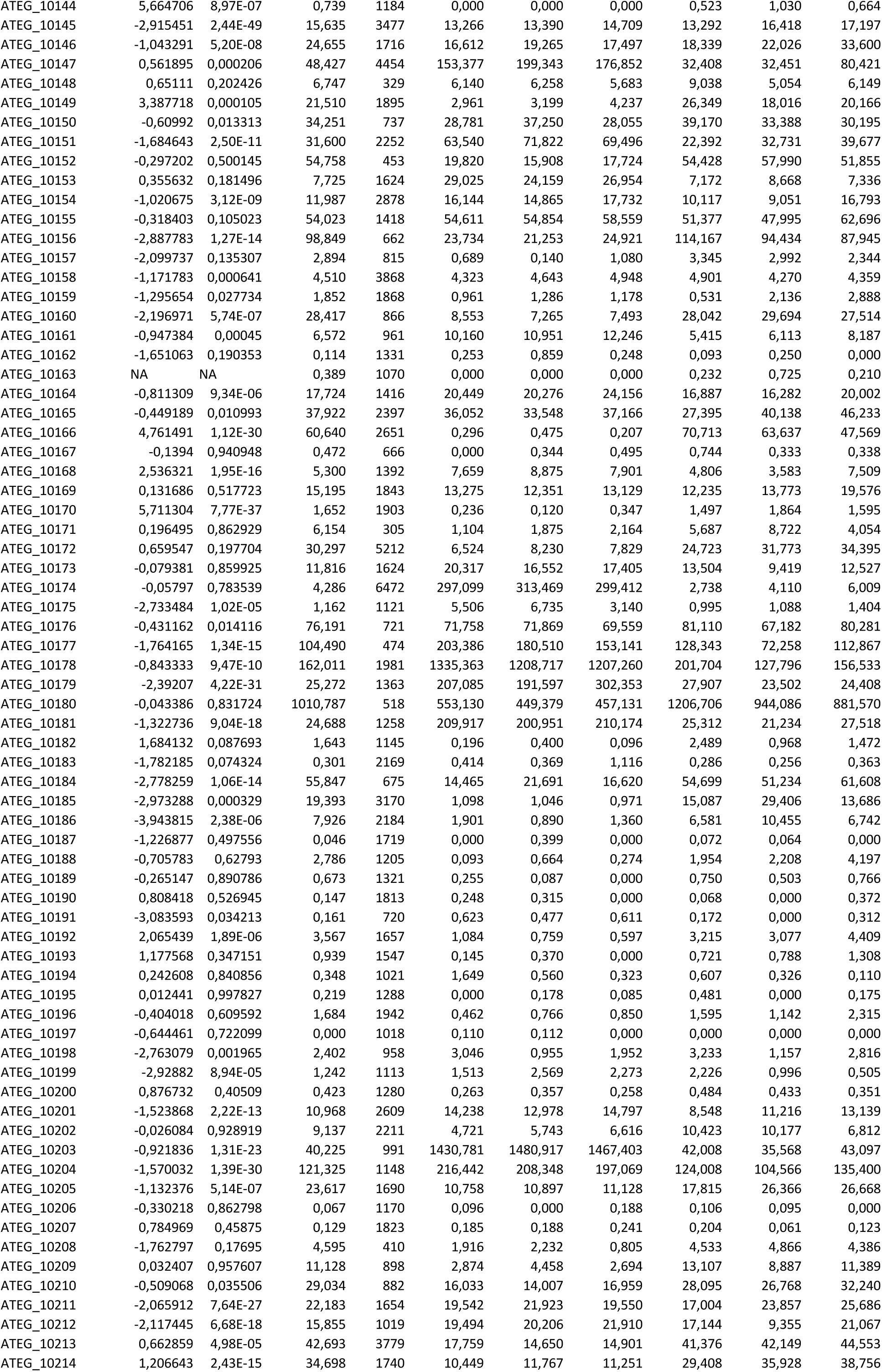
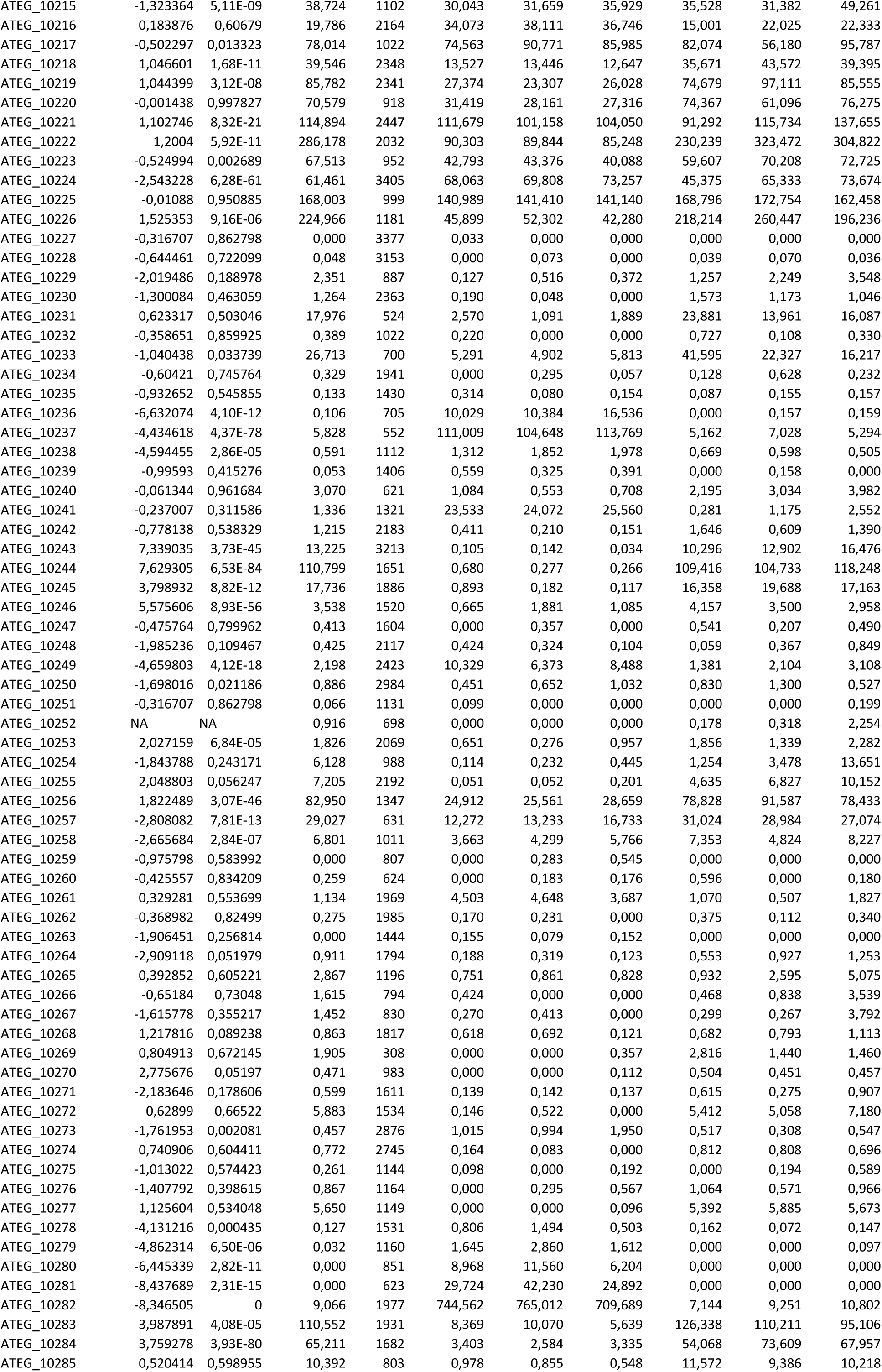
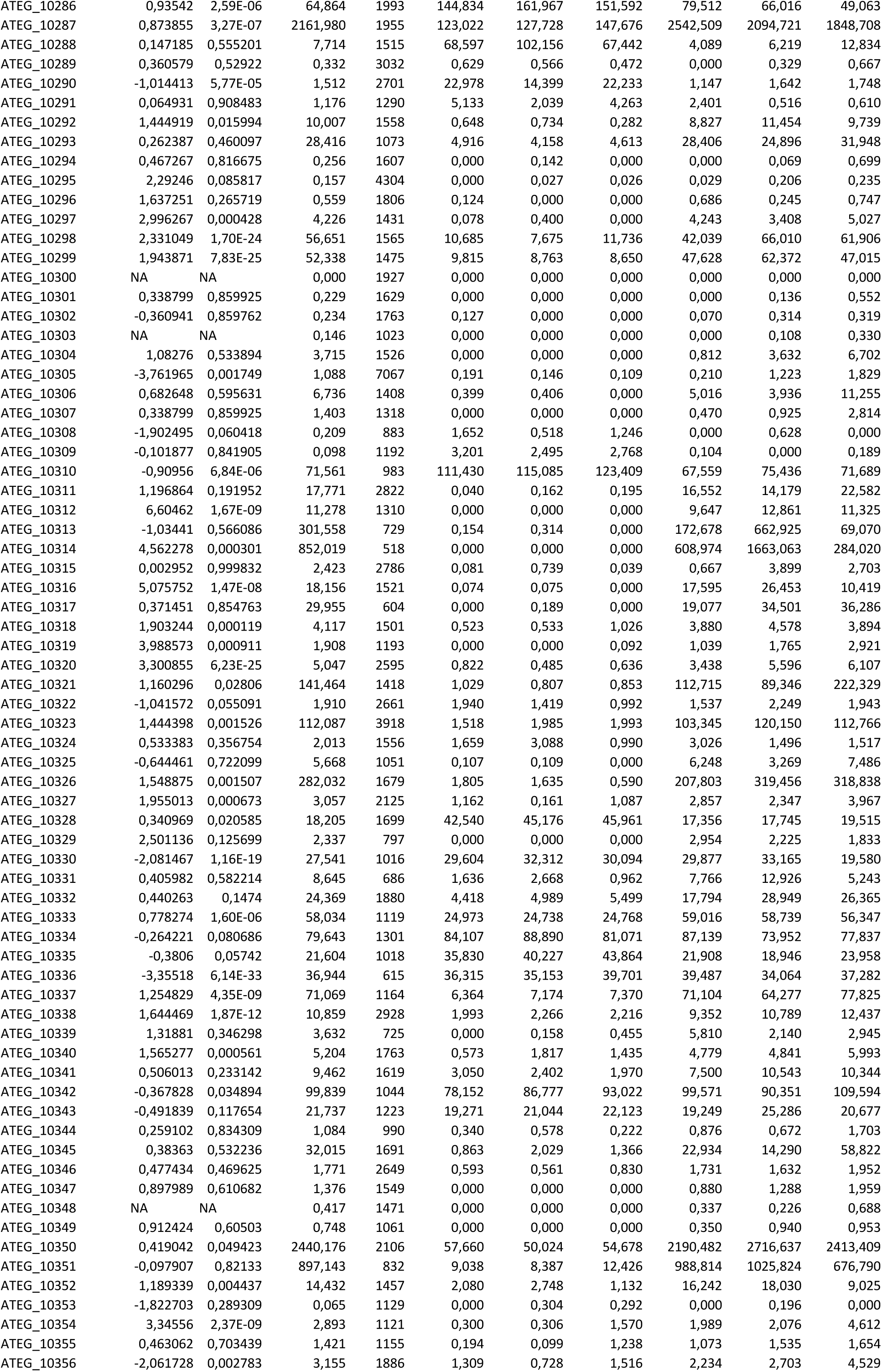
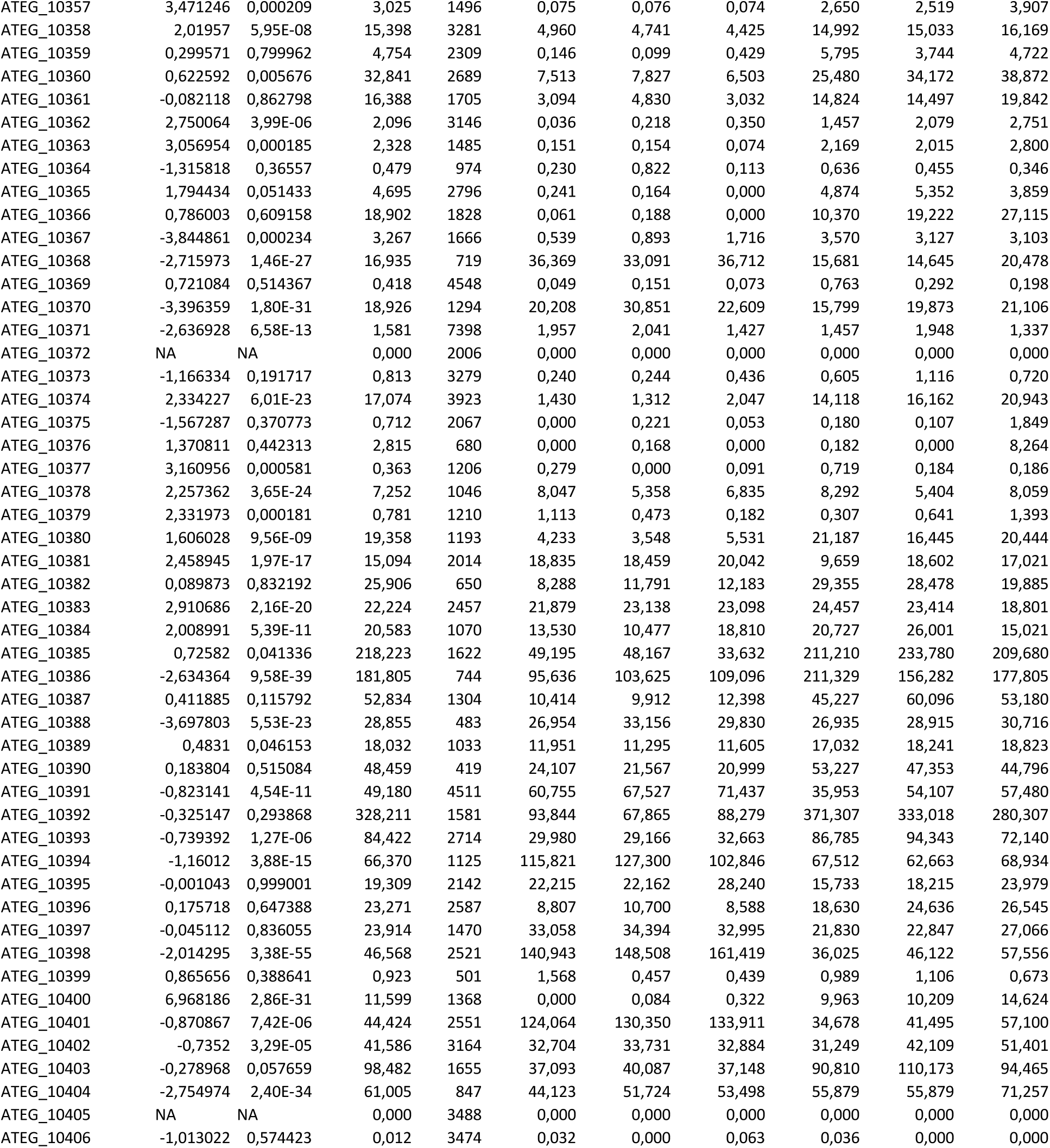
*Aspergillus terreus* RNA-seq gene expression data.

**Table S7.**
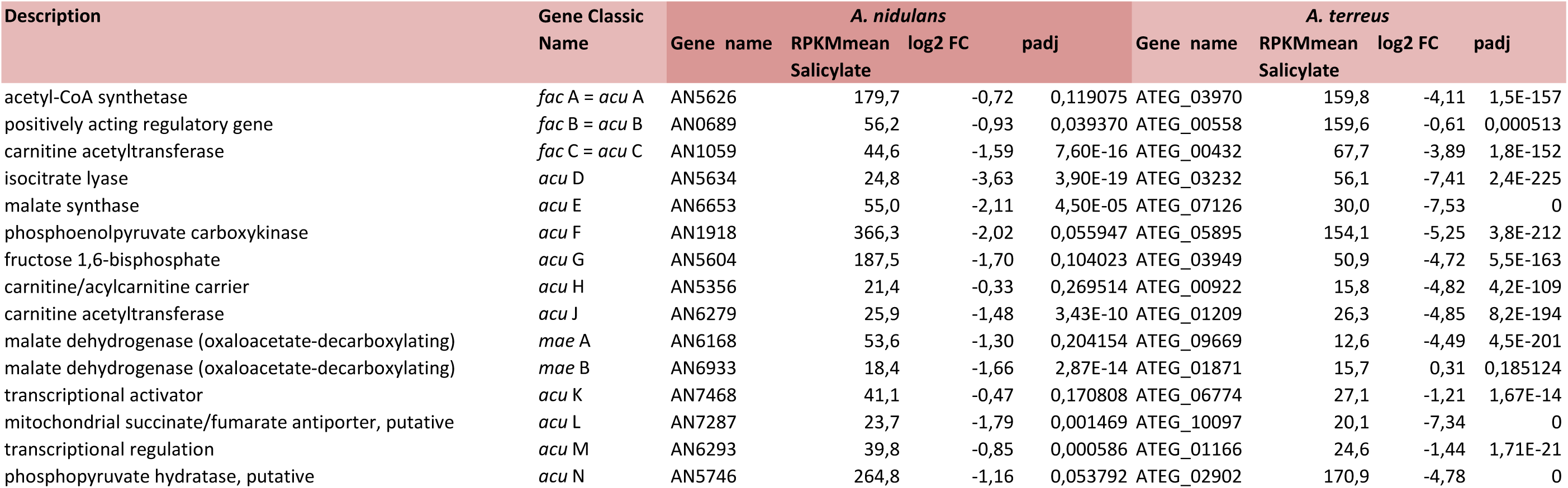
Expression levels of genes coding in the acetate metabolism in response to salicylate in *Aspergillus nidulans* and *Aspergillus terreus* (RNA-seq data).

**Table S8.**
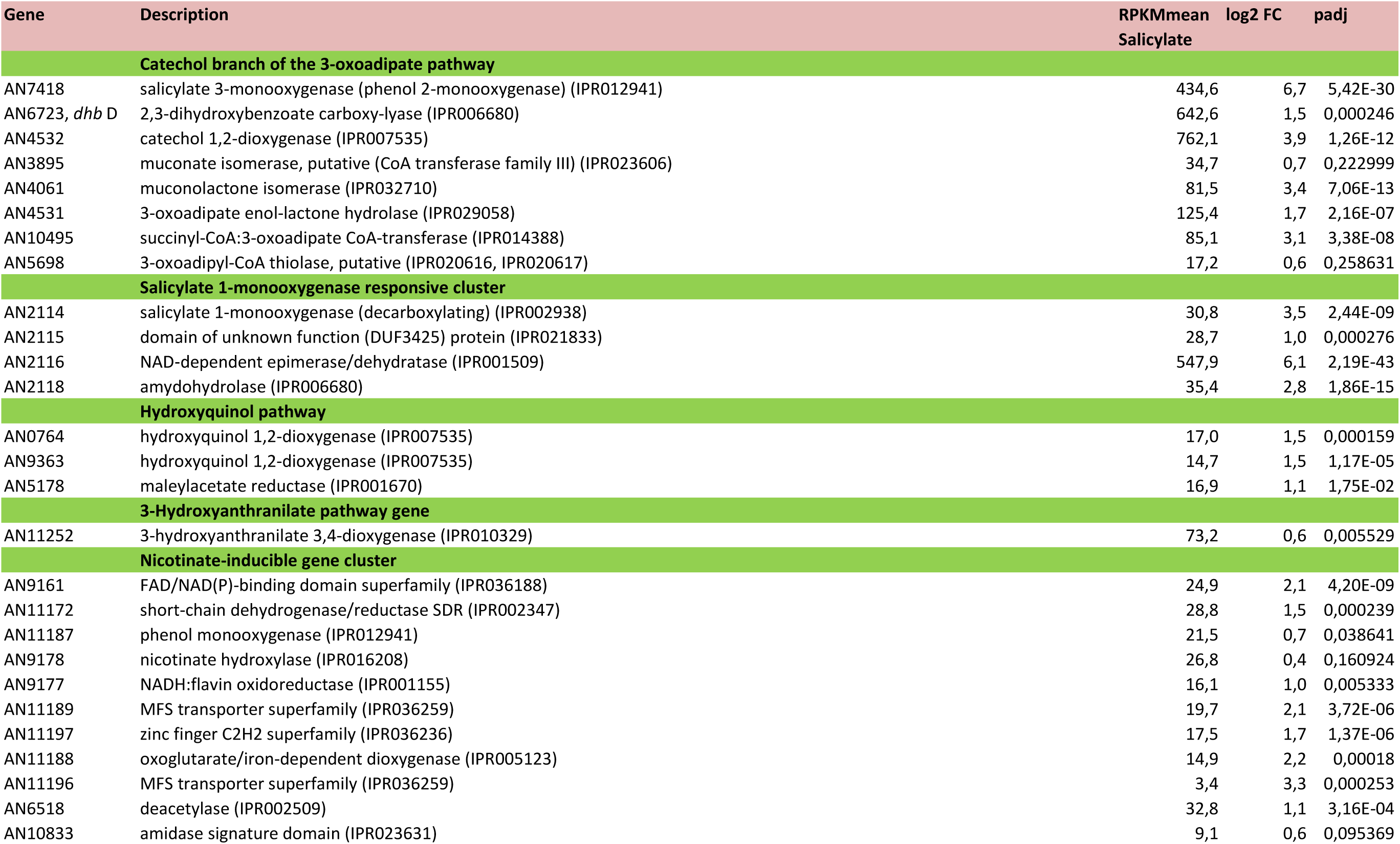
Expression levels of genes of the catechol branch of the 3-oxoadipate pathway (and of other relevant pathways) in response to salicylate in *Aspergillus nidulans* (RNA-seq data).

**Table S9.**
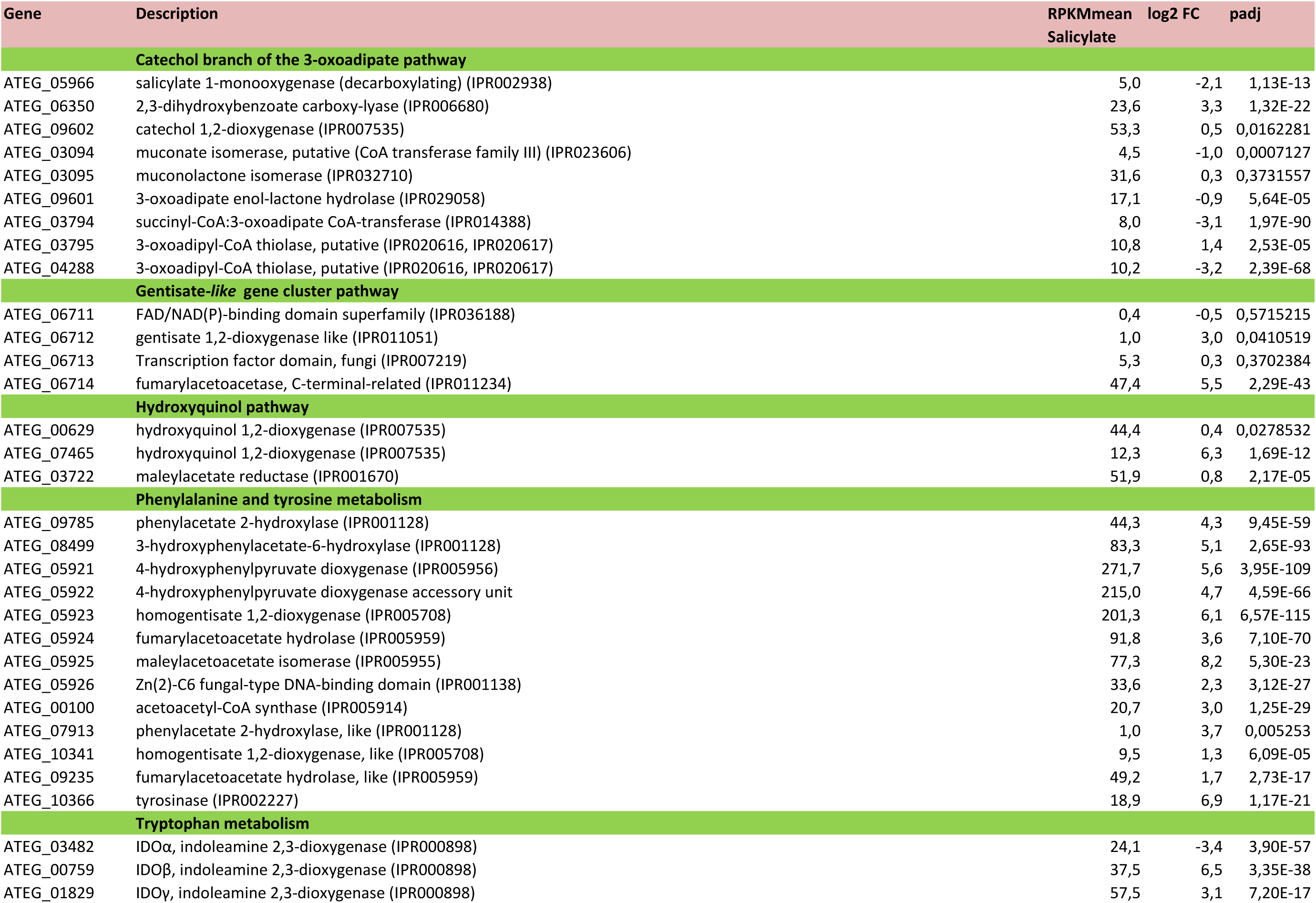
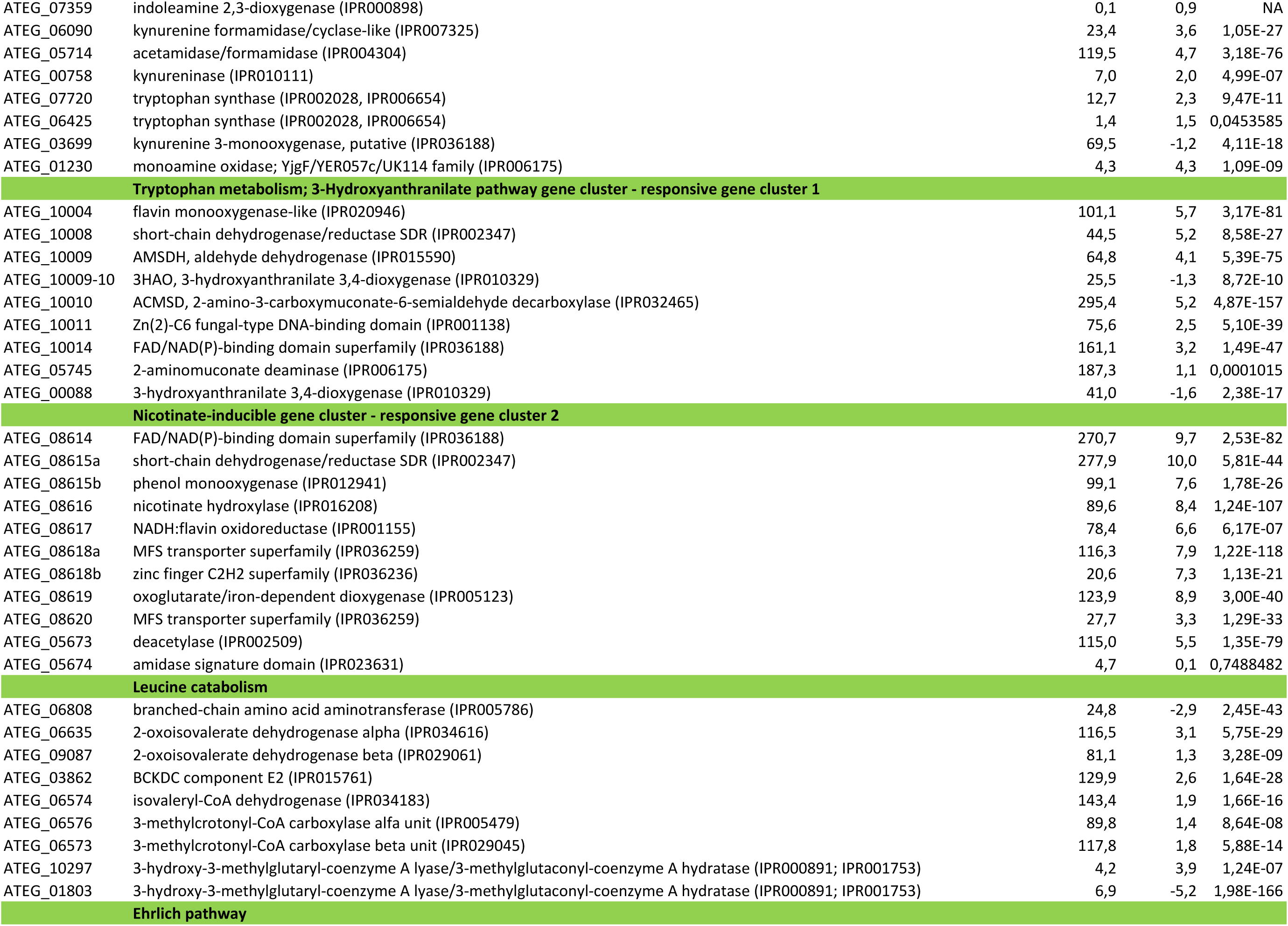
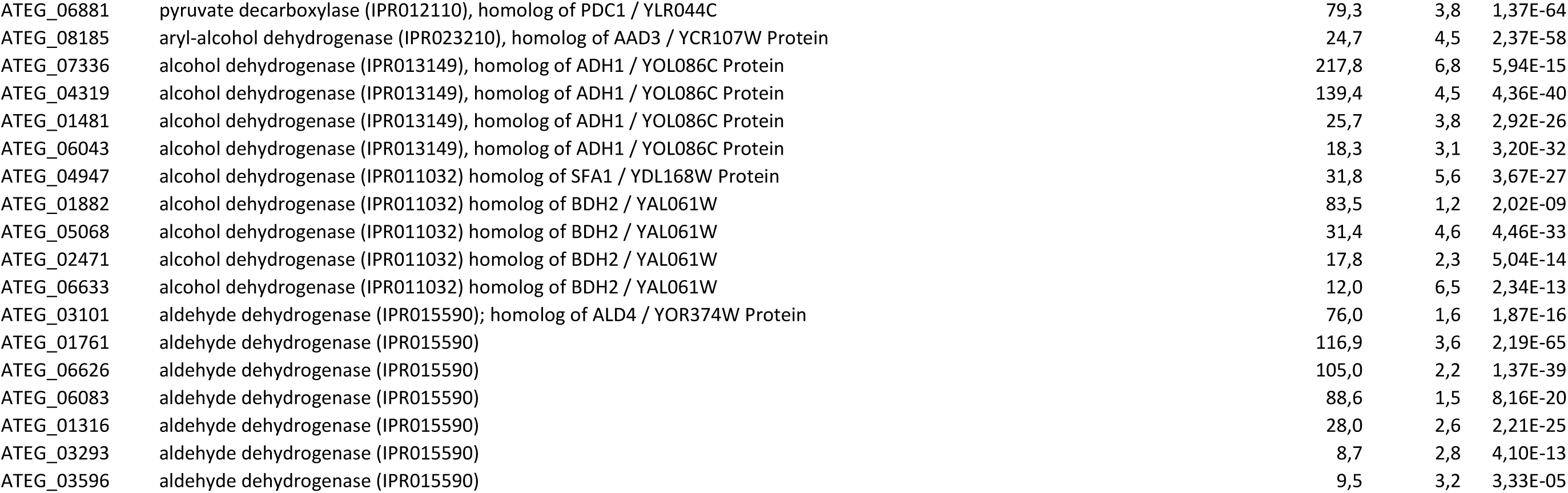
Expression levels of genes of the of the catechol branch of the 3-oxoadipate pathway, the presumed gentisate pathway, the phenylalanine, the tyrosine, the tryptophan metabolism and other two relevant gene clusters, in response to salicylate in *Aspergillus terreus* (RNA-seq data).

**Figure S1.**
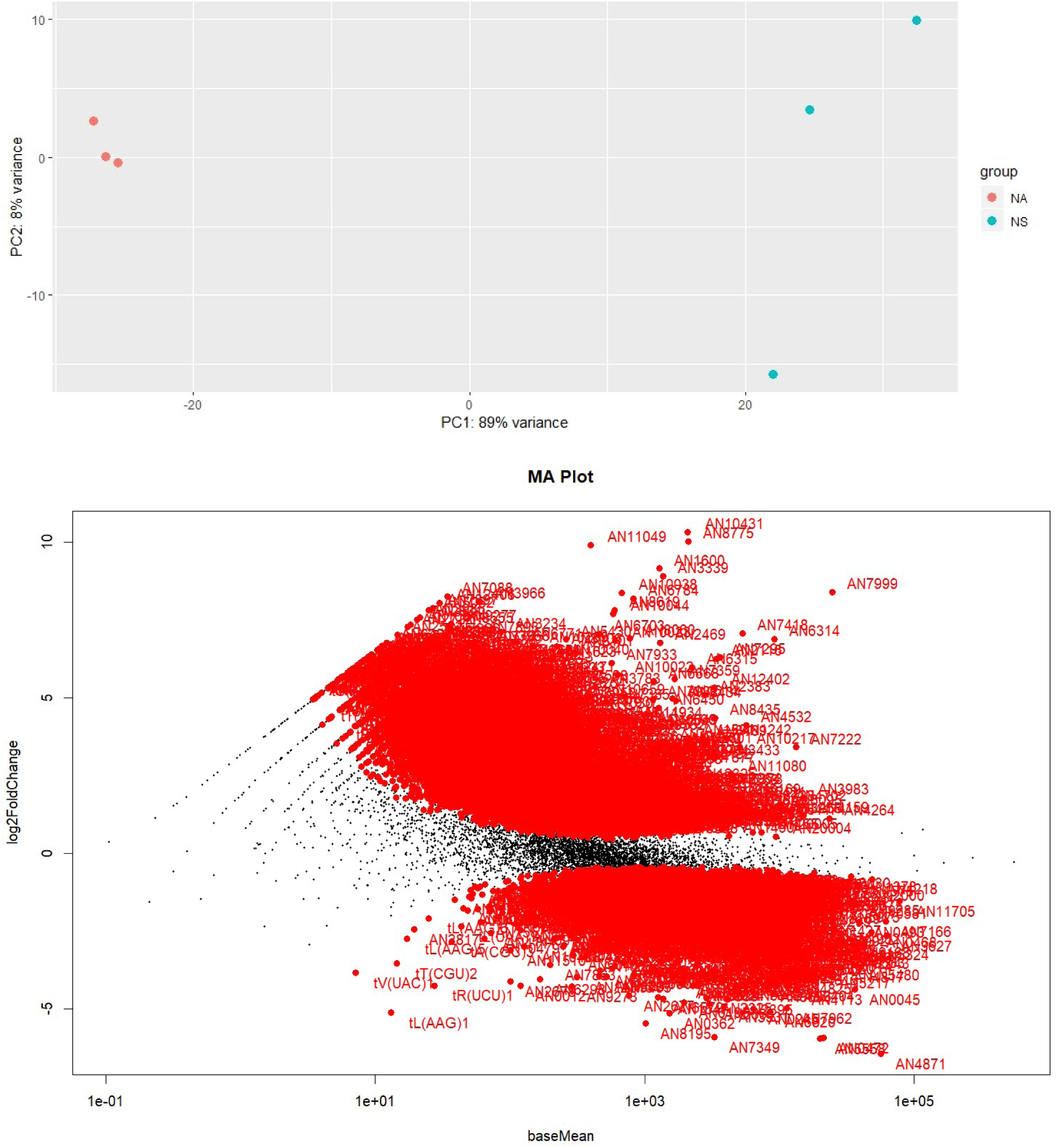
Baseline statistical representation of the RNA-seq data collected for *Aspergillus nidulans* samples. The Principal Component Analysis (top), shows the distribution of the whole-transcriptome reads of each of the analyzed replicas (NA - acetate medium; NS - salicylate medium). In the MA plot (bottom) is possible to visualize the differences between the measurements taken between the expression of genes in acetate medium and salicylate medium. Both plots were computed using the tools embedded in the DEseq2 package.

**Figure S2.**
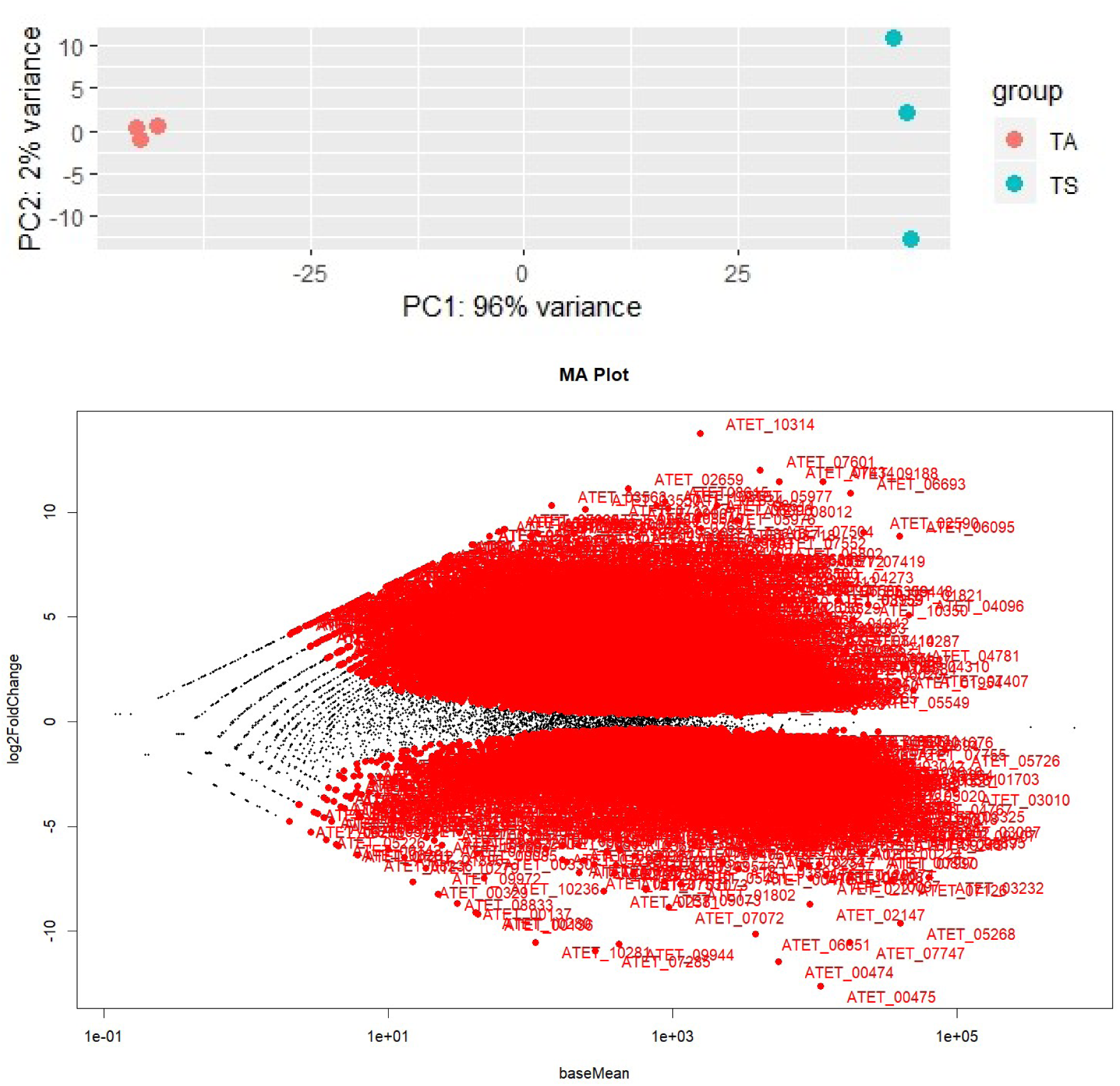
Baseline statistical representation of the RNA-seq data collected for *Aspergillus terreus* samples. The Principal Component Analysis (top), shows the distribution of the whole-transcriptome reads of each of the analyzed replicas (TA - acetate medium; TS - salicylate medium). In the MA plot (bottom) is possible to visualize the differences between the measurements taken between the expression of genes in acetate medium and salicylate medium. Both plots were computed using the tools embedded in the DEseq2 package.

**Figure S3.**
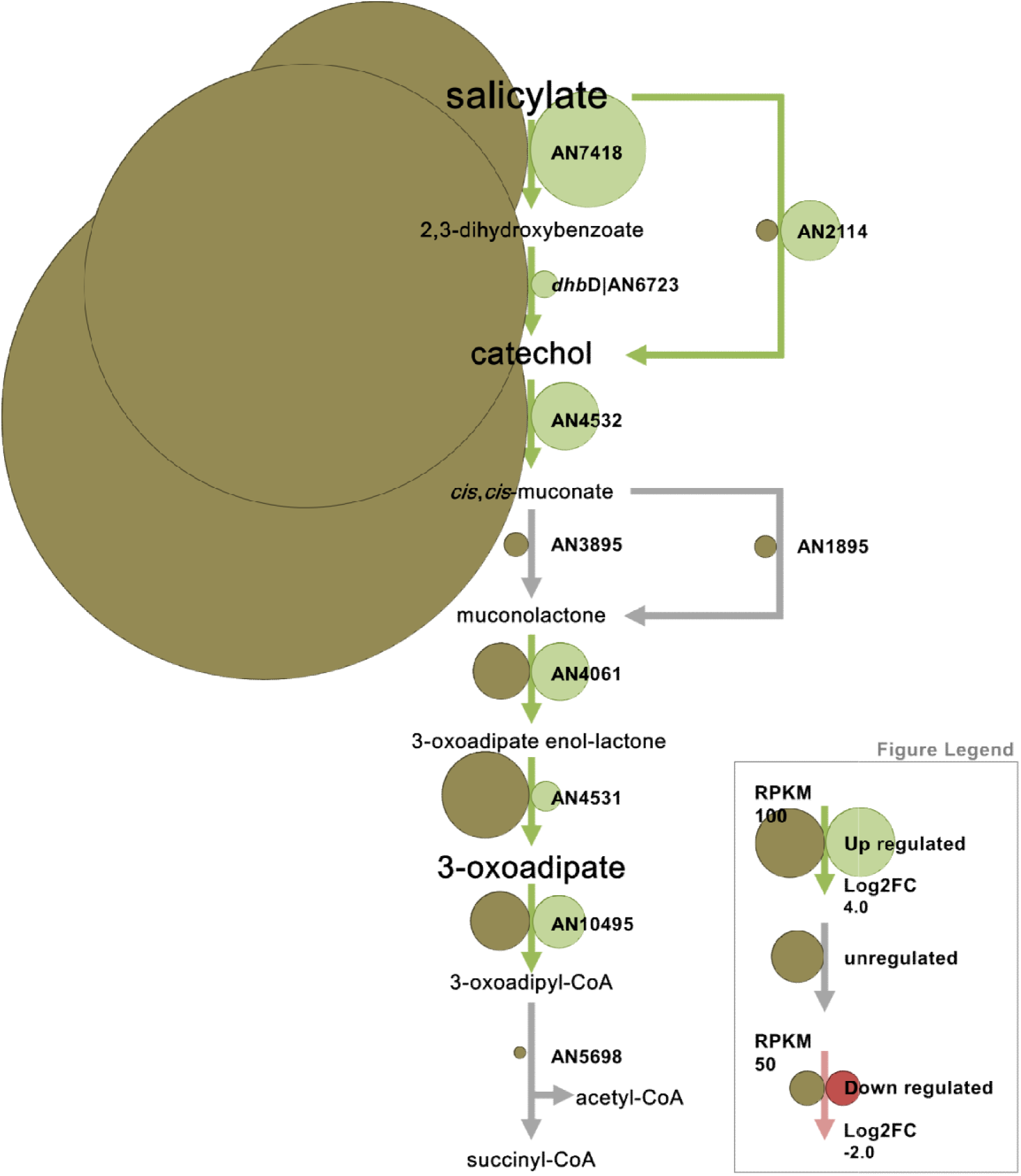
Diagram of the catabolism of salicylate in *A. nidulans*. The differentially expressed genes (RNA-seq) regulation and abundance is represented by circles with variable size and color (see legend).

**Figure S4.**
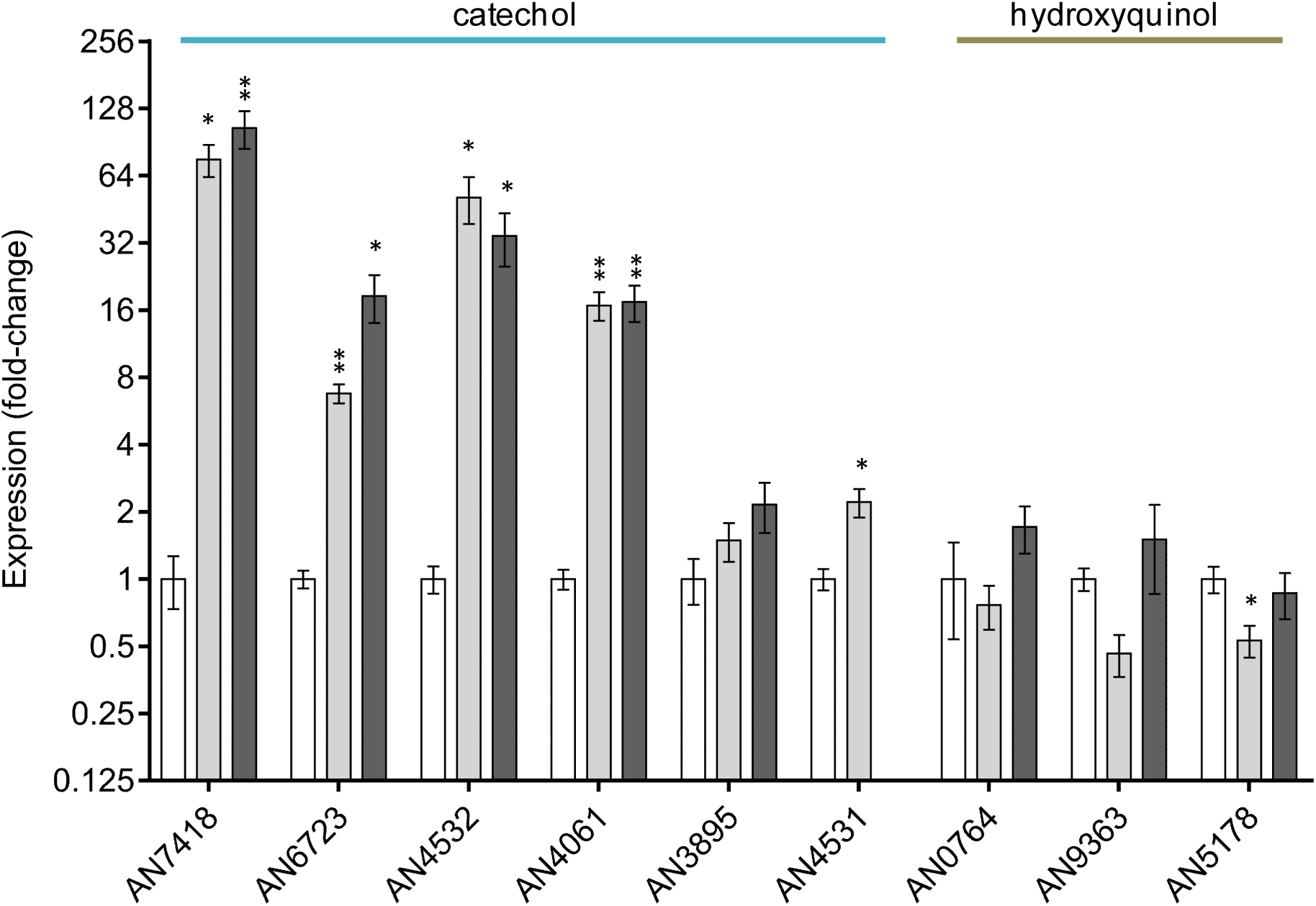
Gene expression of metabolic pathways of *A. nidulans* A1147 wild type (light grey bar) versus ΔAN4531 deletion mutant (dark grey bar) in salicylate (50 mM) compared to the control in acetate (150 mM) (white bar) at 5 days of incubation in solid media. The central intermediates of the respective metabolic pathways are indicated above the corresponding genes expression. Genes analyzed: AN7418 salicylate 1-monooxygenase, AN6723 2,3-dihydroxybenzoate carboxylyase, AN4532 catechol 1,2-dioxygenase, AN4061 muconolactone isomerase, AN3895 muconate cycloisomerase, AN4531 3-oxoadipic enol-lactone hydrolase, AN0764 and AN9363 hydroxyquinol 1,2-dioxygenase and AN5178 maleylacetate reductase. Values represent means ± standard error of the mean of three biological replicates (* p-value < 0.05 and ** < 0.005).

**Figure S5.**
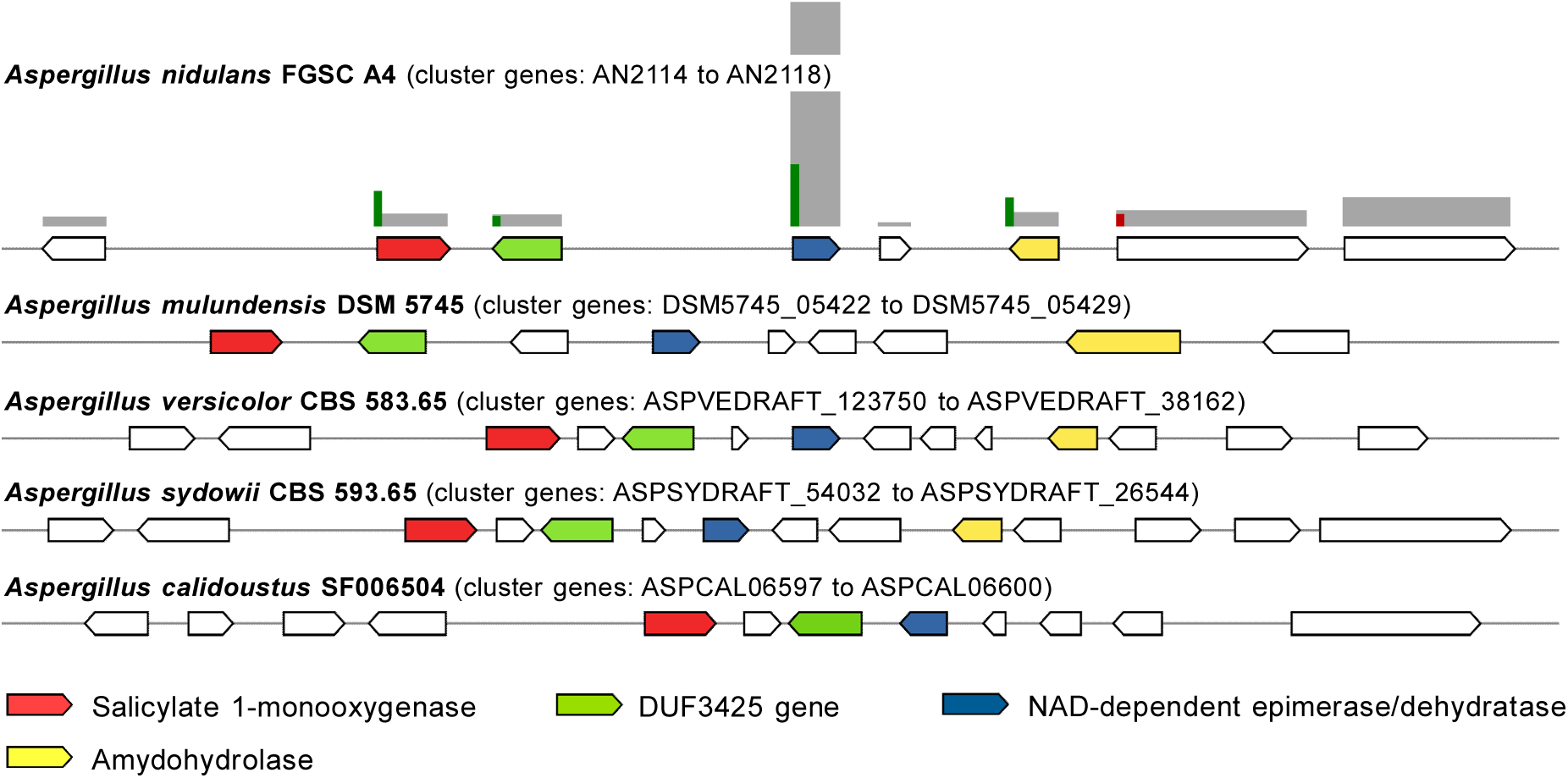
The salicylate 1-monooxygenase gene cluster in *Aspergillus* section *Nidulantes*. Here are depicted selected examples of the cluster identified within annotated genomes through sequence homology searches (MultiGeneBlast software). Genes within the cluster are represented using a color code. The differentially expressed genes (RNA-seq) regulation and abundance is represented by scale sized bars and color (upregulated in green, downregulated in red and RPKM in grey).

**Figure S6.**
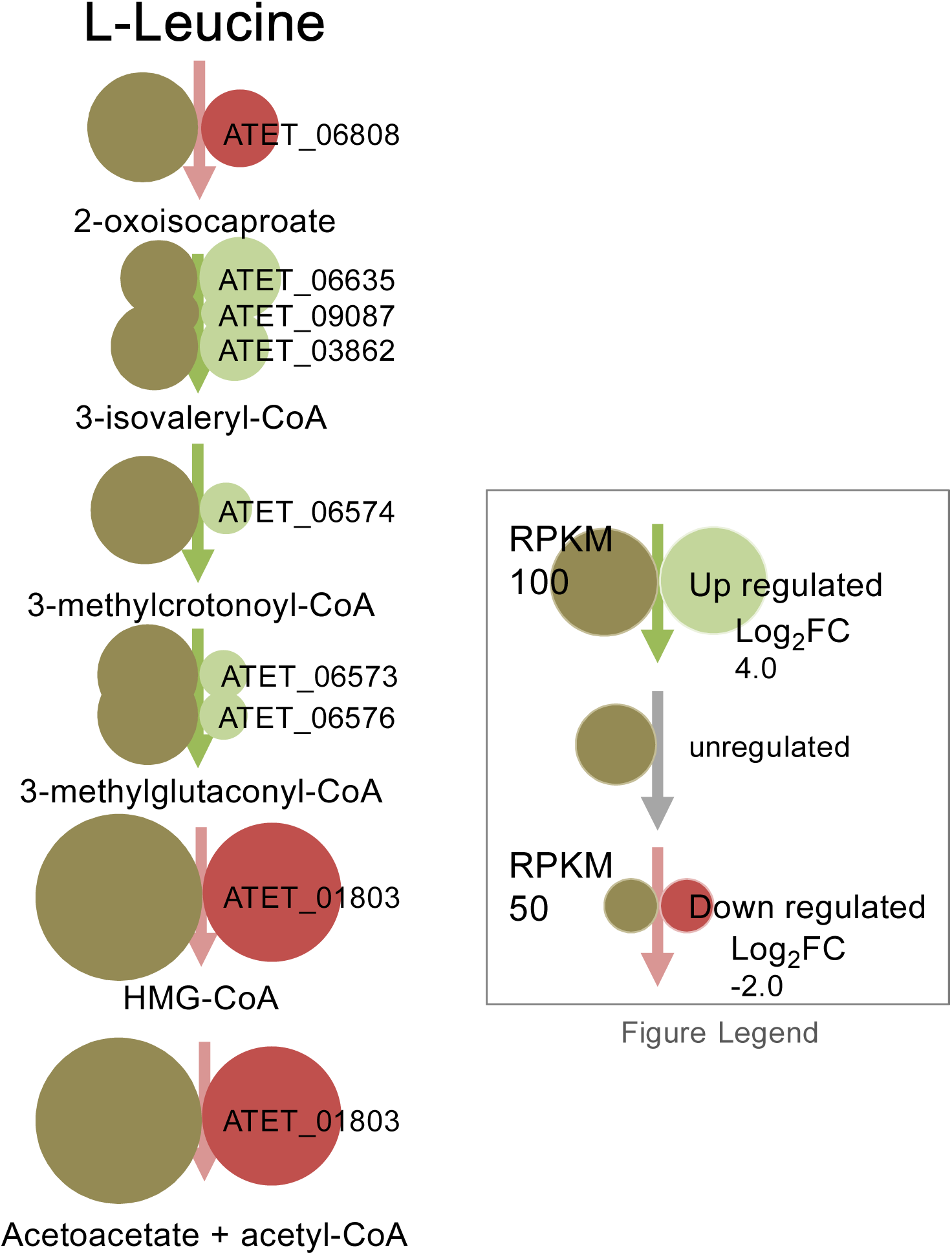
Diagram of the leucine catabolic pathway in *A. terreus*. The differentially expressed genes (RNA-seq) regulation and abundance is represented by circles with variable size and color (see legend).

